# Tunable nucleofugality in carbamoyl-bearing covalent cholinesterase inhibitors

**DOI:** 10.1101/2025.08.20.671340

**Authors:** Anže Meden, Damijan Knez, Xi Chen, Linhui Li, Xavier Brazzolotto, Maša Zorman, Anja Pišlar, Andrej Perdih, Fabrice Modeste, Celine Dalle, Milica Denic, Fabien Chantegreil, José Dias, Janez Ilaš, Janez Košmrlj, Jure Stojan, Florian Nachon, Chang-Guo Zhan, Stanislav Gobec

## Abstract

A handful of carbamate warheads is utilised in chemical biology to target serine hydrolases. The following case study on cholinesterases is the first comprehensive structure-*reactivity* exploration of the carbamoyl warhead, rather than one-target-oriented structure-*activity* study, with in-depth profiling of diverse halogen, chalcogen, and nitrogen-based leaving groups (nucleofuges) that can tune warhead reactivity. With computational tools we correlated the experimentally observed reactivities with steric and electronic factors of the investigated warheads. QM/MM simulations considering the enzymatic environment explained how substitution of carbon for nitrogen in the leaving groups of compounds **26** and **28** through resonance stabilisation, inductive bond polarization, and acidity amplification lowered the reaction barrier and increased the reaction rate >360 million times, making compound **28** a covalent inhibitor. Our findings underline the complexity of covalent inhibition and demonstrate that multiple complementary methods are required to interpret and predict covalent behaviour. Additionally, even though carbamates typically act as slow substrates, we were able to slow down decarbamoylation to a point where inhibition became *de facto* irreversible. The most interesting *O*-isoxazol-3-yl carbamate warhead was further profiled against the wider human proteome and showed low off-target reactivity, making it useful in further drug discovery. By establishing structure-*reactivity* principles for carbamoyl warhead, this study provides a generalisable framework for the development of selective covalent inhibitors and activity-based probes across diverse targets.

## Introduction

In the last three decades, activity-based protein profiling (ABPP) has proven invaluable in the field of serine hydrolases (SHs) and beyond.^1–3^ In medicinal chemistry, ABPP is a versatile tool not only for the discovery of enzyme inhibitors, but also enzyme activators, as well as for confirming the *in vivo* target engagement, for target validation, and selectivity profiling.^4–8^ Beyond catalytic Ser and Cys, the ABPP toolbox also covers Lys, Met, and other non-catalytic residues (usually through trapping and photoaffinity labelling), and even electrophilic post-translational modifications.^4,9–12^

Among more than 200 known SHs that represent about 1% of all human proteins, half of them can be classified as serine proteases (divided into trypsin, chymotrypsin, and subtilisin families), while other are metabolic SHs that preferably hydrolyse (thio)ester or amide substrates. Their active site contains a catalytic dyad (Ser-Lys/Asp) or triad (Ser-His-Asp/Glu, Ser-Ser-Lys), typically with catalytic Ser within a GXSXG consensus sequence and α/β hydrolase fold for tertiary structure.^13^ An ABPP probe for SHs typically consists of a reactive functional group (warhead) that mimics (thio)ester/amide moieties in natural substrates of SHs, a linker, and a detection tag (fluorophore, biotin, radionuclide). Some SH-selective warheads identified so far are fluorophosphonates (e.g., FP-biotin – widely used, covers >80% of SHs^5^), *O*- and *S*-carbamates, benzoxathiazin-3-one-1,1-dioxides, *N*-heterocyclic ureas (*N*-carbamoyl pyr-, tri-, and tetrazoles), phenyl and peptidyl esters, pyrrolopyrazinediones, diphenyl phosphonates, sulfonyl fluorides, β-lactams, 4-chloroisocoumarins, 4-oxo-β-lactams, 3-oxo-β-sultams, β-lactones, aza-β-lactams, oxime esters, boronic acids, etc.^4,14–16^ Useful chemical probes can be developed from electrophilic moieties, as these often display unexpected proteome selectivity in binding to proteins.^17^ For example, fine-tuning of the reactivity and selectivity of S^VI^–F electrophiles was demonstrated,^18^ and in the last decade, sulfonyl fluorides and sulfonyl fluoride exchange (SuFEX) reactions have earned their place in chemical biology, ABPP, and drug discovery.^19,20^

Human acetylcholinesterase (hAChE) and human butyrylcholinesterase (hBChE) are two metabolic SHs responsible mainly for terminating cholinergic neurotransmission, in addition to performing several other functions in the human body. Cholinesterase inhibitors (ChEIs) have various uses, especially for treatment of neurodegenerative diseases, such as dementia and myasthenia gravis, and as pesticides in agricultural chemistry.^21,22^ The majority of covalent ChEIs from the literature are either *O*-(hetero)aryl carbamates (e.g., the two-centuries-old first ChEI – physostigmine), carbamoyl halides, sulfonyl fluorides, and organophosphorous compounds (see List S1 for a comprehensive overview of all reported chemical classes). Beyond FP-biotin, the reactivities of ABPP probes on cholinesterases (ChEs) were rarely reported, though.^5,23–25^ Designated as “pseudo-irreversible” inhibitors, carbamates in fact act as slow substrates of ChEs – i.e., after the initial reversible, non-covalent complex is formed, the leaving group (LG) exits, and catalytic Ser (i.e., Ser198 in hBChE, or Ser203 in hAChE) becomes carbamoylated. The enzyme is finally regenerated through hydrolysis with a water molecule – however, the decarbamoylation step is slower and takes from minutes to several days, depending on the sterics and electronics of the carbamoyl moiety (Scheme 1).

**Scheme 1.**
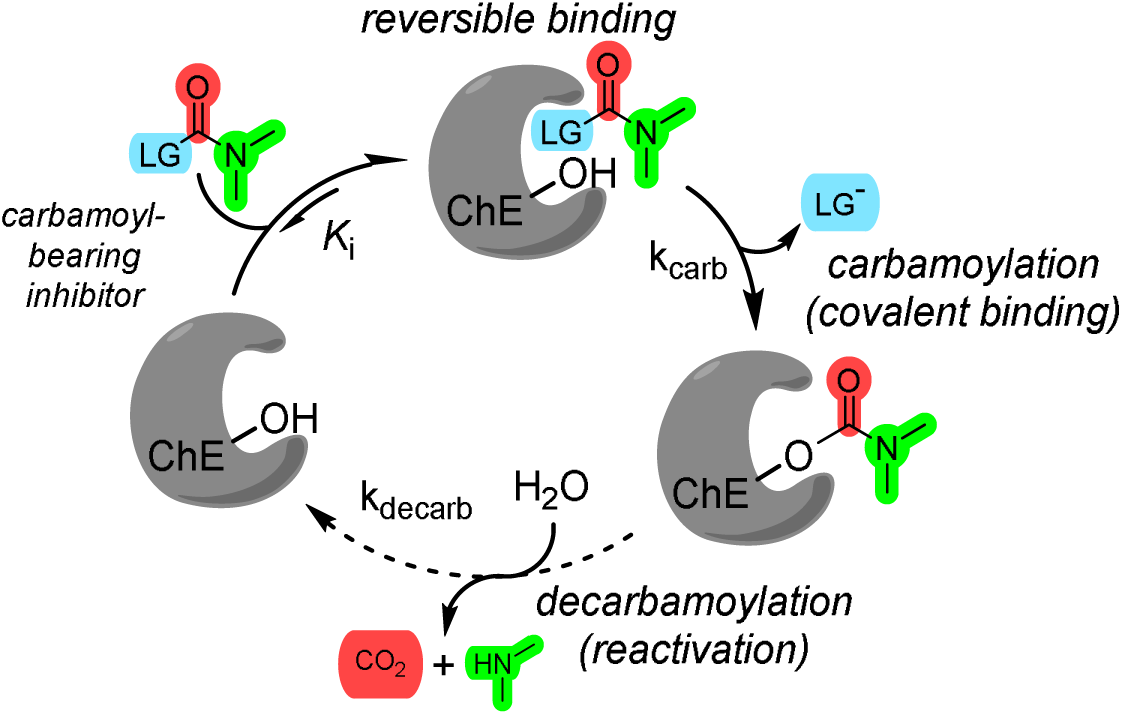
Inhibition of ChEs by the carbamate- and carbamoyl-bearing inhibitors – pseudo-catalytic cycle.

Carbamates and carbamoyl-bearing compounds have been recognised as a “privileged scaffold” due to their tunable selectivity for various SHs, which can be modulated by optimizing their intrinsic reactivity or affinity inside the active site.^26–28^ While non-covalent interactions of the carbamoyl *N*-substituents in the active site are important for providing affinity and selectivity, they only make a minor contribution to the intrinsic reactivity of the carbamoyl group. The latter is mainly influenced by the leaving group (LG), or more specifically, nucleofuge – i.e., LG that takes over the freed bonding electron pair.^29^

Therefore, we have thoroughly investigated various halogen-, chalcogen-, and nitrogen-based LGs for the carbamoyl warhead in a case study on ChEs, systematically investigating the structural factors that determine whether a chemical moiety will function as a LG or not, and thus also determine the reactivity and ultimately the selectivity of the carbamoyl group. The covalent nature of binding was evaluated and confirmed using a range of complementary techniques with increasing reliability. Several cases of compounds with puzzling reactivities were tackled by combining kinetic, computational, and crystallographic studies that partly separated the effects of sterics and electronics that govern the covalent bonding event. Additionally, an association was found between the warhead reactivity and several quantum-chemical molecular descriptors describing the electrophilicity of the carbamoyl group and reaction energetics. For the first time, a multiscale QM/MM study of the carbamoylating ChEIs was performed, and we successfully showed how a minimal change in the leaving group, such as a carbon-nitrogen substitution, changed mode of inhibition from non-covalent to covalent. Furthermore, as we increased the size of the carbamoyl *N*-substituents and made LGs as small as possible, we were able to significantly prolong decarbamoylation phase to the point where inhibition became *de facto* irreversible – this phenomenon has not yet been described for carbamate BChEIs. Finally, a promising *O*-isoxazol-3-yl carbamate warhead was activity-profiled against human proteome, and only a handful of targets other than ChEs were identified.

## Results and discussion

We wanted to investigate diverse LGs to elucidate the factors that determine whether a chemical moiety will function as a leaving group. First and foremost, as one widely accepted tenet in organic chemistry teaches us, the ability of the moiety to act as a nucleofuge (i.e., nucleofugality) is proportional to its ability to stabilise the additional seized electron density that results from heterolytic bond cleavage, and inversely proportional to the p*K*_a_ of its conjugate acid. Therefore, using a hBChE-selective *O*-7*-*indolyl carbamate inhibitor scaffold^30^ (compound **1**, Table 1), analogues of **1** (compounds **2–33**, Tables 1–2) with different phenol isosteres as LGs – that is, isosteres in terms of comparable acidity (p*K*_a_ = 10 ± 5) and molecular size – were synthesised and tested for ChE inhibitory activity.

**Table 1.**
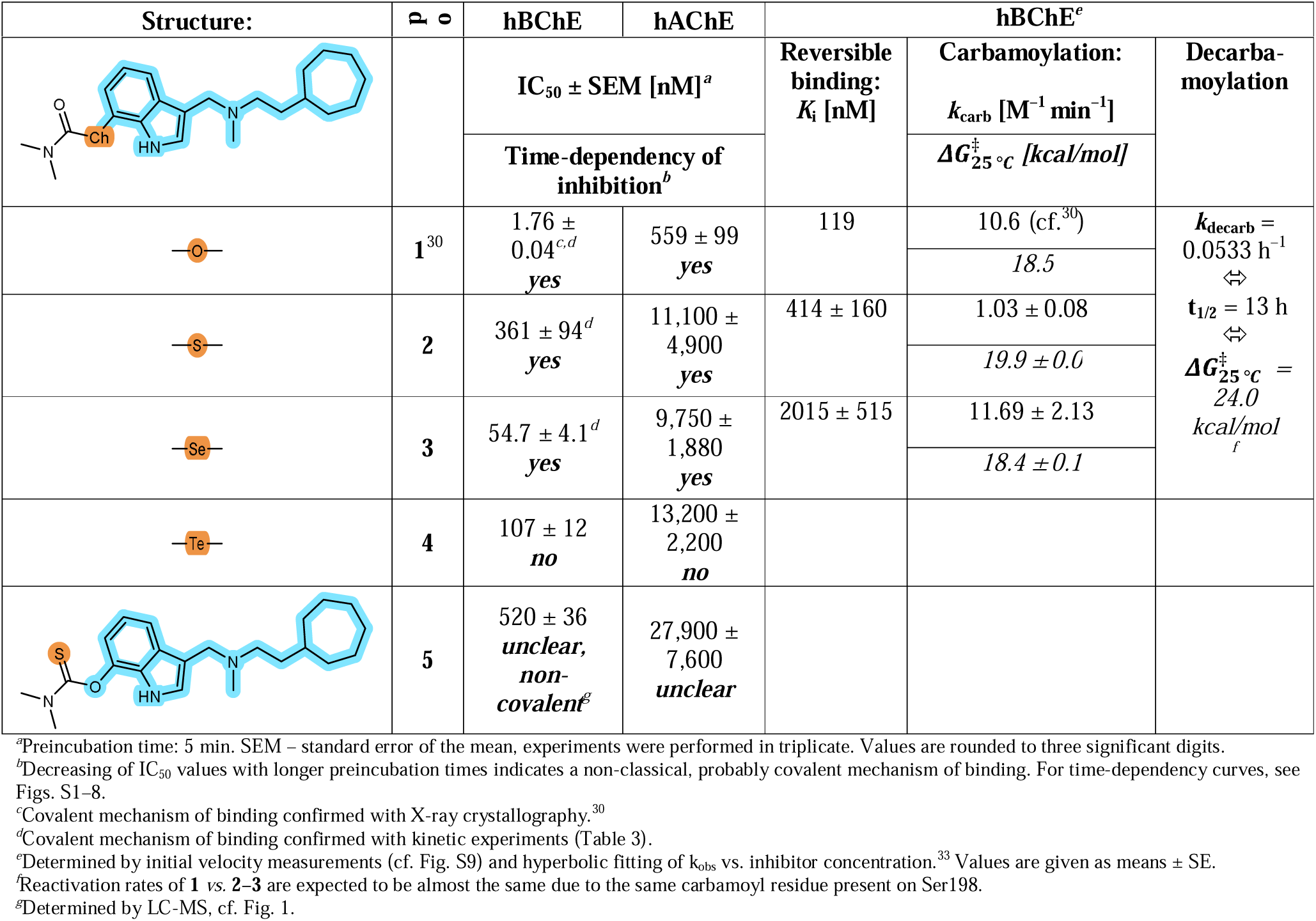
ChE inhibition by dimethyl *Ch*-carbamates featuring chalcogen-based LGs.

**Table 2.**
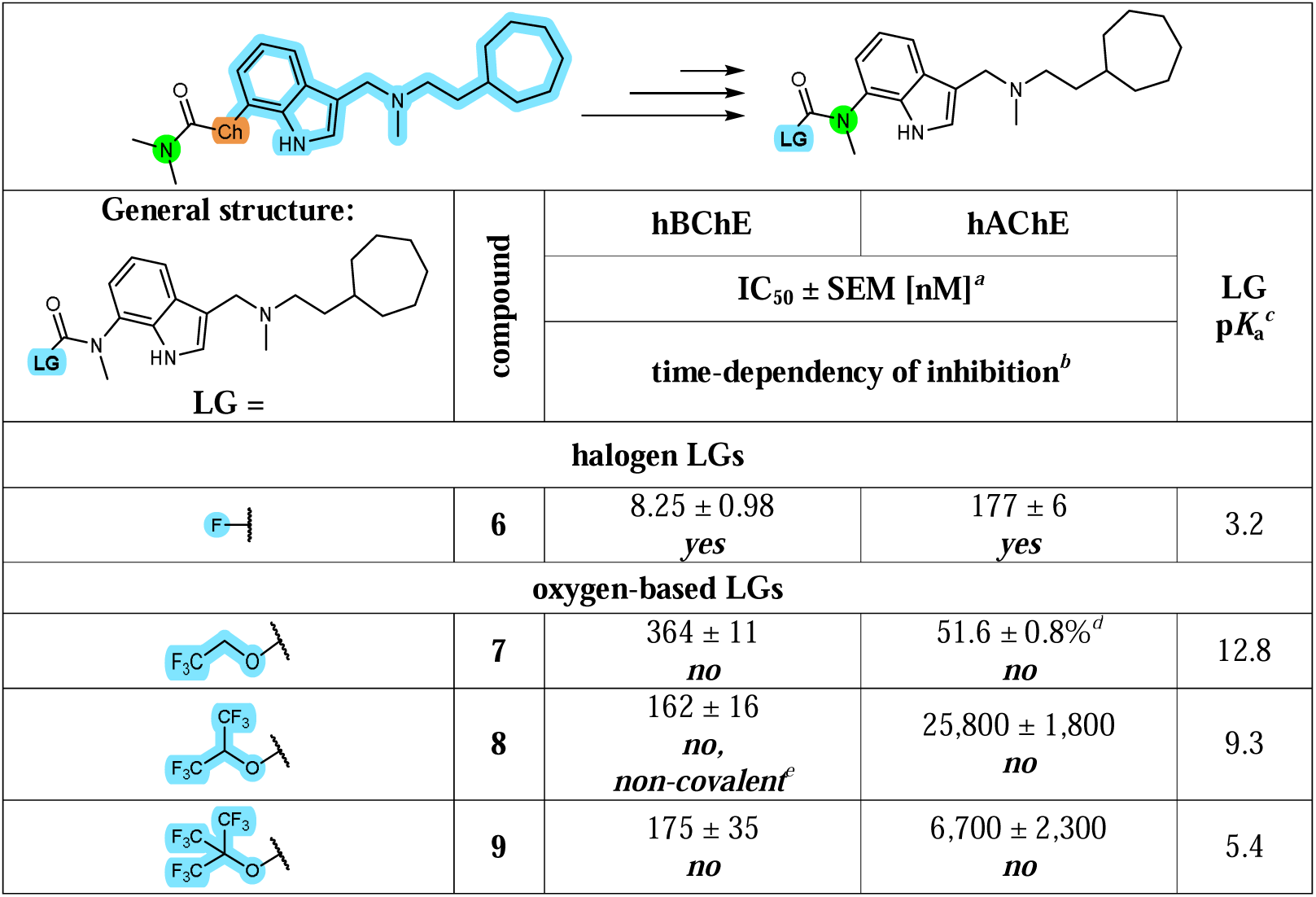
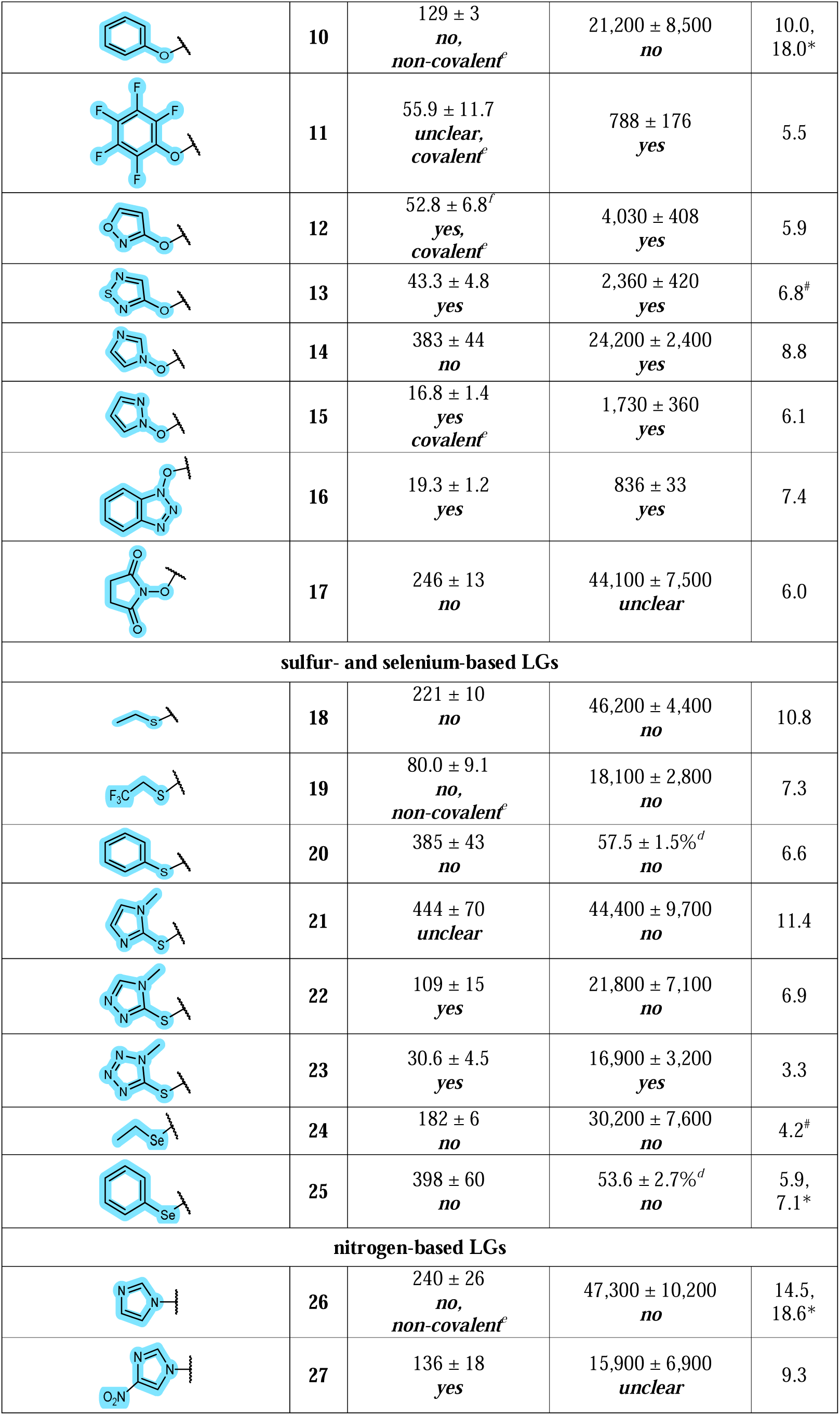
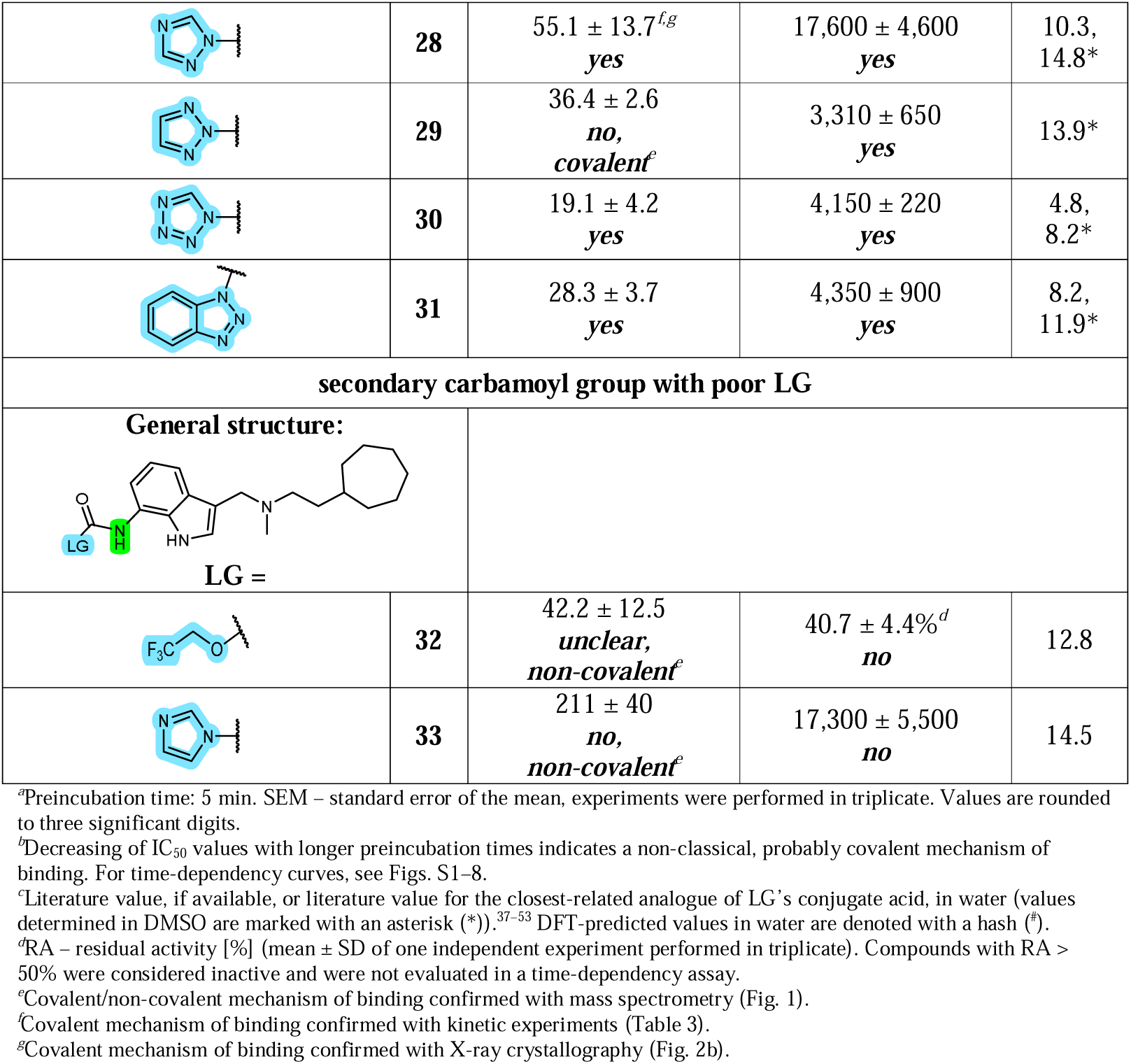
ChE inhibition by carbamoyl-bearing compounds **6–33** featuring small LGs (in blue)

|  |  |  |  |  |
| --- | --- | --- | --- | --- |
| **General structure:**<br>**LG =** | **compound** | **hBChE** | **hAChE** | **LG**<br>**p*K*<sub>a</sub><sup>c</sup>** |
|  |  | **IC<sub>50</sub> ± SEM [nM]<sup>a</sup>** |  |  |
|  |  | **time-dependency of inhibition<sup>b</sup>** |  |  |
| **halogen LGs** |  |  |  |  |
|  | **6** | 8.25 ± 0.98<br>*yes* | 177 ± 6<br>*yes* | 3.2 |
| **oxygen-based LGs** |  |  |  |  |
|  | **7** | 364 ± 11<br>*no* | 51.6 ± 0.8%<sup>d</sup><br>*no* | 12.8 |
|  | **8** | 162 ± 16<br>*no*,<br>*non-covalent*<sup>e</sup> | 25,800 ± 1,800<br>*no* | 9.3 |
|  | **9** | 175 ± 35<br>*no* | 6,700 ± 2,300<br>*no* | 5.4 |
|  | 10 | 129 ± 3<br>*no*,<br>*non-covalent*<sup>e</sup> | 21,200 ± 8,500<br>*no* | 10.0,<br>18.0* |
|  | 11 | 55.9 ± 11.7<br>*unclear*,<br>*covalent*<sup>e</sup> | 788 ± 176<br>*yes* | 5.5 |
|  | 12 | 52.8 ± 6.8<sup>f</sup><br>*yes*,<br>*covalent*<sup>e</sup> | 4,030 ± 408<br>*yes* | 5.9 |
|  | 13 | 43.3 ± 4.8<br>*yes* | 2,360 ± 420<br>*yes* | 6.8<sup>#</sup> |
|  | 14 | 383 ± 44<br>*no* | 24,200 ± 2,400<br>*yes* | 8.8 |
|  | 15 | 16.8 ± 1.4<br>*yes*<br>*covalent*<sup>e</sup> | 1,730 ± 360<br>*yes* | 6.1 |
|  | 16 | 19.3 ± 1.2<br>*yes* | 836 ± 33<br>*yes* | 7.4 |
|  | 17 | 246 ± 13<br>*no* | 44,100 ± 7,500<br>*unclear* | 6.0 |
| **sulfur- and selenium-based LGs** |  |  |  |  |
|  | 18 | 221 ± 10<br>*no* | 46,200 ± 4,400<br>*no* | 10.8 |
|  | 19 | 80.0 ± 9.1<br>*no*,<br>*non-covalent*<sup>e</sup> | 18,100 ± 2,800<br>*no* | 7.3 |
|  | 20 | 385 ± 43<br>*no* | 57.5 ± 1.5%<sup>d</sup><br>*no* | 6.6 |
|  | 21 | 444 ± 70<br>*unclear* | 44,400 ± 9,700<br>*no* | 11.4 |
|  | 22 | 109 ± 15<br>*yes* | 21,800 ± 7,100<br>*no* | 6.9 |
|  | 23 | 30.6 ± 4.5<br>*yes* | 16,900 ± 3,200<br>*yes* | 3.3 |
|  | 24 | 182 ± 6<br>*no* | 30,200 ± 7,600<br>*no* | 4.2<sup>#</sup> |
|  | 25 | 398 ± 60<br>*no* | 53.6 ± 2.7%<sup>d</sup><br>*no* | 5.9,<br>7.1* |
| **nitrogen-based LGs** |  |  |  |  |
|  | 26 | 240 ± 26<br>*no*,<br>*non-covalent*<sup>e</sup> | 47,300 ± 10,200<br>*no* | 14.5,<br>18.6* |
|  | 27 | 136 ± 18<br>*yes* | 15,900 ± 6,900<br>*unclear* | 9.3 |
|  | **28** | 55.1 ± 13.7<sup>f,g</sup><br>**yes** | 17,600 ± 4,600<br>**yes** | 10.3,<br>14.8* |
|  | **29** | 36.4 ± 2.6<br>**no,**<br>**covalent<sup>e</sup>** | 3,310 ± 650<br>**yes** | 13.9* |
|  | **30** | 19.1 ± 4.2<br>**yes** | 4,150 ± 220<br>**yes** | 4.8,<br>8.2* |
|  | **31** | 28.3 ± 3.7<br>**yes** | 4,350 ± 900<br>**yes** | 8.2,<br>11.9* |
| **secondary carbamoyl group with poor LG** |  |  |  |  |
| **General structure:**<br>**LG =** |  |  |  |  |
|  | **32** | 42.2 ± 12.5<br>**unclear,**<br>**non-covalent<sup>e</sup>** | 40.7 ± 4.4%<sup>d</sup><br>**no** | 12.8 |
|  | **33** | 211 ± 40<br>**no,**<br>**non-covalent<sup>e</sup>** | 17,300 ± 5,500<br>**no** | 14.5 |

<sup>a</sup>Preincubation time: 5 min. SEM – standard error of the mean, experiments were performed in triplicate. Values are rounded to three significant digits.
<sup>b</sup>Decreasing of IC<sub>50</sub> values with longer preincubation times indicates a non-classical, probably covalent mechanism of binding. For time-dependency curves, see Figs. S1–8.
<sup>c</sup>Literature value, if available, or literature value for the closest-related analogue of LG's conjugate acid, in water (values determined in DMSO are marked with an asterisk (\*)).<sup>37–53</sup> DFT-predicted values in water are denoted with a hash (#).
<sup>d</sup>RA – residual activity [%] (mean ± SD of one independent experiment performed in triplicate). Compounds with RA > 50% were considered inactive and were not evaluated in a time-dependency assay.
<sup>e</sup>Covalent/non-covalent mechanism of binding confirmed with mass spectrometry (Fig. 1).
<sup>f</sup>Covalent mechanism of binding confirmed with kinetic experiments (Table 3).
<sup>g</sup>Covalent mechanism of binding confirmed with X-ray crystallography (Fig. 2b).

We would like to emphasize that our study focused primarily on whether various carbamoyl-bearing warheads achieved covalent inhibition of ChEs, rather than on the absolute values of the inhibitory activity of the investigated compounds. There are two reasons for this approach: first, the extent of inhibition is strongly dependent on the rate of covalent reaction, which is, roughly speaking, governed by steric and electronic factors. Since significant steric hindrance can effectively prevent even a thermodynamically favourable reaction, we also evaluated the respective fragment-sized precursors of final compounds (i.e., **2a**–**31a**, Scheme 2, Table S1) to somewhat account for the effects of steric repulsion on warhead reactivity. Secondly, the non-covalent interactions of close analogues should be similar, but not entirely. A different LG moiety could, at least in principle, offer better or additional non-covalent interactions with the active site residues, leading to a better inhibition than a slow-reacting, covalent inhibitor with suboptimal non-covalent interactions (cf. Scheme 1).

**Scheme 2.**
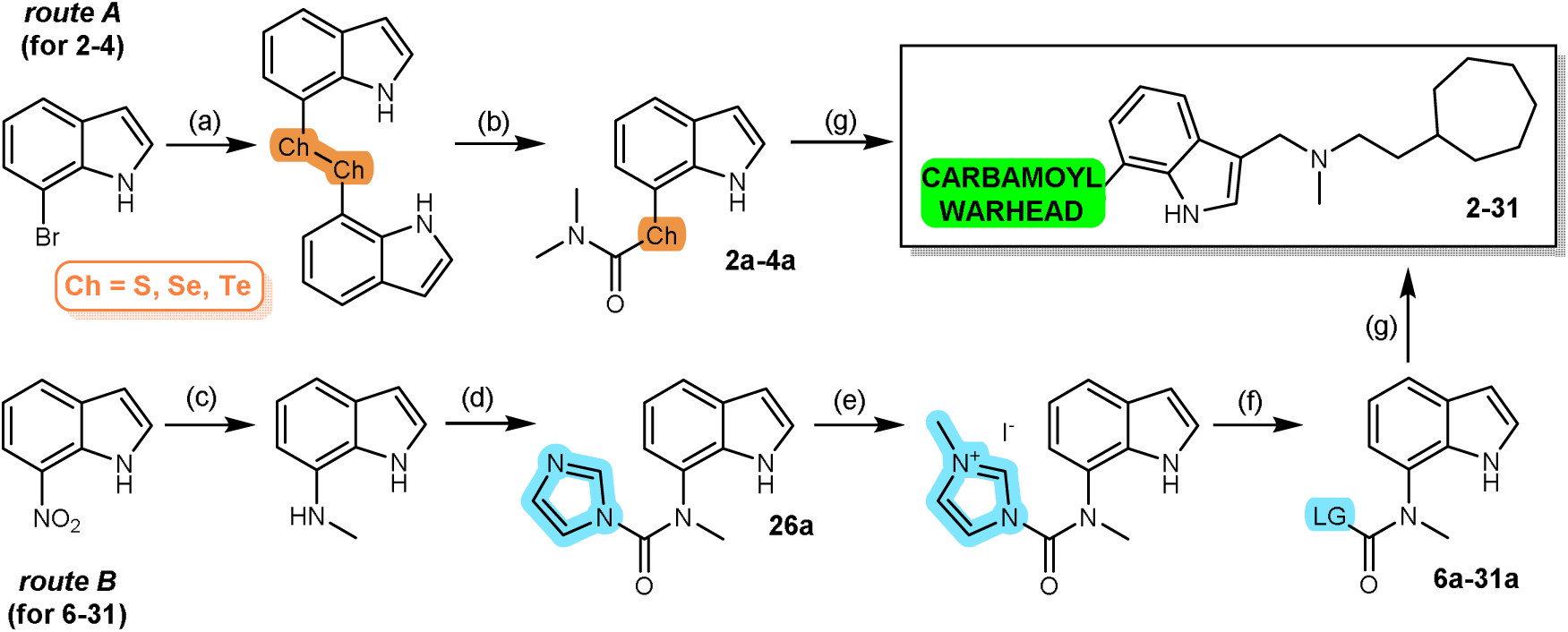
General synthetic routes to carbamoyl-bearing final compounds **2–31**. *Reagents and conditions*: (a) i. *n*-butyllithium, THF, –80 °C, ii. *tert*-butyllithium, –80 °C to –40 °C, iii. elementary chalcogen, –40 °C to rt; (b) i. sodium borohydride, THF/EtOH, rt, ii. dimethylcarbamoyl chloride, triethylamine; (c) i. Pd/C, sodium borohydride, MeOH, 0 °C; ii. ethyl formate, 100 °C; iii. lithium aluminium hydride, THF, 60 °C; (d) CDI, THF, 60 °C; (e) iodomethane, MeCN, rt; (f) LG-H, DIPEA, MeCN, rt; (g) formalin, *N*-methyl-2-cycloheptylethylamine, AcOH, rt.

The following triage of experiments was used to ascertain if compounds bound to ChEs covalently. First, time-dependency profiles of IC_50_ values were determined for all compounds since decreasing IC_50_ values with longer preincubation time indicate a covalent mechanism of binding.^31,32^ Then, for a selection of hits, further kinetic analysis, LC-MS, and X-ray crystallography provided unequivocal proof of covalent binding (*vide infra*).

### 1 Periodic trends and chalcogens

The hydrides’ p*K*_a_ values decrease, the atomic radii increase, and electronegativities decrease upon descending a certain group within the periodic table of elements – these are some of the periodic trends. Accordingly, we wondered how these trends within group 16 (VI) of the periodic table (the chalcogens, Ch) will affect the carbamate reactivity. Therefore, we synthesised thio-, seleno- and telluro-analogues of the *O*-aryl carbamate **1** (compounds **2**–**4**, Table 1) *via* lithium-halogen exchange of 7-bromoindole and quenching with elementary chalcogens that enabled preparation of di(7-indolyl) dichalcogenides. These were *in situ* reduced and carbamoylated with dimethylcarbamoyl chloride to afford fragment-sized dimethyl *Ch*-indolyl carbamates **2a–4a** (Scheme 2, route A), followed by a Mannich reaction, which facilely furnished final compounds **2–4** under mild reaction conditions.

Despite increased LG acidities, the inhibitory potencies of **2**–**4** were lower compared to their oxygen predecessor **1**. Furthermore, a local maximum in experimentally determined energy barrier for carbamoylation was observed at *S*-indolyl carbamate **2**, while *O*- and *Se*-indolyl carbamates **1** and **3** had comparable reaction rates (Table 1, row “carbamoylation”). This trend in activation energies was also mirrored in DFT-predicted activation barriers ΔG^‡^ for the model reaction – the addition of methoxide to chalcogen carbamates **1a–4a** in solution (*vide infra*, Table S9). Going down group 16 (VI), the significant decrease in electronegativity of heavier chalcogens, which lowers the carbamate electrophilicity, is counteracted by increased polarizability, which stabilises the negative charge of the nucleofuge. Interestingly, time dependency was not observed for the *Te*-aryl carbamate **4**, however, its fragment-sized precursor **4a** was time-dependent (Table S1). Since tellurium has approx. 2.3 times greater atomic and 1.4 times greater van der Waals radius than oxygen, the increased atom size may have skewed the binding of **4** and prevented covalent bond formation, even though the warhead itself was reactive enough. Despite the simplicity of the idea, this is the first time that all non-radioactive chalcogen carbamates, i.e., *O*-, *S*-, *Se*-, and *Te*-aryl carbamates, were synthesised and reported as covalent cholinesterase inhibitors. Additionally, in line with the double bond rule,^34^ the replacement of a smaller, harder, and strongly electron-withdrawing oxygen in the carbonyl group with a softer, larger, and more polarisable sulfur atom in thiocarbonyl group precluded covalent inhibition by *O*-thiocarbamate **5** (confirmed by LC-MS, as well – cf. Fig. 1b), either due to lower electrophilicity or steric inability to form activating hydrogen bonds with the oxyanion hole residues.

**Fig. 1.**
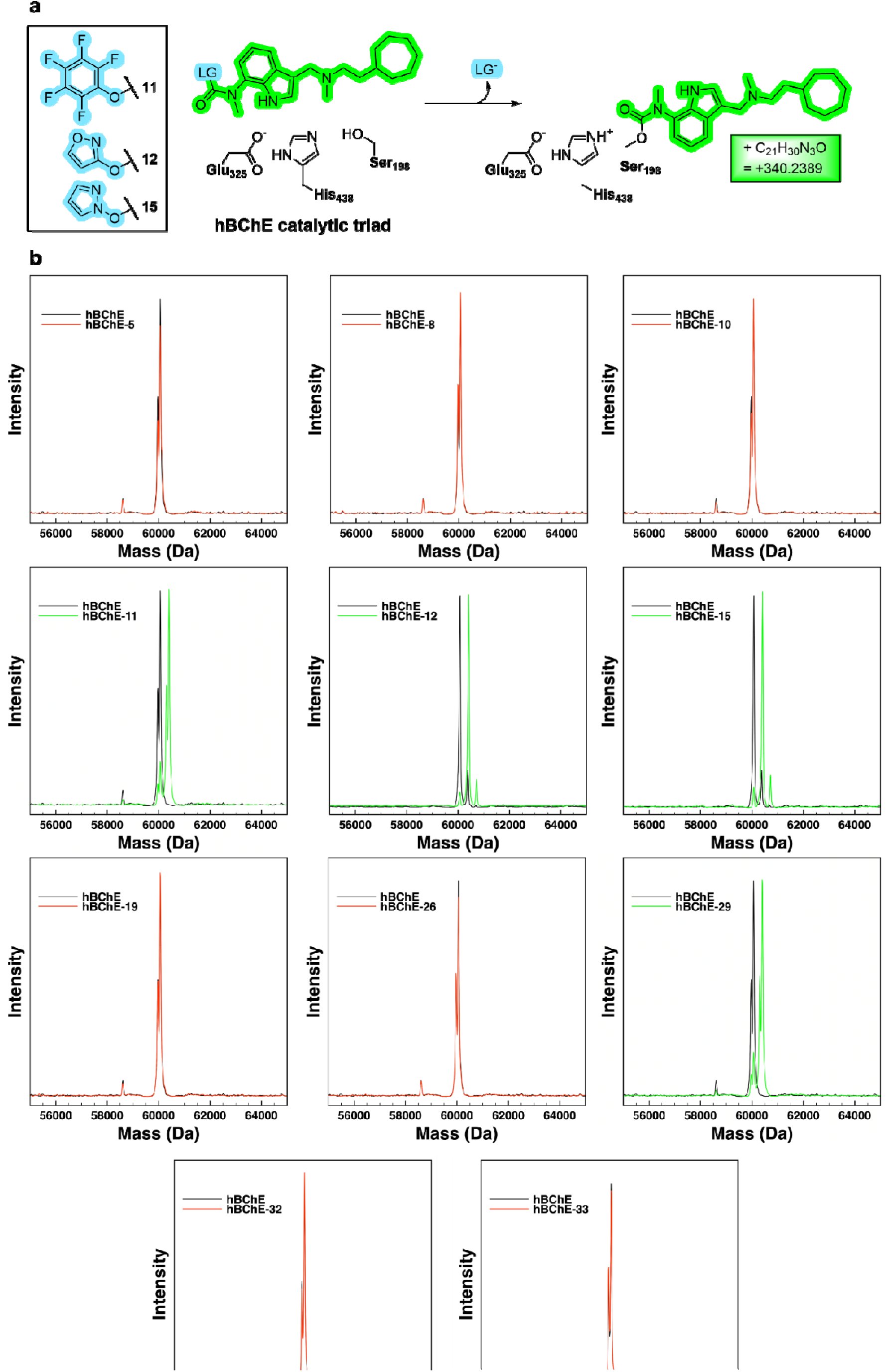
Mass-spectrometry carbamoylation study of hBChE. **a**, Carbamoylation scheme of hBChE by covalent inhibitors **11**, **12**, and **15**. The carbamoyl moiety is coloured green, the leaving group in blue, and the enzyme in black. **b**, Overlays of deconvoluted MS spectra for control sample (in black) and inhibitor-treated samples (in green – covalent adduct detected, in red – no adduct detected). Left to right, inhibitors **5**, **8**, **10** (first row), **11**, **12**, **15** (second row), **19**, **26**, **29** (third row), **32** and **33** (fourth row). The native hBChE in the control sample featured a mass peak of 60073 ± 1 Da (4 Da difference to the ProtParam^60^-calculated mass). **11**, **12**, **15**, and **29**-incubated samples displayed a peak at 60413 ± 1 Da, indicative of covalent binding.

### 2 Making nucleofuge as small as possible

Leaving group has thus far occupied a larger part of the inhibitor structure, and it would remain inside the active site after carbamoylation of catalytic Ser.^30,35^ But if LG could be made sufficiently small, then the bound carbamoyl moiety would occupy a significant portion of the active site, and perhaps hinder the access of water molecules, effectively retarding decarbamoylation. With that in mind, we designed and synthesised “inverted” compounds **6**– **31** (Table 2) with the *N*-methyl-*N*-indol-7-ylcarbamoyl moiety and a comparably small LG. Here, a six-step synthetic procedure starting from 7-nitroindole was used (Scheme 2, route B). Due to the instability of indole-7-amine derivatives, carbamoylation of different leaving groups was easily and cleanly achieved only by using *N*,*N*-dialkylcarbamoylimidazolium salts,^36^ leading to fragment-sized indolyl carbamates **6a–31a**, and, after Mannich reaction, to final compounds **6**–**31**.

#### 2.1 Halide nucleofuges

Featuring the smallest possible, one-atom LG, carbamoyl fluoride **6** was, as expected, a potent covalent ChEI (Table 2). Additionally, in line with the double bond rule,^34^ the fragment-sized thio- and selenocarbamoyl fluorides **6c** and **6d**, respectively, did bind to hBChE covalently, but they exhibited lower potency than their oxygen analogue **6b** (Table S1).

#### 2.2 Oxygen-based nucleofuges

Among the oxygen-based LGs, different carboxylic acid isosteres^54^ and other functional groups of comparable acidity were tested. In contrast with a handful of literature cases,^55–59^ mono-, di- and tri(trifluoro)alcohol derivatives **7**–**9** did not function as covalent ChEIs despite sufficient LG acidity. For example, *O*-hexafluoroisopropyl carbamate probes were described to be proteome-wide selective for endocannabinoid hydrolases,^26^ therefore it is not surprising that compound **8**’s inhibition was non-covalent – in a mass spectrometry (MS) study, the expected increase in molecular mass (340 ± 1 Da for the carbamoylated hBChE) was not observed (Fig. 1). The steric bulk combined with significant repulsion by electronegative fluorine substituents could prevent **7**–**9** from accessing the catalytic Ser residue in ChEs.

Surprisingly, the phenol derivative **10** displayed time-independent behaviour and no carbamoylation in LC-MS experiments (Fig. 1b). Since *O*-aryl carbamates are well-known ChEIs, this would indicate that the access of the warhead to Ser198 must have been impeded due to steric hindrance – molecular dynamics (MD) simulations of **10**-hBChE complex support this conclusion, as the pre-reaction pose was not stable during the simulated time (Fig. S14). However, the more reactive pentafluorophenyl analogue **11**, whose initial time-dependency profile was ambiguous (Table 2, Fig. S3), was indeed able to produce a covalent adduct, as demonstrated by LC-MS (Fig. 1b).

Meanwhile, smaller heterocycles with pronounced electron-withdrawing character^61^ acted as effective LGs (compounds **12**–**15**). For **12**, which has a 3-hydroxyisoxazole LG, the covalent mechanism of binding was determined by progress curve analysis (Fig. S10). The initial reversible binding constant *K*_i_ was determined to be 230 ± 12 nM, followed by carbamoylation with the rate of 5.28 ± 0.54 M^−1^ min^−1^, which equals to the reaction barrier ΔG^‡^ 18.9 ± 0.1 kcal/mol. The covalent binding of **12** was also confirmed by LC-MS (Fig. 1b).

Compound **15** with its *N*-hydroxypyrazole nucleofuge exhibited time-dependency and produced an appropriate MS adduct (Fig. 1b). Interestingly, its positional isomer, *N*-hydroxyimidazole-derived **14** could not produce a stable pre-reaction pose in the hBChE active site *in silico* (Fig. S14), which could be the reason behind its failure to covalently inhibit hBChE, but not hAChE. As for **16**, its bulky *N*-hydroxybenzotriazole (HOBt) moiety effectively replaced indole in the acyl-binding pocket *in silico*, leading to an entirely different binding mode while retaining covalent binding. *N*-Hydroxysuccinimide (NHS)-derived **17** was inactive, perhaps due to repulsion of the NHS moiety with catalytic site residues.

#### 2.3 Thiol- and selenol nucleofuges

Among the sulfur- and selenium-based LGs, a similar trend was observed. Despite sufficient LGs’ acidity, aliphatic derivatives **18**, **19** (confirmed by LC-MS, cf. Fig. 1b), **24**, and bulky aromatic **20** and **25** were noncovalent, in line with the reports on S and Se analogues of *N*,*N*-diethylcarbamylcholine.^62^ For the three homologous 2-thioazole *S*-thiocarbamates **21**–**23**, at least 3 ring nitrogens were needed for sufficient nucleofugality. The plausible underlying reason for the efficacy of the aromatic *vs*. aliphatic LG counterparts could be either improved electron delocalisation within the aromatic ring or favourable π–π interactions with nearby His438, Phe398, and Phe329 residues in the active site.

#### 2.4 Azolic nucleofuges

In the class of nitrogen-based leaving groups,^14^ a similar trend was observed among the azoles. Imidazole derivative **26** acted non-covalently, however by increasing the electron-withdrawing character of the heterocyclic system (with a nitro group, as in **27**, or with additional nitrogen atoms in compounds **28**–**31**) covalent inhibitors were produced. Unfortunately, these *N*-heterocyclic ureas also displayed pronounced instability upon storage and in aqueous milieu (Table S10). Interestingly, among positional isomers **28** and **29** with 1,2,4-triazole and 1,2,3-triazole nucleofuges, respectively, the time-dependency was evident only for **28**, while only mass spectrometry confirmed that **29** is also able to bind covalently to hBChE (Fig. 1b).

Lastly, another issue worth raising are carbamoyl *N*-substituents. In reactions with nucleophiles, tertiary carbamates follow a disfavoured B_AC_2 mechanism, while secondary carbamates can also be deprotonated and then decompose into more reactive isocyanates *via* E1cB mechanism – effectively meaning up to 10^8^ increases in hydrolysis rates (Scheme S2).^63^ Therefore, if E1cB mechanism could also occur inside the SH active site, secondary and primary carbamoyl compounds would be significantly more reactive. Even though *secondary N*-carbamoylimidazoles can readily dissociate into isocyanates and imidazole,^64^ they were found unreactive against 11 diverse antimicrobial enzyme targets and glutathione, and stable in an aqueous solution for days.^17^ Apart from human leukocyte elastase (HLE) inhibitors,^65,66^ FAAH inhibitors,^67,68^ and insecticides from the patent literature,^69–71^ *N*-carbamoylimidazoles have rarely been reported in drug discovery, except as versatile synthesis intermediates.^36^ In our hands, tertiary *N*-carbamoylimidazole **26** did not bind covalently to hBChE, and neither did its secondary analogue **33**, nor **32** with an alcoholic LG (Table 2), which was confirmed by LC-MS, as well (cf. Fig. 1b). At least for ChEs, the nucleofugality of the LG seems to be more important for determining reactivity than carbamoyl *N*-substitutents.

### 3 An *in-silico* approach to disentangle the steric and electronic effects on the reactivity

We applied *in silico* approaches ranging from classical docking to MD simulations to try to wholesomely explain the influence of sterics on the carbamoylation of hBChE’s catalytic Ser198. In order to successfully covalently bind, the ligand’s warhead must be able to approach and reside near the nucleophilic Ser198 residue for a sufficient amount of time. To obtain these pre-reaction poses, the inhibitors were first docked to the active site of hBChE, where Ser198 was mutated to Ala198 to eliminate a potential steric clash. Additionally, the post-reaction poses were obtained using CovDock covalent docking protocol.^72^ Some compounds (e.g., time-independent **8–9**, **17** or time-dependent inhibitors **16**, **31** with large LGs) were quickly excluded as they did not yield meaningful pre- or post-reaction poses – thereby indicating either a significant repulsion or an entirely different binding mode (e.g., ousting indole moiety from the acyl-binding pocket), respectively. Meanwhile, molecular dynamics of the time-dependent ligand (e.g., **28**)–hBChE pre-reaction complexes revealed that ≥2 hydrogen bonds between the ligand’s carbamoyl oxygen and oxyanion hole (Gly116,

Gly117, Ala199) were formed in most of the frames, thereby increasing the electrophilicity of the carbamoyl group. Additionally, the distance between nucleophilic Ser198 *O*^γ^ and ligand’s electrophilic carbamoyl carbon oscillated between 3–3.5 Å (close enough for the initiation of reaction), and Ser198 and His438 remained at hydrogen bond distance (2.7–3.2 Å), able to support the proton transfer that occurs concomitantly with the C*–*O bond formation. The rest of the inhibitor scaffold was involved in persistent cation–π interactions with Tyr332 at the peripheral aromatic site, π–π interactions of the aromatic LG with either His438 or Trp82 in the choline-binding pocket, and van der Waals interactions of the indole moiety with Trp231 and Phe329 in the acyl-binding pocket (Fig. 2a).

**Fig. 2.**
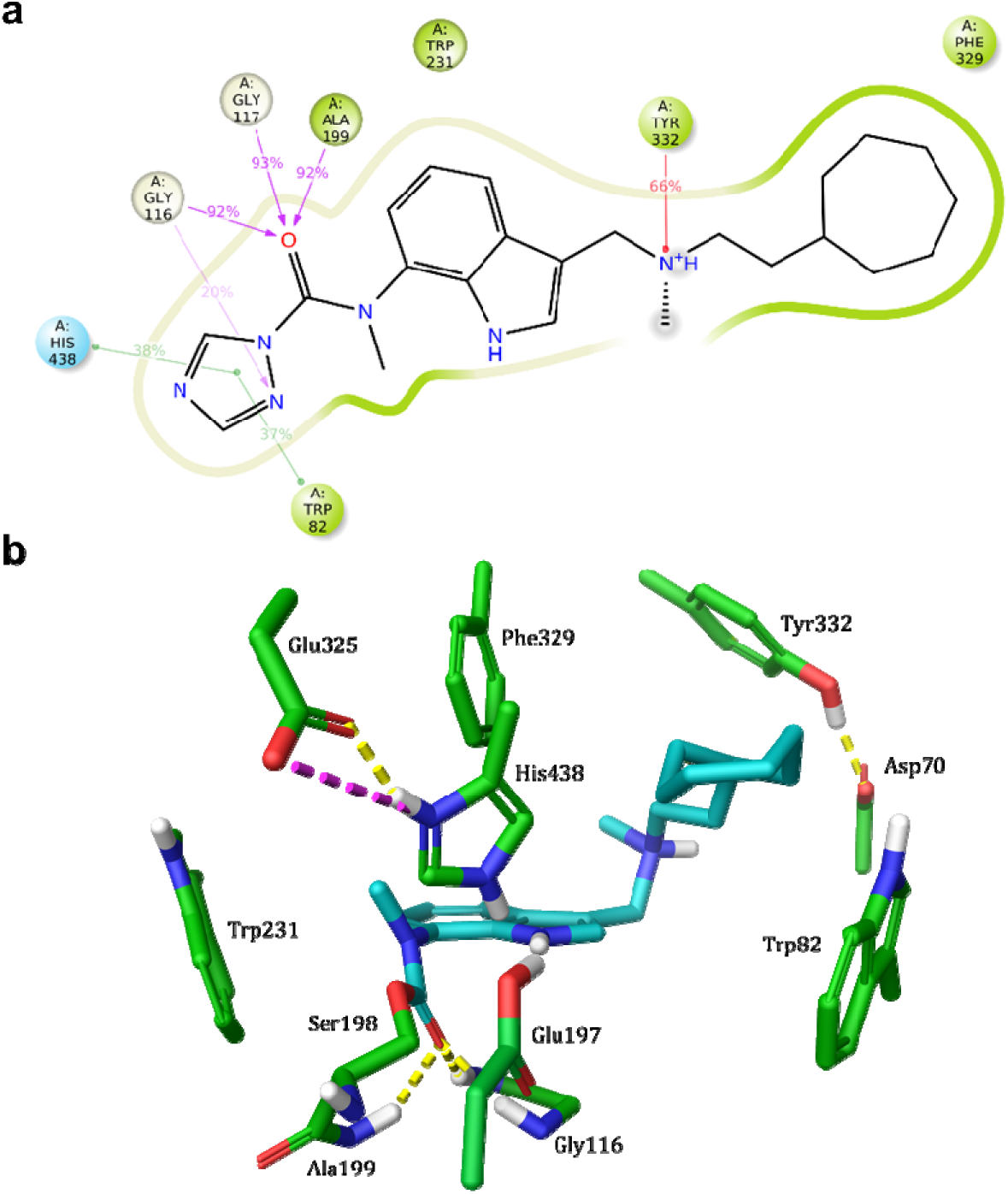
Structural insight in interactions of 28 with hBChE. **a**, Protein-ligand interaction diagram for compound **28** pre-reaction pose during 20 ns of MD simulation. Protein-ligand contacts and interactions present in >20% of the frames are depicted. The π–π interactions are shown as green lines, the cation–π interactions as red lines, and hydrogen bonds as magenta lines. **b**, Crystal structure of the **28**-carbamoylated hBChE (PDB 8QTW, resolution 2.24 Å). The key amino acid residues are shown as green sticks, and **28-**derived moiety as cyan sticks. The binding pose features a characteristic cation-π interaction of the central tertiary amine with Tyr332 at the peripheral aromatic site, while the *N*-methyl-7-aminoindolyl moiety resides in the acyl-binding pocket, forming only van der Waals interactions with the nearby residues. The carbamoyl group oxygen is fixed in the oxyanion hole, forming three hydrogen bonds (shown as yellow dashed lines) with Gly116, Gly117, and Ala199.^30^ His438 and Glu325 of the catalytic triad form a hydrogen bond and a salt bridge (magenta dashed line), while Glu197 is most likely protonated.^73,74^

Since time-independent inhibitors **6**, **18**–**19**, **21** and **26** also produced stable pre-reaction poses in MD (Fig. S14), the underlying reason for their lack of covalent inhibition must lie in the intrinsic reactivity of the warhead itself or the reaction energetics, rather than in the steric discomplementarity. To follow up on this aspect, quantum-mechanical (QM) calculations using density functional theory (DFT) were carried out for the reactions of **6a–31a** with a model nucleophile in solution (Scheme 3). We calculated enthalpy and free energy of the reactions, as well as several quantum-chemical descriptors, and compared them with the observed reactivities. The descriptors were selected based on their prevalence in previous studies^75–78^ and their availability in our computational framework. As previous studies have also shown, no single descriptor is sufficient to universally predict the nucleofugality, especially for chemically diverse compounds, and the interplay of multiple factors must be considered.^79^

**Scheme 3.**
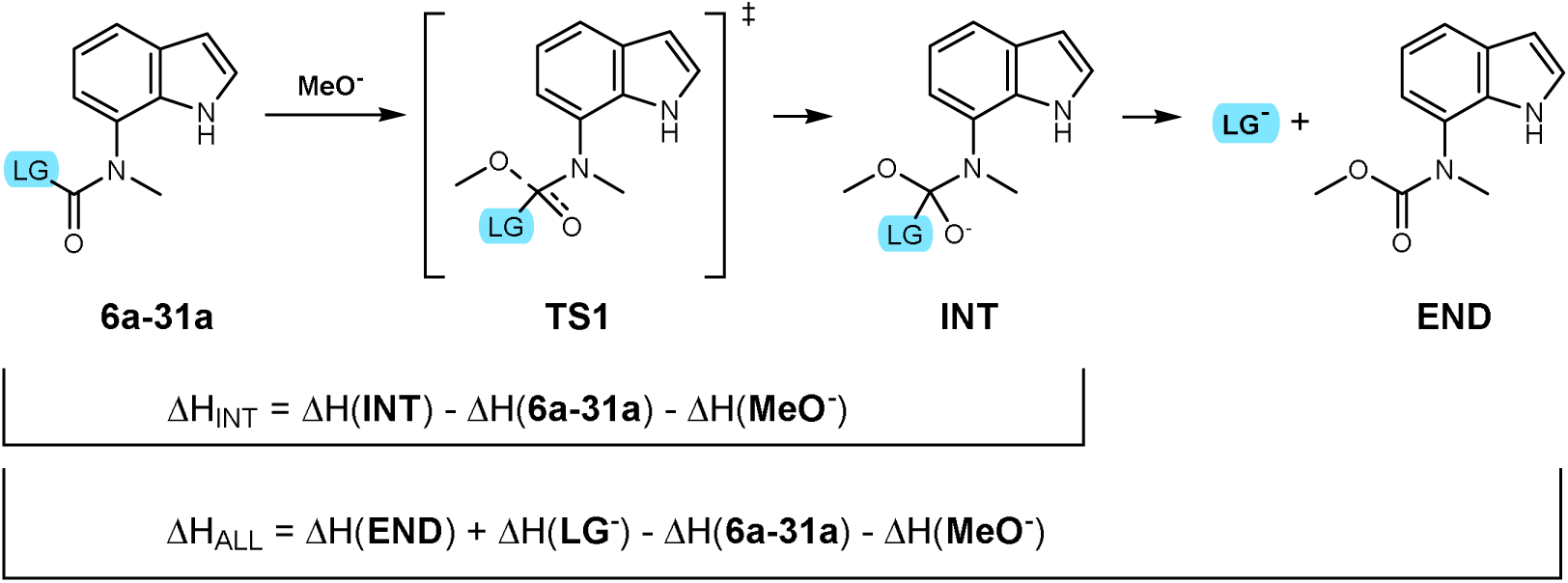
The studied model reaction – to speed up the calculations, minimal representative structures of the compounds **6–31** (i.e., the fragments **6a–31a**) and methoxide anion as catalytic Ser198 surrogate^80^ were used. DFT calculations were done at M06-2X/aug-cc-pVTZ(-f)//M06-2X/6-31G++** level of theory. The molecular descriptors that were used in statistical analysis can be found in Tables S5–S9 and the optimized atom coordinates in Supporting Information.

The statistical analysis of covalent and non-covalent inhibitor groups was done using two-sided non-parametric Mann–Whitney U-tests to identify descriptors that were statistically different between the two groups. Additionally, a non-parametric Spearman correlation of the individual descriptors with pIC_50_ (i.e., covalent inhibition typically meant a higher degree of inhibition, cf. Table 2) was performed (Table S5), followed by principal component regression of the eight descriptors that were identified as significant in both of the previous tests (Table S6). The following molecular descriptors or properties were found to statistically significant (p<0.05) differ between groups of covalent and non-covalent inhibitors: Glide XP docking score for the pre-reaction pose, ^13^C chemical shift (δ) of the carbonyl carbon, enthalpy and free energy of the reaction up to the tetrahedral intermediate **INT** (ΔH_INT_ and ΔG_INT_, Scheme 3), enthalpy of the complete reaction up to the end products (ΔH_ALL_, Scheme 3), highest atomic electrostatic potential (maxESP) for carbonyl carbon, calculated wavenumber for the carbonyl group (ν_C=O_), and electronic chemical potential μ (the arithmetic mean of HOMO and LUMO energies).

All these descriptors (except δ) are energy-related properties and relate to: (1) the steric and electronic complementarity with the active site (Glide pre-reaction score), (2) the electrophilicity of the carbamoyl group (δ, μ, ν_C=O_, maxESP), or (3) to the thermodynamics of the nucleophilic addition to the carbamoyl group (ΔH_INT_, ΔG_INT_, ΔH_ALL_). These three seem to be the major factors determining the reactivity of the carbamoyl warhead and provide a meaningful insight into the underlying chemical driving forces.

Further multiple linear regression or QSAR approaches trying to mathematically relate these descriptors with experimentally observed inhibitory activity (pIC_50_) were unsuccessful. A holistic system integration that would connect the intrinsic reactivity of the warhead, binding affinity of the ligand, and free energy barrier of the covalent binding reaction to fully explain the presence or absence of covalent inhibition is currently not within the realm of possible – a thorough “wet-lab” analysis and use of complementary techniques is needed to accurately describe each individual case, as the example in the next section nicely demonstrates.

### 4 A peculiar case of compound 28 *–* what difference can one atom make?

Covalent inhibition presents an intricate combination of several factors that cannot be described by a simple equation: Apart from certain “rules of thumb”, each case is unique and requires a thorough experimental and computational investigation. To further illustrate this point, let’s take a deep dive into the peculiar case of compound **28**.

Compound **28** inhibited hBChE in a time-dependent manner (Table 2) and its covalent mechanism of binding was further corroborated by kinetic evidence: after a reversible binding step with nanomolar affinity, Ser198 carbamoylation took place with reaction free energy barrier (ΔG^‡^) determined at 19.6 kcal/mol (Table 3). Unequivocally, carbamoylation of hBChE’s catalytic Ser198 was also confirmed by soaking hBChE crystals with **28**, whereupon a crystal structure with **28**-carbamoylated Ser198 (Fig. 2b) was obtained. A second, distinct crystal structure was also obtained that did not present continuous electron density with the Ser198-*O*^γ^ atom and was in accordance with the decarbamoylated and decarboxylated **28**-derived secondary amine in the hBChE active site (Fig. S12), which could likely result from degradation of **28** prior to or upon soaking (cf. Table S10, half-life in PBS: 72 h).

**Table 3.** Kinetic parameters for hBChE inhibition by compound **28**, values are given as means ± SE.*^a^*.

| compound | **28** |
| --- | --- |
| **Reversible binding** |  |
| $K_i$ [nM] | $46.1 \pm 5.0$ |
| **Carbamoylation** |  |
| $k_{\text{carb}}$ [$\text{M}^{-1} \text{min}^{-1}$] | $1.54 \pm 0.32$ |
| $\Delta G_{25^\circ\text{C}}^\ddagger$ [kcal/mol] | $19.6 \pm 0.1$ |
| **Decarbamylation** |  |
| $k_{\text{decarb}}$ [$\text{h}^{-1}$] | $(6.95 \pm 0.68) \times 10^{-4}$ |
| $t_{1/2}$ [days] | $60.0 \pm 5.8^b$ |
| $\Delta G_{25^\circ\text{C}}^\ddagger$ [kcal/mol] | $26.6 \pm 0.1^b$ |

<sup>a</sup>By initial velocity measurements (Fig. S9).
<sup>b</sup>An estimated value since complete reactivation was not observed.

Importantly, the decarbamoylation step was found to be extremely slow, even slower than target turnover – the inhibited enzyme did not fully recover even after several months. The estimated half-life of **28**-carbamoylated hBChE is two months (Table 3), making compound **28** *de facto* an irreversible inhibitor, since the half-life of plasma hBChE is only 12 days.^22^ Beside one example of immeasurable reactivation on electric eel AChE,^81^ this is a unique case when a *carbamoyl-bearing* ChEI turned out to be an *irreversible*, rather than a pseudo-irreversible inhibitor.

However, when comparing **28** with its close analogue, a non-covalent inhibitor **26** with imidazole LG, it was unclear how merely a *carbon-nitrogen* substitution in the leaving group (to 1,2,4-triazole LG in **28**) could make such a difference – especially since there was only a slight difference in predicted reaction energetics [DFT-predicted activation barriers (ΔG^‡^) in solution were 15.0 and 14.3 kcal/mol, respectively. Meanwhile, ΔG_rxn1_ up to **INT** was slightly exergonic only for **28**, while the corresponding ΔG_rxn2_ up to **END** (Scheme 3) were exergonic in both cases (Tables S7, S9)]. To clarify this case with only subtle structural differences in the LG moieties, multiscale QM/MM calculations were pursued to consider the effects of the environment inside the protein, as well.

Two potential energy surface (PES) scanning QM/MM calculations were employed, starting from the optimized pre-reaction non-covalent complexes of hBChE with compounds **26** and **28**, respectively, and using the reaction coordinate driving method.^82^ The carbamoylation of catalytic Ser198 by compound **26** was unsuccessful – as evident from the corresponding potential energy curve (Fig. 3a, in light green), upon the gradual approach of the Ser198’s hydroxyl to the carbonyl *C_INH_* carbon of compound **26** the total energy of the system kept rising, which implies a failed bonding attempt, explaining its experimentally determined non-covalent inhibition. On the other hand, the nucleophilic attack of catalytic Ser198 on the carbonyl *C_INH_* carbon of compound **28** proceeded successfully with a free energy barrier ΔG^‡^ of 19.3 kcal/mol (Fig. 3a in teal and 3c), which is in excellent agreement with the experimentally determined value of 19.6 kcal/mol (Table 3).

**Fig. 3.**
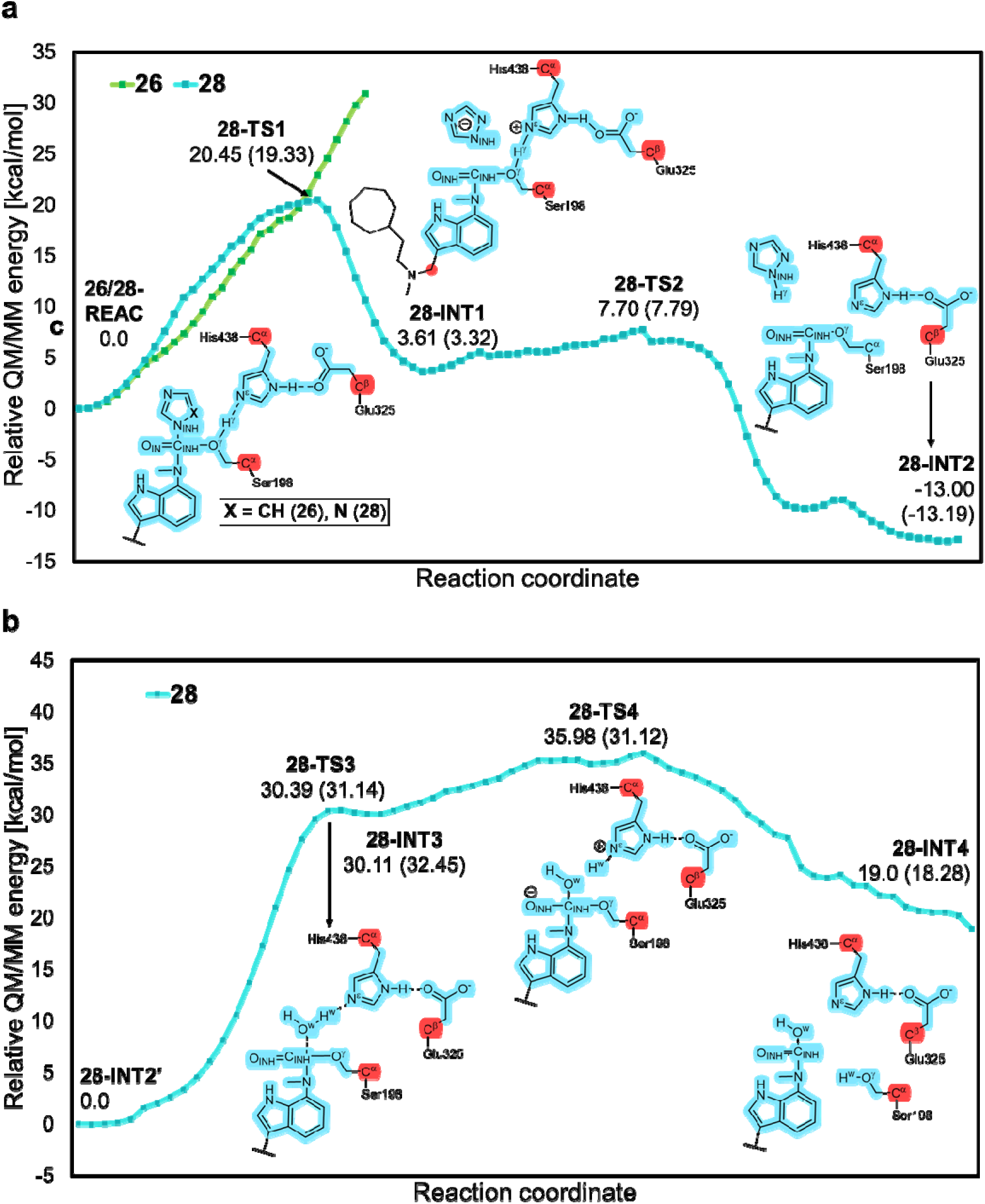
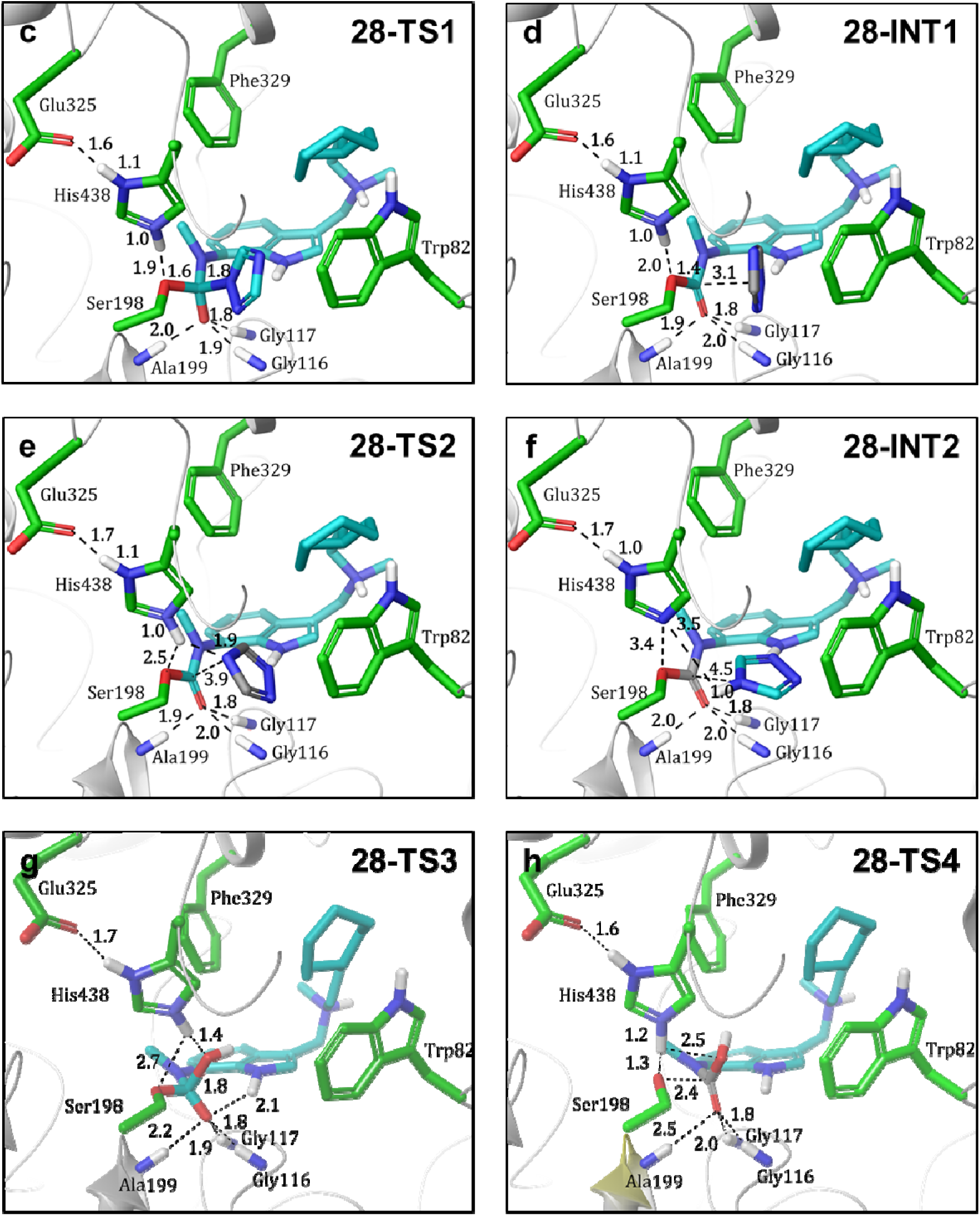
Multiscale QM/MM investigation of reaction of compounds 26 and 28 with hBChE. **a**, The potential energy profile for the carbamoylation of Ser198 by **26** (green) and **28** (teal). The energies were calculated at the B3LYP-D3/6-311G++**:AMBER//B3LYP/6-31G+*:AMBER level of theory. Values in the parentheses are relative Gibbs free energies obtained by the QM/MM total energy plus the Gibbs free energy correction to the QM region. QM atoms are highlighted blue, the boundary carbon atoms are highlighted red, and the MM region is coloured black. Reaction coordinates are defined in *Methods*. **b**, The potential energy profile for the decarbamoylation of carbamoylated **28**-hBChE by a water molecule. **c**, A snapshot of **28-TS1** stationary point along the carbamoylation pathway of compound **28** with hBChE. Amino acid residues are shown as green sticks, **28-**derived moieties are shown as cyan sticks. Key interatom distances (in Å) are shown as labelled black dashed lines. **d**, A snapshot of **28-INT1** stationary point. **e**, A snapshot of **28-TS2** stationary point. **f**, A snapshot of **28-INT2** stationary point. **g**, A snapshot of **28-TS3** stationary point. **h**, A snapshot of **28-TS4** stationary point.

This barrier difference probably arises from distinct electronic effects in the leaving groups of compounds **28** *vs*. **26**. The triazole LG of compound **28** has two »pyridinic« N atoms, which cause a strong electron withdrawal by: (1) resonance stabilisation, which increases the stability of the conjugate base of the LG, (2) inductive polarisation of the carbonyl carbon–LG bond, and (3) acidity amplification, which stabilises the anion departure. Conversely, the imidazole LG of compound **26** with only one »pyridinic« nitrogen exhibits a weaker electron-withdrawing character, a higher carbonyl carbon–LG electron density and bond strength, which effectively elevates the reaction barrier. The energy difference >11.7 kcal/mol translates to a reaction rate ratio of *k***_28_**/*k***_26_** > 3.6 ×10^8^ at 298.15 K, meaning that compound **28** reacts at least 360 million times faster than **26**. This extreme kinetic disparity conclusively explains why the reaction of compound **26** remains experimentally undetectable.

Furthermore, in contrast with QM calculations in solution, no stable tetrahedral intermediate (e.g. structure **INT** from Scheme 3) was observed in QM/MM, as the reaction took place in a concerted way: the 1,2,4-triazole anion was liberated concomitantly with *O*^γ^ of Ser198 approaching the ligand’s carbonyl carbon, leading to an intermediate **28-INT1** (Fig. 3d). The latter progressed *via* a **28-TS2** transition state (Fig. 3e) corresponding to the proton transfer from protonated His438 to 1,2,4-triazole anion, to **28**-carbamoylated Ser198 state **28-INT2** (Fig. 3f).

As expected, the acidic 1,2,4-triazole leaving group did not remain bound in the active site but eventually exited towards bulk solvent, as was observed during 1 µs MD simulation of **28-INT2** (Fig. S15). Therefore, **28-INT2**′ state was constructed by removing the 1,2,4-triazole molecule from the structure of **28-INT2** and relaxing the system by performing a MD simulation. Afterwards, a water molecule close to the *C_INH_* was selected as the nucleophile, and another QM/MM reaction coordinate study was initiated, trying to locate the minimum energy pathway for the decarbamoylation of **28-INT2**′. First, nucleophilic attack of a water molecule on the carbonyl carbon *C_INH_* had to surmount a high activation barrier of 31.1 kcal/mol (**28-TS3**, Fig. 3b and 3g), followed by collapse of the tetrahedral intermediate into carbamic acid and catalytic Ser198, which was regenerated by simultaneous proton transfer from ionized His438 (**28-TS4** to **28-END**, Fig. 3b and 3h). This roughly agrees with the extremely slow, but measurable decarbamoylation observed *in vitro* (estimated ΔG^‡^ = 26.l kcal/mol, 60 days half-life).

Although the goal of systematically integrating all computational and experimental data into a single model for predicting the reactivity of covalent inhibitors is conceptually appealing, it remains a major challenge in practice. The limitations in computational feasibility (e.g., time-consuming QM/MM reaction pathway calculations), the lack of reliable binding affinity predictions by docking or MD alone, and the scarcity of generalisable QM descriptors for reactivity currently prevent such unification. Our exploration of the peculiar case of compound **28** highlights the need for multiple, complementary methods to capture the nuanced interplay of structural and electronic factors that govern the complex world of covalent inhibition.

### 5 Reactivity of the carbamate warhead at the proteome level

Ultimately, it is important to take a step back and consider the broader biological context beyond the role of cholinesterases. Comparing small nucleofuges from sections 2.1–2.4, although interesting from a scientific viewpoint, carbamoyl halides are probably too reactive and unstable to be of use in chemical biology and medicinal chemistry. Carbamoyl thiols and selenols did not inhibit ChEs, unless an aromatic and highly acidic nucleofuge (such as 1-methyl-1*H*-tetrazole-5-thiol in compound **23**) was used, rendering the inhibitor prone to hydrolytic decomposition (Table S10). Carbamoylazoles are already known as inhibitors of various SHs and compounds **27**–**31** were found to be quite unstable both in solution (Table S10) and as solid compounds. Therefore, we turned to *O*-aryl carbamates, which have been already known as ChE inhibitors. Here, the 3-isooxazolyl leaving group (e.g., as in compound **12**) was located in the sweet spot between reactivity, selectivity, and stability. Incorporation of the carbamoyl warhead with this leaving group onto other scaffolds could endow them with pseudo-irreversible cholinesterase inhibition. However, as different *O*-carbamates have been reported to inhibit various SHs across the proteome, and some have been used as ABPP probes,^26^ the off-target selectivity of *O*-carbamoyl isoxazol-3-ols needed to be investigated.

Accordingly, 3-isoxazolyl carbamoyl warheads were connected *via* a long linker to either a rhodamine B derivative^30^ (**P** probes, Fig. 4a) or biotin (**B** probes, Fig. 4a), enabling fluorescent detection of covalently modified proteins or pull-down using (strept)avidin, respectively. Both the minimal (only *N*-methyl carbamate) probes **P3** and **B3**, as well as the “whole inhibitor”, *N*-(3-(((2-cycloheptylethyl)(methyl)amino)methyl)-1*H*-indol-7-yl)-containing probe **P4**, and *O*-phenyl carbamate probe **P5** were synthesised and evaluated. The probes **P1**–**P5** were incubated with human hepatocellular carcinoma HepG2 cell lysate (Fig. 4b) and human neuroblastoma SH-SY5Y cell lysate (Fig. 4c), followed by protein separation using SDS-PAGE and transfer to PVDF membranes by western blot to improve fluorescent detection. Heat-inactivated control samples^83^ and comparison with warhead-less probe **P2**-treated samples were used to account for non-specific labelling.

**Fig. 4.**
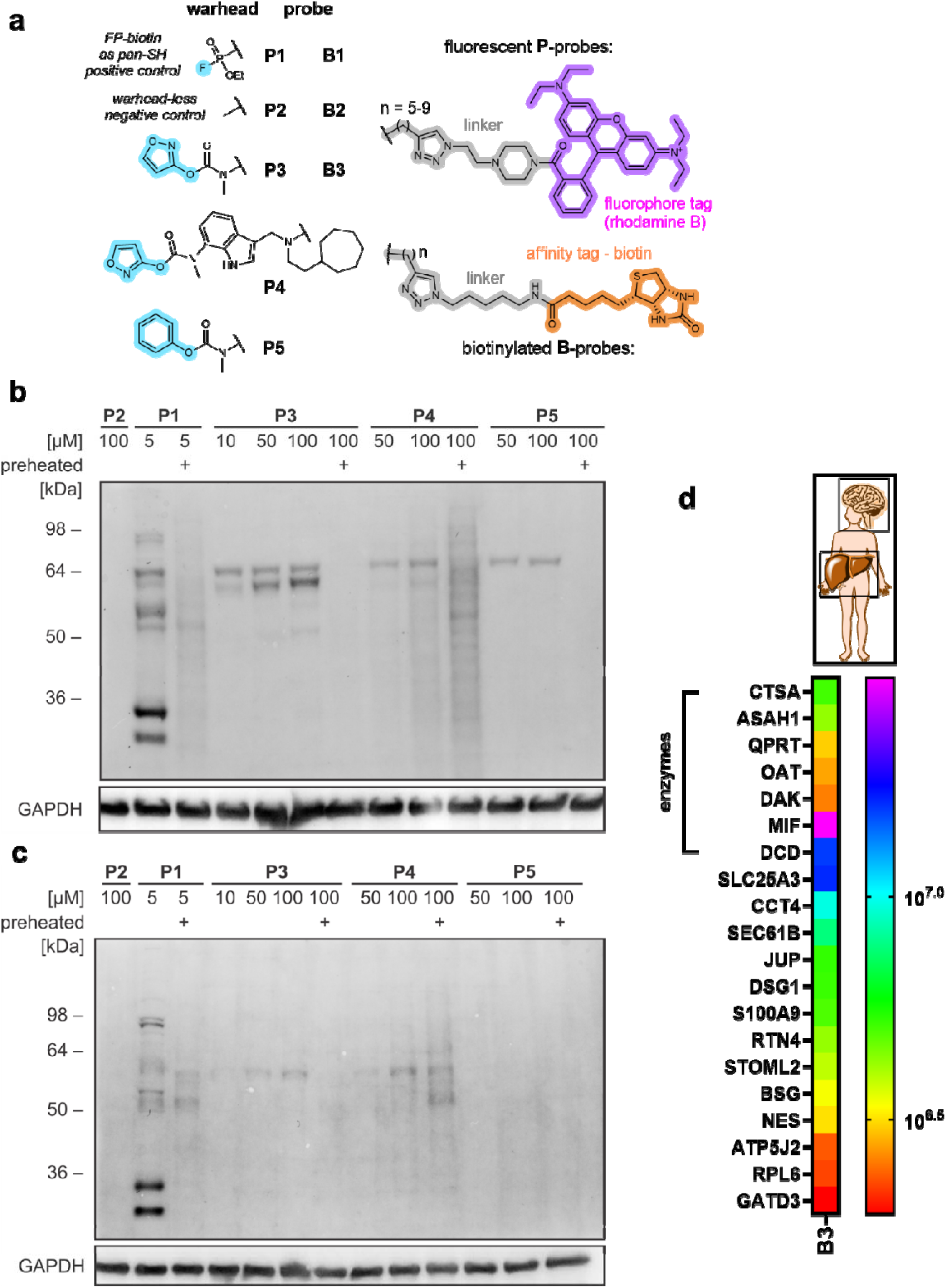
*O*-(hetero)aryl carbamate ABPP probes and their proteome-wide interactions. **a**, Synthesised fluorescent **P** and biotinylated **B** ABPP probes with carbamoyl warheads, a broad-spectrum fluorophosphonate probes **P1** and **B1**, and warhead-less probes **P2** and **B2** as negative controls. **b**–**c**, Western blot membranes following SDS-PAGE separation of **P** probe-treated proteins analysed by fluorescent detection at 595 nm. Unspecific binding is accounted for in heat-inactivated control samples (labelled “preheated”) and samples treated with warhead-less probe **P2**. **b**, Probe comparison on HepG2 lysate. **c**, Probe comparison on SH-SY5Y lysate. **d**, A heatmap of human proteins from neuroblastoma SH-SY5Y and hepatocellular carcinoma HepG2 cell lines that were labelled with biotinylated probe **B3** and identified by LC-MS/MS (Table S4). The intensities ratios that satisfied the inclusion criterion (>10-fold enrichment *vs.* warhead-less probe **B2**) are shown in the log rainbow scale. The full list of identified proteins and raw data can be found in Supplementary Materials.

Probe **P3** activity-dependently labelled three major bands in HepG2 lysate (Fig. 4b), two of which were also labelled by the positive control (a pan-SH fluorophoshonate FP-rhodamine probe **P1**). The band around 64 kDa was detected by probes **P4** and **P5**, as well. The “whole inhibitor” probe **P4** displayed significant non-specific binding, observed in the heat-treated sample. Meanwhile, only one weak band was detected by probes **P3** and **P4** in SH-SY5Y lysate (Fig. 4c). *O*-Phenyl probe **P5** did not appear to label any SH-SY5Y proteins, while the positive control FP-rhodamine **P1** labelled several bands. This points to different reactivity of *O*-aryl and *O*-*hetero*aryl carbamate warheads, with the latter being more reactive not only in terms of warhead electrophilicity but also proteomic reactivity. Furthermore, probes **P1–P3** were incubated with recombinant hBChE and human plasma. For unknown reasons, the successful, activity-dependent binding of **P1** and **P3** was observed for recombinant hBChE, but not for human plasma, which also contains hBChE (Fig. S18).

However, since the number and pattern of fluorescent bands on an SDS-PAGE gel offers only qualitative information about the labelled proteins, unequivocal MS identification of the labelled proteins was sought. To this effect, biotinylated probe **B3** (Fig. 4a) was reacted with mixed human cell lysate (from both SH-SY5Y and HepG2 lines, to ensure a better coverage of the human proteome). The bound proteins were affinity-enriched on an avidin resin, digested, and analysed by LC-MS/MS. The identified proteins were filtered to remove potential contaminants from sample preparation (keratins, HSA, filaggrin, hornerin, etc.), and only the proteins that were >10-fold enriched in the sample incubated with probe **B3** compared to the warhead-less probe **B2** were retained (Fig. 4d, Table S4). In total, 20 proteins were identified, and 7 of them possessed enzymatic activity – lysosomal protective protein CTSA was the only SH, while acid ceramidase ASAH1 was the only Cys hydrolase identified. Among these 20 proteins, only S100-A9 was also picked-up by FP-biotin probe **B1** (data not shown). Neither of ChEs was detected, which could be the result of enzymes’ degradation in the lysate or their low expression – based on literature evidence, the latter seems to be the case.^84,85^

## Conclusion

We presented a comprehensive structure-*reactivity* study of the carbamoyl-bearing warheads and showed how the nature of the leaving group determines covalent inhibition. In a case study on ChEs, we explored the requirements for an ideal leaving group to tune the reactivity of the carbamoyl warhead. A wide spectrum of halogen-, chalcogen- and nitrogen-based nucleofuges of various sizes, with or without aromatic character, were evaluated and a few general “rules of thumb” for nucleofugality were deduced, although a combination of different factors, both favourable and opposing, are in play to influence and confound the binding affinity of each inhibitor. Namely, besides sufficient acidity of the nucleofuge, an aromatic moiety that can delocalise the liberated electron pair and/or form additional stabilising interactions in the active site is another structural element that clearly favours covalent binding. Rudimentary intrinsic chemical reactivity and spatial complementarity could be inferred from the “rough and cheap” DFT calculations in solution combined with molecular docking and dynamics, respectively. However, only multiscale QM/MM studies in the context of topology and electrostatics of the enzymatic environment were able to address the subtle differences in the reactivity of the close analogues **26** and **28**, demonstrating how the substitution of a single atom significantly affected the inhibition mechanism. Namely, the triazole LG of compound **28** has an additional »pyridinic« nitrogen that exerts strong electron withdrawal through resonance stabilisation, inductive bond polarisation, and acidity amplification, thereby lowering the reaction barrier and accelerating the reaction by more than 360 million times compared to compound **26**.

The goal of systematically integrating all computational and experimental data into a single model to predict the reactivity of covalent inhibitors is conceptually appealing, but poses a major challenge in practice. The limitations in computational feasibility (e.g. time-consuming QM/MM reaction pathway calculations), the lack of reliable binding affinity predictions by docking or MD alone, and the scarcity of generalisable QM descriptors for reactivity currently prevent such unification. Instead, our results highlight the need for multiple, complementary methods to capture the nuanced interplay of structural and electronic factors that govern the complex world of covalent inhibition.

Furthermore, by enlarging the *N*-carbamoyl moiety, the decarbamoylation process was slowed down to such an extent that the hBChE inhibition by otherwise pseudo-irreversible carbamoyl-bearing inhibitor became *de facto* irreversible. Looking at the bigger picture beyond ChEs, the off-targets of the promising *O*-isoxazol-3-yl carbamate warhead were discovered by pull-down of a biotinylated **B3** probe and LC-MS/MS identification. As this warhead demonstrated a relatively low off-target binding in the proteome, its incorporation into unrelated reversible ligand scaffolds may be ideal for conferring pseudo-irreversible ChE inhibitory activity to them. Overall, our comprehensive paper has demonstrated the tunable reactivity of the carbamoyl warhead and presented a wide spectrum of available leaving groups that could be further developed into ABPP probes for different SHs.

## Acknowledgments

This research was funded by the Slovenian Research and Innovation Agency (ARIS), Research Core Funding № P1-0208, P1-0012, P4-0127, P4-0053, P1-0230, bilateral travel grant BI-US/22-24-004 (granted to D.K.), and a young researcher grant to A.M. The authors would like to thank the Ažman Computing Centre at National Institute of Chemistry, Slovenia for computational resources, and Assoc. Prof. dr. Igor Locatelli (UL FFA) for help with statistical analysis. FC, FM, MD, JD, FN, and XB were supported by the French Ministry of Armed Forces (Direction Générale de l’Armement and Service de Santé des Armées; grant NBC-5-C-2316), and acknowledge the ESRF for the provision of synchrotron radiation facilities during long-term beamline access (IBS BAG), and would like to thank the ID23-1 beamline staff for their support.

## Data Availability

All the necessary data that support the findings of this study are included in the Supplementary Information. Source Data are available on figshare repository under DOI: 10.6084/m9.figshare.29458835. The mass spectrometry proteomics data have been deposited to the ProteomeXchange Consortium via the PRIDE partner repository with the dataset identifier PXD065683. The crystallography data were deposited on the PDB database under accession codes 8QTW and 8QTX. Additional raw data are available from the corresponding author upon reasonable request.

## References

1. Kam, C. M., Abuelyaman, A. S., Li, Z., Hudig, D. & Powers, J. C. Biotinylated isocoumarins, new inhibitors and reagents for detection, localization, and isolation of serine proteases. Bioconjug. Chem. 4, 560–567 (1993).

2. Fang, H., Peng, B., Ong, S. Y., Wu, Q., Li, L. & Yao, S. Q. Recent advances in activity-based probes (ABPs) and affinity-based probes (AfBPs) for profiling of enzymes. Chem. Sci. 12, 8288–8310 (2021).

3. Cravatt, B. F. Activity-Based Protein Profiling – Finding General Solutions to Specific Problems. Isr. J. Chem. 63, e202300029 (2023).

4. Evans, M. J. & Cravatt, B. F. Mechanism-Based Profiling of Enzyme Families. Chem. Rev. 106, 3279–3301 (2006).

5. Bachovchin, D. A., Ji, T., Li, W., Simon, G. M., Blankman, J. L., Adibekian, A., Hoover, H., Niessen, S. & Cravatt, B. F. Superfamily-wide portrait of serine hydrolase inhibition achieved by library-versus-library screening. Proc. Natl. Acad. Sci. 107, 20941–20946 (2010).

6. Adibekian, A., Martin, B. R., Chang, J. W., Hsu, K.-L., Tsuboi, K., Bachovchin, D. A., Speers, A. E., Brown, S. J., Spicer, T., Fernandez-Vega, V., Ferguson, J., Hodder, P. S., Rosen, H. & Cravatt, B. F. Confirming Target Engagement for Reversible Inhibitors in Vivo by Kinetically Tuned Activity-Based Probes. J. Am. Chem. Soc. 134, 10345–10348 (2012).

7. Kok, B. P., Ghimire, S., Kim, W., Chatterjee, S., Johns, T., Kitamura, S., Eberhardt, J., Ogasawara, D., Xu, J., Sukiasyan, A., Kim, S. M., Godio, C., Bittencourt, J. M., Cameron, M., Galmozzi, A., Forli, S., Wolan, D. W., Cravatt, B. F., Boger, D. L. & Saez, E. Discovery of small-molecule enzyme activators by activity-based protein profiling. Nat. Chem. Biol. 16, 997–1005 (2020).

8. Deng, H., Lei, Q., Wu, Y., He, Y. & Li, W. Activity-based protein profiling: Recent advances in medicinal chemistry. Eur. J. Med. Chem. 191, 112151 (2020).

9. Abbasov, M. E., Kavanagh, M. E., Ichu, T.-A., Lazear, M. R., Tao, Y., Crowley, V. M., am Ende, C. W., Hacker, S. M., Ho, J., Dix, M. M., Suciu, R., Hayward, M. M., Kiessling, L. L. & Cravatt, B. F. A proteome-wide atlas of lysine-reactive chemistry. Nat. Chem. 13, 1081–1092 (2021).

10. Gonzalez-Valero, A., Reeves, A. G., Page, A. C. S., Moon, P. J., Miller, E., Coulonval, K., Crossley, S. W. M., Xie, X., He, D., Musacchio, P. Z., Christian, A. H., McKenna, J. M., Lewis, R. A., Fang, E., Dovala, D., Lu, Y., McGregor, L. M., Schirle, M., Tallarico, J. A., Roger, P. P., Toste, F. D. & Chang, C. J. An Activity-Based Oxaziridine Platform for Identifying and Developing Covalent Ligands for Functional Allosteric Methionine Sites: Redox-Dependent Inhibition of Cyclin-Dependent Kinase 4. J. Am. Chem. Soc. 144, 22890–22901 (2022).

11. Wright, A. T., Song, J. D. & Cravatt, B. F. A Suite of Activity-Based Probes for Human Cytochrome P450 Enzymes. J. Am. Chem. Soc. 131, 10692–10700 (2009).

12. Matthews, M. L., He, L., Horning, B. D., Olson, E. J., Correia, B. E., Yates, J. R., Dawson, P. E. & Cravatt, B. F. Chemoproteomic profiling and discovery of protein electrophiles in human cells. Nat. Chem. 9, 234–243 (2017).

13. Bachovchin, D. A. & Cravatt, B. F. The pharmacological landscape and therapeutic potential of serine hydrolases. Nat. Rev. Drug Discov. 11, 52–68 (2012).

14. Adibekian, A., Martin, B. R., Wang, C., Hsu, K.-L., Bachovchin, D. A., Niessen, S., Hoover, H. & Cravatt, B. F. Click-generated triazole ureas as ultrapotent in vivo –active serine hydrolase inhibitors. Nat. Chem. Biol. 7, 469–478 (2011).

15. Otrubova, K., Chatterjee, S., Ghimire, S., Cravatt, B. F. & Boger, D. L. N-Acyl pyrazoles: Effective and tunable inhibitors of serine hydrolases. Bioorg. Med. Chem. 27, 1693–1703 (2019).

16. Faucher, F., Bennett, J. M., Bogyo, M. & Lovell, S. Strategies for Tuning the Selectivity of Chemical Probes that Target Serine Hydrolases. Cell Chem. Biol. 27, 937–952 (2020).

17. Jöst, C., Nitsche, C., Scholz, T., Roux, L. & Klein, C. D. Promiscuity and Selectivity in Covalent Enzyme Inhibition: A Systematic Study of Electrophilic Fragments. J. Med. Chem. 57, 7590–7599 (2014).

18. Gilbert, K. E., Vuorinen, A., Aatkar, A., Pogány, P., Pettinger, J., Grant, E. K., Kirkpatrick, J. M., Rittinger, K., House, D., Burley, G. A. & Bush, J. T. Profiling Sulfur(VI) Fluorides as Reactive Functionalities for Chemical Biology Tools and Expansion of the Ligandable Proteome. ACS Chem. Biol. 18, 285–295 (2023).

19. Barrow, A. S., Smedley, C. J., Zheng, Q., Li, S., Dong, J. & Moses, J. E. The growing applications of SuFEx click chemistry. Chem. Soc. Rev. 48, 4731–4758 (2019).

20. Carneiro, S. N., Khasnavis, S. R., Lee, J., Butler, T. W., Majmudar, J. D., Ende, C. W. am & Ball, N. D. Sulfur(VI) fluorides as tools in biomolecular and medicinal chemistry. Org. Biomol. Chem. 21, 1356–1372 (2023).

21. Xing, S., Li, Q., Xiong, B., Chen, Y., Feng, F., Liu, W. & Sun, H. Structure and therapeutic uses of butyrylcholinesterase: Application in detoxification, Alzheimer’s disease, and fat metabolism. Med. Res. Rev. 41, 858–901 (2021).

22. Lockridge, O. Review of human butyrylcholinesterase structure, function, genetic variants, history of use in the clinic, and potential therapeutic uses. Pharmacol. Ther. 148, 34–46 (2015).

23. Tuin, A. W., Mol, M. A. E., van den Berg, R. M., Fidder, A., van der Marel, G. A., Overkleeft, H. S. & Noort, D. Activity-Based Protein Profiling Reveals Broad Reactivity of the Nerve Agent Sarin. Chem. Res. Toxicol. 22, 683–689 (2009).

24. Blankman, J. L., Simon, G. M. & Cravatt, B. F. A Comprehensive Profile of Brain Enzymes that Hydrolyze the Endocannabinoid 2-Arachidonoylglycerol. Chem. Biol. 14, 1347–1356 (2007).

25. Planas-Marquès, M., Bernardo-Faura, M., Paulus, J., Kaschani, F., Kaiser, M., Valls, M., van der Hoorn, R. A. L. & Coll, N. S. Protease Activities Triggered by Ralstonia solanacearum Infection in Susceptible and Tolerant Tomato Lines*. Mol. Cell. Proteomics 17, 1112–1125 (2018).

26. Chang, J. W., Cognetta, A. B., Niphakis, M. J. & Cravatt, B. F. Proteome-Wide Reactivity Profiling Identifies Diverse Carbamate Chemotypes Tuned for Serine Hydrolase Inhibition. ACS Chem. Biol. 8, 1590–1599 (2013).

27. Long, J. Z., Jin, X., Adibekian, A., Li, W. & Cravatt, B. F. Characterization of Tunable Piperidine and Piperazine Carbamates as Inhibitors of Endocannabinoid Hydrolases. J. Med. Chem. 53, 1830–1842 (2010).

28. Cognetta, A. B., Niphakis, M. J., Lee, H.-C., Martini, M. L., Hulce, J. J. & Cravatt, B. F. Selective N-Hydroxyhydantoin Carbamate Inhibitors of Mammalian Serine Hydrolases. Chem. Biol. 22, 928–937 (2015).

29. Broeckaert, L., Moens, J., Roos, G., Proft, F. D. & Geerlings, P. Intrinsic Nucleofugality Scale within the Framework of Density Functional Reactivity Theory. J. Phys. Chem. A 112, 12164–12171 (2008).

30. Meden, A., Knez, D., Brazzolotto, X., Modeste, F., Perdih, A., Pišlar, A., Zorman, M., Zorović, M., Denic, M., Pajk, S., Živin, M., Nachon, F. & Gobec, S. Pseudo-irreversible butyrylcholinesterase inhibitors: Structure–activity relationships, computational and crystallographic study of the N-dialkyl O-arylcarbamate warhead. Eur. J. Med. Chem. 247, 115048 (2023).

31. Copeland, R. A. in Eval. Enzyme Inhib. Drug Discov. 345–382 (John Wiley & Sons, Ltd, 2013). doi:10.1002/9781118540398.ch9

32. Thorarensen, A., Balbo, P., Banker, M. E., Czerwinski, R. M., Kuhn, M., Maurer, T. S., Telliez, J.-B., Vincent, F. & Wittwer, A. J. The advantages of describing covalent inhibitor in vitro potencies by IC50 at a fixed time point. IC50 determination of covalent inhibitors provides meaningful data to medicinal chemistry for SAR optimization. Bioorg. Med. Chem. 29, 115865 (2021).

33. Groner, E., Ashani, Y., Schorer-Apelbaum, D., Sterling, J., Herzig, Y. & Weinstock, M. The Kinetics of Inhibition of Human Acetylcholinesterase and Butyrylcholinesterase by Two Series of Novel Carbamates. Mol. Pharmacol. 71, 1610–1617 (2007).

34. Jutzi, P. New Element-Carbon (p-p)π Bonds. Angew. Chem. Int. Ed. Engl. 14, 232–245 (1975).

35. Meden, A., Knez, D., Brazzolotto, X., Nachon, F., Dias, J., Svete, J., Stojan, J., Grošelj, U. & Gobec, S. From tryptophan-based amides to tertiary amines: Optimization of a butyrylcholinesterase inhibitor series. Eur. J. Med. Chem. 234, 114248 (2022).

36. Grzyb, J. A., Shen, M., Yoshina-Ishii, C., Chi, W., Brown, R. S. & Batey, R. A. Carbamoylimidazolium and thiocarbamoylimidazolium salts: novel reagents for the synthesis of ureas, thioureas, carbamates, thiocarbamates and amides. Tetrahedron 61, 7153–7175 (2005).

37. Bordwell, F. G. Equilibrium acidities in dimethyl sulfoxide solution. Acc. Chem. Res. 21, 456–463 (1988).

38. Knunyants, I. L., Dyatkin, B. L., Mochalina, E. P. & Lantseva, L. T. Hexafluoroisopropylhydroxylamine and dissociation constants of some fluorinated hydroxylamines and oximes. Bull. Acad. Sci. USSR Div. Chem. Sci. 15, 164–165 (1966).

39. Filler, R. & Schure, R. M. Highly acidic perhalogenated alcohols. A new synthesis of perfluoro-tert-butyl alcohol. J. Org. Chem. 32, 1217–1219 (1967).

40. Kütt, A., Leito, I., Kaljurand, I., Sooväli, L., Vlasov, V. M., Yagupolskii, L. M. & Koppel, I. A. A Comprehensive Self-Consistent Spectrophotometric Acidity Scale of Neutral Brønsted Acids in Acetonitrile. J. Org. Chem. 71, 2829–2838 (2006).

41. Jørgensen, C. G., Bräuner-Osborne, H., Nielsen, B., Kehler, J., Clausen, R. P., Krogsgaard-Larsen, P. & Madsen, U. Novel 5-substituted 1-pyrazolol analogues of ibotenic acid: Synthesis and pharmacology at glutamate receptors. Bioorg. Med. Chem. 15, 3524–3538 (2007).

42. Stensbøl, T. B., Uhlmann, P., Morel, S., Eriksen, B. L., Felding, J., Kromann, H., Hermit, M. B., Greenwood, J. R., Braüner-Osborne, H., Madsen, U., Junager, F., Krogsgaard-Larsen, P., Begtrup, M. & Vedsø, P. Novel 1-Hydroxyazole Bioisosteres of Glutamic Acid. Synthesis, Protolytic Properties, and Pharmacology. J. Med. Chem. 45, 19–31 (2002).

43. Boyle, F. T. & Jones, R. A. Y. Azole N-oxides. Part I. The tautomerism of benzotriazole 1-oxide and its 4- and 6-nitro-derivatives with the corresponding 1-hydroxybenzotriazoles. J. Chem. Soc. Perkin Trans. 2 160–164 (1973). doi:10.1039/P29730000160

44. Ames, D. E. & Grey, T. F. The synthesis of some N-hydroxyimides. J. Chem. Soc. Resumed 631–636 (1955). doi:10.1039/JR9550000631

45. Di Bussolo, V., Caselli, M., Romano, M. R., Pineschi, M. & Crotti, P. Regio- and Stereoselectivity of the Addition of O-, S-, N-, and C-Nucleophiles to the β Vinyl Oxirane Derived from d-Glucal. J. Org. Chem. 69, 8702–8708 (2004).

46. DeCollo, T. V. & Lees, W. J. Effects of Aromatic Thiols on Thiol−Disulfide Interchange Reactions That Occur during Protein Folding. J. Org. Chem. 66, 4244–4249 (2001).

47. Hanlon, D. P. & Shuman, S. Copper ion binding and enzyme inhibitory properties of the antithyroid drug methimazole. Experientia 31, 1005–1006 (1975).

48. Narisada, M., Terui, Y., Yamakawa, M., Watanabe, F., Ohtani, M. & Miyazaki, H. Thiol-disulfide exchange reactions of bis(methyl-1H-tetrazol-5-yl) disulfide studied by proton nuclear magnetic resonance spectroscopy. J. Org. Chem. 50, 2794–2796 (1985).

49. Sonoda, N. & Ogawa, A. in Encycl. Reag. Org. Synth. (John Wiley & Sons, Ltd, 2005). doi:10.1002/047084289X.rb018

50. Walba, H. & Isensee, R. W. Acidity Constants of Some Arylimidazoles and Their Cations. J. Org. Chem. 26, 2789–2791 (1961).

51. Murłowska, K. & Sadlej-Sosnowska, N. Absolute Calculations of Acidity of C-Substituted Tetrazoles in Solution. J. Phys. Chem. A 109, 5590–5595 (2005).

52. Han, X., Balakrishnan, V. K., vanLoon, G. W. & Buncel, E. Degradation of the Pesticide Fenitrothion as Mediated by Cationic Surfactants and α-Nucleophilic Reagents. Langmuir 22, 9009–9017 (2006).

53. In Compr. Org. Name React. Reag. 9–12 (John Wiley & Sons, Ltd, 2010). doi:10.1002/9780470638859.conrr003

54. Lassalas, P., Gay, B., Lasfargeas, C., James, M. J., Tran, V., Vijayendran, K. G., Brunden, K. R., Kozlowski, M. C., Thomas, C. J., Smith, A. B. I., Huryn, D. M. & Ballatore, C. Structure Property Relationships of Carboxylic Acid Isosteres. J. Med. Chem. 59, 3183–3203 (2016).

55. Matošević, A., Radman Kastelic, A., Mikelić, A., Zandona, A., Katalinić, M., Primožič, I., Bosak, A. & Hrenar, T. Quinuclidine-Based Carbamates as Potential CNS Active Compounds. Pharmaceutics 13, 420 (2021).

56. Pejchal, V., Štěpánková, Š., Pejchalová, M., Královec, K., Havelek, R., Růžičková, Z., Ajani, H., Lo, R. & Lepšík, M. Synthesis, structural characterization, docking, lipophilicity and cytotoxicity of 1-[(1R)-1-(6-fluoro-1,3-benzothiazol-2-yl)ethyl]-3-alkyl carbamates, novel acetylcholinesterase and butyrylcholinesterase pseudo-irreversible inhibitors. Bioorg. Med. Chem. 24, 1560–1572 (2016).

57. Carolan, C. G., Dillon, G. P., Khan, D., Ryder, S. A., Gaynor, J. M., Reidy, S., Marquez, J. F., Jones, M., Holland, V. & Gilmer, J. F. Isosorbide-2-benzyl Carbamate-5-salicylate, A Peripheral Anionic Site Binding Subnanomolar Selective Butyrylcholinesterase Inhibitor. J. Med. Chem. 53, 1190–1199 (2010).

58. Gialih Lin, Gan-Hong Chen, & Hong-Chi Ho. Conformationally restricted carbamate inhibitors of horse serum butyrylcholinesterase. Bioorg. Med. Chem. Lett. 8, 2747–2750 (1998).

59. Kovářová, M., Komers, K., Štěpánková, Š. & Čegan, A. Inhibition of acetylcholinesterase by 14 achiral and five chiral imidazole derivates. Bioresour. Technol. 101, 6281–6283 (2010).

60. ExPASy - ProtParam tool. at <https://web.expasy.org/protparam/>

61. Verma, A., Wong, D. M., Islam, R., Tong, F., Ghavami, M., Mutunga, J. M., Slebodnick, C., Li, J., Viayna, E., Lam, P. C.-H., Totrov, M. M., Bloomquist, J. R. & Carlier, P. R. 3-Oxoisoxazole-2(3H)-carboxamides and isoxazol-3-yl carbamates: Resistance-breaking acetylcholinesterase inhibitors targeting the malaria mosquito, Anopheles gambiae. Bioorg. Med. Chem. 23, 1321–1340 (2015).

62. Lindgren, B., Lindgren, G., Artursson, E., Puu, G., Fredriksson, J. & Andersson, M. Acetylcholinesterase Inhibition by Sulphur and Selenium Heterosubstituted Isomers of N, N-Diethylcarbamyl Choline and Carbaryl. J. Enzym. Inhib. 1, 1–11 (1985).

63. Hegarty, A. F. & Frost, L. N. Isocyanate intermediates in Elcb mechanism of carbamate hydrolysis. J. Chem. Soc. Chem. Commun. 500–501 (1972). doi:10.1039/C39720000500

64. Staab, H. A. New Methods of Preparative Organic Chmistry IV. Syntheses Using Heterocyclic Amides (Azolides). Angew. Chem. Int. Ed. Engl. 1, 351–367 (1962).

65. Groutas, W. C., Abrams, W. R., Theodorakis, M. C., Kasper, A. M., Rude, S. A., Badger, R. C., Ocain, T. D., Miller, K. E. & Moi, M. K. Amino acid derived latent isocyanates: irreversible inactivation of porcine pancreatic elastase and human leukocyte elastase. J. Med. Chem. 28, 204–209 (1985).

66. Groutas, W. C., Brubaker, M. J., Zandler, M. E., Mazo-Gray, V., Rude, S. A., Crowley, J. P., Castrisos, J. C., Dunshee, D. A. & Giri, P. K. Inactivation of leukocyte elastase by aryl azolides and sulfonate salts. Structure-activity relationship studies. J. Med. Chem. 29, 1302–1305 (1986).

67. Rocha, J.-F., Santos, A., Gama, H., Moser, P., Falcão, A., Pressman, P., Wallace Hayes, A. & Soares-da-Silva, P. Safety, Tolerability, and Pharmacokinetics of FAAH Inhibitor BIA 10-2474: A Double-Blind, Randomized, Placebo-Controlled Study in Healthy Volunteers. Clin. Pharmacol. Ther. 111, 391–403 (2022).

68. van Esbroeck, A. C. M., Janssen, A. P. A., Cognetta, A. B., Ogasawara, D., Shpak, G., van der Kroeg, M., Kantae, V., Baggelaar, M. P., de Vrij, F. M. S., Deng, H., Allarà, M., Fezza, F., Lin, Z., van der Wel, T., Soethoudt, M., Mock, E. D., den Dulk, H., Baak, I. L., Florea, B. I., Hendriks, G., De Petrocellis, L., Overkleeft, H. S., Hankemeier, T., De Zeeuw, C. I., Di Marzo, V., Maccarrone, M., Cravatt, B. F., Kushner, S. A. & van der Stelt, M. Activity-based protein profiling reveals off-target proteins of the FAAH inhibitor BIA 10-2474. Science 356, 1084–1087 (2017).

69. Baker, M. W., Kerry, J. C., Kozlik, A., Marshall, J. R., Nichol, K. J. & Weighton, D. Imidazolderivate. (1973). at <https://patents.google.com/patent/DE2260025A1/de>

70. Baker, M. W., Kerry, J. C., Nichol, K. J., Marshall, J. R., Weighton, D. M. & Kozlik, A. 1-Carbonamido imidazoles. (1977). At <https://patents.google.com/patent/US4048188/en>

71. Kurahashi, Y., Shiokawa, K., Goto, T., Kagabu, S., Kamochi, A., Moriya, K. & Hayakawa, H. Tetrahydroquinolin-1-yl-carbonyl-imidazole derivatives. (1986). at <https://patents.google.com/patent/EP0173208A1/en>

72. Zhu, K., Borrelli, K. W., Greenwood, J. R., Day, T., Abel, R., Farid, R. S. & Harder, E. Docking Covalent Inhibitors: A Parameter Free Approach To Pose Prediction and Scoring. J. Chem. Inf. Model. 54, 1932–1940 (2014).

73. Wan, X., Yao, Y., Fang, L. & Liu, J. Unexpected protonation state of Glu197 discovered from simulations of tacrine in butyrylcholinesterase. Phys. Chem. Chem. Phys. 20, 14938–14946 (2018).

74. Zlobin, A., Smirnov, I. & Golovin, A. Dynamic interchange between two protonation states is characteristic of active sites of cholinesterases. Protein Sci. 33, e5100 (2024).

75. Hermann, M. R., Pautsch, A., Grundl, M. A., Weber, A. & Tautermann, C. S. Covalent inhibitor reactivity prediction by the electrophilicity index—in and out of scope. J. Comput. Aided Mol. Des. 35, 531–539 (2021).

76. Keeley, A., Ábrányi-Balogh, P. & Keserű, G. M. Design and characterization of a heterocyclic electrophilic fragment library for the discovery of cysteine-targeted covalent inhibitors. MedChemComm 10, 263–267 (2019).

77. Shokhen, M., Traube, T., Vijayakumar, S., Hirsch, M., Uritsky, N. & Albeck, A. Differentiating Serine and Cysteine Protease Mechanisms by New Covalent QSAR Descriptors. ChemBioChem 12, 1023–1026 (2011).

78. Karelson, M., Lobanov, V. S. & Katritzky, A. R. Quantum-Chemical Descriptors in QSAR/QSPR Studies. Chem. Rev. 96, 1027–1044 (1996).

79. Anderson, J. S. M., Liu, Y., Thomson, J. W. & Ayers, P. W. Predicting the quality of leaving groups in organic chemistry: Tests against experimental data. J. Mol. Struct. THEOCHEM 943, 168–177 (2010).

80. Traube, T., Vijayakumar, S., Hirsch, M., Uritsky, N., Shokhen, M. & Albeck, A. EMBM − A New Enzyme Mechanism-Based Method for Rational Design of Chemical Sites of Covalent Inhibitors. J. Chem. Inf. Model. 50, 2256–2265 (2010).

81. Perola, E., Cellai, L., Lamba, D., Filocamo, L. & Brufani, M. Long chain analogs of physostigmine as potential drugs for Alzheimer’s disease: new insights into the mechanism of action in the inhibition of acetylcholinesterase. Biochim. Biophys. Acta BBA - Protein Struct. Mol. Enzymol. 1343, 41–50 (1997).

82. Zhang, Y., Liu, H. & Yang, W. Free energy calculation on enzyme reactions with an efficient iterative procedure to determine minimum energy paths on a combined ab initio QM/MM potential energy surface. J. Chem. Phys. 112, 3483–3492 (2000).

83. Kidd, D., Liu, Y. & Cravatt, B. F. Profiling Serine Hydrolase Activities in Complex Proteomes. Biochemistry 40, 4005–4015 (2001).

84. Gok, M., Zeybek, N. D. & Bodur, E. Butyrylcholinesterase expression is regulated by fatty acids in HepG2 cells. Chem. Biol. Interact. 259, 276–281 (2016).

85. Kovalevich, J. & Langford, D. in Neuronal Cell Cult. Methods Protoc. (eds. Amini, S. & White, M. K.) 9–21 (Humana Press, 2013). doi:10.1007/978-1-62703-640-5_2

