## Supplementary Information for "Tunable nucleofugality in carbamoyl-bearing covalent cholinesterase inhibitors"

[a]*University of Ljubljana, Faculty of Pharmacy, Department of Pharmaceutical Chemistry. Aškerčeva 7, SI-1000 Ljubljana, Slovenia.*
***
[b]*South-Central Minzu University, College of Chemistry and Material Science, Wuhan 430079, China.*
[c]*Institut de Recherche Biomédicale des Armées, Département de Toxicologie et Risques Chimiques, Unité Neurotoxiques. 91223 Brétigny sur Orge, France.*
[d]*National Institute of Chemistry, Hajdrihova ulica 19, SI-1000 Ljubljana, Slovenia.*
[e]*University of Ljubljana, Faculty of Pharmacy, Department of Pharmaceutical Biology. Aškerčeva 7, SI-1000 Ljubljana, Slovenia.*
[f]*Institut de Recherche Biomédicale des Armées, Département des Plateformes et Recherches Technologiques, Unité Développements Analytiques et Bioanalyse. 91223 Brétigny sur Orge, France.*
[g]*University of Ljubljana, Faculty of Chemistry and Chemical Technology, Večna pot 113, SI-1000 Ljubljana, Slovenia.*
[h]*University of Ljubljana, Faculty of Medicine, Institute of Biochemisty and Molecular Genetics, Vrazov trg 2, SI-1000 Ljubljana, Slovenia.*
[i]*Molecular Modeling and Biopharmaceutical Center, College of Pharmacy, University of Kentucky, 789 South Limestone Street, Lexington, Kentucky 40536, United States.*
*Department of Pharmaceutical Sciences, College of Pharmacy, University of Kentucky, 789 South Limestone Street, Lexington, Kentucky 40536, United States.*
***

### Supporting Information

### Supplementary Results

**List S1 │** Covalent ChEI warhead chemotypes from the literature

(database – PubMed; search phrase – (cholinesterase OR butyrylcholinesterase OR acetylcholinesterase OR AChE OR BChE) AND (covalent OR time-dependent OR irreversible OR warhead OR electrophile OR reactive) AND (inhibitor OR drug OR compound); date – 1. 12. 2023; number of results – 9,954; only compounds with experimentally confirmed irreversible inhibition or with a probable warhead were retained; leaving groups are coloured in blue):

#### 1. Carbamoyl compounds

1.1. (hetero)aryl *O*-carbamates (e.g., pesticides carbofuran, dimetilan, isolan)[1–12,12–23]

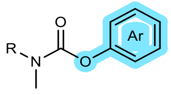

1.2. aryl *O*-thiocarbamates[24]

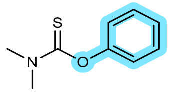

1.3. oxime *O*-carbamates (e.g., pesticides aldicarb, methomyl)[23,25,26]

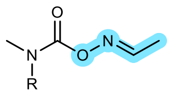

1.4. carbamoyl fluorides, chlorides[27–30]

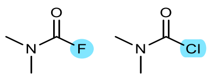

1.5. alkyl *O*-carbamates[31–34]

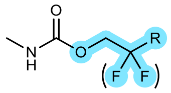

1.6. carbamoylated heterocycles[21,23]

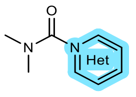

1.7. choline *O*-carbamates[35]

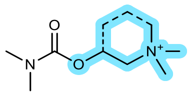

1.8. aliphatic *S*-carbamates[36] (*only after metabolic activation*!)

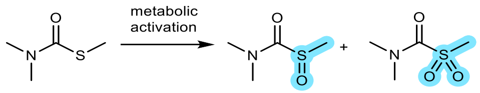

1.9. heteroaromatic *S*-carbamates[37]

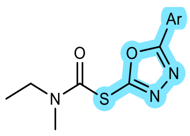

#### 2. Sulfonyl fluorides,[27,38–42] sulfonates[38]

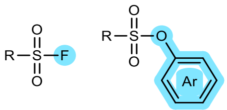

**3. Phosphorous compounds**[43–62]

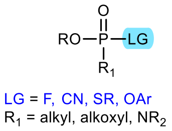

**4. Other electrophiles**

4.1. alkylating agents[63–72]

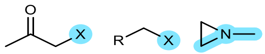

4.2. heterocycles

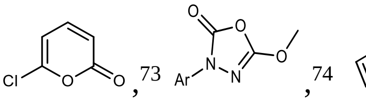

4.3. carbodiimides[76]

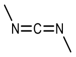

4.4. Michael acceptors[77,78]

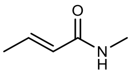

4.5. reversible covalent[79–82]

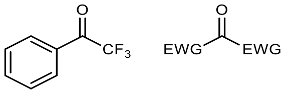

**5. Noncovalent, but with irreversible or slow kinetics**

5.1. arsenite[83]

5.2. dioxopyrimidines spanning $\omega$-tetraalkylammoniums[84]

5.3. DMSO[85]

5.4. territrem B[86]

5.5. (+)-arisugacin A[87]

**6. Other**

6.1. non-catalytic Cys-targeting for insect AChE[88–90]

6.2. divalent cations,[91,92] Hg[93,94]

6.3. reactive oxygen species (ROS)[95–99]

6.4. ($Ph_4B^-$)[100]

6.5. photoinactivation through ROS generation,[101,102] photoaffinity labelling[103–109]

*Cholinesterase inhibition*

**Table S1 |** Cholinesterase inhibition by fragment-sized carbamates

| Structure | Compound | hBChE | hAChE |
|---|---|---|---|
| | | **RA at 100 µM *or* IC$_{50}$ ± SEM$^a$** | |
| | | time-dependency of inhibition | |
| 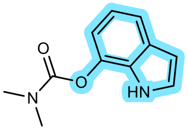 | **1a** | 1.66 ± 0.09 µM<br>***yes*** | 5.55 ± 0.81 µM<br>***yes*** |
| 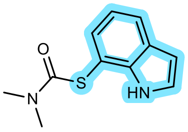 | **2a** | 80.7 ± 1.2% | 91.9 ± 6.7% |
| 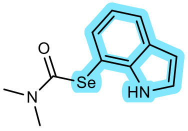 | **3a** | 50.9 ± 2.8% | 88.8 ± 3.1% |
| 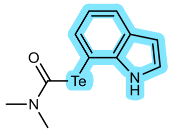 | **4a** | 12.6 ± 1.9 µM<br>***yes*** | 63.3 ± 5.1% |
| 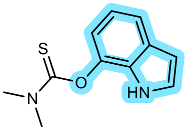 | **5a** | 57.9 ± 1.6% | 84.6 ± 8.5% |
| 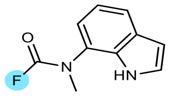 | **6a** | 1.27 ± 0.18 nM<br>***yes*** | 14.9 ± 2.9 nM<br>***yes*** |
| 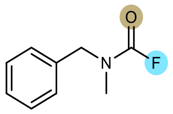 | **6b** | 28.1 ± 3.4 nM<br>***yes*** | 1.13 ± 0.13 µM<br>***yes*** |
| 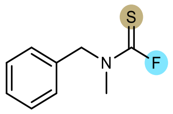 | **6c** | 229 ± 34 µM<br>***yes*** | 88.3 ± 12.7% |
| 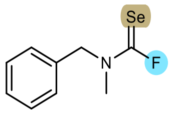 | **6d** | 3.90 ± 0.83 µM<br>***yes*** | 76.2 ± 1.6% |
| 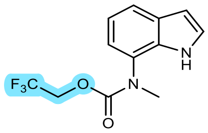 | **7a** | 85.7 ± 8.2% | 76.0 ± 4.4% |
| 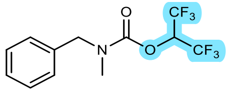 | **8b** | 98.0 ± 4.0% | 97.3 ± 5.1% |
| 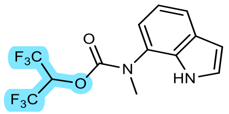 | **9a** | 93.7 ± 3.5% | 54.0 ± 3.1% |

| | | | |
|---|---|---|---|
| 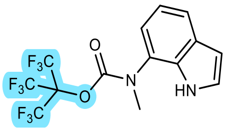 | **10a** | 75.1 ± 8.6% | 85.5 ± 1.7% |
| 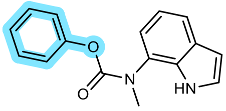 | **11a** | 37.1 ± 4.0% | 77.3 ± 4.0% |
| 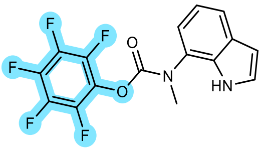 | **12a** | 75.3 ± 2.1% | 21.8 ± 6.7 µM<br>***unclear*** |
| 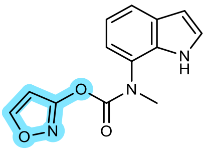 | **12a** | 310 ± 67 nM<br>***yes*** | 97.8 ± 6.0 µM<br>***unclear*** |
| 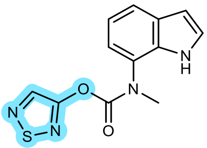 | **13a** | 3.16 ± 0.59 µM<br>***yes*** | 107 ± 12 µM<br>***unclear*** |
| 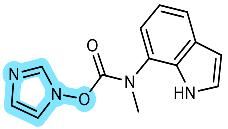 | **14a** | 501 ± 57 nM<br>***yes*** | 70.0 ± 4.6% |
| 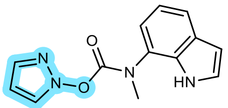 | **15a** | 13.4 ± 0.7 nM<br>***yes*** | 5.46 ± 1.36 µM<br>***yes*** |
| 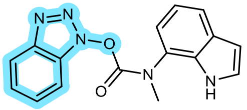 | **16a** | 525 ± 89 nM<br>***yes*** | 624 ± 77 µM<br>***yes*** |
| 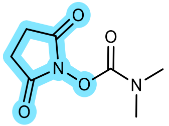 | **17b** | 71.1 ± 1.6% | 88.9 ± 3.2% |
| 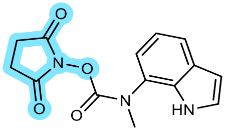 | **17a** | 45.7 ± 1.9% | 79.7 ± 3.6% |
| 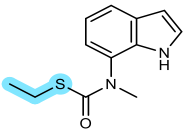 | **18a** | 95.8 ± 1.9% | 97.5 ± 2.0% |
| 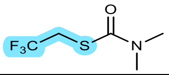 | **19b** | 94.5 ± 1.3% | 93.0 ± 3.5% |
| 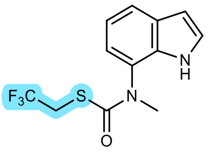 | **19a** | 62.7 ± 4.0% | 83.8 ± 6.1% |

| | | | |
|---|---|---|---|
| 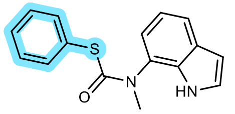 | **20a** | 37.0 ± 4.9% | 42.7 ± 13.6 µM<br>***unclear*** |
| 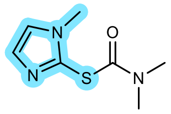 | **21b** | 94.8 ± 3.9% | 96.6 ± 2.5% |
| 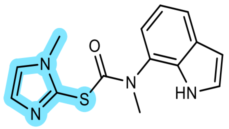 | **21a** | 53.3 ± 1.6% | 92.1 ± 2.2% |
| 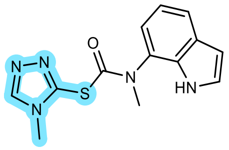 | **22a** | 2.30 ± 0.49 µM<br>***yes*** | 15.5 ± 4.9 µM<br>***yes*** |
| 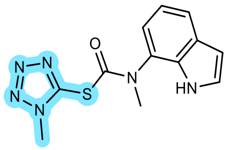 | **23a** | 3.05 ± 0.45 µM<br>***yes*** | 70.0 ± 4.6% |
| 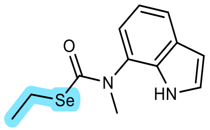 | **24a** | 78.5 ± 2.6% | 78.7 ± 3.1% |
| 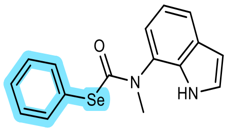 | **25a** | 8.70 ± 2.32 µM<br>***no*** | 64.6 ± 0.9% |
| 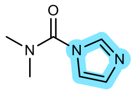 | **26b** | 97.2 ± 1.2% | 93.2 ± 4.2% |
| 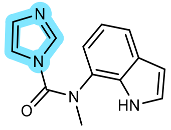 | **26a** | 25.2 ± 6.9 µM<br>***no*** | 57.3 ± 8.7% |
| 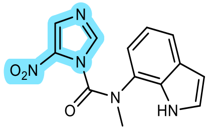 | **27a** | 58.9 ± 3.2% | 51.7 ± 1.4% |
| 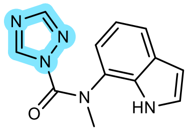 | **28a** | 4.92 ± 0.81 µM<br>***yes*** | 129 ± 23 µM<br>***unclear*** |
| 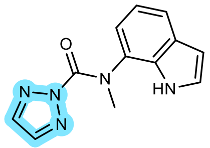 | **29a** | 44.3 ± 2.2 nM<br>***yes*** | 17.6 ± 1.1 µM<br>***yes*** |
| 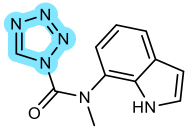 | **30a** | 743 ± 167 nM<br>***yes*** | 12.1 ± 3.5 µM<br>***yes*** |

| | | | |
|---|---|---|---|
| 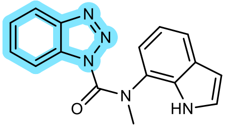 | **31a** | 4.02 ± 1.39 µM<br>***yes*** | 20.6 ± 4.5 µM<br>***yes*** |
| 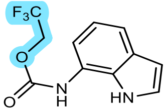 | **32a** | 80.3 ± 2.7% | 88.9 ± 2.3% |
| 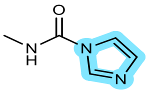 | **33b** | 87.5 ± 4.7% | 83.8 ± 2.9% |

[a]Preincubation time: 30 min. SEM – standard error of the mean, experiments were performed in triplicate. RA – residual activity [%] (mean ± SD of one independent experiment performed in triplicate). Compounds with RA > 50% were considered inactive and were not evaluated in a time-dependency assay.

*Time-dependency experiments – representative examples*

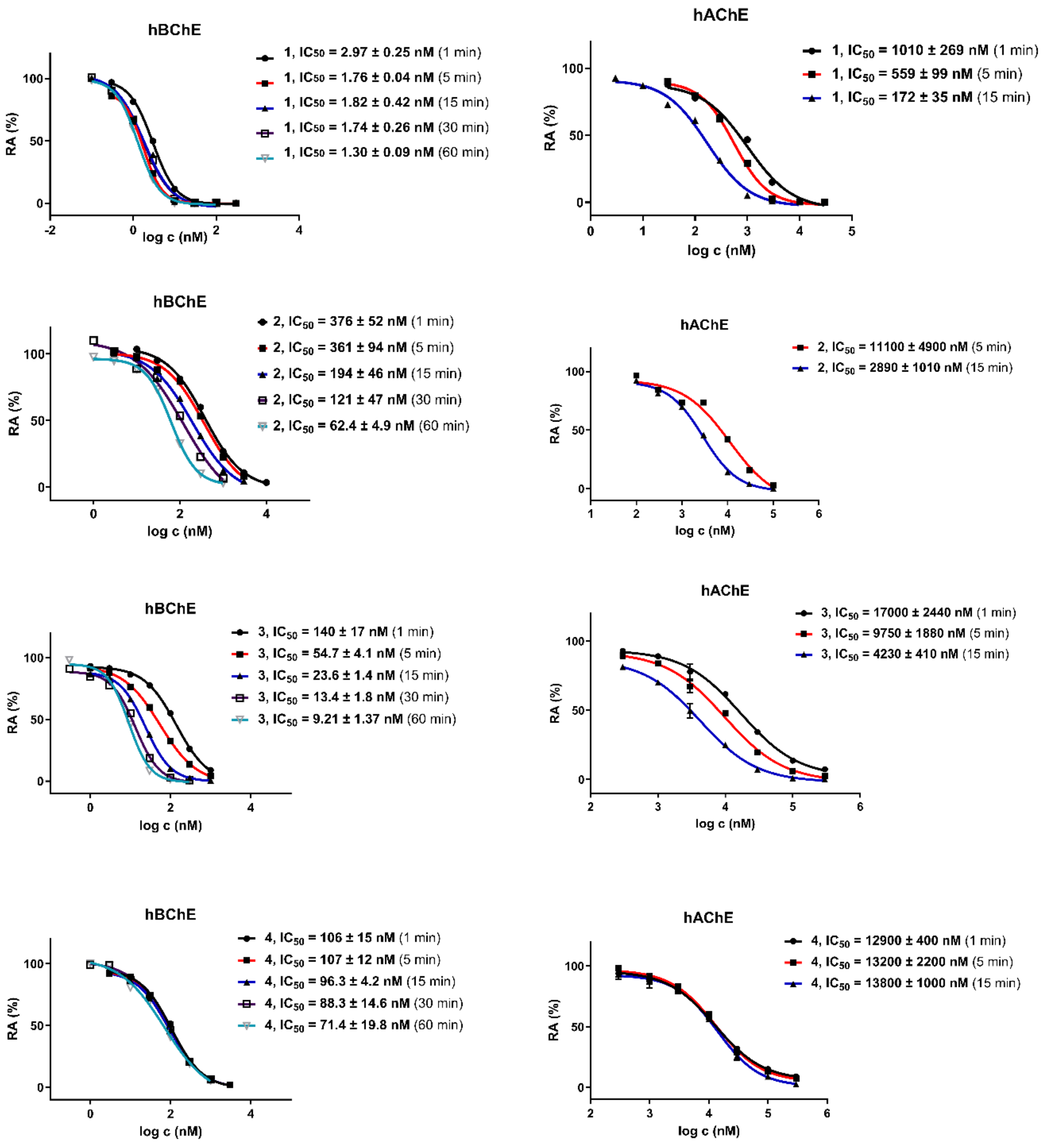

**Fig. S1**│Time-dependency experiments on hBChE and hAChE for compounds **1**–**4**. The residual activities are plotted against log-concentration. The leftward shift, i.e., the decreasing of $IC_{50}$ values with longer preincubation time, suggests covalent mechanism of inhibition. For **1**, the shift for hBChE is less evident, which indicates fast carbamoylation reaction (at >15 min, all inhibitor molecules have reacted, therefore $IC_{50}$ value does not change anymore) – note: independent kinetic measurement confirmed covalent nature of inhibition.[110]

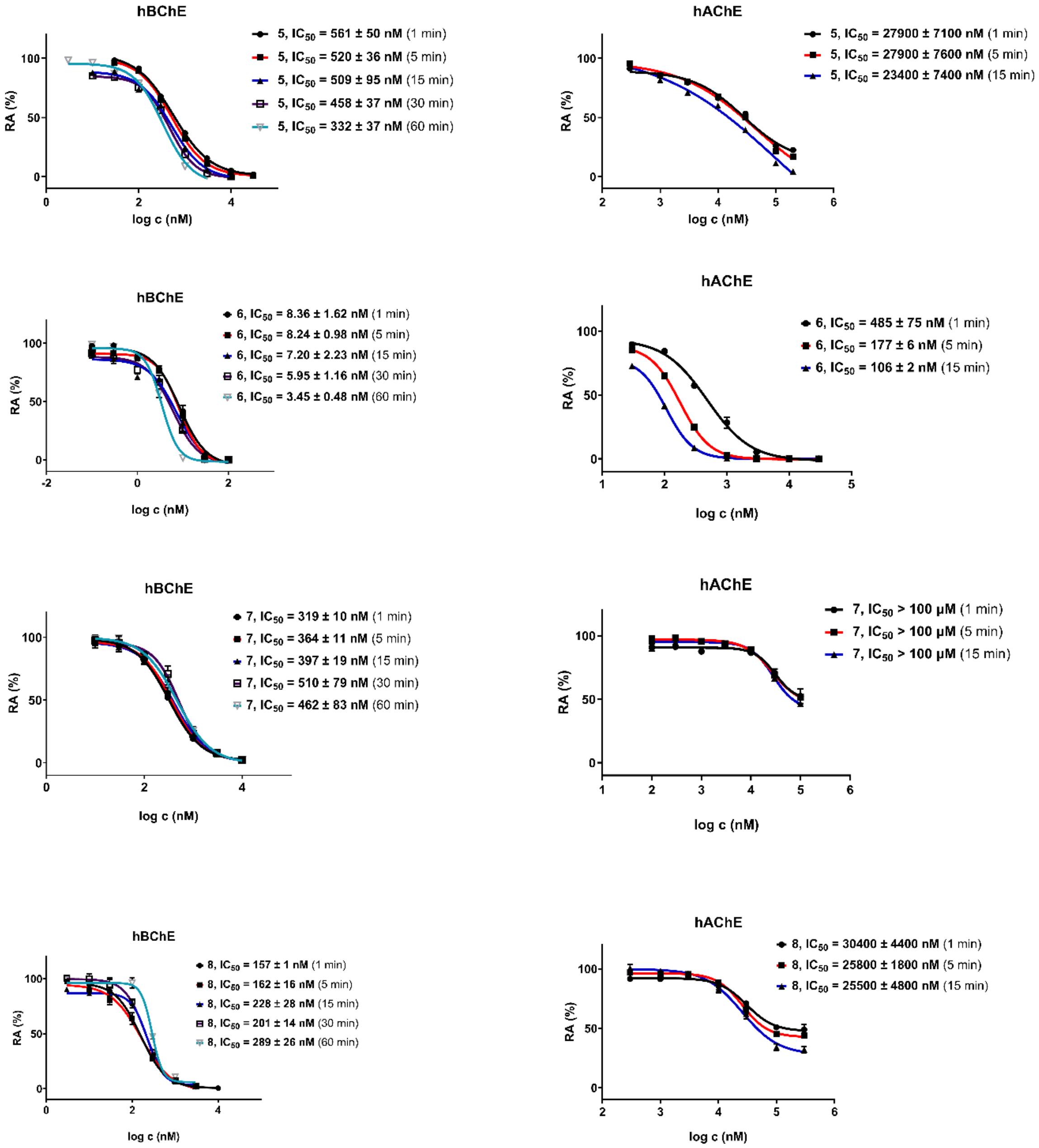

**Fig. S2** | Time-dependency experiments on hBChE and hAChE for compounds **5–8** (*continued*).

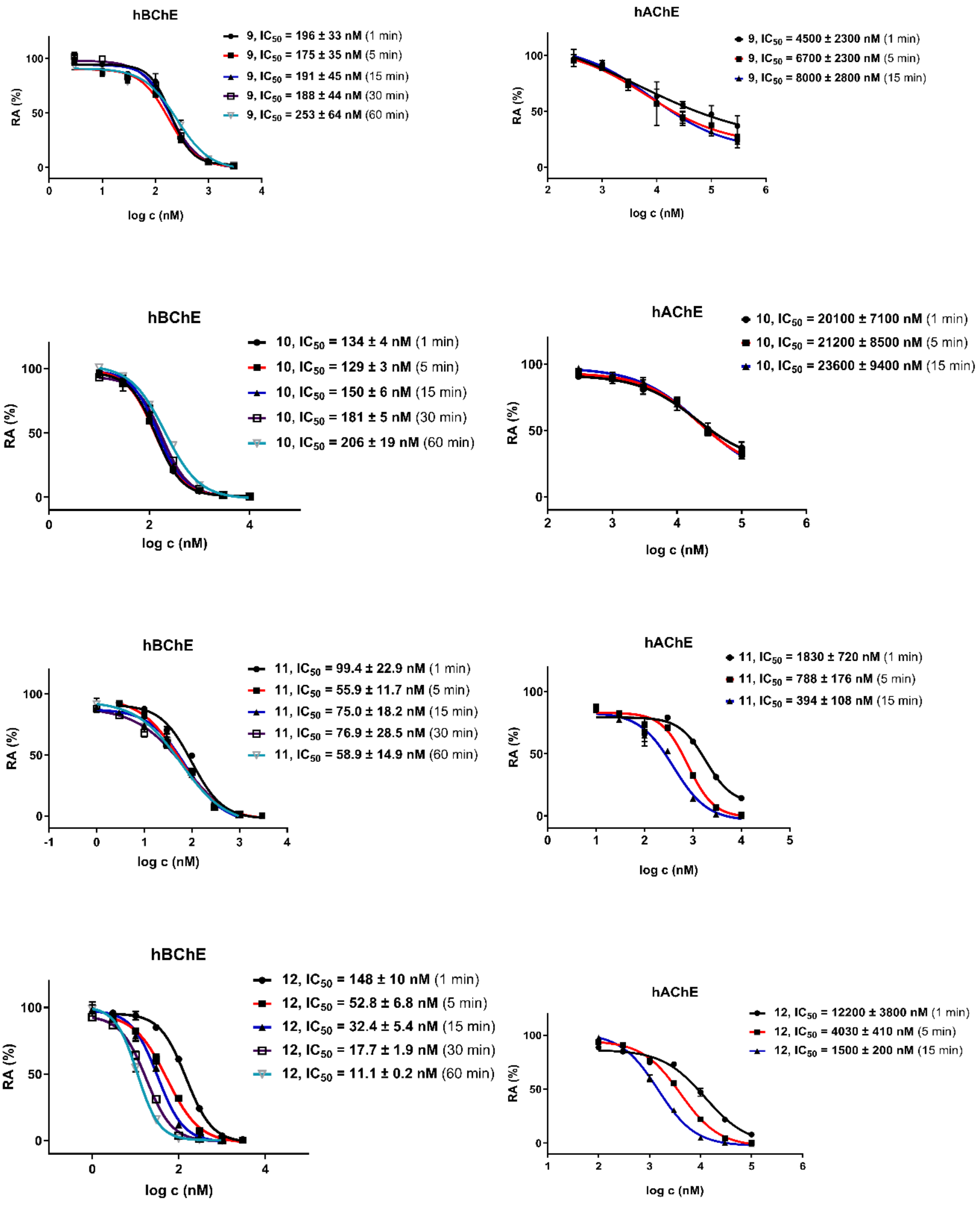

**Fig. S3**│Time-dependency experiments on hBChE and hAChE for compounds **9–12** (*continued*).

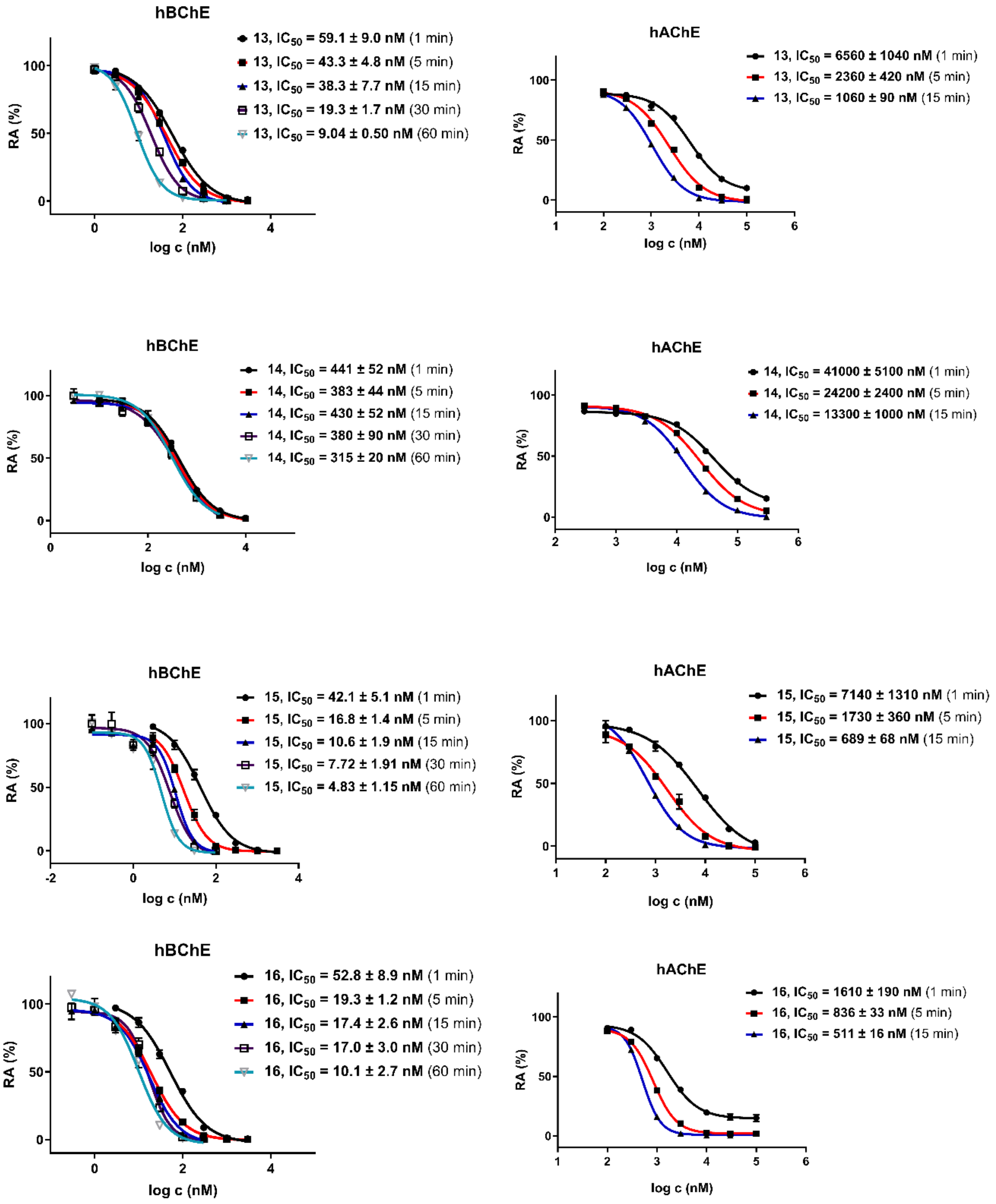

**Fig. S4**│Time-dependency experiments on hBChE and hAChE for compounds **13–16** (*continued*).

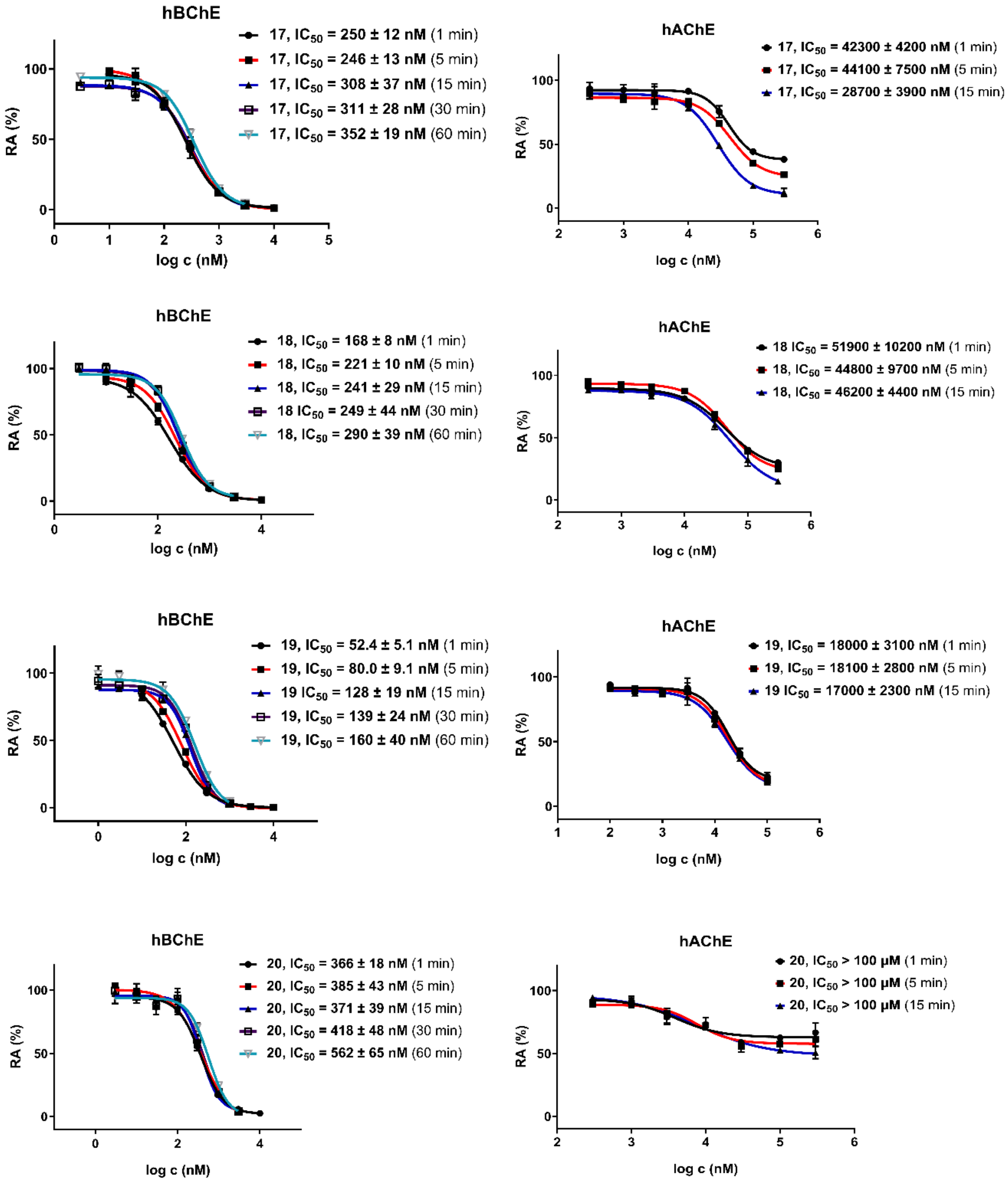

**Fig. S5**│Time-dependency experiments on hBChE and hAChE for compounds **17–20** (*continued*).

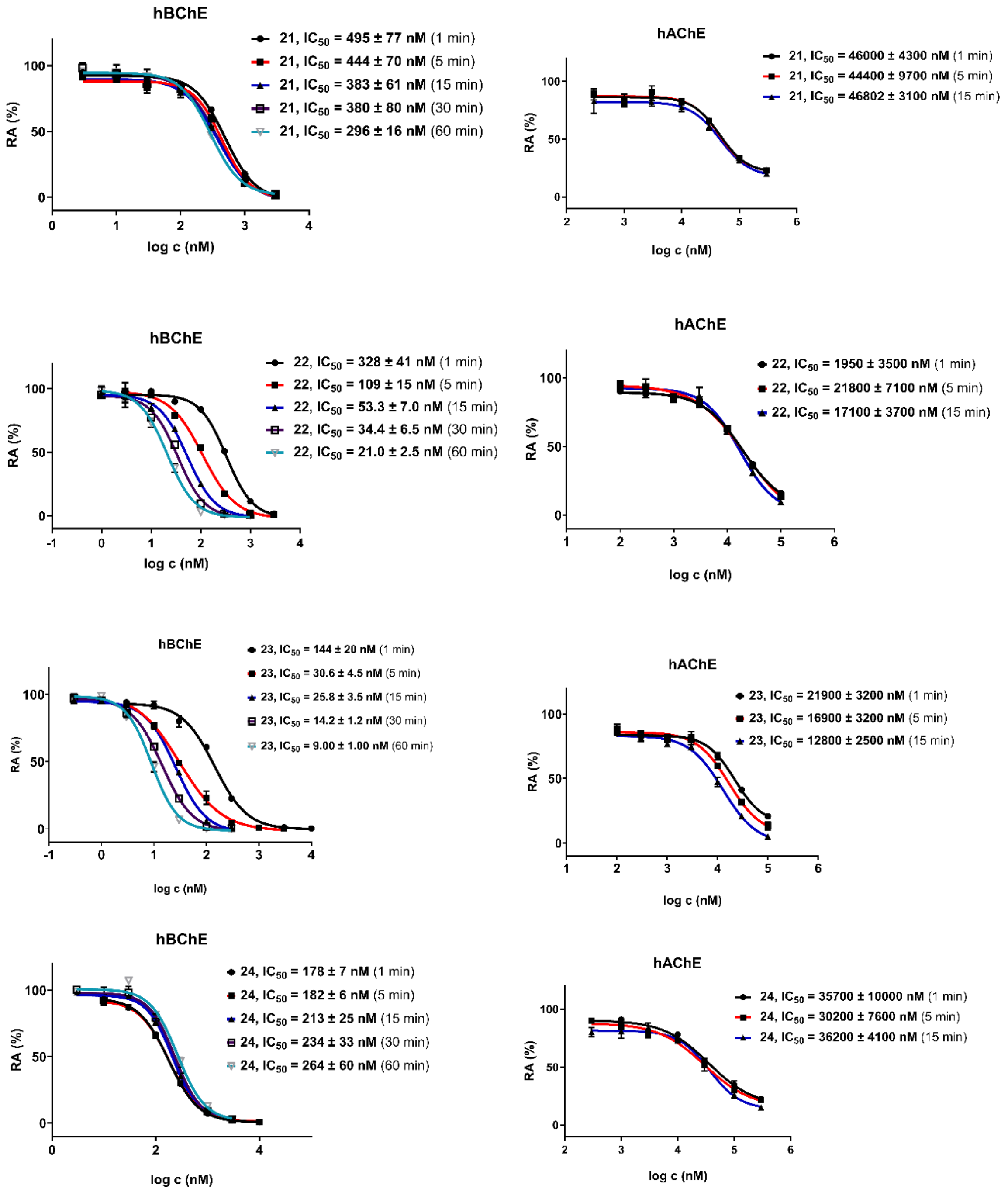

**Fig. S6**│Time-dependency experiments on hBChE and hAChE for compounds **21–24** (*continued*).

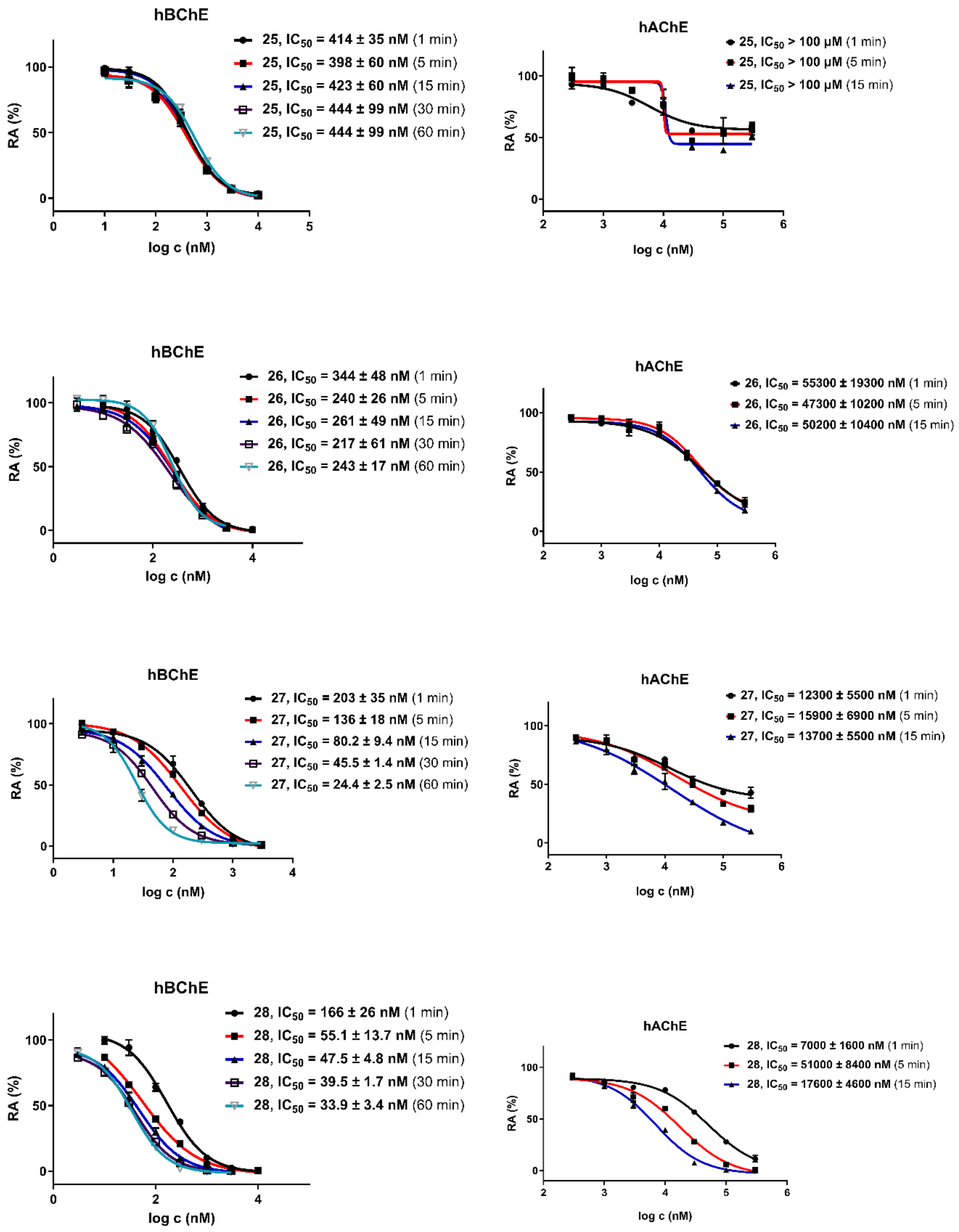

**Fig. S7**│Time-dependency experiments on hBChE and hAChE for compounds **25–28** (*continued*).

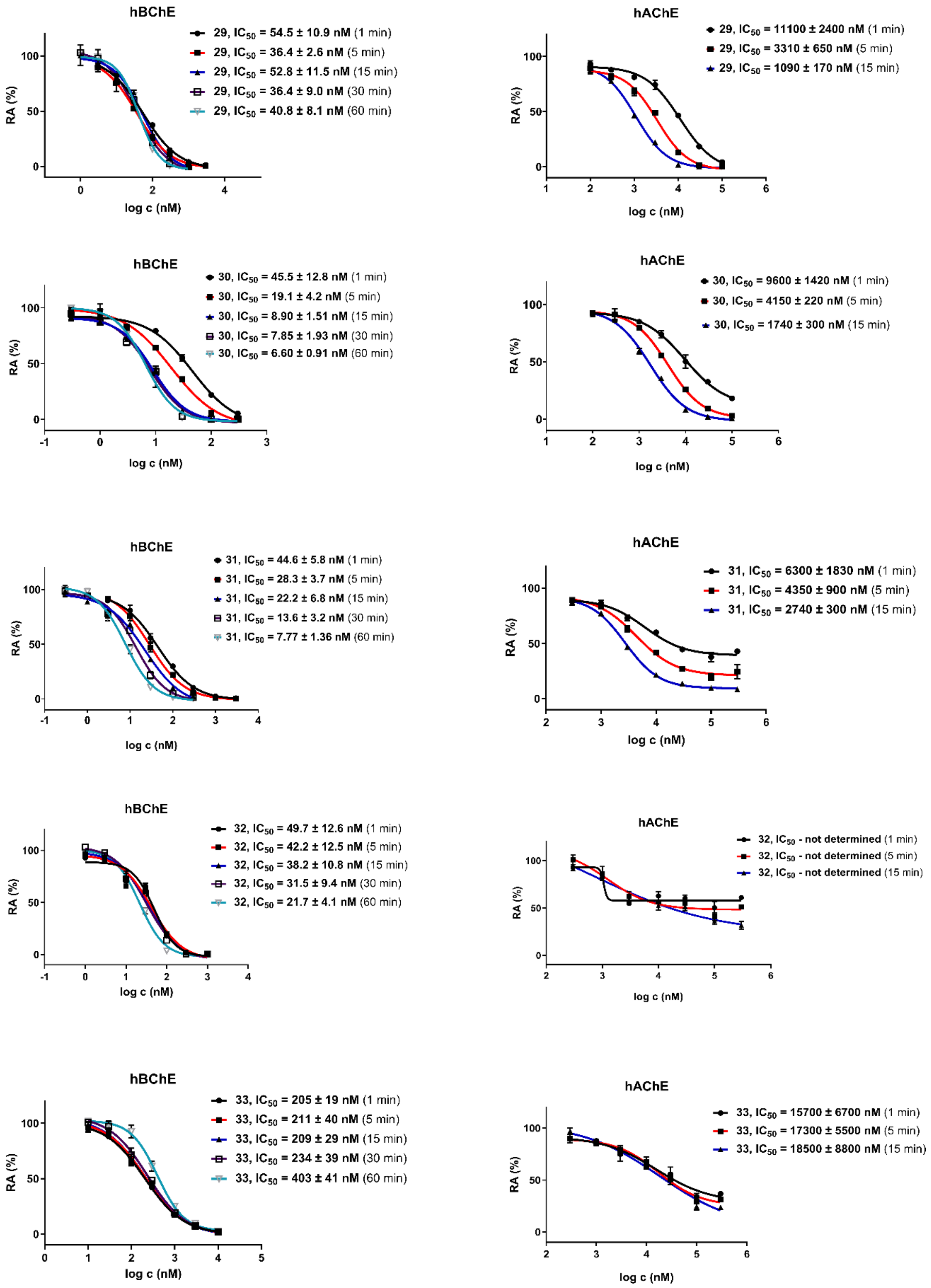

**Fig. S8**│Time-dependency experiments on hBChE and hAChE for compounds **29–33** (*continued*).

*Enzyme kinetics*

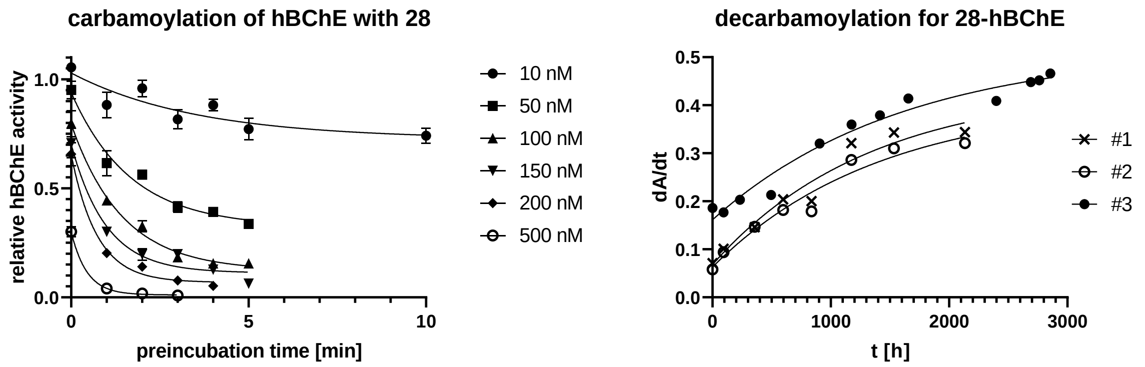

**Fig. S9 |** Initial velocity measurements and decarbamoylation for **28**. Left – hBChE residual activity as a function of preincubation time at different concentrations of **28**. The enzyme and inhibitor were preincubated for a specified amount of time, then the substrate was added, initial velocity measured, and compared with the uninhibited enzyme control. Values are expressed as means ± SEM, measured in triplicates. Right – recovery of enzyme activity over time for **33**-carbamoylated hBChE (in three independent determinations). After 3 months, the spoilage of the dilute enzyme solution occurred – however, a complete reactivation was still not observed.

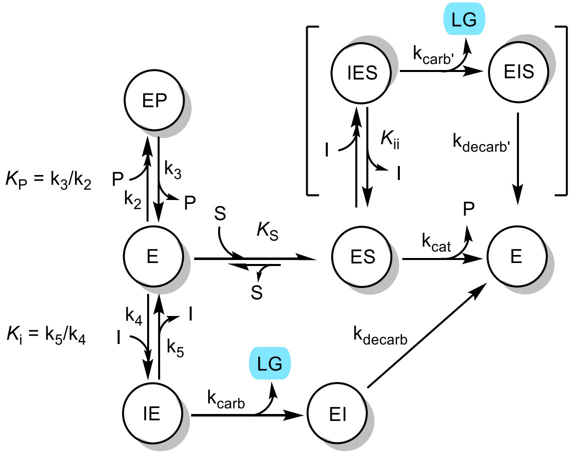

**Scheme S1 |** Reaction scheme for the interaction between hBChE – E and carbamate inhibitor I. Legend: S – substrate (butyrylthiocholine iodide); P – product (mixed disulfide); E, ES, EP, IE, IES, EI, EIS – different enzyme complexes; $K_s$, $K_p$, $K_i$, $K_{ii}$ – dissociation constants; $k_{cat}$ – catalytic constant; $k_{carb}$, $k_{carb'}$, $k_{decarb}$, $k_{decarb'}$ – first-order rate constants for carbamoylation and decarbamoylation. $K_s$, $K_p$ were determined previously.

**Table S2** | Raw data for Fig. S9

| | Initial velocity of uninhibited enzyme | | | | AVERAGE = 0.41075 | | | | | | | | | AVERAGE = 0.4245 | | | | | | | | |
|---|---|---|---|---|---|---|---|---|---|---|---|---|---|---|---|---|---|---|---|---|---|---|
| | 0.392 | 0.423 | 0.391 | 0.437 | | | | | | | | | | 0.417 | 0.459 | 0.406 | 0.416 | | | | | |
| **time [min]** | **STOCK [28]** | **5 µM** | | | **10 µM** | | | **15 µM** | | | **20 µM** | | | **1 µM** | | | **50 µM** | | | **100 µM** | | |
| 0 | **initial velocity** | 0.424 | 0.373 | 0.375 | 0.374 | 0.318 | 0.29 | 0.321 | 0.316 | 0.235 | 0.312 | 0.243 | 0.254 | 0.498 | 0.405 | 0.44 | 0.145 | 0.111 | 0.127 | 0.04 | 0.034 | 0.033 |
| 1 | | 0.208 | 0.29 | 0.26 | 0.171 | 0.188 | 0.188 | 0.114 | 0.124 | 0.133 | 0.074 | 0.091 | 0.085 | 0.347 | 0.353 | 0.424 | 0.013 | 0.019 | 0.02 | 0.005 | 0.005 | 0.006 |
| 2 | | 0.235 | 0.222 | 0.236 | 0.118 | 0.129 | 0.154 | 0.068 | 0.072 | 0.103 | 0.057 | 0.058 | 0.058 | 0.411 | 0.432 | 0.377 | 0.01 | 0.006 | 0.007 | | | |
| 3 | | 0.188 | 0.169 | 0.154 | 0.067 | 0.087 | 0.072 | 0.073 | 0.081 | 0.09 | 0.036 | 0.026 | 0.033 | 0.338 | 0.382 | 0.319 | 0.004 | 0.004 | 0.003 | | | |
| 4 | | 0.155 | 0.169 | 0.159 | 0.081 | 0.057 | 0.054 | 0.048 | 0.043 | 0.067 | 0.024 | 0.016 | 0.025 | 0.388 | 0.351 | 0.383 | | | | | | |
| 5 | | 0.129 | 0.135 | 0.151 | 0.065 | 0.065 | 0.062 | 0.027 | 0.018 | 0.034 | | | | 0.365 | 0.325 | 0.292 | | | | | | |
| 10 | | | | | | | | | | | | | | 0.343 | 0.297 | 0.304 | | | | | | |
| 0 | **residual activity** | 1.0323 | 0.9081 | 0.9130 | 0.9105 | 0.7742 | 0.7060 | 0.7815 | 0.7693 | 0.5721 | 0.7596 | 0.5916 | 0.6184 | 1.1731 | 0.9541 | 1.0365 | 0.3416 | 0.2615 | 0.2992 | 0.0942 | 0.0801 | 0.0777 |
| 1 | | 0.5064 | 0.7060 | 0.6330 | 0.4163 | 0.4577 | 0.4577 | 0.2775 | 0.3019 | 0.3238 | 0.1802 | 0.2215 | 0.2069 | 0.8174 | 0.8316 | 0.9988 | 0.0306 | 0.0448 | 0.0471 | 0.0118 | 0.0118 | 0.0141 |
| 2 | | 0.5721 | 0.5405 | 0.5746 | 0.2873 | 0.3141 | 0.3749 | 0.1656 | 0.1753 | 0.2508 | 0.1388 | 0.1412 | 0.1412 | 0.9682 | 1.0177 | 0.8881 | 0.0236 | 0.0141 | 0.0165 | | | |
| 3 | | 0.4577 | 0.4114 | 0.3749 | 0.1631 | 0.2118 | 0.1753 | 0.1777 | 0.1972 | 0.2191 | 0.0876 | 0.0633 | 0.0803 | 0.7962 | 0.8999 | 0.7515 | 0.0094 | 0.0094 | 0.0071 | | | |
| 4 | | 0.3774 | 0.4114 | 0.3871 | 0.1972 | 0.1388 | 0.1315 | 0.1169 | 0.1047 | 0.1631 | 0.0584 | 0.0390 | 0.0609 | 0.9140 | 0.8269 | 0.9022 | | | | | | |
| 5 | | 0.3141 | 0.3287 | 0.3676 | 0.1582 | 0.1582 | 0.1509 | 0.0657 | 0.0438 | 0.0828 | | | | 0.8598 | 0.7656 | 0.6879 | | | | | | |
| 10 | | | | | | | | | | | | | | 0.8080 | 0.6996 | 0.7161 | | | | | | |
| **inhibitor concentration inside cuvette [M]** | **GraphPad exponential fit $k_{obs}$ [$min^{-1}$]** | **LINEST for 1/[I] = f(1/$k_{obs}$)** | | | ***$K_i$*** | **46.129** | **nM** | | | | | | | | | | | | | | | |
| 1.00E-08 | 0.281 | 2.990E-08 | 6.483E-01 | | $K_i/k_{carb}$ | 0.000 | | | | | | | | | | | | | | | | |
| 5.00E-08 | 0.597 | 3.217E-09 | 1.344E-01 | | $1/k_{carb}$ | 0.648 | | | | | | | | | | | | | | | | |
| 1.50E-07 | 1.015 | 9.558E-01 | 2.777E-01 | | $R^2$ | 0.956 | | | | | | | | | | | | | | | | |
| 2.00E-07 | 1.364 | 8.642E+01 | 4.000E+00 | | **$k_{carb}$=** | **1.543** | **$M^{-1}$ $min^{-1}$** | | | | | | | | | | | | | | | |
| 5.00E-07 | 2.260 | 6.664E+00 | 3.084E-01 | | ΔG‡= | **19.622** | **kcal/mol** | | | | | | | | | | | | | | | |
| 1.00E-06 | 1.900 | | | | | | | | | | | | | | | | | | | | | |

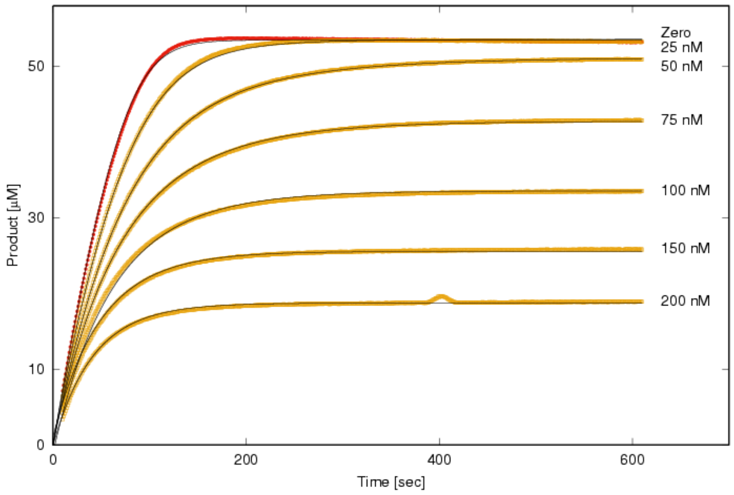

concentrations:
25 nM, 50 nM, 75 nM, 100 nM, 150 nM, 200 nM

$K_i$ = 230 nM
$K_{ii}$ = 326 nM
$k_{carb}$ = 0.088 $s^{-1}$
$k_{carb'}$ = 0.052 $s^{-1}$

**12**

**Fig. S10 |** Left, progress curves for the yellow color formation (thionitrobenzoate as detection species) in the hydrolysis of BTChI by hBChE in the absence and presence of **12**. Right, characteristic kinetic parameters (standard errors were below 1% and are thus not reported). Red curve was measured without inhibitor and orange curves are in presence of different inhibitor concentrations (listed on the right).

Among tested reaction schemes, the one where **12** reacts with the free enzyme and the enzyme-substrate complex, both leading to the carbamoylated enzyme, satisfied the data. The inhibition pattern in Fig. S10 is characteristic of irreversible inhibition in the presence of inhibitor surplus concentrations: while the curve in the absence of inhibitor is mechanistically a typical integrated Michaelis-Menten equation, those in the presence of **12** are exponential. This suggests that an inhibition component of a competitive type must be present. Additionally, kinetics reveals that the ternary complex enzyme–substrate–inhibitor is also present, which increases observed $K_M$ for BTChI. In the Scheme S1, product inhibition is also included, since the disulfide adduct between thiocholine-thionitrobenzoic acid significantly inhibits hBChE at 50 µM concentration.[111] It should be emphasized that no decarbamoylation could be detected at a 24 h-scale, neither upon dilution of the carbamoylated enzyme, nor by the separation of the free **12** by a size exclusion column. Indeed, fully inhibited hBChE, rather than any kind of steady state, is a direct consequence of **12**'s action, since except at the lowest concentration, the plateaus were lower than in the zero curve, suggesting that complete inactivation of the enzyme occurred before it could hydrolyse all substrate.

*hBChE X-ray structural study*

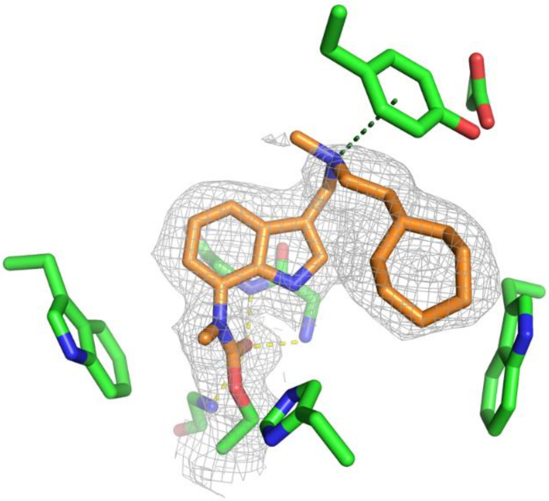

**Fig. S11 |** Crystal structure of **28**-carbamoylated hBChE (PDB 8QTW, resolution 2.24 Å). The key amino acid residues are shown as green sticks, **28-**derived moiety is shown as orange sticks. The grey mesh represents the 1-σ countered phenix-calculated feature-enhanced map around the carbamoylated Ser198 residue.

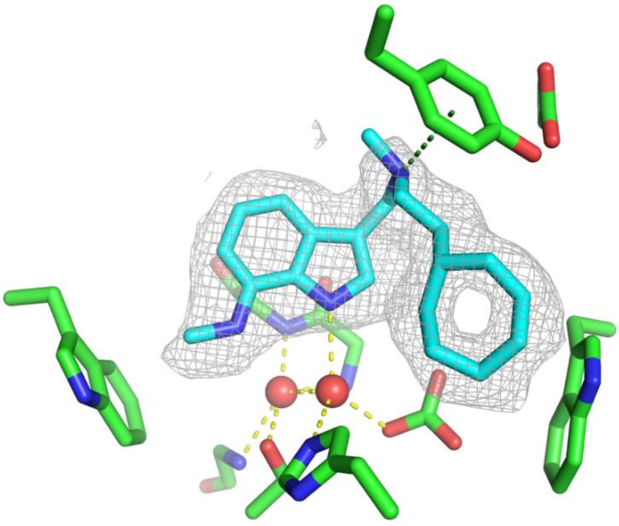

**Fig. S12 |** Crystal structure of hydrolysed and decarboxylated **28** in complex with hBChE (PDB 8QTX, resolution 2.12 Å). The key amino acid residues are shown as green sticks, **28**-derived moiety is shown as cyan sticks, and the water molecules as red spheres. The hydrogen bonds are shown as yellow dashed lines, and the cation-π interaction with Tyr332 as a green dashed line. The grey mesh represents the 1-σ countered phenix-calculated feature-enhanced map around the hydrolysed and decarboxylated ligand **28**. The binding pose features a characteristic cation-π interaction of the central tertiary amine with Tyr332 at the aromatic site, while the *N*-methyl-7-aminoindole moiety resides in the acyl-binding pocket, forming only van der Waals interactions with the nearby residues. The oxyanion hole is occupied by a water molecule. An extra electron density of triangular shape is observed close to Trp82 and was modelled as a carbonate molecule with occupancy around 0.7. Contrarily to cations, anions are not usually observed near Trp82 and such density could have also been modelled as a carbamate from the reversible reaction of the carbonate with the ammonium ion present at crystallisation conditions. However, the current resolution X-ray data cannot discriminate between oxygen and nitrogen atoms.

**Table S3 |** Data collection and refinement statistics of **28**-carbamoylated hBChE (PDB 8QTW) and hBChE in complex with hydrolysed and decarboxylated **28** (PDB 8QTX). Table calculated using Phenix. R-work=Σ |*F*o – |*F*c|| / Σ |*F*o|, where *F*o and *F*c are observed and calculated structure factors, respectively. R-free set uses about 5% randomly chosen reflections. Statistics for the highest-resolution shell are shown in parentheses.

| | **8QTW** | **8QTX** |
|---|---|---|
| ***Data collection*** | | |
| **X-ray source** | ESRF ID23-1 | ESRF ID23-1 |
| **Wavelength [Å]** | 0.8856 | 0.9763 |
| **Resolution range [Å]** | 49.24 - 2.241 (2.321 - 2.241) | 49.22 - 2.12 (2.196 - 2.12) |
| **Space group** | I 4 2 2 | I 4 2 2 |
| **Unit cell [Å; °]** | 154.231 154.231 127.979<br>90 90 90 | 154.639 154.639 127.643<br>90 90 90 |
| **Total reflections** | 274794 (27986) | 353202 (32532) |
| **Unique reflections** | 36701 (3644) | 43198 (4130) |
| **Multiplicity** | 7.5 (7.7) | 8.2 (7.7) |
| **Completeness (%)** | 98.50 (99.21) | 98.05 (95.60) |
| **Mean I/sigma(I)** | 19.79 (2.47) | 13.31 (1.22) |
| **Wilson B-factor [Å$^2$]** | 51.55 | 48.21 |
| **R-merge** | 0.05438 (0.6859) | 0.08076 (1.48) |
| **R-meas** | 0.05833 (0.7359) | 0.08616 (1.589) |
| **R-pim** | 0.02057 (0.2606) | 0.02907 (0.5604) |
| **CC1/2** | 0.999 (0.887) | 0.999 (0.42) |
| **CC*** | 1 (0.97) | 1 (0.769) |
| ***Refinement statistics*** | | |
| **Reflections used in refinement** | 36699 (3636) | 43101 (4130) |
| **Reflections used for R-free** | 1823 (193) | 2092 (202) |
| **R-work** | 0.1778 (0.2892) | 0.1783 (0.3124) |
| **R-free** | 0.2183 (0.3251) | 0.2193 (0.3669) |
| **CC(work)** | 0.961 (0.908) | 0.965 (0.718) |
| **CC(free)** | 0.942 (0.851) | 0.811 (0.687) |
| **Number of non-hydrogen atoms** | 4699 | 4663 |
| **macromolecules** | 4237 | 4248 |
| **ligands** | 300 | 243 |
| **solvent** | 162 | 172 |
| **Protein residues** | 526 | 527 |
| **RMS(bonds)** | 0.009 | 0.008 |
| **RMS(angles)** | 0.96 | 0.88 |
| **Ramachandran favored (%)** | 96.18 | 95.81 |
| **Ramachandran allowed (%)** | 3.63 | 4.19 |
| **Ramachandran outliers (%)** | 0.19 | 0.00 |
| **Rotamer outliers (%)** | 1.31 | 1.53 |
| **Clashscore** | 6.16 | 6.23 |
| **Average B-factor** | 68.11 | 57.66 |
| **macromolecules** | 66.09 | 56.21 |
| **ligands** | 97.70 | 83.94 |
| **solvent** | 66.14 | 56.52 |
| **Number of TLS groups** | 1 | 1 |

*MS proteomic analysis*

**Table S4**│Human proteins from HepG2 and SH-SY5Y cell lines that were activity-dependently labelled with probe **B3**. Proteins with known enzymatic activity are annotated with function and EC, if available (source: UniProtKB[112]). LFQ intensity ratios are ratios of the MS signal LFQ intensity corresponding to the specific protein fragment in the **B3**-treated sample *vs*. in the warhead-less probe **B2**–treated sample. *Raw MS proteomics data are included in Supplementary Materials.*

| UniProtID | Gene name | Protein name | LFQ intensity ratio **B3/B2** |
|---|---|---|---|
| P10619 | CTSA | Lysosomal protective protein<br>3.4.16.5 – *carboxypeptidase (catalytic Ser)* | 5086300 |
| Q13510 | ASAH1 | Acid ceramidase<br>3.5.1.23 – *deacylase (catalytic Cys)* | 4268600 |
| Q15274 | QPRT | Nicotinate-nucleotide pyrophosphorylase [carboxylating]<br>2.4.2.19 – *transferase* | 3040600 |
| P04181 | OAT | Ornithine aminotransferase, mitochondrial<br>2.6.1.13 – *transferase* | 2779600 |
| Q3LXA3 | DAK | Triokinase/FMN cyclase<br>2.7.1.29 – *kinase* | 2557500 |
| P14174 | MIF | Macrophage migration inhibitory factor<br>5.3.2.1 – *tautomerase* | 30910000 |
| Q00325 | SLC25A3 | Solute carrier family 25 member 3<br>*ion transporter* | 16283000 |
| P81605 | DCD | Dermcidin<br>*protease activity* | 15632000 |
| P50991 | CCT4 | T-complex protein 1 subunit delta<br>*chaperone* | 9620200 |
| P60468 | SEC61B | Protein transport protein Sec61 subunit beta | 7678500 |
| P14923 | JUP | Junction plakoglobin | 5270000 |
| Q02413 | DSG1 | Desmoglein-1 | 5187300 |
| P06702 | S100A9 | Protein S100-A9 | 4970600 |
| Q9NQC3 | RTN4 | Reticulon-4 | 4253100 |
| Q9UJZ1 | STOML2 | Stomatin-like protein 2, mitochondrial | 3892300 |
| P35613 | BSG | Basigin | 3435900 |
| P48681 | NES | Nestin | 3166600 |
| P56134 | ATP5J2 | ATP synthase subunit f, mitochondrial | 2353400 |
| Q02878 | RPL6 | Large ribosomal subunit protein eL6 | 2251100 |
| P0DPI2 | GATD3 | Glutamine amidotransferase-like class 1 domain-containing protein 3, mitochondrial | 1944500 |

*Computational studies*

**Glu197** COOH **Glu441** $COO^-$

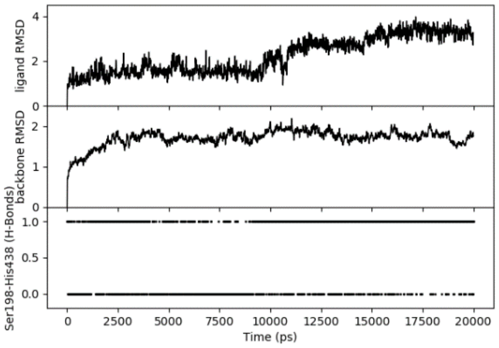

**Glu197** $COO^-$ **Glu441** COOH

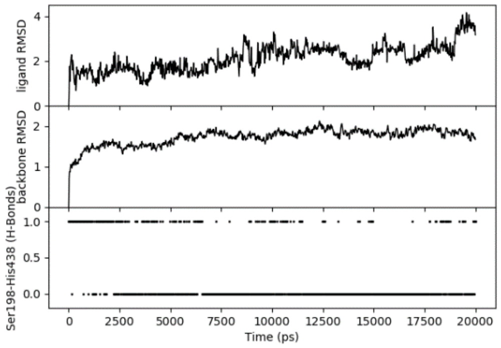

**Glu197** COOH **Glu441** COOH

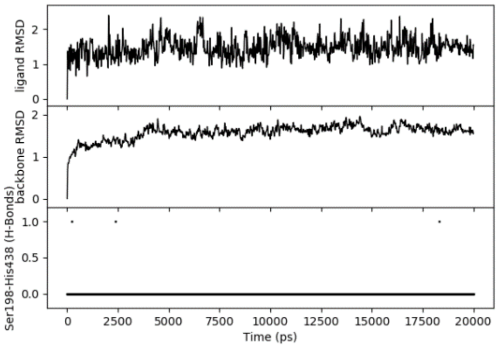

**Glu197** $COO^-$ **Glu441** $COO^-$

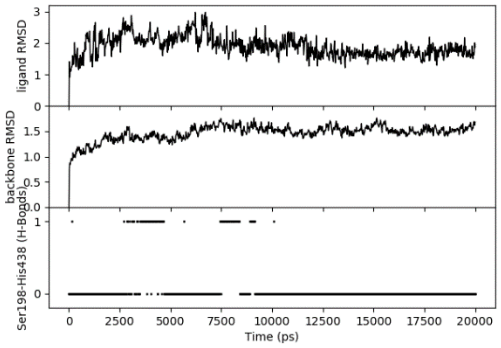

**Fig. S13 |** Molecular dynamics of different protonation states of Glu197 and Glu441 in the 8QTX crystal structure. The top trace shows ligand RMSD (fitted to backbone and compared with frame 0), the middle shows protein backbone RMSD, and the bottom trace depicts the number of hydrogen bonds between Ser198 and His438, all indicating the stability of the protein.

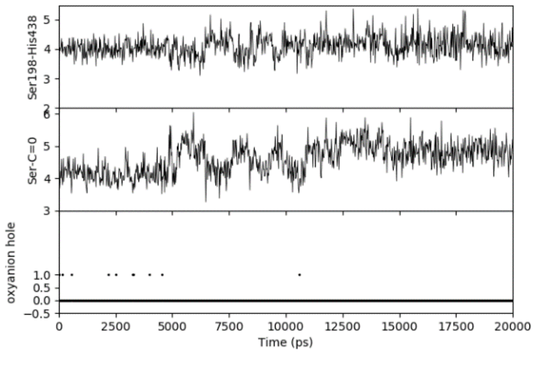

**6**

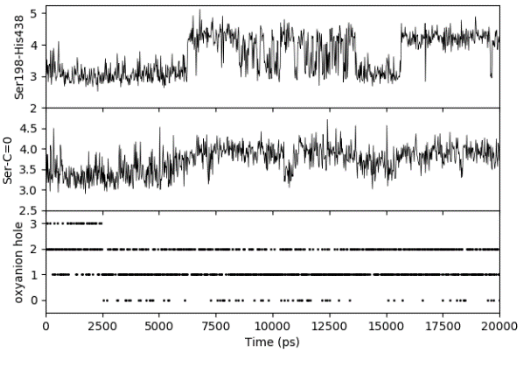

**7**

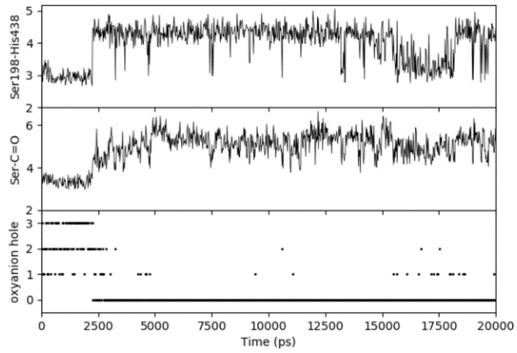

**10**

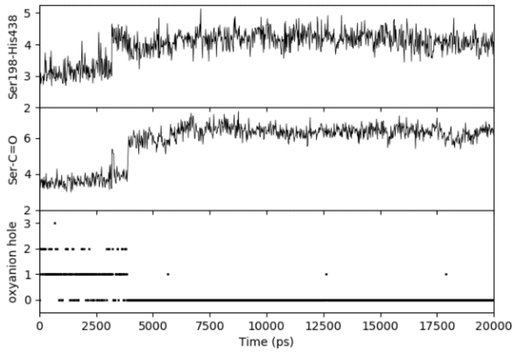

**11**

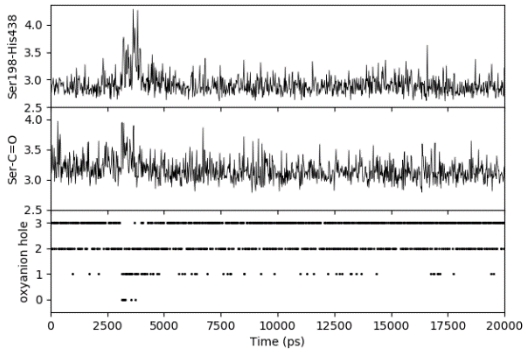

**12**

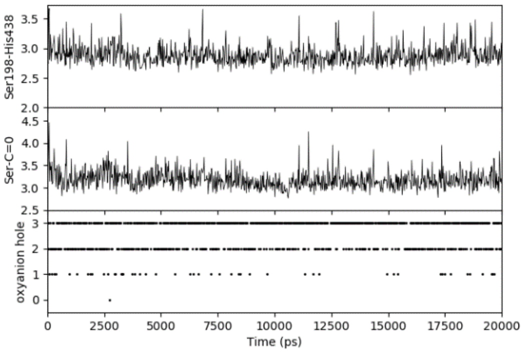

**13**

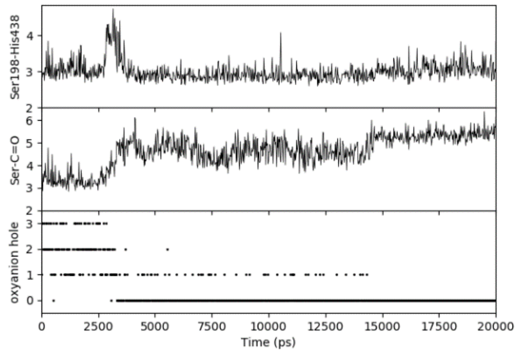

**14**

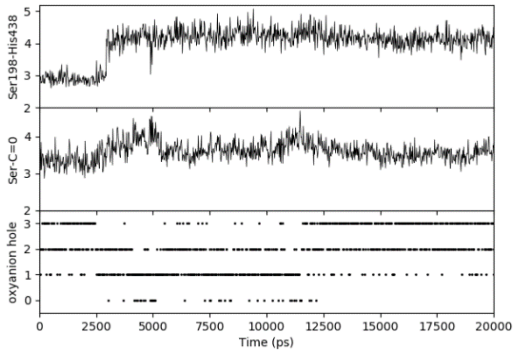

**15**

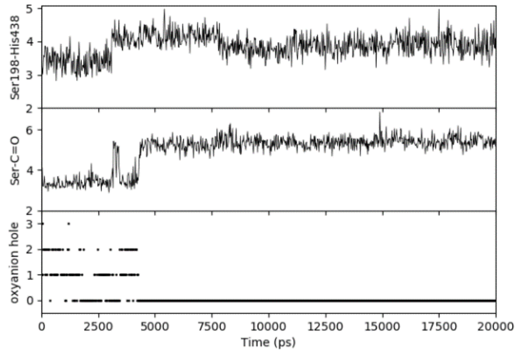

**16**

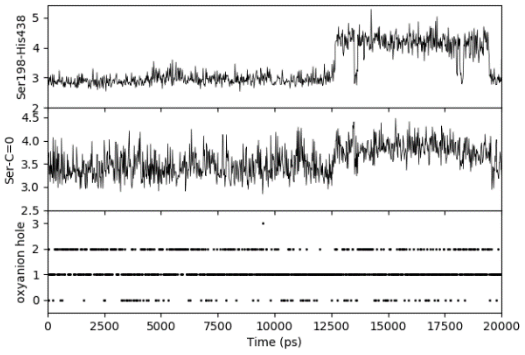

**17**

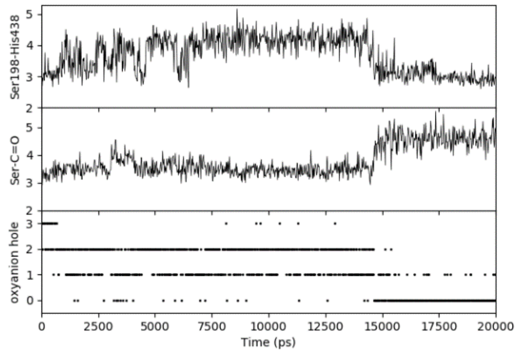

**18**

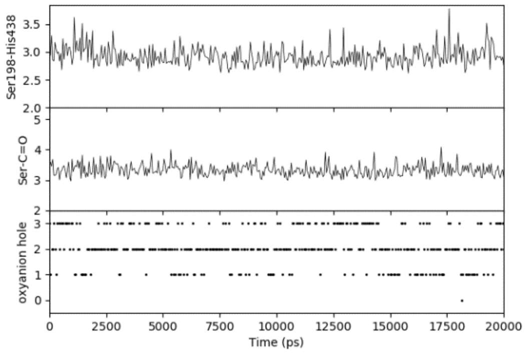

**19**

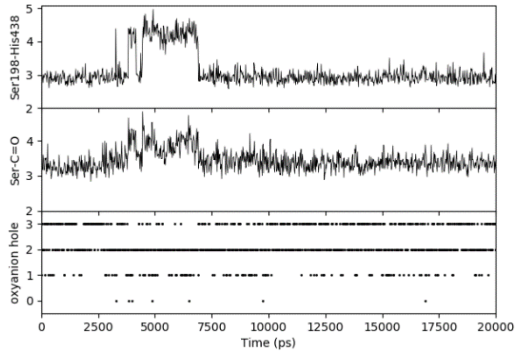

**20**

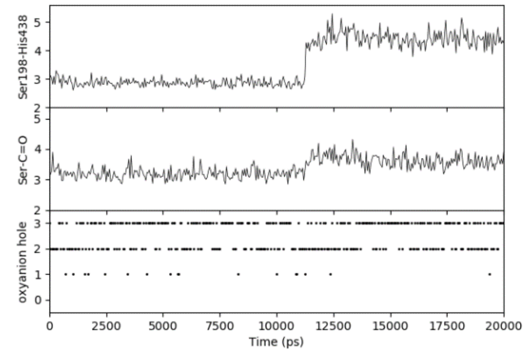

**21**

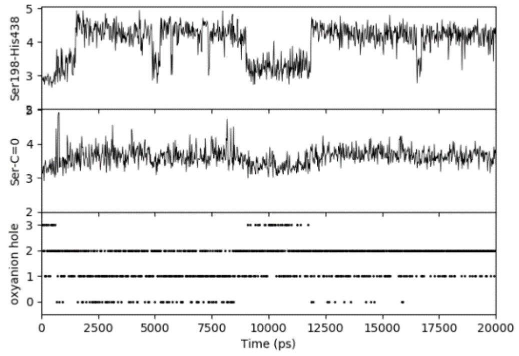

**22**

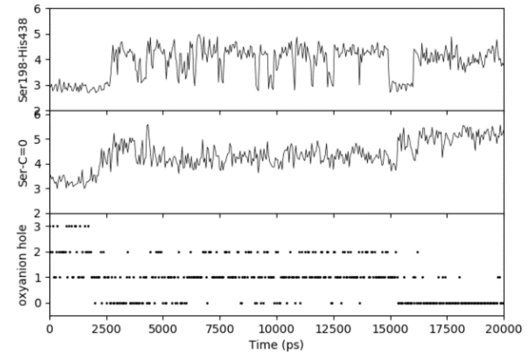

**23**

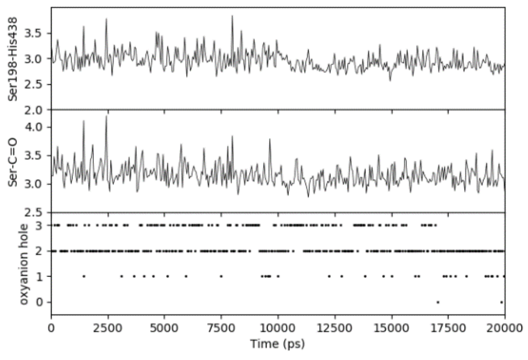

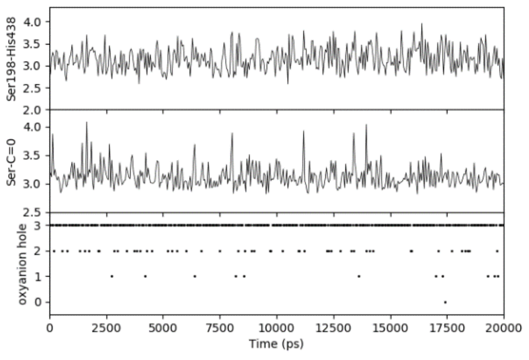

**26** **28**

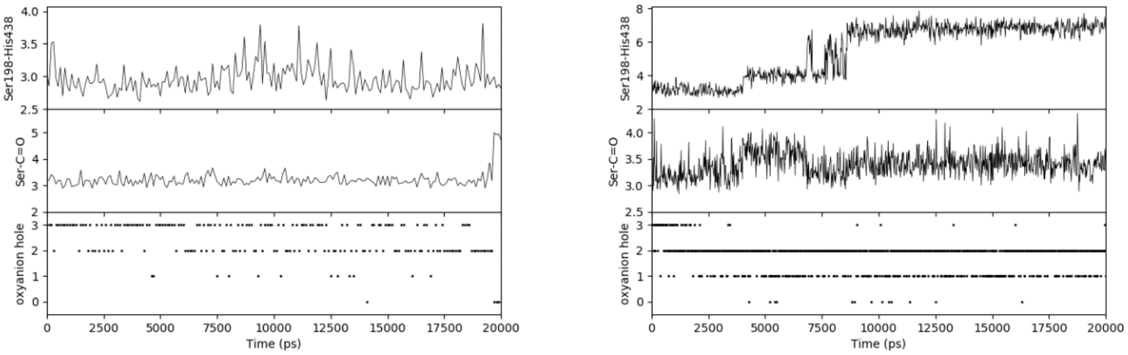

**29** **30**

**Fig. S14│**The analyses of short (20 ns) MD simulations of the pre-reaction noncovalent docking poses in hBChE binding site. The traces from top to bottom: Ser198-His438 – the distance between Ser198 $O^{\gamma}$ and His438 $N^{\varepsilon}$ (in Å, indicating the stability of the triad), Ser-C=O – the distance between Ser198 $O^{\gamma}$ and ligand's carbonyl carbon (in Å, indicating the propensity for covalent bond formation), and oxyanion hole – the number of hydrogen bonds formed with the oxyanion hole residues Gly116, Gly117, Ala199 (indicating the propensity for the activation of the carbonyl group and stabilisation of the intermediate).

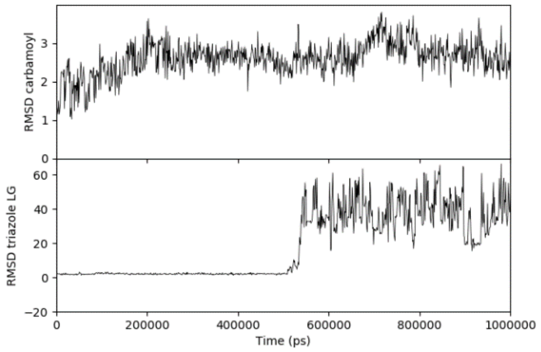

**Fig. S15│**1 µs MD simulation of the compound **28**-carbamoylated hBChE. The 1,2,4-triazole leaving group exits the enzyme active site, as evident from the large increase in its RMSD (and visual trajectory inspection), while the carbamate moiety remains quite stable.

**Table S5 │**The statistical analysis of the relationships between experimental or QM-calculated descriptors and activity ($pIC_{50}$) for compounds **6a–31a**. These were separarted into groups of covalent and non-covalent inhibitors, and Mann-Whitney comparison and Spearman non-parametric correlation were calculated. The statistically significant differences were considered at two-tailed p values <0.05 (*), <0.01 (**), <0.001 (***), or <0.0001 (****) – these descriptors are bolded, while non-significant difference is labelled "ns". Atom-related properties refer to the carbonyl carbon atom, see Scheme 3 for legend.

| **Descriptor** | **Mann-Whitney test** | | **Correlation with $pIC_{50}$** | |
|---|---|---|---|---|
| | **Two-tailed p value:** | **Significance** | **Spearman r** | **Significance** |
| $pIC_{50}$ | <0.0001 | **** | – | – |
| **Glide XP Gscore** | 0.0275 | * | 0.5040 | * |
| cdock affinity | 0.2591 | ns | 0.2548 | ns |
| **$^{13}$C δ** | 0,0407 | * | -0.4379 | * |
| $k_{hydr}$ [$h^{-1}$] | 0.4025 | ns | 0.01001 | ns |
| p$K_a$ | 0.2972 | ns | -0.3402 | ns |
| **$\Delta H_{INT}$** | <0.0001 | **** | -0.6062 | ** |
| **$\Delta G_{INT}$** | <0.0001 | **** | -0.6171 | *** |
| **$\Delta H_{ALL}$** | 0.0108 | * | -0.4003 | * |
| $\Delta G_{ALL}$ | 0.0234 | * | -0.3402 | ns |
| f_NN LUMO | 0.0849 | ns | 0.2677 | ns |
| Mulliken charge | 0.0464 | * | 0.3662 | ns |
| ESP charge | >0.9999 | ns | -0.001026 | ns |
| Stockholder charge | 0.0954 | ns | 0.3562 | ns |
| polarizability | 0.8859 | ns | 0.4826 | * |
| **maxESP** | 0.0004 | *** | 0.5245 | ** |
| minESP | 0.7710 | ns | 0.02838 | ns |
| natural charge | 0.8408 | ns | 0.1287 | ns |
| **ν [$cm^{-1}$]** | 0.0037 | ** | 0.5357 | ** |
| HOMO energy | 0.1147 | ns | -0.1566 | ns |
| LUMO energy | 0.0017 | ** | -0.3682 | ns |
| Chemical Hardness | 0.0202 | * | -0.2937 | ns |
| Electrophilicity index | 0.0020 | ** | 0.3874 | ns |
| **Electronic chemical potential** | 0.0020 | ** | -0.3990 | * |
| Local electrophilicity index | 0.0673 | ns | 0.2944 | ns |

| | | | | |
|---|---|---|---|---|
| Maximum electronic charge | 0.0020 | ** | 0.3593 | ns |

**Table S6 | **The Principal Component Regression analysis of statistically significantly different descriptors from both Mann-Whitney tests and Spearman correlation *vs*. $pIC_{50}$. Atom-related properties refer to the carbonyl carbon atom, see Scheme 3 for legend.

| **Analysis of Variance** | **SS** | **MS** | **F (DFn, DFd)** | **p value** | |
|---|---|---|---|---|---|
| **Regression** | 2.139 | 2.139 | F (1, 21) = 12.86 | p=0.0017 | |
| **Residual** | 3.493 | 0.1664 | | | |
| **Goodness of Fit** | | 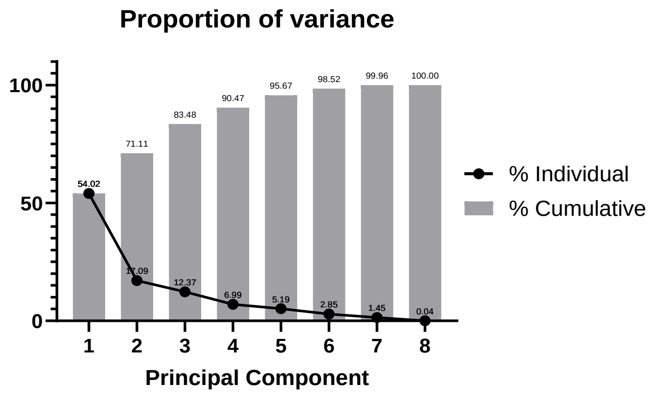<br> | | | |
| **R squared** | 0.3798 | | | | |
| **Sum of Squares** | 3.493 | | | | |
| **Sy.x** | 0.4079 | | | | |
| **Parameter estimates** | **Variable** | **Estimate** | **Standard error** | **p value** | **p value summary** |
| β0 | Intercept | 5.024 | 0.5824 | <0.0001 | **** |
| β1 | Glide XP Gscore | 0.01284 | 0.003580 | 0.0017 | ** |
| β2 | $^{13}C$ δ | -0.008364 | 0.002333 | 0.0017 | ** |
| β3 | $\Delta H_{INT}$ | -0.01146 | 0.003196 | 0.0017 | ** |
| β4 | $\Delta G_{INT}$ | -0.01097 | 0.003060 | 0.0017 | ** |
| β5 | $\Delta H_{ALL}$ | -0.005927 | 0.001653 | 0.0017 | ** |
| β6 | maxESP | 0.1079 | 0.03010 | 0.0017 | ** |
| β7 | ν [$cm^{-1}$] | 0.001414 | 0.0003944 | 0.0017 | ** |
| β8 | Electronic chemical potential | -5.314 | 1.482 | 0.0017 | ** |

**Table S7 | ** The DFT-calculated energies (M06-2X/cc-pVTZ(-f)++) for the species defined in Scheme 3.

| Species / Compound | 6a-31a | | INT | | END | | LG | |
|---|---|---|---|---|---|---|---|---|
| | $H_{tot}$ [Ha] | $G_{tot}$ [Ha] | $H_{tot}$ [Ha] | $G_{tot}$ [Ha] | $H_{tot}$ [Ha] | $G_{tot}$ [Ha] | $H_{tot}$ [Ha] | $G_{tot}$ [Ha] |
| **noncovalent** | | | | | | | | |
| **7a** | -1023.176489 | -1023.235661 | -1138.390826 | -1138.455490 | -686.105003 | -686.157810 | -452.298954 | -452.333907 |
| **8a** | -1360.238060 | -1360.303882 | -1475.458396 | -1475.527703 | -686.105003 | -686.157810 | -789.373760 | -789.416540 |
| **9a** | -1697.277679 | -1697.349883 | -1812.490695 | -1812.566127 | -686.105003 | -686.157810 | -1126.438619 | -1126.489040 |
| **10a** | -877.771194 | -877.829826 | -992.981889 | -993.042966 | -686.105003 | -686.157810 | -306.901514 | -306.935981 |
| **17a** | -1006.172582 | -1006.234352 | -1121.390896 | -1121.455028 | -686.105003 | -686.157810 | -435.306881 | -435.344490 |
| **18a** | -1048.351837 | -1048.408193 | -1163.560227 | -1163.619736 | -686.105003 | -686.157810 | -477.468086 | -477.498986 |
| **19a** | -1346.147049 | -1346.207152 | -1461.357494 | -1461.421242 | -686.105003 | -686.157810 | -775.272186 | -775.308542 |
| **20a** | -1200.736524 | -1200.796545 | -1315.950428 | -1316.012940 | -686.105003 | -686.157810 | -629.865320 | -629.901212 |
| **21a** | -1234.037430 | -1234.100036 | -1349.247438 | -1349.311938 | -686.105003 | -686.157810 | -663.173342 | -663.210546 |
| **24a** | -3051.770743 | -3051.824747 | -3166.981977 | -3167.040815 | -686.105003 | -686.157810 | -2480.901210 | -2480.933190 |
| **25a** | -3204.158627 | -3204.216151 | -3319.370352 | -3319.432484 | -686.105003 | -686.157810 | -2633.295421 | -2633.331540 |
| **26a** | -796.563388 | -796.618003 | -911.787439 | -911.846590 | -686.105003 | -686.157810 | -225.684572 | -225.715027 |
| **covalent** | | | | | | | | |
| **6a** | -670.872389 | -670.921157 | -786.102598 | -786.155526 | -686.105003 | -686.157810 | -100.000041 | -100.016560 |
| **11a** | -1374.006302 | -1374.073755 | -1489.224393 | -1489.294367 | -686.105003 | -686.157810 | -803.155120 | -803.199341 |
| **12a** | -891.619066 | -891.675695 | -1006.840913 | -1006.901564 | -686.105003 | -686.157810 | -320.770125 | -320.802606 |

| | | | | | | | | |
|---|---|---|---|---|---|---|---|---|
| **13a** | -1230.690762 | -1230.748271 | -1345.917188 | -1345.978946 | -686.105003 | -686.157810 | -659.843104 | -659.876541 |
| **14a** | -871.729441 | -871.786570 | -986.957737 | -987.018541 | -686.105003 | -686.157810 | -300.862309 | -300.894908 |
| **15a** | -871.711477 | -871.768633 | -986.940705 | -987.001261 | -686.105003 | -686.157810 | -300.846396 | -300.878913 |
| **16a** | -1041.325345 | -1041.387080 | -1156.555604 | -1156.619647 | -686.105003 | -686.157810 | -470.470384 | -470.508509 |
| **22a** | -1250.080903 | -1250.142002 | -1365.295154 | -1365.360127 | -686.105003 | -686.157810 | -679.226843 | -679.263767 |
| **23a** | -1266.106272 | -1266.167240 | -1381.324145 | -1381.388880 | -686.105003 | -686.157810 | -695.255375 | -695.292194 |
| **27a** | -1001.065336 | -1001.125206 | -1116.299784 | -1116.364342 | -686.105003 | -686.157810 | -430.200031 | -430.236610 |
| **28a** | -812.612661 | -812.667070 | -927.847522 | -927.905054 | -686.105003 | -686.157810 | -241.747327 | -241.777568 |
| **29a** | -812.584146 | -812.638463 | -927.815943 | -927.874201 | -686.105003 | -686.157810 | -241.720237 | -241.749863 |
| **30a** | -828.626747 | -828.681226 | -943.869532 | -943.928169 | -686.105003 | -686.157810 | -257.781244 | -257.811317 |
| **31a** | -966.159217 | -966.218529 | -1081.391011 | -1081.454864 | -686.105003 | -686.157810 | -395.297420 | -395.333611 |
| **methoxide** | -115.2118230 | -115.2369210 | | | | | | |

**Table S8 │** Molecular descriptors of the fragment-sized warheads **6a–31a**. Atom-related properties refer to the carbonyl carbon atom, see Scheme 3 for legend.

| **Compound** | **pIC$_{50}$** | **Glide XP Gscore** | **cdock affinity** | **$^{13}$C δ [ppm]** | **$k_{hydr}$ [h$^{-1}$]** | **p$K_a$** | **Mulliken charge** | **ESP charge** | **Stockholder charge** | **natural charge** | **f_NN LUMO** |
|---|---|---|---|---|---|---|---|---|---|---|---|
| **non-covalent** | | | | | | | | | | | |
| **7a** | 6.438779 | -10.246 | -10.182 | 154.37 | 0.001314 | 12.3 | 6.438779 | 0.94743 | 0.22351 | 0.9344 | -0.0693 |
| **8a** | 6.790485 | -8.313 | -8.371 | 152.72 | 0 | 9.3 | 6.790485 | 0.80331 | 0.22008 | 0.92765 | -0.0468 |
| **9a** | 6.756218 | – | -7.358 | 148.94 | 0.106651 | 5.2 | 6.756218 | 0.66993 | 0.21511 | 0.93214 | -0.0154 |
| **10a** | 6.891097 | -8.941 | -10.815 | 151.36 | 0 | 9.995 | 6.891097 | 0.71984 | 0.2209 | 0.92766 | -0.0407 |

| | | | | | | | | | | | |
|---|---|---|---|---|---|---|---|---|---|---|---|
| **17a** | 6.609065 | -9.034 | -11.991 | 150.51 | – | 6 | 6.609065 | 0.76306 | 0.21714 | 0.91684 | -0.0387 |
| **18a** | 6.654822 | -9.467 | -9.49 | 170.71 | 0.0008 | 10.78 | 6.654822 | 0.53039 | 0.14392 | 0.56877 | -0.0107 |
| **19a** | 7.096693 | -9.429 | -10.609 | 166.94 | 0 | 7.3 | 7.096693 | 0.35905 | 0.14684 | 0.58379 | 0.0279 |
| **20a** | 6.414878 | -9.651 | -10.457 | 169.32 | 0.049407 | 6.61 | 6.414878 | 0.3584 | 0.14595 | 0.58587 | 0.0439 |
| **21a** | 6.352617 | -9.832 | -10.101 | 161.64 | 0.001264 | 12.37 | 6.352617 | 0.38572 | 0.151 | 0.59671 | 0.0348 |
| **24a** | 6.740167 | – | – | 167.94 | 0.01064 | 4.2 | 6.740167 | 0.41145 | – | – | 0.0643 |
| **25a** | 6.400444 | – | – | 167.19 | 0.090731 | 5.9 | 6.400444 | 0.4507 | 0.13003 | – | 0.0608 |
| **26a** | 6.620151 | -9.758 | -9.858 | 150.54 | 0.002123 | 14.52 | 6.620151 | 0.442 | 0.20381 | 0.84042 | -0.0487 |
| **6a** | 8.083336 | -8.406 | -10.262 | 148.93 | 0.006433 | 3.17 | 8.083336 | 0.75665 | 0.23586 | 0.99267 | -0.0386 |
| **11a** | 7.252744 | -3.389 | -10.195 | 151.81 | 0.050804 | 5.53 | 7.252744 | 0.66956 | 0.21999 | 0.92834 | 0.1217 |
| **12a** | 7.277366 | -8.537 | -10.316 | 151.78 | 0.004928 | 5.85 | 7.277366 | 0.74035 | 0.2299 | 0.9274 | 0.0151 |
| **13a** | 7.363211 | -8.583 | -10.434 | 151.04 | 0.001398 | 6.79 | 7.363211 | 0.72332 | 0.22442 | 0.92676 | 0.0082 |
| **14a** | 6.417255 | -9.349 | -9.739 | 153.27 | 0.001641 | 8.82 | 6.417255 | 0.74823 | 0.22095 | 0.9152 | -0.0489 |
| **15a** | 7.775985 | -10.091 | -9.848 | 153.53 | 0.001073 | 6.1 | 7.775985 | 0.64975 | 0.21776 | 0.91525 | -0.0336 |
| **16a** | 7.713993 | -8.424 | -8.424 | 152.27 | 0.012985 | 7.4 | 7.713993 | 0.67307 | 0.22137 | 0.91562 | 0.0133 |
| **22a** | 6.960983 | -7.69 | -10.309 | 166.41 | 0.001156 | 6.89 | 6.960983 | 0.29117 | 0.15435 | 0.60455 | 0.0513 |
| **23a** | 7.514137 | -8.843 | -9.339 | 162.01 | 0.000763 | 3.31 | 7.514137 | 0.29809 | 0.1602 | 0.6085 | 0.0733 |
| **27a** | 6.8671 | -10.067 | -10.067 | 148.51 | 0.017267 | 9.34 | 6.8671 | 0.61605 | 0.21308 | 0.84664 | 0.0161 |
| **28a** | 7.258848 | -9.046 | -9.324 | 149.53 | 0.009685 | 10.26 | 7.258848 | 0.49937 | 0.20483 | 0.83456 | 0.1628 |
| **29a** | 7.43866 | -7.707 | -8.06 | 150.26 | 0.002044 | 13.9 | 7.43866 | 0.46773 | 0.19959 | 0.83049 | 0.1518 |
| **30a** | 7.719649 | -8.423 | -9.35 | 146.77 | 0.143364 | 4.83 | 7.719649 | 0.20848 | 0.21197 | 0.82837 | 0.1416 |

| | | | | | | | | | | | |
|---|---|---|---|---|---|---|---|---|---|---|---|
| **31a** | 7.547907 | -7.381 | -7.381 | 150.95 | 0.007546 | 8.2 | 7.547907 | 0.56992 | 0.20428 | 0.83748 | 0.042 |

**Table S8** (*continued)*

| **Compound** | **polarizability [au]** | **maxESP [eV]** | **ν [cm$^{-1}$]** | **HOMO energy [Ha]** | **LUMO energy [Ha]** | **Chemical Hardness** | **Electrophilicity index** | **Electronic chemical potential** | **Local electrophilicity index** | **Maximum electronic charge** |
|---|---|---|---|---|---|---|---|---|---|---|
| **non-covalent** | | | | | | | | | | |
| **7a** | -0.367 | 0.83 | 1727.618 | -0.26889 | -0.00883 | 0.26006 | 0.037072 | -0.13886 | -0.00257 | 0.533954 |
| **8a** | 1.419 | 1.03 | 1753.821 | -0.27031 | -0.00863 | 0.26168 | 0.037167 | -0.13947 | -0.00174 | 0.532979 |
| **9a** | 1.49 | 0.9 | 1786.864 | -0.27059 | -0.00785 | 0.26274 | 0.036885 | -0.13922 | -0.00057 | 0.529877 |
| **10a** | 0.553 | 0.84 | 1734.314 | -0.2688 | -0.00898 | 0.25982 | 0.037123 | -0.13889 | -0.00151 | 0.534562 |
| **17a** | -0.59 | 0.34 | 1785.973 | -0.27144 | -0.00962 | 0.26182 | 0.037714 | -0.14053 | -0.00146 | 0.536743 |
| **18a** | -0.415 | 0.48 | 1674.558 | -0.27083 | -0.01248 | 0.25835 | 0.038835 | -0.14166 | -0.00042 | 0.548307 |
| **19a** | 0.732 | 0.39 | 1697.435 | -0.27265 | -0.01285 | 0.2598 | 0.039218 | -0.14275 | 0.001094 | 0.549461 |
| **20a** | -0.474 | 0.45 | 1692.583 | -0.27147 | -0.0136 | 0.25787 | 0.039392 | -0.14254 | 0.001729 | 0.55274 |
| **21a** | | 0.4 | 1709.018 | -0.27275 | -0.0142 | 0.25855 | 0.039809 | -0.14348 | 0.001385 | 0.554922 |
| **24a** | | 0.08 | 1672.346 | -0.26995 | -0.01288 | 0.25707 | 0.038896 | -0.14142 | 0.002501 | 0.550103 |
| **25a** | -0.86 | 0.52 | 1680.611 | -0.26744 | -0.01314 | 0.2543 | 0.038697 | -0.14029 | 0.002353 | 0.551671 |
| **26a** | 1.134 | 1.72 | 1720.608 | -0.27171 | -0.00857 | 0.26314 | 0.037317 | -0.14014 | -0.00182 | 0.532568 |
| **covalent** | | | | | | | | | | |
| **6a** | 1.585 | 1.19 | 1808.723 | -0.2714 | -0.00905 | 0.26235 | 0.037475 | -0.14023 | -0.00145 | 0.534496 |
| **11a** | -0.262 | 1.15 | 1767.023 | -0.27077 | -0.02027 | 0.2505 | 0.042268 | -0.14552 | 0.005144 | 0.580918 |
| **12a** | 0.202 | 1.05 | 1755.105 | -0.2685 | -0.00967 | 0.25883 | 0.037369 | -0.13909 | 0.000564 | 0.53736 |

| | | | | | | | | | | |
|---|---|---|---|---|---|---|---|---|---|---|
| **13a** | 0.189 | 0.87 | 1761.629 | -0.27047 | -0.03906 | 0.23141 | 0.051753 | -0.15477 | 0.000424 | 0.668791 |
| **14a** | -0.172 | 1.45 | 1791.615 | -0.27254 | -0.0104 | 0.26214 | 0.038174 | -0.14147 | -0.00187 | 0.539673 |
| **15a** | 0.398 | 1.33 | 1792.11 | -0.27231 | -0.00985 | 0.26246 | 0.037917 | -0.14108 | -0.00127 | 0.53753 |
| **16a** | 0.435 | 0.97 | 1800.906 | -0.2722 | -0.03756 | 0.23464 | 0.051116 | -0.15488 | 0.00068 | 0.660075 |
| **22a** | -0.674 | 0.62 | 1719.441 | -0.27278 | -0.01545 | 0.25733 | 0.040355 | -0.14412 | 0.00207 | 0.56004 |
| **23a** | 0.315 | 1.19 | 1748.668 | -0.27503 | -0.01717 | 0.25786 | 0.041389 | -0.1461 | 0.003034 | 0.566587 |
| **27a** | -0.303 | 1.75 | 1748.822 | -0.27387 | -0.066 | 0.20787 | 0.069461 | -0.16994 | 0.001118 | 0.817506 |
| **28a** | 0.101 | 1.88 | 1736.136 | -0.27077 | -0.0174 | 0.25337 | 0.040969 | -0.14409 | 0.00667 | 0.568674 |
| **29a** | 0.96 | 0.94 | 1747.909 | -0.2707 | -0.02472 | 0.24598 | 0.04435 | -0.14771 | 0.006732 | 0.600496 |
| **30a** | 0.231 | 2.53 | 1758.344 | -0.27267 | -0.02991 | 0.24276 | 0.047143 | -0.15129 | 0.006675 | 0.623208 |
| **31a** | 0.155 | 1.51 | 1737.54 | -0.271 | -0.04184 | 0.22916 | 0.053385 | -0.15642 | 0.002242 | 0.68258 |

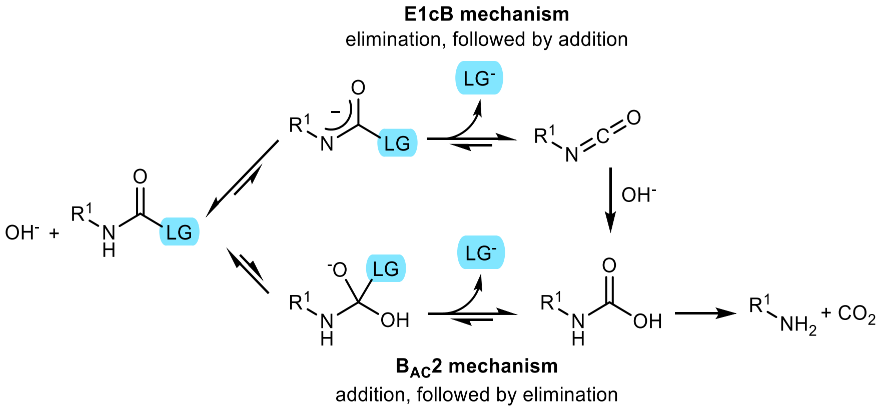

**Scheme S2 |** B$_{AC}$2 and E1cB mechanisms of base-catalysed carbamate hydrolysis.

**Table S9 | ** DFT-calculated reaction energy barriers.

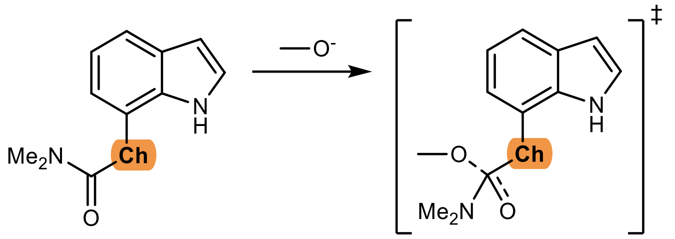

| species | | reactant | | transition state | | | |
|---|---|---|---|---|---|---|---|
| | | $H_{tot}$ [Ha] | $G_{tot}$ [Ha] | $H_{tot}$ [Ha] | $G_{tot}$ [Ha] | $\Delta H^{‡}$ [kcal/mol] | $\Delta G^{‡}$ [kcal/mol] |
| **Ch =** | **Level of theory: M06-2X/LACV3P**++** | | | | | | |
| **O** | **1a** | -686.058542 | -686.112148 | -801.250705 | -801.307328 | 6.96 | 20.82 |
| **S** | **2a** | -1009.022579 | -1009.077658 | -1124.204933 | -1124.264785 | 13.12 | 25.87 |
| **Se** | **3a** | -620.01383 | -620.069304 | -735.200486 | -735.261544 | 10.42 | 22.66 |
| **Te** | **4a** | -618.834255 | -618.89134 | -734.021471 | -734.083408 | 10.07 | 22.77 |
| | **MeO⁻** | -115.203258 | -115.228357 | | | | |

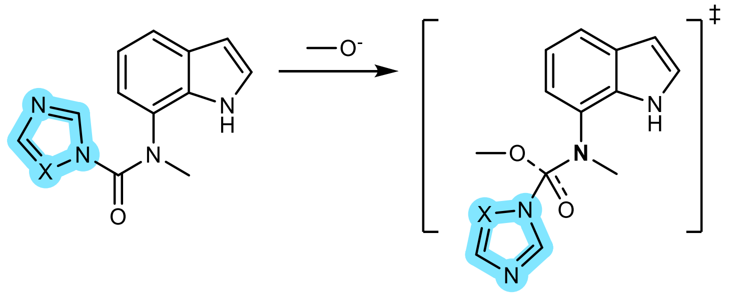

| | | | | | | | |
|---|---|---|---|---|---|---|---|
| **X =** | **Level of theory: M06-2X/cc-pVTZ(-f)++** | | | | | | |
| **CH** | **26a** | -796.563388 | -796.618003 | -911.77338 | -911.83209 | 1.149 | 14.33 |
| **N** | **28a** | -812.612661 | -812.66707 | -927.821 | -927.88 | 1.921 | 15.02 |
| | **MeO⁻** | -115.212 | -115.237 | | | | |

*Aqueous stability*

**Table S10 |** The stability of the investigated compounds in an aqueous solution.

| Compound | $k$ [h$^{-1}$] | $t_{1/2}$ |
|---|---|---|
| **1** | 0.00090 | ≈ 32 days |
| **2** | stable | |
| **3** | stable | |
| **4** | stable | |
| **6** | 0.00643 | ≈ 4 days |
| **6 + 5 mM GSH** | 0.14160 | 4.9 h |
| **8** | stable | |
| **9** | 0.10665 | 6.5 h |
| **10** | stable | |
| **11** | 0.05080 | 13.6 h |
| **12** | 0.00493 | ≈ 6 days |
| **13** | 0.00140 | ≈ 21 days |
| **14** | 0.00164 | ≈ 18 days |
| **15** | 0.00107 | ≈ 27 days |
| **16** | 0.01299 | 53.4 h |
| **18** | 0.00080 | ≈ 36 days |
| **19** | stable | |
| **20** | 0.04941 | 14.0 h |
| **21** | 0.00126 | ≈ 23 days |
| **22** | 0.00116 | ≈ 25 days |
| **23** | 0.00076 | ≈ 38 days |
| **24** | 0.01064 | 65.1 h |
| **25** | 0.09073 | 7.6 h |
| **26** | 0.00212 | ≈ 14 days |
| **27** | 0.01727 | 40.1 h |
| **28** | 0.00968 | 71.6 h |
| **28 + 5 mM GSH** | 0.02729 | 25.4 h |
| **29** | 0.00204 | ≈ 14 days |
| **30** | 0.14336 | 4.8 h |
| **31** | 0.00755 | 91.9 h |

### Supplementary Methods

#### Inhibition of ChEs

##### *$IC_{50}$ determination*

The inhibitory potencies of the compounds against the ChEs were determined using the method of Ellman following the procedure described previously.[21] Briefly, compound stock solutions in DMSO were incubated with Ellman's reagent and the ChEs (final concentrations: 370 µM Ellman's reagent, approximately 1 nM or 50 pM hBChE or hAChE, respectively) in 0.1 M sodium phosphate pH 8.0 for 5 min at 20 °C. For time-dependency measurements, the pre-incubation time was varied (1 min, 5 min, 15 min, 30 min or 60 min). The reactions were started by the addition of the substrate (final concentration, 500 µM butyrylthiocholine iodide (BTChI) or acetylthiocholine iodide for hBChE and hAChE, respectively). The final content of DMSO was always 1%. The increase in absorbance at 412 nm was monitored for 2 min using a microplate reader (Synergy HT, BioTek Instruments, VT, USA). The initial velocities in the presence ($v_i$) and absence ($v_o$) of the test compounds were calculated. The inhibitory potencies were expressed as the residual activities, according to RA = $(v_i - b) / (v_o - b)$, where $b$ is the blank value using phosphate buffer without ChEs. For $IC_{50}$ determinations, at least seven different concentrations for each compound were used. The $IC_{50}$ values were obtained by plotting the residual ChE activities against the applied inhibitor $\log_{10}$ concentrations, with the experimental data fitted to a four-parameter logistic function (GraphPad Prism 9.4). Rivastigmine and donepezil were used as positive controls.

##### *Initial velocity measurements*

Initial velocity measurements were measured on an Agilent Cary 3500 UV-VIS spectrophotometer with Compact Peltier module (thermostatted at 25 °C) at 412 nm using a 0.6 mL 0.5 cm cuvette. 30 mM Tris buffer pH 7.0 with 1 mg/mL bovine serum albumin (BSA) and 0.02% $NaN_3$ was used and all dilutions were done in glass vessels (note: dilute hBChE solutions rapidly lose activity, presumably due to adsorption to plastic, preventing meaningful decarbamoylation assays. In our hands, under the above conditions only, the diluted and stock solutions of hBChE were stable for at least a month at room temperature.). Total volume in the cuvette was always 500 µL, and enzyme solutions that gave upon dilution $v_0 = (dA/dt) \approx 0.5$ (for carbamoylation) and ≈ 1 for (for decarbamoylation) were used. For carbamoylation experiments, 0.5–1 µL of enzyme stock, 5 µL of compound solution, and 484 µL of buffer were preincubated for the specified time, then 10 µL of the mixture of 25 mM DTNB and 20 mM BTChI (large excess to prevent substrate depletion) were added, quickly mixed, and measurement started. For decarbamoylation experiments, 50 µL of the enzyme stock solution and 0.25–0.5 µL of the compound solution in MeOH (concentration chosen to achieve 85–95% inhibition, typically 0.5–1 mM) were incubated for 1 h at room temperature, then the mixture was diluted 1:1000 with the buffer, and 500 µL aliquots were drawn at different timepoint, mixed with 10 µL of the mixture of 25 mM Ellman's reagent (DTNB) and 20 mM butyrylthiocholine, and hBChE activity assayed. The data analysis followed the literature.[1,2,111]

##### *Progress curve analysis*

The progress curves for the time courses of product formation for the hydrolysis of butyrylthiocholine by purified hBChE (~1 nM) were measured on a Perkin-Elmer spectrophotometer (Lambda 45 UV/Vis), using a 0.6 mL cuvette, 1 mM DTNB, 50 µM BTChI at 25 °C in 25 mM phosphate buffer, pH 7.0, at 412 nm using a molar absorption coefficient of thionitrobenzoate anion equal to 13800 L $mol^{-1}$ $cm^{-1}$. A 10 s lag period was considered before the start of each measurement due to technical restrictions of the spectrophotometer. The progress curves were extrapolated to the origin. The conditions were selected so that the curve in the absence of inhibitor always achieved the plateau, indicating complete substrate hydrolysis. The following constants were kept fixed: $K_S$ = 7.5 µM, $k_{cat}$ = 814 $s^{-1}$, $K_P$ = 38 µM, and the rest were fitted using the ENZO program.[111,113] Experimental Gibbs free energy activation barrier $\Delta G^‡$ was calculated from the (de)carbamoylation rates *via* the Eyring equation (taking reference concentration of 1 M, at 298.15 K).

**Activity-based protein profiling**

The human neuroblastoma SH-SY5Y and human hepatocellular carcinoma HepG2 cell lines were purchased from American Type Culture Collection (Manassas, VA, USA). Cells were cultured in Advanced Dulbecco's modified Eagle's medium (Gibco, Thermo Fisher Scientific, Waltham, MA, USA) supplemented with 10% fetal bovine serum (FBS, Gibco), 2 mM L-glutamine, 50 U/mL penicillin and 50 µg/mL streptomycin (Sigma, St. Louis, MO, USA) in a humidified atmosphere of 95% air and 5% $CO_2$ at 37 °C, and grown to 80% confluence. One day prior to cell harvesting, complete medium was replaced with a serum-free medium. The medium was removed, cell layer rinsed once with Ca- and Mg-free Dulbecco’s phosphate buffered saline (PBS), cells harvested in PBS/EDTA for 10 min at 37 °C, and centrifuged at 4 °C, 1000 × *g* for 5 min. The pellet was resuspended in lysis buffer[114] (20 mM HEPES pH 7.2, 2 mM DTT, 1 mM $MgCl_2$, 25 U/mL benzonase), sonicated with a probe sonicator (~30% power) on ice, and centrifuged at 4 °C, 15,000 × g for 15 min. The supernatant was stored at –80 °C. Blood was collected from the author in $K_2$EDTA Vaccutainer tubes, left for 5 min, and centrifuged at room temperature, 3000 × g for 10 min. Plasma was separated and stored at –20 °C. The recombinant hBChE[21] was diluted to 4 µg/mL in lysis buffer. The protein concentrations in each of the samples were quantified by DC Protein Assay (Bio-Rad) according to manufacturer’s instructions.

Preheated control samples were heated at 80 °C for 5 min and cooled on ice before incubation with the respective probe. Probe stock solutions were prepared in DMSO (10 mM), aliquoted, and stored at –20 °C (stable for several weeks). Protein samples (except rhBChE) were adjusted to 2 mg/mL concentration using lysis buffer, and probe stock solution (0.3 µL) in DMSO of the appropriate concentration were added to the protein samples, keeping the total volume of 30 µL and 1% final DMSO concentration.

30 µL of the sample solution and 6 µL of the 6× loading buffer (125 mM Tris pH 6.8, 20% glycerol, 4% SDS, 0.005% bromophenol blue) were thoroughly vortexed, and heated for 5 min at 100 °C before loading onto the gel. Polyacrylamide gels were prepared in the following way: 12% acrylamide separating gel (375 mM Tris pH 8.8, 0.125% SDS, 0.125% ammonium persulfate, 425 ppm TEMED) was poured into 1.5 mm casting molds (8 mL/mold, Mini-

PROTEAN®, BioRad, Hercules, CA, USA), covered with 1 mL of isopropanol, and left to polymerize for 25 min, then 15-well comb was inserted, covered with 5% acrylamide stacking gel (375 mM Tris pH 6.8, 0.1% SDS, 0.1% ammonium persulfate, 1000 ppm TEMED), and left to polymerize for 25 min. 30 µL of samples and 10 µL of protein standard ladder (SeeBlue™ Plus2 Pre-stained Protein Standard, Invitrogen) were loaded into the wells and the electrophoresis was run at 100 V for 2.5 h at room temperature in a Mini-PROTEAN® Tetra cell (BioRad, Hercules, CA, USA) filled with SDS-PAGE buffer (24 mM Tris, 192 mM glycine, 0.1% SDS). The proteins were then transferred onto a nitrocellulose membrane using a dry Western blot system (iBlot 3 Western Blot Transfer System, Thermo Fischer Scientific). Fluorescence was observed under Epi Green illumination (15 s exposure) and a 595 nm filter in a fluorescent imager (Uvitec Cambridge Alliance 9.7, Uvitec, Lodi, NJ, USA).

For immunodetection of GAPDH and hBChE, the membranes were first incubated in 5% non-fat dry milk in Tris buffered saline with Tween 20 (TTBS; 25 mM Tris pH 7.4, 137 mM NaCl, 3 mM KCl, 0.1% Tween 20) for 1 h at room temperature with gentle shaking. The membranes were washed thrice with TTBS buffer for 5 min at room temperature, followed by overnight incubation at 4 °C in the primary antibody solution (1:5000 rabbit anti-GAPDH antibodies, Proteintech 10494-1-AP, in 3% BSA for the HepG2 and SH-SY5Y lysate membranes or 1:1000 rabbit anti-hBChE antibodies, Abcam ab317016, in 3% BSA or the rhBChE and plasma membrane) with gentle shaking. The membranes were then washed three times with TTBS buffer for 5 min at room temperature and incubated in the secondary antibody solution (1:20000 goat anti-rabbit IgG-HRP conjugate, Jackson ImmunoResearch 111-035-045, in 5% milk in TTBS) for 1 h at room temperature with gentle shaking. The membranes were finally washed six times with TTBS buffer for 5 min at room temperature and exposed to 700 µL of the chemiluminescent reagent (SuperSignal™ West Dura Extended Duration Substrate, Thermo Fisher Scientific for the HepG2 and SH-SY5Y lysate membranes and SuperSignal™ West Femto Maximum Sensitivity Substrate, Thermo Fisher Scientific for the rhBChE and plasma membrane) for up to 5 min. Chemiluminescence was observed in an imaging device (G-box, Syngene, 20 s exposure) in the dark.

**MS proteomic analysis**

Human neuroblastoma SH-SY5Y and human hepatocellular carcinoma HepG2 cell lysates, prepared as described above, were mixed in a 1:1 protein ratio, and diluted with equilibration buffer (50 mM Tris, 150 mM NaCl, 0.1% Triton X-100, pH 7.4) to the final concentration of 2 mg/mL. Biotinylated probes (2 mM stock in MeOH, final concentration: 20 µM) were added to 500 µL aliquots of the proteome solution in Eppendorf tubes and incubated for 1 h at room temperature with shaking.

1 mL aliquots of equilibrated SoftLink™ Soft Release Avidin Resin (Promega, WI, USA) (50% slurry in equilibration buffer) were then added and incubated for 1.5 h at room temperature with shaking. The suspensions were transferred to 5 mL columns and the flow-through was discarded. The resin was successively washed with 6 mL of equilibration buffer, 6 mL of glycine pH 3.0 buffer (0.1 M glycine hydrochloride, 1 M NaCl, 0.1% Triton X-100, pH 3.0), and 3 mL equilibration buffer. 500 µL of elution buffer (5 mM biotin, 50 mM Tris, 150 mM

NaCl, 0.1% Triton X-100, pH 7.4) was then added to the resin and incubated for 30 min at room temperature. The eluate was collected, the resin washed with another 500 μL of elution buffer, and the combined eluate was kept at –80 °C until MS analysis. The eluates were concentrated using centrifugation filters (Amicon® Ultra-4 Centrifugal Filter Unit, 10 kDa cutoff), separated on a Novex Tris-Glycine 8–16% gradient gel (Invitrogen), and prepared for mass spectrometry analysis as described previously.[115]

LC-MS/MS analysis was performed on an EASY-nanoLC II HPLC unit (Thermo Fischer Scientific) coupled to an Orbitrap LTQ Velos mass spectrometer (Thermo Fischer Scientific). The samples (3 μL) were loaded onto a C18 trapping column (Proxeon Easy-column, Thermo Fischer Scientific, 0.1 × 20 mm, 5 μm, 120 Å C18-A1) and separated on a C18 PicoFrit Aquasil analytical column (New Objective, 0.075 × 100 mm, 5 μm, 100 Å). The peptides were eluted using a 90 min gradient of 5–40 % acetonitrile with 0.1% formic acid at a flow rate of 0.3 mL/min. The full MS mass spectra were acquired with the Orbitrap mass analyser in the mass range of 300 to 2000 m/z at resolution of 30000 in the profile mode. The MS/MS spectra were obtained by HCD fragmentation of the nine most intense MS precursor ions and recorded at resolution of 7500 in the centroid mode. Only the precursor ions with assigned charge state (> 1) were chosen for MS/MS fragmentation. The dynamic exclusion was set to repeat count of 1, repeat duration of 30 s, and exclusion duration of 20 s.

Database searches were performed using the MaxQuant software package version 2.0.3.0 imbedded with the Andromeda search engine[116–118] against the UniProt-derived human and mouse reference proteome (UniProtKB, *Homo sapiens,* canonical database containing 20 371 entries, obtained on 03. 06. 2022). The peptides were identified using the Try (K, R) cleavage setting. The search engine parameters were set to consider carbamidomethylation of cysteine residues (+57.0215 Da) as fixed modification, oxidation of methionine (+15.9949 Da), and acetylation of protein *N*-termini (+42.0106 Da). The precursor ion and fragment ion mass tolerances were set as 4.5 ppm and 0.5 Da, respectively. A reversed database search was performed with 1% false discovery rate for protein and peptide identification.

Potential contaminants (e.g., keratins and trypsin from sample preparation) and endogenously biotinylated proteins (pyruvate carboxylase, 3-methylcrotonyl CoA carboxylase, propionyl CoA carboxylase, acetyl CoA carboxylase) were removed prior to the analysis, and LFQ intensities of 0 were converted to 0.1 to avoid division by 0. To eliminate nonspecific non-covalent binding due to the probe scaffold, the LFQ intensities for probe-treated samples were divided by the LFQ intensities produced by negative control **B2**, and intensities ratios (enrichments) >10 were considered significant. Protein function was assigned based on the information found on UniProtKB.[112]

##### Carbamoylation mass spectrometry study of hBChE

Compound stock solutions (10 mM) were prepared in 100% MeOH and further diluted in deionized water. To avoid hydrolysis of the bound carbamate during proteolytic sample preparation, whole protein MS analysis using recombinant hBChE produced in *E. coli*[119] was undertaken. The non-glycosylated protein from a prokaryote expression system was needed to produce a uniform multi-charged protein spectrum. hBChE was diluted down to 5 µM in deionized water and incubated with 10 molar excess (50 µM) of carbamate for 15 min before injection. An untreated sample was used as a control. The samples were injected 15 min, 9 h, and 18 after the compound was added – same results were obtained regardless of incubation time.

For compounds **12** and **15**, LC-MS analysis was carried out on an Agilent 1290 Infinity UHPLC system coupled to an iFunnel Q-TOF 6550 HRMS, developing a bioZen™ 3.6 µm Intact XB-C8 LC column 100 × 2.1 mm (Phenomenex) with a water/0.1% formic acid – acetonitrile/0.1% formic acid linear gradient as described previously (the protein eluted at 7.2 min under the experimental conditions).[110] For all other compounds, 2 µL of a 4°C-thermostated sample was injected on a bioZen™ Intact XB-C8 LC column (100 × 2.1 mm, 3.6 µm, Phenomenex) mounted on an Elute UHPLC system (Bruker) coupled to a QTof Impact II HRMS (Bruker). Chromatographic separation followed a linear gradient of water/0.1% formic acid (solvent A) – acetonitrile/0.1% formic acid (solvent B) at 0.4 ml/min under the following conditions: 95% A + 5% B from 0 to 1 min, then a linear gradient from 5% to 100% B from 1 to 18 min, then a plateau at 100% B up to 23 min. A reconditioning step to the initial conditions (95% A + 5% B) was applied from 23 to 24 min and maintained up to 26 min. The main MS parameters were: mass range from 300 to 3000 m/z; spectra rate of 1 Hz; ionization source: VIP-HESI; End Plate Offset: 500 V; Dry Gas: 10.0 L/min; Probe Gas Temp: 350 °C; Exhaust: On; Capillary: 4,500 V; Nebulizer: 3.0 bar; Dry Temp: 260 °C; Probe Gas: 5.0 L/min.

Collected data were converted to mzXML using msconvert from the ProteoWizard toolkit,[120] spectra of interest extracted using OpenChrom® software[121] (v 1.5.13) and deconvoluted with the UniDec software (v 6.0),[122] resulting in one major species for each sample.

##### X-ray structural study

Crystals of hBChE were obtained by vapor diffusion using 0.1 M MES pH 6.5, 2.15 M $(NH_4)SO_4$ as crystallisation buffer, and C-terminus truncated recombinant protein produced in Chinese hamster ovary cells[123] purified by both affinity and size exclusion chromatographies as previously reported.[124] Protein-ligand complexes were obtained after crystal soaking (5 min in case of 8QTW, 15 h in case of 8QTX) with crystallisation buffer containing 2 mM of **28**, where the 0.1 M initial stock compound solution was prepared in 100% MeOH. The crystals were then cryo-protected in the soaking solution containing 20% glycerol before flash cooling into liquid nitrogen. X-ray diffraction data were collected at the European Synchrotron Radiation Facility (ESRF, Grenoble, France) on the ID23-1 beamline at 100 K. Images were automatically processed using the AutoPROC[125] or EDNA[126] pipelines, for the carbamoylated or the hydrolysed and decarboxylated structures, respectively. Indexed and scaled data were analysed with the Phenix software suite.[127] The initial model was obtained by molecular replacement using the hBChE structure (PDB 1P0I) devoid of any ligand, glycan, and water molecules. In each structure, unambiguous extra electron density allowed to fit the different ligands whose restraints were calculated with eLBOW[128] and the AM1 semi-empirical quantum mechanical method. For the carbamoylated structure, additional specific restraints for bond formation, angles, and dihedrals were applied to the Ser198-*O*γ atom. Final structures were obtained after iterative cycles of model building in *Coot* software[129] and refinement (Table S3). Structures of **28**-carbamoylated hBChE or in complex with hydrolysed and decarboxylated **28** were deposited into the Protein Data Bank under accession numbers 8QTW and 8QTX, respectively.

**Computational studies**

Computational experiments were performed on workstations at the Department of Pharmaceutical Chemistry, Faculty of Pharmacy, and facilities of the Ažman Computing Centre at National Institute of Chemistry in Ljubljana, using Schrödinger Small Molecule Discovery Suite Release 2021-1 (Schrödinger, LLC, New York, USA, 2021) and Desmond/Maestro Non-commercial Distribution (Desmond v6.5, D. E. Shaw Research, New York, NY, USA, 2021).[130] The 8QTX crystal structure was prepared using Protein Preparation Wizard[131] using the following protocol: bond orders were assigned using the CCD database, missing hydrogens were added, disulfide bonds were created, termini were capped, missing side chains and loops were modelled with Prime,[132] and het protonation states (pH 7.0 ± 2.0) were modelled with Epik.[133] All hets except the co-crystallised ligands, co-solvent molecules, and fusion partners were removed. The comparison with the available hBChE crystal structures revealed that crystal waters are highly conserved,[134] especially waters 2 and 3 located behind the catalytic triad, which form hydrogen bonds with Gly439 and Ser224 residues, respectively, possess low B-factors, and were later found to be essentially stationary during molecular dynamics (MD) simulations. The crystal waters were therefore retained, except waters 1 and 116 (to avoid clash with the ligand atoms during docking).

Hydrogen bonds and protonation of residues were automatically assigned and optimized using PROPKA[135] (pH 7.0). **NOTE**: the protonation state of the charged residues in the vicinity of the catalytic centre significantly influences the reaction mechanism, as was shown both by mutation experiments and computationally for the hAChE's Glu202 residue. The protonated Glu202 helps maintaining a hydrogen bond network in the catalytic site, while the deprotonated Glu202 disrupts the catalytic His447 residue.[136–142] The same seems to also apply for the corresponding Glu197 residue in hBChE. When Glu197 was modelled as deprotonated by Suárez and Field,[143] they noticed that during their relatively short MD simulation, a $Na^+$ ion quickly moved into its vicinity and that the catalytic Ser198 interacted preferentially with the backbone carbonyl of Ser224, and not His438 of the catalytic triad, as it should. A constant pH MD study[144] indicated that Glu197 (its PROPKA-predicted[135] p$K_a$ value is 10.1) are predominantly in its protonated state, while the catalytic Glu325 (PROPKA p$K_a$ 4.1) and nearby Glu441 residues (PROPKA p$K_a$ 7.0) were predicted to remain charged. The authors observed that protonated Glu197 in hBChE stabilised a highly conserved water molecule, preserved the hydrogen bond network and catalytic triad integrity, and these structural characteristics were essential for a meaningful MD simulation of bound ligands.[144] A recent computational study by Zlobin et al. has also shown that Glu197 and Glu441 can share a proton through a Grotthuss mechanism mediated by the water molecule.[145] Since we have obtained similar results in our 20 ns MD simulations where we modelled Glu197 and Glu441 in 8QTX crystal structure either as neutral or charged (Fig. S13). Namely, the essential hydrogen bond between $O^{\gamma}$ of Ser198 and $N^{\varepsilon}$ of His438 was persistent only in case of neutral Glu197 and charged Glu441. Therefore, we applied this protonation pattern in all subsequent simulations.

*Non-covalent Molecular Docking*

For non-covalent docking calculations, we used the prepared hBChE crystal structure as described above. Additionally, all water molecules and ions were removed, and Ser198 was mutated to Ala198 to remove a potential steric clash. The receptor grid was then generated using Receptor Grid Generation with default settings – van der Waals radii scaling by 1, partial charge cutoff was set to 0.25, and OPLS4 forcefield[146] was used, while the co-crystallised ligand defined the centroid and size of the active site with innerbox of 10 Å$^3$ and outerbox of 24.08 Å$^3$. Additionally, amide nitrogens of Gly116, Gly117, and Ala199 were defined as hydrogen donors. Ligand structures were prepared with LigPrep and ionized with Epik[133] (pH 7 ± 2) using OPLS4 force field.[146] Due to limitations of the forcefield, ligands containing Se and Te atoms had to be excluded. Docking was performed using Glide XP[147] coupled with OPLS4 forcefield[146] with the following modifications of the default settings: (1) a constraint of at least one hydrogen bond formed with the oxyanion hole residues as defined in the grid with a (thio)carbonyl acceptor, and (2) a core constraint of ≤ 3 Å RMSD *vs*. the cognate ligand position (defined by SMARTS pattern c1cccc(c12)[n]cc2CN(C)CCC3CCCCCC3) had to be satisfied. The output poses (2 per ligand) were visualised with Maestro and ranked by Glide composite score, number of hydrogen bonds formed with the oxyanion hole, and overall similarity with the cognate pose. One pose per ligand, merged with the prepared crystal structure from above, was retained for further analysis and calculations.

*Covalent Molecular Docking*

The CovDock protocol[148] was used for covalent docking of carbamoylazoles. The prepared hBChE crystal structure with retained crystal waters was used. His438 residue was manually modified to the charged HIP438 to ensure a proper post-reaction structure, and Ser198 was negatively ionized. Ser198 was selected as the reactive residue, the box was defined around the centroid of the co-crystallised ligand with the size defined by the ligand (innerbox of 10.0 Å$^3$ and outerbox of 24.08 Å$^3$). Reaction type was customized using a custom chemistry file (see below) to correctly produce a negatively charged tetrahedral intermediate, since the available “nucleophilic addition to a double bond” led to a neutral one. No constraints were imposed and the thorough pose prediction docking mode was used. The results were visualised in Maestro and ranked by Glide composite score, the obtained docking poses with the “ex-carbonyl” anionic oxygen located outside the oxyanion hole were removed, and the remaining poses ranked by calculated score and observed favourable interactions with the binding site residues.

*Custom chemistry file for CovDock covalent docking*

```
REACTION_NAME          Custom
CUSTOM_CHEMISTRY         ('<1>', ('charge', 0, 1))
CUSTOM_CHEMISTRY         ('<1>|<2>', ('bond', 1, (1, 2)))
CUSTOM_CHEMISTRY         ('<2>[O,S]', ('charge', -1, 2))
```

LIGAND_SMARTS_PATTERN 1,[C,c]=[O,S]
RECEPTOR_SMARTS_PATTERN 2,[CH2X4]-[O]

*Molecular Dynamics simulations*

The BChE poses generated in docking experiments were transferred back to the crystal structure with crystal waters present (the pre-grid generation crystal structure, same alignment). The systems for molecular dynamics (MD) simulations were prepared with System Builder: TIP4P[149] water molecules were added up to 10 Å from the protein surface to solvate the protein in a orthorhombic box, $Na^+$ and $Cl^-$ ions were added to neutralise the system and produce the final 0.15 M concentration, and OPLS_2005 force field[150,151] was used for parametrization of the macromolecule as well as the ligand. The default Desmond relaxation protocol (desmond_npt_relax.msj) was used for the equilibration stage: (1) 100 ps of Brownian Dynamics NVT, 10 K, small timesteps, with restraints on the solute heavy atoms, (2) 12 ps NVT, 10 K, with small timesteps and restraints on the solute heavy atoms, (3) 12 ps NPT, 10 K, and restraints on the solute heavy atoms, (4) 24 ps unrestrained NPT; followed by the production stage. The following setup was used for the MD production stage: 1.2 ps interval for energy, RESPA integrator with 2 fs time step, cutoff scheme at 9.0 Å, random seed, isothermal-isobaric NPT ensemble at 300 K and 1.013 bar pressure with Nose-Hoover chain thermostat and Martyna-Tobias-Klein barostat (1 and 2 ps relaxation time, respectively, isotropic coupling). The simulation times were 20–1000 ns with 1000 frames per trajectory. The simulation results were analysed using the built-in Desmond tools, focusing on three crucial aspects: (1) the binding and concomitant activation of the carbonyl group in the oxyanion hole via the number of hydrogen bonds formed with the oxyanion hole residues Gly116, Gly117, and Ala199, (2) propensity for covalent bond formation via the distance of catalytic residue Ser198 $O^{\gamma}$ and ligand's carbonyl carbon, and (3) the stability of the catalytic triad, evaluated as the distance between Ser198 $O^{\gamma}$ and His438 $N^{\varepsilon}$.

*Quantum Mechanical Calculations*

The quantum mechanical (QM) density functional theory (DFT) calculations for the model reaction of fragments containing warheads with methoxide were carried out in Jaguar.[152] After initial rough MacroModel minimization, the structures were geometrically optimized at M06-2X/6-31G++** level of theory and solvent (water) effect were modelled by self-consistent reaction field method using the Poisson-Boltzmann (PBF) solver. The structures were checked for the absence of imaginary frequencies to confirm that they correspond to geometric minima. The ultrafine pseudospectral grids were used for geometry optimization, followed by full analytic integrals for a single point frequency calculation at M06-2X/aug-cc-pVTZ(-f) level with PBF solver in water. For anionic intermediates, the stop_rxn=3 and geoconv_mode=standard keywords were applied to successfully model the unstable intermediates. Transition states were located utilizing the QST (quadratic synchronous transit) method and further verified by the presence of a single imaginary frequency oscillating in the direction of the assumed reaction coordinate, as well as by performing an intrinsic reaction coordinate scan (IRC) in direction of both reactants and products. The calculated total energies for reactants, intermediates, and products were compared to calculate the reaction energetics.

For the leaving group p$K_a$ values calculation in water, Jaguar's p$K_a$ prediction module[153] was used.

Several quantum-chemical descriptors were also calculated from the optimized structures of **6a**–**31a**, such as atomic charges (Mulliken, Stockholder, ESP, natural charge from NBO) and α polarizabilities for the carbonyl carbon atom, vibrational wavenumbers and Fukui f_NN LUMO index for the carbonyl group, HOMO and LUMO energies, and their derived descriptors (chemical hardness, electrophilicity index, electronic chemical potential, local electrophilicity index).[154–159]

The inhibitors **6a–31a** were separated into two groups: covalent inhibitors (14 cases) and non-covalent inhibitors (12 cases). The quantum-chemical and experimentally-derived descriptors were compared between both groups using two-sided Mann-Whitney U-tests. Additionally, the correlation between individual descriptors and $pIC_{50}$ was computed using non-parametric Spearman correlation. Two-tailed p values were calculated and considered significant at $p < 0.05$ (*), <0.01 (**), <0.001 (***), or <0.0001 (****). Furthermore, the eight descriptors that were found to significantly differ in both Mann-Whitney and correlation tests were analysed using principal component regression vs. $pIC_{50}$: data were standardised and principal components were selected based on parallel analysis (percentile level 95%, number of simulations: 1000, auto random seed). GraphPad Prism 10.4.1 (GraphPad Software, Boston, MA, USA) was used for statistical analysis.

*Multiscale QM/MM calculations*

The pre-reaction docking poses for the **26** and **28** ligands were merged with the hBChE protein, the system was solvated and ions were added using the Desmond System Builder, then everything beyond 4 Å of the protein atoms was deleted to reduce system size. The QM region, MM region, and pseudobond atoms were defined as shown in Fig. 3. The initial structures were optimized by pseudo-bond QM/MM calculations with an iterative energy minimization procedure. The QM part of the calculations employed the Beck 3-parameter hybrid functional combined with the Lee-Yang-Parr correlation functional (B3LYP)[31] and 6-31G+* basis set, using a modified version of the Gaussian03 program, while the MM calculations were performed with a modified version of the AMBER8 program. For the QM subsystem, the convergence criteria for geometry optimizations used the original Gaussian03 default settings, while the MM geometry convergence criterion was energy gradient RMSD below 0.1 kcal $mol^{-1}$ $Å^{-1}$. The SCF convergence criteria were set to "tight", corresponding to a maximum density change of $\leq 1.0 \times 10^{-6}$ and an RMS density change of $\leq 1.0 \times 10^{-8}$. The boundary carbon atoms were treated with improved pseudo-bond parameters. No cutoff for non-bonded interactions was used, and only atoms within 20 Å of the ligand's carbonyl carbon were allowed to move while the other atoms were frozen during QM/MM calculations. With a reaction coordinate driving method and an iterative energy minimization procedure[160], the enzyme reaction path was determined by the pseudo-bond QM/MM calculations at the same level of theory. For the carbamoylation reaction, the reaction coordinate is set as $d(C_{INH}–N_{INH}) – d(H^{γ}–N^{ε}) – d(C_{INH}–O^{γ})$ for the nucleophilic attack of Ser198 on the ligand's carbonyl $C_{INH}$ carbon and as $d(N^{ε}–H^{γ})$

– d($N_{\text{INH}}$–$H^{\gamma}$) for the proton transfer from His438 to the leaving group. For the decarbamoylation reaction, the reaction coordinate is set as as – d($H^{w}$–$N^{\varepsilon}$) – d($C_{INH}$–$O^{w}$) and as d($C_{INH}$–$O^{\gamma}$) – d($O^{\gamma}$–$H^{w}$) for the proton transfer from His438 to Ser198 (Fig. 3). Normal mode frequency analyses were performed to characterize the reactants, intermediates, transition states, final products, and to obtain the thermodynamic parameters of the complexes. In addition, single-point QM/MM energy calculations were carried out at B3LYP-D3/6-311G++**:AMBER level on the optimized geometries to obtain more accurate energies.

**Aqueous stability**

UPLC analyses were performed as described under Chemistry – General Information. The method used a Waters Acquity UPLC® HSS C18 SB column (2.1 × 50 mm, 1.8 µm) thermostated at 40 °C, with: injection volume, 1 µL; flow rate, 0.4 mL/min; detector λ, 254 nm; mobile phase A: 0.1% TFA (v/v) in water; mobile phase B: MeCN. Gradient: 0–5 min, 20–100% B; 5–6 min, 20% B.

Compound stock solutions (10 mM) were prepared in absolute EtOH, while internal standard (caffeine) was dissolved in PBS (0.1 mg/mL). The aqueous stability solutions were prepared by mixing 800 µL PBS with internal standard, 175 µL absolute EtOH, and 25 µL compound stock solution in a HPLC vial to give 250 µM compound concentration in PBS/EtOH = 80:20, which placed in an autosampler thermostatted at 20 °C, injected immediately and then at hourly intervals for at least 24 hours. The analyte AUCs were normalised on internal standard AUCs and compared with the timepoint zero. For the rate calculation, pseudo-first order kinetics were presumed (*Table S10*). The stability data show that some of these carbamates are stable on the order of hours for the most reactive compounds, while some are stable for weeks in the aqueous solution under physiologically similar conditions. For the two highly reactive compounds **6** and **28**, the addition of GSH at cellular concentrations significantly accelerated the decay.

**Synthetic Chemistry**

*General Information*

The reagents and solvents were used as received from commercial suppliers. Tetrahydrofuran (THF) was distilled from sodium/benzophenone and stored under Ar over 4 Å molecular sieves prior to use. After extraction organic phases were dried over anhydrous sodium sulfate. Reactions were monitored using analytical thin-layer chromatography (TLC) on silica gel 60 $F_{254}$ Al plates. Developed plates were inspected under UV light and, if necessary, visualised with ninhydrin, vanillin/sulfuric acid, Dragendorff's or potassium permanganate stains. Melting points were determined with Büchi 535 Melting Point Apparatus (uncorrected). Nuclear magnetic resonance spectra were recorded with a Bruker Avance III 400 MHz spectrometer at 400 MHz for $^{1}$H, 100 MHz for $^{13}$C, and 376 MHz for $^{19}$F nucleus, respectively, using DMSO-$d_6$, $CDCl_3$, acetone-$d_6$, or MeOH-$d_4$ with TMS as the internal standard, as solvents. Chemical shifts are reported in *parts per million* (ppm), $^{1}$H spectra are calibrated to the TMS peak (0 ppm) or the residual non-perdeuterated solvent peak, $^{13}$C spectra to the central peak of the deuterated solvent, while the $^{19}$F and $^{31}$P spectra were not calibrated. $^{77}$Se (at 95 MHz) and $^{125}$Te NMR (at 158 MHz) spectra were recorded with a Bruker Avance III 500 MHz spectrometer equipped with a 5 mm BBO probe using a 0.5 M solutions of diphenyl diselenide (PhSeSePh) in $CDCl_3$ ($\delta_{Se}$ = 463 ppm)[161] and 0.9 M solutions of diphenyl ditelluride (PhTeTePh) in $CDCl_3$ ($\delta_{Te}$ = 420 ppm)[162], respectively, in a sealed tubes as external standards at 23 °C. The $^{77}$Se and $^{125}$Te chemical shifts were extracted from $^{1}$H–$^{77}$Se *gs*-HMBC and $^{1}$H–$^{125}$Te *gs*-HMBC spectra, respectively. The experiments were optimized for a long-range coupling of 2 Hz. The number of increments for evolution was 512. The number of scans per increment was 4 with 2 s delay between transients. Gradient ratios were 70 : 30 : 59.0715% for ($^{1}$H–$^{77}$Se *gs*-HMBC) and 70 : 30 : 71.5498% (for $^{1}$H–$^{125}$Te *gs*-HMBC). The spectra were recorded with a spectral width of 400 ppm, and to avoid any misinterpretation of potential aliased or folded peaks, each compound was recorded in two different spectral windows. The multiplicities are reported as follows: *s* (singlet), *d* (doublet), *t* (triplet), *q* (quartet), *m* (multiplet), *dd* (doublet of doublets), *ddd* (doublet doublet of doublets), *td* (triplet of doublets), *qd* (quartet of doublets), and *br* (broad), number of equivalent nuclei (by integration), coupling constants (*J*) quoted in Hertz (Hz). Mass spectra were recorded on Agilent 6224 Accurate Mass TOF LC/MS and Thermo Scientific Q Exactive™ Plus LC-MS/MS spectrometers and IR spectra on Thermo Nicolet FT-IR spectrophotometer. Column chromatography was performed on silica gel (Silica gel 60, particle size: 0.035–0.070 mm, Merck). UPLC analyses were performed on Thermo Scientific Dionex UltiMate 3000 modular system (Thermo Fisher Scientific Inc.). The general method (method I) used a Waters Acquity UPLC® HSS C18 SB column (2.1 × 50 mm, 1.8 µm) thermostated at 40 °C, with: injection volume, 1–10 µL; sample, 0.1–0.2 mg/mL in MeOH; flow rate, 0.4 mL/min; detector λ, 220 and 254 nm; mobile phase A: 0.1% TFA (v/v) in water; mobile phase B: MeCN. Gradient: 0–2 min, 20% B; 2–5 min, 20%–90% B; 5–8 min, 90% B. Method II had the same conditions except gradient: 0–5 min, 20–100% B; 5–6 min, 20% B. Method III had the same conditions except gradient: 0–2 min, 0% B; 2–8 min, 0–100% B. The purities of the tested compounds were established to be ≥95%, as determined by UPLC, unless indicated otherwise.

***General procedure 1 (GP1) – carbamoylation***:

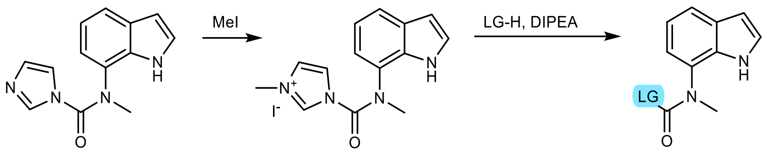

*N*-(1*H*-Indol-7-yl)-*N*-methyl-1*H*-imidazole-1-carboxamide[163] **26a** (2403 mg, 10 mmol) and methyl iodide (1.5 eq., 930 µL) in 15 mL THF were stirred in a pressure tube at 50 °C under Ar for 48 h. The solvent was then removed *in vacuo* and the residue containing 1-((1*H*-indol-7-yl)(methyl)carbamoyl)-3-methyl-1*H*-imidazol-3-ium iodide reconstituted in dry MeCN to 25.0 mL in a volumetric flask. An aliquot of 1.0 mmol was then transferred to a reaction flask, DIPEA (500 µL) and acidic leaving group (heterocycle, alcohol, thiol, etc., 1.5 eq.) were added, and stirred under Ar at 75 °C for 3 h. The volatile components were then removed *in vacuo* and the residue purified by column chromatography on silica.

***General procedure 2 (GP2) – Mannich reaction with 2-cycloheptyl-N-methylethan-1-amine***:

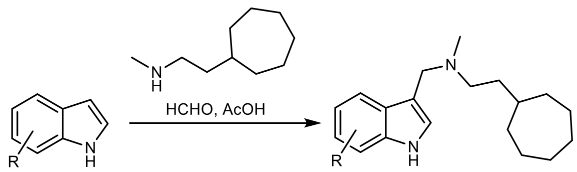

Parent indole (1.0 mmol) was dissolved in EtOAc (1 mL), glacial AcOH (1 mL), formaldehyde (1.2 eq., 37%, 89 µL), and 2-cycloheptyl-*N*-methylethan-1-amine (171 mg, 1.1 eq.) were added, and the reaction mixture was stirred at rt for 18 h. The volatile components were removed *in vacuo* and the product was purified following *GP3*.

***General procedure 3 (GP3) – RP-CC purification***: Compounds were purified by reversed-phase column chromatography (RP-CC) (Isolera Biotage One Flash Chromatography system, Biotage® Sfär C18 Duo 100 Å 30 µm column, 30 g) using a gradient of 0.1% TFA in deionized water and MeCN as eluent (gradient 0–100% MeCN in 6 column volumes (300 mL); 100% MeCN for 2 column volumes (100 mL). After the RP-CC, fractions containing the product were combined and organic volatiles were evaporated *in vacuo*. The remaining aqueous solution was made alkaline (pH 10) with 1 M $NaOH_{(aq)}$ and extracted with DCM (2 × 30 mL). The combined organic phase was dried over anhydrous sodium sulfate, filtered, and volatile components evaporated *in vacuo* to afford pure product.

***General procedure 4 (GP4) – reduction with lithium aluminium hydride***: To a solution of the starting compound (amide, nitrile, ester) (1.0 mmol) in anhydrous THF (1 mL), lithium aluminium hydride solution (2.4 M in THF, 3.0 eq. (1.25 mL) for amides, 1.5 eq. (0.63 mL) for nitriles, 1.0 eq. (0.42 mL) for esters and ketones, 1.0 eq. (0.42 mL) for azides or, if multiple reducible groups were present, total volume was the sum of volumes for individual group) was added slowly with stirring under argon. The reaction mixture was stirred at 50 °C for 12 h and then while being cooled on an icebath, cautiously quenched with brine. The resulting grey solids were suspended in $Et_2O$ (20 mL), sonicated in an ultrasonic cleaning bath for 5 minutes, filtered, and washed with $Et_2O$ (2 × 10 mL). The ethereal extracts were dried over sodium sulfate, filtered, and volatile components evaporated *in vacuo* to afford the product (amine or alcohol).

***General procedure 5 (GP5) – click reaction***: Ascorbic acid (881 mg, 5 mmol) and sodium bicarbonate (336 mg, 4 mmol) in 5 mL water were stirred until effervescence subsided. Azide (1 mmol) and alkyne (1 mmol) in DCM (10 mL) were then added, and in the cases with biotin azide, acetone (30 mL) was also added to improve solubility. Copper sulfate pentahydrate (125 mg, 0.5 mmol) was then added, and the mixture vigorously stirred at rt for 48 h or until complete conversion by LC-MS. Solvent was removed *in vacuo*, and the residue mixed with 1 M $HCl_{(aq)}$ (15 mL) – for B probes, or $NaHCO_{3(aq.,\ sat.)}$ (15 mL) – for P probes, and then extracted with DCM (2 × 20 mL). The organic phases were separated, dried with anhydrous sodium sulfate, the solvent removed *in vacuo*, and the product purified by *GP3*.

***General procedure 6 (GP6) – preparation of N-carbamoylimidazoles***: Secondary amine (1.0 mmol) was dissolved in DCM (5 mL), CDI (1.1 eq., 178 mg) was added, and the reaction mixture stirred at rt for 12 h. The organic phase was extracted with water (2 × 10 mL), dried with sodium sulfate, and solvent removed *in vacuo* to afford *N*-carbamoylimidazole, which was sufficiently pure to be used further.

*Synthesis*

*S-(3-(((2-Cycloheptylethyl)(methyl)amino)methyl)-1H-indol-7-yl) dimethylcarbamothioate* (**2**)

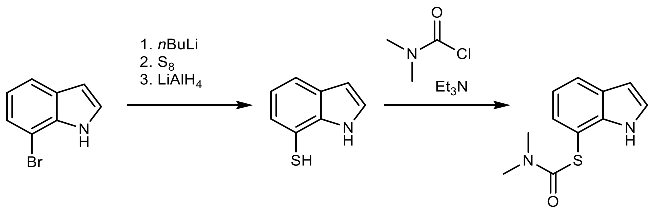

7-Bromo-1*H*-indole (1462 mg, 7.46 mmol) in THF (15 mL) was cooled to –80 °C under Ar, then *n*-butyllithium (2.1 eq., 6.8 mL, 2.3 M in cyclohexane/hexane) was added dropwise, the mixture stirred for 30 min at –80 °C and for 4 h at 0 °C, then elemental sulfur (1.1 eq., 263 mg) was added. After the sulfur dissolved, 2 mL of 2.4 M $LiAlH_4$ in THF were added, the mixture stirred for 1 h at rt, then 10 % citric $acid_{(aq)}$ (20 mL) and 1 M $HCl_{(aq)}$ (10 mL) were added, the solvent removed *in vacuo*, and the residue extracted with $Et_2O$ (30 mL). The organic phase was shaken with 2 M $NaOH_{(aq)}$ (20 mL), the aqueous phase separated, acidified with 6 M $HCl_{(aq)}$ to pH 2, and extracted with $Et_2O$ (30 mL). The organic phase was dried with sodium sulfate, the solvent removed *in vacuo*, and 1*H*-indole-7-thiol isolated by column chromatography on silica (PE/DCM = 5:1). Yield: 213 mg (1.43 mmol, 19.1%) of beige semisolid, which was immediately used further. ESI-HRMS: *m/z* = 148.0217 ($MH^+$); $C_8H_6NS$ requires: *m/z* = 148.0226 ($MH^+$). $\nu_{max}$ 3411, 2964, 2930, 2867, 1540, 1428, 1333, 1285, 1192, 1095, 785 $cm^{-1}$. $^1H$ NMR (400 MHz, $CDCl_3$) δ = 3.24 – 3.26 (m, 1H), 6.58 (dd, *J*=2.1 Hz, 3.2 Hz, 1H), 7.04 (t, *J*=7.6 Hz, 1H), 7.21 – 7.23 (m, 1H), 7.27 – 7.30 (m, 1H), 7.56 – 7.60 (m, 1H), 8.37 (br s, 1H).

$^{13}$C NMR (101 MHz, $CDCl_3$) δ = 103.71, 109.10, 120.50, 120.81, 124.61, 127.61, 127.99, 137.49.

1*H*-Indole-7-thiol (126 mg, 0.844 mmol), *N*,*N*-dimethylcarbamoyl chloride (2.0 eq., 155 µL), and triethylamine (2.0 eq., 235 µL) in DCM (5 mL) were stirred at rt for 18 h. *S*-(1*H*-Indol-7-yl) dimethylcarbamothioate **2a** was isolated by column chromatography on silica (1. PE/EtOAc = 5:1; 2. PE/EtOAc = 2:1; 3. PE/EtOAc = 1:1). Yield: 92 mg (0.418 mmol, 49.5%) of beige semisolid. ESI-HRMS: *m/z* = 221.0740 ($MH^+$); $C_{11}H_{13}N_2OS$ requires: *m/z* = 221.0743 ($MH^+$). $\nu_{max}$ 3253, 3100, 2920, 1645, 1434, 1361, 1332, 1254, 1089, 882, 786, 720, 693, 651 $cm^{-1}$. Purity: UPLC (method I, 220 nm): $t_r$ = 4.093 min, 99.4% total area. $^1$H NMR (400 MHz, $CDCl_3$) δ = 3.03 (s, 3H), 3.17 (s, 3H), 6.59 (dd, *J*=2.1 Hz, 3.2 Hz, 1H), 7.13 – 7.18 (m, 2H), 7.36 (dd, *J*=0.8 Hz, 7.4 Hz, 1H), 7.72 – 7.76 (m, 1H), 8.60 (br s, 1H). $^{13}$C NMR (101 MHz, $CDCl_3$) δ = 37.04, 103.32, 109.86, 120.16, 122.91, 124.73, 128.65, 129.86, 138.37, 166.22.

*S*-(3-(((2-Cycloheptylethyl)(methyl)amino)methyl)-1*H*-indol-7-yl) dimethylcarbamothioate **2** was prepared from *S*-(1*H*-indol-7-yl) dimethylcarbamothioate **2a** (80 mg, 0.363 mmol) following *GP2*. Yield: 83 mg (0.214 mmol, 60.0%) of beige solid. mp 120.8–122.9 °C. ESI-HRMS: *m/z* = 388.2411 ($MH^+$); $C_{22}H_{34}N_3OS$ requires: *m/z* = 388.2417 ($MH^+$). $\nu_{max}$ 2918, 2851, 1657, 1588, 1444, 1364, 1343, 1251, 1098, 912, 744, 691, 652 $cm^{-1}$. Purity: UPLC (method I, 254 nm): $t_r$ = 4.557 min, 95.5% total area. $^1$H NMR (400 MHz, $CDCl_3$) δ = 1.13 – 1.22 (m, 2H), 1.35 – 1.72 (m, 13H), 2.20 (s, 3H), 2.40 – 2.44 (m, 2H), 3.02 and 3.18 (2 × s, 6H), 3.67 (s, 2H), 7.10 – 7.12 (m, 1H), 7.13 (d, *J*=7.7 Hz, 1H), 7.31 (dd, *J*=0.9 Hz, 7.3 Hz, 1H), 7.78 – 7.81 (m, 1H), 8.40 (s, 1H). $^{13}$C NMR (101 MHz, $CDCl_3$) δ = 26.60, 28.63, 34.94, 35.96, 37.10, 37.26, 37.71, 42.44, 53.09, 55.92, 109.74, 114.52, 119.97, 121.93, 123.87, 128.82, 129.94, 138.89, 166.29.

*Se-(3-(((2-Cycloheptylethyl)(methyl)amino)methyl)-1H-indol-7-yl) dimethylcarbamoselenoate* (**3**)

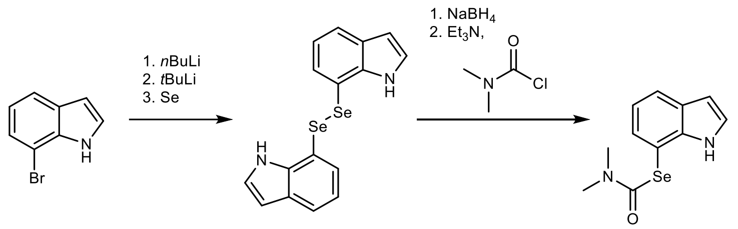

7-Bromo-1*H*-indole (604 mg, 3.08 mmol) was dissolved in dry THF (10 mL) under Ar, cooled to –80 °C, *n*-butyllithium (1.4 mL, 1.05 eq., 2.3 M in cyclohexane/hexane) was added dropwise, the solution stirred for 30 min at –80 °C, then *tert*-butyllithium (3.8 mL, 2.1 eq., 1.7 M in pentane) was added dropwise, the reaction mixture was stirred for 60 min at –40 °C before being cannulated to another flask containing elemental black selenium powder (2.0 eq., 486 mg). The reaction mixture was left to slowly warm to rt and stirred for 4 h. 10% Citric $acid_{(aq)}$ (10 mL) was then added and the reaction mixture stirred unstoppered in air overnight. The solvent was then removed *in vacuo*, 1,2-di(1*H*-indol-7-yl)diselenide extracted into $Et_2O$ (15 mL), and isolated by column chromatography on silica (1. PE/DCM = 5:1; 2. PE/DCM = 1:1).

Yield: 201 mg (0.515 mmol, 16.7%) of yellow-orangish solid. mp 116.1–119.0 °C. $^1$H NMR (400 MHz, $CDCl_3$) δ = 6.48 (dd, *J*=2.1 Hz, 3.2 Hz, 2H), 6.78 (t, *J*=2.9 Hz, 2H), 6.93 (t, *J*=7.7 Hz, 2H), 7.30 (dd, *J*=0.8 Hz, 7.3 Hz, 2H), 7.57 – 7.60 (m, 2H), 7.80 (br s, 2H). $^{13}$C NMR (101 MHz, $CDCl_3$) δ = 103.31, 112.71, 120.34, 123.00, 124.71, 127.91, 130.12 (t, $J_{C–Se}$=5.2 Hz), 137.85. $^{77}$Se NMR (95 MHz, $CDCl_3$) δ = 429.

To 1,2-di(1*H*-indol-7-yl)diselenide (122 mg, 0.313 mmol) in THF (5 mL) and absolute EtOH (2 mL) was added sodium borohydride in 20 mg portions until the orange solution discoloured. *N*,*N*-Dimethylcarbamoylchloride (10 eq., 288 µL) and triethylamine (500 µL) were added, the solution stirred at rt for 1 h, then the solvent was removed *in vacuo*, and *Se*-(1*H*-indol-7-yl) dimethylcarbamoselenoate **3a** isolated by column chromatography on silica (1. PE/EtOAc = 3:1; 3. PE/EtOAc = 1:1). Yield: 108 mg (0.404 mmol, 64.6%) of beige solid. mp 102.8–104.3 °C. ESI-HRMS: *m/z* = 269.0181 ($MH^+$); $C_{11}H_{13}N_2OSe$ requires: *m/z* = 269.0188 ($MH^+$). $\nu_{max}$ 3265, 3056, 2928, 1653, 1361, 1332, 1256, 1113, 1091, 782, 723, 679 $cm^{-1}$. Purity: UPLC (method I, 220 nm): $t_r$ = 4.180 min, 98.4% total area. $^1$H NMR (400 MHz, $CDCl_3$) δ = 3.02 (s, 3H), 3.11 (s, 3H), 6.62 (dd, *J*=2.1 Hz, 3.2 Hz, 1H), 7.12 (dd, *J*=7.4 Hz, 7.8 Hz, 1H), 7.18 – 7.20 (m, 1H), 7.44 (dd, *J*=0.8 Hz, 7.3 Hz, 1H), 7.71 – 7.75 (m, 1H), 8.58 (s, 1H). $^{13}$C NMR (101 MHz, $CDCl_3$) δ = 36.94, 37.73, 103.46, 108.13, 120.32, 122.68, 124.58, 128.32, 130.69 (t, $J_{C–Se}$=6.6 Hz), 138.68, 163.75. $^{77}$Se NMR (95 MHz, $CDCl_3$) δ = 429.

*Se*-(3-(((2-Cycloheptylethyl)(methyl)amino)methyl)-1*H*-indol-7-yl) dimethylcarbamoselenoate **3** was prepared from **3a** (58 mg, 0.22 mmol) following *GP2*. Yield: 56 mg (0.128 mmol, 58.4%) of beige solid. mp 84.9–87.3 °C. ESI-HRMS: *m/z* = 436.1849 ($MH^+$); $C_{22}H_{34}N_3OSe$ requires: *m/z* = 436.1862 ($MH^+$). $\nu_{max}$ 3325, 2920, 2852, 2792, 2758, 1642, 1460, 1360, 1255, 1082, 785, 739, 675, 597 $cm^{-1}$. Purity: UPLC (method I, 220 nm): $t_r$ = 4.597 min, 98.6% total area. $^1$H NMR (400 MHz, $CDCl_3$) δ = 1.15 – 1.24 (m, 2H), 1.36 – 1.73 (m, 13H), 2.22 (s, 3H), 2.41 – 2.46 (m, 2H), 3.00 (s, 3H), 3.10 (s, 3H), 3.68 (s, 2H), 7.08 – 7.13 (m, 2H), 7.39 – 7.43 (m, 1H), 7.79 (d, *J*=7.9 Hz, 1H), 8.48 (s, 1H). $^{13}$C NMR (101 MHz, $CDCl_3$) δ = 26.54, 28.58, 34.88, 35.91, 36.92, 37.64, 37.72, 42.38, 53.11, 55.87, 108.00, 114.40, 119.98, 121.56, 123.77, 128.46, 130.67 (t, $^3J$=6.3 Hz ($^{13}$C–$^{77}$Se)), 139.10, 163.79. $^{77}$Se NMR (95 MHz, $CDCl_3$) δ = 427.

*Te-(3-(((2-Cycloheptylethyl)(methyl)amino)methyl)-1H-indol-7-yl) dimethylcarbamotelluroate* (**4**)

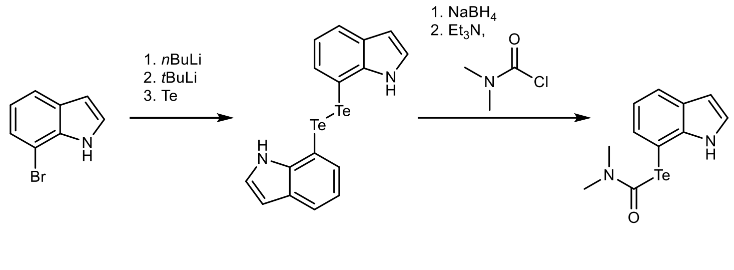

7-Bromo-1*H*-indole (800 mg, 4.08 mmol) was dissolved in dry THF (10 mL) under Ar, cooled to –80 °C, *n*-butyllithium (1.8 mL, 1.0 eq., 2.3 M in cyclohexane) was added dropwise, the solution stirred for 30 min at –80 °C, then *tert*-butyllithium (4.8 mL, 2.0 eq., 1.7 M in pentane)

was added dropwise, the reaction mixture was stirred for 60 min at –40 °C before being cannulated to another flask containing elemental tellurium (1.0 eq., 530 mg). The reaction mixture was left to slowly warm to rt and stirred for 6 h. 10% citric acid$_{(aq)}$ (10 mL) was then added and the reaction mixture stirred unstoppered in air overnight to complete the oxidation of tellurol to ditelluride. The solvent was then removed *in vacuo*, 1,2-di(1*H*-indol-7-yl)ditelluride extracted into $Et_2O$ (15 mL), and isolated by column chromatography on silica (1. PE/DCM = 10:1; 2. PE/DCM = 5:1). Yield: 726 mg (1.49 mmol, 73.0%) of dark orange solid. $\nu_{max}$ 3436, 3380, 3125, 3090, 3047, 1556, 1420, 1407, 1323, 1277, 1215, 1192, 1091, 1064, 780, 723 $cm^{-1}$. $^{1}$H NMR (400 MHz, $CDCl_3$) δ = 6.63 (dd, *J*=2.1 Hz, 3.2 Hz, 2H), 6.93 (t, *J*=2.9 Hz, 2H), 6.95 – 6.99 (m, 2H), 7.62 (dd, *J*=0.9 Hz, 7.2 Hz, 2H), 7.63 – 7.67 (m, 2H), 7.81 (s, 2H). $^{13}$C NMR (101 MHz, $CDCl_3$) δ = 89.42, 103.77, 120.74, 123.19, 124.52, 126.78, 135.22 (t, $J_{C–Te}$=9.6 Hz), 140.71. $^{125}$Te (158 MHz, $CDCl_3$) δ = 292.

1,2-Di(1*H*-indol-7-yl)ditelluride (400 mg, 0.821 mmol) was dissolved in THF (9 mL) and EtOH (1 mL), sodium borohydride was added in 10 mg portions with stirring at rt until the solution turned colourless, then *N*,*N*-dimethylcarbamoyl chloride (1 mL, 6.6 eq.) and triethylamine (1 mL) was added, and the reaction mixture stirred at rt for 1 h. The solvent was then removed *in vacuo*, 10% citric acid$_{(aq)}$ (10 mL) was added, *Te*-(1*H*-indol-7-yl) dimethylcarbamotelluroate **4a** extracted into $Et_2O$ (15 mL), and isolated by column chromatography on silica (1. PE/EtOAc = 5:1; 2. PE/EtOAc = 3:1). Yield: 355 mg (1.12 mmol, 68.2%) of orangeish semisolid. ESI-HRMS: *m/z* = 319.0092 ($MH^+$); $C_{11}H_{13}N_2OTe$ requires: *m/z* = 319.0085 ($MH^+$). $\nu_{max}$ 3270, 3048, 2920, 1639, 1599, 1355, 1328, 1254, 1112, 1084, 884, 780, 726, 673, 587 $cm^{-1}$. Purity: UPLC (method II, 220 nm): $t_r$ = 2.793 min, 97.3% total area. $^{1}$H NMR (400 MHz, $CDCl_3$) δ = 2.94 (s, 3H), 3.00 (s, 3H), 6.68 (dd, *J*=2.1 Hz, 3.2 Hz, 1H), 7.08 (dd, *J*=7.3 Hz, 7.8 Hz, 1H), 7.19 – 7.22 (m, 1H), 7.67 (dd, *J*=0.8 Hz, 7.2 Hz, 1H), 7.76 (d, *J*=7.9 Hz, 1H), 8.78 (s, 1H). $^{13}$C NMR (101 MHz, $CDCl_3$) δ = 35.95, 38.61, 95.79, 103.55, 120.61, 122.79, 124.45, 127.39, 134.48 (t, $^{3}J$=16.2 Hz ($^{13}$C–$^{125}$Te)), 140.32, 156.94. $^{125}$Te (158 MHz, $CDCl_3$) δ = 597.

*Te*-(3-(((2-Cycloheptylethyl)(methyl)amino)methyl)-1*H*-indol-7-yl) dimethylcarbamotelluroate **4** was prepared following *GP2* from **4a** (171 mg, 0.543 mmol). Yield: 118 mg (0.244 mmol, 45.0%) of yellowish semisolid. ESI-HRMS: *m/z* = 486.1772 ($MH^+$); $C_{22}H_{34}N_3OTe$ requires: *m/z* = 486.1759 ($MH^+$). $\nu_{max}$ 3391, 3360, 3052, 2978, 2918, 2852, 2786, 2761, 1650, 1637, 1605, 1428, 1356, 1330, 1255, 1079, 877, 741, 670 $cm^{-1}$. Purity: UPLC (method I, 254 nm): $t_r$ = 4.797 min, 96.1% total area. $^{1}$H NMR (400 MHz, $CDCl_3$) δ = 1.14 – 1.23 (m, 2H), 1.35 – 1.72 (m, 13H), 2.21 (s, 3H), 2.40 – 2.44 (m, 2H), 2.99 (s, 3H), 3.01 (s, 3H), 3.67 (s, 2H), 7.01 – 7.05 (m, 1H), 7.14 (d, *J*=2.3 Hz, 1H), 7.61 (dd, *J*=0.8 Hz, 7.2 Hz, 1H), 7.77 – 7.80 (m, 1H), 8.58 (s, 1H). $^{13}$C NMR (101 MHz, $CDCl_3$) δ = 26.59, 28.63, 34.93, 35.96, 36.06, 37.70, 38.83, 42.44, 53.23, 55.90, 95.80, 114.65, 120.42, 121.86, 123.73, 127.69, 134.54, 140.90, 157.06. $^{125}$Te (158 MHz, $CDCl_3$) δ = 592.

*O-(3-(((2-Cycloheptylethyl)(methyl)amino)methyl)-1H-indol-7-yl) dimethylcarbamothioate* (**5**)

1*H*-Indol-7-ol[110] (140 mg, 1.05 mmol), dimethylthiocarbamoyl chloride (1.1 eq., 143 mg), and caesium carbonate (1.0 eq., 313 mg) in THF (3 mL) were stirred at rt under Ar for 18 h. Impure *O*-(1*H*-indol-7-yl) dimethylcarbamothioate **5a** was isolated by *GP3* (37 mg) and immediately used further following *GP2* to afford *O*-(3-(((2-cycloheptylethyl)(methyl)amino)methyl)-1*H*-indol-7-yl) dimethylcarbamothioate **5**. Yield: 24 mg (0.0619 mmol, 5.9% over two steps) of beige semisolid. ESI-HRMS: *m/z* = 388.2409 ($MH^+$); $C_{22}H_{34}N_3OS$ requires: *m/z* = 388.2417 ($MH^+$). $\nu_{max}$ 3206, 3108, 3058, 3027, 2924, 2849, 1631, 1495, 1449, 1333, 1303, 1221, 1170, 1074, 732, 693 $cm^{-1}$. Purity: UPLC (method I, 254 nm): $t_r$ = 4.530 min, 99.6% total area. $^1H$ NMR (400 MHz, $CDCl_3$) δ = 1.13 – 1.23 (m, 2H), 1.34 – 1.72 (m, 13H), 2.22 (s, 3H), 2.40 – 2.46 (m, 2H), 3.40 (s, 3H), 3.48 (s, 3H), 3.67 (s, 2H), 6.89 (d, *J*=7.7 Hz, 1H), 7.08 – 7.13 (m, 2H), 7.60 (d, *J*=7.9 Hz, 1H), 8.19 (s, 1H). $^{13}C$ NMR (101 MHz, $CDCl_3$) δ = 26.58, 28.61, 34.89, 35.88, 37.66, 38.96, 42.46, 43.56, 52.99, 55.94, 114.50, 115.11, 117.63, 119.63, 124.14, 129.44, 130.95, 139.32, 186.90.

*(3-(((2-Cycloheptylethyl)(methyl)amino)methyl)-1H-indol-7-yl)(methyl)carbamic fluoride* (**6**)

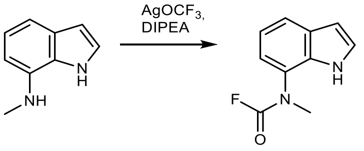

(1*H*-Indol-7-yl)(methyl)carbamic fluoride **6a** was prepared from *N*-methyl-1*H*-indol-7-amine (206 mg, 1.41 mmol) according to the reported procedure[164] and isolated by column chromatography on silica (PE/EtOAc = 10:1). Yield: 119 mg (0.619 mmol, 43.9%) of beige semisolid. ESI-HRMS: *m/z* = 193.0777 ($MH^+$); $C_{10}H_{10}FN_2O$ requires: *m/z* = 193.0772 ($MH^+$). $\nu_{max}$ 3361, 3310, 3113, 1792, 1589, 1497, 1421, 1390, 1339, 1294, 1279, 1187, 1138, 1068, 1012, 915, 801, 732, 641, 565, 515 $cm^{-1}$. Purity: UPLC (method I, 220 nm): $t_r$ = 3.887 min, 99.5% total area. $^1H$ NMR (400 MHz, $CDCl_3$) δ = 3.39 (s, 3H), 6.55 and 6.57 (2 × dd, *J*=2.0 Hz, 3.1 Hz, 1H), 7.02 – 7.11 (m, 2H), 7.12 – 7.14 and 7.18 – 7.20 (2 × m, 1H), 7.59 – 7.63 (m, 1H), 8.95 and 9.47 (2 × s, 1H). $^{13}C$ NMR (101 MHz, $CDCl_3$) δ = 38.71 (d, *J*=2.5 Hz), 38.82 (d, *J*=2.0 Hz), 103.14, 103.24, 118.18, 119.16, 119.84, 119.93, 120.93, 121.28, 124.52, 125.55, 125.60, 125.67, 130.30, 130.50, 130.62, 131.56, 147.11 (d, *J*=292.0 Hz), 147.84 (d, *J*=286.6 Hz). $^{19}F$ NMR (376 MHz, $CDCl_3$) δ = –16.99, –15.92. Two sets of conformer signals.

(3-(((2-Cycloheptylethyl)(methyl)amino)methyl)-1*H*-indol-7-yl)(methyl)carbamic fluoride **6** was prepared from **6a** (60 mg, 0.314 mmol) following *GP2*. Yield: 51 mg (0.143 mmol, 45.5%) of beige semisolid. ESI-HRMS: *m/z* = 360.2438 ($MH^+$); $C_{21}H_{31}ON_3F$ requires: *m/z* = 360.2446 ($MH^+$). $\nu_{max}$ 3149, 2918, 2851, 1787, 1629, 1586, 1459, 1347, 1291, 1199, 1122, 728 $cm^{-1}$. Purity: UPLC (method I, 220 nm): $t_r$ = 4.437 min, 99.5% total area. $^1H$ NMR (400 MHz, $CDCl_3$) δ = 1.12 – 1.22 (m, 2H), 1.34 – 1.70 (m, 13H), 2.21 and 2.22 (2 × s, 3H), 2.41 – 2.46 (m, 2H), 3.43 and 3.47 (2 × s, 3H), 3.68 (s, 2H), 7.04 – 7.19 (m, 3H), 7.69 – 7.75 (m, 1H), 8.48 and 8.88 (2 × s, 1H). $^{13}C$ NMR (101 MHz, $CDCl_3$) δ = 26.61, 28.61, 34.92, 35.93, 37.62, 37.66, 38.87,

42.47, 52.99, 55.96, 114.77, 117.96, 119.32, 119.73, 120.23, 124.53, 125.72, 130.34, 130.64, 131.00, 131.93, 147.52 (d, *J*=286.9 Hz). $^{19}$F NMR (376 MHz, $CDCl_3$) δ = –16.31, –15.78.

*Benzyl(methyl)carbamic fluoride (**6b**)*

Prepared according to literature.[165] ESI-HRMS: *m/z* = 168.0823 ($MH^+$); $C_9H_{11}NOF$ requires: *m/z* = 168.0819 ($MH^+$). $\nu_{max}$ 3065, 3032, 2935, 1776, 1455, 1402, 1221, 1112, 999, 748, 697 $cm^{-1}$. Purity: UPLC (method II, 220 nm): $t_r$ = 2.407 min, 99.1% total area. $^1$H NMR (400 MHz, $CDCl_3$) δ = 2.88 and 2.93 (s, 3H), 4.43 and 4.47 (s, 2H), 7.22 – 7.26 (m, 1H), 7.27 – 7.41 (m, 4H). $^{13}$C NMR (101 MHz, $CDCl_3$) δ = 33.9 (d, *J*=3.8 Hz), 35.0 (d, *J*=1.5 Hz), 53.0 (d, *J*=3.6 Hz), 53.5 (d, *J*=1.0 Hz), 127.5, 128.1, 128.2, 128.9, 129.0, 135.4, 147.6 (d, *J*=286.1 Hz), 148.2 (d, *J*=286.9 Hz). $^{19}$F NMR (376 MHz, $CDCl_3$) δ = –23.4, –21.8.

*Benzyl(methyl)carbamothioic fluoride* (**6c**)

Prepared according to literature.[166] ESI-HRMS: *m/z* = 184.0595 ($MH^+$); $C_9H_{11}FNS$ requires: *m/z* = 184.0591 ($MH^+$). $\nu_{max}$ 3063, 3031, 2944, 1529, 1453, 1405, 1260, 1238, 1134, 1086, 701 $cm^{-1}$. Purity: UPLC (method II, 220 nm): $t_r$ = 2.893 min, 99.2% total area. $^1$H NMR (400 MHz, $CDCl_3$) δ = 3.07 (d, *J*=2.3 Hz) and 3.27 (s, 3H), 4.68 and 4.99 (2 × s, 2H), 7.20 – 7.24 (m, 1H), 7.32 – 7.41 (m, 4H). $^{19}$F NMR (376 MHz, $CDCl_3$) δ = 13.24, 17.22.

*Benzyl(methyl)carbamoselenoic fluoride* (**6d**)

Prepared according to literature.[167] Briefly, tetramethylammonium trifluoromethylselenate[168] (1.5 eq., 250 mg) and *N*-methylbenzylamine (0.75 mmol, 97 µL) in MeCN (2 mL) were stirred at rt under Ar for 1 h, the solvent was removed *in vacuo* and benzyl(methyl)carbamoselenoic fluoride isolated by column chromatography on silica (PE/DCM = 5:1). Yield: 95 mg (0.41 mmol, 55.0%) of beige-yellowish oil. ESI-HRMS: *m/z* = 232.0041 ($MH^+$); $C_9H_{11}FNSe$ requires: *m/z* = 232.0035 ($MH^+$).

$\nu_{max}$ 3063, 3031, 2946, 1786, 1529, 1453, 1401, 1261, 1223, 1109, 1072, 698 $cm^{-1}$. Purity: UPLC (method II, 254 nm): $t_r$ = 2.843 min, 98.4% total area. $^1$H NMR (400 MHz, $CDCl_3$) δ = 3.04 (d, *J*=2.8 Hz) and 3.35 (s, 3H), 4.71 and 5.12 (s, 2H), 7.20 – 7.23 (m, 1H), 7.35 – 7.43 (m, 4H). $^{13}$C NMR (101 MHz, $CDCl_3$) δ = 35.66 (d, *J*=8.7 Hz), 42.92 (d, *J*=7.2 Hz), 55.15 (d, *J*=7.3 Hz), 60.71 (d, *J*=6.9 Hz), 127.73, 128.13, 128.75, 129.06, 129.24, 133.07, 133.26, 184.44 (d, *J*=152.7 Hz), 187.91 (d, *J*=152.7 Hz). $^{19}$F NMR (376 MHz, $CDCl_3$) δ = 24.68, 29.13. $^{77}$Se NMR (95 MHz, $CDCl_3$) δ = 258, 287.

*2,2,2-Trifluoroethyl (3-(((2-cycloheptylethyl)(methyl)amino)methyl)-1H-indol-7-yl)(methyl)carbamate* (**7**)

Following *GP1*, 2,2,2-trifluoroethyl (1*H*-indol-7-yl)(methyl)carbamate **7a** was prepared from sodium trifluoroethanolate and isolated by column chromatography on silica (PE/EtOAc = 5:1). Yield: 238 mg (0.876 mmol, 87.6%) of beige solid. mp = 82.4–84.6 °C. ESI-HRMS: *m/z* = 273.0842 ($MH^+$); $C_{12}H_{12}N_2O_2F_3$ requires: *m/z* = 273.0845 ($MH^+$). $\nu_{max}$ 3284, 1707, 1687, 1439, 1384, 1337, 1280, 1160, 1130, 964, 798, 730, 673 $cm^{-1}$. Purity: UPLC (method I, 254 nm): $t_r$ = 4.420 min, 96.7% total area. $^1$H NMR (400 MHz, $CDCl_3$) δ = 3.29 – 3.38 (m, 3H), 4.33 – 4.44 and 4.54 – 4.66 (2 × m, 2H), 6.50 – 6.53 (m, 1H), 6.96 – 7.01 (m, 1H), 7.01 (t, *J*=2.8 Hz, 1H),

7.06 (t, *J*=7.7 Hz, 1H), 7.55 (d, *J*=7.8 Hz, 1H), 8.39 and 8.86 (2 × s, 1H). $^{13}$C NMR (101 MHz, $CDCl_3$) δ = 38.10, 61.38 (q, *J*=36.5 Hz), 103.03, 118.14, 119.37, 119.89, 120.41, 120.68, 121.55, 124.31, 124.98, 126.01, 127.27, 129.41, 129.93, 130.18, 131.01, 131.79, 153.86, 154.47. $^{19}$F NMR (376 MHz, $CDCl_3$) δ = –74.01 (t, *J*=8.2 Hz). Two sets of conformer signals.

2,2,2-Trifluoroethyl (3-(((2-cycloheptylethyl)(methyl)amino)methyl)-1*H*-indol-7-yl)(methyl)carbamate **7** was prepared following *GP2* from **7a** (65 mg, 0.239 mmol). Yield: 74 mg (0.168 mmol, 70.4%) of beige solid. mp 80.6–82.1 °C. ESI-HRMS: *m/z* = 440.2513 ($MH^+$); $C_{23}H_{33}N_3O_2F_3$ requires: *m/z* = 440.2519 ($MH^+$). $\nu_{max}$ 3289, 2921, 2853, 2793, 2761, 1706, 1441, 1279, 1161, 742, 675 cm$^{-1}$. Purity: UPLC (method I, 254 nm): $t_r$ = 4.637 min, 95.3% total area. $^1$H NMR (400 MHz, $CDCl_3$) δ = 1.14 – 1.23 (m, 2H), 1.35 – 1.72 (m, 13H), 2.22 (s, 3H), 2.42 – 2.47 (m, 2H), 3.37 – 3.47 (m, 3H), 3.68 (s, 2H), 4.41 – 4.51 and 4.57 – 4.67 (2 × m, 2H), 7.00 – 7.07 (m, 1H), 7.08 – 7.14 (m, 2H), 7.68 (d, *J*=7.8 Hz, 1H), 8.27 and 8.43 (2 × s, 1H). $^{13}$C NMR (101 MHz, $CDCl_3$) δ = 26.57, 28.58, 34.89, 35.89, 37.62, 38.30, 42.41, 52.98, 55.92, 61.46 (q, *J*=37.2 Hz), 114.42, 118.01, 118.92, 119.38, 119.68, 121.62, 124.10, 125.98, 127.29, 130.00, 130.32, 131.56, 132.23, 153.69, 154.40. $^{19}$F NMR (376 MHz, $CDCl_3$) δ = –74.07 (t, *J*=8.4 Hz). Two sets of rotamer signals. The $CF_3$ quartet in the aromatic region is unresolved, hence the peaks are listed as such.

*1,1,1,3,3,3-Hexafluoropropan-2-yl (3-(((2-cycloheptylethyl)(methyl)amino)methyl)-1H-indol-7-yl)(methyl)carbamate* (**8**)

Following *GP1*, 1,1,1,3,3,3-hexafluoropropan-2-yl (1*H*-indol-7-yl)(methyl)carbamate **8a** was prepared from hexafluoroisopropanol and isolated by column chromatography on silica (PE/EtOAc = 10:1). Yield: 197 mg (0.522 mmol, 52.2%) of beige semisolid. ESI-HRMS: *m/z* = 341.0726 ($MH^+$); $C_{13}H_{11}F_6N_2O_2$ requires: *m/z* = 341.0719 ($MH^+$). $\nu_{max}$ 3503, 3346, 2985, 1748, 1719, 1386, 1279, 1228, 1194, 1135, 1107, 801, 731 cm$^{-1}$. Purity: UPLC (method II, 220 nm): $t_r$ = 3.460 min, 97.1% total area. $^1$H NMR (400 MHz, $CDCl_3$) δ = 3.35 and 3.42 (2 × s, 3H), 5.78 and 6.01 (2 × dt, *J*=5.8 Hz, 11.8 Hz, 1H), 6.52 and 6.56 (s, 1H), 6.86 – 6.90 and 6.96 – 7.00 and 7.04 – 7.13 (3 × m, 3H), 7.55 – 7.62 (m, 1H), 8.12 and 8.37 (s, 1H). $^{13}$C NMR (101 MHz, $CDCl_3$) δ = 38.45, 38.52, 67.75 – 69.58 (m), 103.30, 103.62, 118.37, 119.18, 119.63, 120.11, 120.24, 120.64, 120.94, 122.00, 122.34, 124.91, 125.14, 125.28, 126.74, 130.04, 130.49, 130.75, 131.65, 152.22, 152.85. $^{19}$F NMR (376 MHz, $CDCl_3$) δ = –73.67 and –73.55 (2 × d, *J*=6.1 Hz). Overlain signals of two conformers in a 60:40 ratio.

1,1,1,3,3,3-Hexafluoropropan-2-yl (3-(((2-cycloheptylethyl)(methyl)amino)methyl)-1*H*-indol-7-yl)(methyl)carbamate **8** was prepared following *GP2* from **8a** (130 mg, 0.382 mmol). Yield: 102 mg (0.201 mmol, 52.6%) of beige semisolid. ESI-HRMS: *m/z* = 508.2402 ($MH^+$); $C_{24}H_{32}F_6N_3O_2$ requires: *m/z* = 508.2393 ($MH^+$). $\nu_{max}$ 3313, 2922, 2852, 2776, 1720, 1446, 1386, 1281, 1222, 1194, 1136, 1108, 911, 738, 688 cm$^{-1}$. Purity: UPLC (method I, 220 nm): $t_r$ = 4.880 min, 99.5% total area. $^1$H NMR (400 MHz, $CDCl_3$) δ 1.14 – 1.23 (m, 2H), 1.33 – 1.72 (m, 13H), 2.23 (s, 3H), 2.44 (t, *J* = 7.2 Hz, 2H), 3.40 and 3.49 (2 × s, 3H), 3.68 (s, 2H), 5.76 and 5.93 (2 × hept, *J* = 6.0 Hz, 1H), 6.97 – 7.19 (m, 3H), 7.71 (d, *J* = 7.8 Hz, 1H), 8.12 and 8.29 (2 × s, 1H). $^{13}$C NMR (101 MHz, $CDCl_3$) δ = 26.47, 28.50, 34.78, 35.76, 37.51, 38.43, 38.55, 42.30,

52.87, 55.80, 67.62 – 69.26 (m), 114.49, 118.14, 119.03, 119.37, 119.54, 119.77, 121.84, 122.16, 124.00, 124.37, 124.94, 126.56, 129.97, 130.46, 131.13, 131.97, 151.89, 152.63. $^{19}$F NMR (376 MHz, $CDCl_3$) δ –73.63 and –73.49 (2 × d, *J* = 6.0 Hz). A mixture of two conformers in a ratio of 60:40.

*1,1,1,3,3,3-Hexafluoropropan-2-yl benzyl(methyl)carbamate* (**8b**)

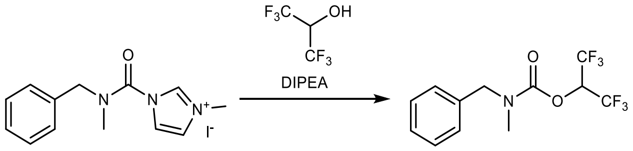

1-(Benzyl(methyl)carbamoyl)-3-methyl-1*H*-imidazol-3-ium iodide (1052 mg, 2.95 mmol), hexafluoroisopropanol (1 mL), and DIPEA (2 mL) in MeCN (10 mL) were stirred at 80 °C for 6 h. The solvent was removed *in vacuo* and 1,1,1,3,3,3-hexafluoropropan-2-yl benzyl(methyl)carbamate **8b** isolated by column chromatography on silica (PE/DCM = 10:1). Yield: 881 mg (2.79 mmol, 94.7%) of colourless oil. ESI-HRMS: *m/z* = 316.0774 ($MH^+$); $C_{12}H_{12}F_6NO_2$ requires: *m/z* = 316.0767 ($MH^+$). $\nu_{max}$ 3034, 2967, 1734, 1389, 1283, 1230, 1190, 1127, 1106, 896, 753, 730, 689 $cm^{-1}$. Purity: UPLC (method II, 254 nm): $t_r$ = 3.400 min, 99.9% total area. $^1$H NMR (400 MHz, $CDCl_3$) δ = 2.91 and 2.96 (2 × s, 3H), 4.51 and 4.53 (2 × s, 2H), 5.83 (heptd, *J*=3.8 Hz, 6.2 Hz, 1H), 7.19 – 7.25 (m, 2H), 7.28 – 7.38 (m, 3H). $^{13}$C NMR (101 MHz, $CDCl_3$) δ = 33.75, 35.23, 52.77, 53.57, 68.32 and 68.40 (2 × sept, *J*=34.7 Hz), 120.89 (q, *J*=283.7 Hz), 127.59, 127.89, 128.08, 128.10, 128.99, 135.94, 136.09, 152.57, 153.36. $^{19}$F NMR (376 MHz, $CDCl_3$) δ = –73.75, –73.57.

*Perfluoro-tert-butyl (3-(((2-cycloheptylethyl)(methyl)amino)methyl)-1H-indol-7-yl)(methyl)carbamate* (**9**)

Following *GP1*, perfluoro-*tert*-butyl (1*H*-indol-7-yl)(methyl)carbamate **9a** was prepared from perfluoro-*tert*-butanol and isolated by column chromatography on silica (PE/EtOAc = 10:1). Yield: 104 mg (0.252 mmol, 25.2%) of beige solid. mp 97.2–98.4 °C. ESI-HRMS: *m/z* = 409.0599 ($MH^+$); $C_{14}H_{10}F_9N_2O_2$ requires: *m/z* = 409.0593 ($MH^+$). $\nu_{max}$ 3383, 2922, 1759, 1587, 1496, 1377, 1246, 1133, 975, 800, 728 $cm^{-1}$. Purity: UPLC (method I, 220 nm): $t_r$ = 5.103 min, 96.3% total area. $^1$H NMR (400 MHz, $CDCl_3$) δ = 3.43 and 3.51 (2 × s, 3H), 6.60 – 6.64 (m, 1H), 7.02 – 7.16 (m, 2H), 7.21 – 7.25 (m, 1H), 7.61 – 7.65 (m, 1H), 8.13 and 8.24 (2 × s, 1H). $^{13}$C NMR (101 MHz, $CDCl_3$) δ = 38.52, 39.32, 78.99 – 80.81 (m), 103.67, 115.23, 115.26, 115.59, 118.14, 118.19, 118.50, 119.51, 120.18, 120.66, 121.10, 121.40, 123.94, 124.89, 125.39, 126.56, 130.08, 130.54, 130.68, 131.55, 148.26, 149.00. $^{19}$F NMR (376 MHz, $CDCl_3$) δ = –69.89, –69.37. Rotamer signal duplication and overlay does not allow the unambiguous assignment of $CF_3$ multiplet, the peaks are reported as such.

Perfluoro-*tert*-butyl (3-(((2-cycloheptylethyl)(methyl)amino)methyl)-1*H*-indol-7-yl)(methyl)carbamate was prepared following *GP2* from **9a** (77 mg, 0.189 mmol). Yield: 69 mg (0.120 mmol, 63.6%) of beige semisolid. ESI-HRMS: *m/z* = 576.2276 ($MH^+$); $C_{25}H_{31}F_9N_3O_2$ requires: *m/z* = 576.2267 ($MH^+$). $\nu_{max}$ 2922, 2853, 2792, 1755, 1445, 1371, 1249, 1132, 1000, 976, 728 $cm^{-1}$. Purity: UPLC (method I, 220 nm): $t_r$ = 5.057 min, 99.7% total area.

$^1H$ NMR (400 MHz, $CDCl_3$) δ = 1.12 – 1.22 (m, 2H), 1.33 – 1.72 (m, 13H), 2.20 (s, 3H), 2.38 – 2.45 (m, 2H), 3.41 and 3.49 (2 × s, 3H), 3.68 (s, 2H), 6.99 – 7.15 (m, 3H), 7.67 – 7.73 (m, 1H), 8.02 and 8.16 (2 × s, 1H). $^{13}C$ NMR (101 MHz, $CDCl_3$) δ = 26.59, 28.62, 34.90, 35.91, 37.70, 38.54, 39.34, 42.42, 52.97, 55.82, 55.93, 79.41 – 80.34, 114.79, 114.88, 115.18, 118.12, 118.22, 118.51, 119.43, 119.61, 119.83, 120.05, 121.03, 121.40, 124.01, 124.56, 125.26, 126.41, 130.16, 130.63, 131.18, 131.99, 148.22, 148.95. $^{19}F$ NMR (376 MHz, $CDCl_3$) δ = –69.86, –69.36. Rotamer signal duplication and overlay does not allow the unambiguous assignment of $CF_3$ multiplet, the peaks are reported as such. Two conformers in a 72:18 ratio.

*Phenyl (3-(((2-cycloheptylethyl)(methyl)amino)methyl)-1H-indol-7-yl)(methyl)carbamate* (**10**)

Following *GP1,* phenyl (1*H*-indol-7-yl)(methyl)carbamate **10a** was prepared from phenol, and isolated by column chromatography on silica (PE/EtOAc = 5:1). Yield: 220 mg (0.826 mmol, 82.6%) of beige solid. mp 101.7–103.4 °C. ESI-HRMS: *m/z* = 267.1116 ($MH^+$); $C_{16}H_{15}N_2O_2$ requires: *m/z* = 267.1128 ($MH^+$). $\nu_{max}$ 3254, 3115, 3072, 2925, 1708, 1692, 1588, 1495, 1433, 1379, 1339, 1288, 1199, 1163, 1150, 1121, 923, 797, 758, 717, 690 $cm^{-1}$. Purity: UPLC (method I, 254 nm): $t_r$ = 4.560 min, 97.8% total area. $^1H$ NMR (400 MHz, $CDCl_3$) δ = 3.45 and 3.63 (2 × s, 3H), 6.56 (s, 1H), 6.97 – 7.33 (m, 8H), 7.59 – 7.65 (m, 1H), 8.58 and 9.38 (2 × s, 1H). $^{13}C$ NMR (101 MHz, $CDCl_3$) δ = 38.20, 102.80, 119.56, 119.86, 120.35, 121.75, 125.22, 125.55, 126.78, 129.27, 130.03, 132.15, 151.35, 155.14.

Phenyl (3-(((2-cycloheptylethyl)(methyl)amino)methyl)-1*H*-indol-7-yl)(methyl)carbamate was prepared following *GP2* from **10a** (87 mg, 0.327 mmol). Yield: 88 mg (0.203 mmol, 62.0%) of beige solid. mp 98.3–100.1 °C. ESI-HRMS: *m/z* = 434.2780 ($MH^+$); $C_{27}H_{36}N_3O_2$ requires: *m/z* = 434.2802 ($MH^+$). $\nu_{max}$ 3274, 2917, 2850, 2764, 1701, 1494, 1432, 1380, 1210, 1165, 752, 738 $cm^{-1}$. Purity: UPLC (method I, 254 nm): $t_r$ = 4.680 min, 96.3% total area. $^1H$ NMR (400 MHz, $CDCl_3$) δ = 1.13 – 1.23 (m, 2H), 1.33 – 1.71 (m, 13H), 2.23 (s, 3H), 2.40 – 2.50 (m, 2H), 3.43 and 3.57 (2 × s, 3H), 3.69 (s, 2H), 6.97 – 7.43 (m, 8H), 7.64 – 7.69 (m, 1H), 8.44 and 8.89 (2 × s, 1H). $^{13}C$ NMR (101 MHz, $CDCl_3$) δ = 26.59, 28.62, 34.91, 35.83, 37.64, 38.27, 42.38, 52.94, 55.87, 113.88, 118.53, 119.20, 119.74, 121.76, 124.46, 125.50, 126.79, 129.30, 130.06, 132.57, 151.37, 154.87.

*Perfluorophenyl (3-(((2-cycloheptylethyl)(methyl)amino)methyl)-1H-indol-7-yl)(methyl)carbamate* (**11**)

Following *GP1,* perfluorophenyl (1*H*-indol-7-yl)(methyl)carbamate **11a** was prepared from pentafluorophenol, and isolated by column chromatography on silica (1. PE/EtOAc = 10:1; 2. PE/EtOAc = 5:1). Yield: 283 mg (0.794 mmol, 79.4%) of beige solid. mp 137.8–139.7 °C. ESI-HRMS: *m/z* = 357.0645 ($MH^+$); $C_{16}H_{10}N_2O_2F_5$ requires: *m/z* = 357.0657 ($MH^+$), *m/z* = 355.0513 ($(M–H)^-$); $C_{16}H_8N_2O_2F_5$ requires: *m/z* = 355.0511 ($(M–H)^-$). $\nu_{max}$ 3364, 1732, 1518, 1427, 1377, 1290, 1142, 1033, 994, 977, 801, 728 $cm^{-1}$. Purity: UPLC (method I, 220 nm): $t_r$ = 4.953 min, 98.8% total area. $^1H$ NMR (400 MHz, $CDCl_3$) δ = 3.40 and 3.59 (2 × s, 3H), 6.51 – 6.58 (m, 1H), 7.05 – 7.17 (m, 3H), 7.52 – 7.62 (m, 1H), 8.24 and 8.53 (2 × s, 1H). $^{13}C$ NMR (101 MHz, CDCl3) δ = 38.62, 39.12, 103.40, 117.98, 119.70, 120.19, 120.36, 120.96, 125.06, 125.30,

125.78, 125.94, 126.08, 126.78, 130.05, 130.40, 130.65, 131.65, 136.30 – 136.78 (m), 137.81 – 138.26 (m), 138.85 – 139.26 (m), 140.21 – 140.79 (m), 142.69 – 143.00 (m), 151.07, 151.85. $^{19}$F NMR (376 MHz, $CDCl_3$) δ = –162.85 (m, major isomer), –162.59 – –162.44 (m, minor isomer), –158.70 – –158.54 (m, major isomer), –158.42 – –158.27 (m, minor isomer), –153.72 – –153.58 (m, both isomers). Two conformers in an 82:18 ratio.

Perfluorophenyl (3-(((2-cycloheptylethyl)(methyl)amino)methyl)-1*H*-indol-7-yl)(methyl)carbamate was prepared following *GP2* from **11a** (105 mg, 0.295 mmol). Yield: 90 mg (0.172 mmol, 58.3%) of beige oil. ESI-HRMS: *m/z* = 524.2317 ($MH^+$); $C_{27}H_{31}N_3O_2F_5$ requires: *m/z* = 524.2331 ($MH^+$). $\nu_{max}$ 3355, 2921, 2852, 2793, 1735, 1519, 1446, 1375, 1285, 995, 734 $cm^{-1}$. Purity: UPLC (method I, 254 nm): $t_r$ = 4.890 min, 95.8% total area. $^1$H NMR (400 MHz, $CDCl_3$) δ = 1.14 – 1.24 (m, 2H), 1.35 – 1.74 (m, 13H), 2.23 and 2.25 (2 × s, 3H), 2.41 – 2.51 (m, 2H), 3.46 and 3.66 (2 × s, 3H), 3.69 (s, 2H), 7.11 – 7.21 (m, 3H), 7.68 – 7.78 (m, 1H), 8.21 and 8.52 (2 × s, 1H). $^{13}$C NMR (101 MHz, $CDCl_3$) δ = 26.57, 28.58, 34.89, 35.90, 37.56, 38.76, 42.45, 52.98, 56.00, 114.49, 114.92, 117.89, 119.28, 119.74, 119.91, 119.97, 124.21, 124.50, 125.65, 125.86 – 126.26 (m), 126.71, 130.15, 130.55, 131.17, 132.12, 136.18 – 136.84 (m), 137.82 – 138.36 (m), 138.74 – 139.36 (m), 140.20 – 140.82 (m), 142.74 – 143.17 (m), 150.93, 151.81. $^{19}$F NMR (376 MHz, $CDCl_3$) δ = –162.86 – –162.68 and –162.55 – –162.39 (2 × m), –158.58 and –158.33 (2 × t, *J*=21.7 Hz), –153.61 – –153.47 and –153.47 – –153.35 (2 × m).

*Isoxazol-3-yl (3-(((2-cycloheptylethyl)(methyl)amino)methyl)-1H-indol-7-yl)(methyl)carbamate* (**12**)

Following *GP1*, isoxazol-3-yl (1*H*-indol-7-yl)(methyl)carbamate **12a** was prepared from 3-hydroxyisoxazole[169] and isolated by column chromatography on silica (PE/EtOAc = 1:1). Yield: 142 mg (0.553 mmol, 55.3%) of beige solid. mp 146.8–148.7 °C. ESI-HRMS: *m/z* = 258.0866 ($MH^+$); $C_{13}H_{12}N_3O_3$ requires: *m/z* = 258.0873 ($MH^+$). $\nu_{max}$ 3302, 3179, 3111, 2921, 1752, 1573, 1416, 1363, 1264, 1114, 1041, 949, 784, 728, 635 $cm^{-1}$. Purity: UPLC (method I, 220 nm): $t_r$ = 4.090 min, 95.7% total area. $^1$H NMR (400 MHz, $CDCl_3$) δ = 3.43 and 3.49 (2 × s, 3H), 6.31 – 6.33 and 6.55 – 6.60 (2 × m, 2H), 7.09 – 7.14 (m, 3H), 7.59 – 7.66 (m, 1H), 8.15 – 8.19 and 8.26 – 8.30 (2 × m, 1H), 8.95 and 9.25 (2 × s, 1H). $^{13}$C NMR (101 MHz, $CDCl_3$) δ = 38.52, 38.77, 99.52, 103.08, 118.39, 119.51, 119.92, 120.45, 120.85, 125.39, 125.77, 126.71, 130.17, 130.30, 131.07, 131.77, 150.84, 151.79, 160.39, 166.39, 166.54. A mixture of conformers in a 67:33 ratio.

Isoxazol-3-yl (3-(((2-cycloheptylethyl)(methyl)amino)methyl)-1*H*-indol-7-yl)(methyl)carbamate was prepared following *GP2* from **12a** (79 mg, 0.307 mmol). Yield: 99 mg (0.233 mmol, 76.0%) of beige solid. mp = 142.7–144.5 °C. ESI-HRMS: *m/z* = 425.2526 ($MH^+$); $C_{24}H_{33}N_4O_3$ requires: *m/z* = 425.2547 ($MH^+$). $\nu_{max}$ 3276, 2914, 2852, 2790, 1765, 1578, 1458, 1425, 1363, 1349, 1278, 1135, 1028, 943, 887, 770, 742, 656 $cm^{-1}$. Purity: UPLC (method I, 254 nm): $t_r$ = 4.543 min, 100.0% total area. $^1$H NMR (400 MHz, $CDCl_3$) δ = 1.14 – 1.23 (m, 2H), 1.35 – 1.73 (m, 13H), 2.20 and 2.22 (2 × s, 3H), 2.41 – 2.49 (m, 2H), 3.41 and 3.47 (2 × s, 3H), 3.67 (s, 2H), 6.30 – 6.33 and 6.55 – 6.58 (2 × m, 1H), 7.03 – 7.07 (m, 1H),

7.08 – 7.15 (m, 2H), 7.67 – 7.73 (m, 1H), 8.14 – 8.19 and 8.25 – 8.30 (2 × m, 1H), 8.87 and 9.19 (2 × s, 1H). $^{13}$C NMR (101 MHz, $CDCl_3$) δ = 26.52, 28.54, 34.85, 35.87, 37.55, 38.47, 38.73, 42.36, 52.96, 55.94, 99.48, 114.01, 114.18, 118.37, 119.32, 119.54, 119.72, 124.53, 125.58, 126.53, 130.19, 130.35, 131.49, 132.16, 150.78, 151.78, 160.29, 160.36, 166.34, 166.50. A mixture of conformers in a 70:30 ratio.

*1,2,5-Thiadiazol-3-yl (3-(((2-cycloheptylethyl)(methyl)amino)methyl)-1H-indol-7-yl)(methyl)carbamate* (**13**)

Following *GP1*, 1,2,5-thiadiazol-3-yl (1*H*-indol-7-yl)(methyl)carbamate **13a** was prepared from 1,2,5-thiadiazol-3-ol[170] and isolated by column chromatography on silica (PE/EtOAc = 3:1). Yield: 251 mg (0.915 mmol, 91.5%) of beige solid. mp 108.7–111.4 °C. ESI-HRMS: *m/z* = 275.0591 ($MH^+$); $C_{12}H_{11}N_4O_2S$ requires: *m/z* = 275.05972 ($MH^+$). $\nu_{max}$ 3305, 1734, 1712, 1494, 1430, 1355, 1324, 1286, 1190, 1116, 1032, 901, 798, 771, 731, 520 cm$^{-1}$. Purity: UPLC (method I, 220 nm): $t_r$ = 4.267 min, 96.6% total area. $^1$H NMR (400 MHz, $CDCl_3$) δ = 3.45 and 3.57 (2 × s, 3H), 6.57 – 6.59 (m, 1H), 7.08 – 7.17 (m, 3H), 7.60 – 7.67 (m, 1H), 8.31 and 8.60 (2 × s, 1H), 8.73 and 9.22 (2 × s, 1H). $^{13}$C NMR (101 MHz, $CDCl_3$) δ = 38.50, 38.95, 103.19, 118.25, 119.46, 119.96, 120.47, 120.95, 125.24, 125.64, 130.12, 131.69, 142.27, 152.27, 157.95. A mixture of conformers in a 76:24 ratio.

1,2,5-Thiadiazol-3-yl (3-(((2-cycloheptylethyl)(methyl)amino)methyl)-1*H*-indol-7-yl)(methyl)carbamate was prepared following *GP2* from **13a** (163 mg, 0.594 mmol). Yield: 171 mg (0.387 mmol, 65.2%) of yellowish semisolid. ESI-HRMS: *m/z* = 442.2251 ($MH^+$); $C_{23}H_{32}N_5O_2S$ requires: *m/z* = 442.2271 ($MH^+$). $\nu_{max}$ 3301, 2918, 2849, 2769, 1726, 1449, 1357, 1324, 1189, 1127, 1032, 738, 518 cm$^{-1}$. Purity: UPLC (method I, 254 nm): $t_r$ = 4.623 min, 98.4% total area. $^1$H NMR (400 MHz, $CDCl_3$) δ = 1.14 – 1.24 (m, 2H), 1.35 – 1.72 (m, 13H), 2.21 and 2.24 (2 × s, 3H), 2.43 – 2.49 (m, 2H), 3.42 and 3.55 (2 × s, 3H), 3.68 (s, 2H), 7.00 – 7.04 (m, 1H), 7.11 – 7.15 (m, 2H), 7.69 – 7.75 (m, 1H), 8.31 and 8.58 (2 × s, 1H), 8.71 and 9.22 (2 × s, 1H). $^{13}$C NMR (101 MHz, $CDCl_3$) δ = 26.48, 28.49, 34.80, 35.80, 37.49, 38.45, 42.34, 52.90, 55.86, 114.13, 114.24, 118.24, 119.41, 119.56, 119.84, 124.41, 125.44, 126.41, 130.16, 131.33, 132.08, 142.23, 152.26, 157.93. A mixture of conformers in a 78:22 ratio.

*1H-Imidazol-1-yl (3-(((2-cycloheptylethyl)(methyl)amino)methyl)-1H-indol-7-yl)(methyl)carbamate* (**14**)

Following *GP1*, 1*H*-imidazol-1-yl (1*H*-indol-7-yl)(methyl)carbamate **14a** was prepared from *N*-hydroxyimidazole[171] and isolated by column chromatography on silica (1. PE/EtOAc = 1:1; 2. EtOAc). Yield: 246 mg (0.958 mmol, 95.8%) of beige semisolid. $\nu_{max}$ 3136, 2927, 1764, 1655, 1584, 1496, 1440, 1375, 1339, 1283, 1263, 1139, 1099, 1015, 905, 799, 723 cm$^{-1}$. Purity: UPLC (method I, 220 nm): $t_r$ = 2.370 min, 94.6% total area. $^1$H NMR (400 MHz, $CDCl_3$) δ = 3.32 and 3.48 (2 × s, 3H), 6.49 – 6.52 and 6.55 – 6.58 (2 × m, 1H), 6.90 (d, *J*=13.0 Hz, 2H), 7.06 – 7.12 (m, 2H), 7.19 – 7.22 (m, 1H), 7.55 (s, 1H), 7.63 (d, *J*=7.2 Hz, 1H), 10.74 and 10.95 (2 × s, 1H). $^{13}$C NMR (101 MHz, $CDCl_3$) δ = 38.27, 39.00, 102.82, 117.29, 119.23, 119.63,

121.14, 124.40, 124.63, 125.75, 130.31, 131.90, 132.50, 152.97. A mixture of conformers in a 82:18 ratio.

1*H*-Imidazol-1-yl (3-(((2-cycloheptylethyl)(methyl)amino)methyl)-1*H*-indol-7-yl)(methyl)carbamate was prepared following *GP2* from **14a** (144 mg, 0.562 mmol). Yield: 42 mg (0.1 mmol, 17.6%) of beige semisolid. ESI-HRMS: $m/z$ = 424.2686 ($MH^+$); $C_{24}H_{34}N_5O_2$ requires: $m/z$ = 424.2707 ($MH^+$). $\nu_{max}$ 3142, 2919, 2851, 2788, 1767, 1458, 1345, 1286, 1263, 1102, 1015, 970, 884, 731, 657 $cm^{-1}$. Purity: UPLC (method I, 220 nm): $t_r$ = 3.983 min, 96.0% total area. $^1H$ NMR (400 MHz, $CDCl_3$) δ = 1.03 – 1.13 (m, 2H), 1.22 – 1.62 (m, 13H), 2.13 (s, 3H), 2.31 – 2.38 (m, 2H), 3.28 and 3.46 (2 × s, 3H), 3.59 (s, 2H), 6.85 and 6.94 (2 × s, 2H), 6.98 – 7.12 (m, 3H), 7.41 and 7.52 (2 × s, 1H), 7.65 (d, *J*=7.3 Hz, 1H), 10.01 and 10.39 (2 × s, 1H). $^{13}C$ NMR (101 MHz, $CDCl_3$) δ = 26.57, 28.58, 34.89, 35.92, 37.58, 39.14, 42.47, 53.03, 56.02, 114.42, 117.28, 119.49, 119.53, 120.24, 124.60, 124.95, 125.05, 130.45, 132.39, 132.49, 153.28.

*1H-Pyrazol-1-yl (3-(((2-cycloheptylethyl)(methyl)amino)methyl)-1H-indol-7-yl)(methyl)carbamate* (**15**)

Following *GP1*, 1*H*-pyrazol-1-yl (1*H*-indol-7-yl)(methyl)carbamate **15a** was prepared from 1-hydroxypyrazole and isolated by column chromatography on silica (PE/EtOAc = 3:1). Yield: 146 mg (0.571 mmol, 57.1%) of beige solid. mp 185.4–186.8 °C. ESI-HRMS: $m/z$ = 257.1037 ($MH^+$); $C_{13}H_{13}N_4O_2$ requires: $m/z$ = 257.1033 ($MH^+$). $\nu_{max}$ 3314, 3128, 2949, 1763, 1743, 1588, 1495, 1433, 1389, 1375, 1337, 1289, 1255, 1111, 1041, 1014, 967, 873, 800, 748, 726, 645 $cm^{-1}$. Purity: UPLC (method I, 220 nm): $t_r$ = 4.100 min, 98.2% total area. $^1H$ NMR (400 MHz, $CDCl_3$) δ = 3.48 and 3.57 (2 × s, 3H), 6.20 – 6.25 and 6.30 – 6.36 (2 × m, 1H), 6.50 – 6.55 and 6.56 – 6.59 (2 × m, 1H), 7.12 – 7.17 (m, 3H), 7.31 (s, 1H), 7.40 (s, 1H), 7.59 – 7.68 (m, 1H), 8.84 and 9.35 (2 × s, 1H). $^{13}C$ NMR (101 MHz, $CDCl_3$) δ = 39.41, 103.11, 104.60, 119.29, 119.98, 121.09, 124.63, 125.03, 125.79, 130.35, 131.48, 134.19, 153.47.

1*H*-Pyrazol-1-yl (3-(((2-cycloheptylethyl)(methyl)amino)methyl)-1*H*-indol-7-yl)(methyl)carbamate was prepared following *GP2* from **15a** (72 mg, 0.281 mmol). Yield: 54 mg (0.127 mmol, 45.4%) of beige semisolid. ESI-HRMS: $m/z$ = 424.2716 ($MH^+$); $C_{24}H_{34}N_5O_2$ requires: $m/z$ = 424.2707 ($MH^+$). $\nu_{max}$ 3307, 2917, 2852, 2781, 1760, 1742, 1448, 1435, 1387, 1374, 1252, 1197, 1117, 1040, 976, 801, 741 $cm^{-1}$. Purity: UPLC (method I, 220 nm): $t_r$ = 4.453 min, 96.7% total area. $^1H$ NMR (400 MHz, $CDCl_3$) δ = 1.13 – 1.22 (m, 2H), 1.34 – 1.71 (m, 13H), 2.18 and 2.21 (2 × s, 3H), 2.41 – 2.46 (m, 2H), 3.46 and 3.55 (2 × s, 3H), 3.66 (s, 2H), 6.19 – 6.22 and 6.29 – 6.34 (2 × m, 1H), 6.95 and 7.03 (2 × s, 1H), 7.08 – 7.17 (m, 2H), 7.29 (s, 1H), 7.41 (s, 1H), 7.69 – 7.75 (m, 1H), 8.79 and 9.25 (2 × s, 1H). $^{13}C$ NMR (101 MHz, $CDCl_3$) δ = 26.57, 28.58, 34.89, 35.93, 37.58, 39.34, 42.43, 53.00, 55.93, 104.53, 114.23, 119.33, 119.59, 120.01, 124.57, 124.80, 124.93, 130.36, 131.93, 134.11, 153.54.

*1H-Benzotriazol-1-yl (3-(((2-cycloheptylethyl)(methyl)amino)methyl)-1H-indol-7-yl)(methyl)carbamate* (**16**)

Following *GP1*, 1*H*-benzotriazol-1-yl (1*H*-indol-7-yl)(methyl)carbamate **16a** was prepared from HOBt and isolated by column chromatography on silica (PE/EtOAc = 4:1). Yield: 226 mg (0.735 mmol, 73.5%) of dark gray solid. mp 160.5–162.3 °C. ESI-HRMS: *m/z* = 308.1128 ($MH^+$); $C_{16}H_{14}N_5O_2$ requires: *m/z* = 308.1142 ($MH^+$). $\nu_{max}$ 3377, 3069, 2956, 1762, 1586, 1495, 1442, 1423, 1360, 1337, 1278, 1141, 1083, 993, 905, 883, 769, 746, 725, 636 $cm^{-1}$. $^1$H NMR (400 MHz, $CDCl_3$) δ = 3.43 – 3.50 (m, 3H), 6.49 – 6.53 and 6.58 – 6.63 (2 × m, 1H), 7.01 – 7.52 (m, 6H), 7.57 – 7.61 (m) and 7.69 (d, *J*=7.8 Hz, 1H), 7.99 (d, *J*=8.3 Hz) and 8.05 – 8.11 (m, 1H), 9.08 and 9.73 (2 × s, 1H). $^{13}$C NMR (101 MHz, $CDCl_3$) δ = 39.43, 103.25, 108.75, 119.61, 120.02, 120.16, 121.46, 124.54, 124.94, 125.83, 128.70, 130.49, 131.63, 143.27, 152.29.

1*H*-Benzotriazol-1-yl (3-(((2-cycloheptylethyl)(methyl)amino)methyl)-1*H*-indol-7-yl)(methyl)carbamate **16** was prepared following *GP2* from **16a** (143 mg, 0.465 mmol). Yield: 119 mg (0.251 mmol, 53.9%) of beige semisolid. ESI-HRMS: *m/z* = 475.2795 ($MH^+$); $C_{27}H_{35}N_6O_2$ requires: *m/z* = 475.2816 ($MH^+$). $\nu_{max}$ 3191, 2920, 2852, 2794, 1779, 1616, 1446, 1349, 1279, 1090, 907, 728 $cm^{-1}$. Purity: UPLC (method I, 254 nm): $t_r$ = 4.617 min, 96.1% total area. $^1$H NMR (400 MHz, $CDCl_3$) δ = 1.11 – 1.20 (m, 2H), 1.29 – 1.70 (m, 13H), 2.16 and 2.21 (2 × s, 3H), 2.37 – 2.48 (m, 2H), 3.42 and 3.48 (2 × s, 3H), 3.62 and 3.67 (2 × s, 2H), 7.06 – 7.57 (m, 6H), 7.65 – 7.68 (m) and 7.76 (d, *J*=7.8 Hz, 1H), 7.95 (d, *J*=8.3 Hz) and 8.01 – 8.06 (m, 1H), 9.01 and 9.69 (2 × s, 1H). $^{13}$C NMR (101 MHz, $CDCl_3$) δ = 26.51, 28.52, 34.83, 35.85, 37.51, 39.43, 42.39, 52.98, 55.90, 108.75, 114.36, 119.59, 119.67, 120.15, 120.41, 124.40, 124.87, 125.02, 128.64, 128.71, 130.53, 132.08, 143.26, 152.28.

*2,5-Dioxopyrrolidin-1-yl (3-(((2-cycloheptylethyl)(methyl)amino)methyl)-1H-indol-7-yl)(methyl)carbamate* (**17**)

Following *GP1*, 2,5-dioxopyrrolidin-1-yl (1*H*-indol-7-yl)(methyl)carbamate **17a** was prepared from *N*-hydroxysuccinimide and isolated by column chromatography on silica (1. PE/EtOAc = 3:1; 2. PE/EtOAc = 1:1). Yield: 240 mg (0.835 mmol, 83.5%) of beige foam. mp 197.1–198.2 °C. ESI-HRMS: *m/z* = 288.0985 ($MH^+$); $C_{14}H_{14}N_3O_4$ requires: *m/z* = 288.0979 ($MH^+$). $\nu_{max}$ 3350, 1768, 1726, 1586, 1494, 1425, 1385, 1337, 1295, 1277, 1209, 1097, 990, 905, 883, 798, 728 $cm^{-1}$. Purity: UPLC (method I, 254 nm): $t_r$ = 3.797 min, 99.7% total area. $^1$H NMR (400 MHz, $CDCl_3$) δ = 2.55 – 2.73 (m, 4H), 3.42 and 3.50 (2 × s, 3H), 6.53 – 6.55 (m, 1H), 7.07 – 7.10 (m, 2H), 7.13 – 7.16 (m) and 7.21 (t, *J*=2.7 Hz, 1H), 7.58 (p, *J*=3.7 Hz, 1H), 8.94 and 9.59 (2 × s, 1H). $^{13}$C NMR (101 MHz, $CDCl_3$) δ = 25.35, 39.75, 102.93, 118.70, 119.84, 120.57, 125.06, 125.61, 130.09, 130.94, 150.48, 169.41, 170.24, 171.35.

2,5-Dioxopyrrolidin-1-yl (3-(((2-cycloheptylethyl)(methyl)amino)methyl)-1*H*-indol-7-yl)(methyl)carbamate **17** was prepared following *GP2* from **17a** (114 mg, 0.397 mmol). Yield: 56 mg (0.123 mmol, 31.0%) of beige semisolid. ESI-HRMS: *m/z* = 455.2642 ($MH^+$); $C_{25}H_{35}N_4O_4$ requires: *m/z* = 455.2653 ($MH^+$). $\nu_{max}$ 3403, 2916, 2852, 1763, 1734, 1584, 1458,

1378, 1342, 1296, 1205, 1099, 736 cm$^{-1}$. Purity: UPLC (method I, 254 nm): $t_r$ = 4.310 min, 98.3% total area. $^1$H NMR (400 MHz, $CDCl_3$) δ = 1.12 – 1.22 (m, 2H), 1.33 – 1.70 (m, 13H), 2.20 (s, 3H), 2.40 – 2.45 (m, 2H), 2.66 – 2.84 (m, 4H), 3.46 and 3.55 (2 × s, 3H), 3.64 (s, 2H), 7.09 – 7.12 (m, 2H), 7.15 (d, *J*=2.1 Hz, 1H), 7.68 (p, *J*=3.8 Hz, 1H), 8.64 and 9.43 (2 × s, 1H). $^{13}$C NMR (101 MHz, $CDCl_3$) δ = 25.51, 26.56, 28.57, 34.88, 35.87, 37.57, 39.98, 42.39, 52.89, 55.95, 114.17, 118.71, 119.61, 124.94, 124.99, 130.28, 131.43, 150.52, 169.15, 170.13, 171.52.

*2,5-Dioxopyrrolidin-1-yl dimethylcarbamate* (**17b**)

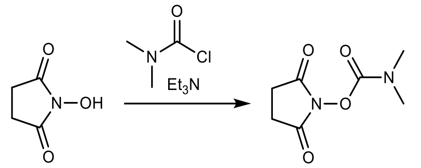

*N*-Hydroxysuccinimide (1175 mg, 10.2 mmol), *N,N*-dimethylcarbamoyl chloride (1.0 eq., 939 µL), triethylamine (5 mL) in DCM (20 mL) were stirred at rt for 3 h. Saturated sodium bicarbonate (50 mL) was then added, organic phase dried with sodium sulfate, and the volatiles removed *in vacuo* to afford 2,5-dioxopyrrolidin-1-yl dimethylcarbamate **17b** as white solid. Yield: 1008 mg (5.41 mmol, 53.1%). mp 136.0–137.1 °C. ESI-HRMS: *m/z* = 187.0718 ($MH^+$); $C_7H_{11}N_2O_4$ requires: *m/z* = 187.0713 ($MH^+$). $\nu_{max}$ 2951, 1724, 1423, 1394, 1208, 1141, 1082, 997, 963, 832, 815, 751, 739, 652, 605 cm$^{-1}$. Purity: UPLC (method II, 220 nm): $t_r$ = 0.660 min, 98.1% total area. $^1$H NMR (400 MHz, DMSO-$d_6$) δ = 2.77 (s, 4H), 2.92 and 3.01 (2 × s, 6H). $^{13}$C NMR (101 MHz, DMSO-$d_6$) δ = 25.27, 35.63, 37.17, 151.18, 170.72.

*S-Ethyl (3-(((2-cycloheptylethyl)(methyl)amino)methyl)-1H-indol-7-yl)(methyl)carbamothioate* (**18**)

Following *GP1*, *S*-ethyl (1*H*-indol-7-yl)(methyl)carbamothioate **18a** was prepared from ethyl mercaptan and isolated by column chromatography on silica (PE/EtOAc = 3:1). Yield: 56 mg (0.239 mmol, 23.9%) of beige semisolid. ESI-HRMS: *m/z* = 235.0898 ($MH^+$); $C_{12}H_{15}N_2OS$ requires: *m/z* = 235.0900 ($MH^+$). $\nu_{max}$ 3299, 3271, 2970, 2928, 1639, 1584, 1446, 1336, 1280, 1270, 1185, 1130, 1068, 1052, 1041, 960, 895, 864, 800, 731, 656, 637, 598 cm$^{-1}$. Purity: UPLC (method II, 254 nm): $t_r$ = 3.110 min, 96.3% total area. $^1$H NMR (400 MHz, $CDCl_3$) δ = 1.20 (t, *J*=7.4 Hz, 3H), 2.62 – 2.73 (m, 1H), 2.87 – 2.99 (m, 1H), 3.50 (s, 3H), 6.66 (dd, *J*=2.0 Hz, 3.1 Hz, 1H), 7.13 – 7.20 (m, 2H), 7.35 (t, *J*=2.7 Hz, 1H), 7.73 (d, *J*=7.3 Hz, 1H), 10.15 (s, 1H). $^{13}$C NMR (101 MHz, $CDCl_3$) δ = 14.81, 25.40, 37.62, 103.09, 119.98, 121.72, 121.89, 125.38, 125.53, 130.24, 132.98, 170.93.

*S*-Ethyl (3-(((2-cycloheptylethyl)(methyl)amino)methyl)-1*H*-indol-7-yl)(methyl)carbamothioate **18** was prepared *GP2* from **18a**. Yield: 45 mg (0.112 mmol, 46.9%) of beige semisolid. ESI-HRMS: *m/z* = 402.2568 ($MH^+$); $C_{23}H_{36}N_3OS$ requires: *m/z* = 402.2574 ($MH^+$). $\nu_{max}$ 3277, 3167, 3124, 2917, 2851, 2781, 1636, 1433, 1344, 1202, 1161, 1151, 1049, 960, 817, 808, 758, 645 cm$^{-1}$. Purity: UPLC (method I, 220 nm): $t_r$ = 4.597 min, 99.9% total area. $^1$H NMR (400 MHz, $CDCl_3$) δ = 1.13 – 1.23 (m, 5H), 1.36 – 1.72 (m, 13H), 2.25 (s, 3H), 2.42 – 2.48 (m, 2H), 2.64 – 2.72 (m, 1H), 2.83 – 2.92 (m, 1H), 3.45 (s, 3H), 3.63 – 3.78 (m, 2H), 7.07 – 7.17 (m, 2H), 7.21 (d, *J*=1.8 Hz, 1H), 7.77 (d, *J*=7.5 Hz, 1H), 9.65 (s, 1H). $^{13}$C

NMR (101 MHz, $CDCl_3$) δ = 14.91, 25.39, 26.59, 28.61, 34.89, 35.95, 37.59, 37.62, 42.56, 53.07, 55.95, 114.40, 119.72, 120.60, 121.94, 124.50, 125.45, 130.28, 133.35, 170.71.

*S-(2,2,2-Trifluoroethyl) (3-(((2-cycloheptylethyl)(methyl)amino)methyl)-1H-indol-7-yl)(methyl)carbamothioate* (**19**)

Following *GP1*, *S*-(2,2,2-trifluoroethyl) (1*H*-indol-7-yl)(methyl)carbamothioate **19a** was prepared from 2,2,2-trifluoroethanethiol and isolated by column chromatography on silica (1. PE/EtOAc = 10:1; 2. PE/EtOAc = 5:1). Yield: 247 mg (0.856 mmol, 85.6%) of white solid. mp 128.4–130.6 °C. ESI-HRMS: *m/z* = 289.0610 ($MH^+$); $C_{12}H_{12}N_2OF_3S$ requires: *m/z* = 289.0617 ($MH^+$). $\nu_{max}$ 3336, 3293, 2999, 2946, 1671, 1659, 1585, 1362, 1336, 1273, 1238, 1117, 1080, 958, 860, 843, 731, 638 $cm^{-1}$. Purity: UPLC (method I, 220 nm): $t_r$ = 4.600 min, 99.6% total area. $^1$H NMR (400 MHz, $CDCl_3$) δ = 3.39 – 3.64 (m, 2H), 3.48 (s, 3H), 6.65 – 6.68 (m, 1H), 7.14 – 7.21 (m, 2H), 7.30 (t, *J*=2.7 Hz, 1H), 7.74 (d, *J*=7.1 Hz, 1H), 9.05 (s, 1H). $^{13}$C NMR (101 MHz, $CDCl_3$) δ = 33.07 (q, *J*=33.6 Hz), 38.24, 103.74, 120.42, 122.45, 122.50, 124.25, 125.05 (q, *J*=275.8 Hz), 125.45, 130.45, 132.77, 167.00. $^{19}$F NMR (376 MHz, $CDCl_3$) δ = –66.52 (t, *J*=9.9 Hz).

*S*-(2,2,2-Trifluoroethyl) (3-(((2-cycloheptylethyl)(methyl)amino)methyl)-1*H*-indol-7-yl)(methyl)carbamothioate **19** was prepared following *GP2* from **19a** (133 mg, 0.461 mmol). Yield: 137 mg (0.301 mmol, 65.2%) of white solid. mp 79.0–81.8 °C. ESI-HRMS: *m/z* = 456.2269 ($MH^+$); $C_{23}H_{33}N_3OF_3S$ requires: *m/z* = 456.2291 ($MH^+$). $\nu_{max}$ 3341, 3309, 2920, 2852, 2786, 1667, 1444, 1345, 1308, 1283, 1243, 1130, 1090, 640 $cm^{-1}$. Purity: UPLC (method I, 254 nm): $t_r$ = 4.770 min, 99.6% total area. $^1$H NMR (400 MHz, $CDCl_3$) δ = 1.14 – 1.23 (m, 2H), 1.36 – 1.72 (m, 13H), 2.25 (s, 3H), 2.44 – 2.49 (m, 2H), 3.48 – 3.58 (m, 2H), 3.45 (s, 3H), 3.65 – 3.76 (m, 2H), 7.11 – 7.20 (m, 3H), 7.82 (d, *J*=7.3 Hz, 1H), 9.15 (s, 1H). $^{13}$C NMR (101 MHz, $CDCl_3$) δ = 26.56, 28.57, 33.04 (q, *J*=33.4 Hz), 34.85, 35.87, 37.54, 38.16, 42.46, 53.01, 55.93, 114.91, 119.96, 121.41, 122.36, 124.05, 124.59, 125.01 (q, *J*=275.9 Hz), 130.48, 133.18, 166.94. $^{19}$F NMR (376 MHz, $CDCl_3$) δ = –66.47 (t, *J*=10.0 Hz).

*S-(2,2,2-Trifluoroethyl) dimethylcarbamothioate* (**19b**)

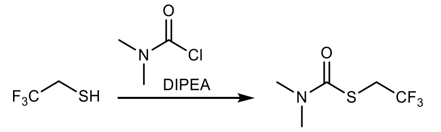

2,2,2-Trifluoroethanethiol (1 mL, 11.24 mmol), *N*,*N*-dimethylcarbamoyl chloride (1.0 eq., 1.04 mL), and DIPEA (1.2 eq., 1.92 mL) in DCM (10 mL) were stirred at rt under Ar for 2 h, the solvent was removed *in vacuo*, the residue suspended in $Et_2O$ (15 mL), sonicated for 2 min, and filtered through a 10 cm column of silica. The solvent was removed *in vacuo* to afford **19b** as a yellowish oil. Yield: 2079 mg (11.1 mmol, 98.8%). ESI-HRMS: *m/z* = 188.0352 ($MH^+$); $C_5H_9F_3NOS$ requires: *m/z* = 188.0352 ($MH^+$). $\nu_{max}$ 2947, 1740, 1668, 1369, 1307, 1273, 1240, 1127, 1081, 911, 846, 686, 639 $cm^{-1}$. Purity: UPLC (method II, 220 nm): $t_r$ = 2.193 min, 98.1% total area. $^1$H NMR (400 MHz, $CDCl_3$) δ = 3.02 (br s, 6H), 3.65 (q, *J*=10.0 Hz, 2H). $^{13}$C NMR (101 MHz, $CDCl_3$) δ = 32.58 (q, *J*=33.8 Hz), 36.56, 37.36, 125.13 (q, *J*=275.6 Hz), 164.60. $^{19}$F NMR (376 MHz, $CDCl_3$) δ = –66.85 (t, *J*=10.0 Hz).

*S-Phenyl (3-(((2-cycloheptylethyl)(methyl)amino)methyl)-1H-indol-7-yl)(methyl)carbamothioate* (**20**)

Following *GP1*, *S*-phenyl (1*H*-indol-7-yl)(methyl)carbamothioate **20a** was prepared from thiophenol and isolated by column chromatography on silica (PE/EtOAc = 5:1). Yield: 113 mg (0.400 mmol, 40.0%) of beige solid. mp 151.0–152.9 °C. ESI-HRMS: *m/z* = 283.0906 ($MH^+$); $C_{16}H_{15}N_2OS$ requires: *m/z* = 283.0900 ($MH^+$). $\nu_{max}$ 3304, 3073, 2939, 1664, 1649, 1584, 1439, 1334, 1272, 1182, 1133, 1061, 956, 893, 856, 800, 735, 724, 639 $cm^{-1}$. Purity: UPLC (method I, 254 nm): $t_r$ = 4.760 min, 95.1% total area. $^1H$ NMR (400 MHz, $CDCl_3$) δ = 3.36 (s, 3H), 6.50 (dd, *J*=2.0 Hz, 3.1, 1H), 6.83 (t, *J*=2.5 Hz, 1H), 7.10 – 7.28 (m, 5H), 7.43 – 7.47 (m, 2H), 7.67 (d, *J*=7.6 Hz, 1H), 9.82 (s, 1H). $^{13}C$ NMR (101 MHz, $CDCl_3$) δ = 38.00, 102.39, 119.73, 121.84, 125.01, 125.99, 128.92, 129.18, 129.43, 130.32, 132.73, 135.60, 169.49. $^{13}C$ NMR (101 MHz, $CDCl_3$) δ = 38.20, 102.80, 119.56, 119.86, 120.35, 121.75, 125.22, 125.55, 126.78, 129.29, 130.03, 132.15, 151.35, 155.14.

*S*-Phenyl (3-(((2-cycloheptylethyl)(methyl)amino)methyl)-1*H*-indol-7-yl)(methyl)carbamothioate **20** was prepared following *GP2* from **20a** (75 mg, 0.266 mmol). Yield: 79 mg (0.176 mmol, 66.0%) of beige oil. ESI-HRMS: *m/z* = 450.2561 ($MH^+$); $C_{27}H_{36}N_3OS$ requires: *m/z* = 450.2574 ($MH^+$). $\nu_{max}$ 3300, 2919, 2849, 2781, 1651, 1441, 1341, 1277, 1099, 908, 803, 736, 652 $cm^{-1}$. Purity: UPLC (method I, 254 nm): $t_r$ = 4.770 min, 98.6% total area. $^1H$ NMR (400 MHz, $CDCl_3$) δ = 1.13 – 1.23 (m, 2H), 1.33 – 1.72 (m, 13H), 2.23 (s, 3H), 2.44 (t, *J*=6.9 Hz, 2H), 3.35 (s, 3H), 3.63 (2 × d, *J*=12.3 Hz, 2H), 6.81 (s, 1H), 7.11 – 7.30 (m, 5H), 7.44 (d, *J*=7.3 Hz, 2H), 7.77 (d, *J*=7.2 Hz, 1H), 9.53 (s, 1H). $^{13}C$ NMR (101 MHz, $CDCl_3$) δ = 26.59, 28.60, 34.90, 35.99, 37.56, 38.01, 42.46, 53.08, 55.94, 113.71, 119.51, 120.84, 121.93, 124.91, 125.13, 128.91, 129.07, 129.59, 130.34, 133.21, 135.59, 169.33.

*S-(1-Methyl-1H-imidazol-2-yl) (3-(((2-cycloheptylethyl)(methyl)amino)methyl)-1H-indol-7-yl)(methyl)carbamothioate* (**21**)

Following *GP1*, *S*-(1-methyl-1*H*-imidazol-2-yl) (1*H*-indol-7-yl)(methyl)carbamothioate **21a** was prepared from 1-methyl-1*H*-imidazole-2-thiol and isolated by column chromatography on silica (PE/EtOAc = 1:1). Yield: 113 mg (0.395 mmol, 39.5%) of light greenish solid. mp 153.1–156.0 °C. ESI-HRMS: *m/z* = 287.0953 ($MH^+$); $C_{14}H_{15}N_4OS$ requires: *m/z* = 287.0961 ($MH^+$). $\nu_{max}$ 3197, 3138, 3110, 2937, 1716, 1450, 1369, 1336, 1289, 1211, 1133, 1111, 804, 740, 667 $cm^{-1}$. Purity: UPLC (method I, 220 nm): $t_r$ = 3.753 min, 95.3% total area. $^1H$ NMR (400 MHz, $CDCl_3$) δ = 3.26 and 3.38 (2 × s, 3H), 3.48 and 3.57 (2 × s, 3H), 6.11 and 7.27 (2 × s, 1H), 6.34 and 6.55 (2 × s, 1H), 6.43 and 6.72 (2 × s, 1H), 6.92 – 7.16 (m, 3H), 7.44 – 7.49 and 7.57 – 7.65 (2 × m, 1H), 9.74 and 10.30 (2 × s, 1H). $^{13}C$ NMR (101 MHz, $CDCl_3$) δ = 34.41, 34.88, 38.76, 40.36, 102.67, 114.98, 114.98, 118.25, 118.25, 119.07, 119.66, 120.16, 120.85, 125.37, 126.03, 129.83, 130.05, 131.11, 150.72, 150.84, 161.21, 162.47. A mixture of two conformers in a ratio of 66:34.

*S*-(1-Methyl-1*H*-imidazol-2-yl) (3-(((2-cycloheptylethyl)(methyl)amino)methyl)-1*H*-indol-7-yl)(methyl)carbamothioate **21** was prepared following *GP2* from **21a** (71 mg, 0.248 mmol).

Yield: 67 mg (0.148 mmol, 59.6%) of greenish semisolid. ESI-HRMS: *m/z* = 454.2613 ($MH^+$); $C_{25}H_{36}N_5OS$ requires: *m/z* = 454.2635 ($MH^+$). $\nu_{max}$ 3294, 2918, 2849, 2782, 1709, 1447, 1371, 1282, 1214, 1146, 1103, 734 $cm^{-1}$. Purity: UPLC (method I, 254 nm): $t_r$ = 4.310 min, 100.0% total area. $^1H$ NMR (400 MHz, $CDCl_3$) δ = 1.10 – 1.22 (m, 2H), 1.31 – 1.71 (m, 13H), 2.14 and 2.20 (2 × s, 3H), 2.33 – 2.46 (m, 2H), 3.28 – 3.71 (m, 8H), 6.17 and 6.78 (2 × s, 1H), 6.37 (s) and 6.91 – 7.21 (m, 4H), 7.54 – 7.74 (m, 1H), 9.58 and 10.10 (2 × s, 1H). $^{13}C$ NMR (101 MHz, $CDCl_3$) δ = 26.52, 28.54, 34.84, 35.03, 35.90, 37.55, 40.49, 42.38, 52.93, 55.84, 113.79, 113.99, 115.12, 118.27, 119.08, 119.39, 119.94, 120.09, 124.70, 125.27, 125.93, 130.00, 130.20, 131.65, 131.94, 150.68, 150.85, 161.53. A mixture of two conformers in a ratio of 70:30.

*S-(1-Methyl-1H-imidazol-2-yl) dimethylcarbamothioate* (**21b**)

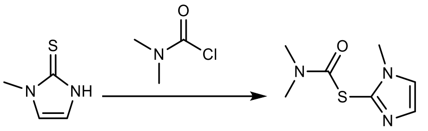

Methimazole (550 mg, 4.82 mmol), dimethylcarbamoyl chloride (1.2 eq., 532 µL), 1-methylimidazole (100 µL), DIPEA (1 mL) in MeCN (5 mL) were stirred for 8 h at 60 °C, and *S*-(1-methyl-1*H*-imidazol-2-yl) dimethylcarbamothioate isolated following *GP3*. Yield: 481 mg (2.60 mmol, 53.9%) of beige solid. ESI-HRMS: *m/z* = 186.0693 ($MH^+$); $C_7H_{12}N_3OS$ requires: *m/z* = 186.0696 ($MH^+$). $\nu_{max}$ 3164, 3118, 3090, 2938, 1699, 1452, 1375, 1291, 1258, 1221, 1148, 1133, 770, 730, 716, 669 $cm^{-1}$. Purity: UPLC (method II, 254 nm): $t_r$ = 0.637 min, 99.7% total area. $^1H$ NMR (400 MHz, $CDCl_3$) δ = 2.98 (s, 3H), 3.12 (s, 3H), 3.54 (s, 3H), 6.72 (d, *J*=2.5 Hz, 1H), 6.79 (d, *J*=2.5 Hz, 1H). $^{13}C$ NMR (101 MHz, $CDCl_3$) δ = 34.88, 37.12, 38.76, 114.78, 119.47, 151.48, 162.27.

*S-(4-Methyl-4H-1,2,4-triazol-3-yl) (3-(((2-cycloheptylethyl)(methyl)amino)methyl)-1H-indol-7-yl)(methyl)carbamothioate* (**22**)

Following *GP1*, *S*-(4-methyl-4*H*-1,2,4-triazol-3-yl) (1*H*-indol-7-yl)(methyl)carbamothioate **22a** was prepared from 4-methyl-4*H*-1,2,4-triazole-3-thiol and isolated by *GP3*. Yield: 196 mg (0.682 mmol, 68.2%) of beige solid. mp 193.5 °C (decomp.). ESI-HRMS: *m/z* = 288.0906 ($MH^+$); $C_{13}H_{14}N_5OS$ requires: *m/z* = 288.914 ($MH^+$). $\nu_{max}$ 3254, 2931, 1716, 1377, 1343, 1295, 1217, 1183, 1138, 1094, 1010, 849, 801, 726 $cm^{-1}$. Purity: UPLC (method I, 220 nm): $t_r$ = 3.937 min, 98.4% total area. $^1H$ NMR (400 MHz, DMSO-$d_6$) δ = 3.25 and 3.30 (2 × s, 3H), 3.43 and 3.44 and 3.56 (3 × s, 3H), 6.44 – 6.47 and 6.54 – 6.57 (2 × m, 1H), 6.83 and 7.09 (2 × t, *J*=7.7 Hz, 1H), 6.95 and 7.18 (2 × d, *J*=7.3 Hz, 1H), 7.38 (t, *J*=2.6 Hz) and 7.58 – 7.62 (m, 1H), 7.45 (d, *J*=7.8 Hz, 1H), 8.17 and 8.77 (2 × s, 1H), 11.49 (s, 1H). $^{13}C$ NMR (101 MHz, DMSO-$d_6$) δ = 31.23, 31.74, 37.69, 101.77, 118.05, 118.54, 120.46, 125.82, 126.19, 129.67, 131.01, 142.01, 142.67, 149.88, 166.43.

*S*-(4-Methyl-4*H*-1,2,4-triazol-3-yl) (3-(((2-cycloheptylethyl)(methyl)amino)methyl)-1*H*-indol-7-yl)(methyl)carbamothioate **22** was prepared following *GP2* from **22a** (115 mg, 0.400 mmol). Yield: 60 mg (0.132 mmol, 33.0%) of beige solid. mp 145.7–148.4 °C. ESI-HRMS: *m/z* = 455.2566 ($MH^+$); $C_{24}H_{35}N_6OS$ requires: *m/z* = 455.2588 ($MH^+$). $\nu_{max}$ 3366, 3120, 2920, 2850, 2769, 1713, 1388, 1358, 1286, 1210, 1097, 1053, 998, 809, 738, 629 $cm^{-1}$. Purity: UPLC (method I, 220 nm): $t_r$ = 4.427 min, 97.9% total area. $^1$H NMR (400 MHz, $CDCl_3$) δ = 1.11 – 1.21 (m, 2H), 1.33 – 1.70 (m, 13H), 2.17 (br s, 3H), 2.40 (br s, 2H), 3.21 – 3.70 (m, 8H), 6.97 – 7.21 (m, 3H), 7.35 and 7.95 (2 × s, 1H), 7.56 – 7.75 (m, 1H), 8.96 and 9.60 (2 × s, 1H). $^{13}$C NMR (101 MHz, $CDCl_3$) δ = 26.53, 28.54, 32.18, 32.59, 34.85, 35.89, 37.54, 38.92, 39.95, 42.41, 52.93, 55.94, 114.17, 114.41, 118.58, 119.58, 119.91, 124.76, 125.64, 130.07, 131.36, 131.94, 140.41, 142.34, 149.34, 150.30, 166.82, 167.43. A mixture of conformers.

*S-(1-Methyl-1H-tetrazol-5-yl) (3-(((2-cycloheptylethyl)(methyl)amino)methyl)-1H-indol-7-yl)(methyl)carbamothioate* (**23**)

Following *GP1, S*-(1-methyl-1*H*-tetrazol-5-yl) (1*H*-indol-7-yl)(methyl)carbamothioate **23a** was prepared from 1-methyl-1*H*-tetrazole-5-thiol and isolated by *GP3*. Yield: 146 mg (0.506 mmol, 50.6%) of beige solid. mp 180.5–184.0 °C. ESI-HRMS: *m/z* = 289.0873 ($MH^+$); $C_{12}H_{13}N_6OS$ requires: *m/z* = 289.0866 ($MH^+$). $\nu_{max}$ 3309, 2925, 1686, 1585, 1493, 1443, 1334, 1267, 1177, 1134, 1061, 891, 855, 800, 726, 649 $cm^{-1}$. Purity: UPLC (method II, 220 nm): $t_r$ = 2.503 min, 95.9% total area. $^1$H NMR (400 MHz, $CDCl_3$) δ = 3.30 (s, 3H), 3.98 (s, 3H), 6.61 (dd, *J*=2.0 Hz, 3.0 Hz, 1H), 7.13 – 7.19 (m, 2H), 7.23 (t, *J*=2.7 Hz, 1H), 7.74 (dd, *J*=1.1 Hz, 7.2 Hz, 1H), 9.97 (s, 1H). $^{13}$C NMR (101 MHz, $CDCl_3$) δ = 34.74, 38.12, 103.32, 120.09, 122.29, 123.08, 123.63, 126.33, 130.73, 132.54, 148.39, 161.88.

*S*-(1-Methyl-1*H*-tetrazol-5-yl) (3-(((2-cycloheptylethyl)(methyl)amino)methyl)-1*H*-indol-7-yl)(methyl)carbamothioate **23** was prepared following *GP2* from **23a** (74 mg, 0.257 mmol). Yield: 63 mg (0.138 mmol, 53.8%) of beige semisolid. ESI-HRMS: *m/z* = 456.2517 ($MH^+$); $C_{23}H_{34}N_7OS$ requires: *m/z* = 456.2540 ($MH^+$). $\nu_{max}$ 3331, 2919, 2850, 2789, 1694, 1447, 1341, 1267, 1098, 1033, 908, 801, 730, 648 $cm^{-1}$. Purity: UPLC (method II, 254 nm): $t_r$ = 2.767 min, 98.7% total area. $^1$H NMR (400 MHz, $CDCl_3$) δ = 1.12 – 1.21 (m, 2H), 1.31 – 1.69 (m, 13H), 2.22 (s, 3H), 2.41 – 2.46 (m, 2H), 3.32 (s, 3H), 3.64 – 3.73 (m, 2H), 4.01 (s, 3H), 7.14 – 7.20 (m, 3H), 7.82 – 7.89 (m, 1H), 9.71 (s, 1H). $^{13}$C NMR (101 MHz, $CDCl_3$) δ = 26.56, 28.57, 34.77, 34.86, 35.89, 37.54, 38.19, 42.47, 53.01, 55.97, 114.80, 119.84, 122.19, 122.31, 123.50, 125.40, 130.81, 133.04, 148.39, 162.02.

*Se-Ethyl (3-(((2-cycloheptylethyl)(methyl)amino)methyl)-1H-indol-7-yl)(methyl)carbamoselenoate* (**24**)

Following *GP1, Se*-ethyl (1*H*-indol-7-yl)(methyl)carbamoselenoate **24a** was prepared from magnesium ethaneselenolate (3.0 eq., prepared by stirring elemental black selenium and 1.1 eq. 1 M ethylmagnesium bromide in THF at rt for 18 h) and isolated by column chromatography on silica (PE/EtOAc = 10:1). Yield: 40 mg (0.142 mmol, 14.2%) of yellowish solid. ESI-HRMS: *m/z* = 283.0352 ($MH^+$); $C_{12}H_{15}N_2OSe$ requires: *m/z* = 283.0344 ($MH^+$). $\nu_{max}$ 3297, 3102, 2962, 2921, 2865, 1639, 1583, 1494, 1446, 1332, 1276, 1239, 1183, 1128, 1066, 1047, 950, 893, 855, 798, 730, 674, 623, 594 $cm^{-1}$. Purity: UPLC (method II, 220 nm): $t_r$ = 3.047 min, 99.5% total area. $^1$H NMR (400 MHz, $CDCl_3$) δ = 1.33 (t, *J*=7.5 Hz, 3H), 2.69 (dq, *J*=7.5 Hz, 14.9 Hz, 1H), 2.87 (dq, *J*=7.4 Hz, 14.9 Hz, 1H), 3.48 (s, 3H), 6.64 (dd, *J*=2.1 Hz, 3.0 Hz, 1H), 7.12 – 7.18 (m, 2H), 7.34 (t, *J*=2.6 Hz, 1H), 7.70 – 7.75 (m, 1H), 9.79 (s, 1H). $^{13}$C NMR (101

MHz, $CDCl_3$) δ = 15.70, 21.02, 37.46, 103.28, 120.06, 122.10, 122.46, 125.41, 125.52, 130.32, 133.16, 168.17. $^{77}$Se NMR (95 MHz, $CDCl_3$) δ = 440.

*Se*-Ethyl (3-(((2-cycloheptylethyl)(methyl)amino)methyl)-1*H*-indol-7-yl)(methyl)carbamoselenoate **24** was prepared following *GP2* from **24a** (40 mg, 0.142 mmol). Yield: 39 mg (0.087 mmol, 61.2%) of beige semisolid. ESI-HRMS: *m/z* = 450.1998 [M + H]+; $C_{23}H_{36}N_3OSe$ requires: *m/z* = 450.2018 ($MH^+$). $\nu_{max}$ 3299, 2919, 2850, 2782, 1642 1583, 1446, 1339, 1278, 1233, 1192, 1099, 799, 736 $cm^{-1}$. Purity: UPLC (method II, 254 nm): $t_r$ = 3.430 min, 99.1% total area. $^1$H NMR (400 MHz, $CDCl_3$) δ = 1.12 – 1.21 (m, 2H), 1.31 (t, *J*=7.4 Hz, 3H), 1.34 – 1.71 (m, 13H), 2.24 (s, 3H), 2.42 – 2.47 (m, 2H), 2.69 (dq, *J*=7.4 Hz, 14.7 Hz, 1H), 2.84 (dq, *J*=6.2 Hz, 6.8 Hz, 13.7 Hz, 1H), 3.44 (s, 3H), 3.63 – 3.76 (m, 2H), 7.08 – 7.15 (m, 2H), 7.18 – 7.23 (m, 1H), 7.76 – 7.81 (m, 1H), 9.39 (s, 1H). $^{13}$C NMR (101 MHz, $CDCl_3$) δ = 15.76, 20.99, 26.61, 28.62, 34.90, 35.96, 37.44, 37.59, 42.58, 53.09, 55.94, 114.60, 119.77, 121.01, 122.48, 124.53, 125.38, 130.36, 133.55, 167.95. $^{77}$Se NMR (95 MHz, $CDCl_3$) δ = 441.

*Se-Phenyl (3-(((2-cycloheptylethyl)(methyl)amino)methyl)-1H-indol-7-yl)(methyl)carbamoselenoate* (**25**)

Following *GP1*, *Se*-phenyl (1*H*-indol-7-yl)(methyl)carbamoselenoate **25a** was prepared from selenophenol and isolated by column chromatography on silica (PE/EtOAc = 4:1). Yield: 138 mg (0.419 mmol, 41.9%) of beige solid. ESI-HRMS: *m/z* = 331.0350 ($MH^+$); $C_{16}H_{15}N_2OSe$ requires: *m/z* = 331.0344 ($MH^+$). $\nu_{max}$ 3321, 3048, 1665, 1585, 1331, 1274, 1129, 798, 729 $cm^{-1}$. $^1$H NMR (400 MHz, $CDCl_3$) δ = 3.36 (s, 3H), 6.57 (dd, *J*=2.0 Hz, 3.1 Hz, 1H), 7.01 (t, *J*=2.7 Hz, 1H), 7.13 – 7.32 (m, 5H), 7.54 – 7.58 (m, 2H), 7.73 (d, *J*=7.6 Hz, 1H), 9.54 (s, 1H). $^{13}$C NMR (101 MHz, $CDCl_3$) δ = 37.79, 102.81, 119.90, 122.31, 122.40, 124.94, 125.93, 127.98, 128.86, 129.06, 130.41, 132.95, 136.62 (t, $^3J$=5.1 Hz, $^{13}$C–$^{77}$Se), 167.25.

*Se*-Phenyl (3-(((2-cycloheptylethyl)(methyl)amino)methyl)-1*H*-indol-7-yl)(methyl)carbamoselenoate **25** was prepared following *GP2* from **25a** (107 mg, 0.325 mmol). Yield: 75 mg (0.151 mmol, 46.5%) of beige semisolid. ESI-HRMS: *m/z* = 498.2031 ($MH^+$); $C_{27}H_{36}N_3OSe$ requires: *m/z* = 498.2018 ($MH^+$). $\nu_{max}$ 3321, 1665, 1585, 1332, 1274, 1129, 1022, 947, 850, 798, 729, 689 $cm^{-1}$. Purity: UPLC (method I, 220 nm): $t_r$ = 4.840 min, 99.2% total area. $^1$H NMR (400 MHz, $CDCl_3$) δ = 1.13 – 1.23 (m, 2H), 1.35 – 1.72 (m, 13H), 2.23 (s, 3H), 2.41 – 2.48 (m, 2H), 3.31 (s, 3H), 3.60 – 3.72 (m, 2H), 6.85 (s, 1H), 7.09 – 7.27 (m, 5H), 7.53 (d, *J*=7.2 Hz, 2H), 7.80 (d, *J*=7.5 Hz, 1H), 9.51 (s, 1H). $^{13}$C NMR (101 MHz, $CDCl_3$) δ = 26.59, 28.60, 34.89, 36.00, 37.54, 37.75, 42.50, 53.11, 55.93, 113.86, 119.45, 121.26, 122.29, 124.69, 125.19, 128.08, 128.69, 128.98, 130.40, 133.36, 136.58, 167.19. $^{77}$Se NMR (95 MHz, $CDCl_3$) δ = 538.

*N-(3-(((2-Cycloheptylethyl)(methyl)amino)methyl)-1H-indol-7-yl)-N-methyl-1H-imidazole-1-carboxamide* (**26**)

*N*-(3-(((2-Cycloheptylethyl)(methyl)amino)methyl)-1*H*-indol-7-yl)-*N*-methyl-1*H*-imidazole-1-carboxamide **26** was prepared following *GP2* from *N*-(1*H*-indol-7-yl)-*N*-methyl-1*H*-

imidazole-1-carboxamide[163] **26a** (69 mg, 0.287 mmol). Yield: 67 mg (0.164 mmol, 57.3%) of beige solid. mp 140.5–143.0 °C. ESI-HRMS: *m/z* = 408.2752 ($MH^+$); $C_{24}H_{34}N_5O$ requires: *m/z* = 408.2758 ($MH^+$). $\nu_{max}$ 3119, 3027, 2917, 2849, 2795, 2761, 1692, 1585, 1459, 1416, 1382, 1346, 1293, 1203, 1119, 1103, 1048, 1000, 827, 743, 665, 652 $cm^{-1}$. Purity: UPLC (method I, 254 nm): $t_r$ = 3.823 min, 98.0% total area. $^1H$ NMR (400 MHz, $CDCl_3$) δ = 1.12 – 1.21 (m, 2H), 1.32 – 1.69 (m, 13H), 2.19 (s, 3H), 2.39 – 2.44 (m, 2H), 3.40 (s, 3H), 3.64 (s, 2H), 6.61 – 6.65 (m, 2H), 6.89 – 6.92 (m, 1H), 7.05 (t, *J*=7.7 Hz, 1H), 7.13 (d, *J*=2.0 Hz, 1H), 7.54 – 7.56 (m, 1H), 7.73 (d, *J*=7.9 Hz, 1H), 9.97 (s, 1H). $^{13}C$ NMR (101 MHz, $CDCl_3$) δ = 26.55, 28.56, 34.85, 35.87, 37.53, 39.43, 42.46, 52.96, 56.02, 114.92, 118.36, 119.70, 120.17, 120.76, 125.16, 126.23, 128.87, 130.66, 132.02, 137.75, 150.55.

*N-(3-(((2-Cycloheptylethyl)(methyl)amino)methyl)-1H-indol-7-yl)-N-methyl-4-nitro-1H-imidazole-1-carboxamide* (**27**)

Following *GP1*, *N*-(1*H*-indol-7-yl)-*N*-methyl-4-nitro-1*H*-imidazole-1-carboxamide **27a** was prepared from 4(5)-nitro-1*H*-imidazole and isolated by column chromatography on silica (1. PE/EtOAc = 3:1; 2. PE/EtOAc = 1:1). Yield: 98 mg (0.344 mmol, 34.4%) of greenish solid. mp 166.6–167.7 °C. ESI-HRMS: *m/z* = 286.0927 ($MH^+$); $C_{13}H_{12}N_5O_3$ requires: *m/z* = 286.0935 ($MH^+$). $\nu_{max}$ 3232, 3145, 1698, 1557, 1514, 1484, 1376, 1279, 1145, 1018, 870, 821, 725 $cm^{-1}$. Purity: UPLC (method II, 220 nm): $t_r$ = 2.383 min, 97.2% total area. $^1H$ NMR (400 MHz, $CDCl_3$) δ = 3.54 (s, 3H), 6.65 (dd, *J*=2.0 Hz, 3.2 Hz, 1H), 6.93 – 6.98 (m, 1H), 7.11 (t, *J*=7.7 Hz, 1H), 7.24 (d, *J*=1.5 Hz, 1H), 7.27 – 7.29 (m, 1H), 7.58 (d, *J*=1.6 Hz, 1H), 7.70 (d, *J*=8.0 Hz, 1H), 8.90 (s, 1H). $^{13}C$ NMR (101 MHz, $CDCl_3$) δ = 39.80, 104.51, 118.48, 119.85, 121.04, 122.86, 125.07, 126.34, 131.02, 131.09, 136.11, 147.38, 148.50.

*N*-(3-(((2-Cycloheptylethyl)(methyl)amino)methyl)-1*H*-indol-7-yl)-*N*-methyl-4-nitro-1*H*-imidazole-1-carboxamide **27** was prepared following *GP2* from **27a** (68 mg, 0.238 mmol). Yield: 43 mg (0.0950 mmol, 39.9%) of yellowish semisolid. ESI-HRMS: *m/z* = 453.2586 ($MH^+$); $C_{24}H_{33}N_6O_3$ requires: *m/z* = 453.2609 ($MH^+$). $\nu_{max}$ 3135, 2919, 2850, 2781, 1721, 1583, 1554, 1514, 1488, 1456, 1371, 1286, 1262, 1233, 1097, 821, 741 $cm^{-1}$. Purity: UPLC (method I, 254 nm): $t_r$ = 4.330 min, 99.7% total area. $^1H$ NMR (400 MHz, $CDCl_3$) δ = 1.11 – 1.21 (m, 2H), 1.33 – 1.70 (m, 13H), 2.18 (s, 3H), 2.40 – 2.45 (m, 2H), 3.52 (s, 3H), 3.63 (br s, 1H), 3.67 (br s, 1H), 6.94 (d, *J*=7.3 Hz, 1H), 7.10 (t, *J*=7.8 Hz, 1H), 7.18 – 7.20 (m, 1H), 7.25 (d, *J*=1.4 Hz, 1H), 7.56 (d, *J*=1.4 Hz, 1H), 7.79 (d, *J*=7.8 Hz, 1H), 8.89 (s, 1H). $^{13}C$ NMR (101 MHz, $CDCl_3$) δ = 26.58, 28.58, 34.88, 35.82, 37.59, 39.78, 42.35, 52.87, 56.11, 115.55, 118.49, 119.77, 120.60, 121.78, 124.97, 125.48, 131.05, 131.55, 136.15, 147.32, 148.51.

*N-(3-(((2-Cycloheptylethyl)(methyl)amino)methyl)-1H-indol-7-yl)-N-methyl-1H-1,2,4-triazole-1-carboxamide* (**28**)

Following *GP1*, *N*-(1*H*-indol-7-yl)-*N*-methyl-1*H*-1,2,4-triazole-1-carboxamide **28a** was prepared from 1*H*-1,2,4-triazole and isolated by column chromatography on silica (1. PE/EtOAc = 2:1; 2. PE/EtOAc = 1:1). Yield: 70 mg (0.290 mmol, 29.0%) of beige solid. mp = 192.2–193.5 °C. ESI-HRMS: *m/z* = 242.1030 ($MH^+$); $C_{12}H_{12}N_5O$ requires: *m/z* = 242.1036

(MH$^+$). $\nu_{max}$ 3187, 3125, 2928, 1705, 1587, 1516, 1424, 1397, 1339, 1282, 999, 798, 735 cm$^{-1}$. Purity: UPLC (method I, 254 nm): $t_r$ = 3.170 min, 99.0% total area. $^1$H NMR (400 MHz, $CDCl_3$) δ = 3.57 (s, 3H), 6.60 (dd, *J*=2.0 Hz, 3.2 Hz, 1H), 6.95 (d, *J*=6.8 Hz, 1H), 7.06 (t, *J*=7.7 Hz, 1H), 7.24 (t, *J*=2.8 Hz, 1H), 7.63 (d, *J*=7.9 Hz, 2H), 8.31 (s, 1H), 9.21 (s, 1H). $^{13}$C NMR (101 MHz, $CDCl_3$) δ = 39.94, 103.68, 119.40, 120.28, 121.53, 125.67, 126.84, 130.45, 131.73, 145.46, 149.67, 151.83.

*N*-(3-(((2-Cycloheptylethyl)(methyl)amino)methyl)-1*H*-indol-7-yl)-*N*-methyl-1*H*-1,2,4-triazole-1-carboxamide **28** was prepared following *GP2* from **28a** (51 mg, 0.211 mmol). Yield: 40 mg (0.0979 mmol, 46.4%) of yellowish semisolid. ESI-HRMS: *m/z* = 409.2698 (MH$^+$); $C_{23}H_{33}N_6O$ requires: *m/z* = 409.2710 (MH$^+$). $\nu_{max}$ 3197, 2918, 2850, 2790, 1719, 1702, 1624, 1584, 1456, 1423, 1397, 1285, 1185, 1127, 1050, 993, 802, 738 cm$^{-1}$. Purity: UPLC (method I, 254 nm): $t_r$ = 4.210 min, 97.2% total area. $^1$H NMR (400 MHz, $CDCl_3$) δ = 1.12 – 1.21 (m, 2H), 1.33 – 1.69 (m, 13H), 2.19 (s, 3H), 2.40 – 2.45 (m, 2H), 3.53 (s, 3H), 3.65 (s, 2H), 6.94 (d, *J*=6.8 Hz, 1H), 7.05 (t, *J*=7.7 Hz, 1H), 7.13 (d, *J*=1.7 Hz, 1H), 7.60 – 7.65 (m, 1H), 7.71 (d, *J*=7.9 Hz, 1H), 8.19 (s, 1H), 9.44 (s, 1H). $^{13}$C NMR (101 MHz, $CDCl_3$) δ = 26.57, 28.58, 34.87, 35.87, 37.57, 39.86, 42.47, 52.97, 56.00, 114.72, 119.32, 119.85, 120.50, 124.97, 126.47, 130.54, 132.10, 145.34, 149.54, 151.65.

*N-(3-(((2-Cycloheptylethyl)(methyl)amino)methyl)-1H-indol-7-yl)-N-methyl-2H-1,2,3-triazole-2-carboxamide* (**29**)

Following *GP1*, *N*-(1*H*-indol-7-yl)-*N*-methyl-2*H*-1,2,3-triazole-2-carboxamide **29a** was prepared from 1,2,3-triazole and isolated by column chromatography on silica (PE/EtOAc = 2:1). Yield: 90 mg (0.373 mmol, 37.3%) of beige solid. ESI-HRMS: *m/z* = 242.1041 (MH$^+$); $C_{12}H_{12}N_5O$ requires: *m/z* = 242.1036 (MH$^+$). $\nu_{max}$ 3262, 3133, 3122, 2926, 1707, 1584, 1497, 1421, 1386, 1338, 1266, 1180, 1130, 1060, 961, 937, 854, 799, 726 cm$^{-1}$. Purity: UPLC (method I, 254 nm): $t_r$ = 3.727 min, 96.6% total area. $^1$H NMR (400 MHz, $CDCl_3$) δ = 3.61 (s, 3H), 6.57 (dd, *J*=2.0 Hz, 3.2 Hz, 1H), 6.90 (d, *J*=6.5 Hz, 1H), 6.99 (t, *J*=7.7 Hz, 1H), 7.23 (t, *J*=2.7 Hz, 1H), 7.48 (s, 2H), 7.55 (d, *J*=7.9 Hz, 1H), 8.64 (s, 1H). $^{13}$C NMR (101 MHz, $CDCl_3$) δ = 39.77, 103.55, 119.40, 120.10, 120.82, 125.28, 127.59, 130.07, 131.73, 136.05, 150.40.

*N*-(3-(((2-Cycloheptylethyl)(methyl)amino)methyl)-1*H*-indol-7-yl)-*N*-methyl-2*H*-1,2,3-triazole-2-carboxamide **29** was prepared following *GP2* from **29a** (62 mg, 0.257 mmol). Yield: 50 mg (0.122 mmol, 47.6%) of beige semisolid. ESI-HRMS: *m/z* = 409.2719 (MH$^+$); $C_{23}H_{33}N_6O$ requires: *m/z* = 409.2710 (MH$^+$). z$\nu_{max}$ 3307, 2917, 2848, 2794, 2764, 1715, 1584, 1451, 1423, 1391, 1268, 1099, 960, 801 cm$^{-1}$. Purity: UPLC (method I, 220 nm): $t_r$ = 4.340 min, 99.4% total area. $^1$H NMR (400 MHz, $CDCl_3$) δ = 1.11 – 1.20 (m, 2H), 1.33 – 1.69 (m, 13H), 2.20 (s, 3H), 2.38 – 2.44 (m, 2H), 3.58 (s, 3H), 3.64 (s, 2H), 6.88 (d, *J*=6.7 Hz, 1H), 6.98 (t, *J*=7.7 Hz, 1H), 7.13 (d, *J*=2.1 Hz, 1H), 7.43 (s, 2H), 7.62 (d, *J*=7.9 Hz, 1H), 8.98 (s, 1H). $^{13}$C NMR (101 MHz, $CDCl_3$) δ = 26.57, 28.59, 34.87, 35.85, 37.59, 39.83, 42.47, 52.96, 55.93, 114.28, 119.25, 119.55, 119.60, 124.73, 127.40, 130.14, 132.15, 135.91, 150.32.

*N-(3-(((2-Cycloheptylethyl)(methyl)amino)methyl)-1H-indol-7-yl)-N-methyl-1H-tetrazole-1-carboxamide* (**30**)

Following *GP1*, *N*-(1*H*-indol-7-yl)-*N*-methyl-1*H*-tetrazole-1-carboxamide **30a** was prepared from 1*H*-tetrazole and isolated by column chromatography on silica (1. PE/EtOAc = 5:1; 2. PE/EtOAc = 3:1; 3. PE/EtOAc = 1:1). Yield: 153 mg (0.632 mmol, 63.2%) of greenish solid. mp 142.1 °C (decomp.). ESI-HRMS: *m/z* = 241.0843 ($(M–H)^–$); $C_{11}H_9N_6O$ requires: *m/z* = 241.0843 ($(M–H)^–$). $\nu_{max}$ 3250, 3153, 3121, 2946, 1735, 1585, 1498, 1374, 1336, 1264, 1189, 1158, 1081, 997, 877, 799, 738 $cm^{–1}$. Purity: UPLC (method I, 220 nm): $t_r$ = 3.533 min, 97.0% total area. $^1H$ NMR (400 MHz, $CDCl_3$) δ = 3.58 (s, 3H), 6.61 (dd, *J*=2.0 Hz, 3.0 Hz, 1H), 6.89 – 6.95 (m, 1H), 7.02 – 7.08 (m, 1H), 7.25 (t, *J*=2.9 Hz, 1H), 7.65 (d, *J*=7.9 Hz, 1H), 8.57 (s, 1H), 9.09 (s, 1H). $^{13}C$ NMR (101 MHz, $CDCl_3$) δ = 39.72, 104.04, 119.65, 120.44, 122.51, 125.10, 126.08, 130.84, 131.52, 142.57, 146.99.

*N*-(3-(((2-Cycloheptylethyl)(methyl)amino)methyl)-1*H*-indol-7-yl)-*N*-methyl-1*H*-tetrazole-1-carboxamide **30** was prepared following *GP2* from *N*-(1*H*-indol-7-yl)-*N*-methyl-1*H*-tetrazole-1-carboxamide **30a** (88 mg, 0.364 mmol). Yield: 84 mg (0.205 mmol, 56.4%) of brownish semisolid. ESI-HRMS: *m/z* = 410.2651 ($MH^+$); $C_{22}H_{32}N_7O$ requires: *m/z* = 410.2663 ($MH^+$). $\nu_{max}$ 3216, 2920, 2852, 1734, 1614, 1485, 1441, 1390, 1343, 1101, 731 $cm^{–1}$. Purity: UPLC (method I, 254 nm): $t_r$ = 4.440 min, 95.4% total area. $^1H$ NMR (400 MHz, $CDCl_3$) δ = 1.11 – 1.20 (m, 2H), 1.34 – 1.69 (m, 13H), 2.19 (s, 3H), 2.40 – 2.48 (m, 2H), 3.51 (s, 3H), 3.58 – 3.71 (m, 2H), 6.90 (d, *J*=6.2 Hz, 1H), 7.01 (t, *J*=7.3 Hz, 1H), 7.13 (s, 1H), 7.71 (d, *J*=8.1 Hz, 1H), 8.46 (s, 1H), 10.02 (s, 1H). $^{13}C$ NMR (101 MHz, $CDCl_3$) δ = 26.50, 28.52, 34.79, 35.57, 37.50, 39.53, 42.13, 52.74, 55.87, 114.15, 119.60, 119.92, 121.23, 124.78, 125.77, 130.73, 131.89, 142.44, 146.78.

*N-(3-(((2-Cycloheptylethyl)(methyl)amino)methyl)-1H-indol-7-yl)-N-methyl-1H-benzotriazole-1-carboxamide* (**31**)

Following *GP1*, *N*-(1*H*-indol-7-yl)-*N*-methyl-1*H*-benzotriazole-1-carboxamide **31a** was prepared from 1*H*-benzotriazole sodium salt and isolated by column chromatography on silica (1. PE/EtOAc = 5:1; 2. PE/EtOAc = 3:1). Yield: 275 mg (0.944 mmol, 94.4%) of grayish solid. mp 180.2–184.5 °C. ESI-HRMS: *m/z* = 292.1185 ($MH^+$); $C_{16}H_{14}N_5O$ requires: *m/z* = 292.1193 ($MH^+$). $\nu_{max}$ 3339, 1698, 1585, 1487, 1420, 1301, 1186, 1082, 1043, 964, 801, 731 $cm^{–1}$. Purity: UPLC (method I, 220 nm): $t_r$ = 4.443 min, 97.0% total area. $^1H$ NMR (400 MHz, $CDCl_3$) δ = 3.49 (s, 3H), 6.51 (dd, *J*=2.0 Hz, 3.0 Hz, 1H), 6.74 – 6.80 (m, 1H), 6.84 – 6.89 (m, 1H), 7.03 (t, *J*=2.6 Hz, 1H), 7.23 – 7.28 (m, 1H), 7.41 (t, *J*=7.6 Hz, 1H), 7.51 (d, *J*=7.8 Hz, 1H), 7.84 (d, *J*=8.3 Hz, 1H), 7.93 (d, *J*=8.3 Hz, 1H), 9.57 (s, 1H). $^{13}C$ NMR (101 MHz, $CDCl_3$) δ = 39.30, 103.05, 113.09, 118.80, 119.52, 119.60, 120.76, 125.14, 125.44, 127.70, 129.20, 130.20, 131.95, 132.64, 144.75, 150.60.

*N*-(3-(((2-Cycloheptylethyl)(methyl)amino)methyl)-1*H*-indol-7-yl)-*N*-methyl-1*H*-benzotriazole-1-carboxamide **31** was prepared following *GP2* from **31a** (114 mg, 0.390 mmol). Yield: 81 mg (0.177 mmol, 45.3%) of greenish semisolid. ESI-HRMS: *m/z* = 459.2852 ($MH^+$);

$C_{27}H_{35}N_6O$ requires: *m/z* = 459.2867 ($MH^+$). $\nu_{max}$ 3303, 2917, 2849, 2763, 1673, 1449, 1394, 1163, 1024, 740 $cm^{-1}$. Purity: UPLC (method I, 254 nm): $t_r$ = 4.610 min, 98.2% total area. $^1H$ NMR (400 MHz, $CDCl_3$) δ = 1.13 – 1.22 (m, 2H), 1.33 – 1.71 (m, 13H), 2.21 (s, 3H), 2.42 – 2.46 (m, 2H), 3.60 (s, 3H), 3.66 (s, 2H), 6.82 – 6.86 (m, 1H), 6.93 (t, *J*=7.4 Hz, 1H), 7.12 (d, *J*=2.1 Hz, 1H), 7.34 – 7.39 (m, 1H), 7.53 (ddd, *J*=0.9 Hz, 7.1 Hz, 8.2 Hz, 1H), 7.64 (d, *J*=7.9 Hz, 1H), 7.92 (d, *J*=8.3 Hz, 1H), 8.00 (d, *J*=8.3 Hz, 1H), 8.89 (s, 1H). $^{13}C$ NMR (101 MHz, $CDCl_3$) δ = 26.57, 28.60, 34.89, 35.90, 37.61, 39.68, 42.47, 53.00, 56.03, 113.27, 114.70, 119.15, 119.67, 119.85, 119.93, 124.60, 125.26, 127.75, 129.40, 130.42, 132.56, 132.82, 145.08, 150.96.

*2,2,2-Trifluoroethyl (3-(((2-cycloheptylethyl)(methyl)amino)methyl)-1H-indol-7-yl)carbamate* (**32**)

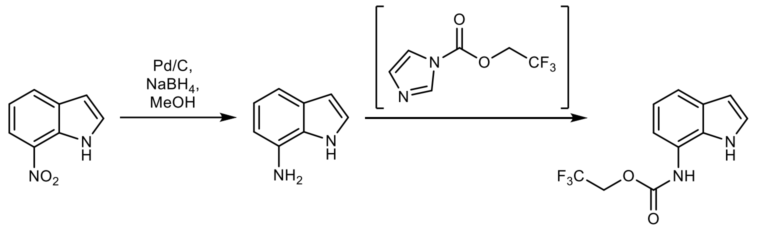

7-Nitro-1*H*-indole (524 mg, 3.23 mmol) and 10% Pd/C (50 mg) in MeOH (20 mL) was cooled in an icebath, and sodium borohydride was added in 50 mg portions, waiting in between for the foaming to subside, until the yellow colour disappeared and conversion was complete (by TLC). The suspension was filtered through Celite, the solvent removed *in vacuo*, residue diluted with water (50 mL), 50% sodium hydroxide$_{(aq)}$ (5 mL) was added, and 7-aminoindole extracted with $Et_2O$ (2 × 30 mL). The organic phase was dried with sodium sulfate, the solvent removed *in vacuo*.

CDI (1000 mg, 1.05 eq.) and 2,2,2-trifluoroethanol (447 µL, 5.92 mmol) in DCM (15 mL) were stirred at rt for 2 h, then crude 7-aminoindole (229 mg, 1.73 mmol) was added, and stirring continued at rt for 18 h. The solvent was removed *in vacuo* and 2,2,2-trifluoroethyl (1*H*-indol-7-yl)carbamate **32a** isolated following *GP3*. Yield: 120 mg (0.465 mmol, 26.9%) of white solid. ESI-HRMS: *m/z* = 259.0687 ($MH^+$); $C_{11}H_{10}O_2N_2F_3$ requires: *m/z* = 259.0689 ($MH^+$). $\nu_{max}$ 3401, 3272, 1713, 1538, 1434, 1286, 1235, 1167, 1115, 995, 957, 777, 722 $cm^{-1}$. Purity: UPLC (method II, 220 nm): $t_r$ = 2.737 min, 99.9% total area. $^1H$ NMR (400 MHz, $CDCl_3$) δ = 4.62 (q, *J*=8.4 Hz, 2H), 6.57 (dd, *J*=2.1 Hz, 3.1 Hz, 1H), 6.78 (d, *J*=7.4 Hz, 1H), 7.04 (t, *J*=7.7 Hz, 2H), 7.22 (t, *J*=2.8 Hz, 1H), 7.51 (d, *J*=7.9 Hz, 1H), 9.43 (br s, 1H). $^{13}C$ NMR (101 MHz, $CDCl_3$) δ = 61.65 (q, *J*=36.8 Hz), 102.97, 113.60, 118.53, 119.75, 120.98, 123.05 (q, *J*=277.6 Hz), 125.17, 127.95, 130.95, 152.56. $^{19}F$ NMR (376 MHz, $CDCl_3$) δ = –74.04.

2,2,2-Trifluoroethyl (3-(((2-cycloheptylethyl)(methyl)amino)methyl)-1*H*-indol-7-yl)carbamate **32** was prepared from **32a** (64 mg, 0.248 mmol) following *GP2*. Yield: 44 mg (0.103 mmol, 41.7%) of beige semisolid. ESI-HRMS: *m/z* = 426.2354 ($MH^+$); $C_{22}H_{31}O_2N_3F_3$ requires: *m/z* = 426.2363 ($MH^+$). $\nu_{max}$ 3378, 2920, 2852, 2796, 1723, 1633, 1585, 1530, 1443, 1278, 1164, 1101, 956, 735 $cm^{-1}$. Purity: UPLC (method II, 254 nm): $t_r$ = 3.110 min, 98.7% total area. $^1H$ NMR (400 MHz, $CDCl_3$) δ = 1.12 – 1.21 (m, 2H), 1.33 – 1.69 (m, 13H), 2.21 (s, 3H), 2.40 – 2.45 (m, 2H), 3.67 (s, 2H), 4.60 (q, *J*=8.4 Hz, 2H), 6.80 (d, *J*=7.4 Hz, 1H), 7.02 (t,

$J$=7.7 Hz, 1H), 7.08 – 7.10 (m, 1H), 7.39 (s, 1H), 7.56 (d, $J$=7.9 Hz, 1H), 9.27 (s, 1H). $^{13}$C NMR (101 MHz, $CDCl_3$) δ = 26.60, 28.64, 34.93, 35.80, 37.79, 42.40, 52.84, 55.95, 61.56 (q, $J$=36.7 Hz), 113.61, 113.75, 117.18, 119.45, 121.00, 123.11 (q, $J$=276.6 Hz), 124.49, 128.48, 130.86, 152.58. $^{19}$F NMR (376 MHz, $CDCl_3$) δ = –74.02.

*N-Methyl-1H-imidazole-1-carboxamide* (**33a**)

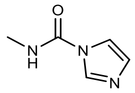

Prepared according to literature procedure.[172] ESI-HRMS: $m/z$ = 126.0663 ($MH^+$); $C_5H_8ON_3$ requires: $m/z$ = 126.0662 ($MH^+$). $\nu_{max}$ 3202, 3145, 3111, 3040, 2945, 1713, 1552, 1480, 1421, 1363, 1331, 1290, 1251, 1108, 1070, 1045, 910, 837, 806, 720 cm$^{-1}$. Purity: UPLC (method II, 220 nm): $t_r$ = 0.380 min, 99.96% total area. $^1$H NMR (400 MHz, DMSO-$d_6$) δ = 2.80 and 2.81 (2 × s, 3H), 7.02 (dd, $J$=0.9 Hz, 1.5, 1H), 7.63 (t, $J$=1.4 Hz, 1H), 8.20 (t, $J$=1.1 Hz, 1H), 8.45 (br s, 1H). $^{13}$C NMR (101 MHz, DMSO-$d_6$) δ = 26.78, 116.50, 129.56, 135.85, 149.26.

*N-(3-(((2-Cycloheptylethyl)(methyl)amino)methyl)-1H-indol-7-yl)-1H-imidazole-1-carboxamide* (**33**)

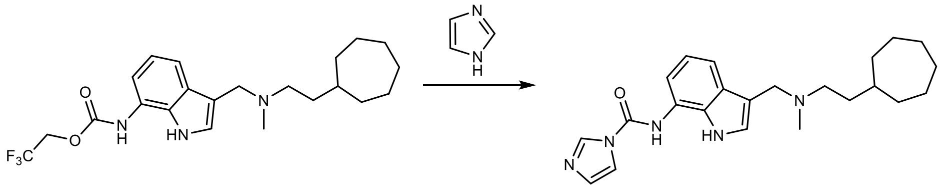

**32** (60 mg, 0.14 mmol) and imidazole (500 mg) in THF (5 mL) were heated in a pressure tube at 80 °C for 5 days. Volatiles were removed *in vacuo* and *N*-(3-(((2-cycloheptylethyl)(methyl)amino)methyl)-1*H*-indol-7-yl)-1*H*-imidazole-1-carboxamide **33** isolated by *GP3*. Yield: 11 mg (0.0280 mmol, 19.8%) of beige semisolid. ESI-HRMS: $m/z$ = 392.2457 ($(M–H)^-$); $C_{23}H_{30}ON_5$ requires: $m/z$ = 392.2456 ($(M–H)^-$). $\nu_{max}$ 3283, 2919, 2852, 1633, 1504, 1433, 1353, 1303, 1268, 1204, 1076, 796, 755, 735, 721, 626 cm$^{-1}$. Purity: UPLC (method II, 220 nm): $t_r$ = 3.007 min, 99.8% total area. $^1$H NMR (400 MHz, $CDCl_3$) δ = 1.20 – 1.30 (m, 2H), 1.38 – 1.78 (m, 13H), 3.05 (s, 3H), 3.37 – 3.43 (m, 2H), 5.26 (s, 2H), 6.68 – 6.72 (m, 2H), 6.92 – 6.94 (m, 1H), 6.97 (t, $J$=7.7 Hz, 1H), 7.00 – 7.02 (m, 1H), 7.13 – 7.18 (m, 2H), 7.54 (s, 1H), 10.27 (s, 1H). $^{13}$C NMR (101 MHz, $CDCl_3$) δ = 26.55, 28.53, 34.72, 35.15, 36.11, 37.04, 42.89, 48.08, 110.35, 113.13, 114.28, 119.20, 120.09, 124.52, 124.53, 128.99, 129.08, 129.30, 137.14, 155.51.

**P1**

Ethyl undec-10-yn-1-ylphosphonofluoridate was prepared as described in literature.[114] ESI-HRMS: $m/z$ = 263.1569 ($MH^+$); $C_{13}H_{25}O_2FP$ requires: $m/z$ = 263.1571 ($MH^+$). $^1$H NMR (400 MHz, $CDCl_3$) δ = 1.25 – 1.43 (m, 13H), 1.47 – 1.55 (m, 2H), 1.59 – 1.71 (m, 2H), 1.82 – 1.92 (m, 2H), 1.93 (t, $J$=2.7 Hz, 1H), 2.17 (td, $J$=2.6 Hz, 7.1 Hz, 2H), 4.20 – 4.31 (m, 2H). $^{13}$C NMR (101 MHz, $CDCl_3$) δ = 16.50 (d, $J$=5.7 Hz), 18.52, 22.03 (d, $J$=5.5 Hz), 24.45 (dd, $J$=22.3 Hz, 143.2 Hz), 28.57, 28.79, 29.04, 29.12, 29.27, 30.32, 30.49, 63.16 (d, $J$=7.2 Hz), 68.24, 84.85.

$^{19}F$ NMR (376 MHz, $CDCl_3$) δ = –64.70 (d, *J*=1069.3 Hz). $^{31}P$ NMR (162 MHz, $CDCl_3$) δ = 31.91 (d, *J*=1069.4 Hz).

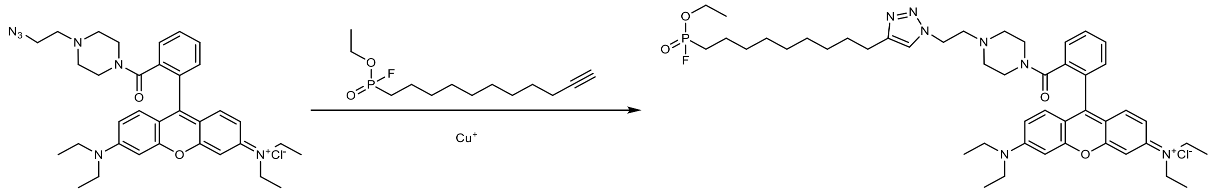

*N*-(6-(Diethylamino)-9-(2-(4-(2-(4-(9-(ethoxyfluorophosphoryl)nonyl)-1*H*-1,2,3-triazol-1-yl)ethyl)piperazine-1-carbonyl)phenyl)-3*H*-xanthen-3-ylidene)-*N*-ethylethanaminium chloride **P1** was prepared following *GP5* from *N*-(9-(2-(4-(2-azidoethyl)piperazine-1-carbonyl)phenyl)-6-(diethylamino)-3*H*-xanthen-3-ylidene)-*N*-ethylethanaminium chloride[110] (90 mg, 0.146 mmol) and ethyl undec-10-yn-1-ylphosphonofluoridate (1.0 eq., 38 mg). Yield: 40 mg (0.0455 mmol, 31.2%) of pink-violet semisolid. ESI-HRMS: *m/z* = 842.4878 ($M^+$); $C_{47}H_{66}N_7O_4FP$ requires: *m/z* = 842.4892 ($M^+$), *m/z* = 421.7479 ($(M+H)^{2+}$); $C_{47}H_{67}N_7O_4FP$ requires: *m/z* = 421.7483 ($(M+H)^{2+}$). $\nu_{max}$ 3434, 2975, 2930, 2855, 2362, 1692, 1632, 1589, 1529, 1468, 1414, 1340, 1276, 1181, 1133, 1075, 683 $cm^{-1}$. Purity: UPLC (method II, 254 nm): $t_r$ = 3.353 min, 97.2% total area. $^1H$ NMR (400 MHz, $CDCl_3$) δ = 1.26 – 1.40 (m, 24H), 1.58 – 1.68 (m, 4H), 1.83 – 1.93 (m, 2H), 2.29 (br s, 2H), 2.41 (br s, 2H), 2.63 – 2.69 (m, 2H), 2.78 (t, *J*=6.5 Hz, 2H), 2.85 (s, 1H), 3.38 (br s, 4H), 3.58 – 3.70 (m, 8H), 4.20 – 4.30 (m, 2H), 4.44 (t, *J*=6.5 Hz, 2H), 6.77 (d, *J*=2.4 Hz, 2H), 6.99 (dd, *J*=1.9 Hz, 9.5 Hz, 2H), 7.23 (d, *J*=9.5 Hz, 2H), 7.31 – 7.34 (m, 1H), 7.51 – 7.54 (m, 1H), 7.57 (s, 1H), 7.65 – 7.68 (m, 2H). $^{13}C$ NMR (101 MHz, $CDCl_3$) δ = 12.60, 16.32 (d, *J*=5.7 Hz), 21.85 (d, *J*=5.6 Hz), 24.22 (dd, *J*= 22.2 Hz, 142.8 Hz), 25.63, 28.88, 29.14, 29.16, 29.19, 29.47, 30.11, 30.28, 41.74, 46.09, 47.32, 47.59, 52.69 – 52.81 (m), 57.34, 63.03 (d, *J*=7.2 Hz), 96.20, 113.74, 114.20, 121.80, 127.62, 129.82, 130.04, 130.13, 130.70, 132.12, 135.37, 148.03, 155.60, 156.09, 157.70, 167.19. $^{31}P$ NMR (162 MHz, $CDCl_3$) δ = 31.97 (d, *J*=1069.1 Hz). $^{19}F$ NMR (376 MHz, $CDCl_3$) δ = –74.43, –64.83 (dt, *J*=3.9 Hz, 1068.9 Hz).

**P2**

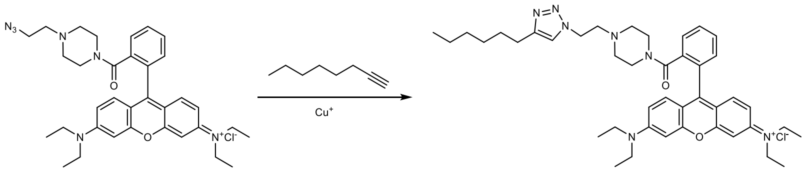

*N*-(6-(Diethylamino)-9-(2-(4-(2-(4-hexyl-1*H*-1,2,3-triazol-1-yl)ethyl)piperazine-1-carbonyl)phenyl)-3*H*-xanthen-3-ylidene)-*N*-ethylethanaminium chloride **P2** was prepared following *GP5* from *N*-(9-(2-(4-(2-azidoethyl)piperazine-1-carbonyl)phenyl)-6-(diethylamino)-3*H*-xanthen-3-ylidene)-*N*-ethylethanaminium chloride (125 mg, 0.203 mmol) and 1-octyne (3.5 eq., 104 µL). Yield: 97 mg (0.134 mmol, 65.8%) of pink-violet semisolid. ESI-HRMS: *m/z* = 690.4480 ($M^+$); $C_{42}H_{56}N_7O_2$ requires: *m/z* = 690.4490 ($M^+$), *m/z* = 345.7280 ($(M+H)^{2+}$); $C_{42}H_{57}N_7O_2$ requires: *m/z* = 345.7281 ($(M+H)^{2+}$). $\nu_{max}$ 3471, 2930, 2859, 2359, 2340, 1691, 1631, 1588, 1468, 1414, 1339, 1181, 1132, 1075, 1011, 827 $cm^{-1}$. Purity: UPLC (method II, 254 nm): $t_r$ = 3.303 min, 97.0% total area. $^1H$ NMR (400 MHz, $CDCl_3$) δ = 0.84 – 0.88 (m, 3H), 1.17 (t, *J*=7.0 Hz, 1H), 1.25 – 1.36 (m, 16H), 1.59 – 1.67 (m, 2H), 2.29 (br s, 2H), 2.38 (br s, 2H), 2.66 (t, *J*=7.6 Hz, 2H), 2.77 (t, *J*=6.4 Hz, 2H), 2.93 (br s, 1H), 3.32 – 3.42 (m,

4H), 3.55 – 3.70 (m, 8H), 4.43 (t, *J*=6.4 Hz, 2H), 6.78 (d, *J*=2.4 Hz, 2H), 6.96 (dd, *J*=2.2 Hz, 9.6 Hz, 2H), 7.23 (d, *J*=9.5 Hz, 2H), 7.31 – 7.34 (m, 1H), 7.51 – 7.55 (m, 2H), 7.65 – 7.68 (m, 2H).$^{13}$C NMR (101 MHz, $CDCl_3$) δ = 12.50, 12.56, 14.04, 22.52, 25.63, 28.86, 29.43, 31.51, 41.73, 44.47, 46.04, 47.29, 47.55, 52.70, 57.30, 96.21, 113.72, 114.09, 121.72, 127.59, 128.42, 128.54, 128.93, 129.20, 129.81, 130.02, 130.13, 130.67, 131.96, 132.06, 132.08, 135.33, 148.09, 155.58, 156.05, 157.69, 167.17.

**P3**

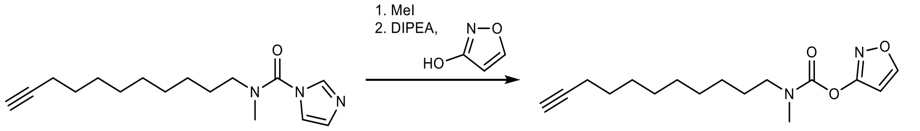

3-Methyl-1-(methyl(undec-10-yn-1-yl)carbamoyl)-1*H*-imidazol-3-ium iodide was prepared by stirring *N*-methyl-*N*-(undec-10-yn-1-yl)-1*H*-imidazole-1-carboxamide (438 mg, 1.59 mmol) and methyl iodide (1.5 eq., 99 µL) in MeCN (3 mL) under Ar for 24 h. The solvent was removed *in vacuo*, the residue dissolved in MeCN (3 mL), isoxazol-3-ol (1.5 eq., 203 mg), and DIPEA (500 µL) were added, and the reaction mixture stirred at rt for 18 h. The solvent was removed *in vacuo* and isoxazol-3-yl methyl(undec-10-yn-1-yl)carbamate isolated by column chromatography on silica (1. PE/DCM = 3:1; 2. DCM). Yield: 200 mg (0.684 mmol, 43.0%) of beige oil, which was directly used further.

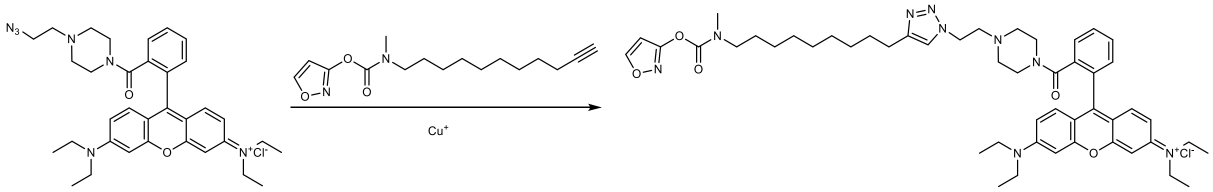

*N*-(6-(Diethylamino)-9-(2-(4-(2-(4-(9-(((isoxazol-3-yloxy)carbonyl)(methyl)amino)nonyl)-1*H*-1,2,3-triazol-1-yl)ethyl)piperazine-1-carbonyl)phenyl)-3*H*-xanthen-3-ylidene)-*N*-ethylethanaminium chloride **P3** was prepared following *GP5* from *N*-(9-(2-(4-(2-azidoethyl)piperazine-1-carbonyl)phenyl)-6-(diethylamino)-3*H*-xanthen-3-ylidene)-*N*-ethylethanaminium chloride (70 mg, 0.114 mmol) and isoxazol-3-yl methyl(undec-10-yn-1-yl)carbamate (1.0 eq., 33 mg). Yield: 35 mg (0.0385 mmol, 33.8%) of pink-violet semisolid. ESI-HRMS: *m/z* = 872.5189 ($M^+$); $C_{50}H_{66}N_9O_5$ requires: *m/z* = 872.5181 ($M^+$). $\nu_{max}$ 3454, 2971, 2930, 2855, 1739, 1691, 1589, 1467, 1414, 1340, 1276, 1181, 1132, 1075 cm$^{-1}$. Purity: UPLC (method II, 254 nm): $t_r$ = 3.720 min, 95.5% total area. $^1$H NMR (400 MHz, $CDCl_3$) δ = 1.21 – 1.32 (m, 22H), 1.51 – 1.64 (m, 4H), 2.22 – 2.27 (m, 2H), 2.31 – 2.37 (m, 2H), 2.61 (t, *J*=7.8 Hz, 2H), 2.73 (t, *J*=6.5 Hz, 2H), 2.95 and 3.03 (2 × s, 3H), 3.26 – 3.38 (m, 6H), 3.51 – 3.66 (m, 8H), 4.38 (t, *J*=6.5 Hz, 2H), 6.48 (dd, *J*=1.8 Hz, 5.3 Hz, 1H), 6.73 (d, *J*=2.4 Hz, 2H), 6.93 (dd, *J*=2.0 Hz, 9.5 Hz, 2H), 7.19 (d, *J*=9.5 Hz, 2H), 7.26 – 7.29 (m, 1H), 7.47 – 7.50 (m, 2H), 7.60 – 7.64 (m, 2H), 8.27 (t, *J*=1.7 Hz, 1H). $^{13}$C NMR (101 MHz, $CDCl_3$) δ = 12.63, 25.69, 26.54, 26.57, 27.15, 27.93, 29.22, 29.26, 29.41, 29.51, 34.72, 35.09, 46.11, 47.35, 47.63, 49.61, 49.79, 52.78, 57.38, 96.27, 99.35, 99.39, 113.80, 114.20, 121.79, 127.68, 129.87, 130.10, 130.19, 130.74, 132.18, 135.44, 148.15, 151.52, 151.62, 155.66, 156.15, 157.77, 159.99, 160.05, 166.63, 167.25.

**P4**

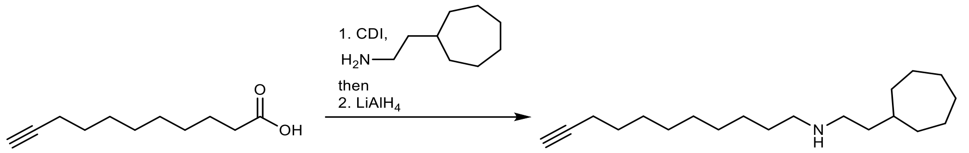

*N*-(2-Cycloheptylethyl)undec-10-ynamide was prepared following *GP1* from undec-10-ynoic acid (260 mg, 1.427 mmol), CDI (1.1 eq., 254 mg), and 2-cycloheptylethan-1-amine (1.1 eq., 222 mg) in THF (5 mL), isolated by column chromatography on silica (1. PE/EtOAc = 5:1; 2. PE/EtOAc = 3:1, 278 mg, 63.8% yield), and immediately reduced following *GP4* to afford *N*-(2-cycloheptylethyl)undec-10-yn-1-amine. Yield: 227 mg (0.779 mmol, 54.6% over two steps) of colourless oil, which used without further purification. ESI-HRMS: *m/z* = 292.2994 ($MH^+$); $C_{20}H_{38}N$ requires: *m/z* = 292.2999 ($MH^+$). $^1$H NMR (400 MHz, $CDCl_3$) δ = 1.14 – 1.72 (m, 29H), 1.94 (t, *J*=2.6 Hz, 1H), 2.12 – 2.20 (m, 4H), 3.22 – 3.29 (m, 2H), 5.31 (s, 1H). $^{13}$C NMR (101 MHz, $CDCl_3$) δ = 18.54, 25.95, 26.50, 28.60, 28.63, 28.84, 29.10, 29.37, 29.39, 34.57, 37.06, 37.09, 37.91, 37.96, 68.24, 84.90.

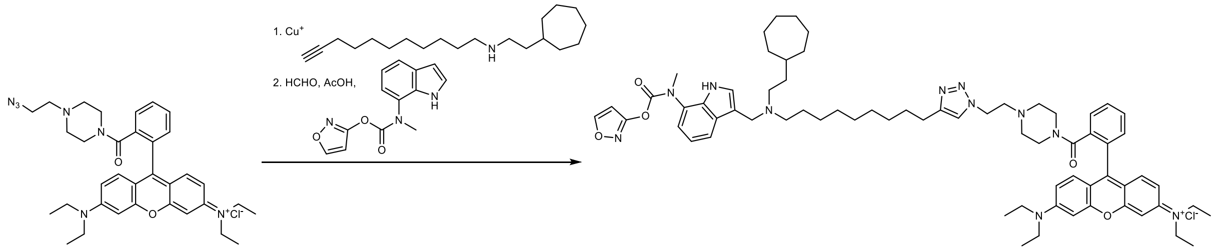

*N*-(9-(2-(4-(2-(4-(9-((2-Cycloheptylethyl)amino)nonyl)-1*H*-1,2,3-triazol-1-yl)ethyl)piperazine-1-carbonyl)phenyl)-6-(diethylamino)-3*H*-xanthen-3-ylidene)-*N*-ethylethanaminium chloride **P4a** was prepared following *GP5* from *N*-(9-(2-(4-(2-azidoethyl)piperazine-1-carbonyl)phenyl)-6-(diethylamino)-3*H*-xanthen-3-ylidene)-*N*-ethylethanaminium chloride (585 mg, 0.950 mmol) and *N*-(2-cycloheptylethyl)undec-10-yn-1-amine (1.0 eq., 277 mg). Yield: 318 mg (0.350 mmol, 36.9%) of pink-violet semisolid, which was immediately used further.

*N*-(9-(2-(4-(2-(4-(9-((2-Cycloheptylethyl)((7-(((isoxazol-3-yloxy)carbonyl)(methyl)amino)-1*H*-indol-3-yl)methyl)amino)nonyl)-1*H*-1,2,3-triazol-1-yl)ethyl)piperazine-1-carbonyl)phenyl)-6-(diethylamino)-3*H*-xanthen-3-ylidene)-*N*-ethylethanaminium chloride **P4** was prepared following *GP2* from **12a** (1.2 eq., 68 mg) and **P4a** (200 mg, 0.220 mmol), and purified by column chromatography on silica (DCM/MeOH = 15:1). Yield: 69 mg (0.0586 mmol, 26.6%) of violet-pink semisolid. ESI-HRMS: *m/z* = 1140.7098 ($M^+$); $C_{68}H_{90}N_{11}O_5$ requires: *m/z* = 1140.7121 ($M^+$), *m/z* = 570.8592 ($(M+H)^{2+}$); $C_{68}H_{91}N_{11}O_5$ requires: *m/z* = 570.8597 ($(M+H)^{2+}$). $\nu_{max}$ 3413, 2927, 2854, 1744, 1690, 1589, 1466, 1414, 1339, 1275, 1181, 1132, 1075 $cm^{-1}$. Purity: UPLC (method II, 210 nm): $t_r$ = 3.773 min, 90.1% total area.$^1$H NMR (400 MHz, $CDCl_3$) δ = 1.08 – 1.73 (m, 41H), 2.23 – 2.33 (m, 4H), 2.63 (t, *J*=7.6 Hz, 2H), 2.67 – 2.95 (m, 6H), 3.26 – 3.38 (m, 6H), 3.40 (s, 2H), 3.51 – 3.61 (m, 8H), 4.33 – 4.40 (m, 2H), 6.71 – 6.74 (m, 2H), 6.89 (d, *J*=9.6 Hz, 2H), 7.05 – 7.09 (m, 2H), 7.18 (d, *J*=9.6 Hz, 2H), 7.26 – 7.30 (m, 1H), 7.38 (s, 1H), 7.44 – 7.66 (m, 5H). $^{13}$C NMR (101 MHz, $CDCl_3$) δ = 12.60, 12.66, 24.91, 25.62, 26.29, 26.35, 26.90, 28.42, 28.98, 29.04, 29.10, 29.35, 34.24, 34.47, 37.29, 37.53, 38.50, 41.86, 44.62, 46.15, 47.41, 47.61, 47.73, 51.39, 52.65, 52.94, 57.36, 96.31, 97.54, 100.15, 100.24, 106.52, 108.41, 115.95, 118.89, 119.99, 120.12, 121.74, 124.58, 125.15,

127.76, 129.24, 129.29, 129.92, 130.12, 130.18, 130.21, 130.75, 132.20, 132.57, 134.17, 135.44, 148.27, 149.93, 151.40, 153.72, 155.68, 156.20, 157.80, 159.88, 161.29, 161.62, 161.95, 166.81, 167.25, 170.10. A mixture of conformers.

**P5**

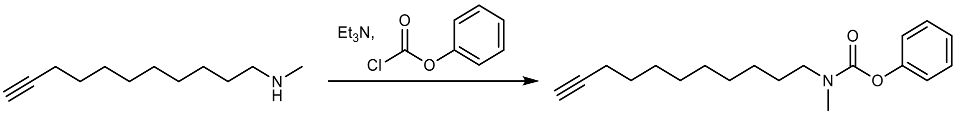

*N*-Methylundec-10-yn-1-amine (544 mg, 3.02 mmol) and triethylamine (1.2 eq., 505 µL) were dissolved in DCM (10 mL), cooled in an icebath, phenyl chloroformate (1.1 eq., 416 µL) was added dropwise, and the reaction mixture stirred at rt for 12 h. The solvent was removed *in vacuo* and phenyl methyl(undec-10-yn-1-yl)carbamate isolated by column chromatography on silica (1. PE/DCM = 10:1; 2. PE/DCM = 5:1). Yield: 438 mg (1.45 mmol, 48 %) of colorless oil. ESI-HRMS: *m/z* = 302.2111 ($MH^+$); $C_{19}H_{28}NO_2$ requires: *m/z* = 302.2115 ($MH^+$). $\nu_{max}$ 3308, 2928, 2855, 1717, 1594, 1456, 1398, 1307, 1203, 1164, 750, 690, 625 $cm^{-1}$. Purity: UPLC (method II, 254 nm): $t_r$ = 4.410 min, 99.1% total area. $^1$H NMR (400 MHz, $CDCl_3$) δ = 1.26 – 1.43 (m, 10H), 1.48 – 1.68 (m, 4H), 1.94 (t, *J*=2.6 Hz, 1H), 2.18 (td, *J*=2.6 Hz, 7.1 Hz, 2H), 2.99 and 3.07 (2 × s, 3H), 3.34 (t, *J*=7.4 Hz, 1H), 3.41 (t, *J*=7.3 Hz, 1H), 7.11 (t, *J*=7.2 Hz, 2H), 7.18 (t, *J*=7.3 Hz, 1H), 7.35 (t, *J*=7.8 Hz, 2H). $^{13}$C NMR (101 MHz, $CDCl_3$) δ = 18.50, 26.75, 27.48, 28.12, 28.56, 28.82, 29.14, 29.40, 29.53, 34.54, 34.88, 49.34, 49.50, 68.21, 84.84, 121.85, 125.16, 125.22, 129.29, 129.33, 151.68, 154.81, 154.92. A mixture of conformers.

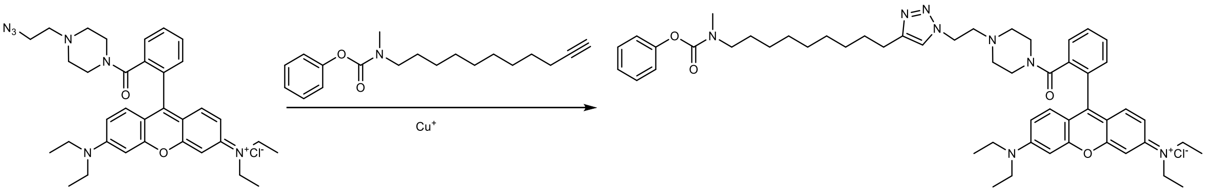

*N*-(6-(diethylamino)-9-(2-(4-(2-(4-(9-(methyl(phenoxycarbonyl)amino)nonyl)-1*H*-1,2,3-triazol-1-yl)ethyl)piperazine-1-carbonyl)phenyl)-3*H*-xanthen-3-ylidene)-*N*-ethylethanaminium chloride **P5** was prepared following *GP5* from *N*-(9-(2-(4-(2-azidoethyl)piperazine-1-carbonyl)phenyl)-6-(diethylamino)-3*H*-xanthen-3-ylidene)-*N*-ethylethanaminium chloride (400 mg, 0.649 mmol) and phenyl methyl(undec-10-yn-1-yl)carbamate (1.5 eq., 195 mg). Yield: 234 mg (0.255 mmol, 39.3%) of pink-violet semisolid. ESI-HRMS: *m/z* = 881.5445 ($M^+$); $C_{53}H_{69}N_8O_4$ requires: *m/z* = 881.5436 ($M^+$). $\nu_{max}$ 3416, 2971, 2929, 2855, 1716, 1692, 1646, 1589, 1467, 1414, 1339, 1276, 1248, 1181, 1133, 1074, 1010, 683 $cm^{-1}$. Purity: UPLC (method II, 254 nm): $t_r$ = 3.987 min, 98.7% total area. $^1$H NMR (400 MHz, $CDCl_3$) δ = 1.26 – 1.36 (m, 22H), 1.58 – 1.69 (m, 4H), 2.29 (br s, 2H), 2.39 (br s, 2H), 2.66 (t, *J*=7.5 Hz, 2H), 2.77 (t, *J*=5.9 Hz, 2H), 2.98 and 3.07 (s, 3H), 3.32 – 3.43 (m, 6H), 3.56 – 3.66 (m, 8H), 4.37 – 4.47 (m, 2H), 6.77 (s, 2H), 6.96 (d, *J*=9.5, 2H), 7.10 (t, *J*=6.8 Hz, 2H), 7.16 (t, *J*=7.4 Hz, 1H), 7.22 (d, *J*=9.4 Hz, 2H), 7.30 – 7.36 (m, 3H), 7.50 – 7.55 (m, 1H), 7.58 (s, 1H), 7.62 – 7.70 (m, 2H). $^{13}$C NMR (101 MHz, $CDCl_3$) δ = 12.30, 25.37, 26.30, 27.01, 27.67, 28.91, 28.96, 28.97, 29.12, 29.14, 29.21, 29.34, 34.16, 34.44, 41.46, 45.76, 46.76, 46.97, 47.30, 48.89, 49.02, 52.42, 52.67, 53.38, 57.04, 95.92, 113.41, 113.82, 121.41, 121.44, 121.58, 124.66, 124.73, 127.31, 128.81, 128.86, 129.54, 129.72, 129.87, 130.44, 131.77, 135.06, 147.68, 151.26, 154.31, 154.42, 155.27, 155.70, 157.38, 166.86.

**B1**

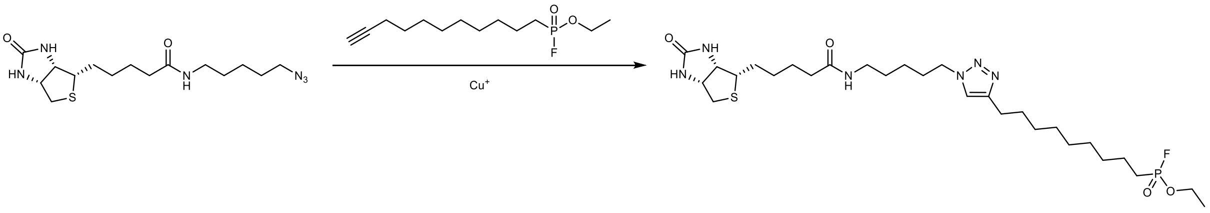

Ethyl (9-(1-(5-(5-((3a*S*,4*S*,6a*R*)-2-oxohexahydro-1*H*-thieno[3,4-*d*]imidazol-4-yl)pentanamido)pentyl)-1*H*-1,2,3-triazol-4-yl)nonyl)phosphonofluoridate **B1** was prepared following *GP5* from *N*-(5-azidopentyl)-5-((3a*S*,4*S*,6a*R*)-2-oxohexahydro-1*H*-thieno[3,4-*d*]imidazol-4-yl)pentanamide (135 mg, 0.381 mmol) and ethyl undec-10-yn-1-ylphosphonofluoridate (100 mg, 1.0 eq.). Yield: 21 mg (0.034 mmol, 8.9%) of beige semisolid. ESI-HRMS: *m/z* = 617.3388 ($MH^+$); $C_{28}H_{51}FN_6O_4PS$ requires: *m/z* = 617.3409 ($MH^+$). $\nu_{max}$ 3295, 2928, 2855, 1664, 1464, 1192, 1164, 1038, 961, 717, 626 $cm^{-1}$. $^1H$ NMR (400 MHz, $CDCl_3$) δ = 1.24 – 1.76 (m, 27H), 1.82 – 1.94 (m, 6H), 2.18 (t, *J*=7.3 Hz, 2H), 2.68 (t, *J*=7.5 Hz, 2H), 2.74 (d, *J*=12.8 Hz, 1H), 2.91 (dd, *J*=4.9 Hz, 12.8 Hz, 1H), 3.12 – 3.17 (m, 1H), 3.20 (q, *J*=6.6 Hz, 2H), 4.31 (t, *J*=7.0 Hz, 3H), 4.50 – 4.54 (m, 1H), 5.49 (s, 1H), 6.35 (t, *J*=5.7 Hz, 1H), 6.42 (s, 1H), 7.27 (s, 1H). $^{13}C$ NMR (101 MHz, $CDCl_3$) δ = 16.46, 16.52, 21.98, 22.03, 23.59, 23.77, 23.81, 25.01, 25.23, 25.80, 28.14, 28.22, 28.94, 29.04, 29.28, 29.31, 29.37, 29.59, 29.97, 30.30, 30.46, 36.00, 39.10, 40.72, 49.98, 55.78, 60.30, 61.90, 63.15, 63.23, 120.69, 148.58, 164.00, 173.41. $^{19}F$ NMR (376 MHz, $CDCl_3$) δ = –64.71 (d, *J*=1069.6 Hz). $^{31}P$ NMR (162 MHz, $CDCl_3$) δ = 31.99 (d, *J*=1069.4 Hz).

**B2**

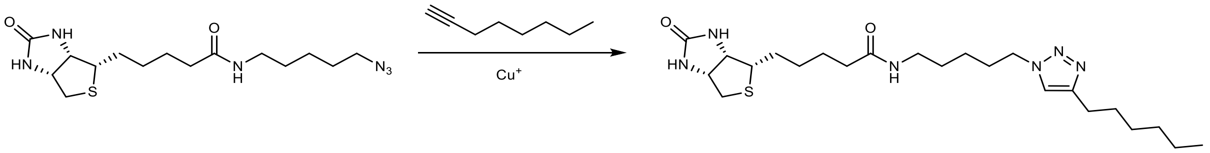

*N*-(5-(4-Hexyl-1*H*-1,2,3-triazol-1-yl)pentyl)-5-((3a*S*,4*S*,6a*R*)-2-oxohexahydro-1*H*-thieno[3,4-*d*]imidazol-4-yl)pentanamide **B2** was prepared following *GP5* from *N*-(5-azidopentyl)-5-((3a*S*,4*S*,6a*R*)-2-oxohexahydro-1*H*-thieno[3,4-*d*]imidazol-4-yl)pentanamide (58 mg, 0.164 mmol) and 1-octyne (10 eq., 241 µL). Yield: 15 mg (0.0323 mmol, 19.7%) of white solid. mp 134.2–137.1 °C. ESI-HRMS: *m/z* = 465.3001 ($MH^+$); $C_{23}H_{41}N_6O_2S$ requires: *m/z* = 465.3006 ($MH^+$). $\nu_{max}$ 3267, 3113, 2925, 2857, 1692, 1625, 1552, 1480, 1466, 1264, 1215, 1150, 1053, 683 $cm^{-1}$. Purity: UPLC (method II, 220 nm): $t_r$ = 2.947 min, 97.5% total area. $^1H$ NMR (400 MHz, DMSO-$d_6$) δ = 0.85 (t, *J*=7.0 Hz, 3H), 1.13 – 1.66 (m, 18H), 1.77 (p, *J*=7.3 Hz, 2H), 2.03 (t, *J*=7.4 Hz, 2H), 2.55 – 2.61 (m, 3H), 2.81 (dd, *J*=5.1 Hz, 12.4 Hz, 1H), 2.99 (q, *J*=6.7 Hz, 2H), 3.06 – 3.12 (m, 1H), 4.10 – 4.14 (m, 1H), 4.24 – 4.32 (m, 3H), 6.36 (s, 1H), 6.43 (s, 1H), 7.74 (t, *J*=5.6 Hz, 1H), 7.82 (s, 1H). $^{13}C$ NMR (101 MHz, DMSO-$d_6$) δ = 13.94, 22.03, 23.29, 25.04, 25.32, 28.04, 28.22, 28.26, 28.54, 28.98, 29.40, 31.01, 35.21, 38.09, 39.85, 49.05, 55.44, 59.19, 61.04, 121.53, 146.83, 162.70, 171.81.

**B3**

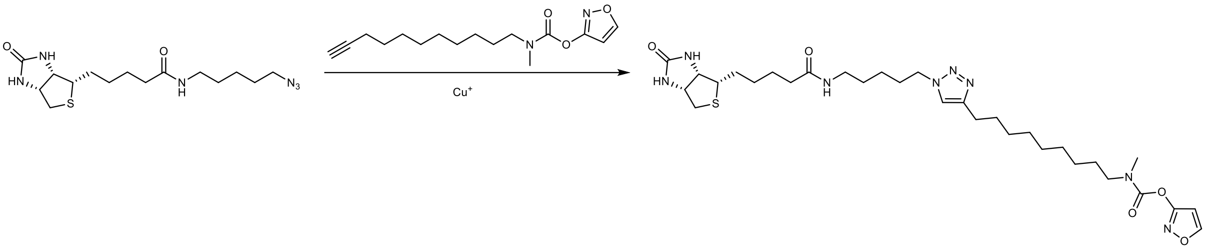

Isoxazol-3-yl methyl(9-(1-(5-(5-((3a*S*,4*S*,6a*R*)-2-oxohexahydro-1*H*-thieno[3,4-*d*]imidazol-4-yl)pentanamido)pentyl)-1*H*-1,2,3-triazol-4-yl)nonyl)carbamate **B3** was prepared following *GP5* from *N*-(5-azidopentyl)-5-((3a*S*,4*S*,6a*R*)-2-oxohexahydro-1*H*-thieno[3,4-*d*]imidazol-4-yl)pentanamide (89 mg, 0.251 mmol) and isoxazol-3-yl methyl(undec-10-yn-1-yl)carbamate (1.0 eq., 73 mg). Yield: 82 mg (0.141 mmol, 46.2%) of white solid. ESI-HRMS: *m/z* = 647.3685 ($MH^+$); $C_{31}H_{51}N_8O_5S$ requires: *m/z* = 647.3698 ($MH^+$). $\nu_{max}$ 3240, 3066, 2925, 2851, 1722, 1690, 1626, 1579, 1398, 1275, 1155, 1095, 774 $cm^{-1}$. Purity: UPLC (method II, 220 nm): $t_r$ = 3.273 min, 99.8% total area. $^1$H NMR (400 MHz, DMSO-$d_6$) δ = 1.13 – 1.65 (m, 25H), 1.77 (p, *J*=7.2 Hz, 2H), 2.03 (t, *J*=7.4 Hz, 2H), 2.54 – 2.60 (m, 3H), 2.81 (dd, *J*=5.1 Hz, 12.4 Hz, 1H), 2.91 and 3.01 (2 × s, 3H), 2.98 (q, *J*=6.3 Hz, 2H), 3.06 – 3.12 (m, 1H), 3.27 (t, *J*=7.2 Hz, 1H), 4.10 – 4.14 (m, 1H), 4.23 – 4.32 (m, 3H), 6.36 (s, 1H), 6.42 (s, 1H), 6.62 (d, *J*=1.8 Hz, 1H), 7.74 (t, *J*=5.6 Hz, 1H), 7.81 (s, 1H), 8.86 (d, *J*=1.8 Hz, 1H). $^{13}$C NMR (101 MHz, DMSO-$d_6$) δ = 23.30, 25.03, 25.33, 25.92, 26.00, 26.54, 27.27, 28.04, 28.23, 28.54, 28.58, 28.64, 28.71, 28.89, 29.01, 29.41, 34.31, 34.68, 35.22, 38.11, 39.86, 48.74, 48.95, 49.06, 55.45, 59.20, 61.05, 99.73, 99.81, 121.53, 146.84, 151.09, 151.11, 162.10, 162.15, 162.71, 166.42, 171.84.

### ABPP: Western-blotted membranes and raw images

The images of the membranes were taken in the same position under white light and then in either fluorescence (595 nm) or chemiluminescence mode, to detect probe-labelled proteins or GAPDH and hBChE, respectively. The pictures were then overlaid, cropped, and treated in the same manner. Due to high fluorescence of the marker lane and the dye front, these areas were covered with aluminium foil before capturing the fluorescent signal.

**HepG2**

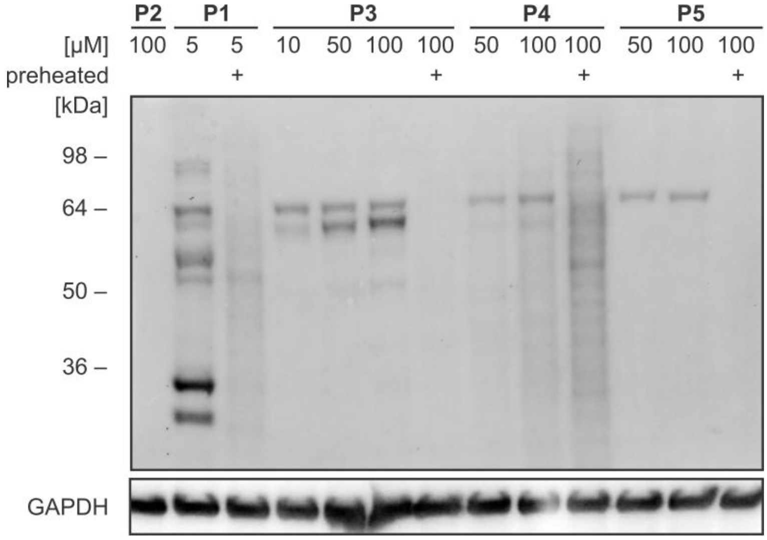

Detection mode: fluorescence (UVITEC)

white light | epi green filter

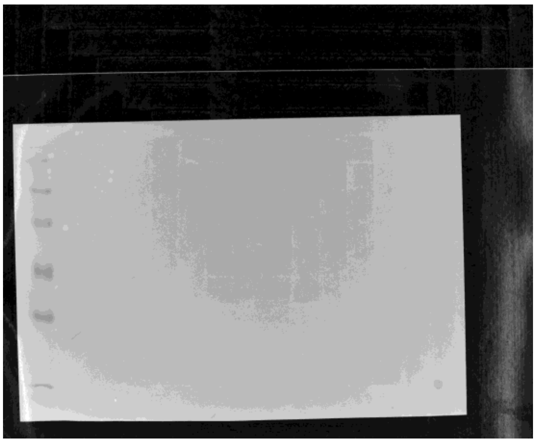

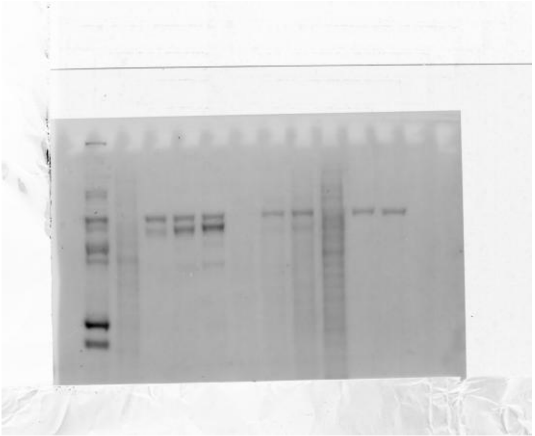

Detection mode: chemiluminescence (G-box)
Primary antibody: anti-GAPDH

white light | Chemiluminescence

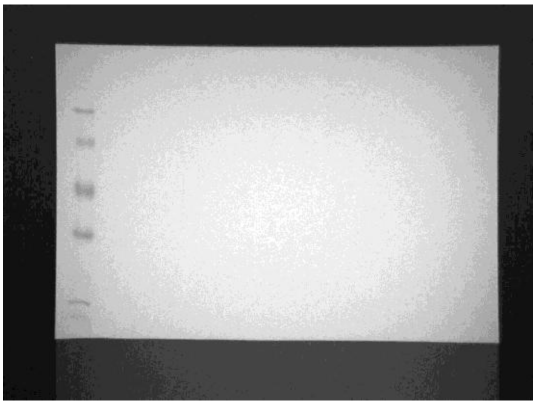

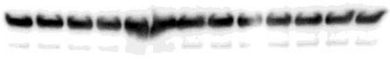

**Fig. S16** | Western-blotted membrane and raw images of the probe-treated HepG2 lysate.

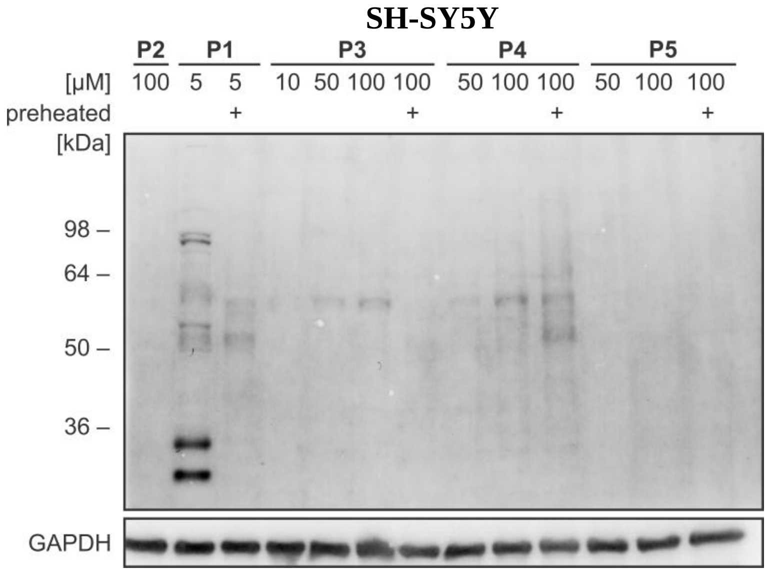

Detection mode: fluorescence (UVITEC)

white light

epi green filter

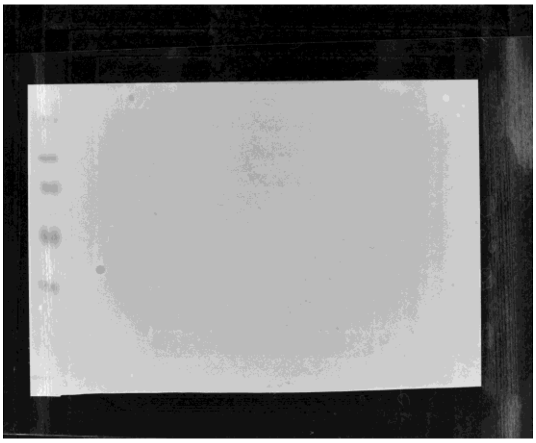

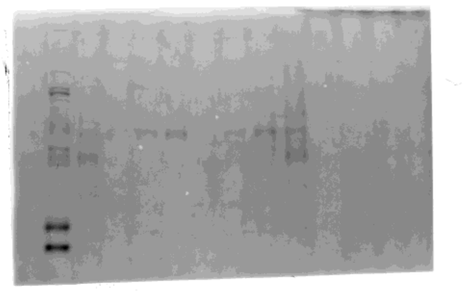

Detection mode: chemiluminescence (G-box)

Primary antibody: anti-GAPDH

white light

chemiluminescence

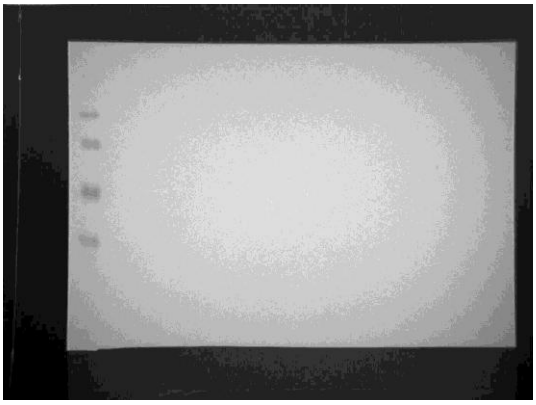

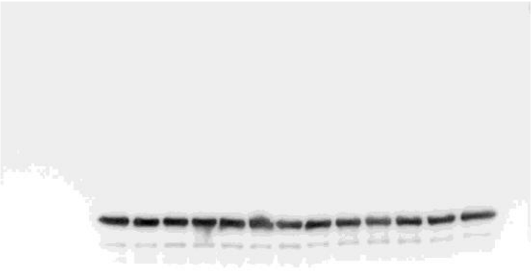

**Fig. S17│** Western-blotted membrane and raw images of the probe-treated SH-SY5Y lysate.

**rhBChE and human plasma**

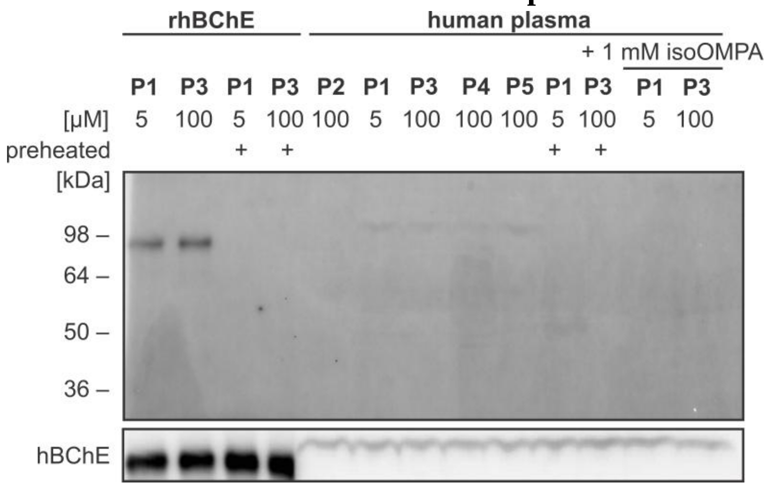

Detection mode: fluorescence (UVITEC)

white light

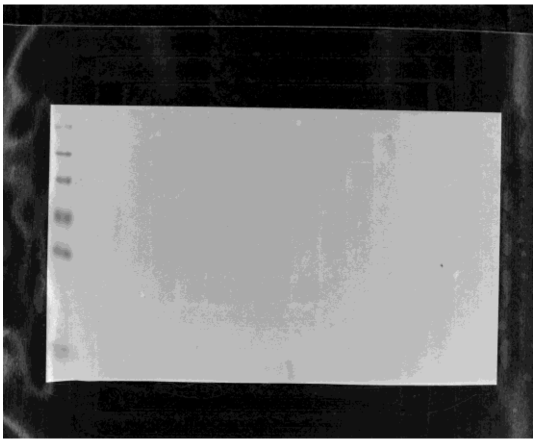

epi green filter

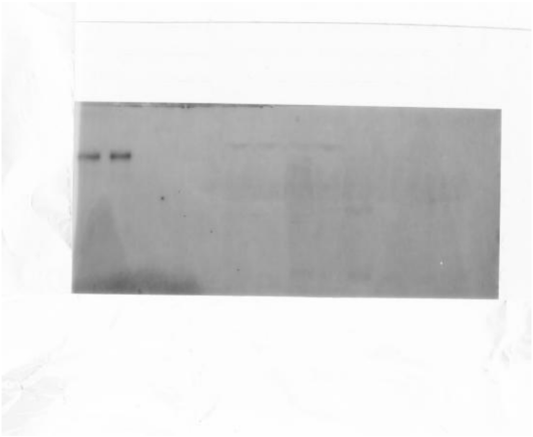

Detection mode: chemiluminescence (G-box)

Primary antibody: anti-hBChE

white light

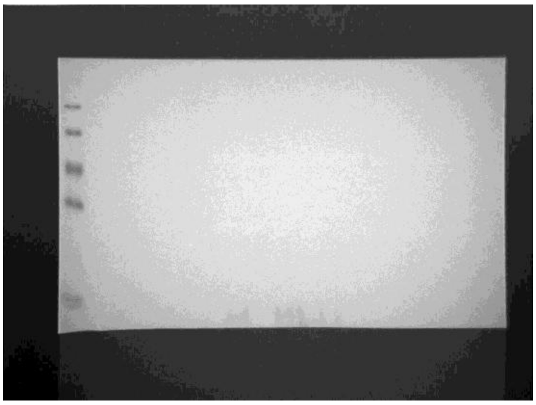

Chemiluminescence

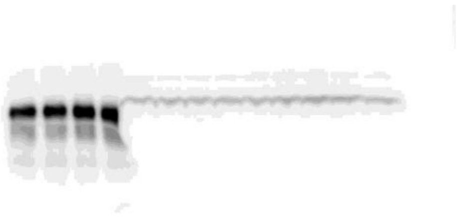

**Fig. S18 |** Western-blotted membrane and raw images of the probe-treated recombinant human BChE and human plasma. Probes **P1** and **P3** activity-dependently label recombinant hBChE, whereas the binding cannot be seen in human plasma (despite enzymatically active hBChE, determined by Ellman method).

### NMR spectra

## 2

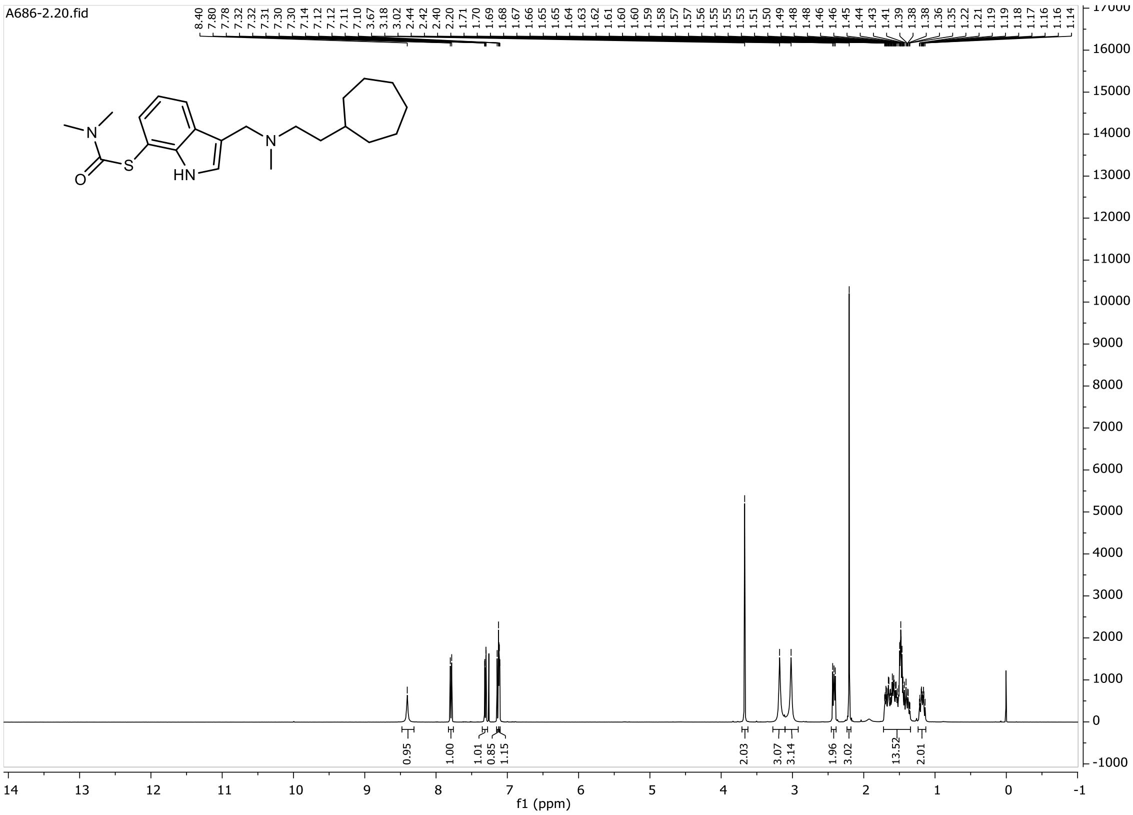

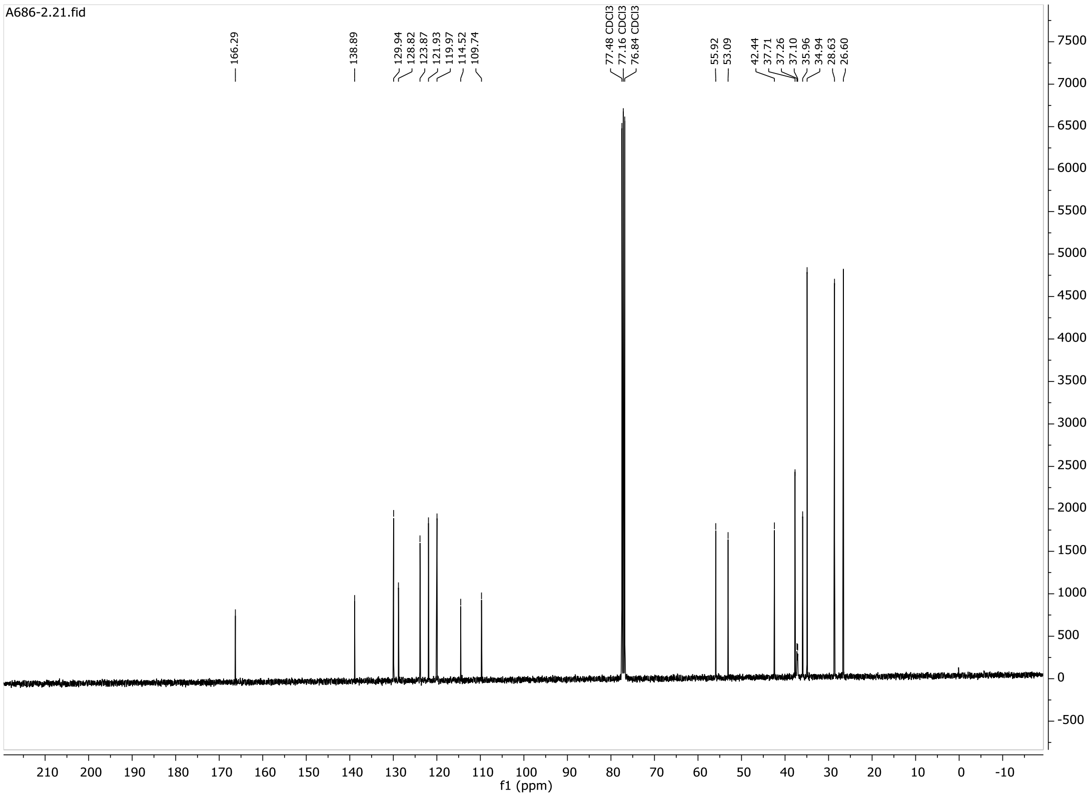

# 3

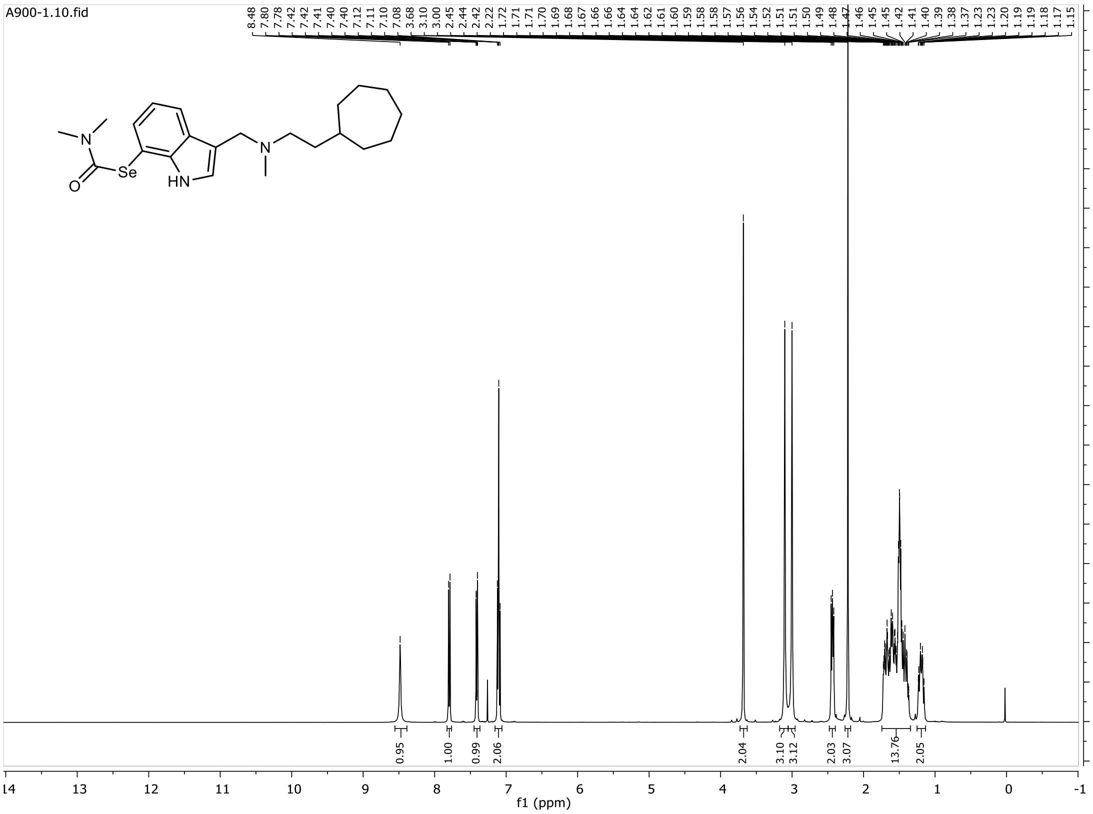
A900-1.10.fid
8.48 7.80 7.78 7.42 7.42 7.41 7.40 7.12 7.11 7.10 7.08 3.68 3.10 3.00 2.45 2.44 2.42 2.22 1.72 1.71 1.71 1.70 1.69 1.68 1.67 1.66 1.66 1.64 1.62 1.61 1.60 1.59 1.58 1.58 1.57 1.56 1.54 1.52 1.51 1.51 1.50 1.49 1.48 1.47 1.46 1.45 1.45 1.42 1.41 1.40 1.39 1.38 1.37 1.23 1.23 1.20 1.19 1.19 1.18 1.17 1.15
Se
HN
N
O
0.95 1.00 0.99 2.06 2.04 3.10 3.12 2.03 3.07 13.76 2.05
f1 (ppm)

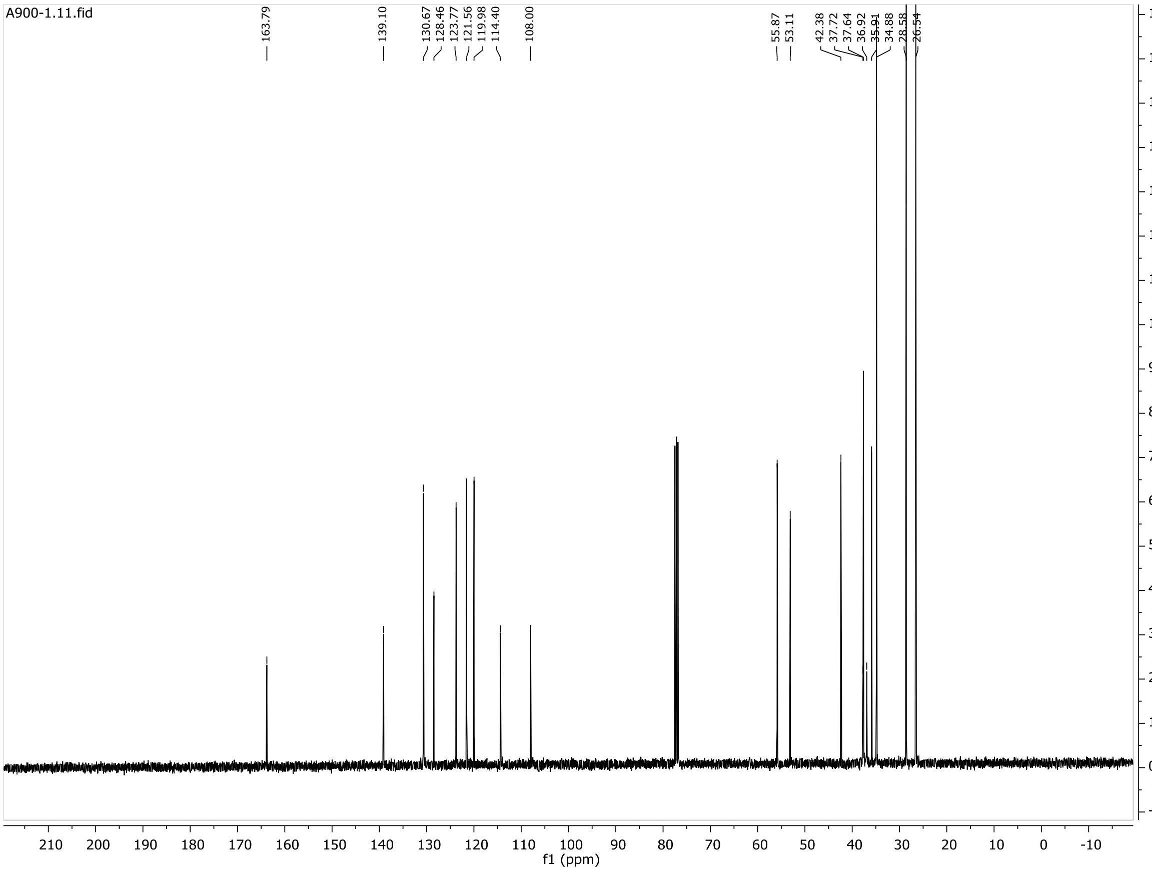
A900-1.11.fid
163.79 139.10 130.67 128.46 123.77 121.56 119.98 114.40 108.00 55.87 53.11 42.38 37.72 37.64 36.92 35.91 34.88 28.58 26.54
f1 (ppm)

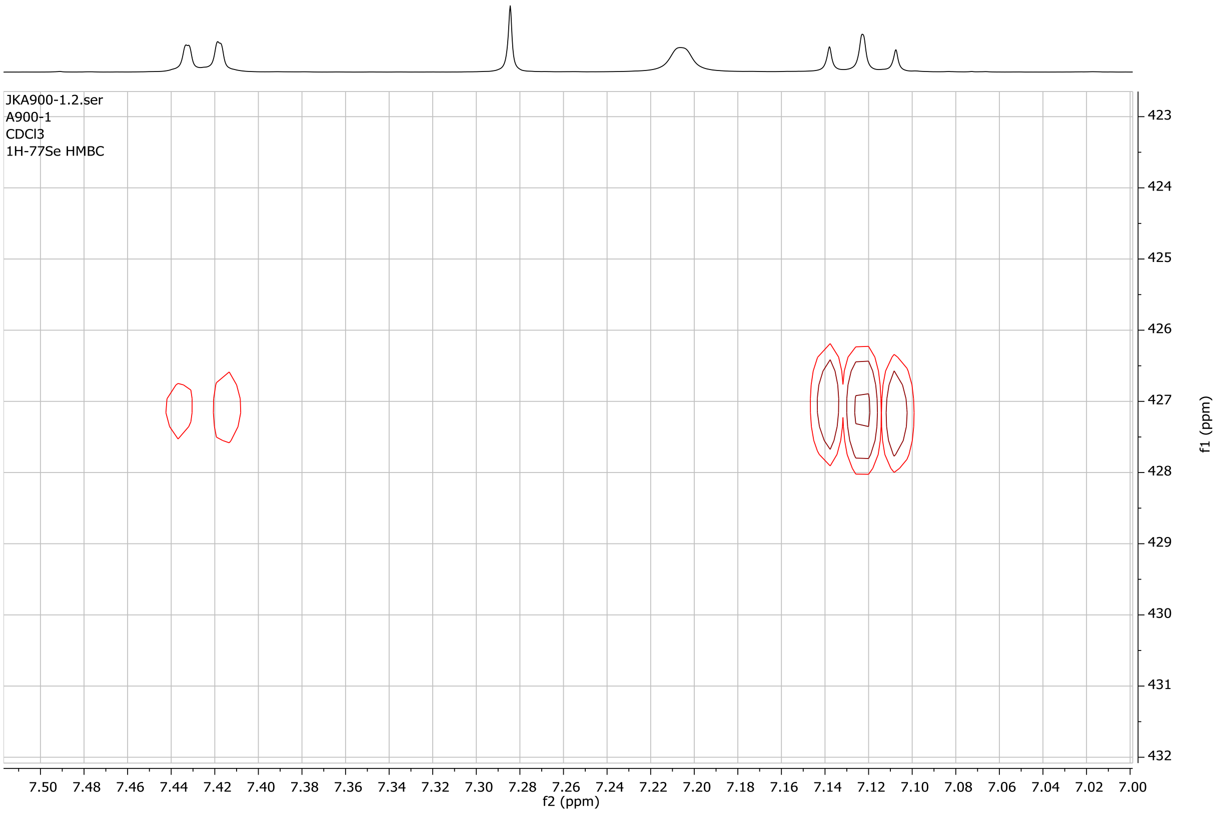

JKA900-1.2.ser
A900-1
CDCl3
1H-77Se HMBC
f2 (ppm)
f1 (ppm)
7.50
7.48
7.46
7.44
7.42
7.40
7.38
7.36
7.34
7.32
7.30
7.28
7.26
7.24
7.22
7.20
7.18
7.16
7.14
7.12
7.10
7.08
7.06
7.04
7.02
7.00

**4**

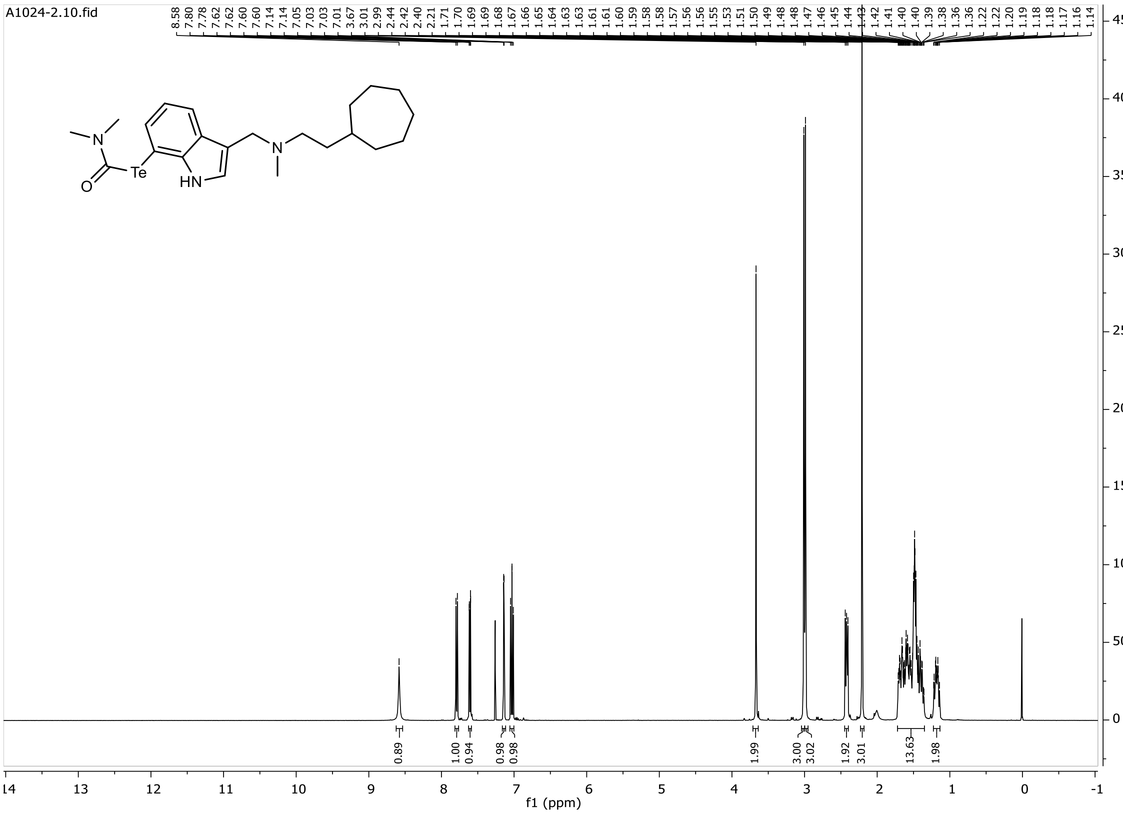
A1024-2.10.fid
8.58 7.80 7.78 7.62 7.62 7.60 7.60 7.14 7.05 7.03 7.03 7.01 3.67 3.01 2.99 2.44 2.42 2.40 2.21 1.71 1.70 1.69 1.69 1.68 1.67 1.66 1.65 1.64 1.63 1.63 1.61 1.61 1.60 1.59 1.58 1.58 1.57 1.56 1.56 1.55 1.53 1.51 1.50 1.49 1.48 1.48 1.47 1.46 1.45 1.44 1.43 1.42 1.41 1.40 1.40 1.39 1.38 1.36 1.36 1.22 1.22 1.20 1.19 1.18 1.18 1.17 1.16 1.14
Te
HN
N
O
0.89 1.00 0.94 0.98 0.98 1.99 3.00 3.02 1.92 3.01 13.63 1.98
f1 (ppm)

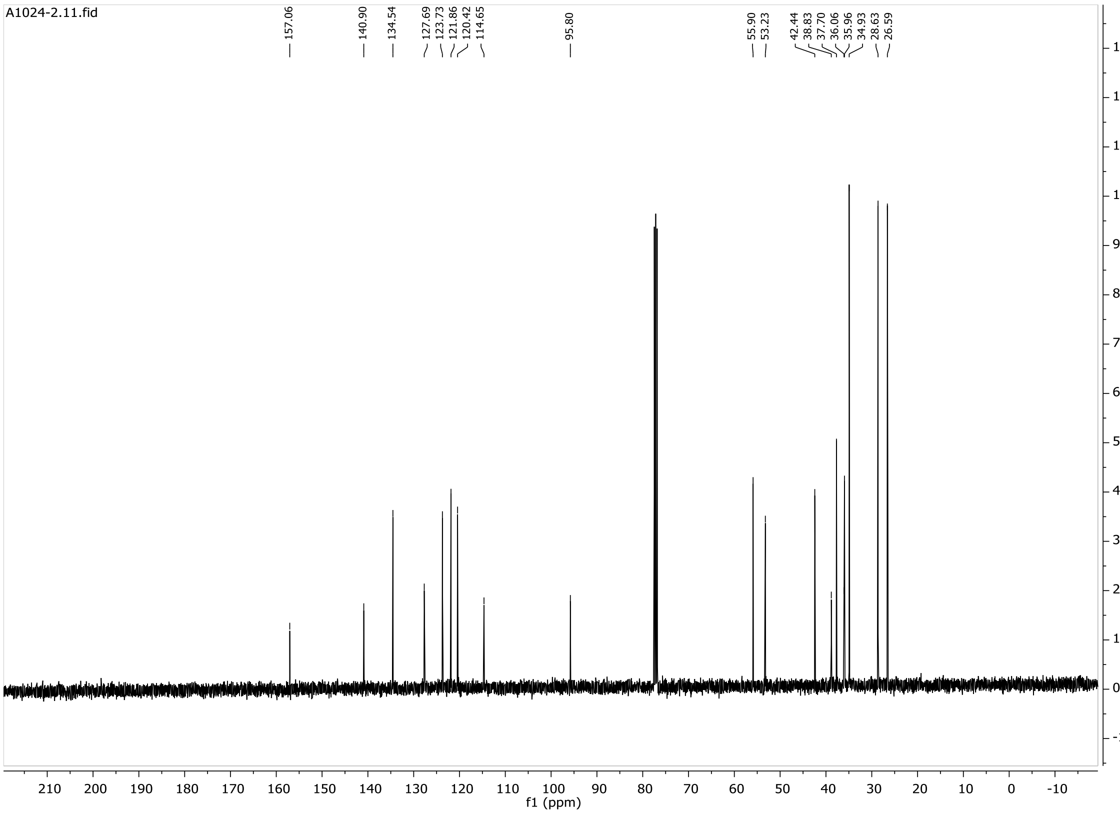
A1024-2.11.fid
157.06 140.90 134.54 127.69 123.73 121.86 120.42 114.65 95.80 55.90 53.23 42.44 38.83 37.70 36.06 35.96 34.93 28.63 26.59
f1 (ppm)

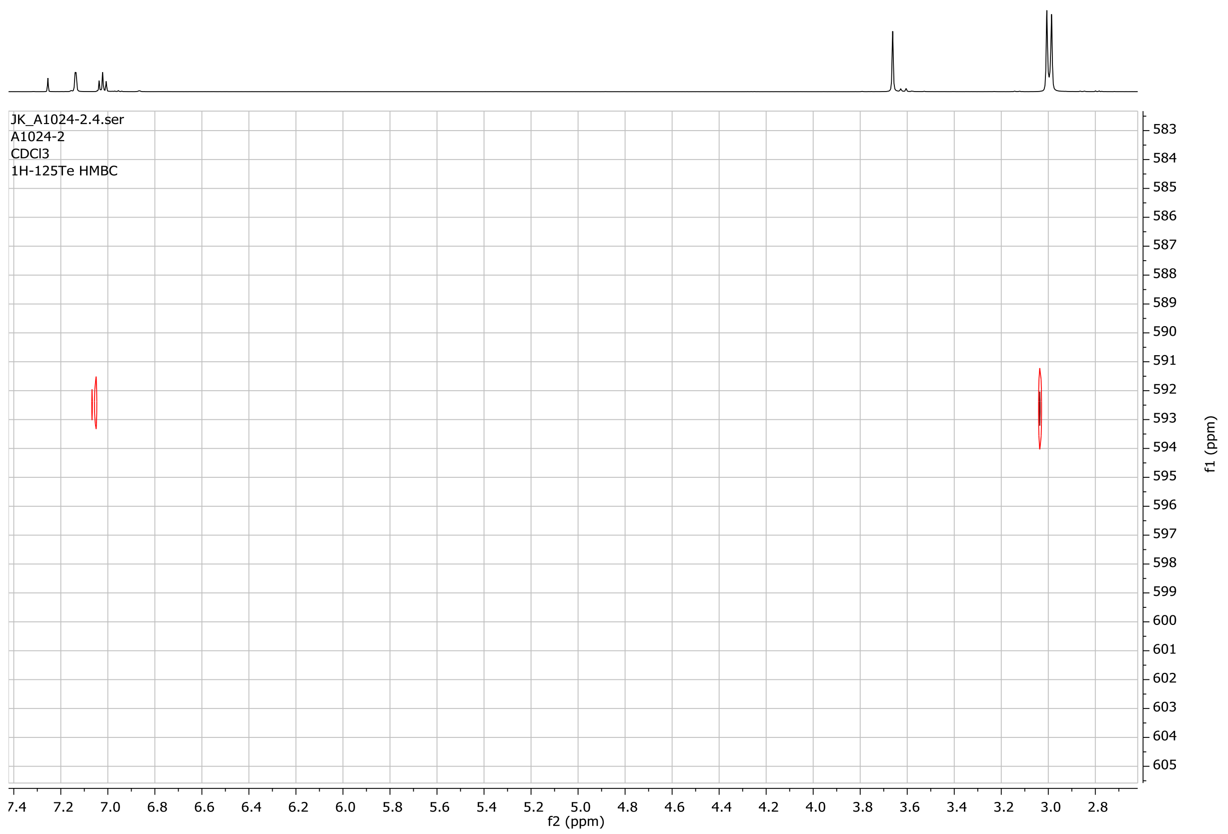
JK_A1024-2.4.ser
A1024-2
CDCl3
1H-125Te HMBC
f2 (ppm)
f1 (ppm)

**5**

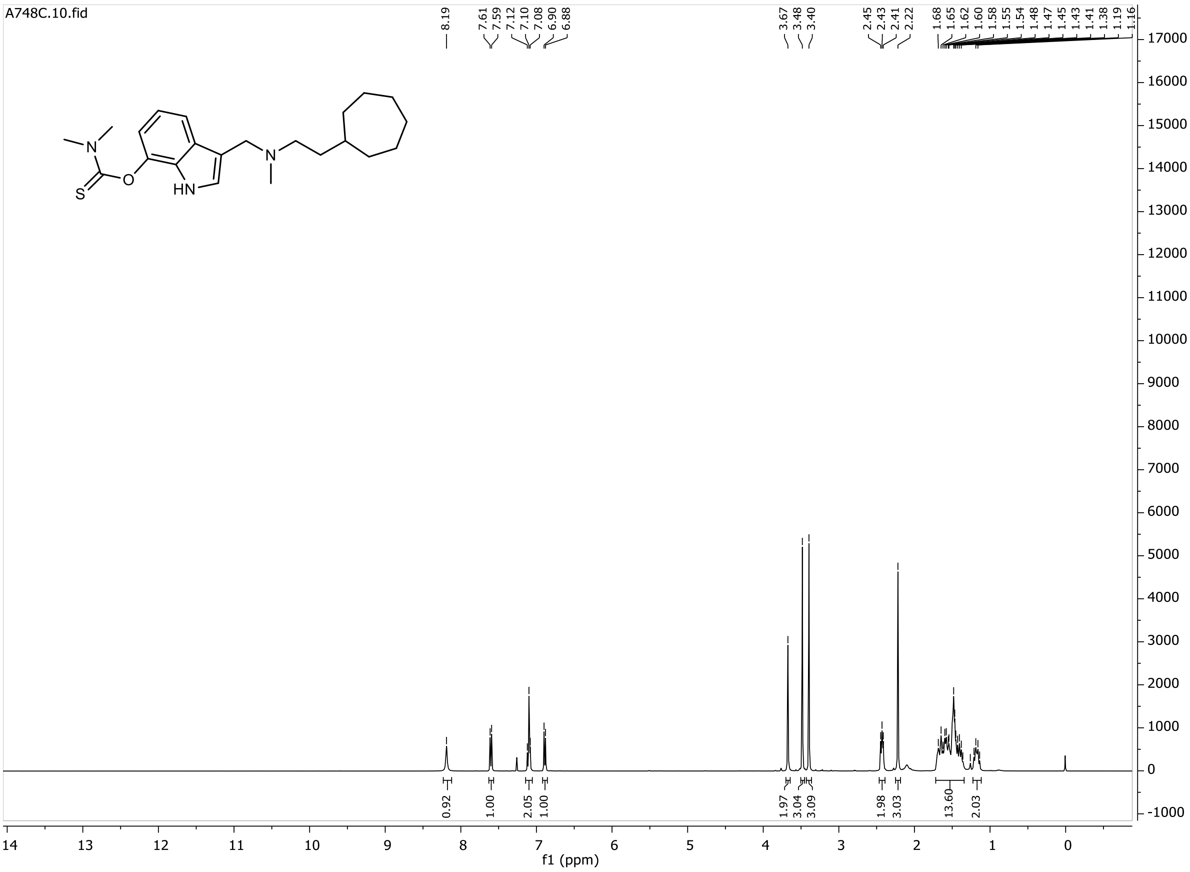
A748C.10.fid
8.19
7.61 7.59 7.12 7.10 7.08 6.90 6.88
3.67 3.48 3.40
2.45 2.43 2.41 2.22
1.68 1.65 1.62 1.60 1.58 1.55 1.54 1.48 1.47 1.45 1.43 1.41 1.38 1.19 1.16
0.92 1.00 2.05 1.00 1.97 3.04 3.09 1.98 3.03 13.60 2.03
HN
N
O
S
f1 (ppm)

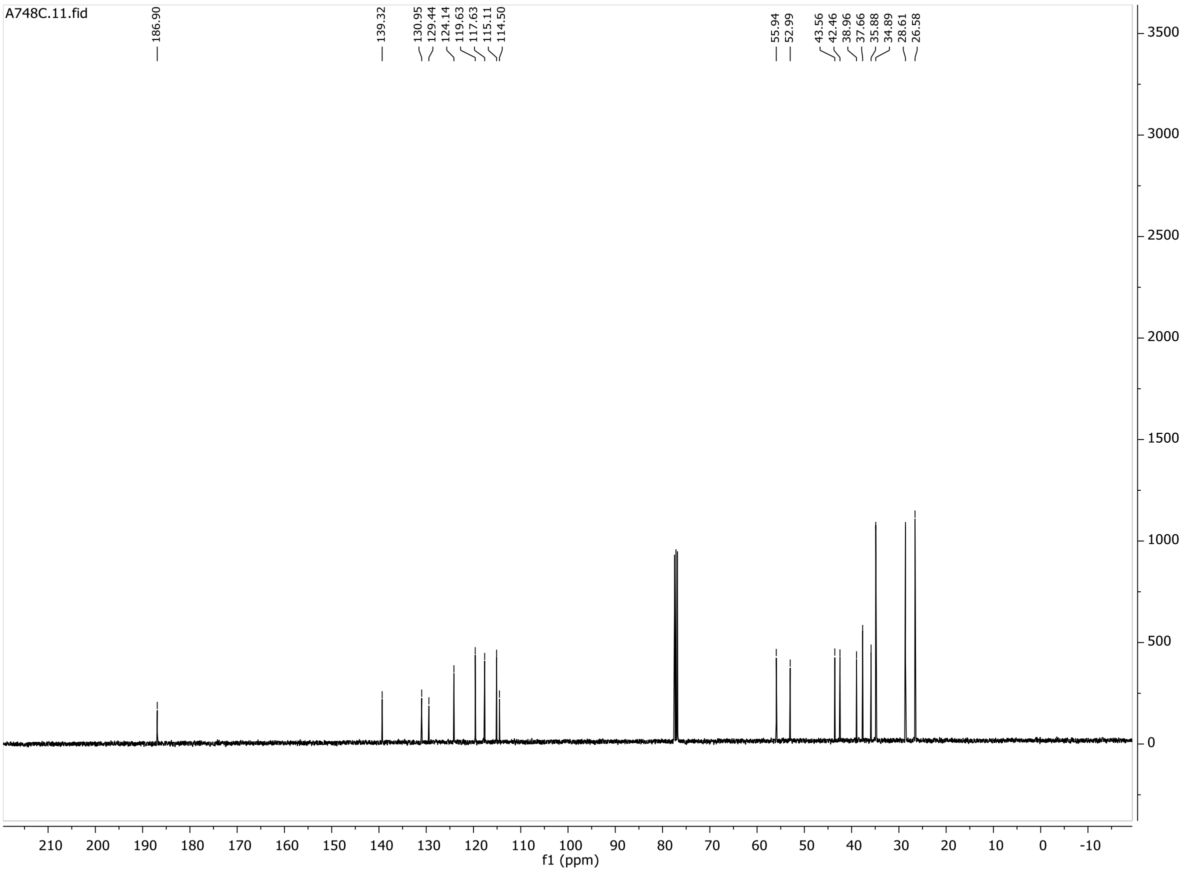
A748C.11.fid
186.90
139.32
130.95 129.44 124.14 119.63 117.63 115.11 114.50
55.94 52.99
43.56 42.46 38.96 37.66 35.88 34.89 28.61 26.58
f1 (ppm)

**6**

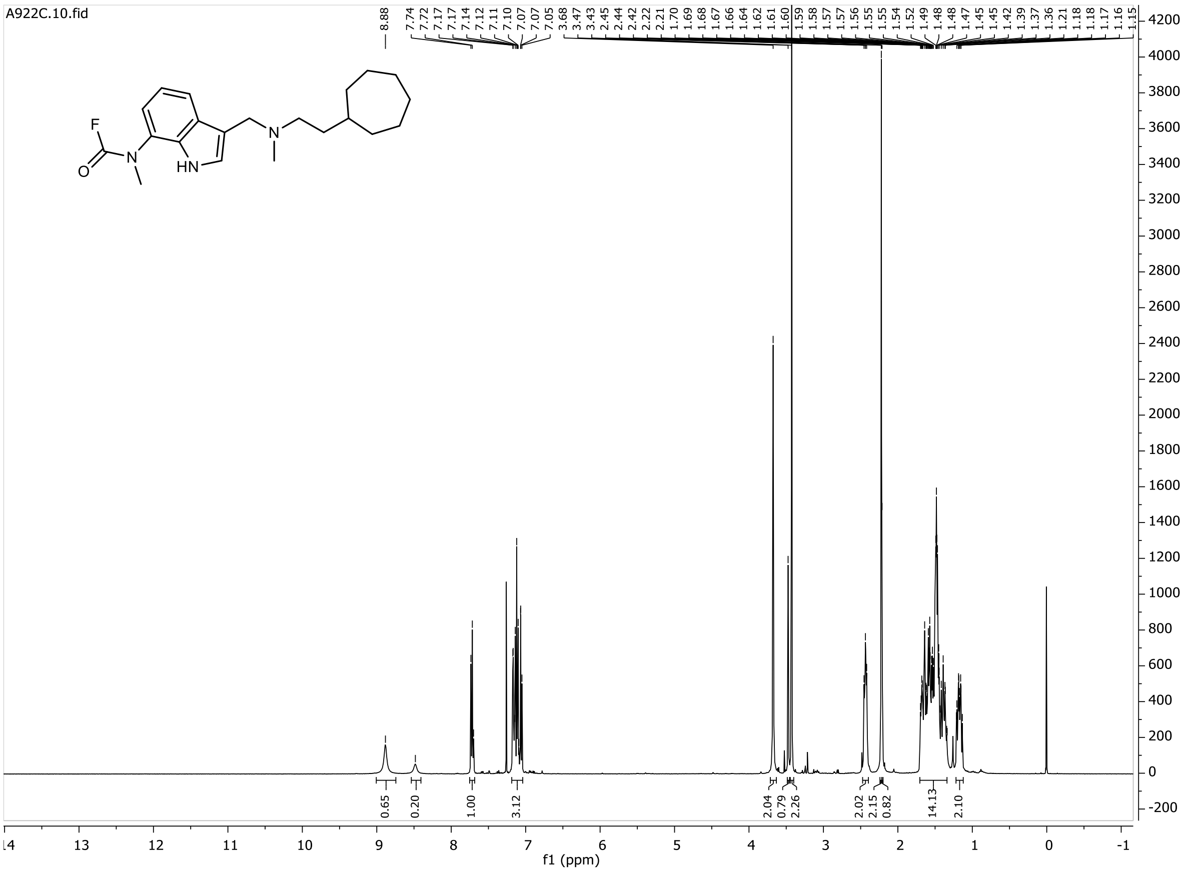
A922C.10.fid
8.88
7.74 7.72 7.17 7.17 7.14 7.12 7.11 7.10 7.07 7.07 7.05 3.68 3.47 3.43 2.45 2.44 2.42 2.22 2.21 1.70 1.69 1.68 1.67 1.66 1.64 1.62 1.61 1.60 1.59 1.58 1.57 1.57 1.56 1.55 1.55 1.54 1.52 1.49 1.48 1.48 1.47 1.45 1.45 1.42 1.39 1.37 1.36 1.21 1.18 1.18 1.17 1.16 1.15
F
O
N
HN
N
0.65 0.20 1.00 3.12 2.04 0.79 2.26 2.02 2.15 0.82 14.13 2.10
f1 (ppm)

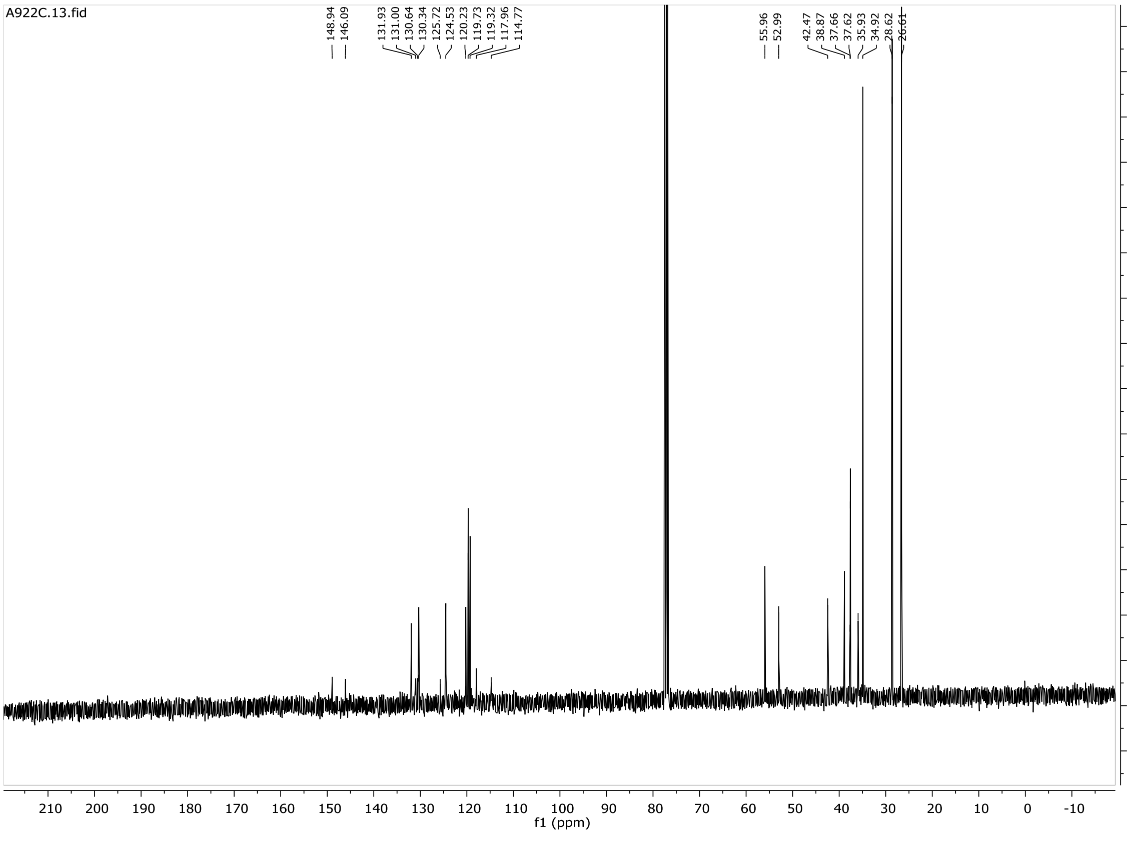
A922C.13.fid
148.94 146.09
131.93 131.00 130.64 130.34 125.72 124.53 120.23 119.73 119.32 117.96 114.77
55.96 52.99
42.47 38.87 37.66 37.62 35.93 34.92 28.62 26.61
f1 (ppm)

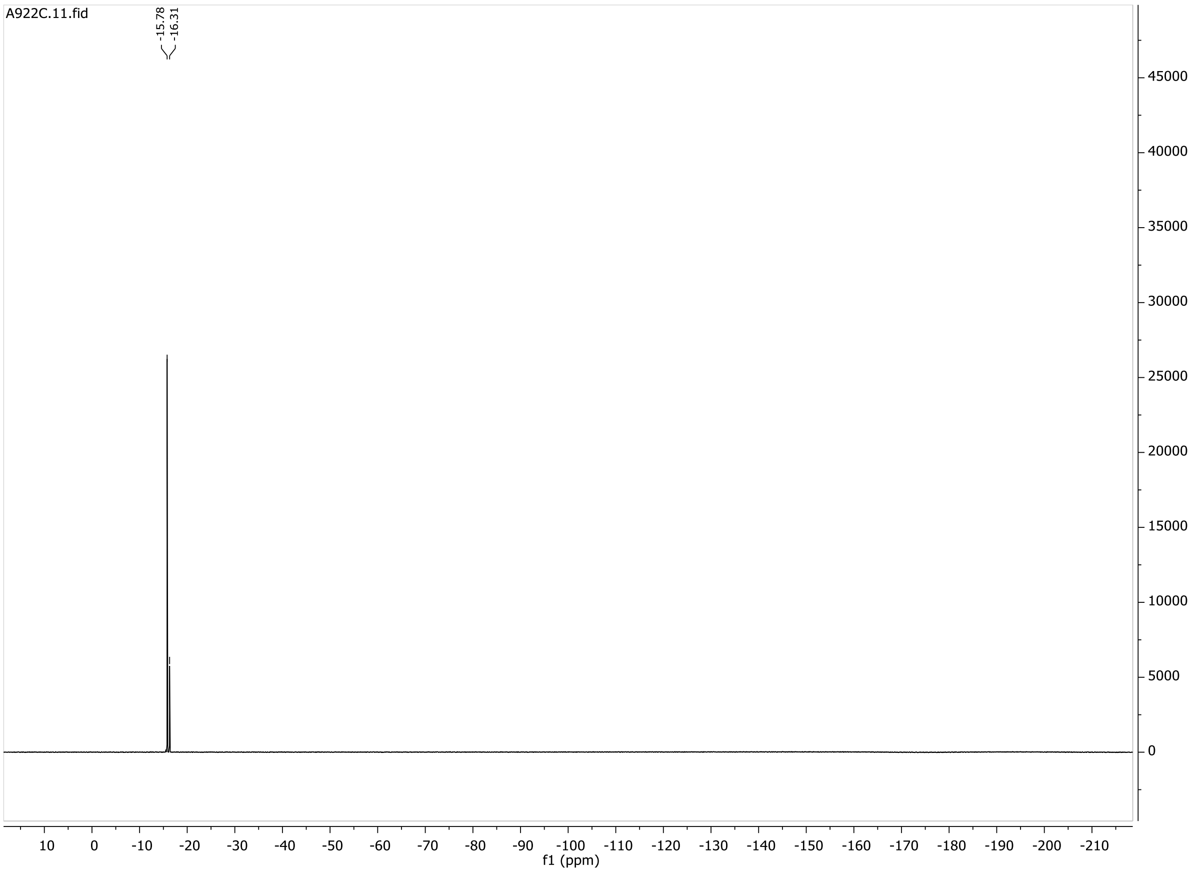
A922C.11.fid
-15.78
-16.31
45000
40000
35000
30000
25000
20000
15000
10000
5000
0
10 0 -10 -20 -30 -40 -50 -60 -70 -80 -90 -100 -110 -120 -130 -140 -150 -160 -170 -180 -190 -200 -210
f1 (ppm)

**6b**

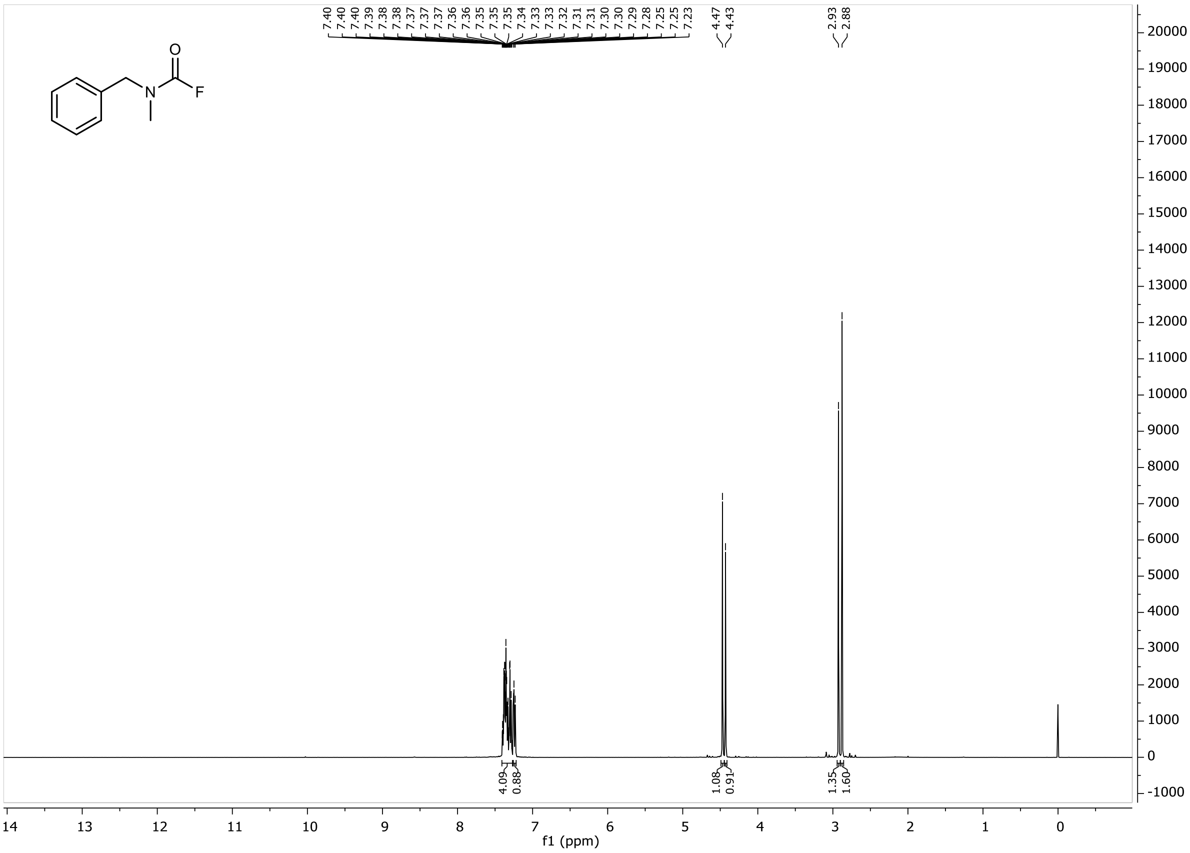
O
N
F
7.40
7.40
7.40
7.39
7.38
7.38
7.37
7.37
7.37
7.36
7.36
7.35
7.35
7.35
7.34
7.33
7.33
7.32
7.31
7.31
7.30
7.30
7.29
7.28
7.25
7.25
7.23
4.47
4.43
2.93
2.88
4.09
0.88
1.08
0.91
1.35
1.60
f1 (ppm)

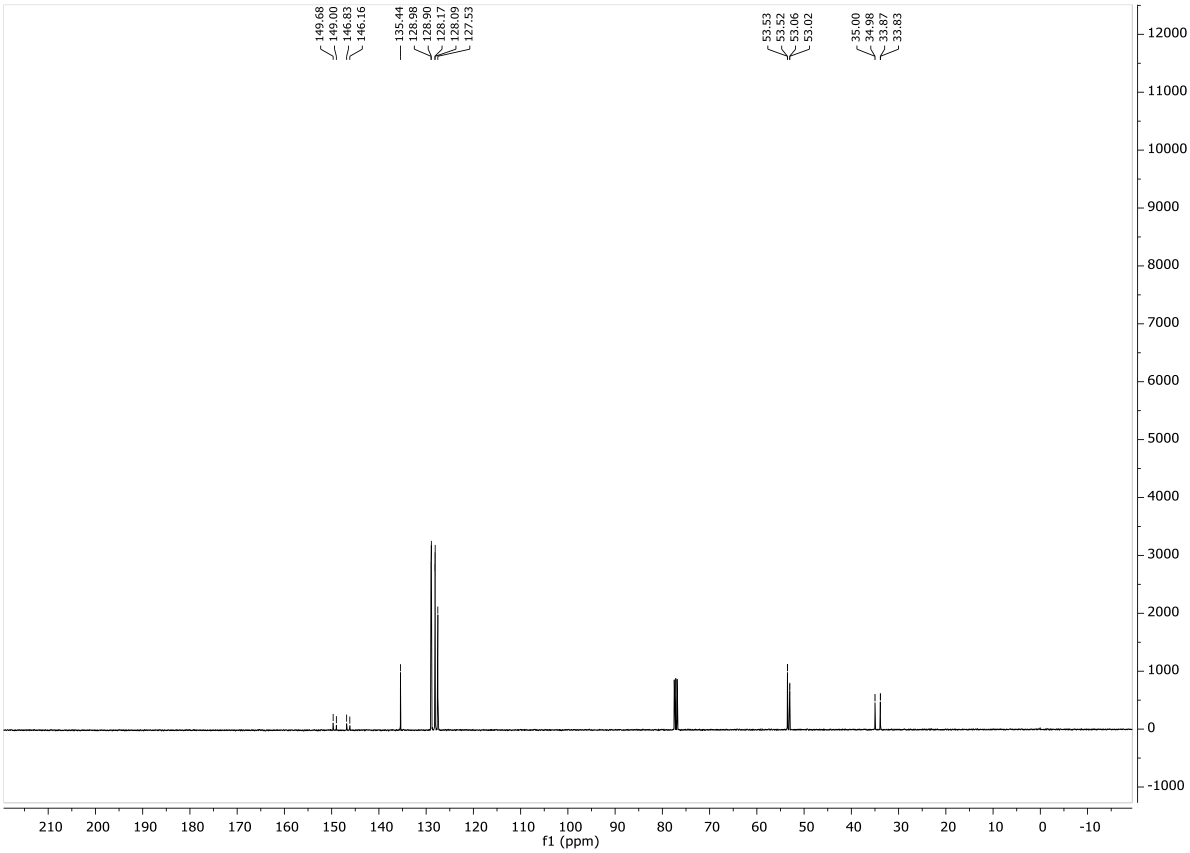
149.68
149.00
146.83
146.16
135.44
128.98
128.90
128.17
128.09
127.53
53.53
53.52
53.06
53.02
35.00
34.98
33.87
33.83
f1 (ppm)

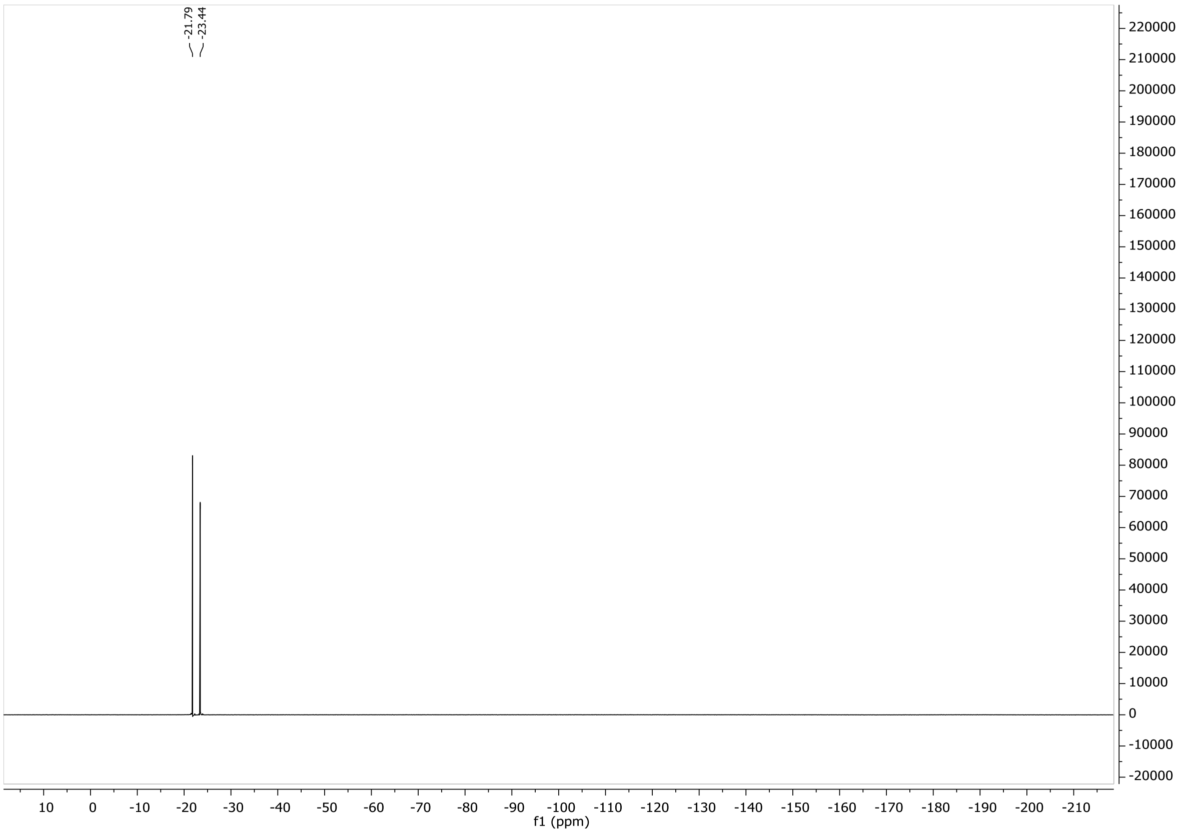
-21.79
-23.44
f1 (ppm)

**6c**

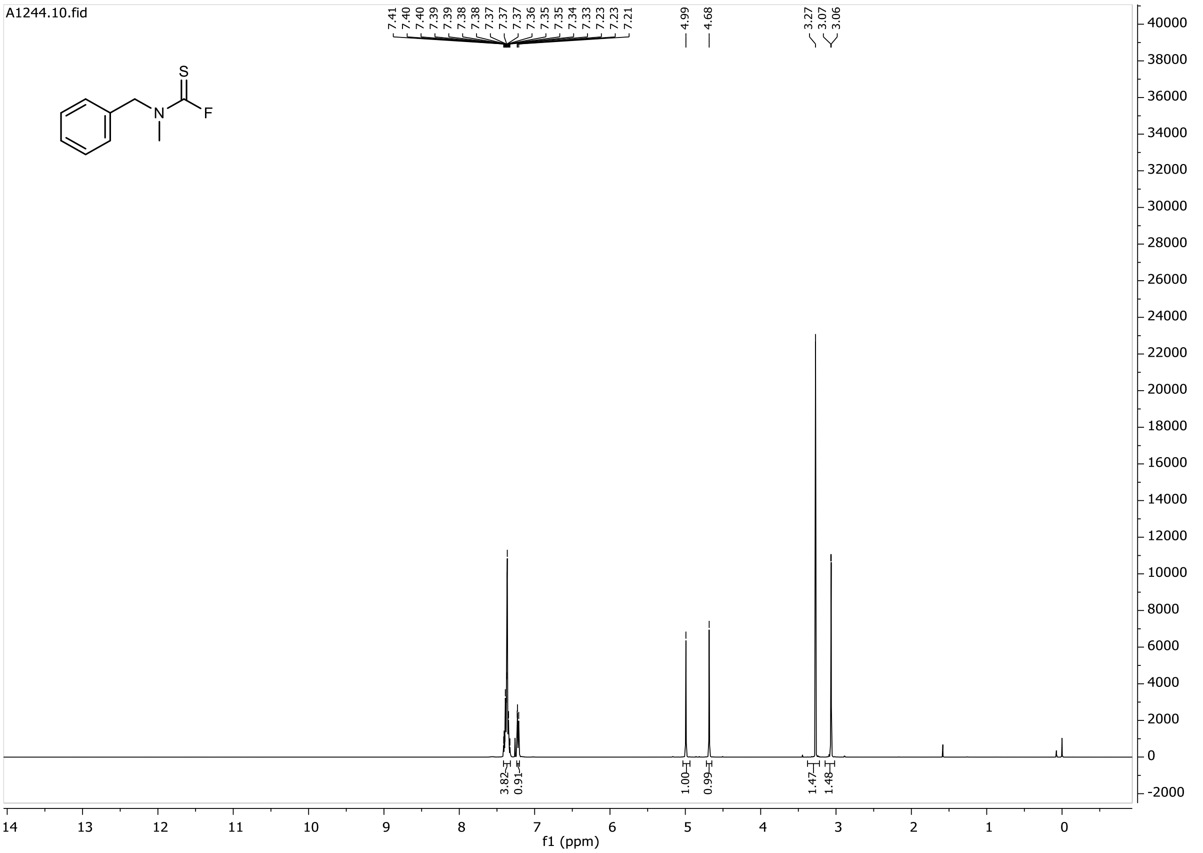
A1244.10.fid
7.41 7.40 7.40 7.39 7.39 7.38 7.38 7.37 7.37 7.37 7.37 7.36 7.35 7.35 7.34 7.33 7.23 7.23 7.21
4.99
4.68
3.27 3.07 3.06
S
N
F
3.82
0.91
1.00
0.99
1.47
1.48
f1 (ppm)

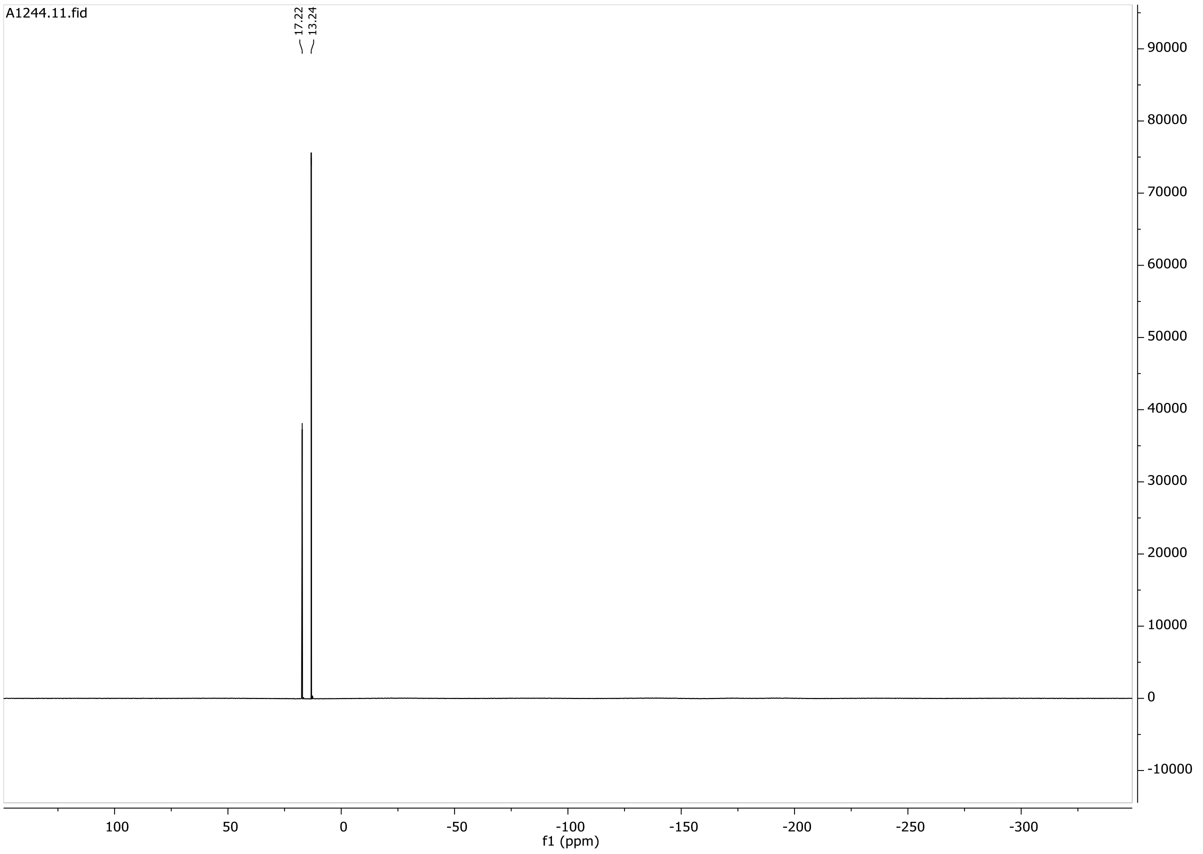
A1244.11.fid
17.22
13.24
f1 (ppm)

**6d**

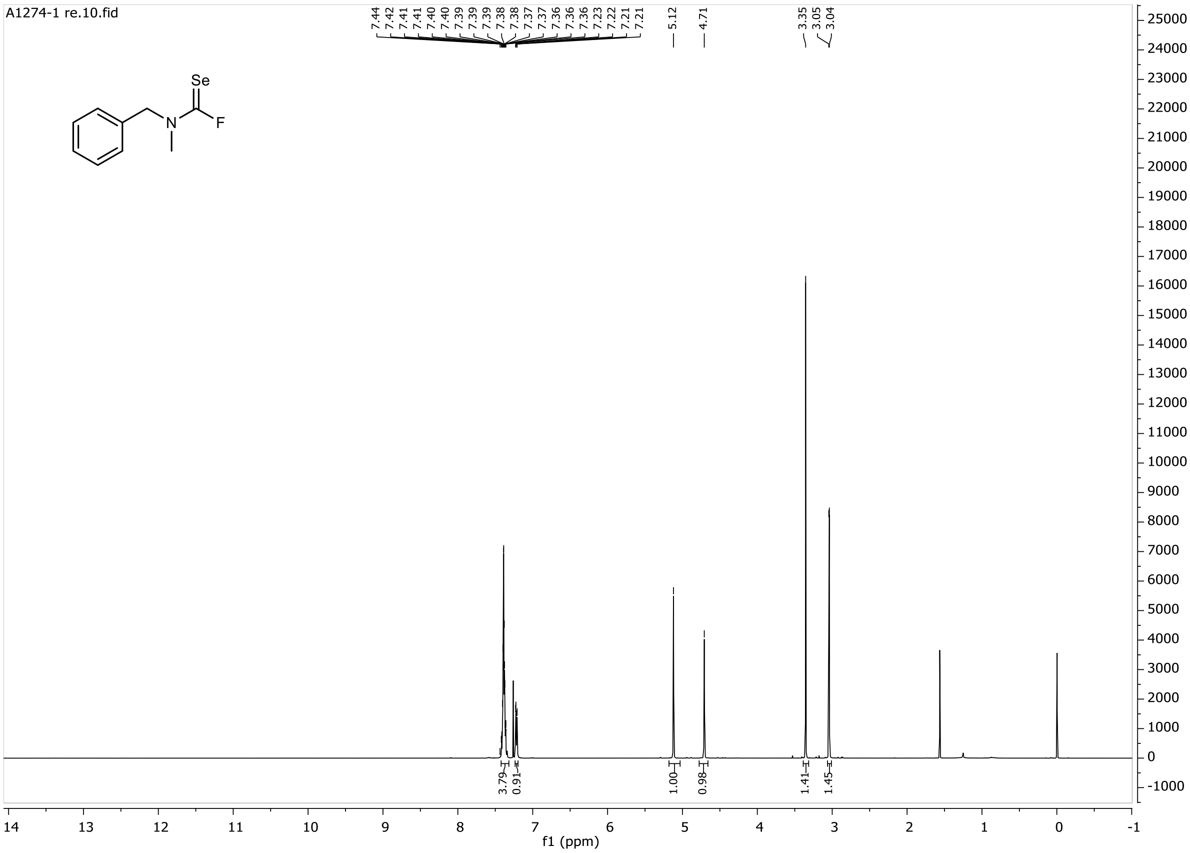
A1274-1 re.10.fid
7.44 7.42 7.41 7.41 7.40 7.40 7.39 7.39 7.39 7.38 7.38 7.37 7.37 7.36 7.36 7.36 7.23 7.22 7.21 7.21
5.12
4.71
3.35 3.05 3.04
Se
N
F
3.79 0.91 1.00 0.98 1.41 1.45
f1 (ppm)

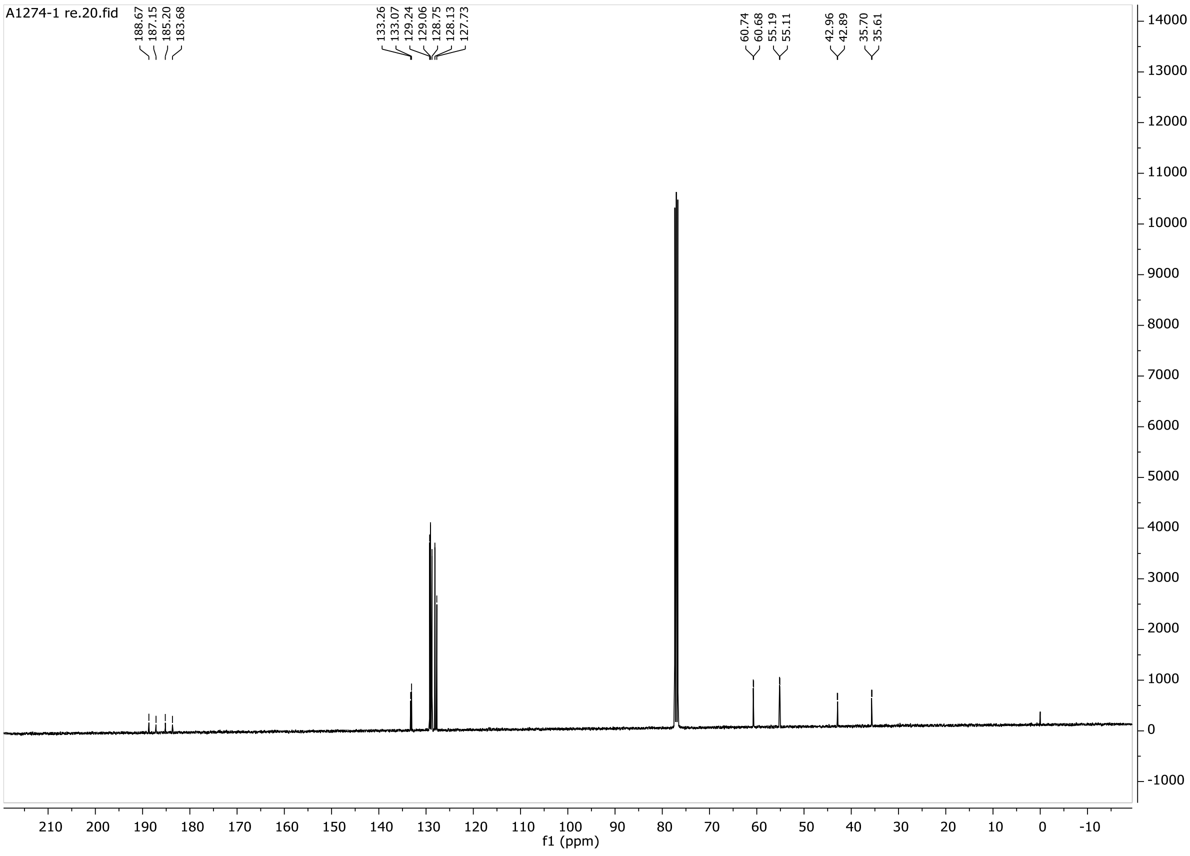
A1274-1 re.20.fid
188.67 187.15 185.20 183.68
133.26 133.07 129.24 129.06 128.75 128.13 127.73
60.74 60.68 55.19 55.11
42.96 42.89 35.70 35.61
f1 (ppm)

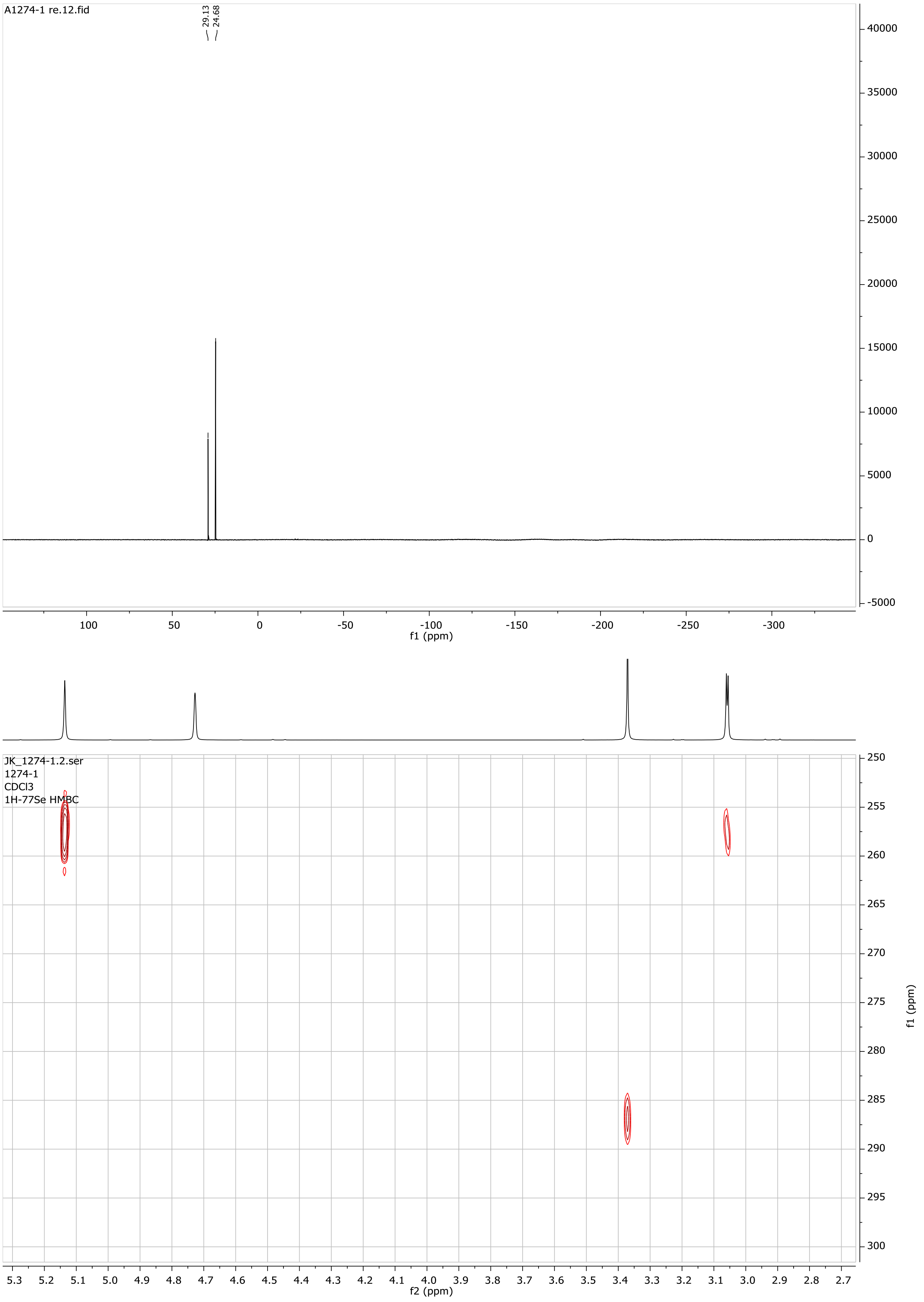
A1274-1 re.12.fid
29.13
24.68
f1 (ppm)
JK_1274-1.2.ser
1274-1
CDCl3
1H-77Se HMBC
f2 (ppm)
f1 (ppm)

**7**

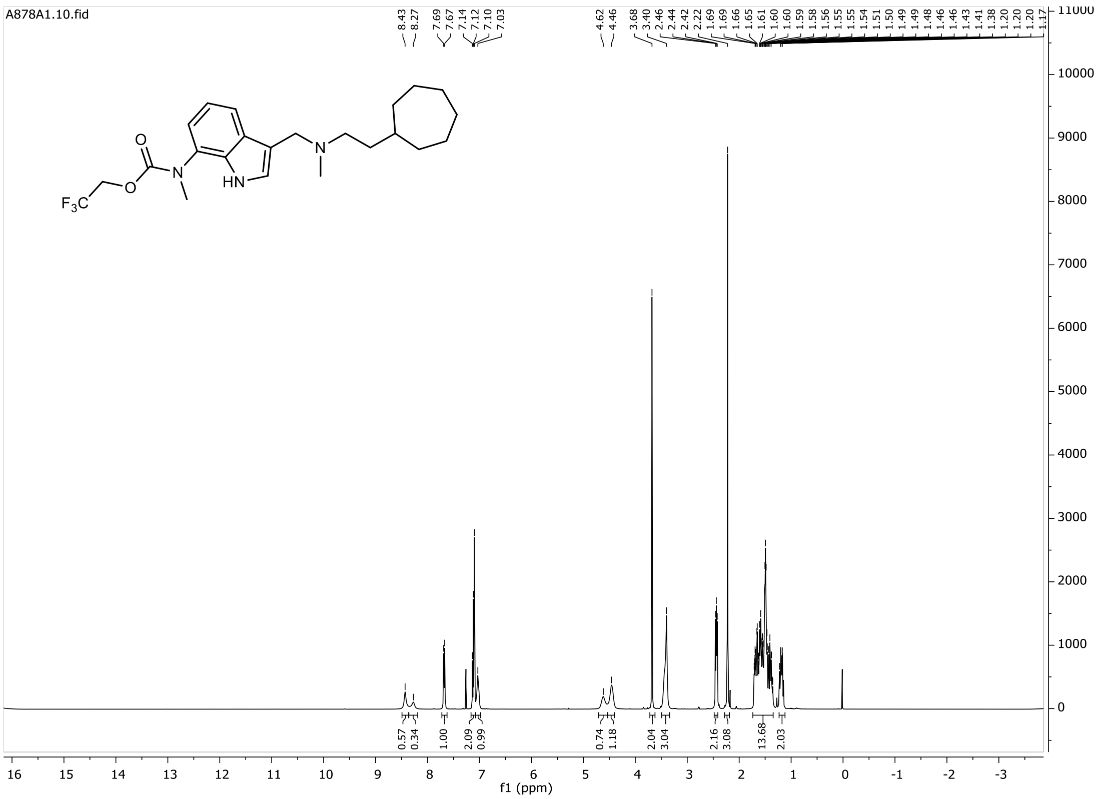

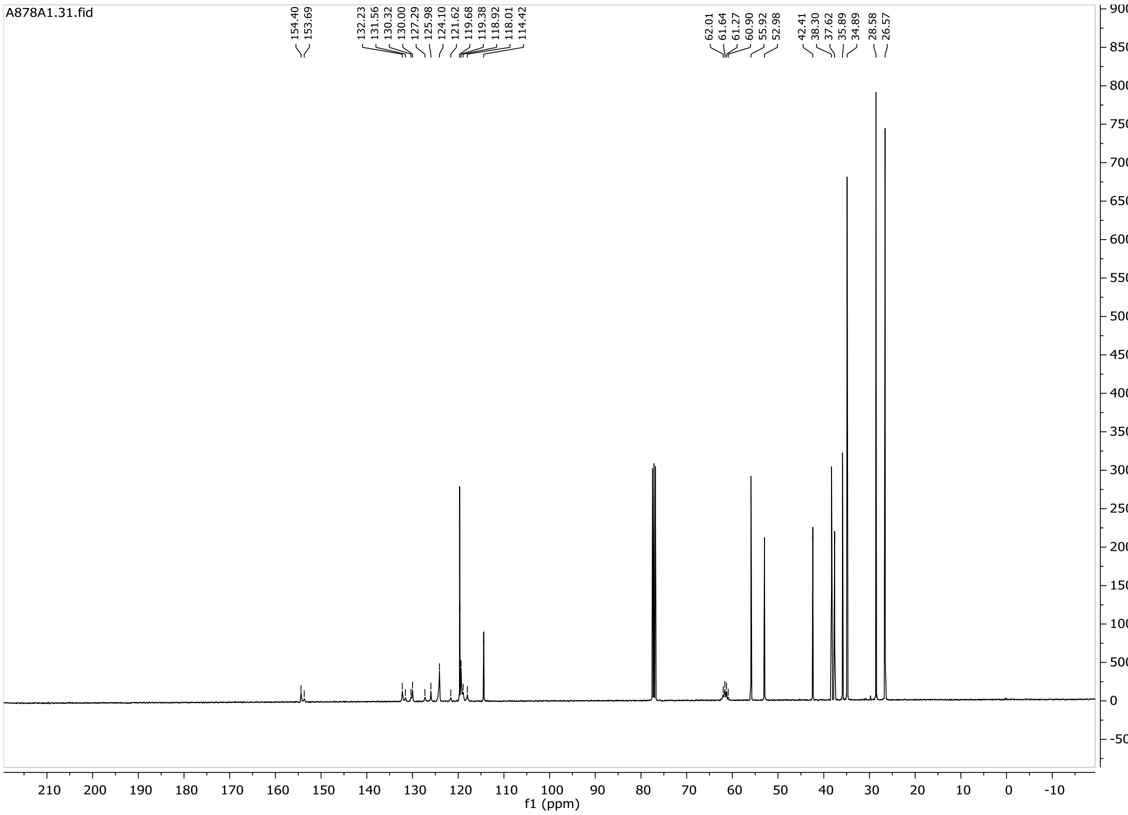

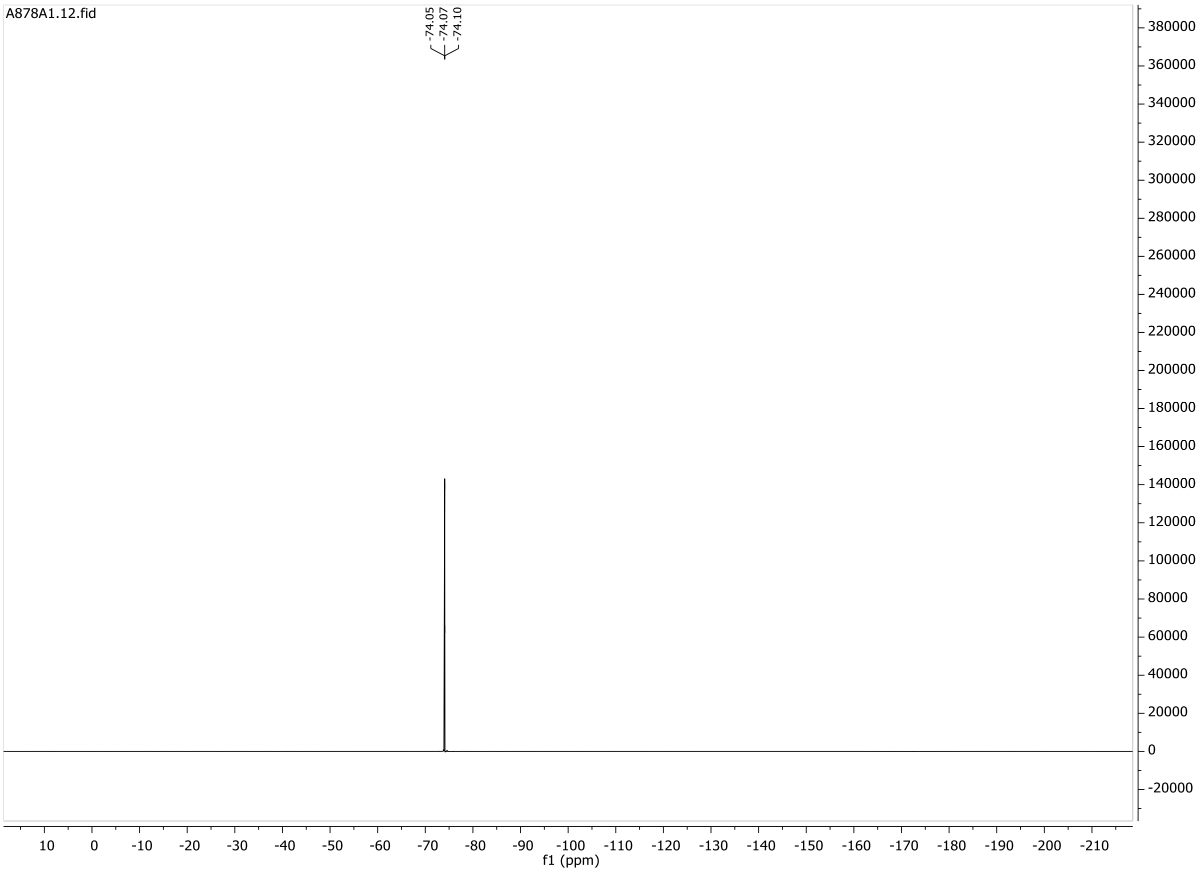

A878A1.12.fid
-74.05
-74.07
-74.10
380000
360000
340000
320000
300000
280000
260000
240000
220000
200000
180000
160000
140000
120000
100000
80000
60000
40000
20000
0
-20000
10 0 -10 -20 -30 -40 -50 -60 -70 -80 -90 -100 -110 -120 -130 -140 -150 -160 -170 -180 -190 -200 -210
f1 (ppm)

**8**

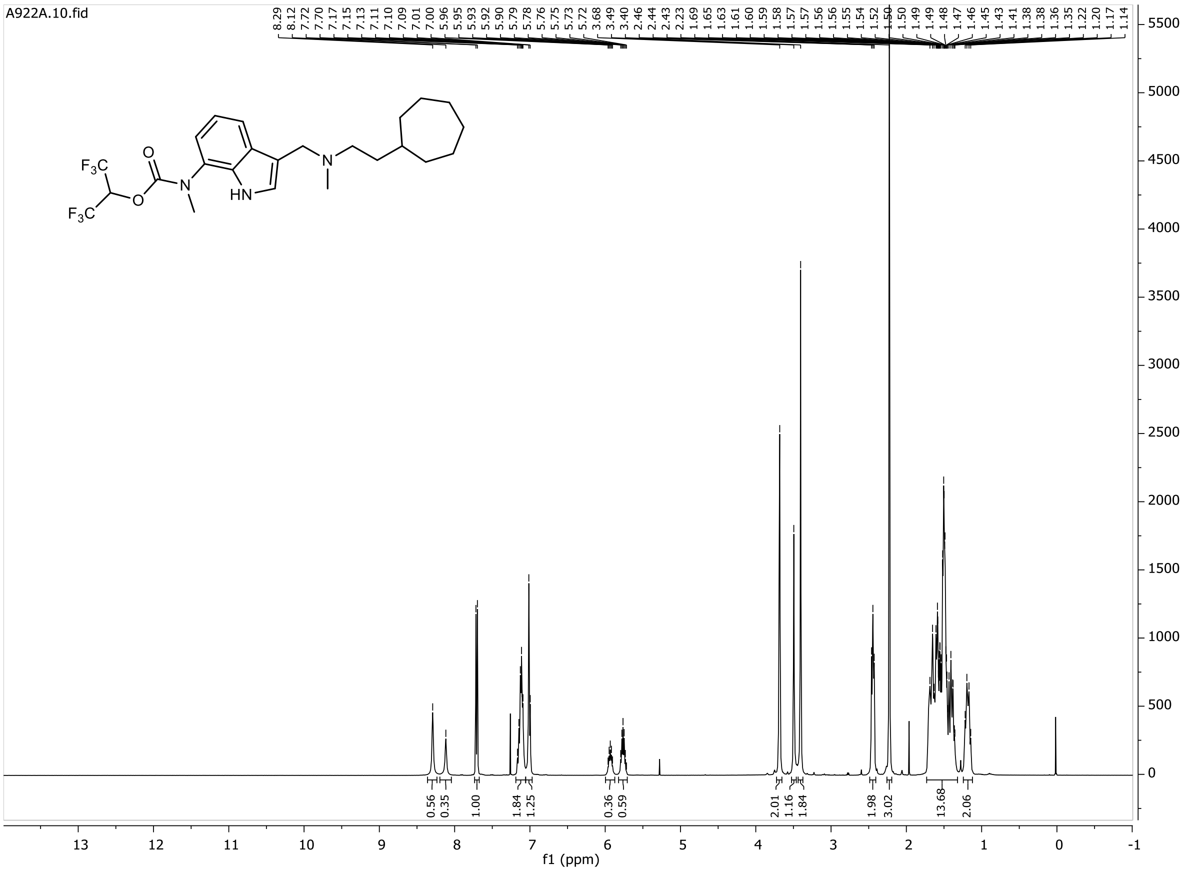
A922A.10.fid
8.29
8.12
7.72
7.70
7.17
7.15
7.13
7.11
7.10
7.09
7.01
7.00
5.96
5.95
5.93
5.92
5.90
5.79
5.78
5.76
5.75
5.73
5.72
3.68
3.49
3.40
2.46
2.44
2.43
2.23
1.69
1.65
1.63
1.61
1.60
1.59
1.58
1.57
1.57
1.56
1.56
1.55
1.54
1.52
1.50
1.49
1.49
1.48
1.47
1.46
1.45
1.43
1.41
1.38
1.38
1.36
1.35
1.22
1.20
1.17
1.14
F3C
F3C
O
O
N
HN
N
0.56
0.35
1.00
1.84
1.25
0.36
0.59
2.01
1.16
1.84
1.98
3.02
13.68
2.06
f1 (ppm)

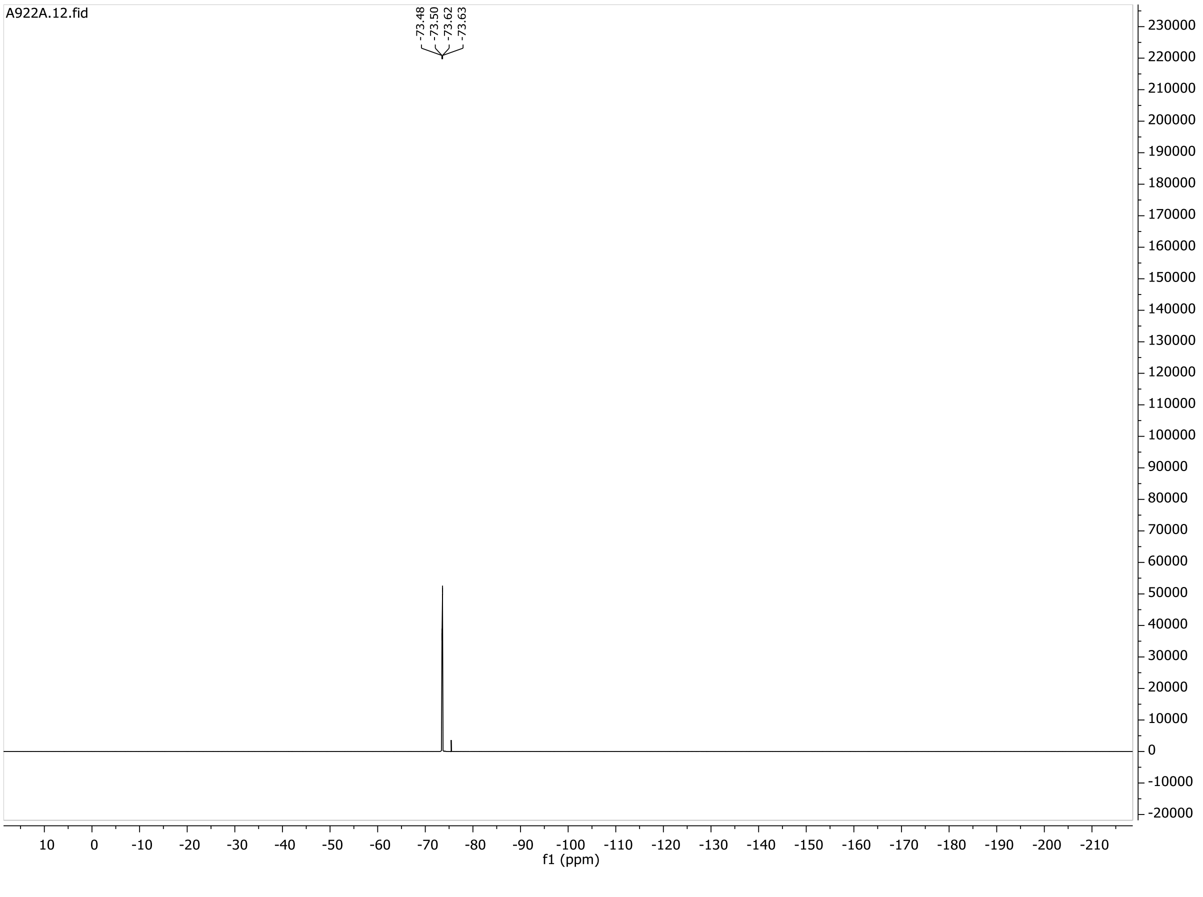
A922A.12.fid
-73.48
-73.50
-73.62
-73.63
f1 (ppm)

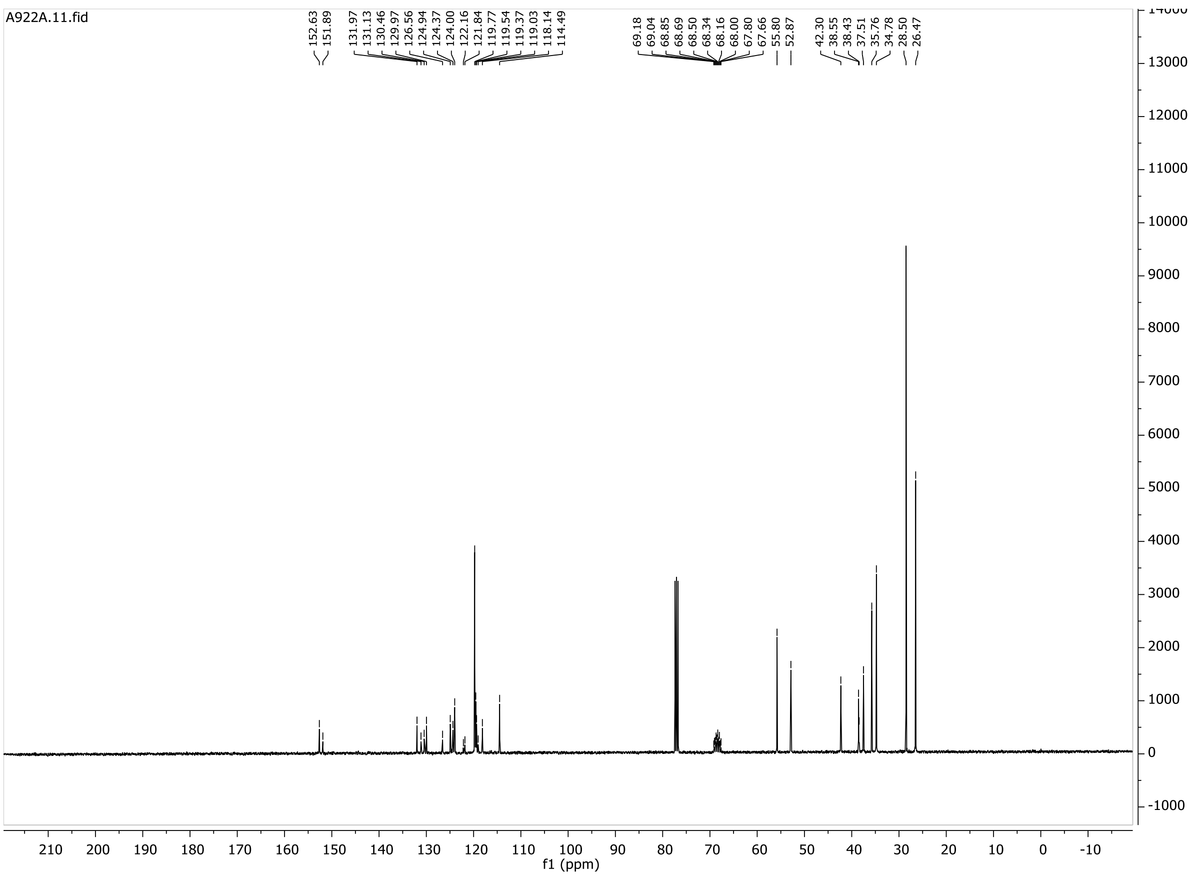
A922A.11.fid
152.63
151.89
131.97
131.13
130.46
129.97
126.56
124.94
124.37
124.00
122.16
121.84
119.77
119.54
119.37
119.03
118.14
114.49
69.18
69.04
68.85
68.69
68.50
68.34
68.16
68.00
67.80
67.66
55.80
52.87
42.30
38.55
38.43
37.51
35.76
34.78
28.50
26.47
f1 (ppm)

**8b**

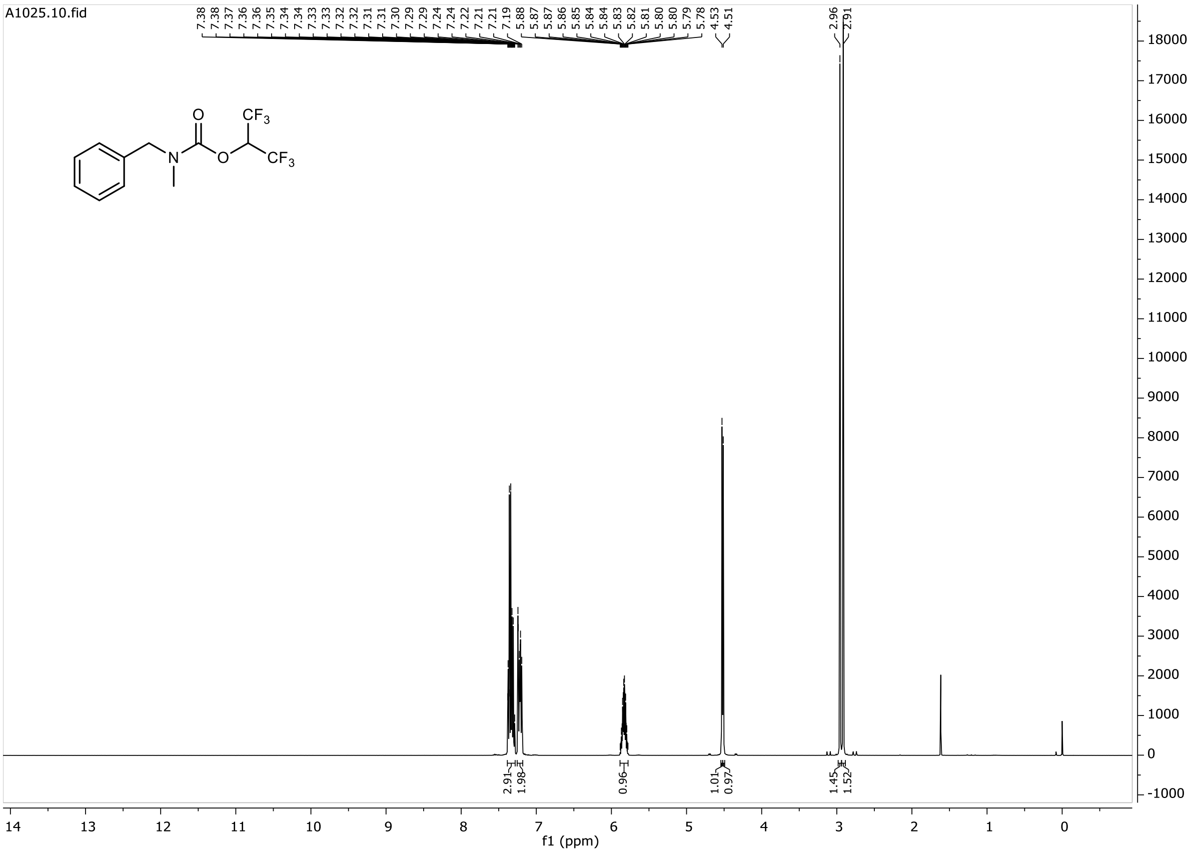
A1025.10.fid
7.38 7.38 7.37 7.36 7.36 7.35 7.34 7.34 7.33 7.33 7.32 7.32 7.31 7.31 7.30 7.29 7.29 7.24 7.24 7.22 7.21 7.21 7.19 5.88 5.87 5.87 5.86 5.85 5.84 5.84 5.83 5.82 5.81 5.80 5.80 5.79 5.78 4.53 4.51 2.96 2.91
O
N
O
$CF_3$
$CF_3$
2.91 1.98 0.96 1.01 0.97 1.45 1.52
f1 (ppm)

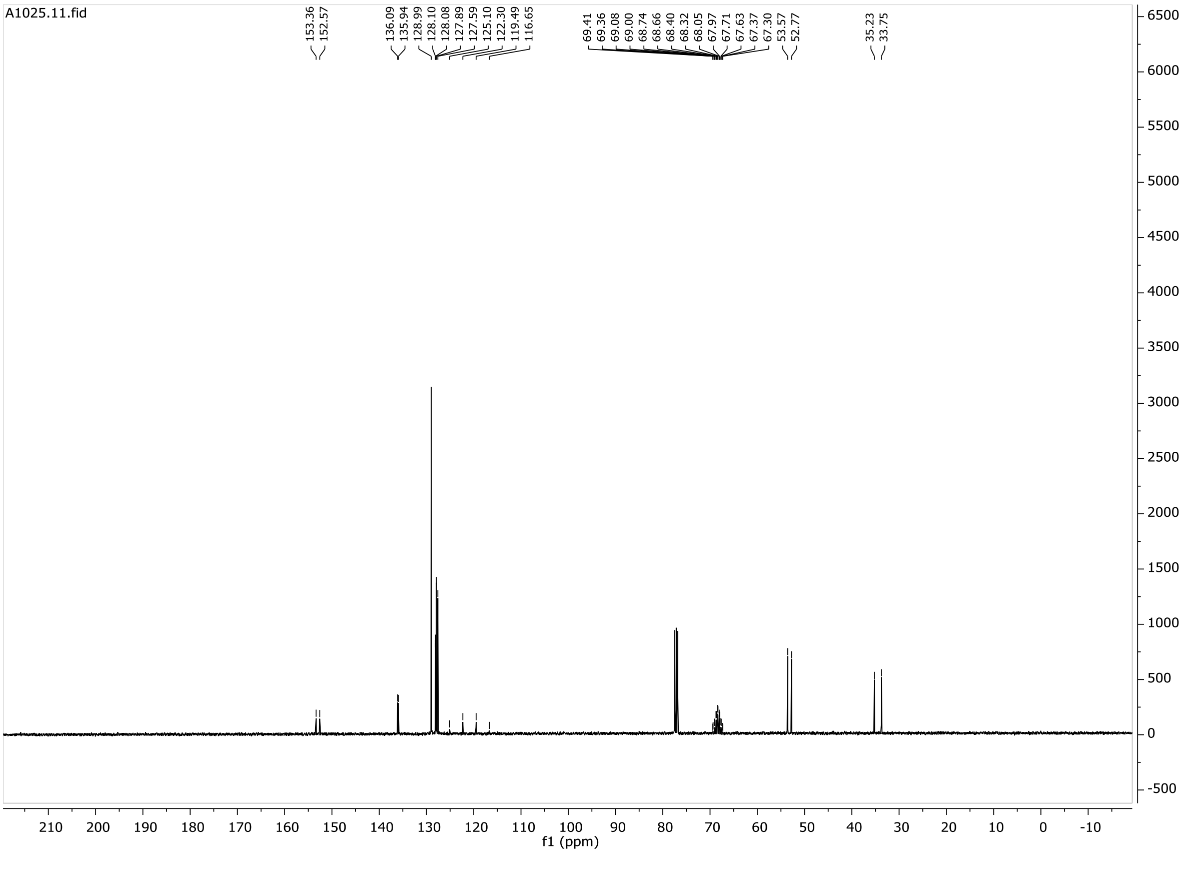
A1025.11.fid
153.36 152.57 136.09 135.94 128.99 128.10 128.08 127.89 127.59 125.10 122.30 119.49 116.65 69.41 69.36 69.08 69.00 68.74 68.66 68.40 68.32 68.05 67.97 67.71 67.63 67.37 67.30 53.57 52.77 35.23 33.75
f1 (ppm)

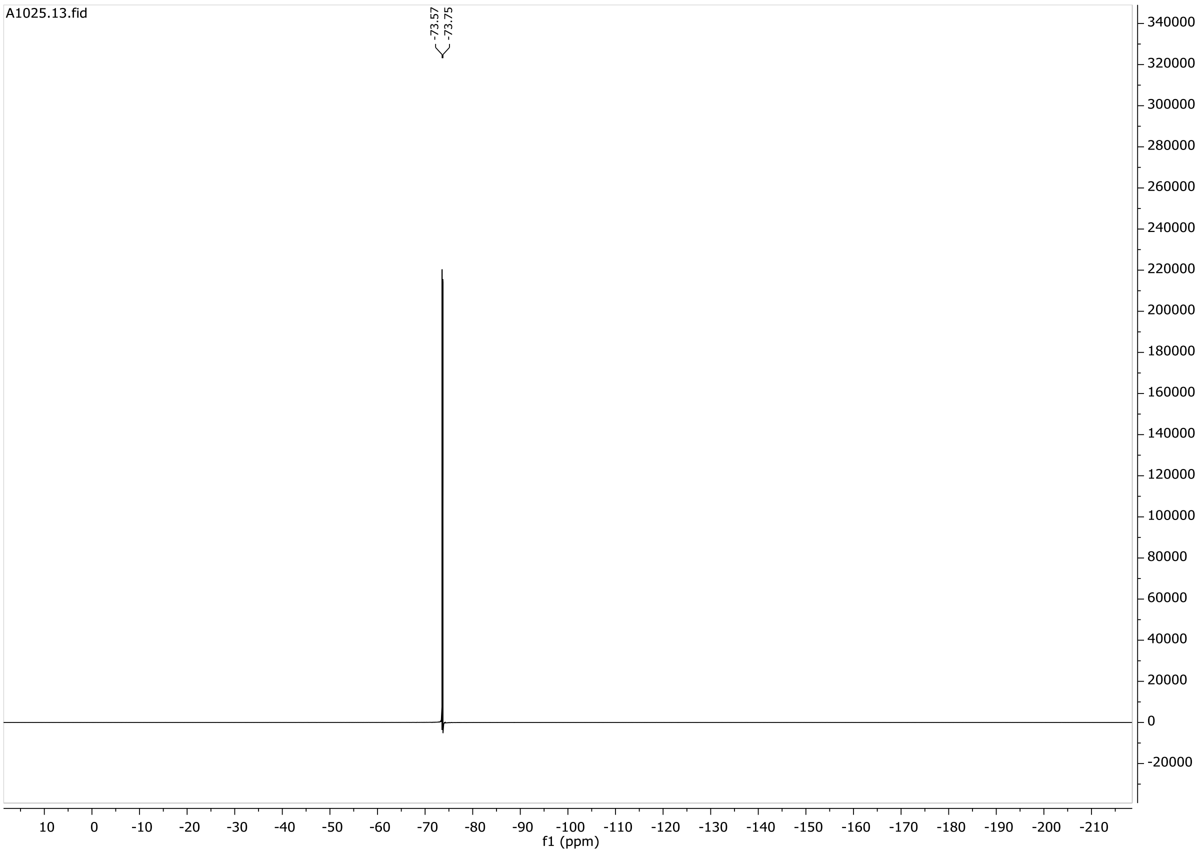

**9**

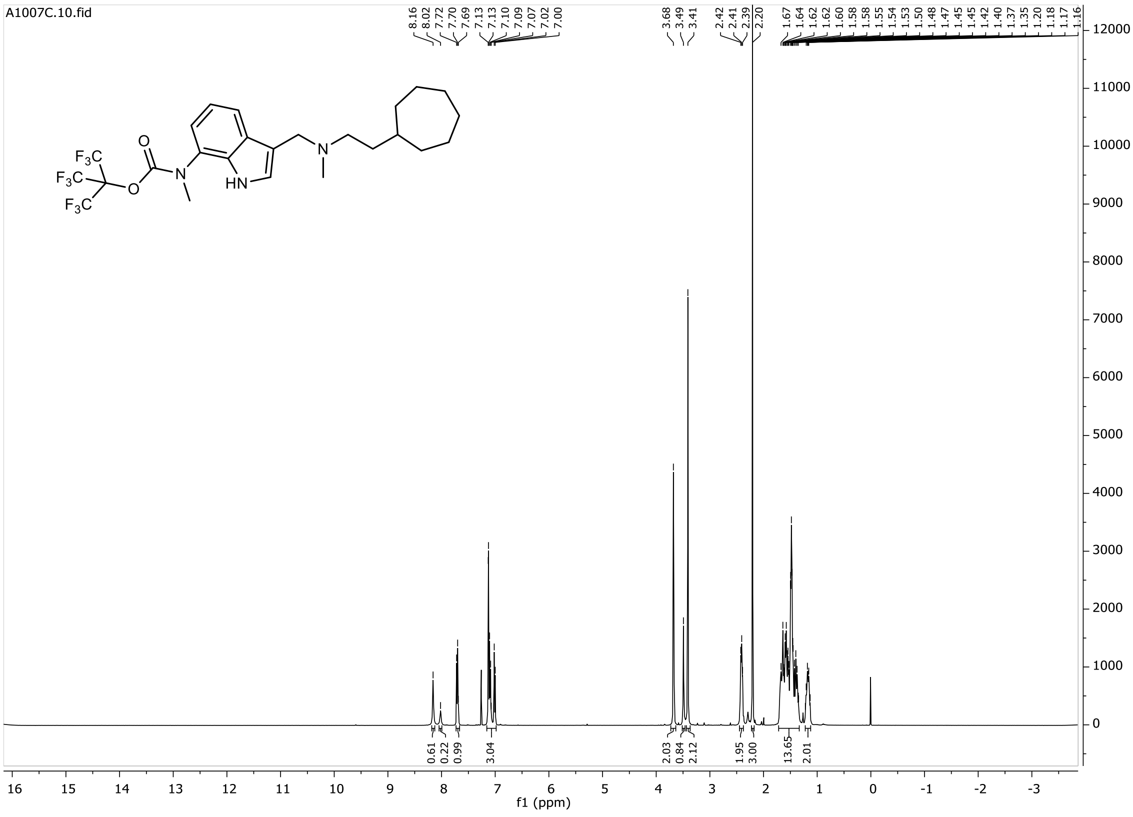

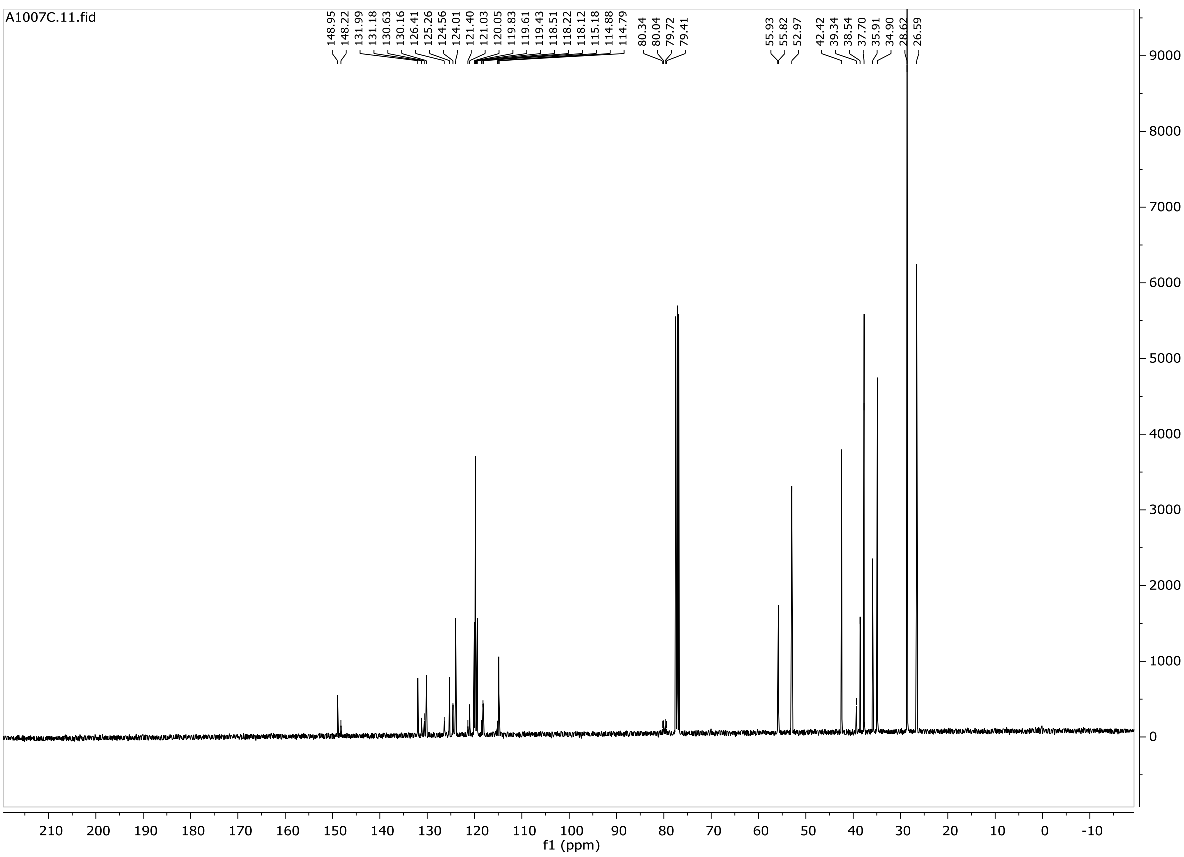

A1007C.11.fid
148.95
148.22
131.99
131.18
130.63
130.16
126.41
125.26
124.56
124.01
121.40
121.03
120.05
119.83
119.61
119.43
118.51
118.22
118.12
115.18
114.88
114.79
80.34
80.04
79.72
79.41
55.93
55.82
52.97
42.42
39.34
38.54
37.70
35.91
34.90
28.62
26.59
f1 (ppm)

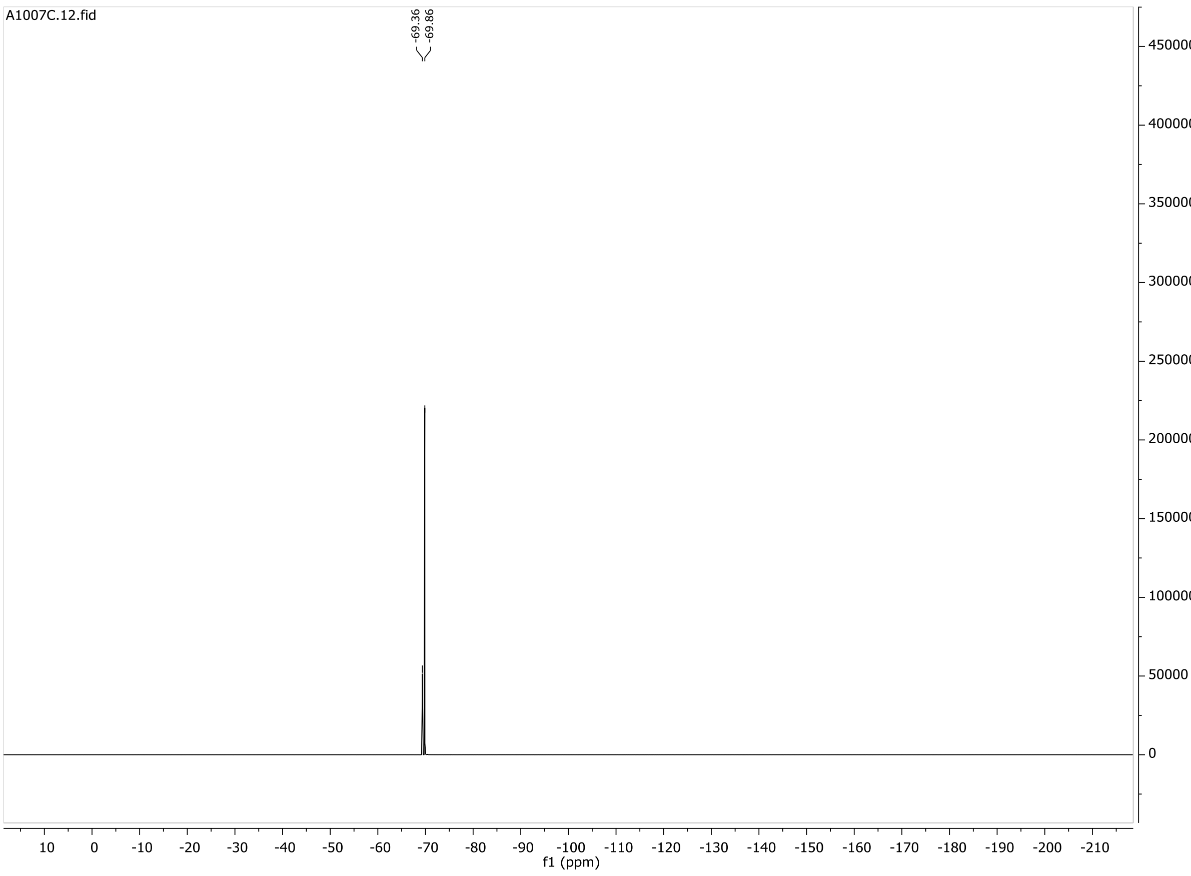

A1007C.12.fid
-69.36
-69.86
f1 (ppm)

**10**

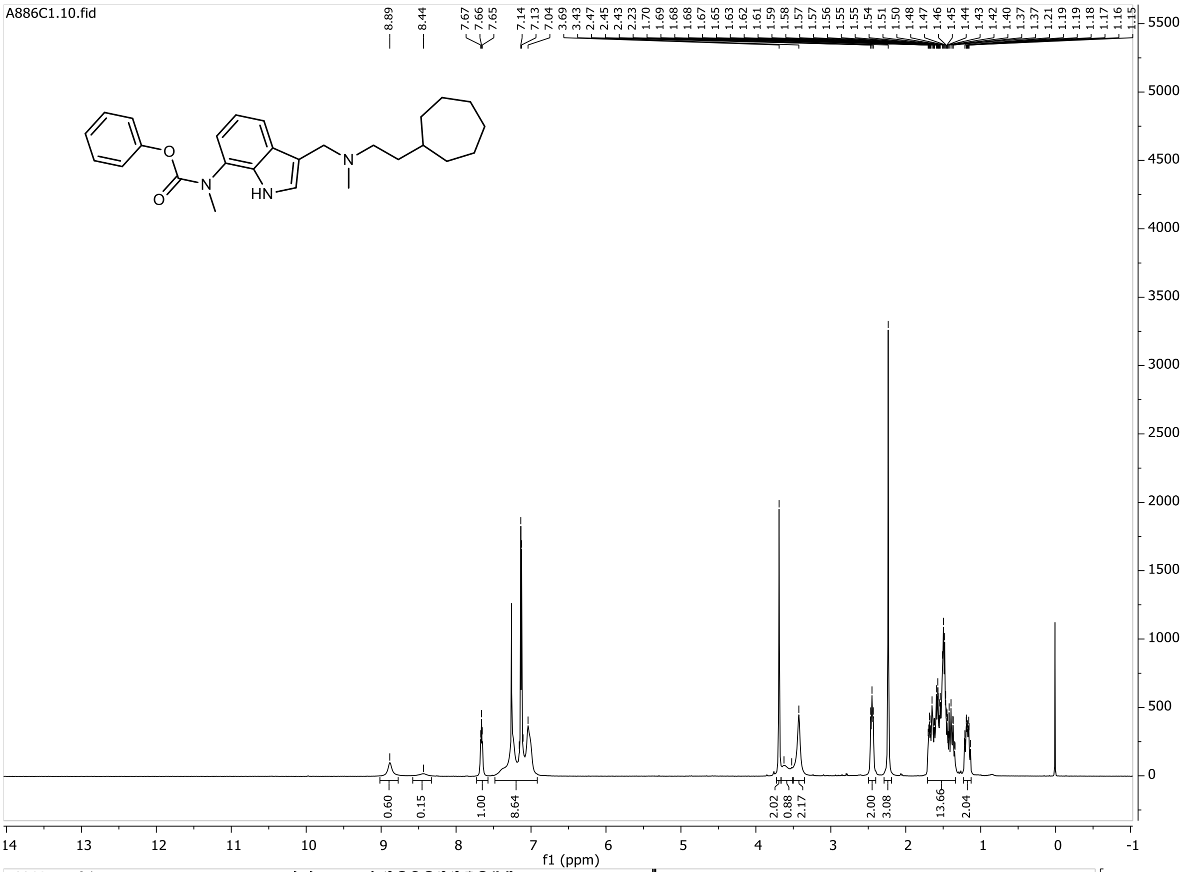

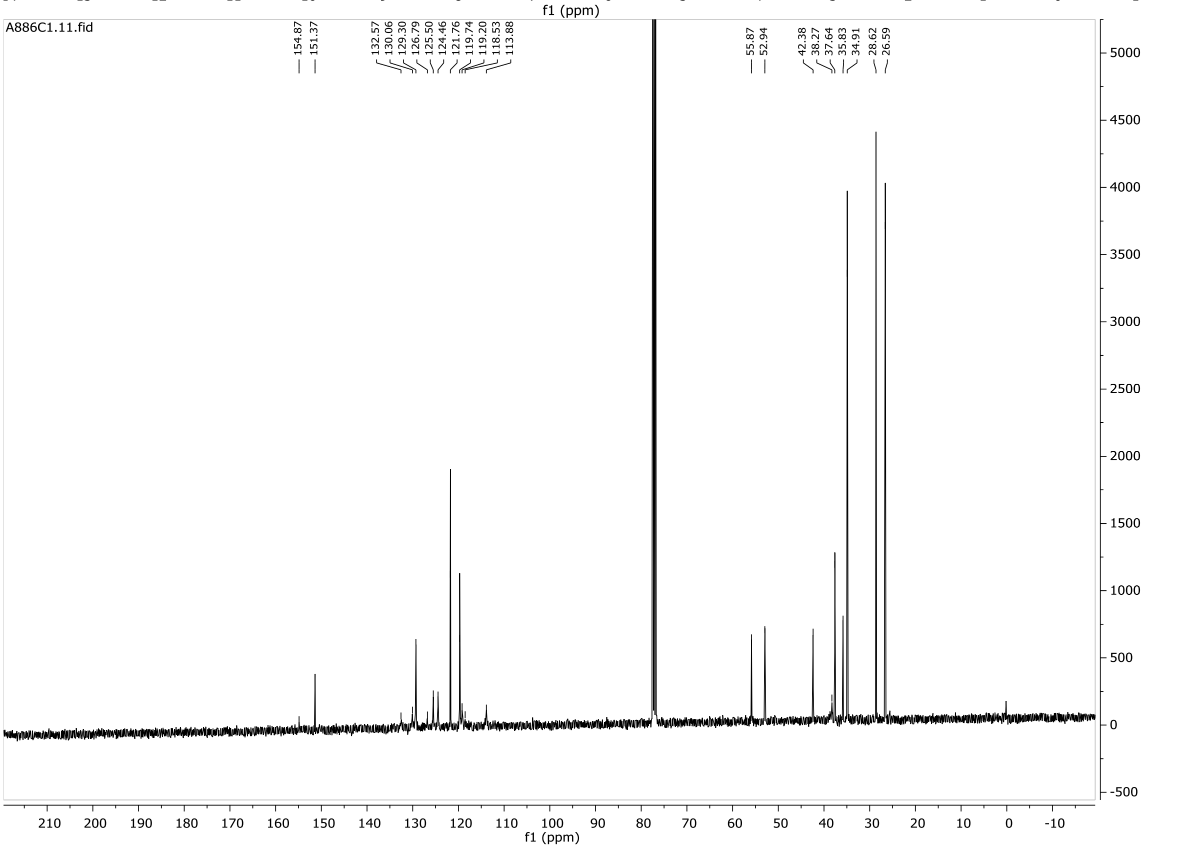

**11**

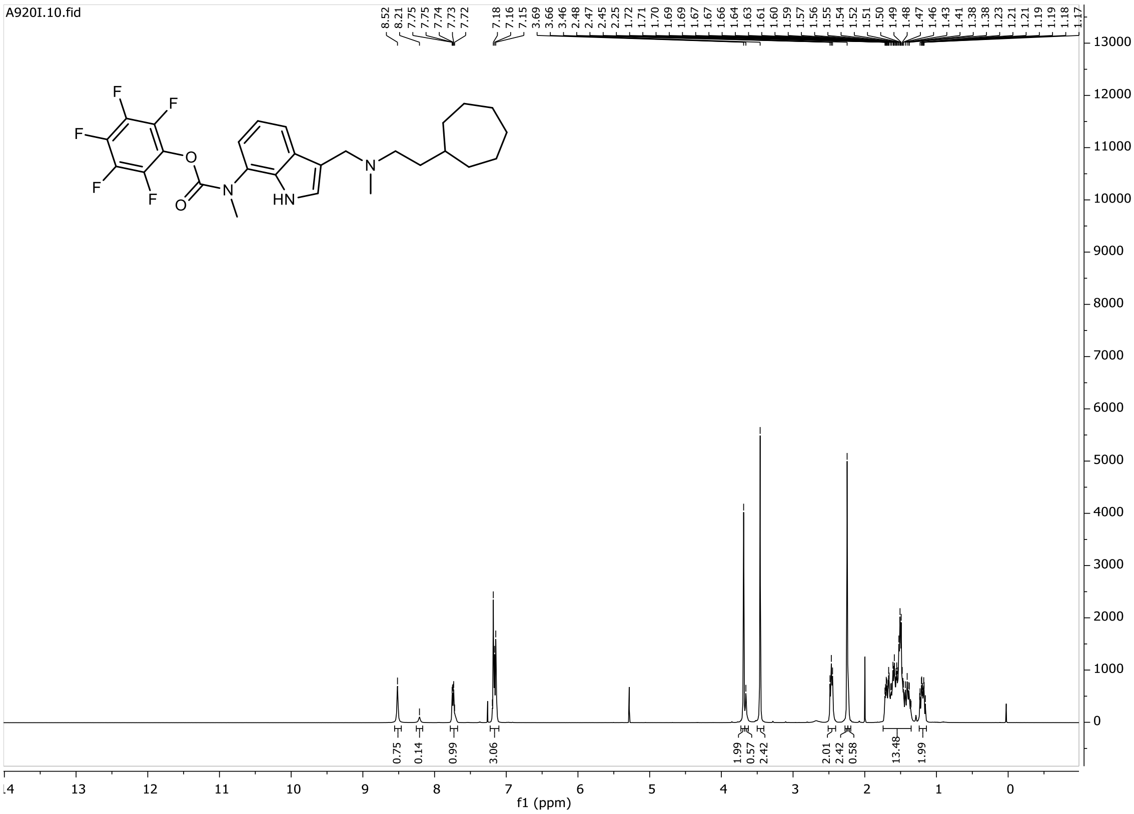
A920I.10.fid
8.52 8.21 7.75 7.75 7.74 7.73 7.72 7.18 7.16 7.15 3.69 3.66 3.46 2.48 2.47 2.45 2.25 1.72 1.71 1.70 1.69 1.69 1.67 1.67 1.66 1.64 1.63 1.61 1.60 1.59 1.57 1.56 1.55 1.54 1.52 1.51 1.50 1.49 1.48 1.47 1.46 1.43 1.41 1.38 1.38 1.23 1.21 1.21 1.19 1.19 1.18 1.17
0.75 0.14 0.99 3.06 1.99 0.57 2.42 2.01 2.42 0.58 13.48 1.99
f1 (ppm)

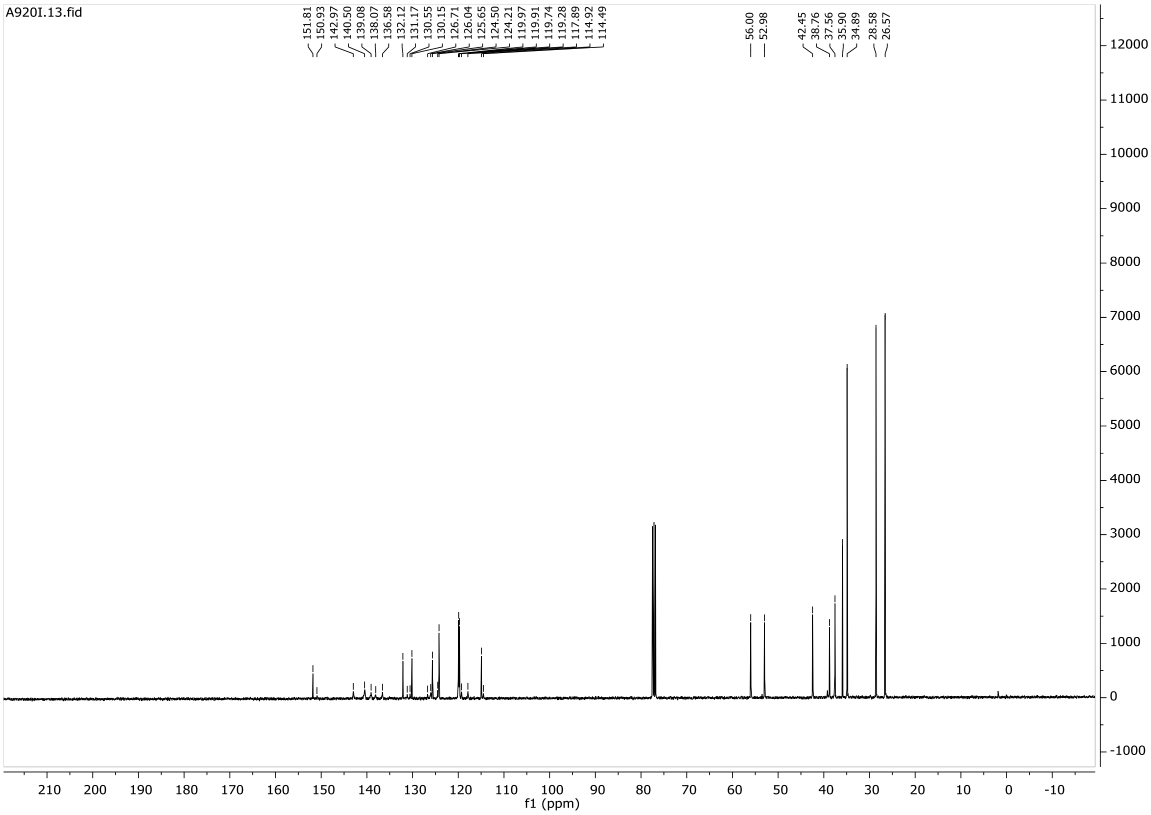
A920I.13.fid
151.81 150.93 142.97 140.50 139.08 138.07 136.58 132.12 131.17 130.55 130.15 126.71 126.04 125.65 124.50 124.21 119.97 119.91 119.74 119.28 117.89 114.92 114.49
56.00 52.98
42.45 38.76 37.56 35.90 34.89 28.58 26.57
f1 (ppm)

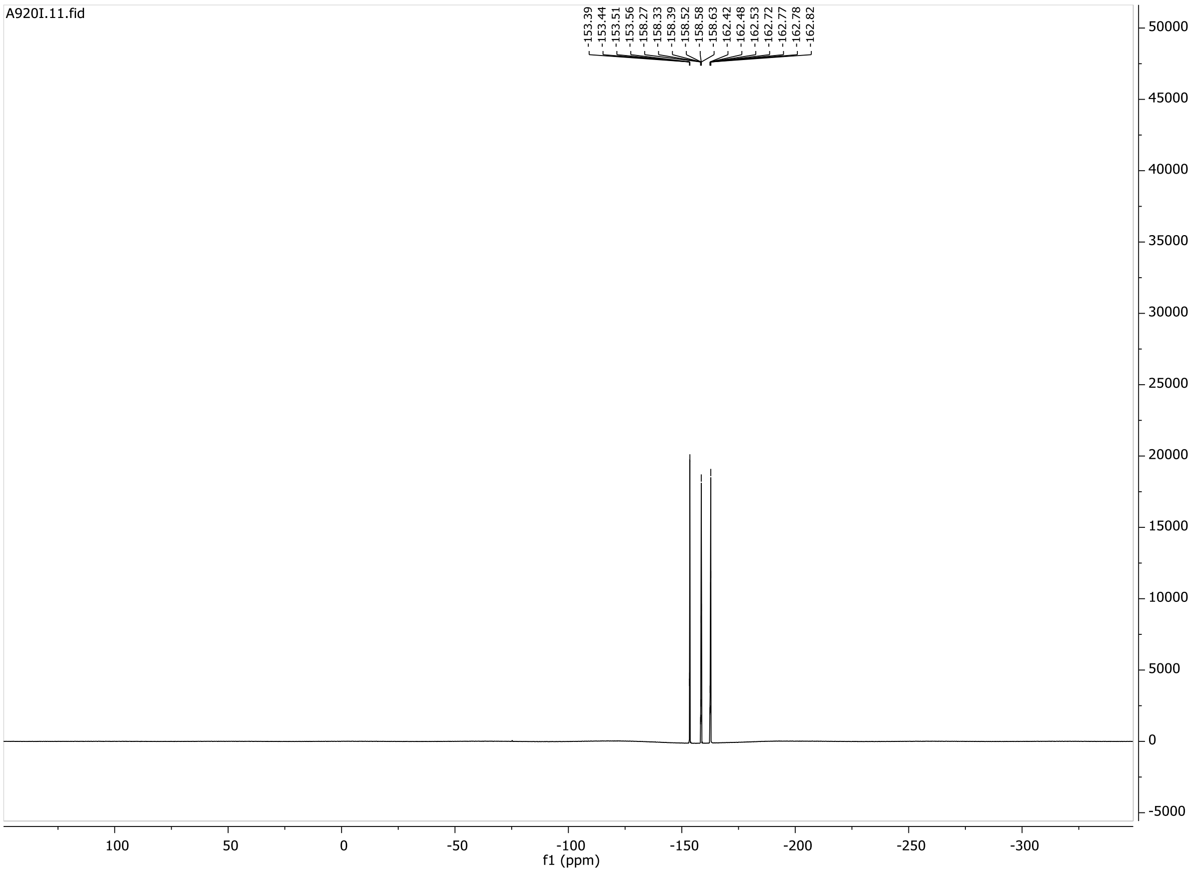
A920I.11.fid
-153.39
-153.44
-153.51
-153.56
-158.27
-158.33
-158.39
-158.52
-158.58
-158.63
-162.42
-162.48
-162.53
-162.72
-162.77
-162.78
-162.82
50000
45000
40000
35000
30000
25000
20000
15000
10000
5000
0
-5000
100
50
0
-50
-100
-150
-200
-250
-300
f1 (ppm)

**12**

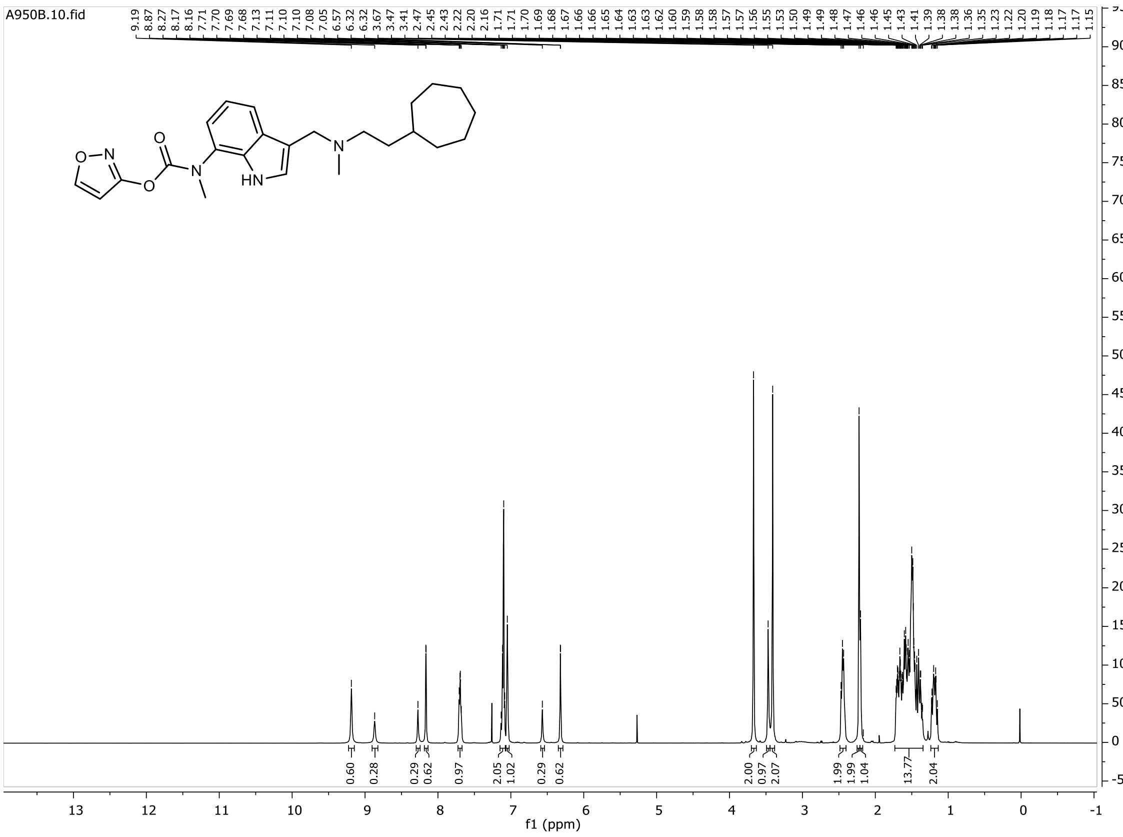

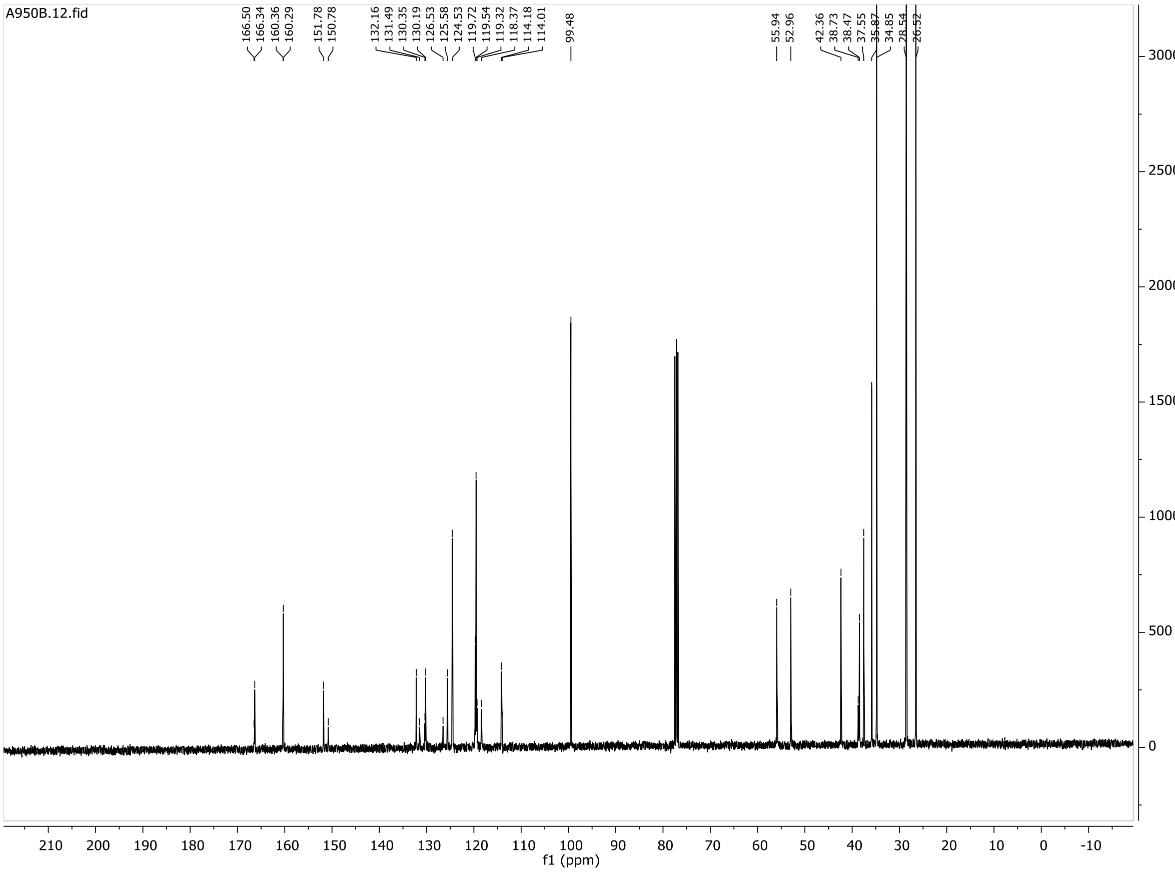

**13**

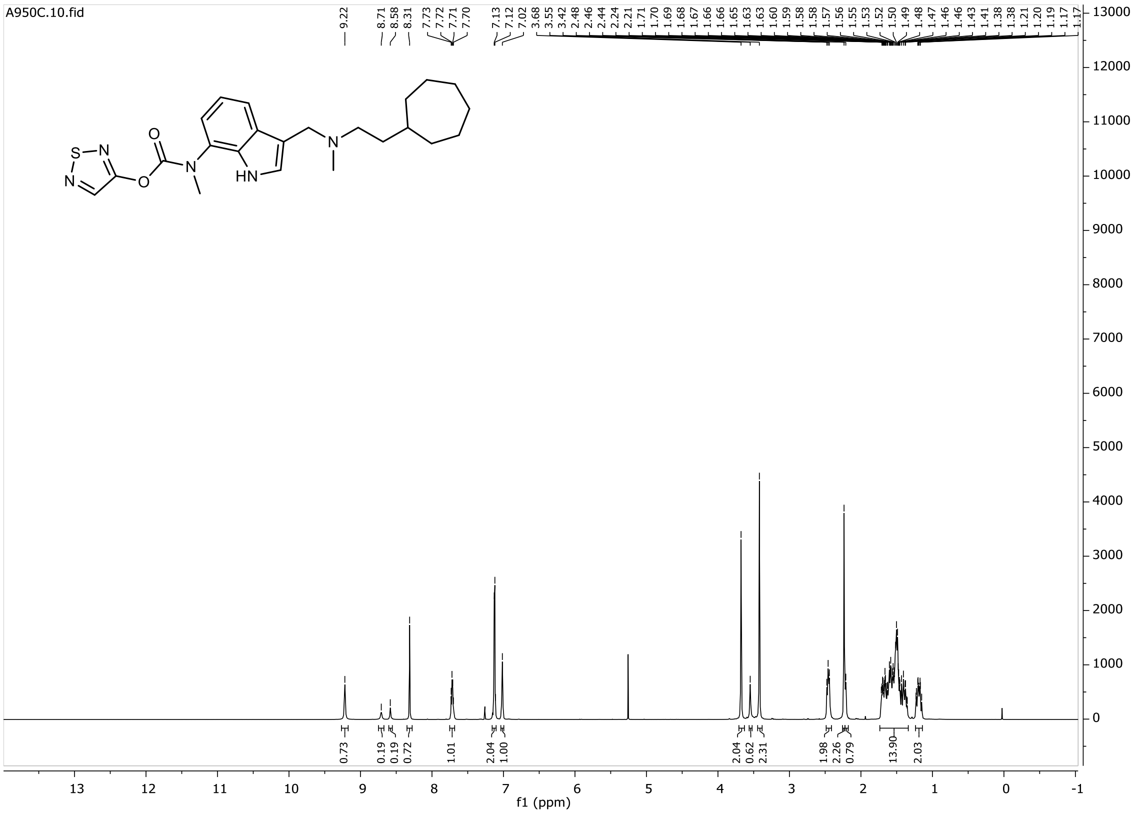

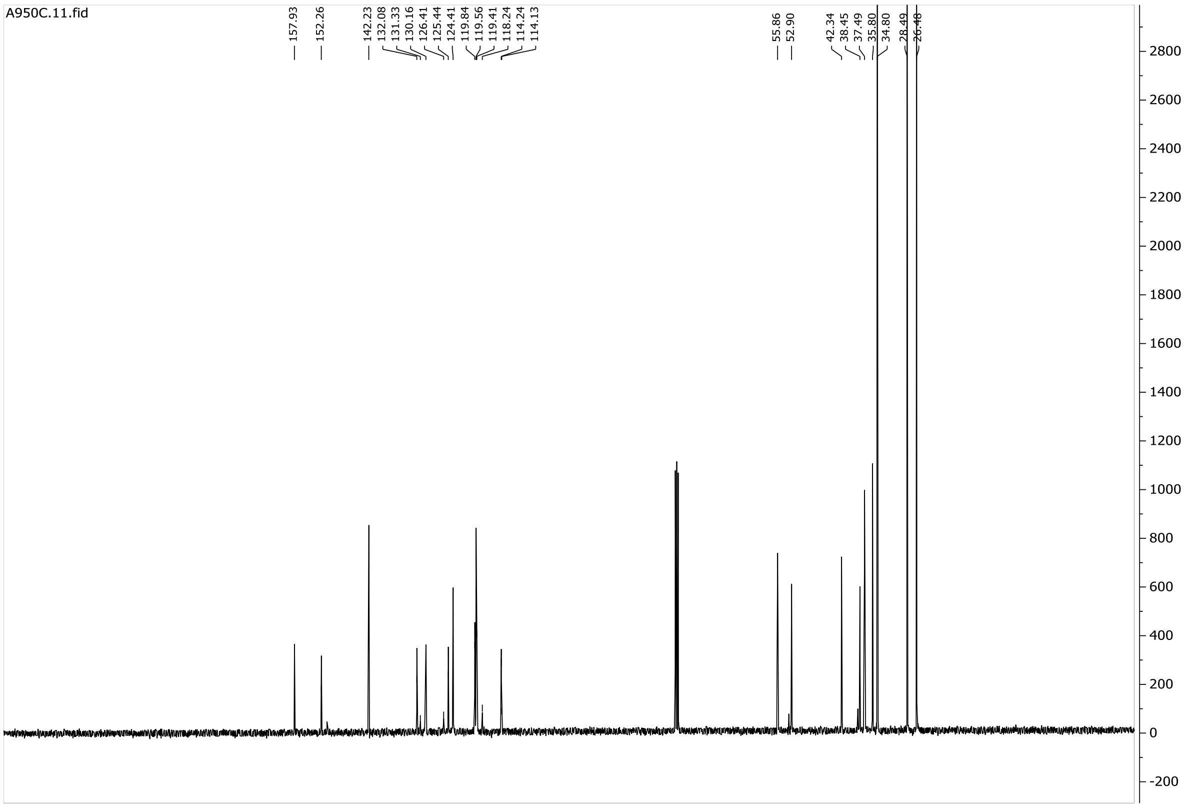

**14**

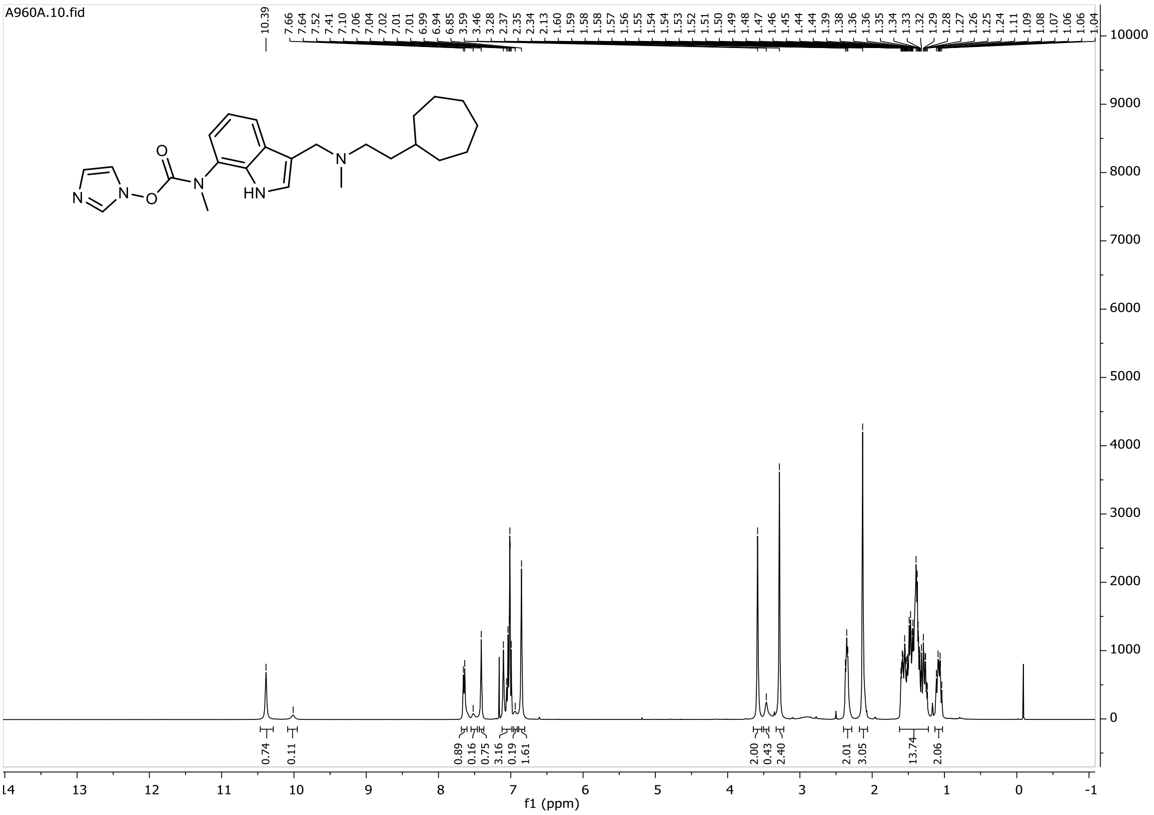

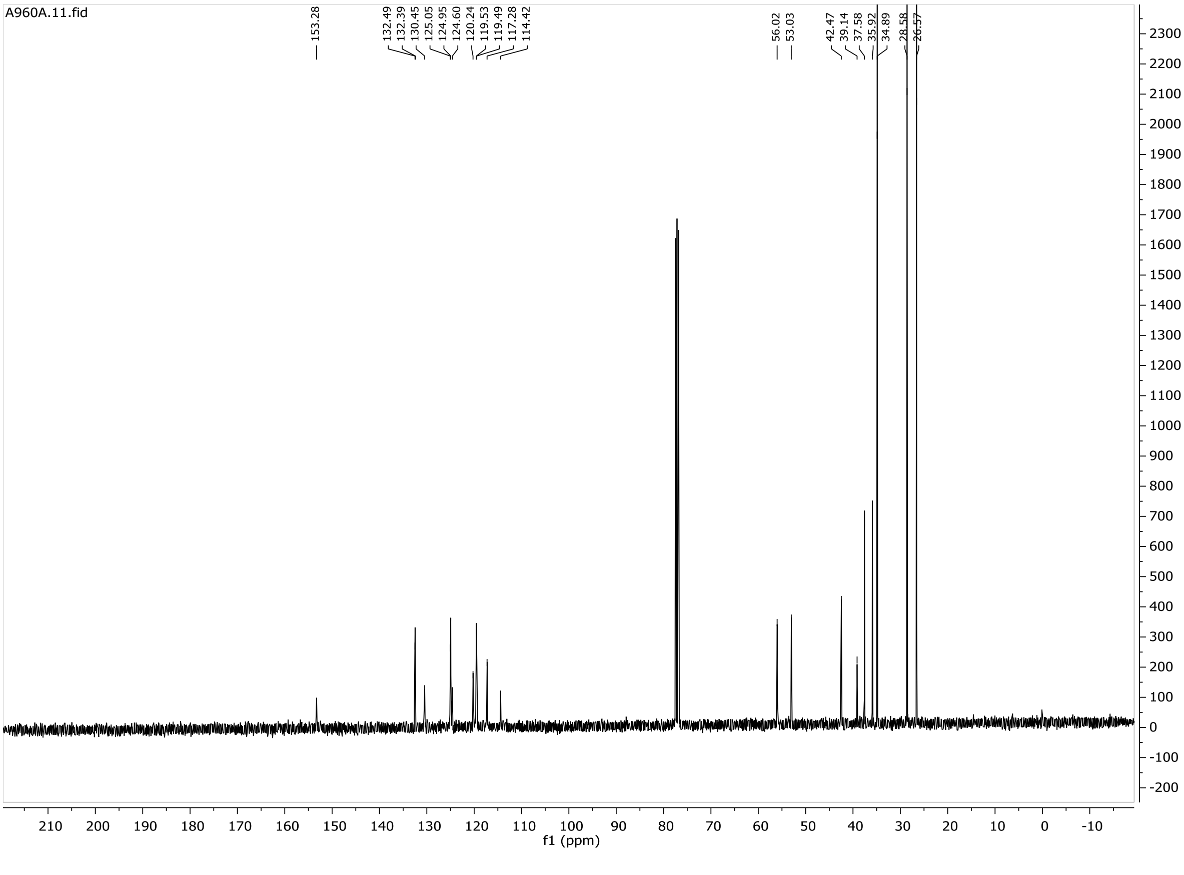

**15**

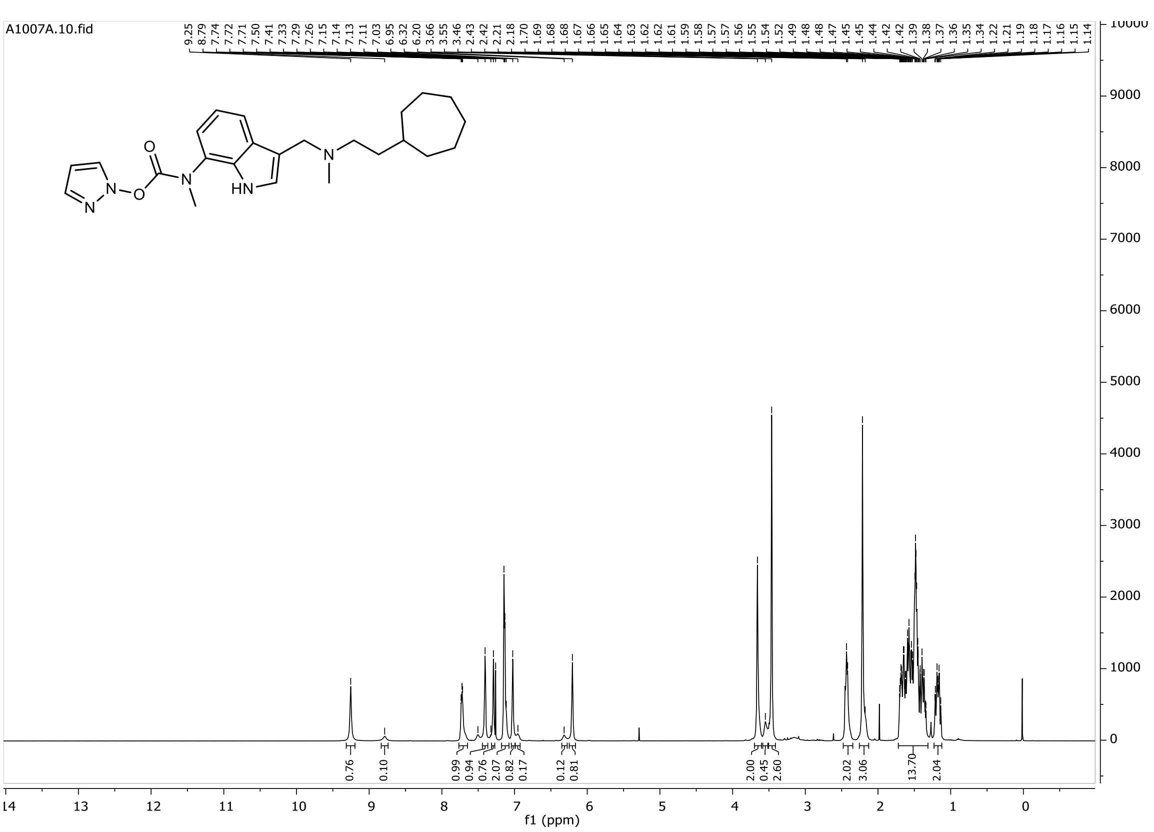

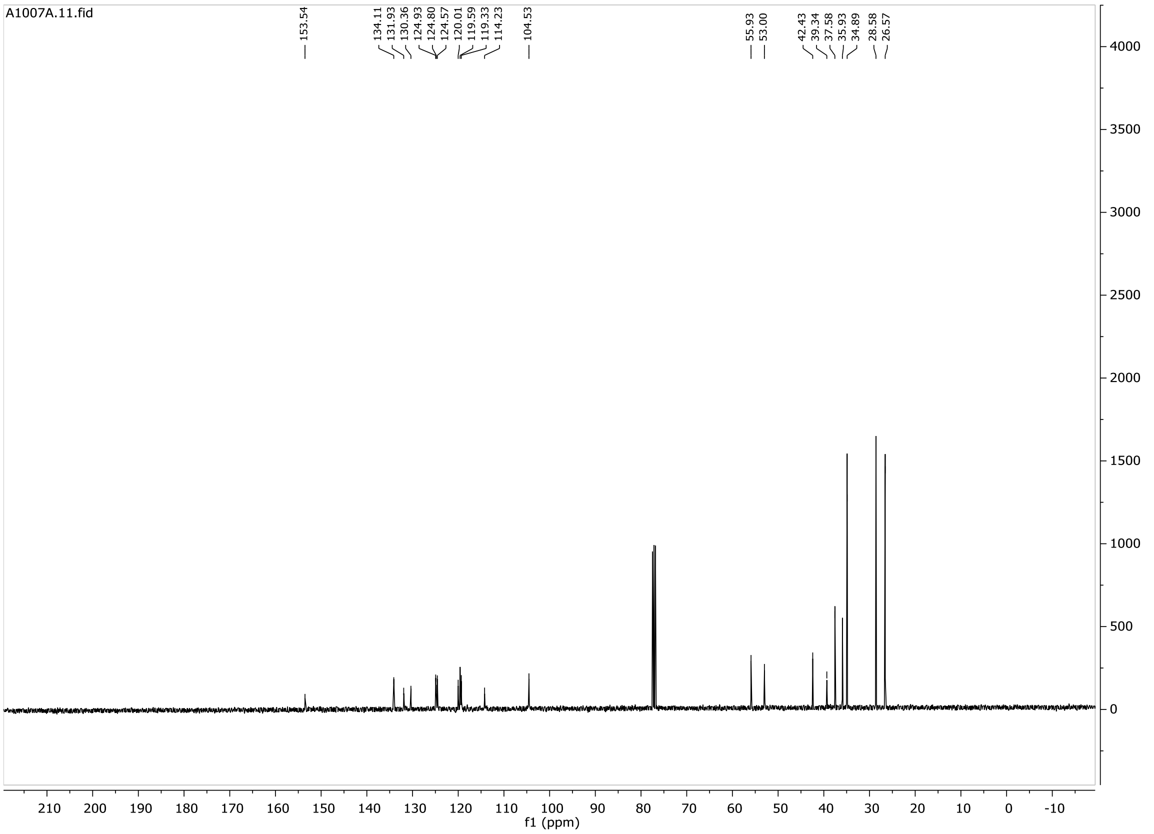

# 16

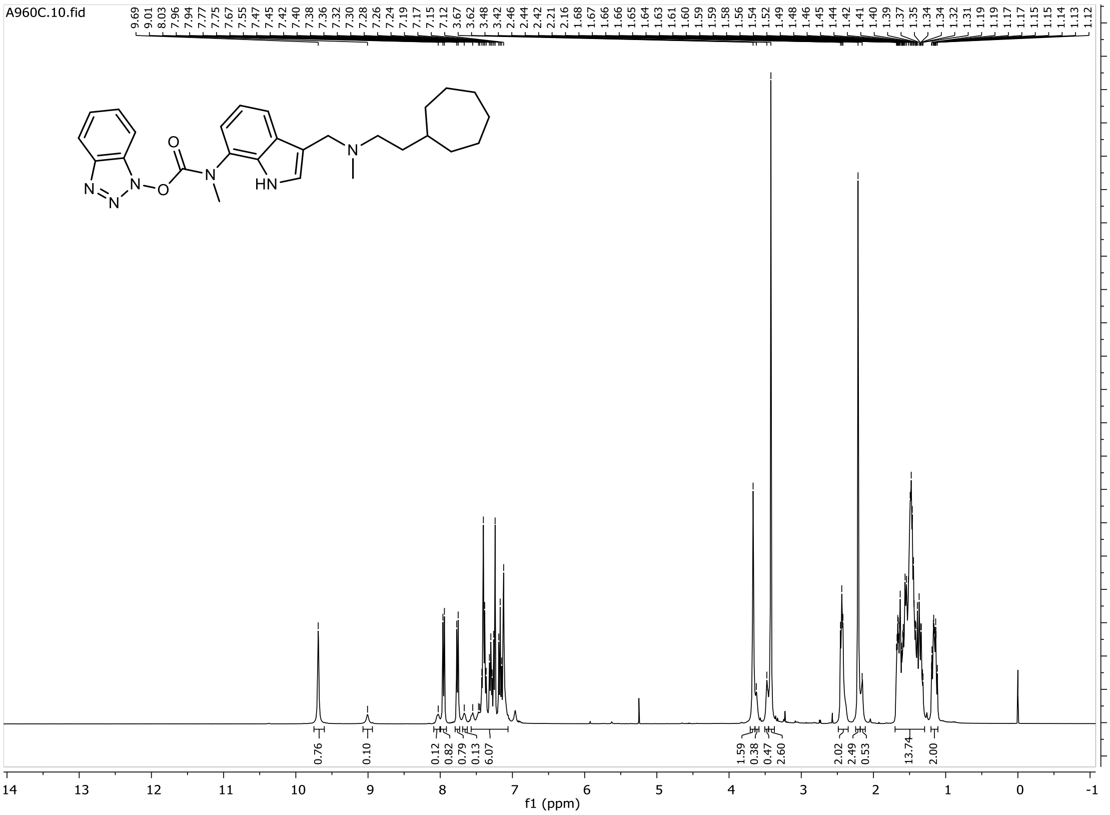
A960C.10.fid
9.69 9.01 8.03 7.96 7.94 7.77 7.75 7.67 7.55 7.47 7.45 7.42 7.40 7.38 7.36 7.32 7.30 7.28 7.26 7.24 7.19 7.17 7.15 7.12 3.67 3.62 3.48 3.42 2.46 2.44 2.42 2.21 2.16 1.68 1.67 1.66 1.65 1.64 1.63 1.61 1.60 1.59 1.59 1.58 1.56 1.54 1.52 1.49 1.48 1.46 1.45 1.44 1.42 1.41 1.40 1.39 1.37 1.35 1.34 1.34 1.32 1.31 1.19 1.19 1.17 1.17 1.15 1.15 1.14 1.13 1.12
0.76 0.10 0.12 0.82 0.79 0.13 6.07 1.59 0.38 0.47 2.60 2.02 2.49 0.53 13.74 2.00
f1 (ppm)

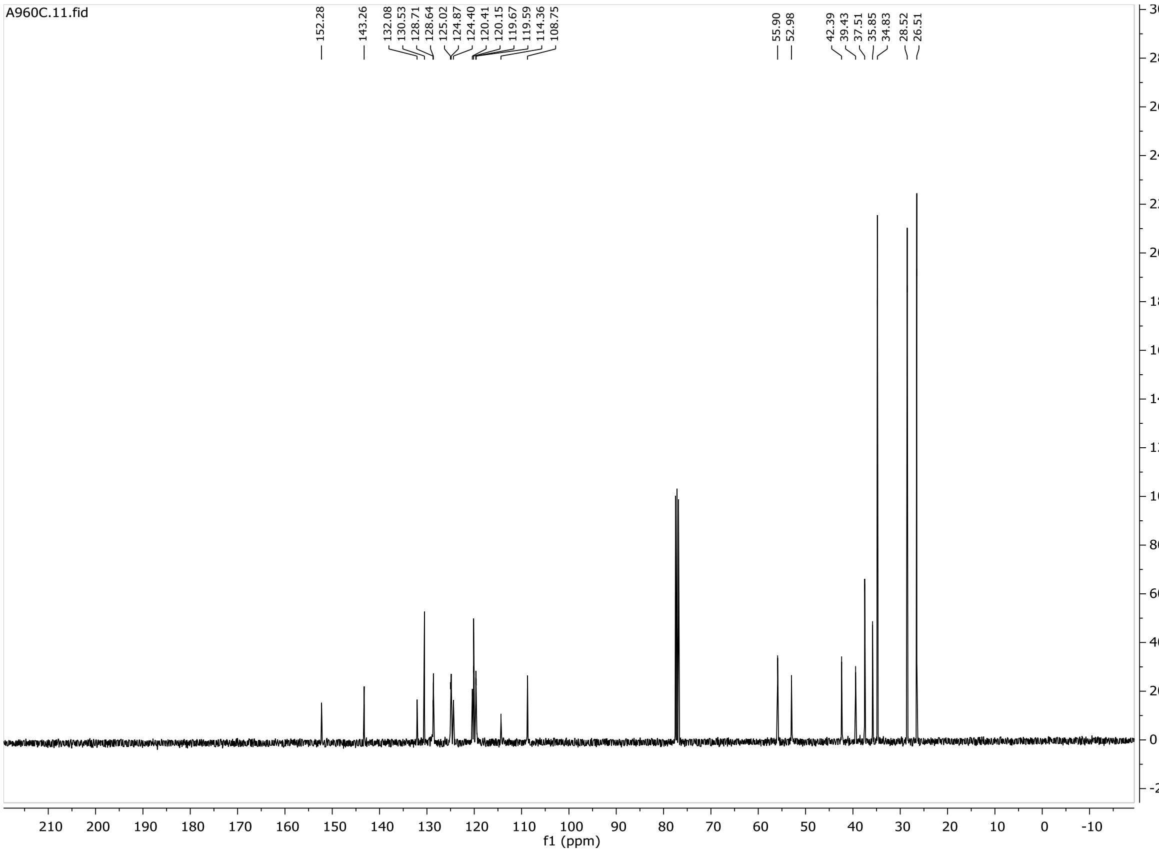
A960C.11.fid
152.28 143.26 132.08 130.53 128.71 128.64 125.02 124.87 124.40 120.41 120.15 119.67 119.59 114.36 108.75 55.90 52.98 42.39 39.43 37.51 35.85 34.83 28.52 26.51
f1 (ppm)

**17**

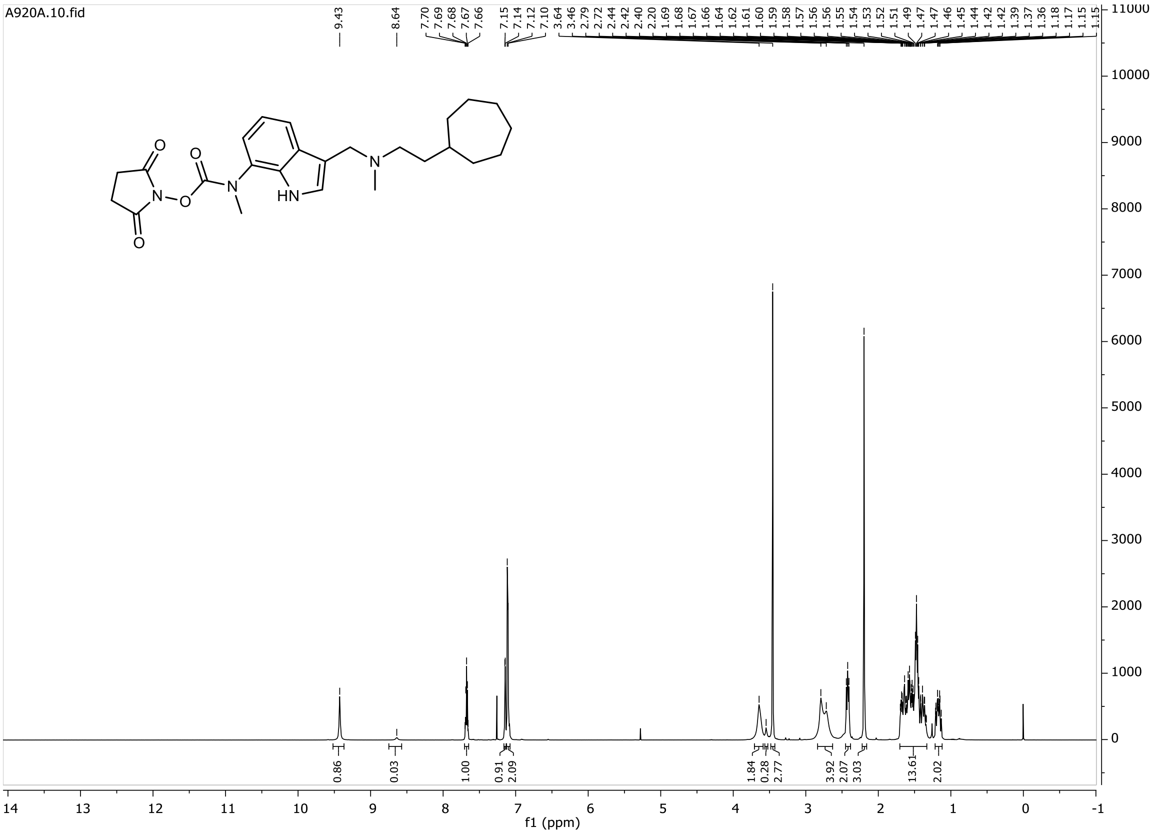
A920A.10.fid
9.43
8.64
7.70 7.69 7.68 7.67 7.66
7.15 7.14 7.12 7.10
3.64 3.46 2.79 2.72 2.44 2.42 2.40 2.20 1.69 1.68 1.67 1.66 1.64 1.62 1.61 1.60 1.59 1.58 1.57 1.56 1.56 1.55 1.54 1.53 1.52 1.51 1.49 1.47 1.47 1.46 1.45 1.44 1.42 1.39 1.37 1.36 1.18 1.17 1.15 1.15
0.86 0.03 1.00 0.91 2.09 1.84 0.28 2.77 3.92 2.07 3.03 13.61 2.02
O
N
HN
f1 (ppm)

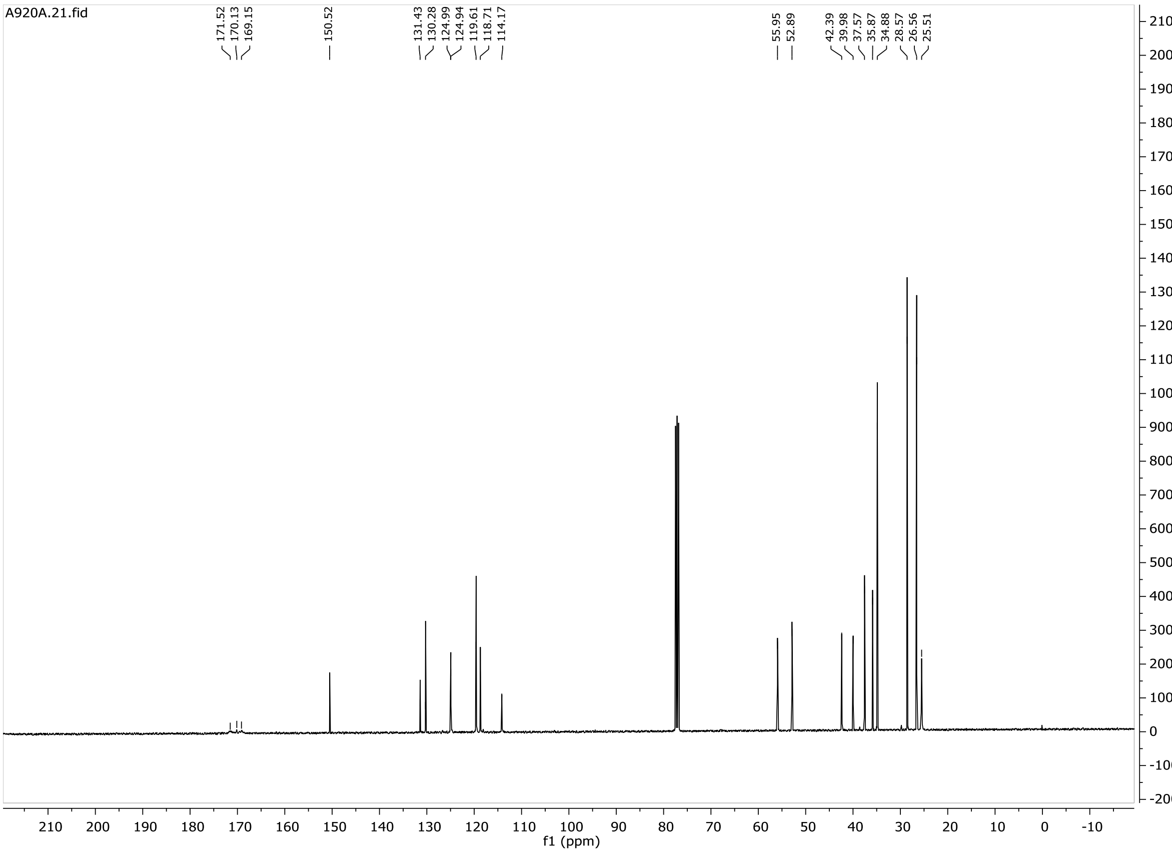
A920A.21.fid
171.52 170.13 169.15
150.52
131.43 130.28 124.99 124.94 119.61 118.71 114.17
55.95 52.89
42.39 39.98 37.57 35.87 34.88 28.57 26.56 25.51
f1 (ppm)

**17b**

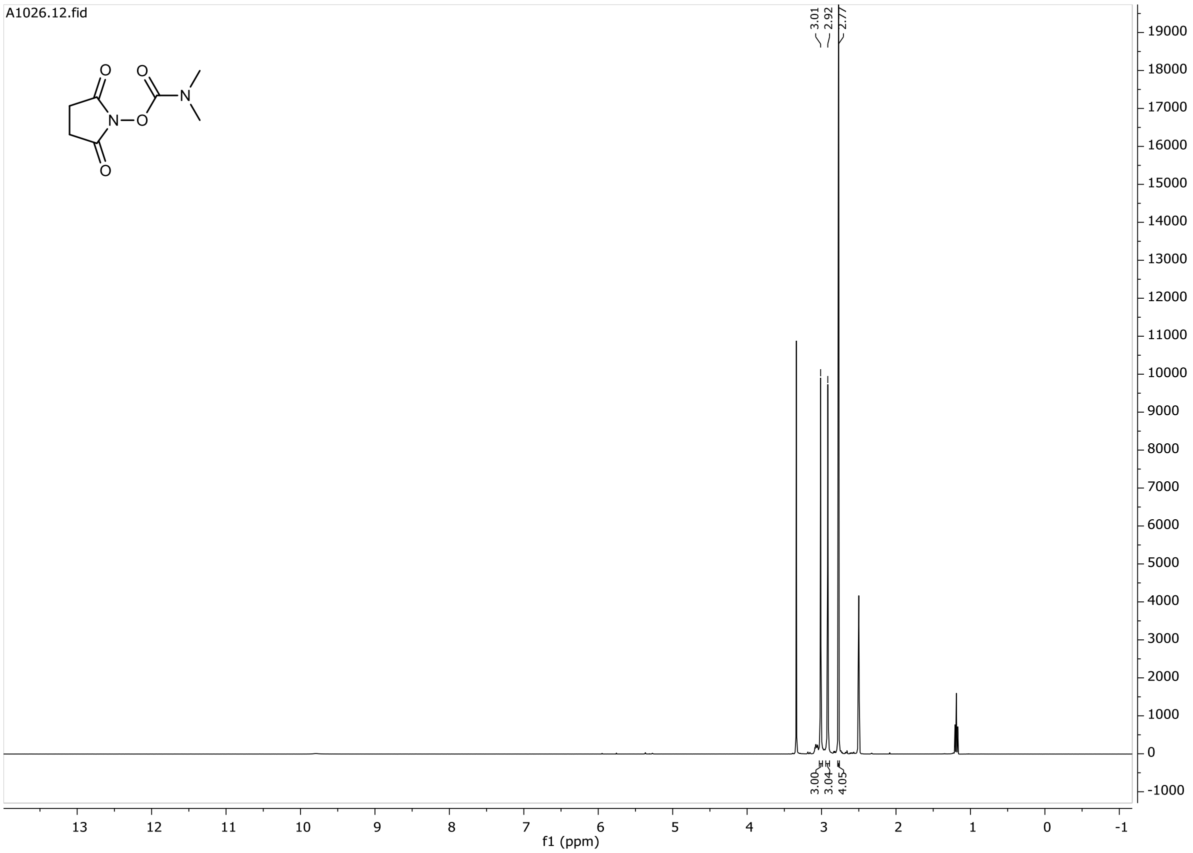
A1026.12.fid
3.01
2.92
2.77
3.00
3.04
4.05
O
N
f1 (ppm)

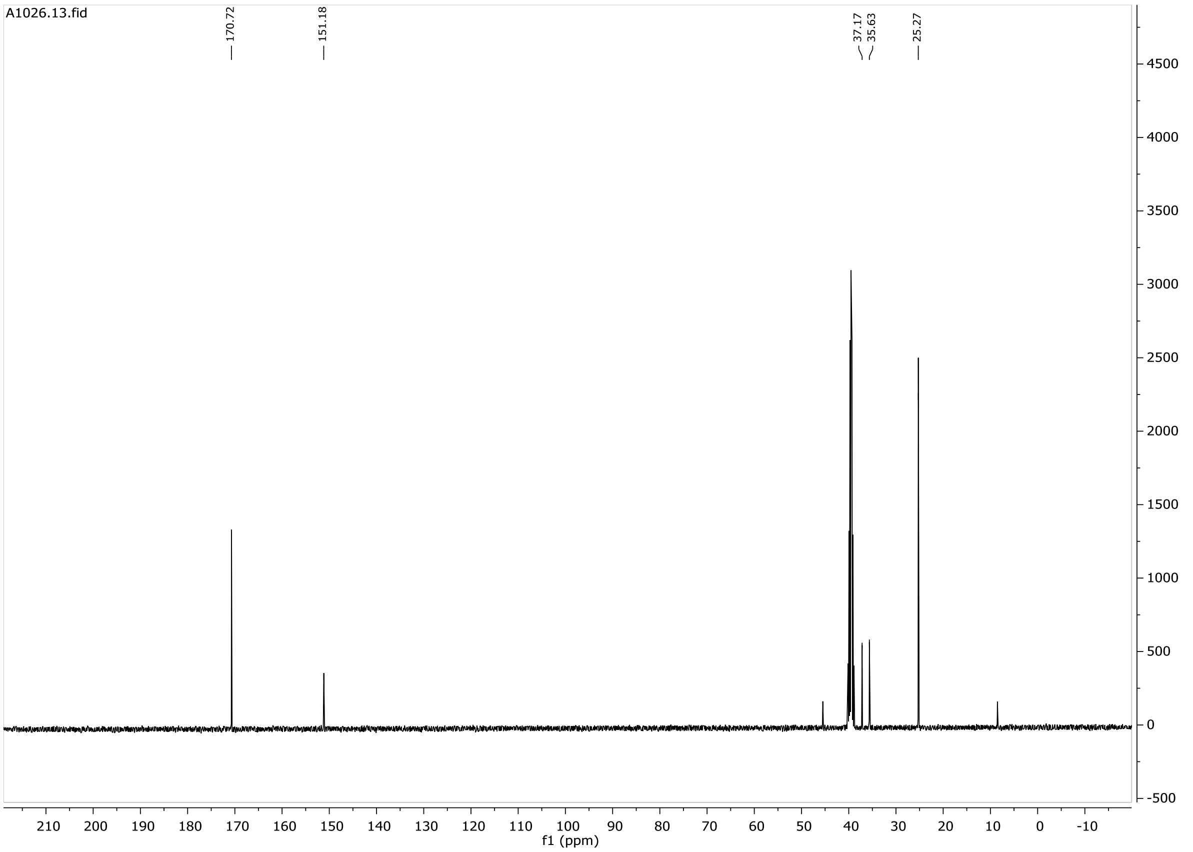
A1026.13.fid
170.72
151.18
37.17
35.63
25.27
f1 (ppm)

**18**

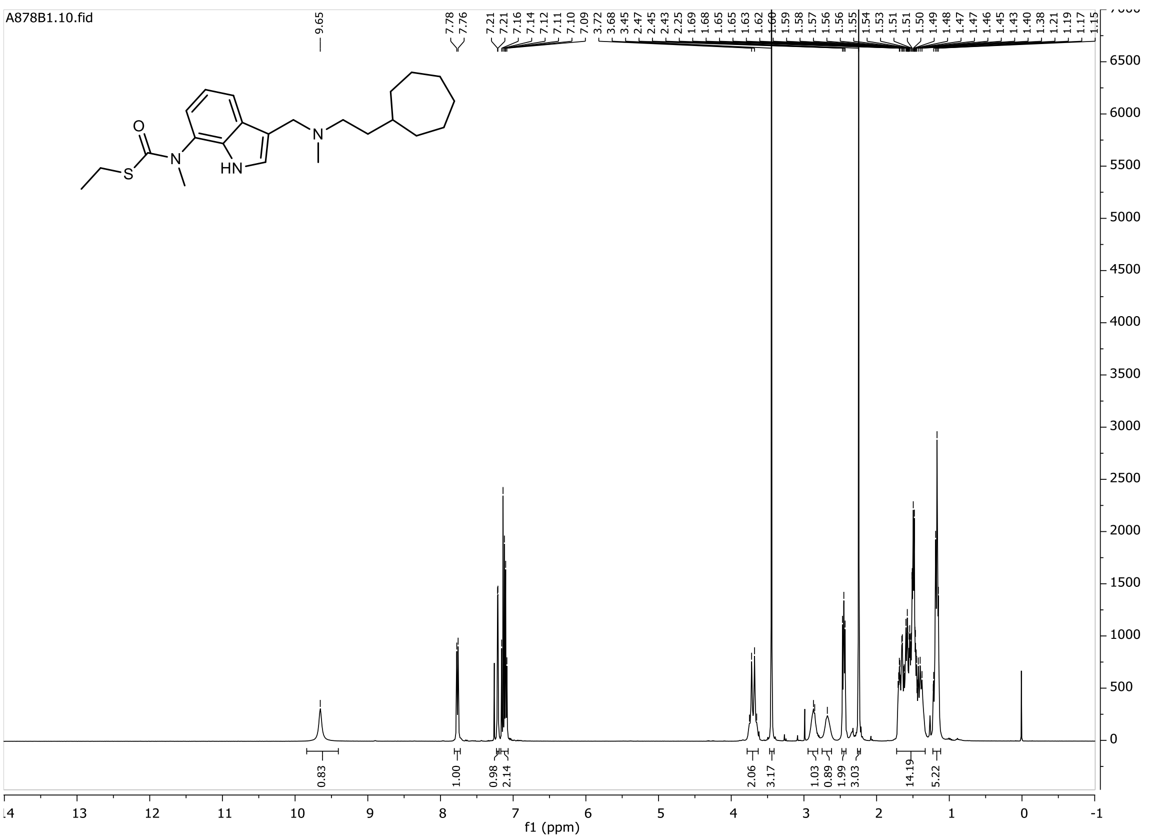
A878B1.10.fid
f1 (ppm)

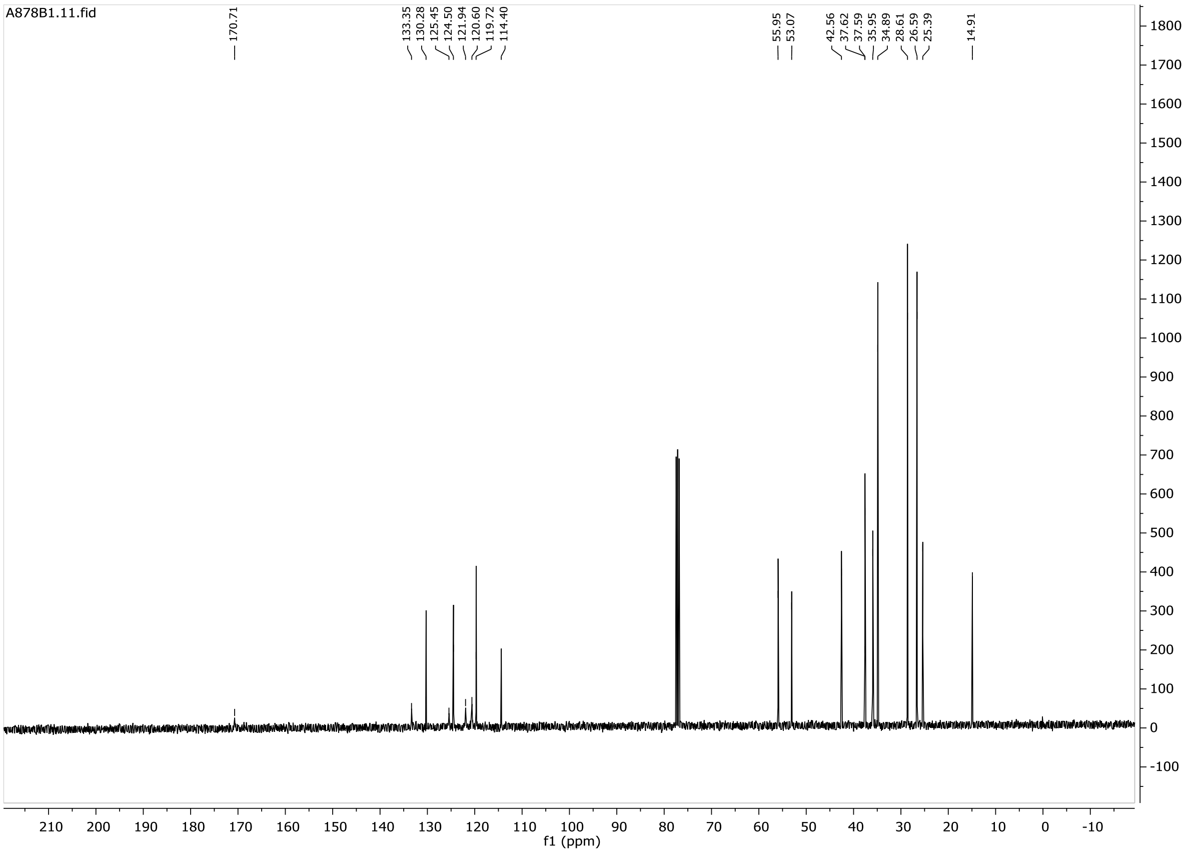
A878B1.11.fid
170.71
133.35
130.28
125.45
124.50
121.94
120.60
119.72
114.40
55.95
53.07
42.56
37.62
37.59
35.95
34.89
28.61
26.59
25.39
14.91
f1 (ppm)

**19**

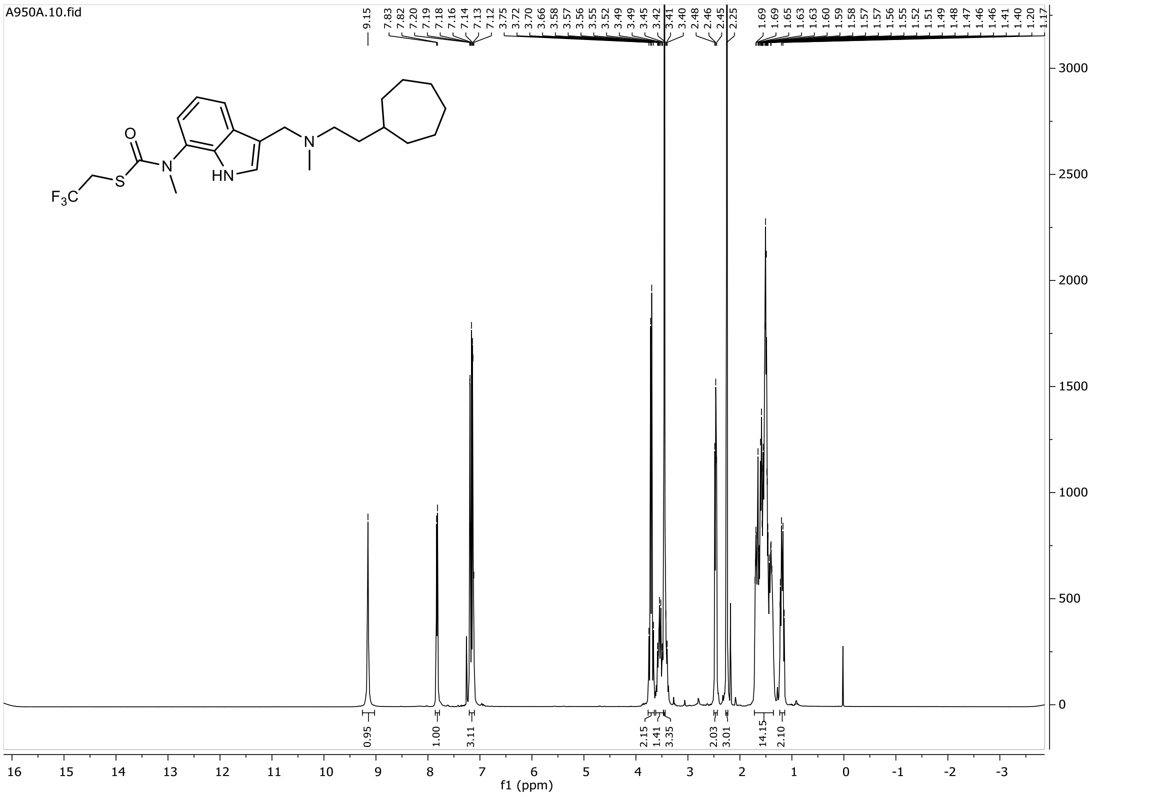
A950A.10.fid
9.15
7.83 7.82 7.20 7.19 7.18 7.16 7.14 7.13 7.12
3.75 3.72 3.70 3.66 3.58 3.57 3.56 3.55 3.52 3.49 3.49 3.45 3.42 3.41 3.40
2.48 2.46 2.45 2.25
1.69 1.69 1.65 1.63 1.63 1.60 1.59 1.58 1.57 1.57 1.56 1.55 1.52 1.51 1.49 1.48 1.47 1.46 1.46 1.41 1.40 1.20 1.17
0.95 1.00 3.11 2.15 1.41 3.35 2.03 3.01 14.15 2.10
f1 (ppm)

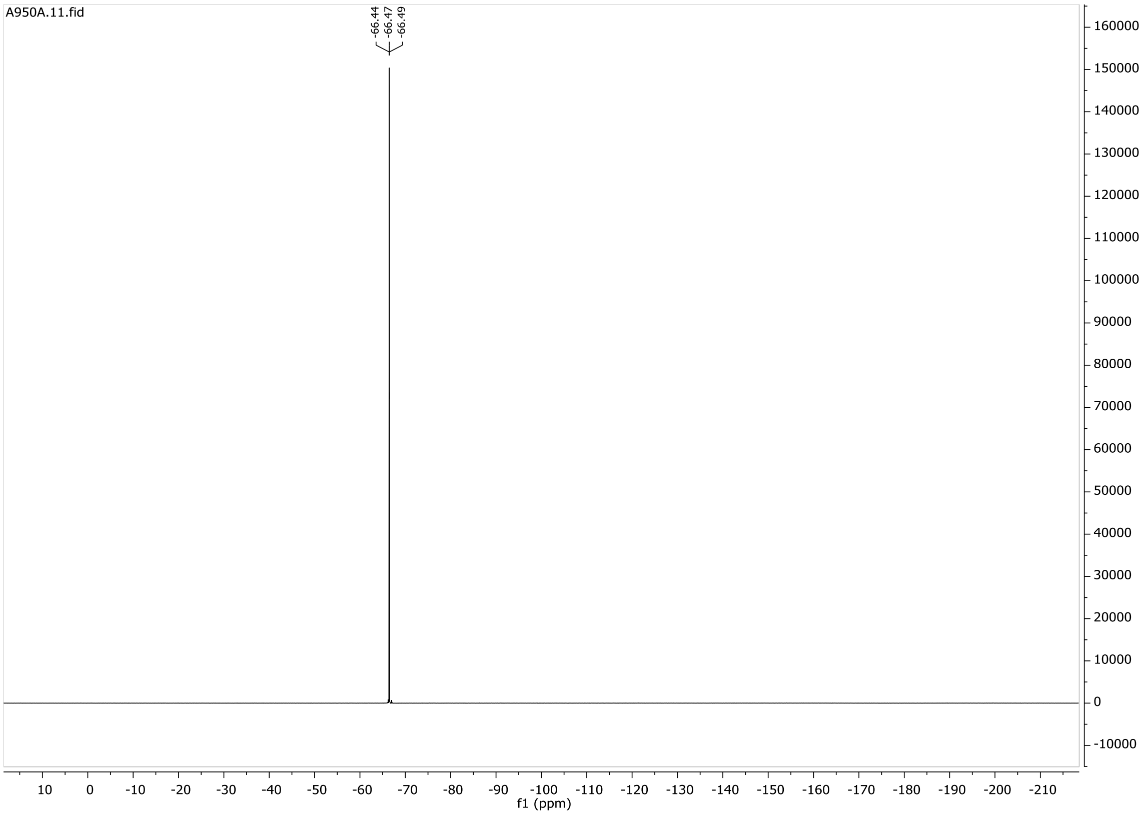
A950A.11.fid
-66.44
-66.47
-66.49
f1 (ppm)

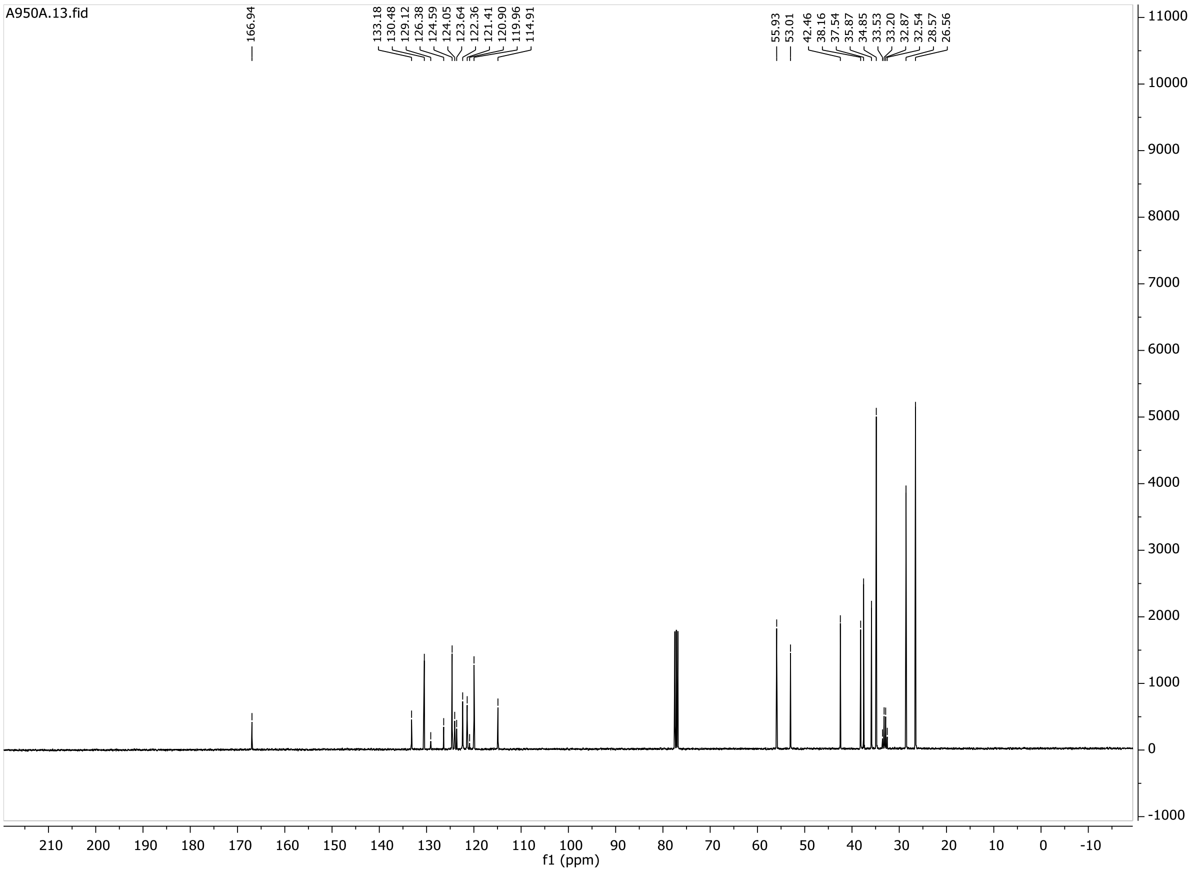
A950A.13.fid
166.94
133.18
130.48
129.12
126.38
124.59
124.05
123.64
122.36
121.41
120.90
119.96
114.91
55.93
53.01
42.46
38.16
37.54
35.87
34.85
33.53
33.20
32.87
32.54
28.57
26.56
f1 (ppm)

# 19b

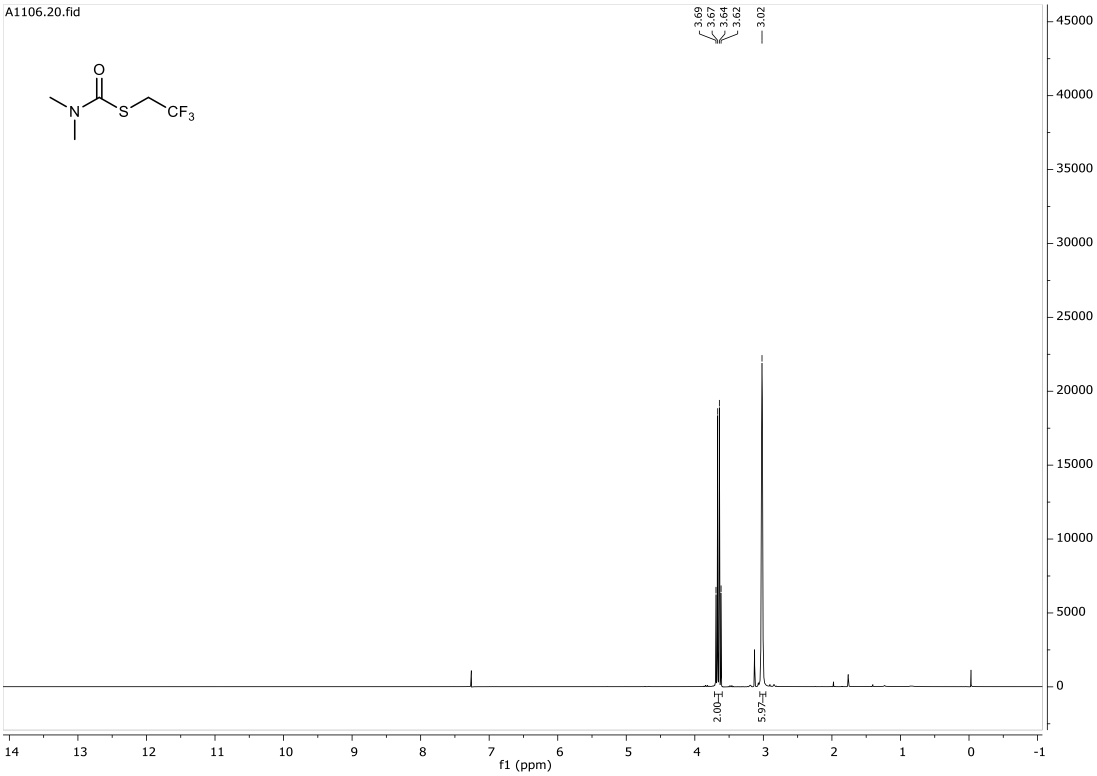
A1106.20.fid
3.69
3.67
3.64
3.62
3.02
2.00
5.97
f1 (ppm)

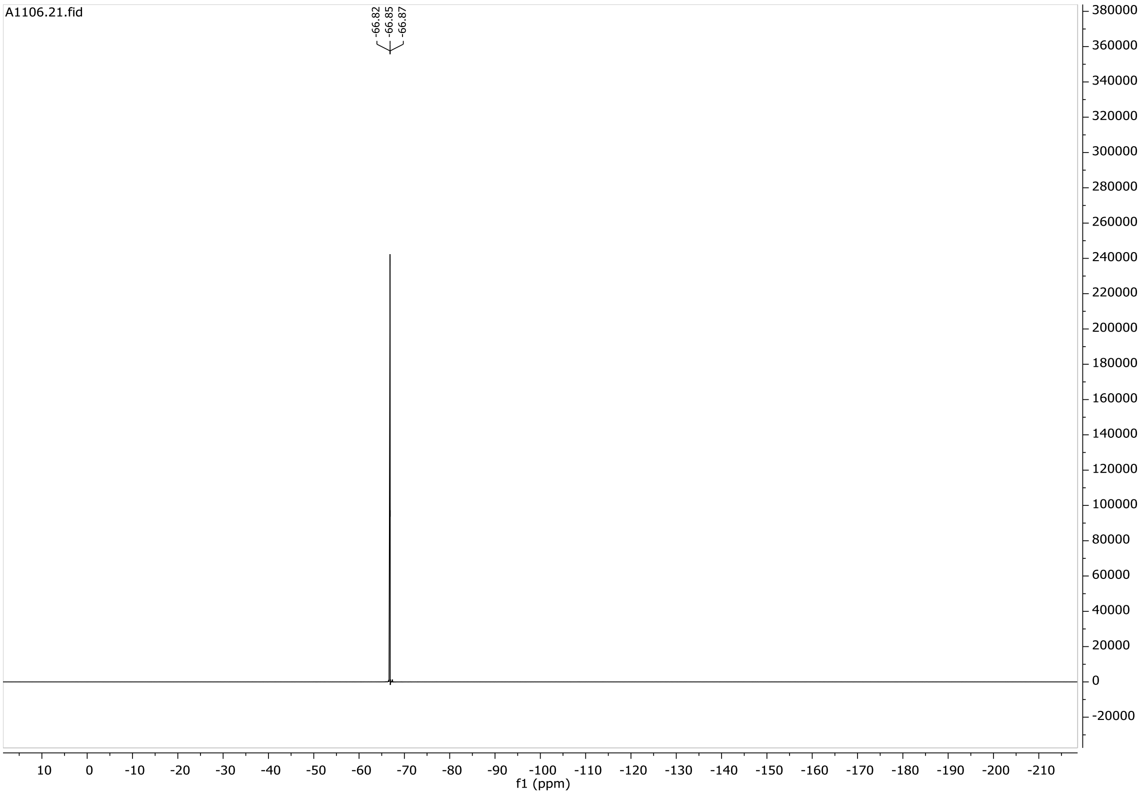
A1106.21.fid
-66.82
-66.85
-66.87
f1 (ppm)

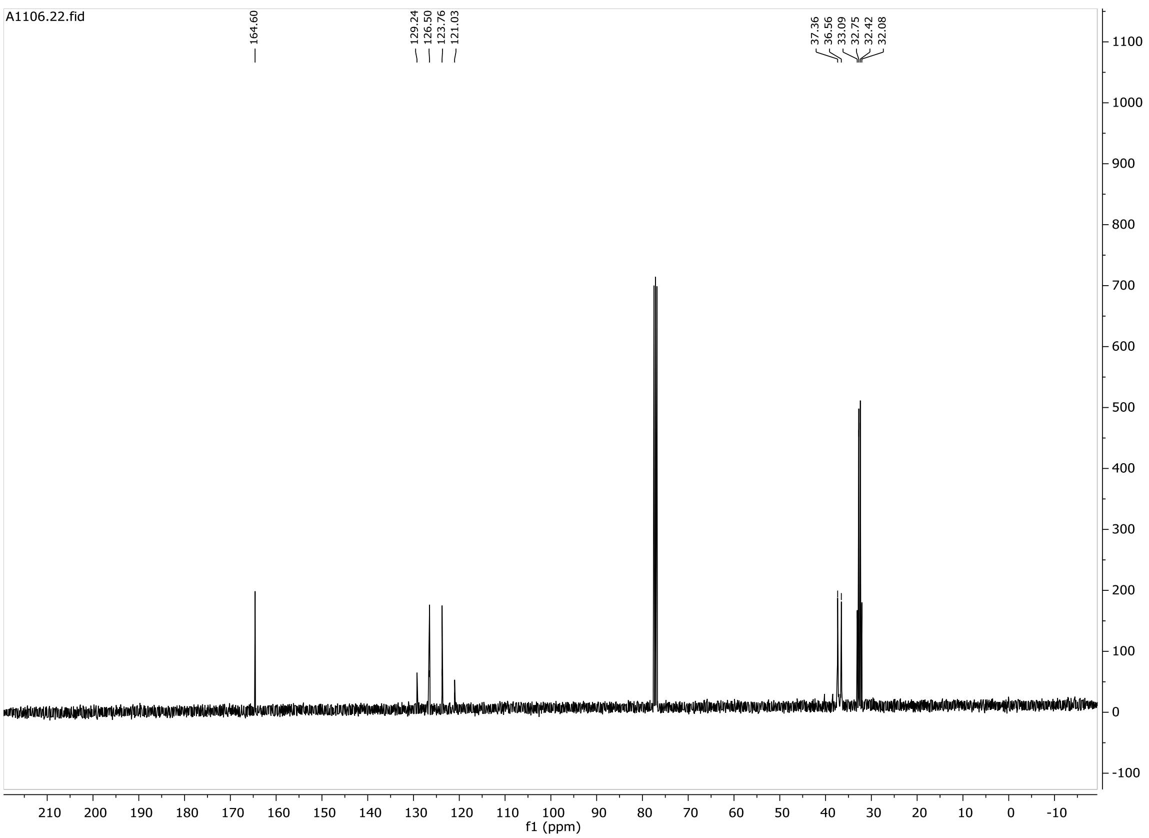

A1106.22.fid
164.60
129.24
126.50
123.76
121.03
37.36
36.56
33.09
32.75
32.42
32.08
1100
1000
900
800
700
600
500
400
300
200
100
0
-100
210 200 190 180 170 160 150 140 130 120 110 100 90 80 70 60 50 40 30 20 10 0 -10
f1 (ppm)

**20**

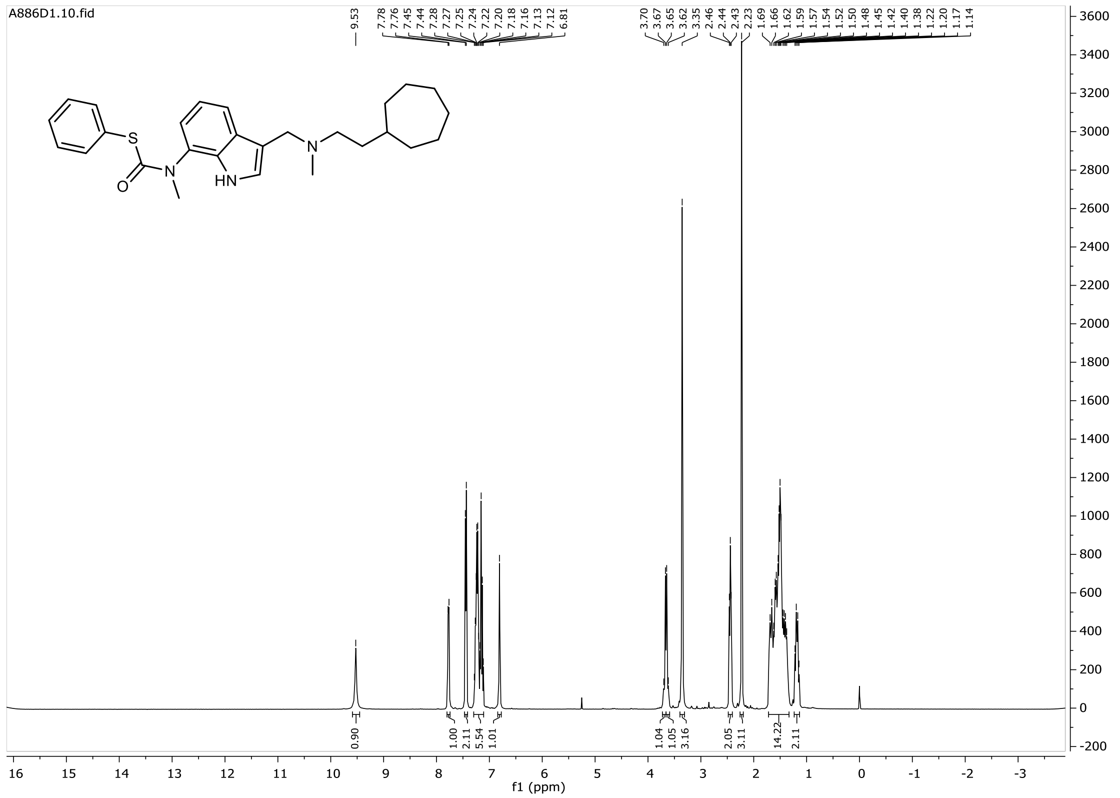
A886D1.10.fid
9.53
7.78 7.76 7.45 7.44 7.28 7.27 7.25 7.24 7.22 7.20 7.18 7.16 7.13 7.12 6.81
3.70 3.67 3.65 3.62 3.35 2.46 2.44 2.43 2.23 1.69 1.66 1.62 1.59 1.57 1.54 1.52 1.50 1.48 1.45 1.42 1.40 1.38 1.22 1.20 1.17 1.14
S
O
N
HN
N
0.90 1.00 2.11 5.54 1.01 1.04 1.05 3.16 2.05 3.11 14.22 2.11
f1 (ppm)

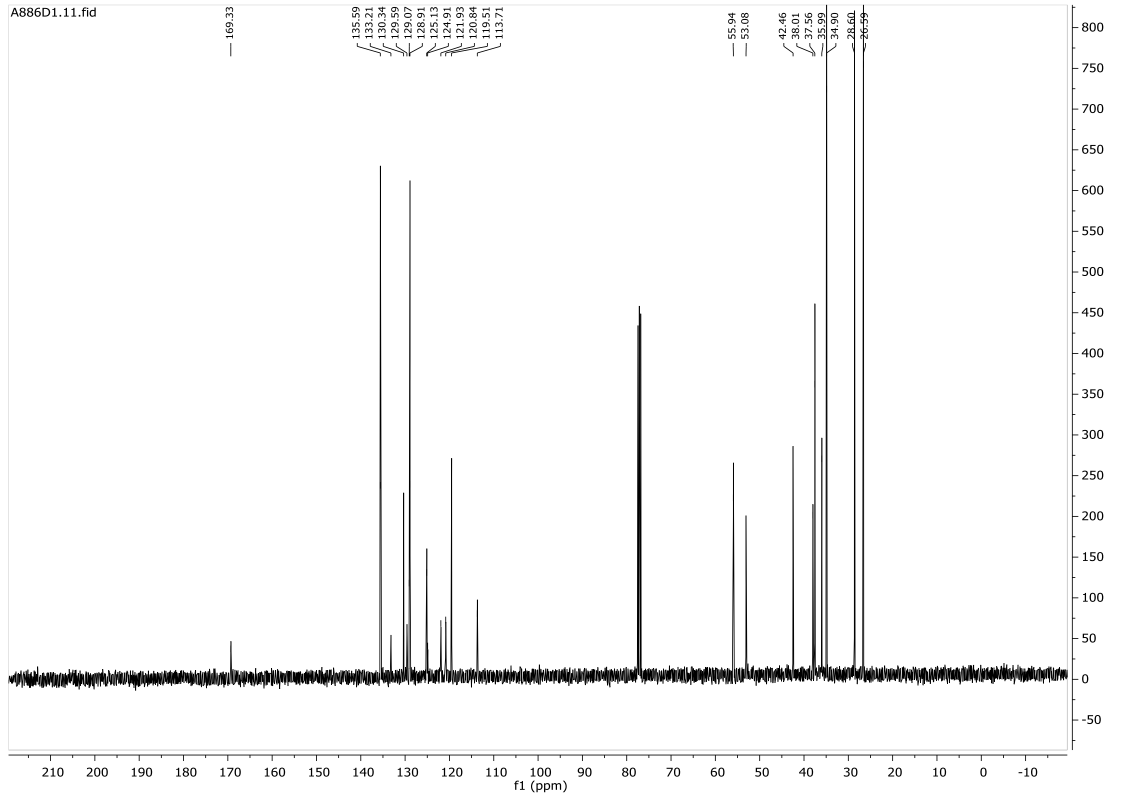
A886D1.11.fid
169.33
135.59 133.21 130.34 129.59 129.07 128.91 125.13 124.91 121.93 120.84 119.51 113.71
55.94 53.08
42.46 38.01 37.56 35.99 34.90 28.60 26.59
f1 (ppm)

**21**

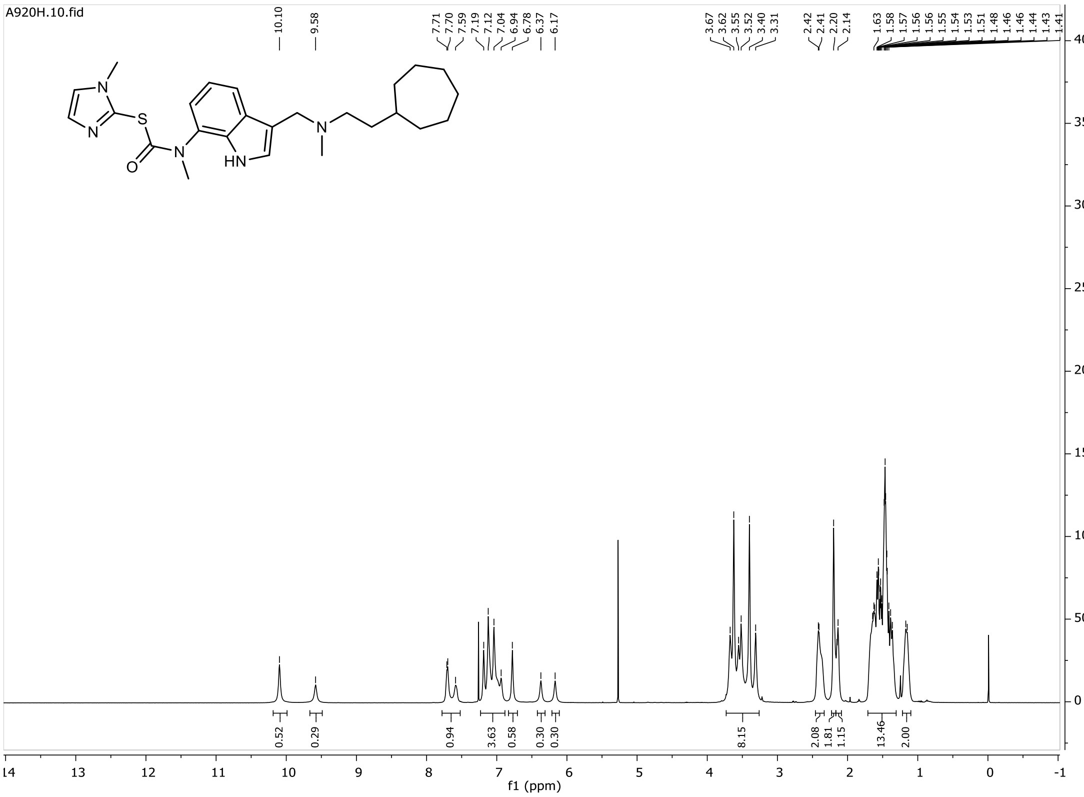

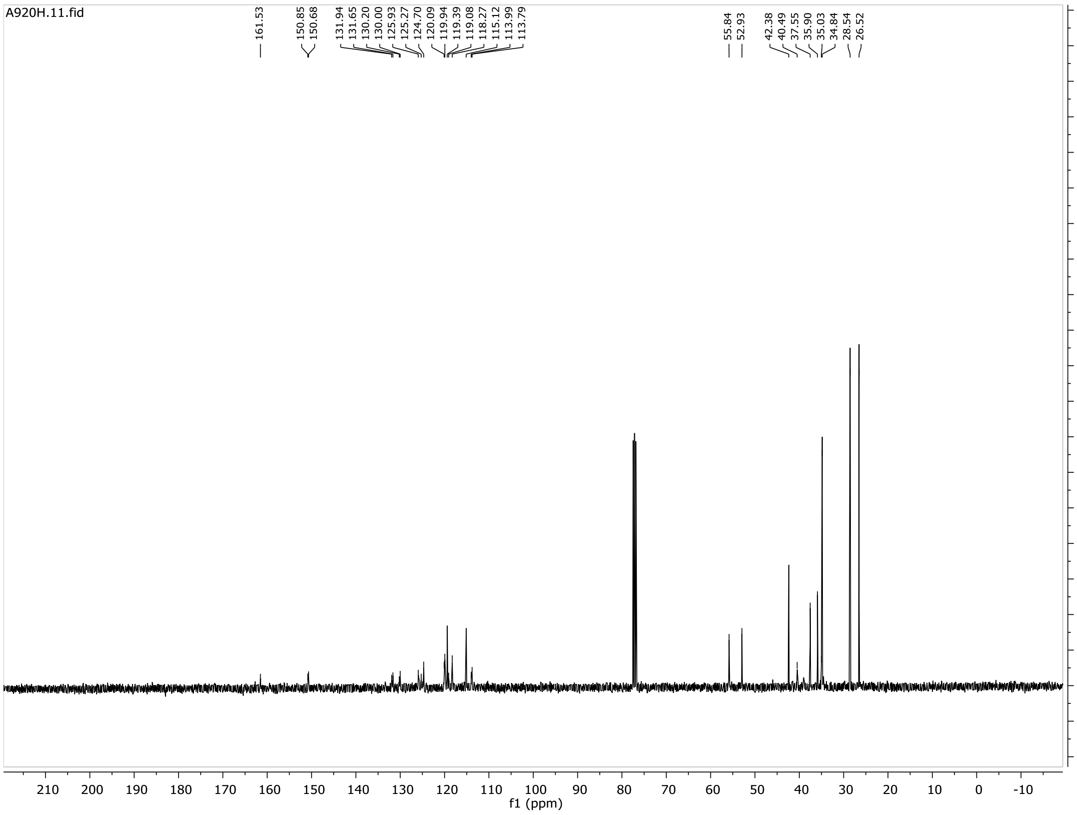

# 21b

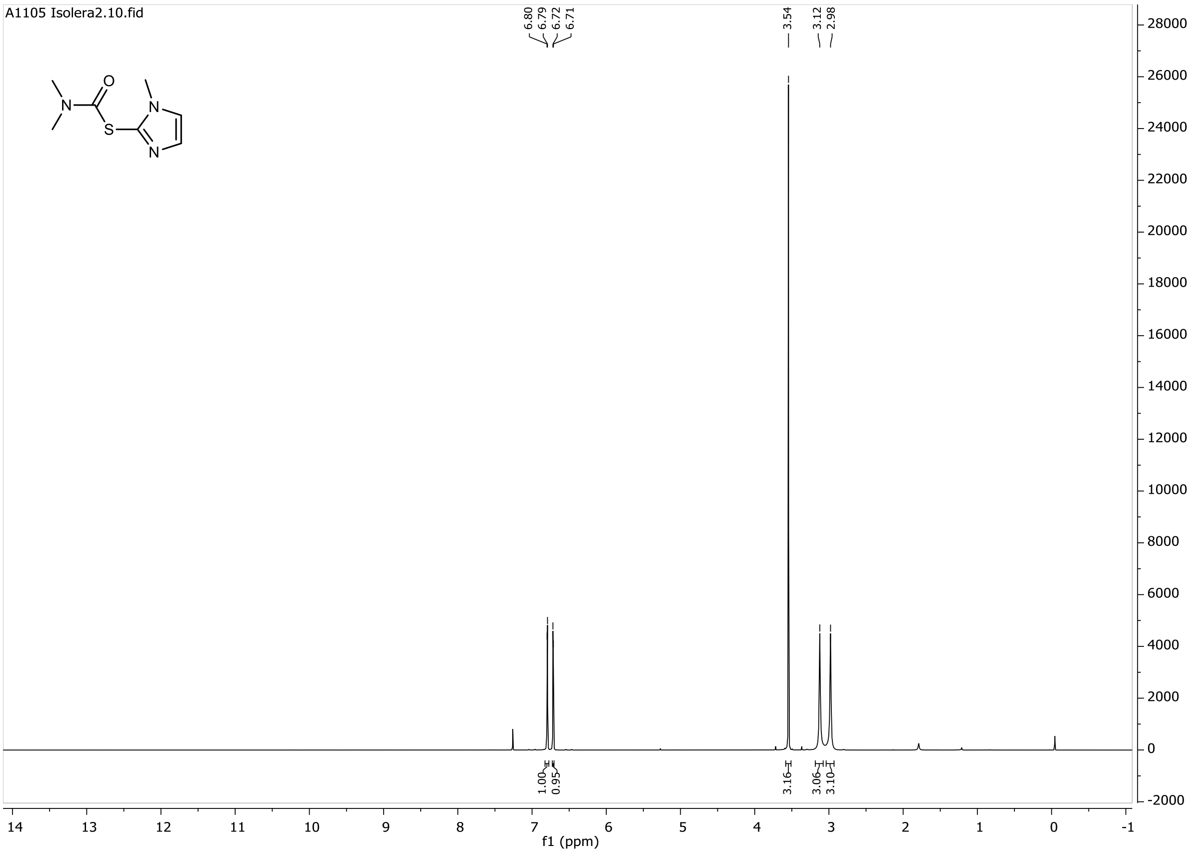
A1105 Isolera2.10.fid
6.80
6.79
6.72
6.71
3.54
3.12
2.98
O
N
S
N
N
1.00
0.95
3.16
3.06
3.10
28000
26000
24000
22000
20000
18000
16000
14000
12000
10000
8000
6000
4000
2000
0
-2000
14 13 12 11 10 9 8 7 6 5 4 3 2 1 0 -1
f1 (ppm)

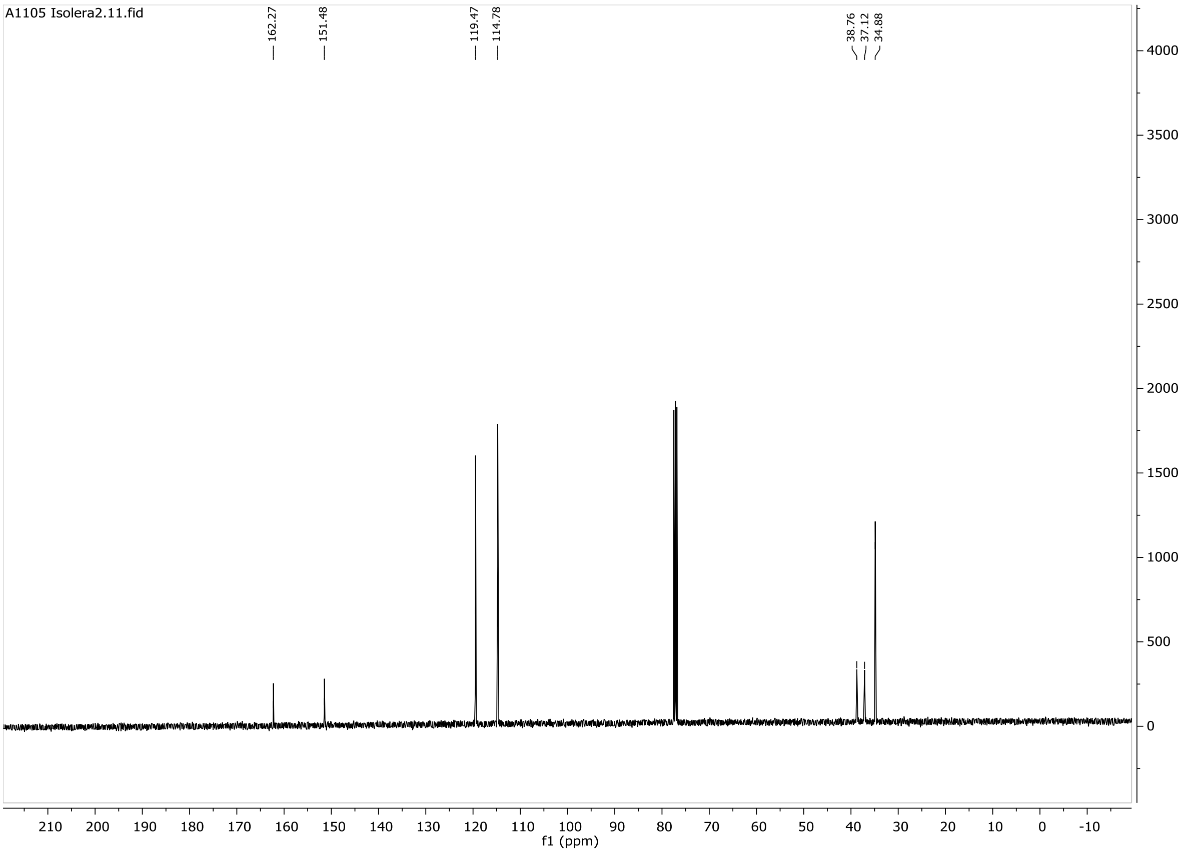
A1105 Isolera2.11.fid
162.27
151.48
119.47
114.78
38.76
37.12
34.88
4000
3500
3000
2500
2000
1500
1000
500
0
210 200 190 180 170 160 150 140 130 120 110 100 90 80 70 60 50 40 30 20 10 0 -10
f1 (ppm)

**22**

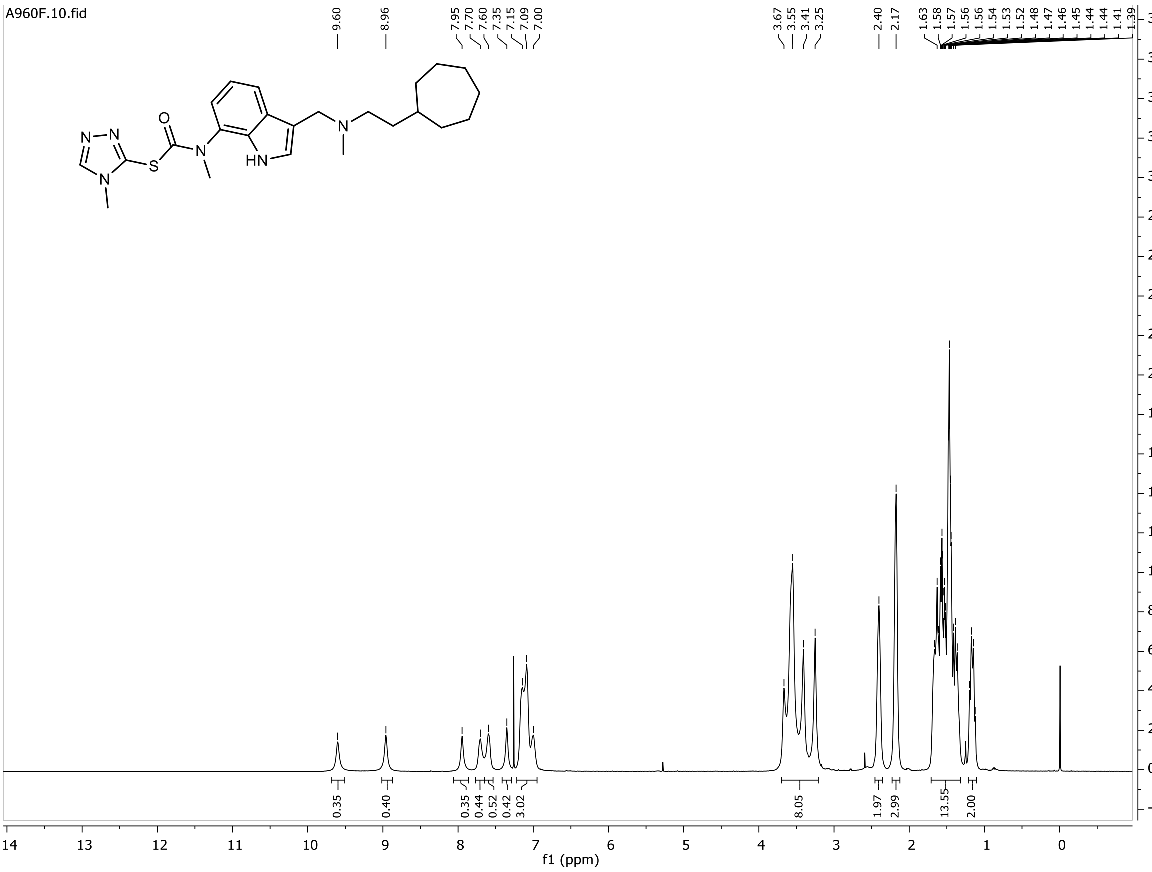
A960F.10.fid
9.60
8.96
7.95 7.70 7.60 7.35 7.15 7.09 7.00
3.67 3.55 3.41 3.25
2.40 2.17
1.63 1.58 1.57 1.56 1.56 1.54 1.53 1.52 1.48 1.47 1.46 1.45 1.44 1.44 1.41 1.39
0.35 0.40 0.35 0.44 0.52 0.42 3.02 8.05 1.97 2.99 13.55 2.00
f1 (ppm)

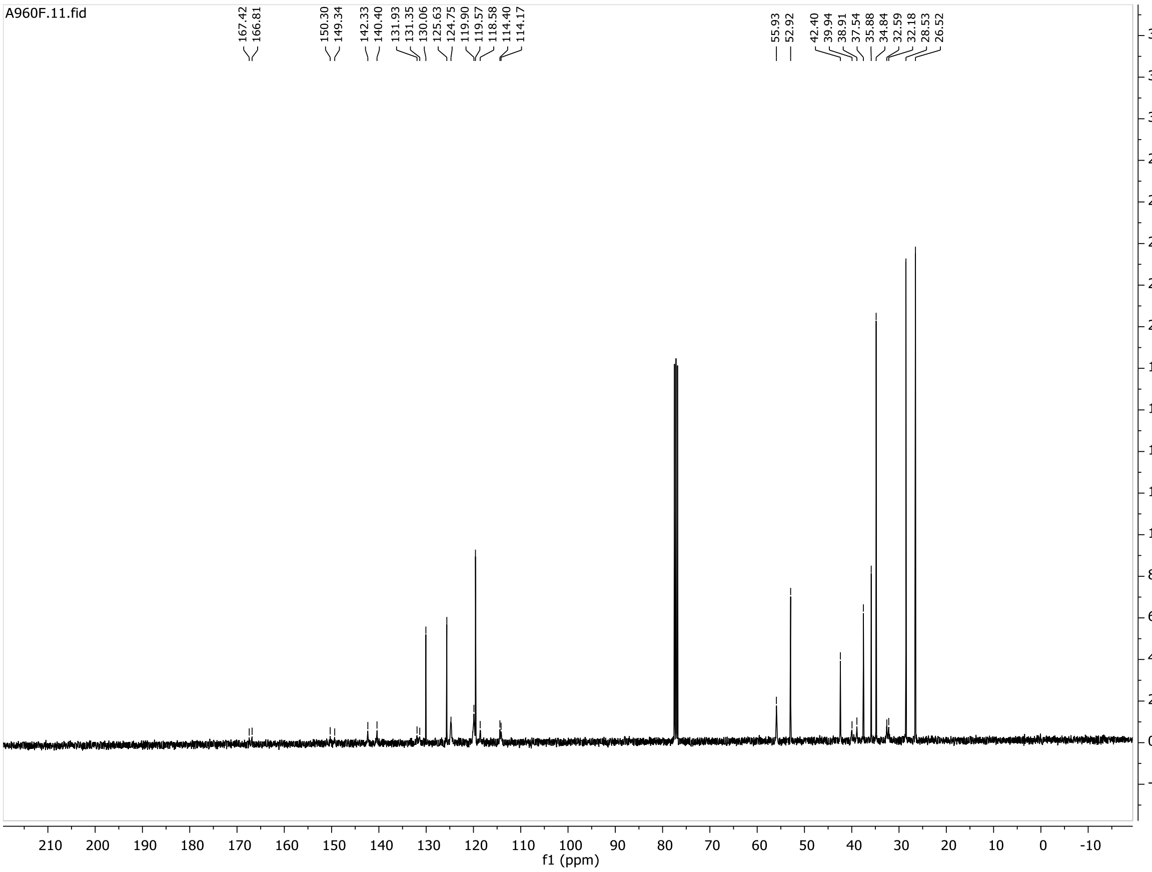
A960F.11.fid
167.42 166.81
150.30 149.34
142.33 140.40
131.93 131.35 130.06 125.63 124.75 119.90 119.57 118.58 114.40 114.17
55.93 52.92
42.40 39.94 38.91 37.54 35.88 34.84 32.59 32.18 28.53 26.52
f1 (ppm)

**23**

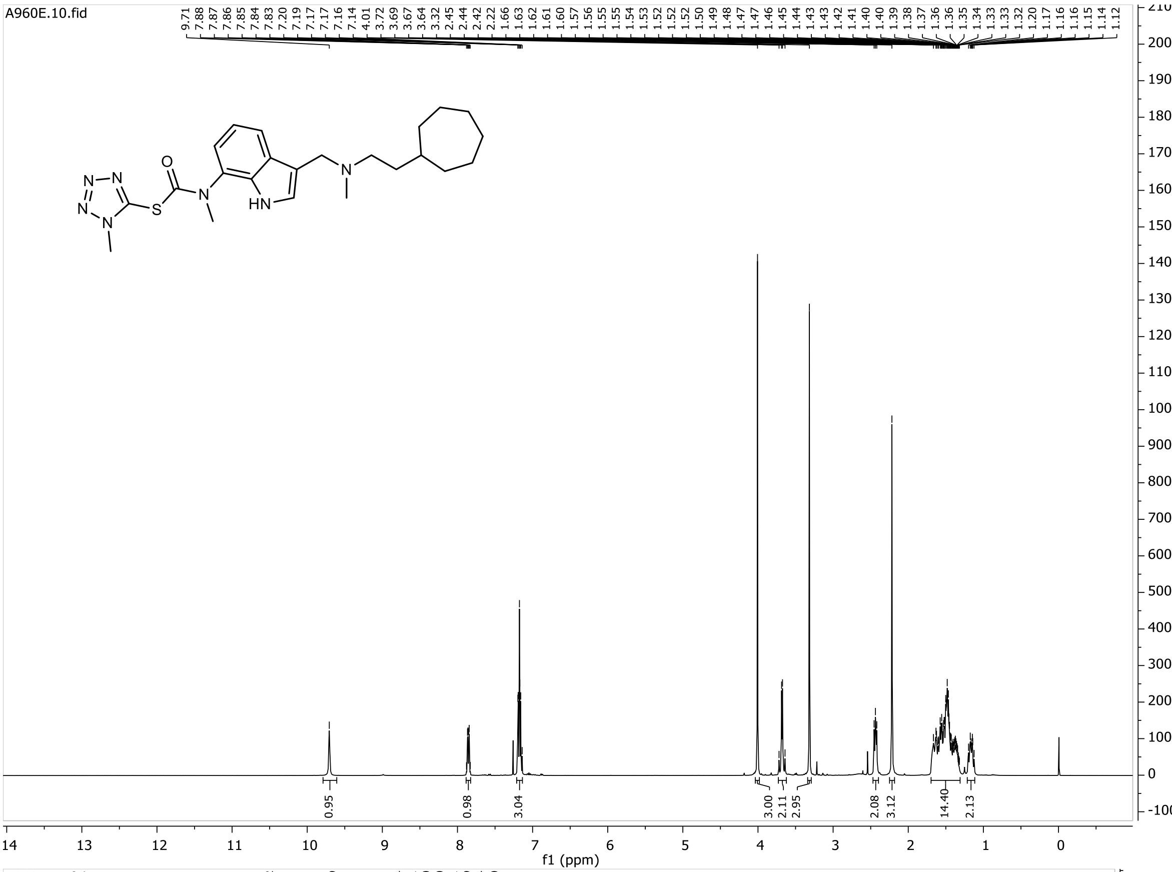

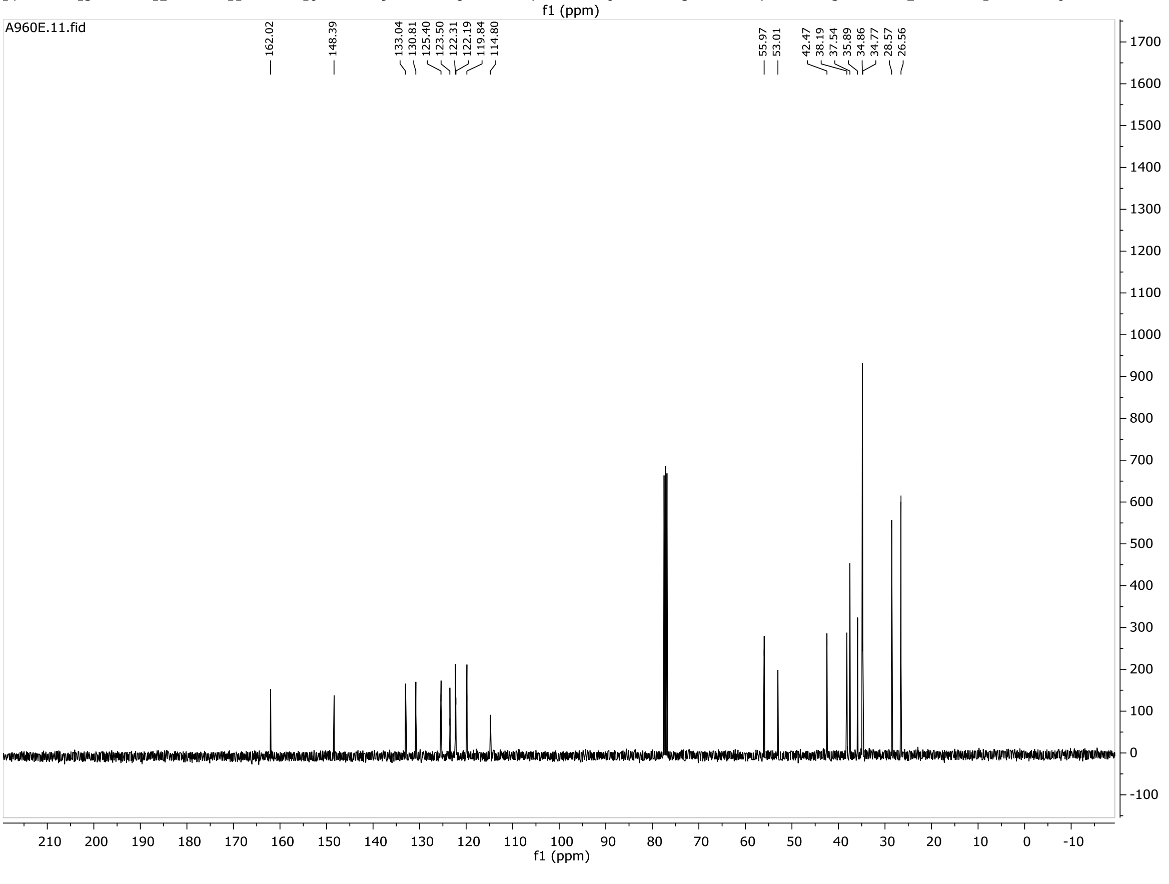

**24**

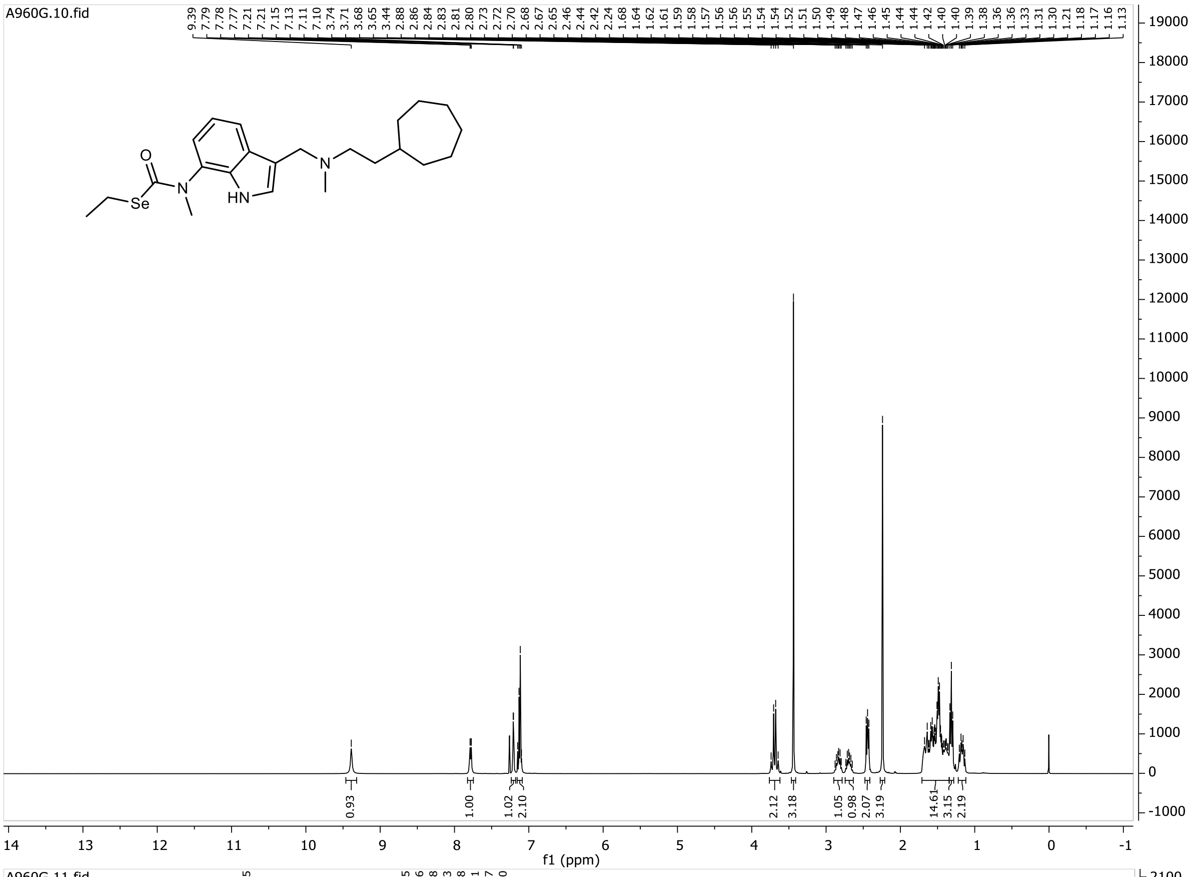
A960G.10.fid
f1 (ppm)

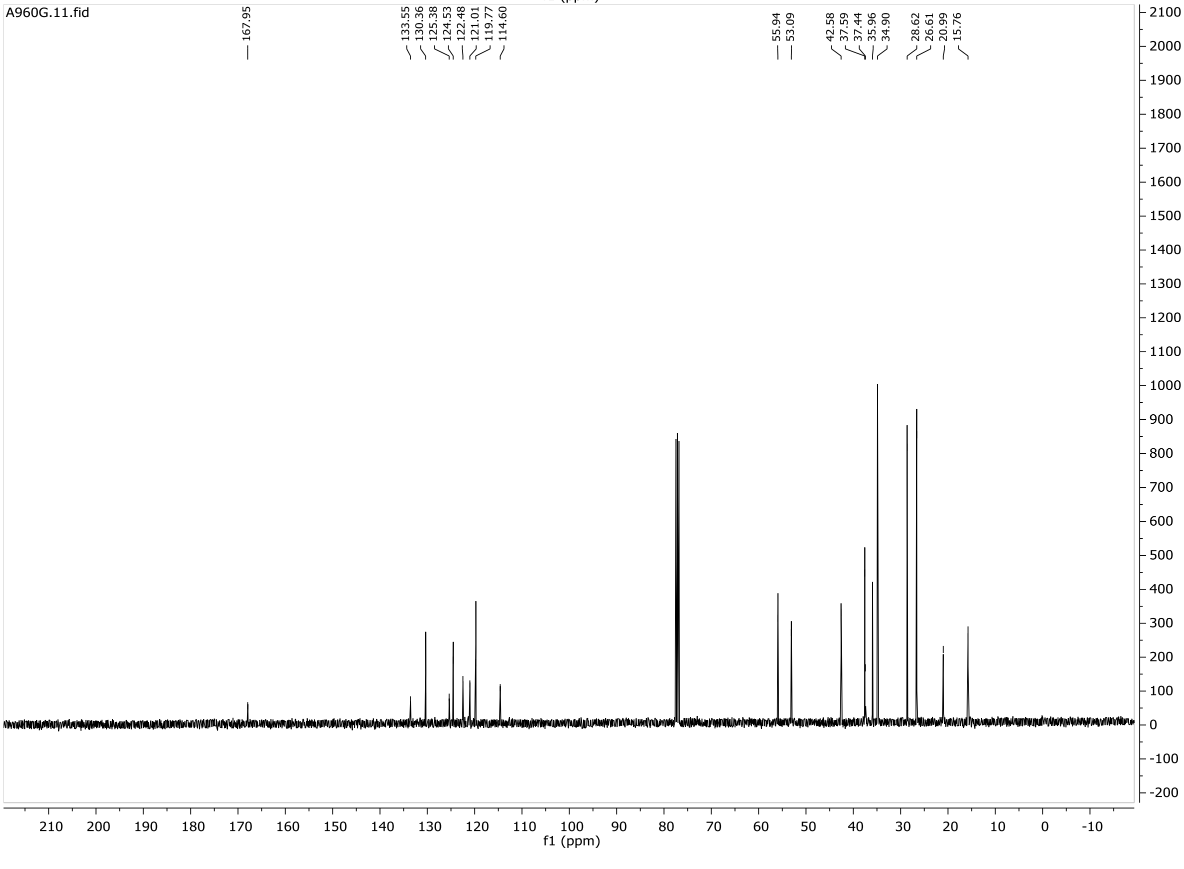
A960G.11.fid
167.95
133.55
130.36
125.38
124.53
122.48
121.01
119.77
114.60
55.94
53.09
42.58
37.59
37.44
35.96
34.90
28.62
26.61
20.99
15.76
f1 (ppm)

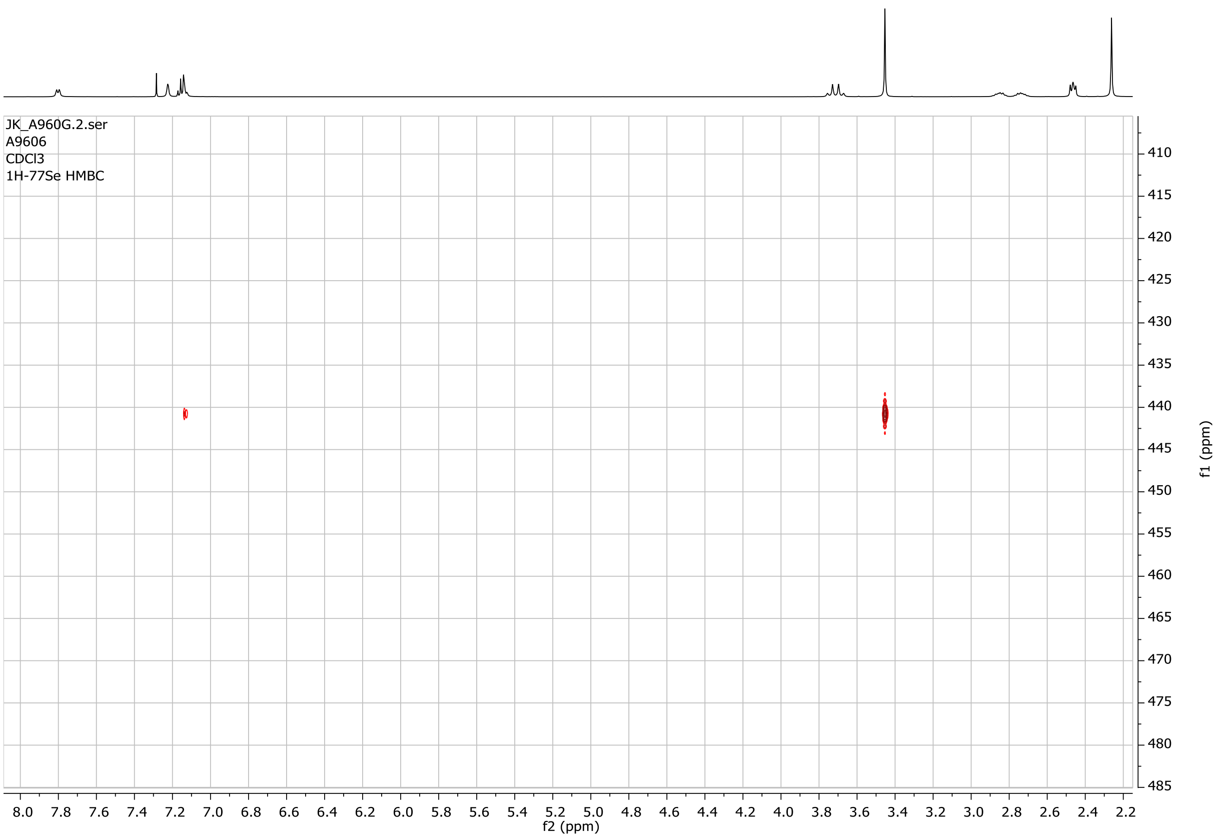
JK_A960G.2.ser
A9606
CDCl3
1H-77Se HMBC
f2 (ppm)
f1 (ppm)

**25**

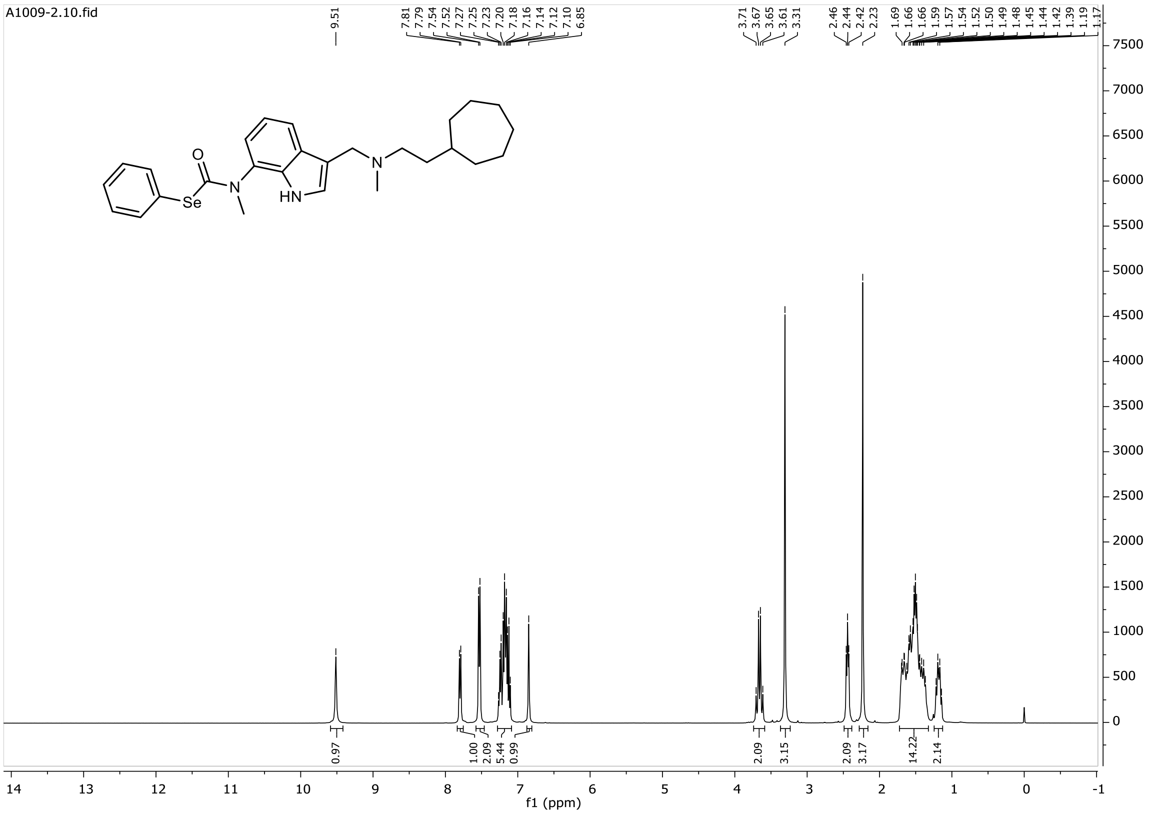
A1009-2.10.fid
9.51
7.81 7.79 7.54 7.52 7.27 7.25 7.23 7.20 7.18 7.16 7.14 7.12 7.10 6.85
3.71 3.67 3.65 3.61 3.31
2.46 2.44 2.42 2.23
1.69 1.66 1.66 1.59 1.57 1.54 1.52 1.50 1.49 1.48 1.45 1.44 1.42 1.39 1.19 1.17
0.97 1.00 2.09 5.44 0.99 2.09 3.15 2.09 3.17 14.22 2.14
Se
HN
N
O
f1 (ppm)

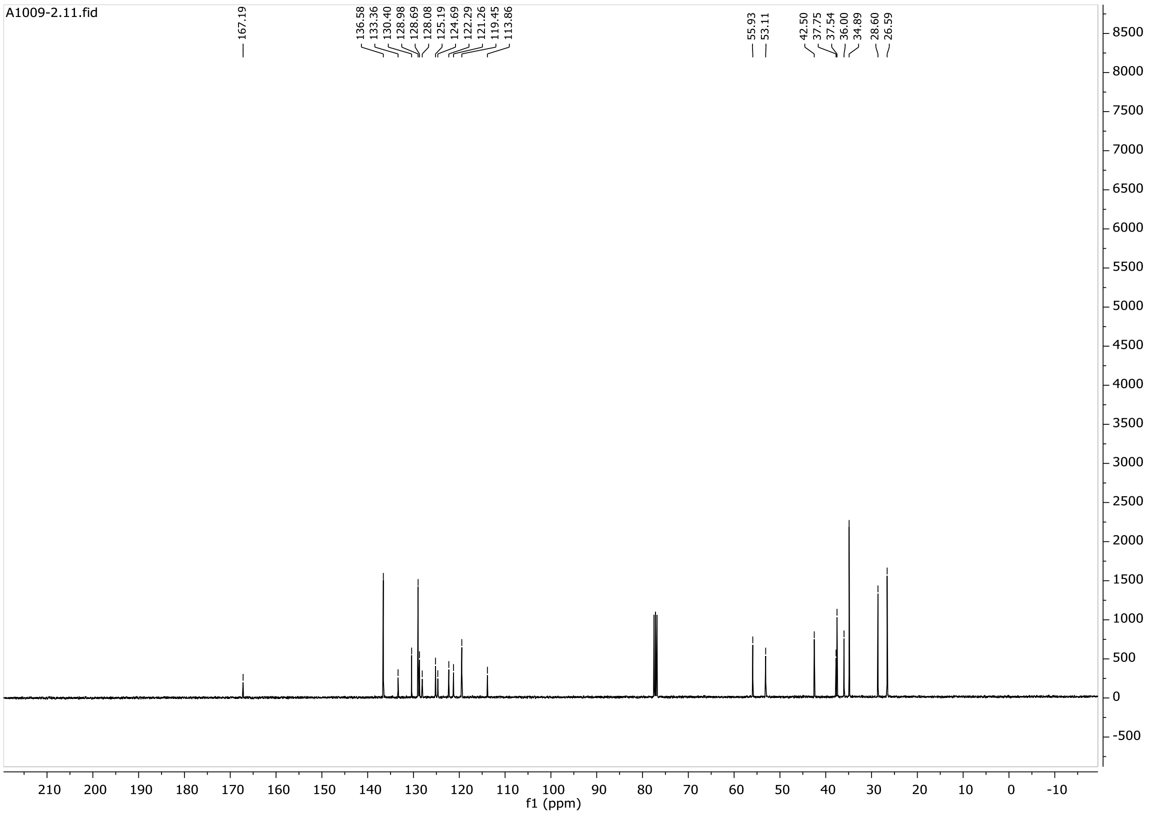
A1009-2.11.fid
167.19
136.58 133.36 130.40 128.98 128.69 128.08 125.19 124.69 122.29 121.26 119.45 113.86
55.93 53.11
42.50 37.75 37.54 36.00 34.89 28.60 26.59
f1 (ppm)

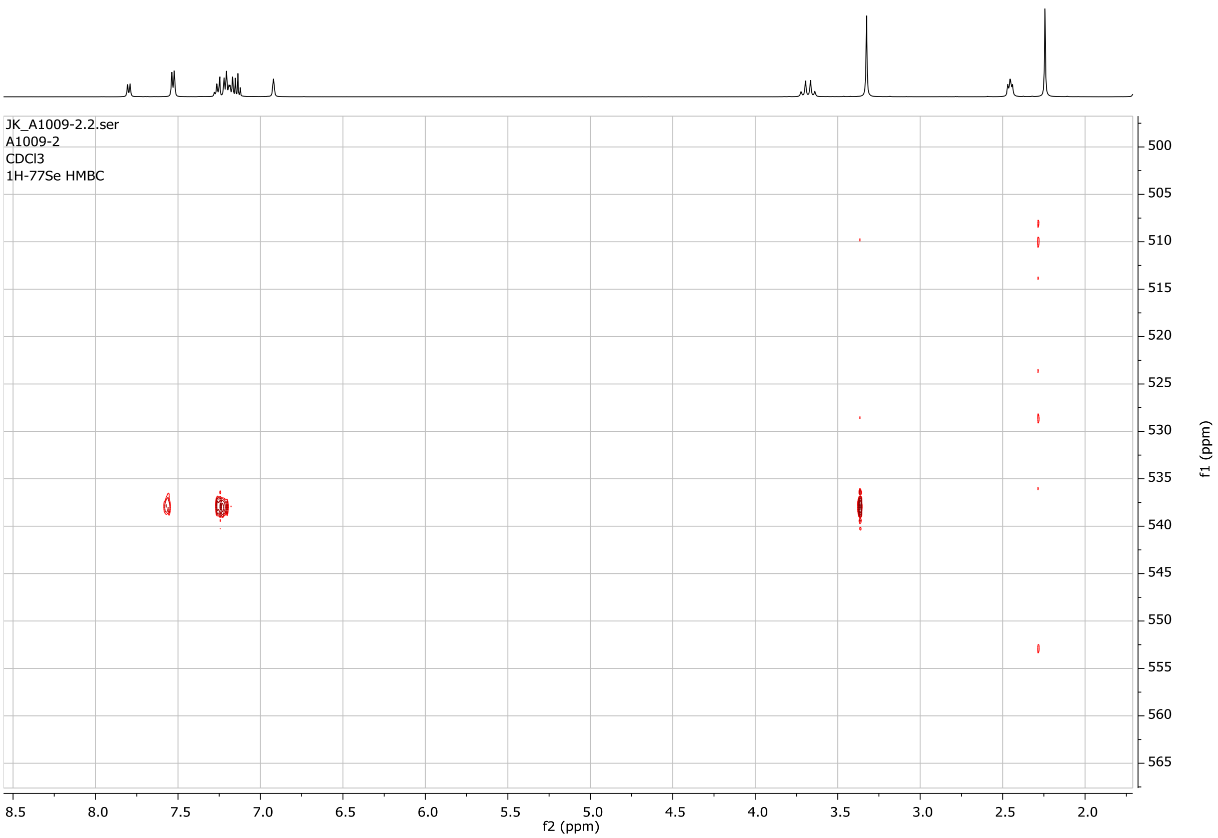
JK_A1009-2.2.ser
A1009-2
CDCl3
1H-77Se HMBC
f2 (ppm)
f1 (ppm)

**26**

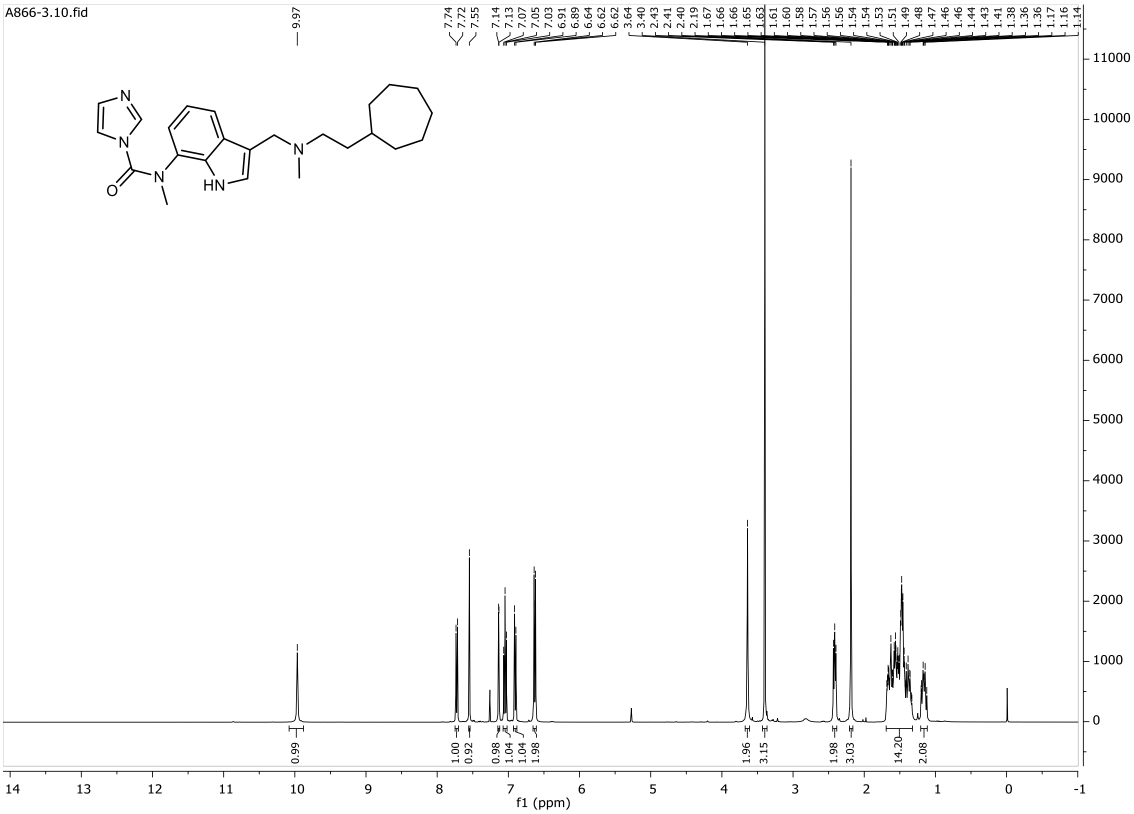
A866-3.10.fid
9.97
7.74 7.72 7.55 7.14 7.13 7.07 7.05 7.03 6.91 6.89 6.64 6.62 6.62 3.64 3.40 2.43 2.41 2.40 2.19 1.67 1.66 1.66 1.65 1.63 1.61 1.60 1.58 1.57 1.56 1.56 1.54 1.54 1.53 1.51 1.49 1.48 1.47 1.46 1.46 1.44 1.43 1.41 1.38 1.36 1.36 1.17 1.16 1.14
0.99 1.00 0.92 0.98 1.04 1.04 1.98 1.96 3.15 1.98 3.03 14.20 2.08
f1 (ppm)

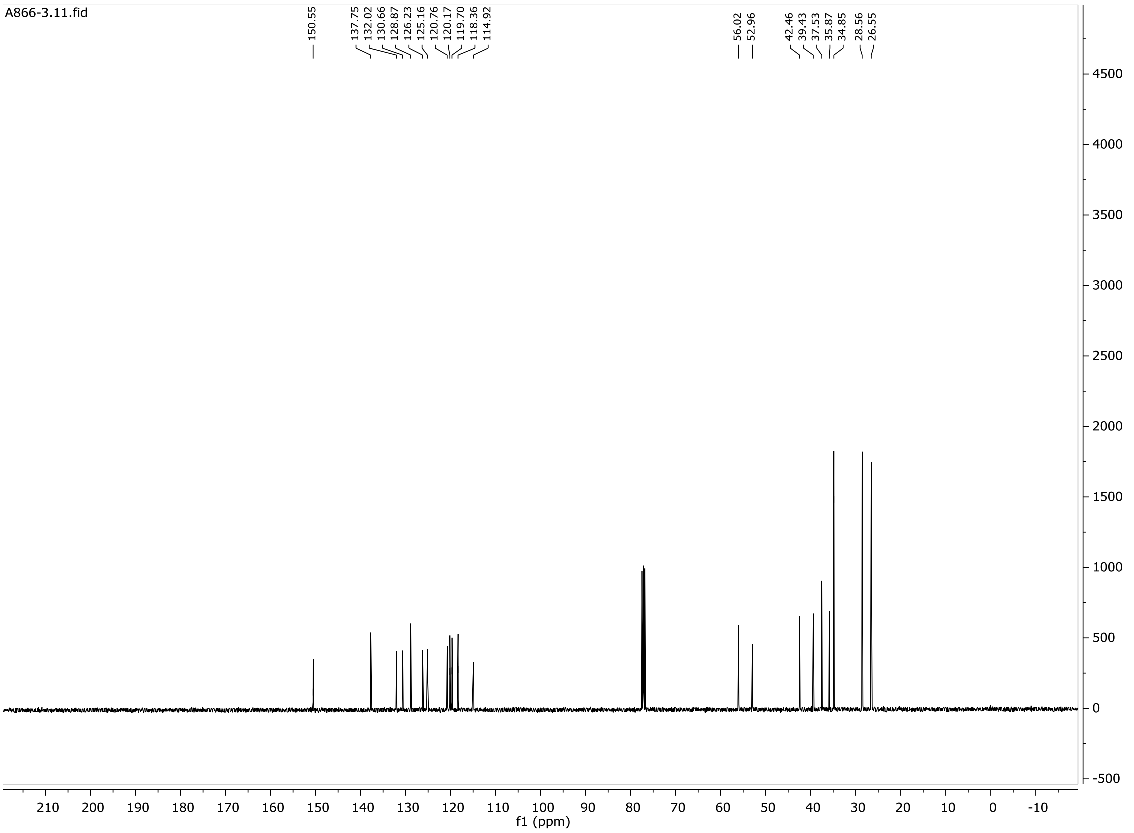
A866-3.11.fid
150.55
137.75 132.02 130.66 128.87 126.23 125.16 120.76 120.17 119.70 118.36 114.92
56.02 52.96
42.46 39.43 37.53 35.87 34.85 28.56 26.55
f1 (ppm)

**27**

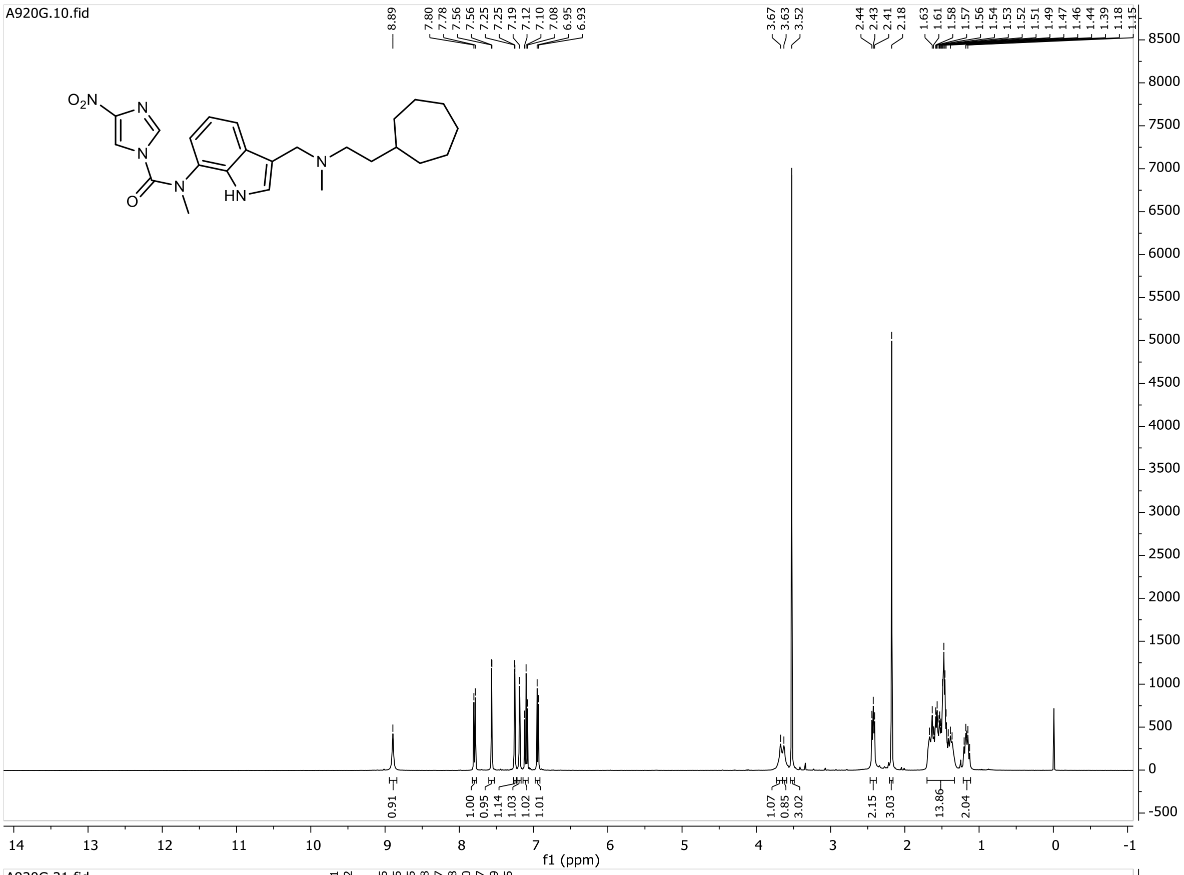

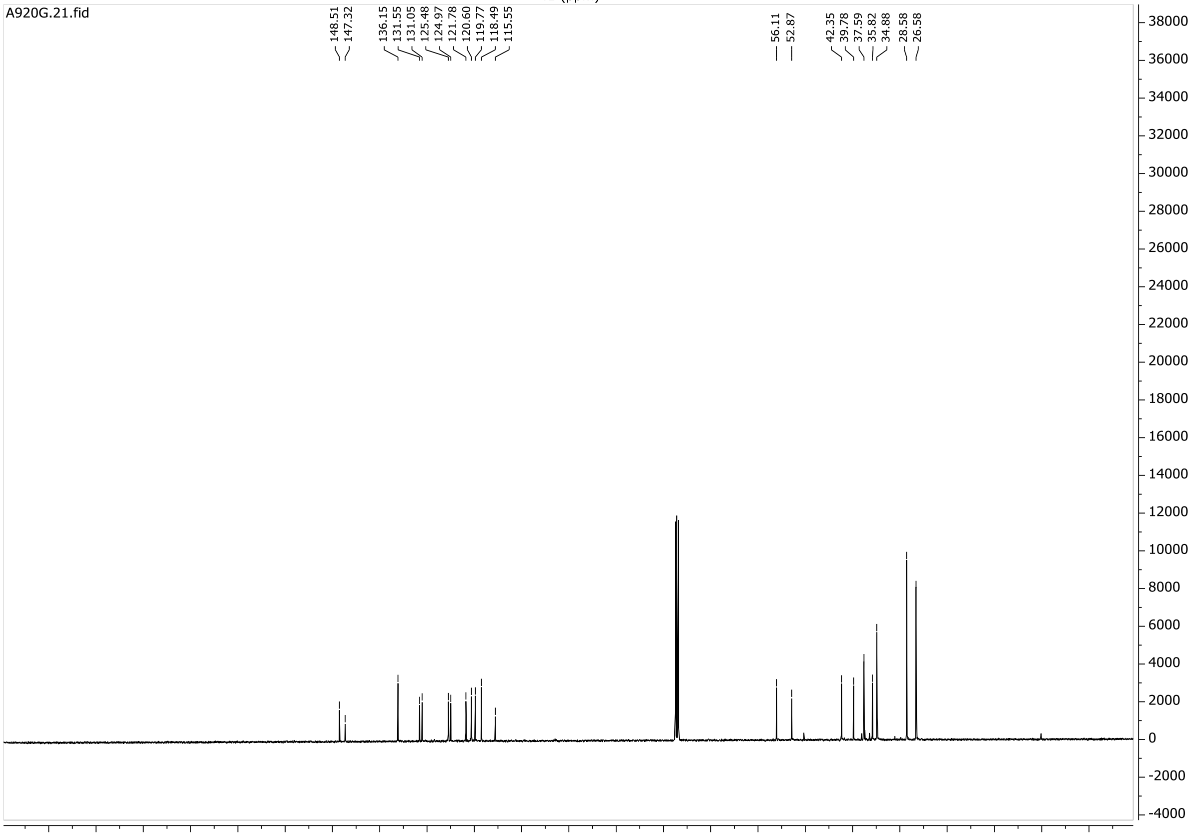

**28**

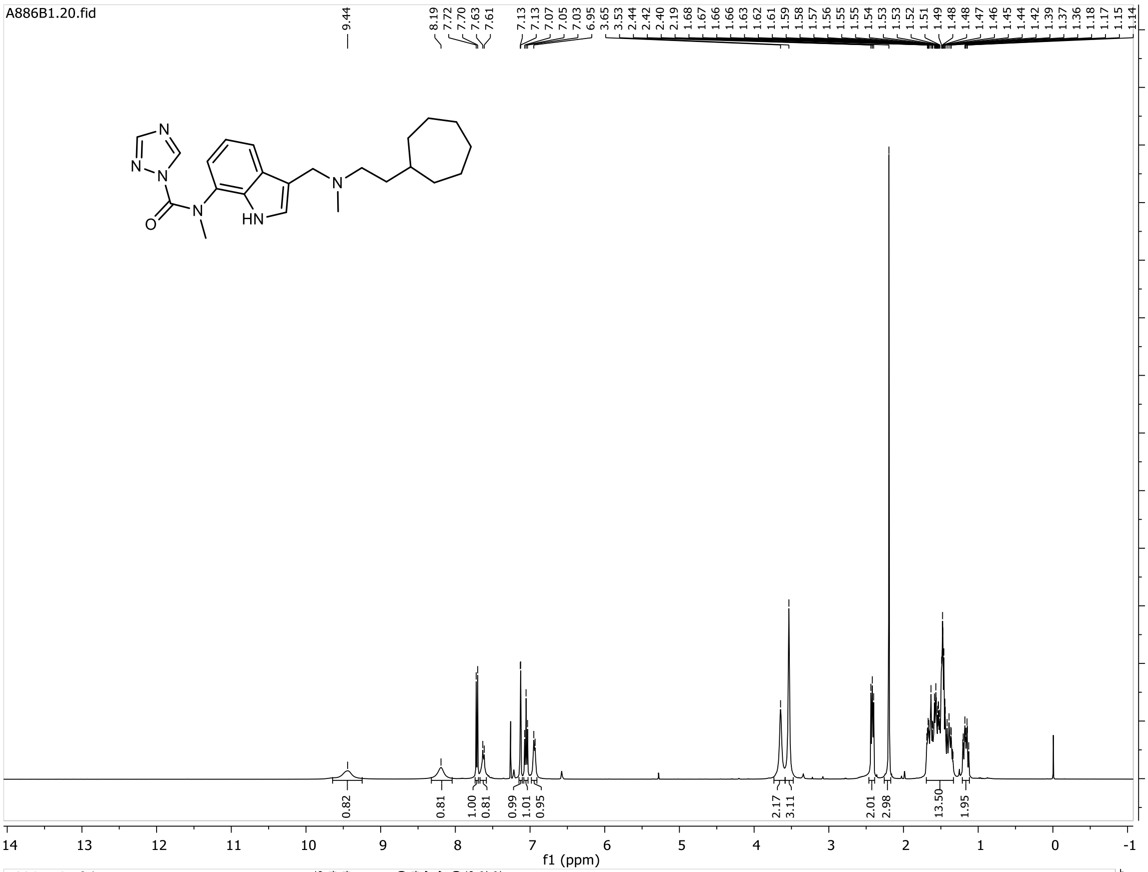
A886B1.20.fid
9.44
8.19 7.72 7.70 7.63 7.61
7.13 7.13 7.07 7.05 7.03 6.95 3.65 3.53 2.44 2.42 2.40 2.19 1.68 1.67 1.66 1.66 1.63 1.62 1.61 1.59 1.58 1.57 1.56 1.55 1.55 1.54 1.53 1.53 1.53 1.52 1.51 1.49 1.48 1.48 1.47 1.46 1.45 1.44 1.42 1.39 1.37 1.36 1.18 1.17 1.15 1.14
N
N
N
O
N
HN
N
0.82 0.81 1.00 0.81 0.99 1.01 0.95 2.17 3.11 2.01 2.98 13.50 1.95
f1 (ppm)

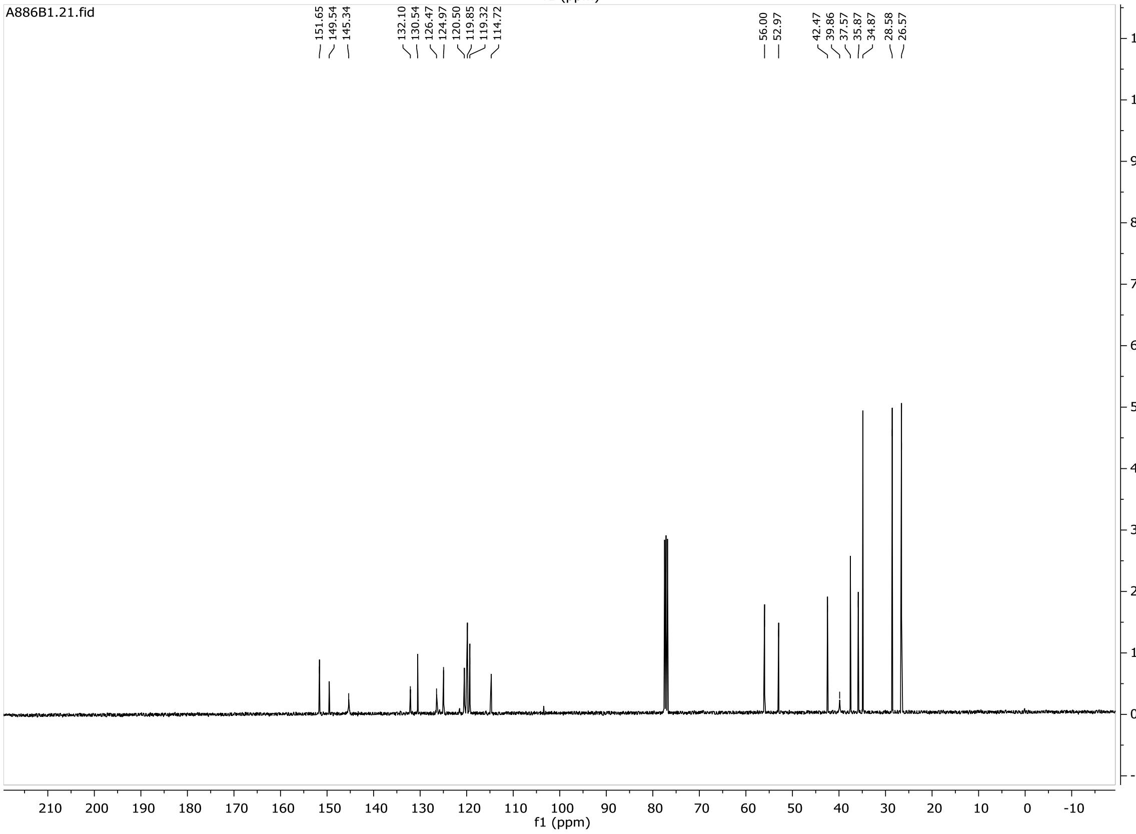
A886B1.21.fid
151.65 149.54 145.34
132.10 130.54 126.47 124.97 120.50 119.85 119.32 114.72
56.00 52.97
42.47 39.86 37.57 35.87 34.87 28.58 26.57
f1 (ppm)

**29**

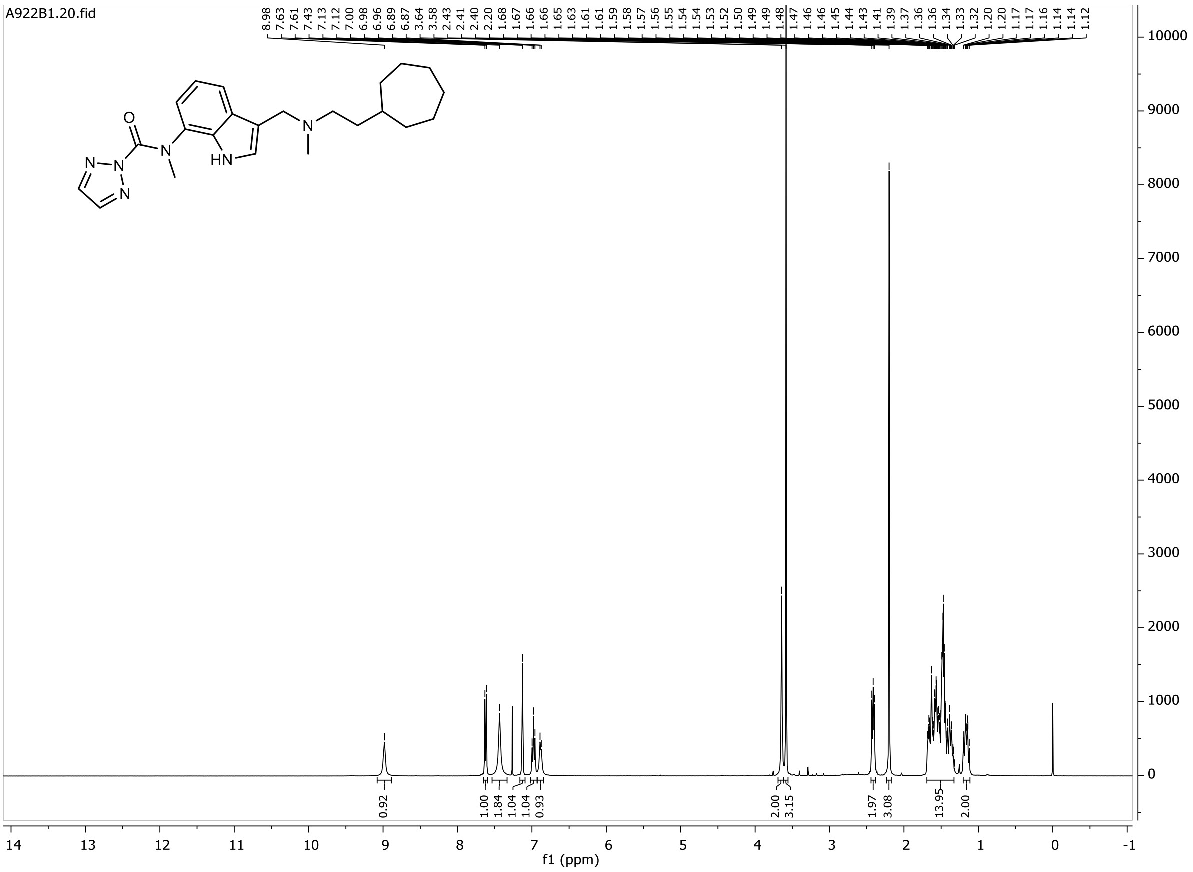
A922B1.20.fid
8.98 7.63 7.61 7.43 7.13 7.12 7.00 6.98 6.96 6.89 6.87 3.64 3.58 2.43 2.41 2.40 2.20 1.68 1.67 1.66 1.66 1.65 1.63 1.61 1.61 1.59 1.58 1.57 1.56 1.55 1.54 1.54 1.53 1.52 1.50 1.49 1.49 1.48 1.47 1.46 1.46 1.45 1.44 1.43 1.41 1.39 1.37 1.36 1.36 1.34 1.33 1.32 1.20 1.20 1.17 1.17 1.16 1.14 1.14 1.12
0.92 1.00 1.84 1.04 1.04 0.93 2.00 3.15 1.97 3.08 13.95 2.00
f1 (ppm)

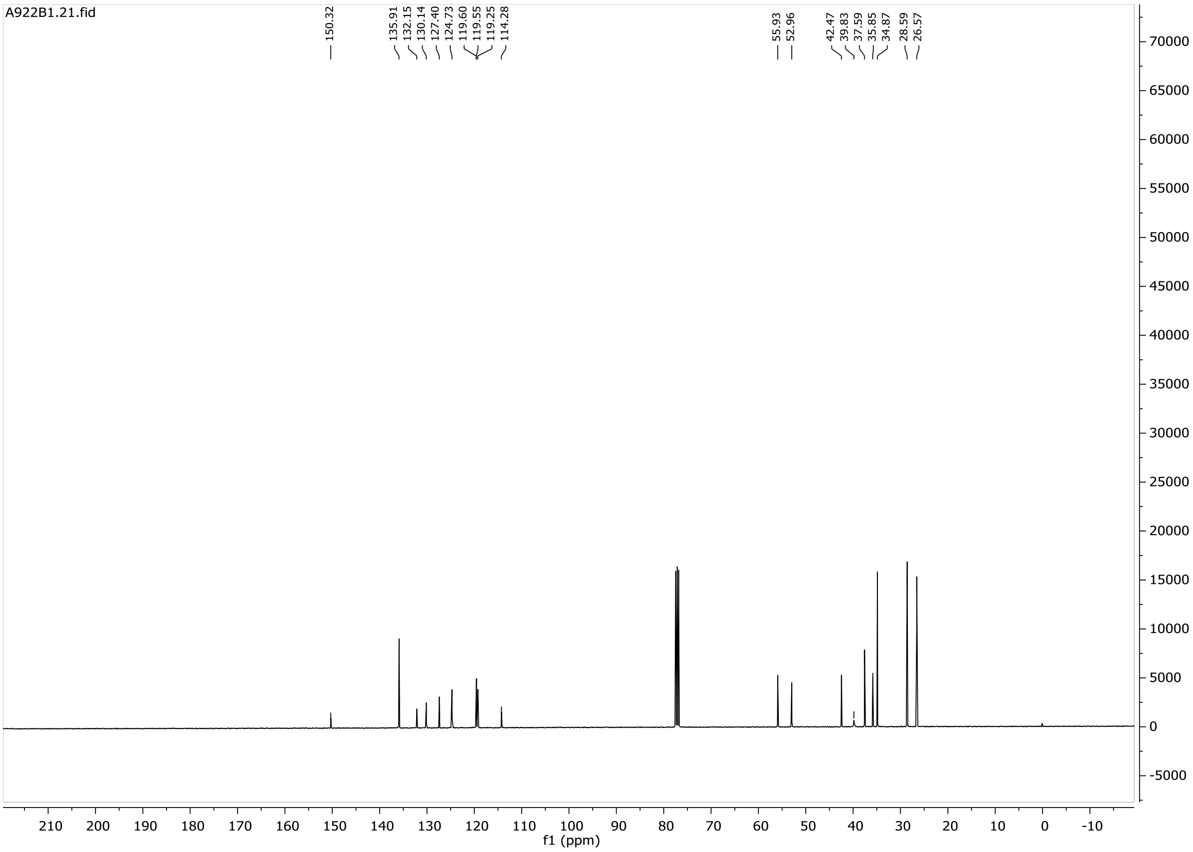
A922B1.21.fid
150.32 135.91 132.15 130.14 127.40 124.73 119.60 119.55 119.25 114.28 55.93 52.96 42.47 39.83 37.59 35.85 34.87 28.59 26.57
f1 (ppm)

**30**

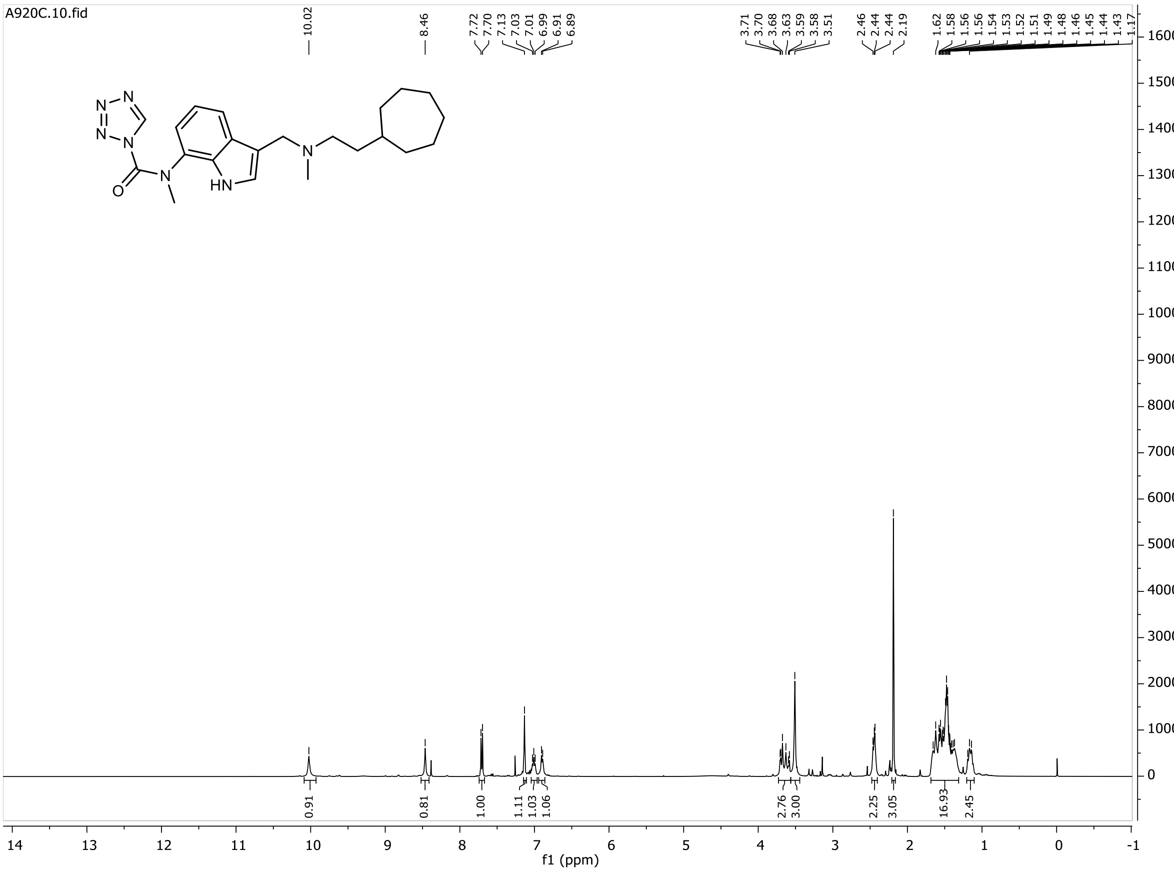
A920C.10.fid
10.02
8.46
7.72 7.70 7.13 7.03 7.01 6.99 6.91 6.89
3.71 3.70 3.68 3.63 3.59 3.58 3.51
2.46 2.44 2.44 2.19
1.62 1.58 1.56 1.56 1.54 1.53 1.52 1.51 1.49 1.48 1.46 1.45 1.44 1.43 1.17
0.91 0.81 1.00 1.11 1.03 1.06 2.76 3.00 2.25 3.05 16.93 2.45
f1 (ppm)

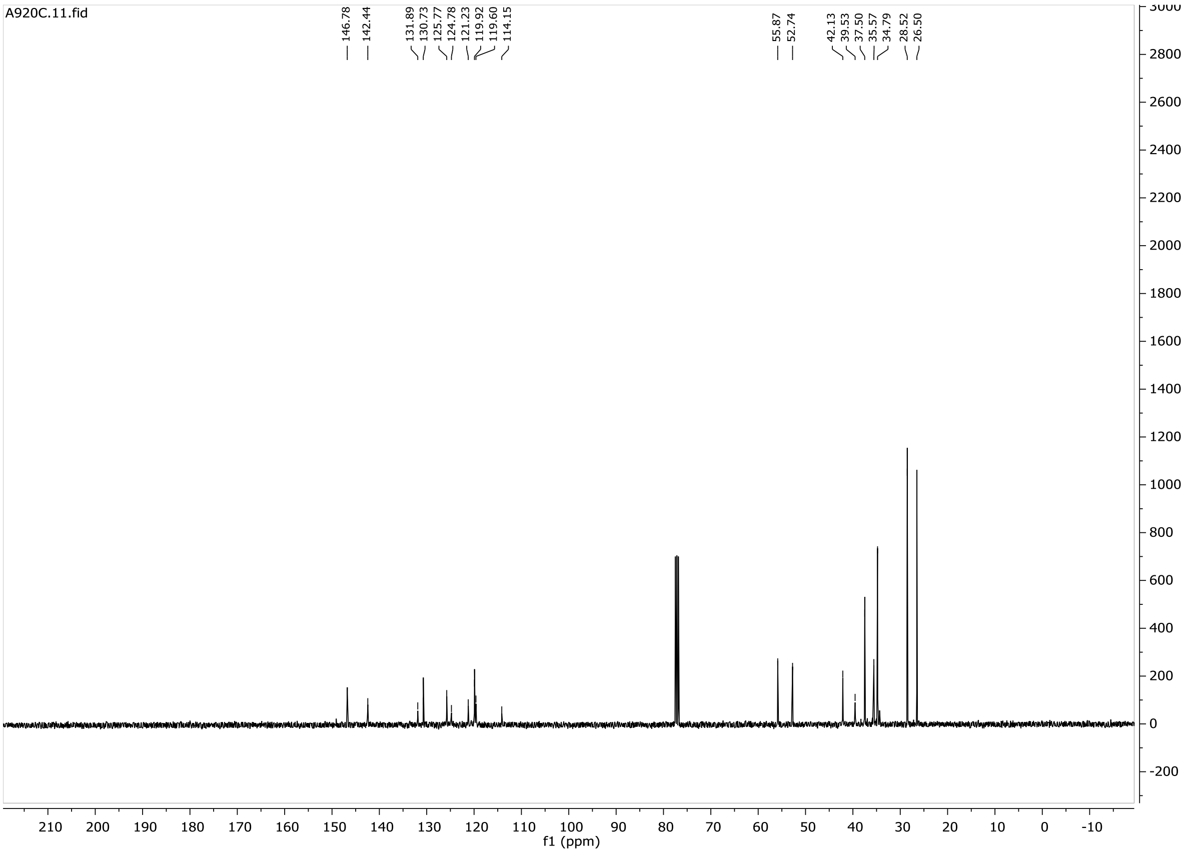
A920C.11.fid
146.78 142.44
131.89 130.73 125.77 124.78 121.23 119.92 119.60 114.15
55.87 52.74
42.13 39.53 37.50 35.57 34.79 28.52 26.50
f1 (ppm)

**31**

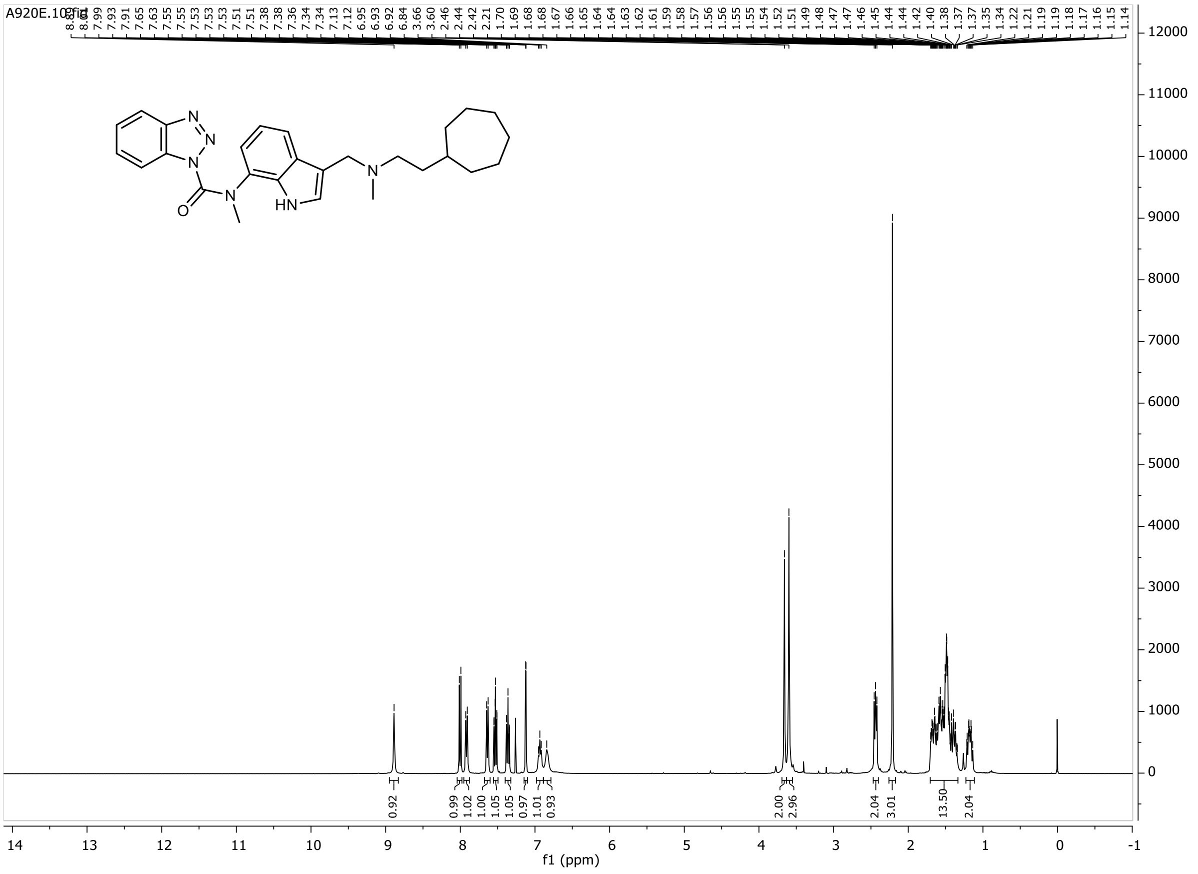
A920E.10.fid
7.99
7.93
7.91
7.65
7.63
7.55
7.55
7.53
7.53
7.53
7.51
7.51
7.38
7.38
7.36
7.34
7.34
7.13
7.12
6.95
6.93
6.92
6.84
3.66
3.60
2.46
2.44
2.42
2.21
1.70
1.69
1.68
1.68
1.67
1.66
1.65
1.64
1.64
1.63
1.62
1.61
1.59
1.58
1.57
1.56
1.56
1.55
1.55
1.54
1.52
1.51
1.49
1.48
1.47
1.47
1.46
1.45
1.44
1.44
1.42
1.40
1.38
1.37
1.37
1.35
1.34
1.22
1.21
1.19
1.19
1.18
1.17
1.16
1.15
1.14
N
N
N
O
N
HN
N
0.92
0.99
1.02
1.00
1.05
1.05
0.97
1.01
0.93
2.00
2.96
2.04
3.01
13.50
2.04
f1 (ppm)

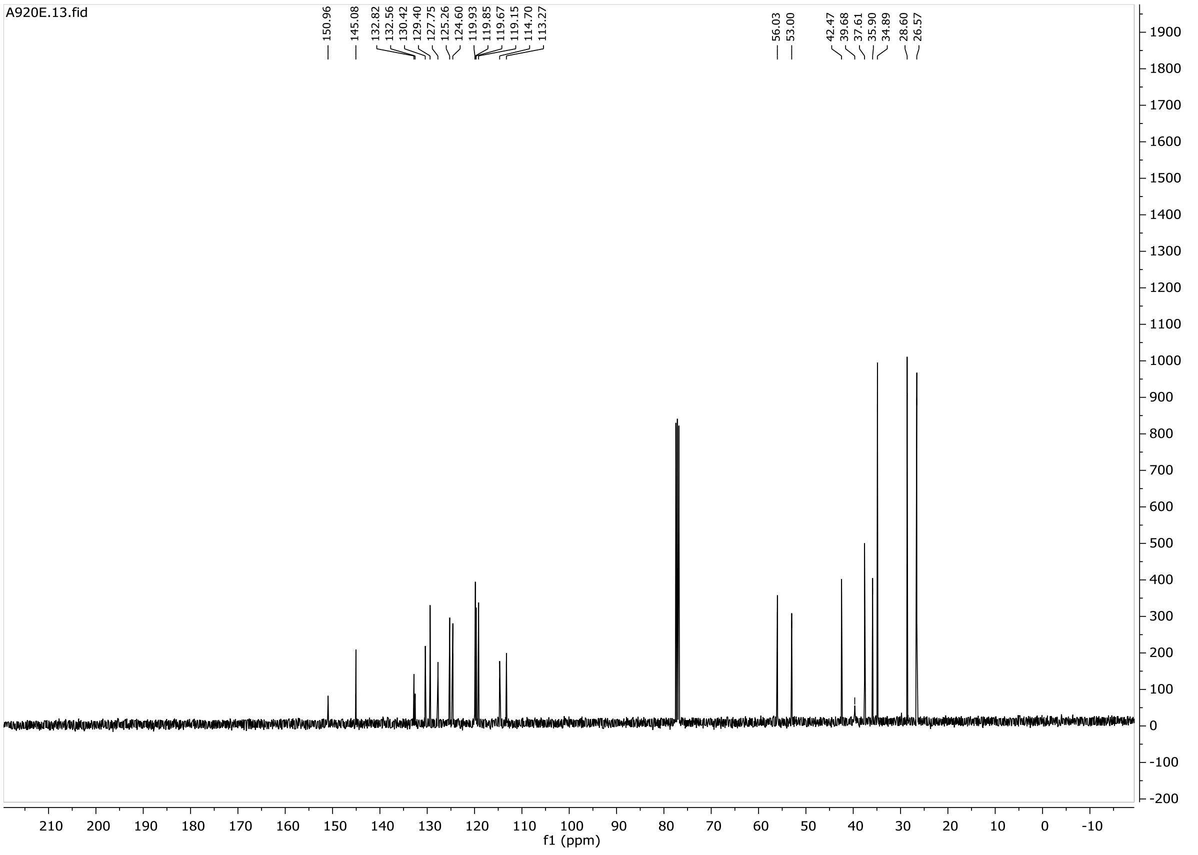
A920E.13.fid
150.96
145.08
132.82
132.56
130.42
129.40
127.75
125.26
124.60
119.93
119.85
119.67
119.15
114.70
113.27
56.03
53.00
42.47
39.68
37.61
35.90
34.89
28.60
26.57
f1 (ppm)

# 32

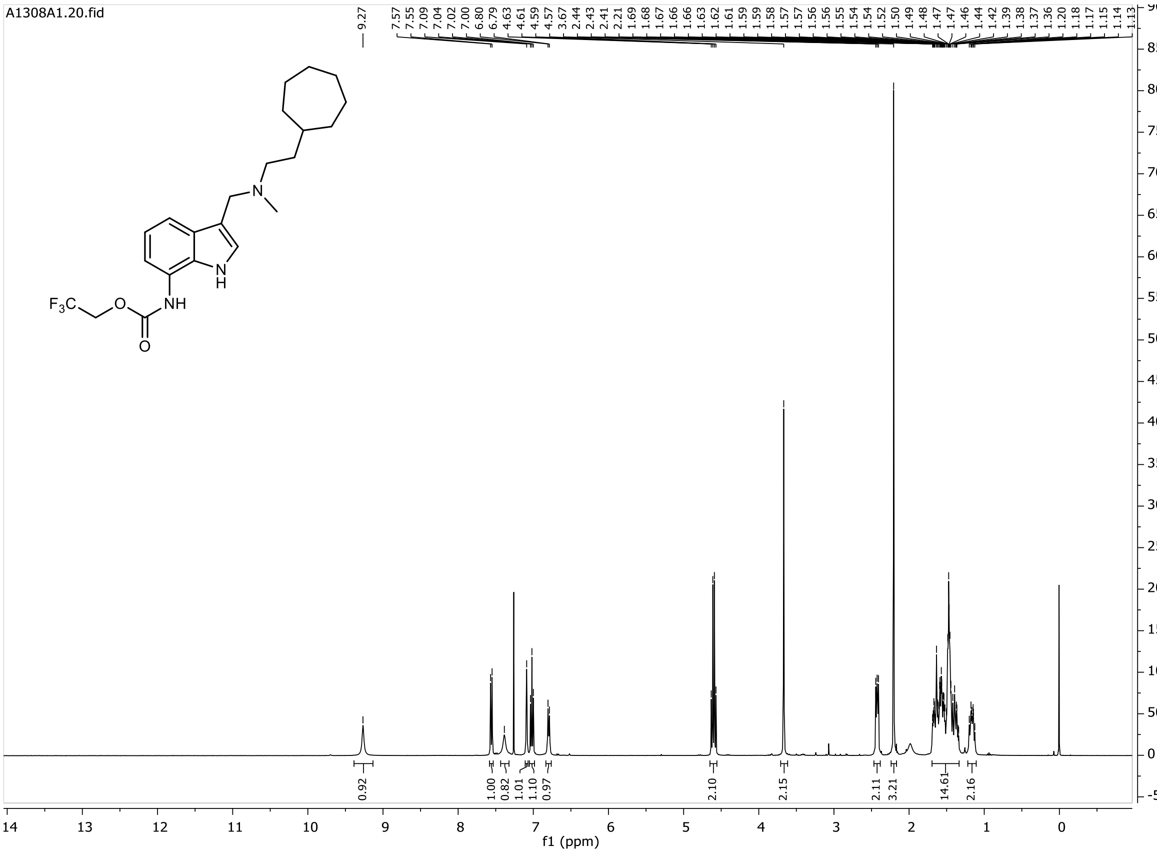

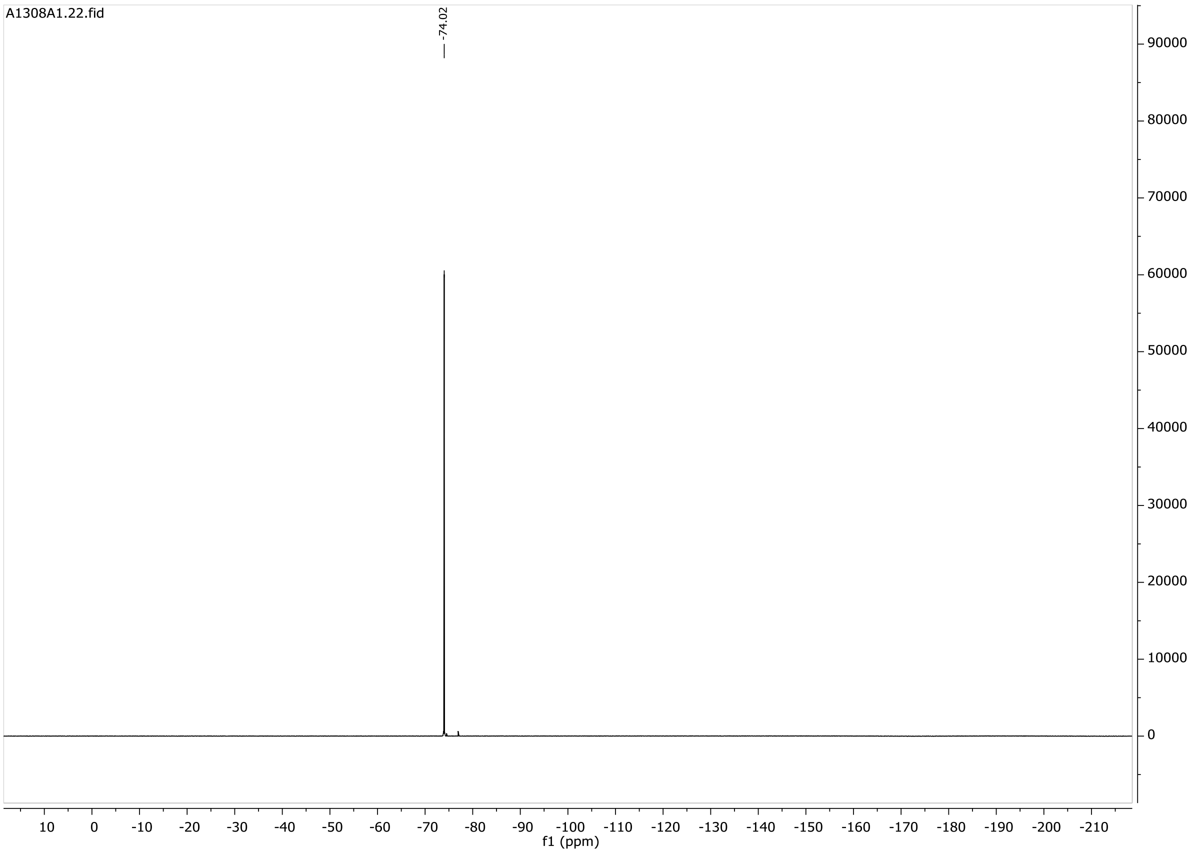
A1308A1.22.fid
-74.02
f1 (ppm)

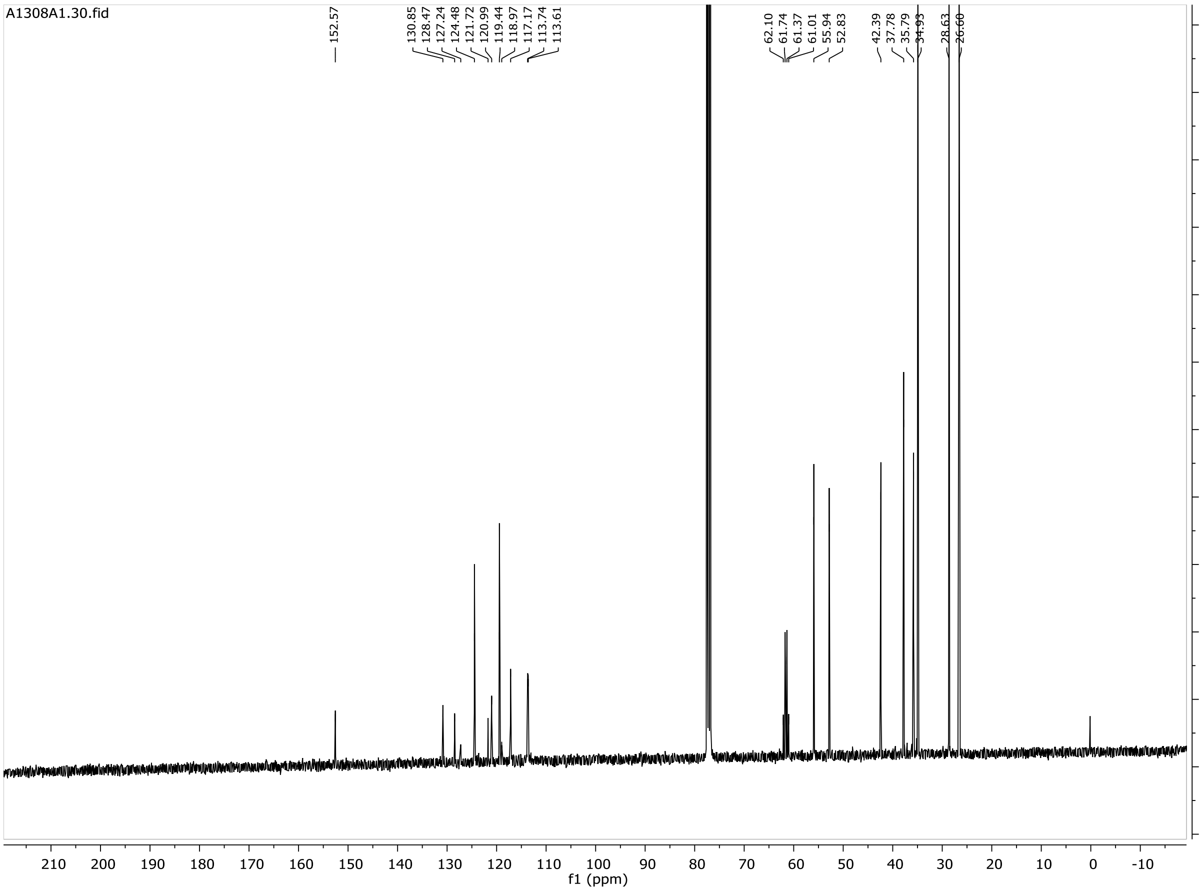
A1308A1.30.fid
152.57
130.85
128.47
127.24
124.48
121.72
120.99
119.44
118.97
117.17
113.74
113.61
62.10
61.74
61.37
61.01
55.94
52.83
42.39
37.78
35.79
34.93
28.63
26.60
f1 (ppm)

# 33a

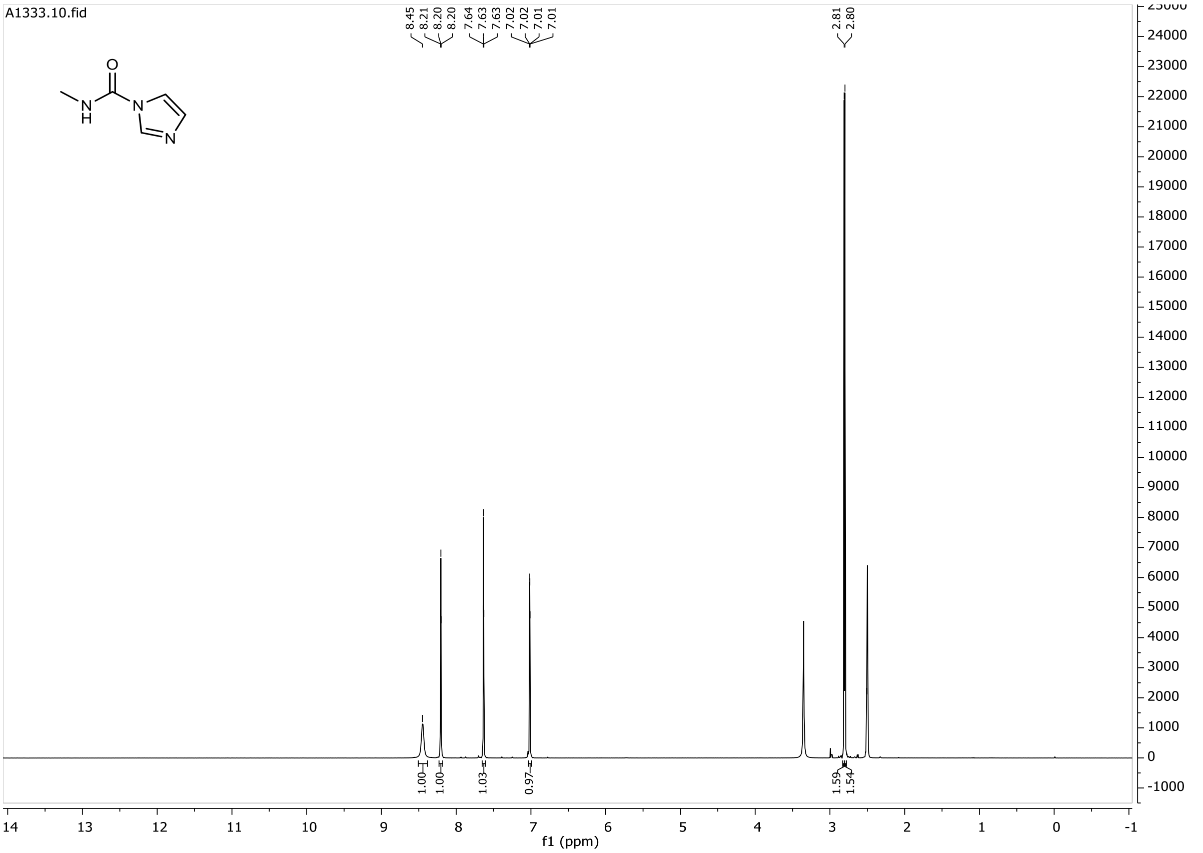

A1333.10.fid
8.45
8.21
8.20
8.20
7.64
7.63
7.63
7.02
7.02
7.01
7.01
2.81
2.80
1.00
1.00
1.03
0.97
1.59
1.54
f1 (ppm)

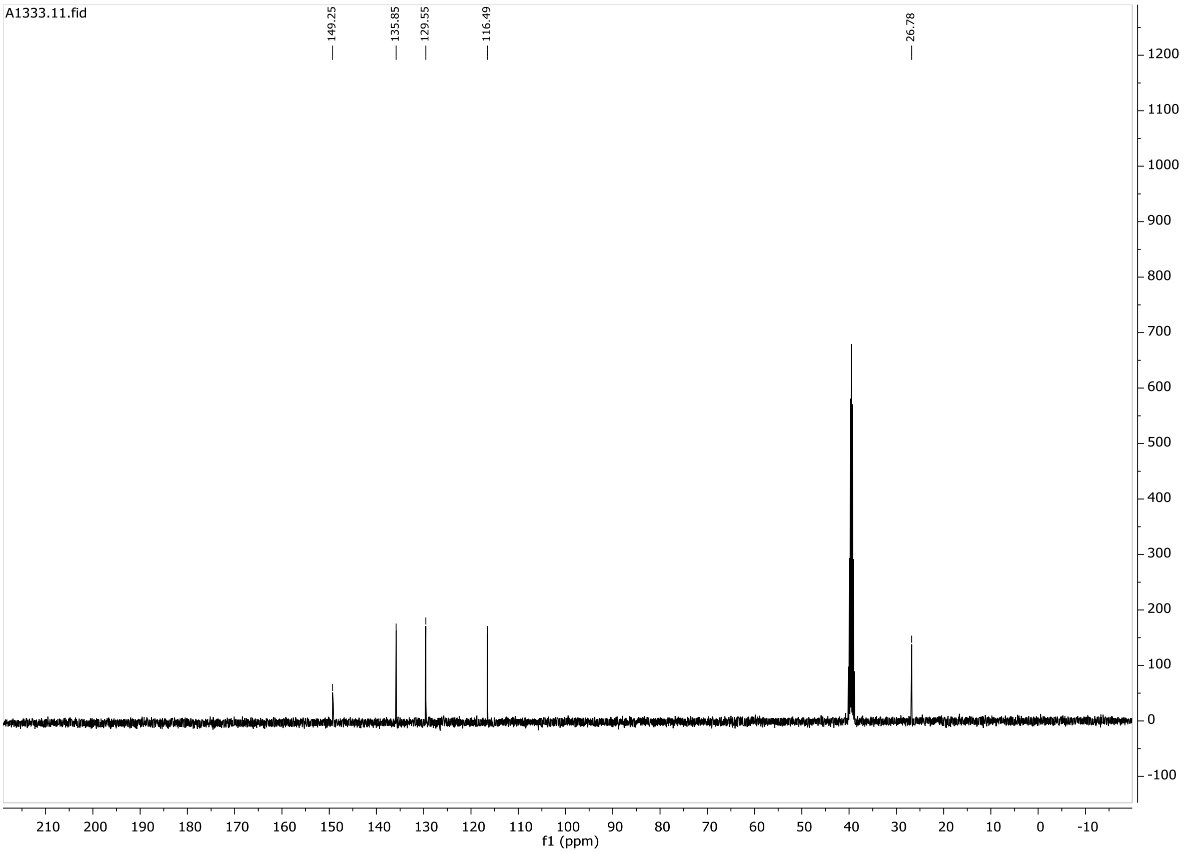

A1333.11.fid
149.25
135.85
129.55
116.49
26.78
f1 (ppm)

**33**

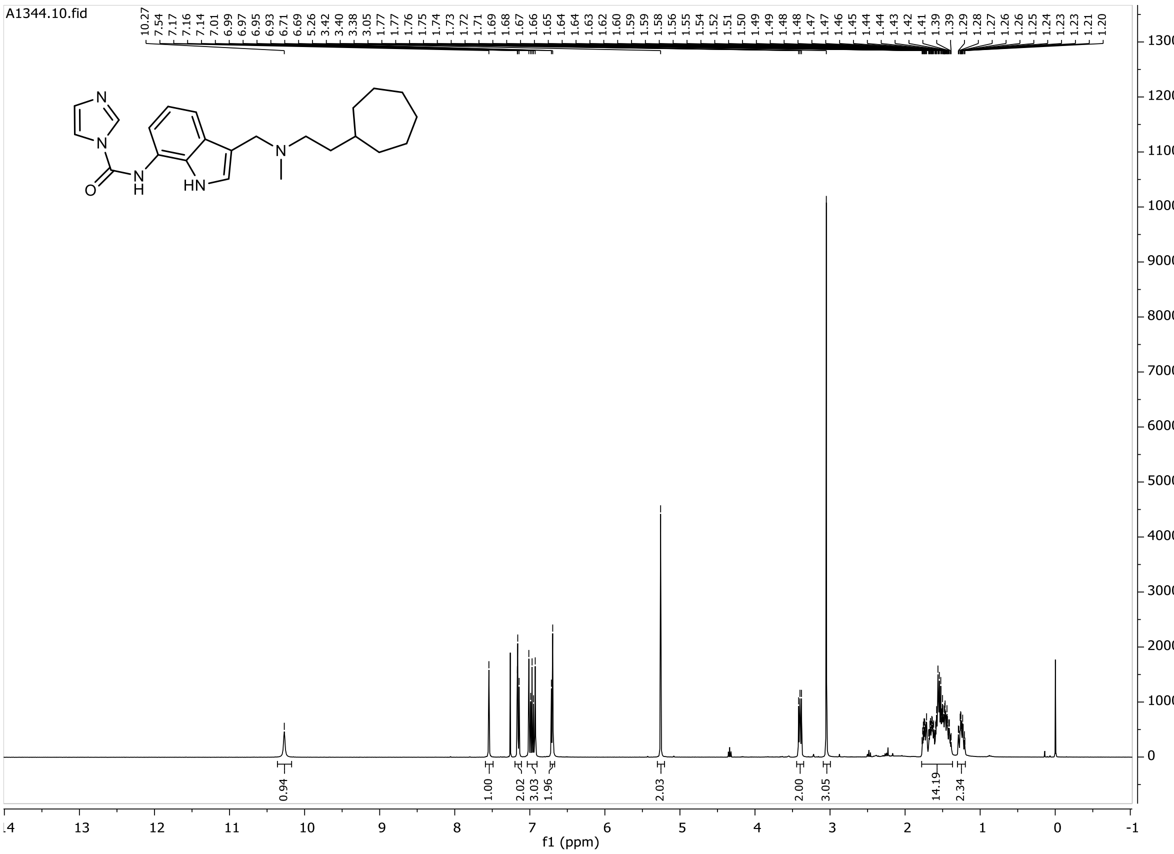

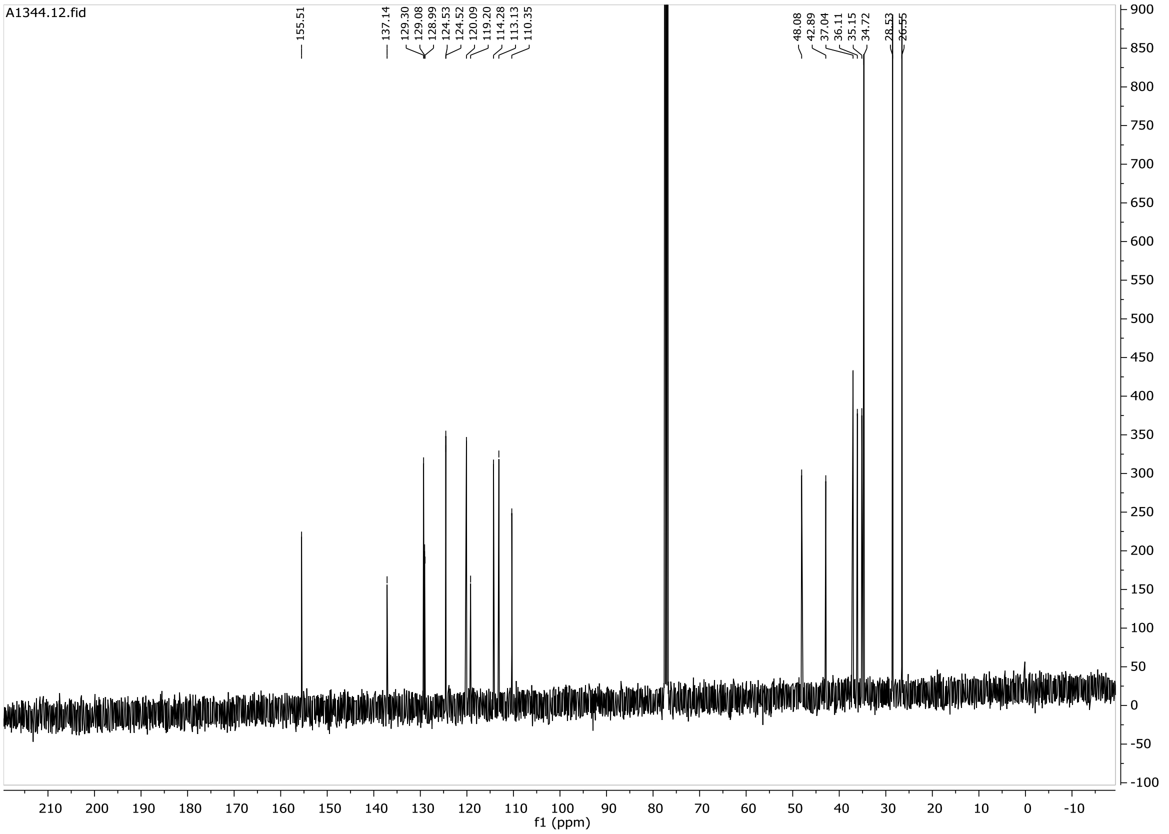

# P1

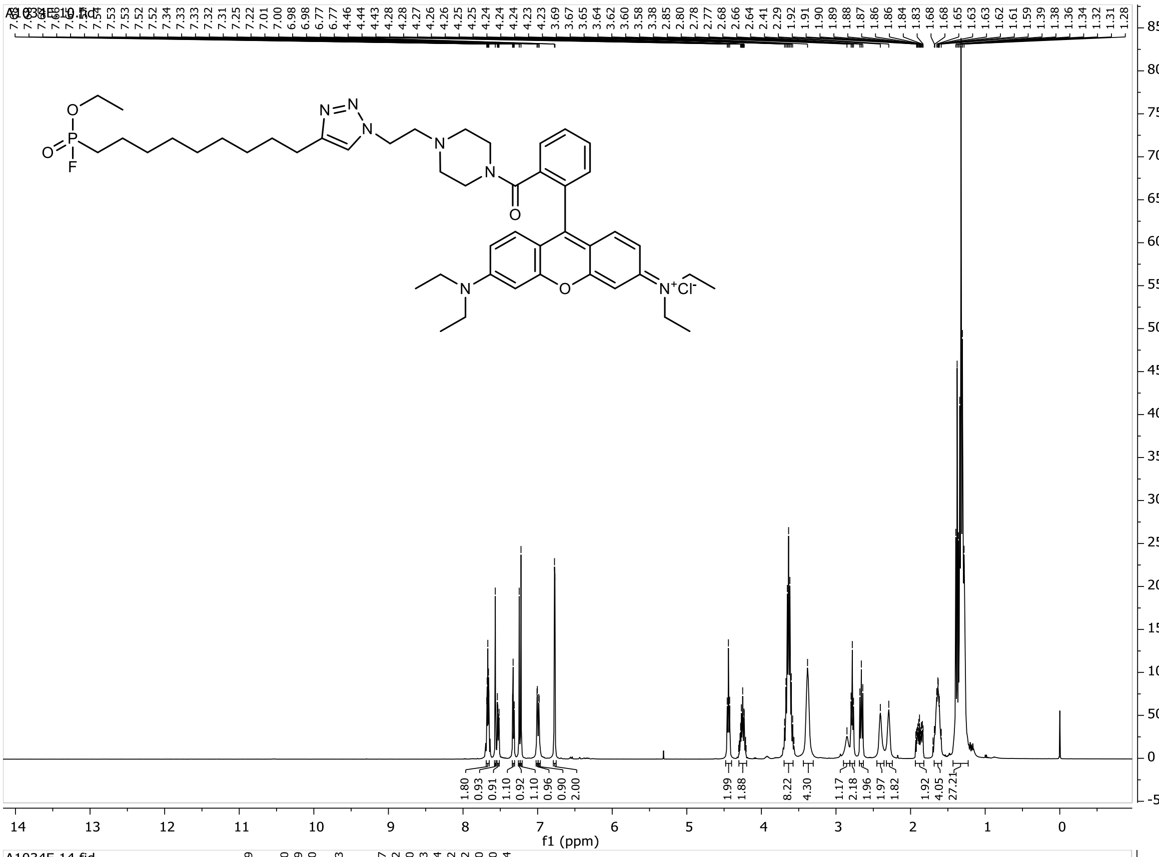

f1 (ppm)

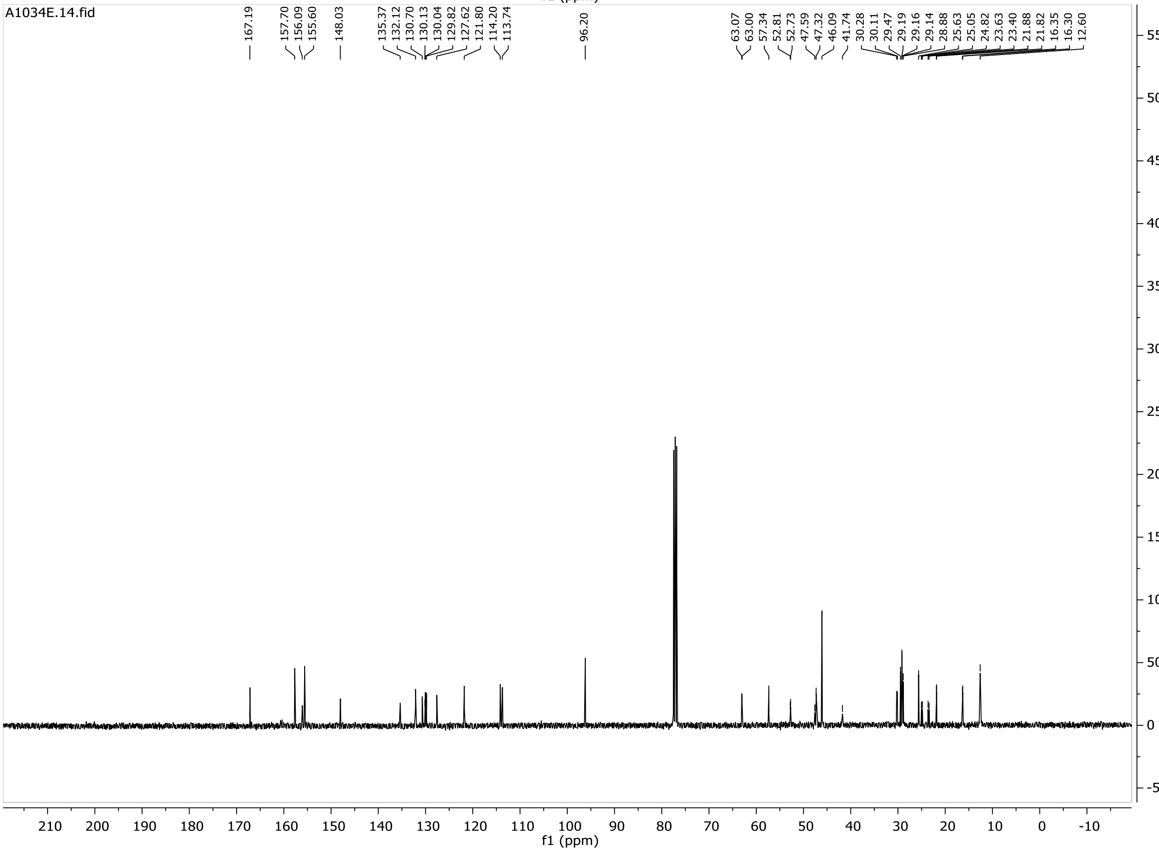

A1034E.14.fid
167.19
157.70
156.09
155.60
148.03
135.37
132.12
130.70
130.13
130.04
129.82
127.62
121.80
114.20
113.74
96.20
63.07
63.00
57.34
52.81
52.73
47.59
47.32
46.09
41.74
30.28
30.11
29.47
29.19
29.16
29.14
28.88
25.63
25.05
24.82
23.63
23.40
21.88
21.82
16.35
16.30
12.60
f1 (ppm)

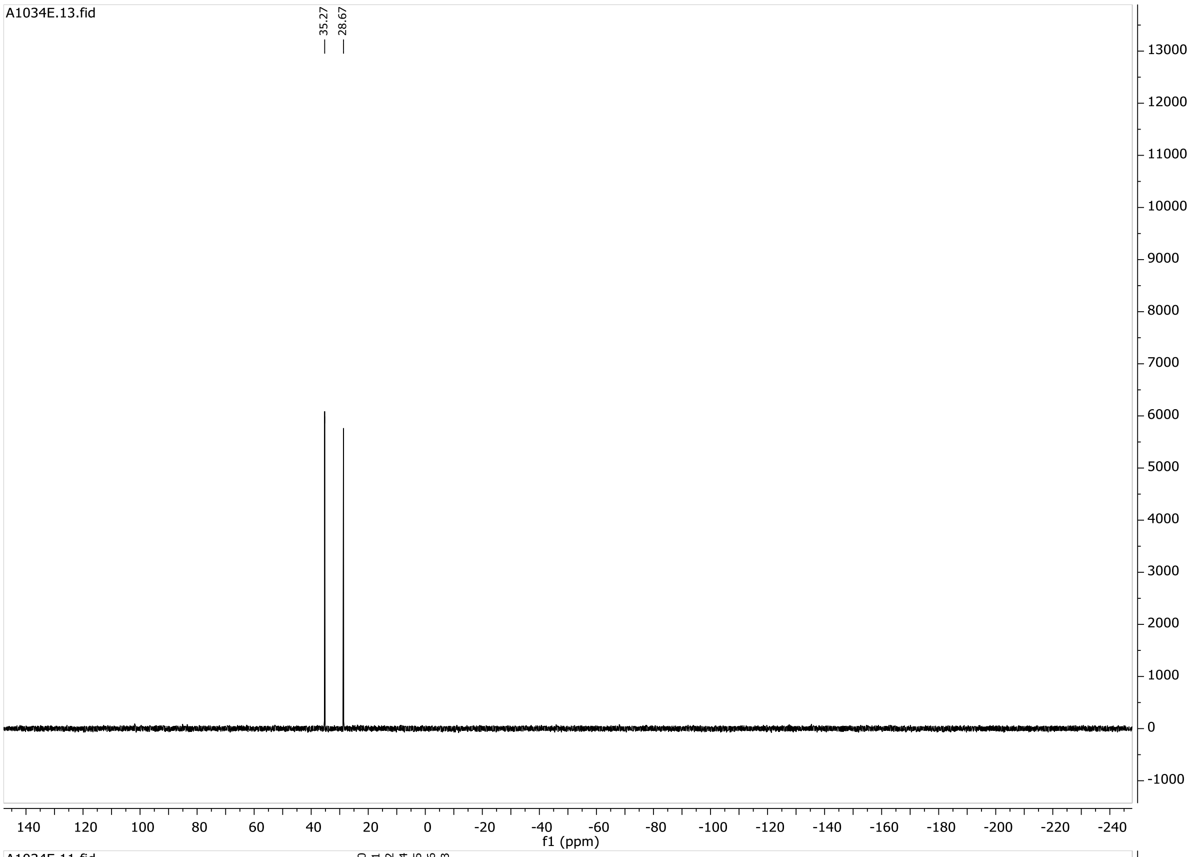
A1034E.13.fid
35.27
28.67
f1 (ppm)

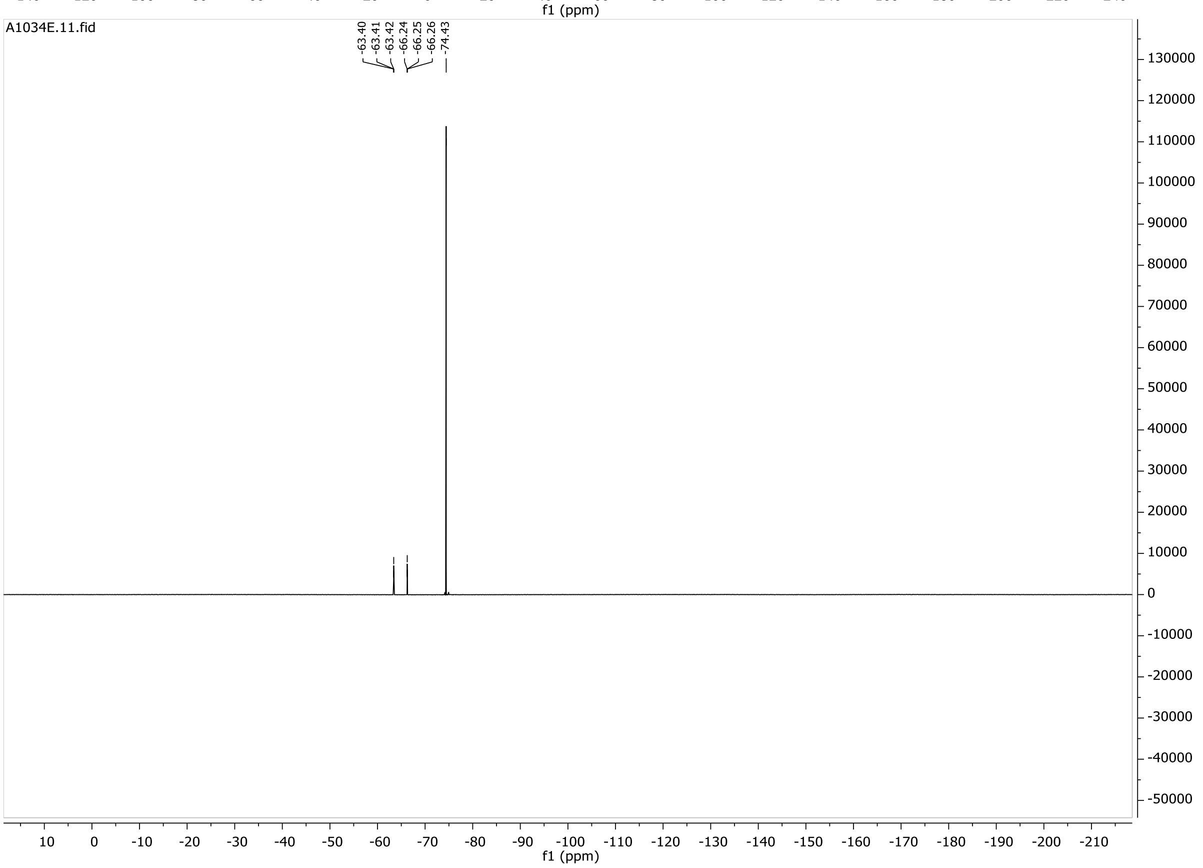
A1034E.11.fid
-63.40
-63.41
-63.42
-66.24
-66.25
-66.26
-74.43
f1 (ppm)

# P2

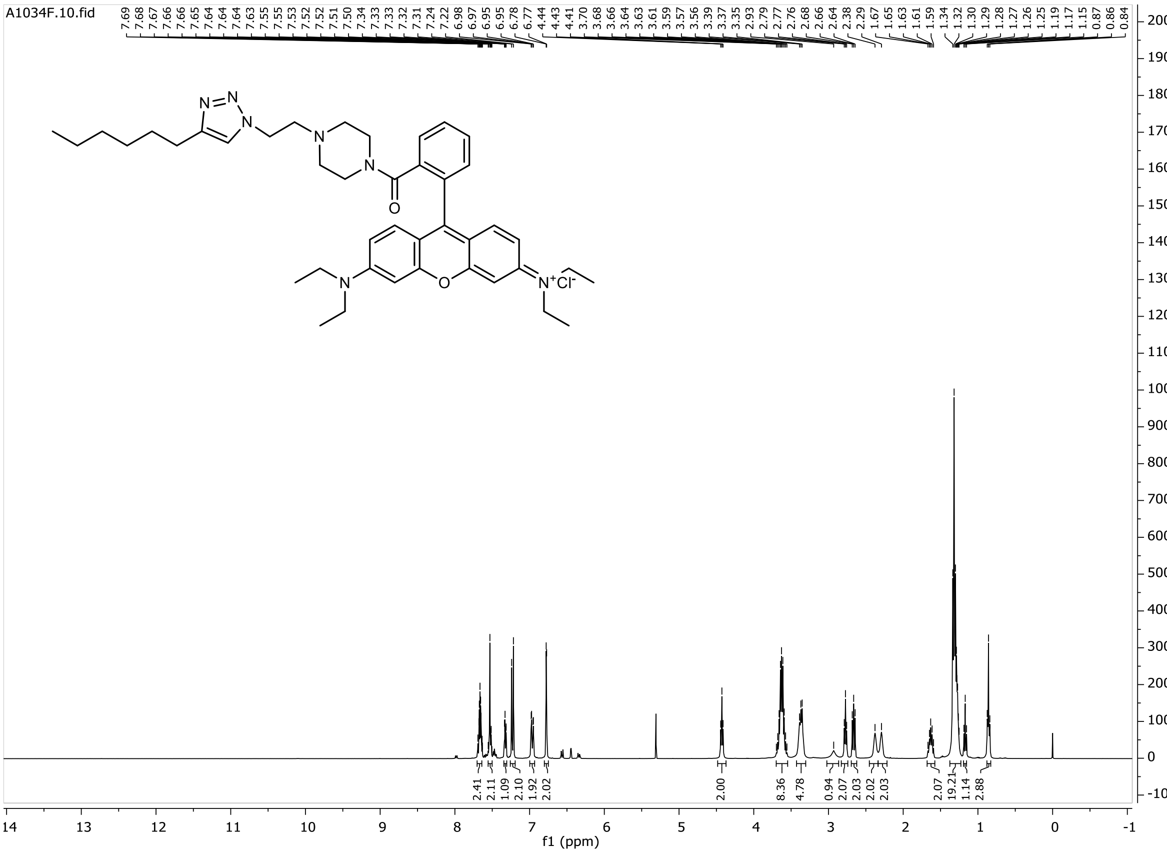

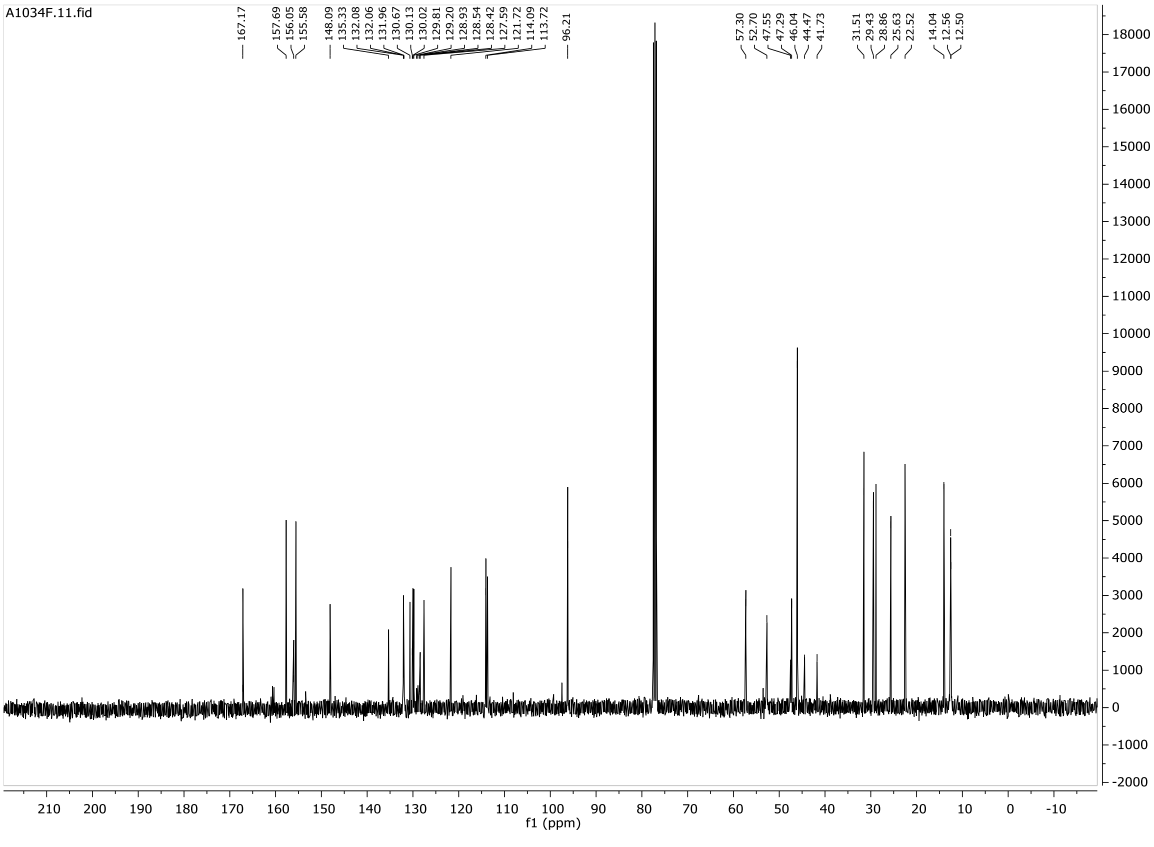

# P3

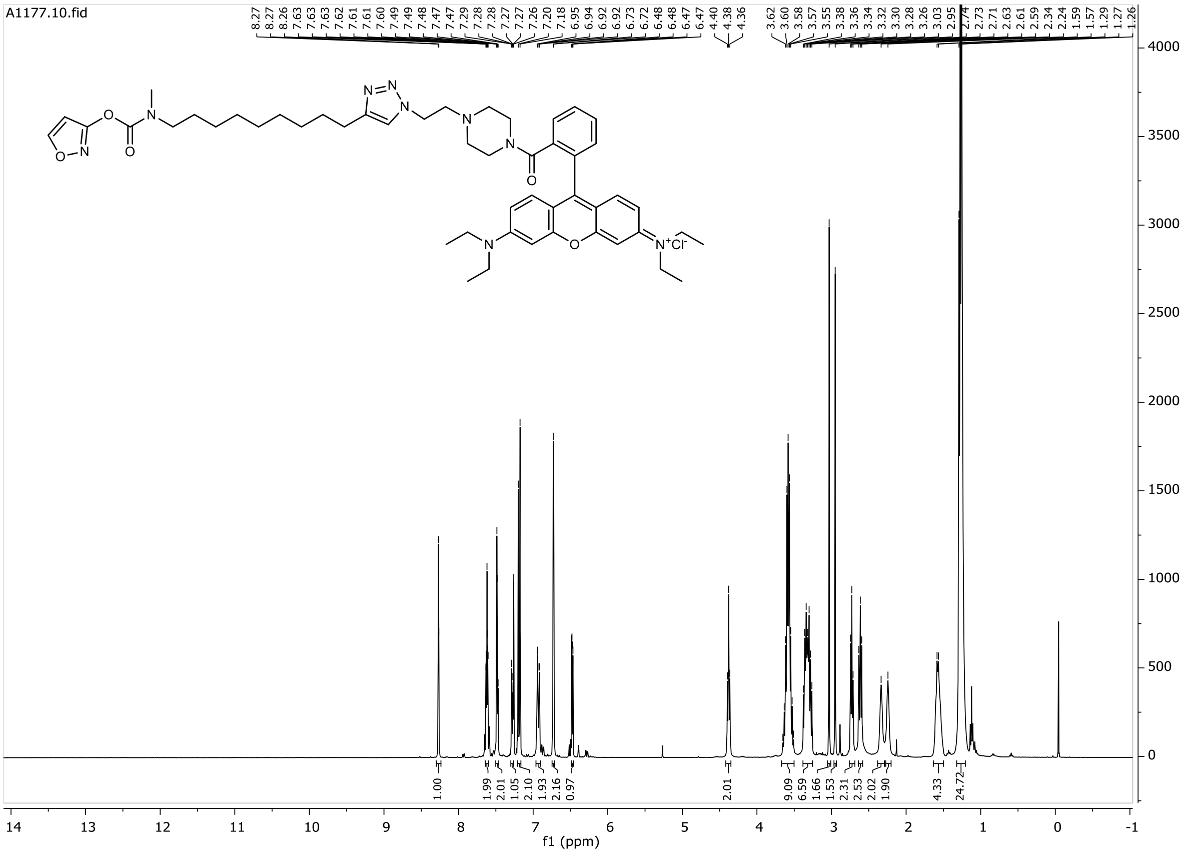

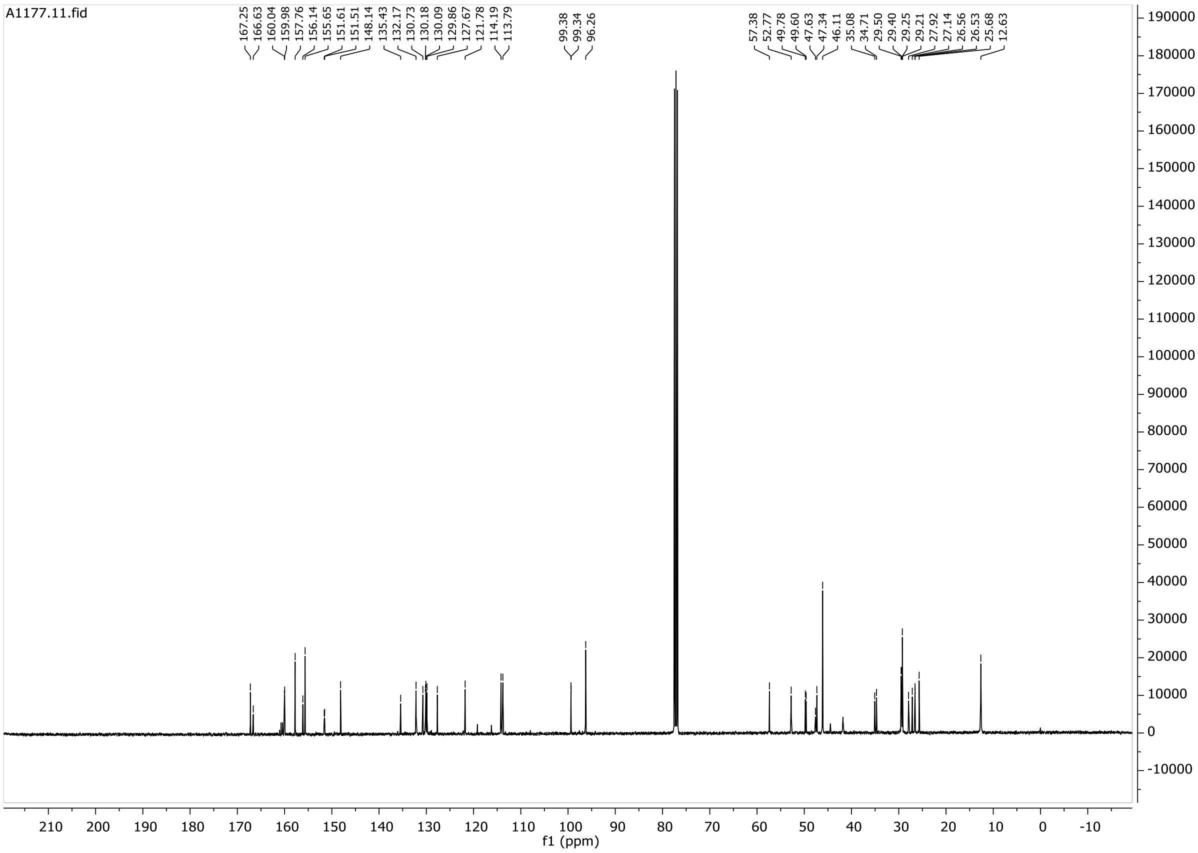

# P4

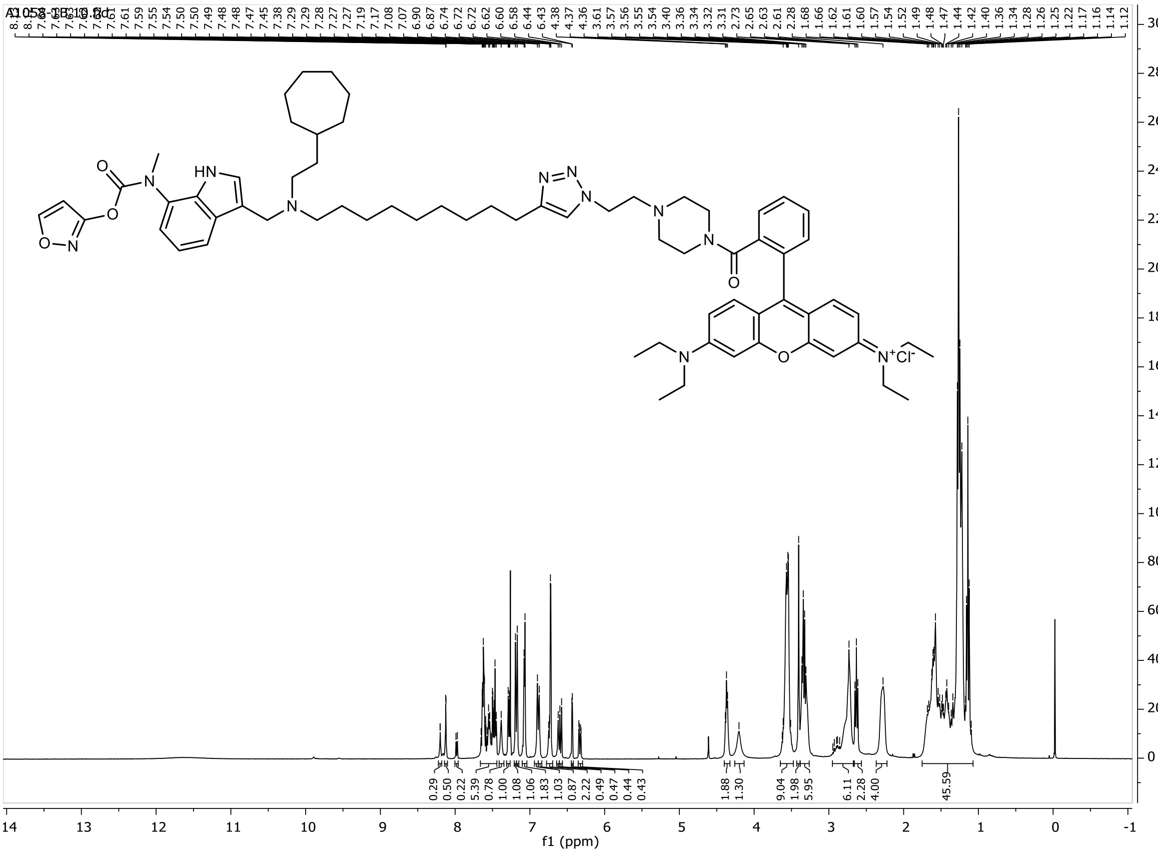

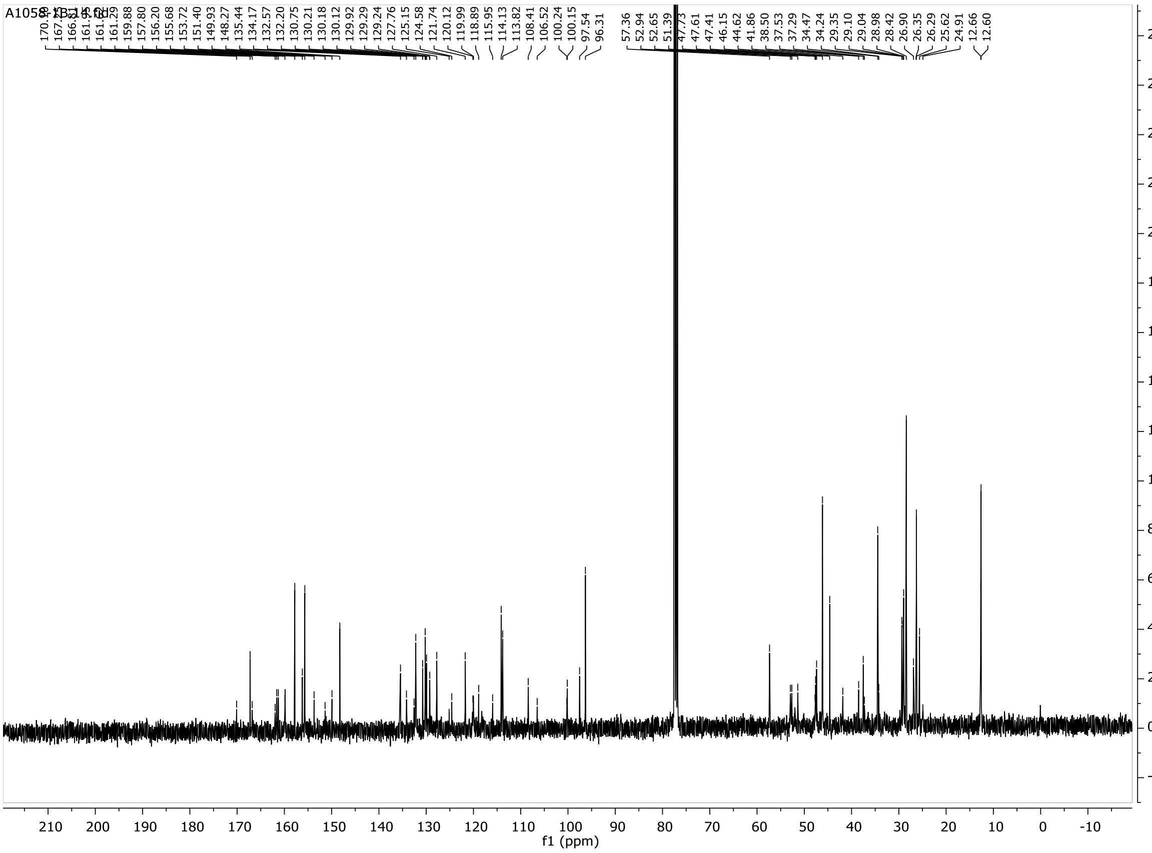

# P5

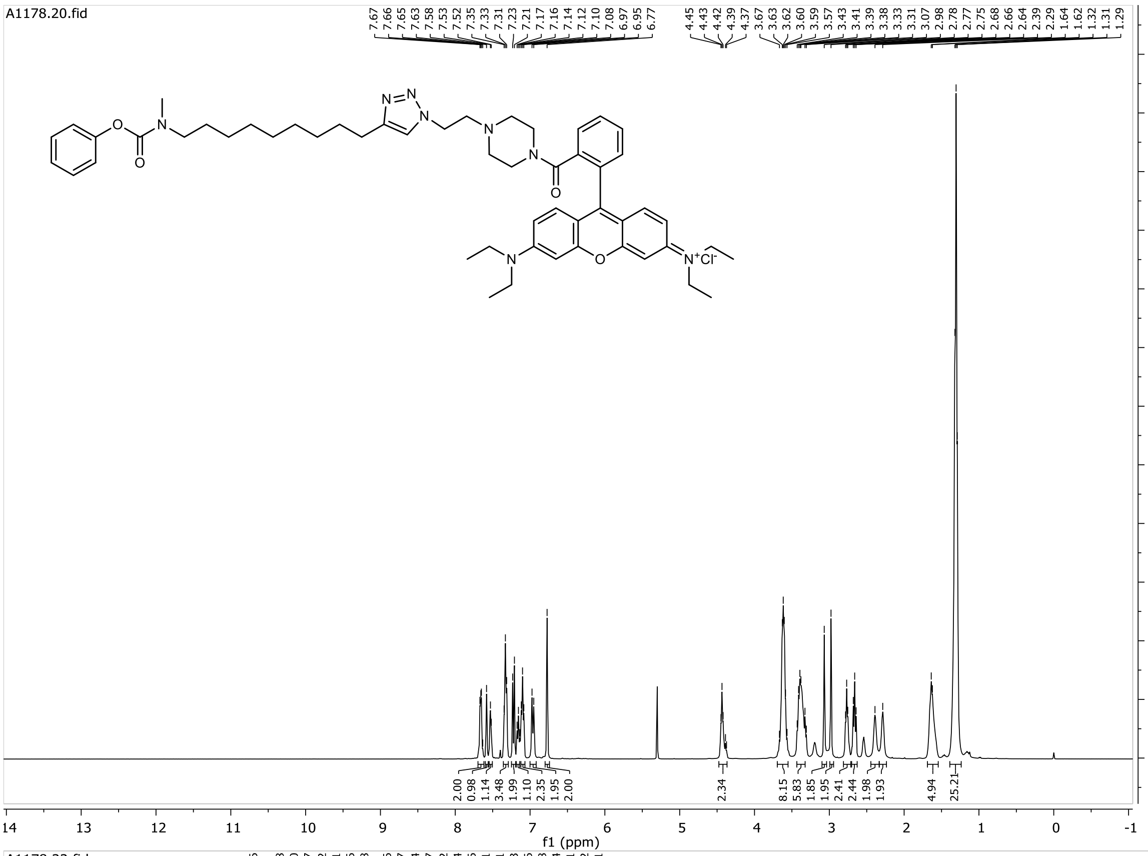
A1178.20.fid
7.67 7.66 7.65 7.63 7.58 7.53 7.52 7.35 7.33 7.31 7.23 7.21 7.17 7.16 7.14 7.12 7.10 7.08 6.97 6.95 6.77
4.45 4.43 4.42 4.39 4.37 3.67 3.63 3.62 3.60 3.59 3.57 3.43 3.41 3.39 3.38 3.33 3.31 3.07 2.98 2.78 2.77 2.75 2.68 2.66 2.64 2.39 2.29 1.64 1.62 1.32 1.31 1.29
2.00 0.98 1.14 3.48 1.99 1.10 2.35 1.95 2.00
2.34
8.15 5.83 1.85 1.95 2.41 2.44 1.98 1.93
4.94 25.21
f1 (ppm)

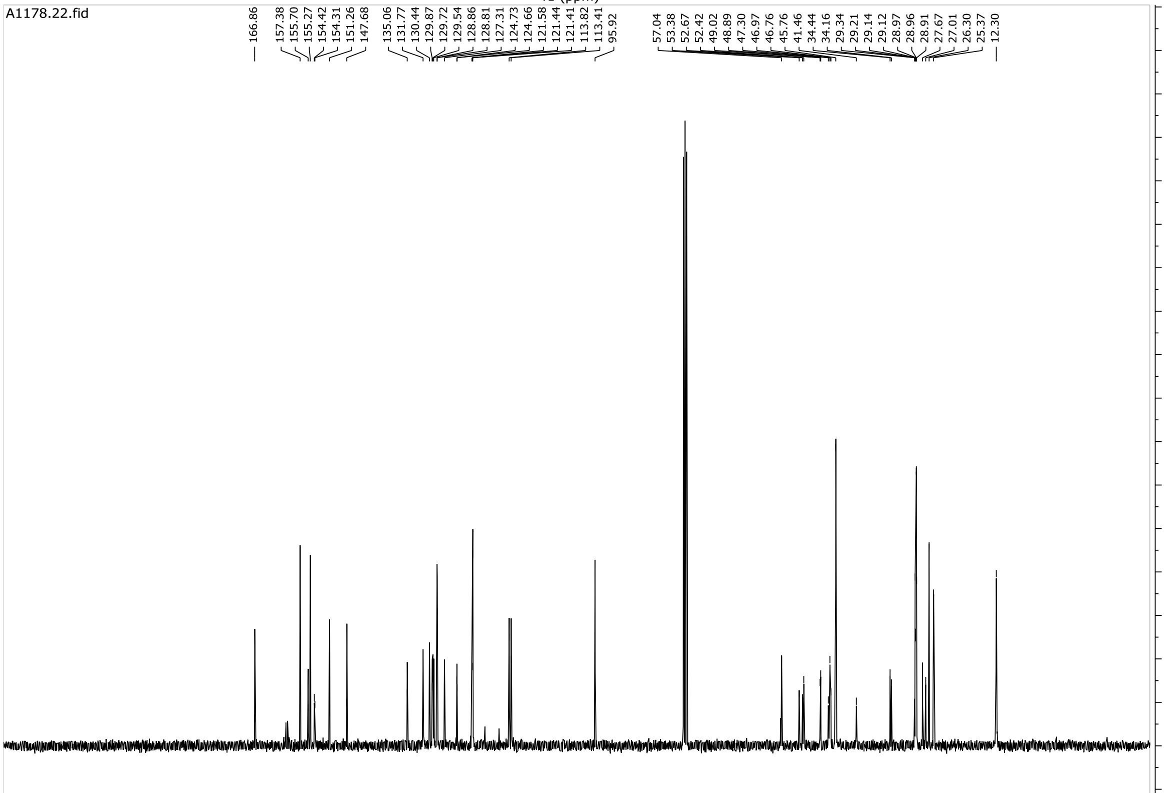
A1178.22.fid
166.86
157.38 155.70 155.27 154.42 154.31 151.26 147.68
135.06 131.77 130.44 129.87 129.72 129.54 128.86 128.81 127.31 124.73 124.66 121.58 121.44 121.41 113.82 113.41 95.92
57.04 53.38 52.67 52.42 49.02 48.89 47.30 46.97 46.76 45.76 41.46 34.44 34.16 29.34 29.21 29.14 29.12 28.97 28.96 28.91 27.67 27.01 26.30 25.37
12.30
f1 (ppm)

# B1

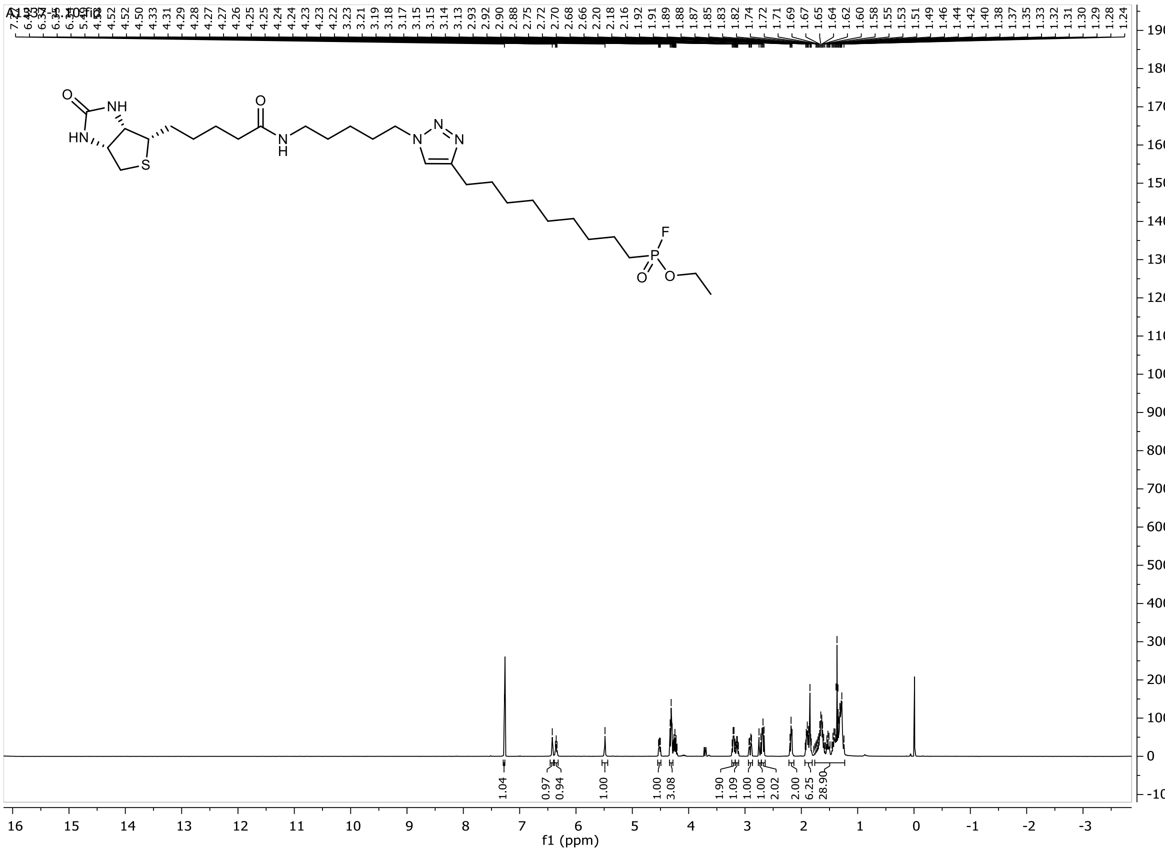
f1 (ppm)
1.04
0.97
0.94
1.00
1.00
3.08
1.90
1.09
1.00
1.00
2.02
2.00
6.25
28.90

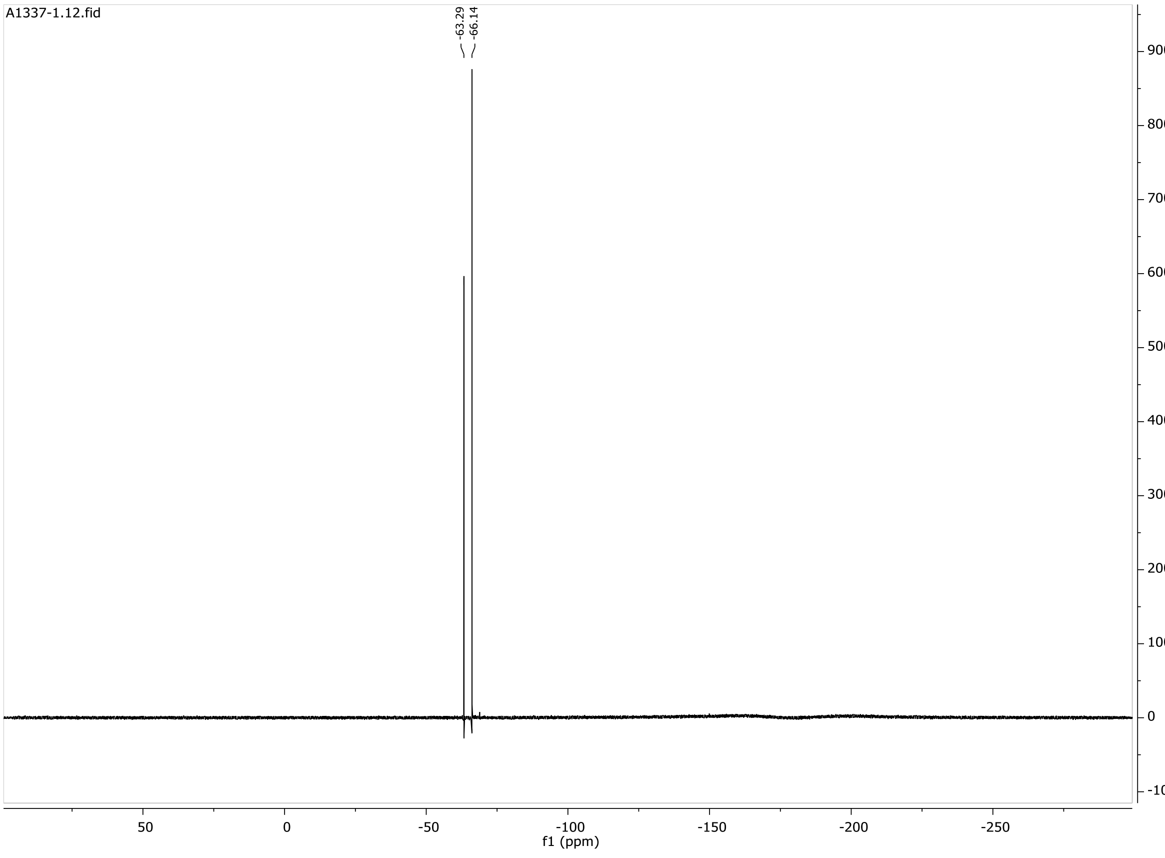
A1337-1.12.fid
-63.29
-66.14
f1 (ppm)

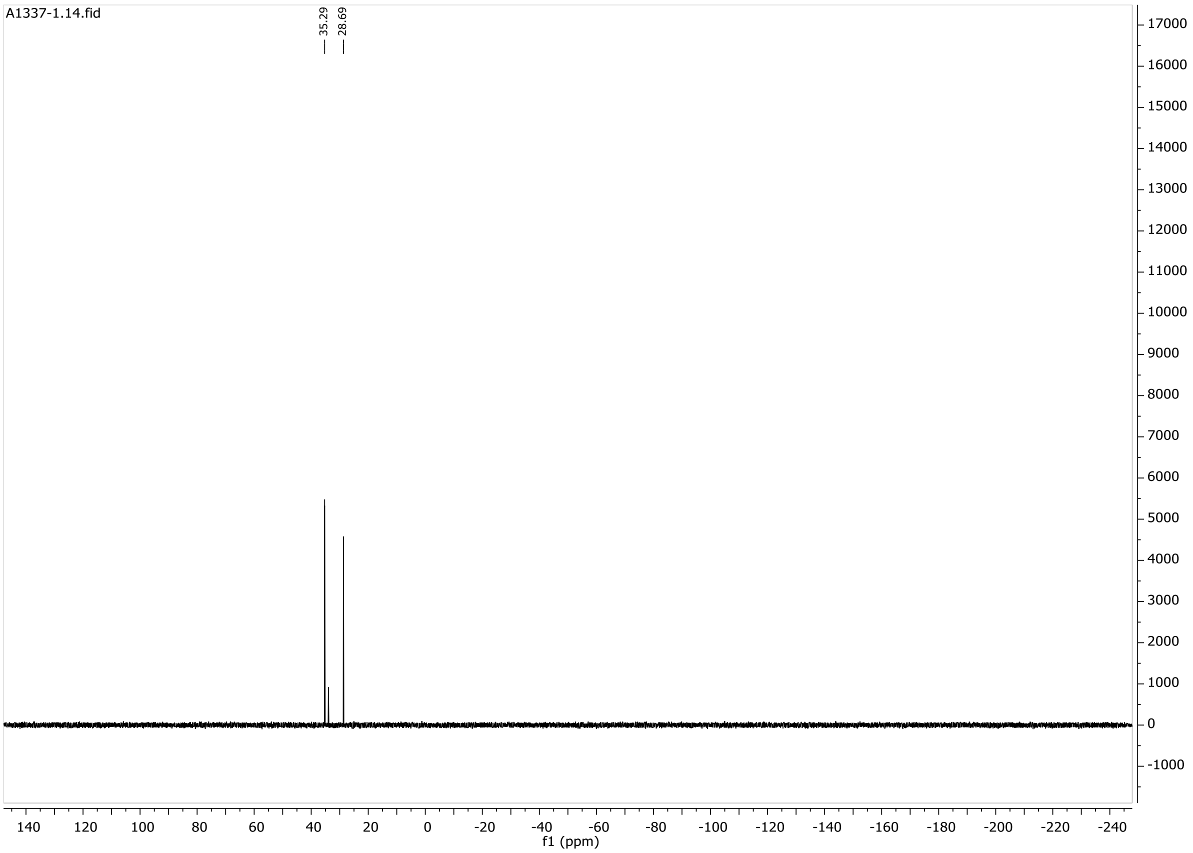
A1337-1.14.fid
35.29
28.69
f1 (ppm)

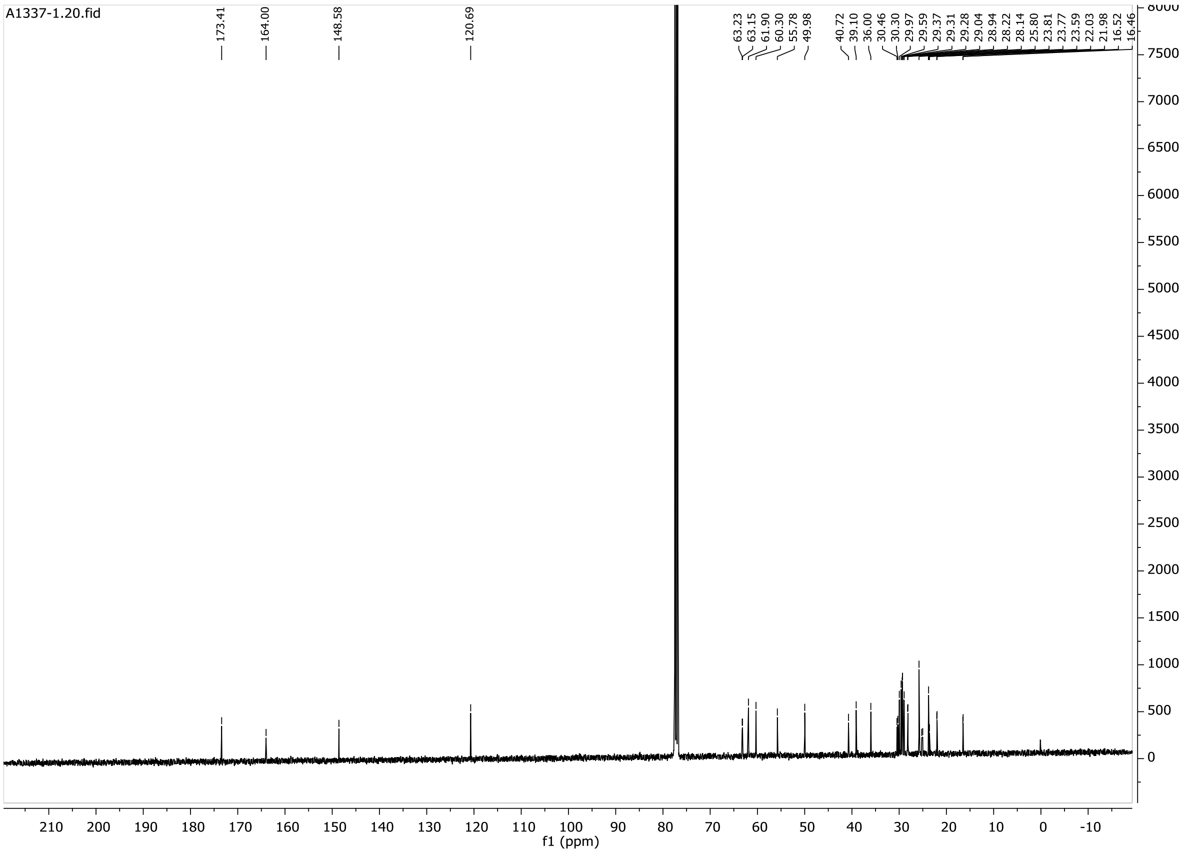
A1337-1.20.fid
173.41
164.00
148.58
120.69
63.23
63.15
61.90
60.30
55.78
49.98
40.72
39.10
36.00
30.46
30.30
29.97
29.59
29.37
29.31
29.28
29.04
28.94
28.22
28.14
25.80
23.81
23.77
23.59
22.03
21.98
16.52
16.46
f1 (ppm)

# B2

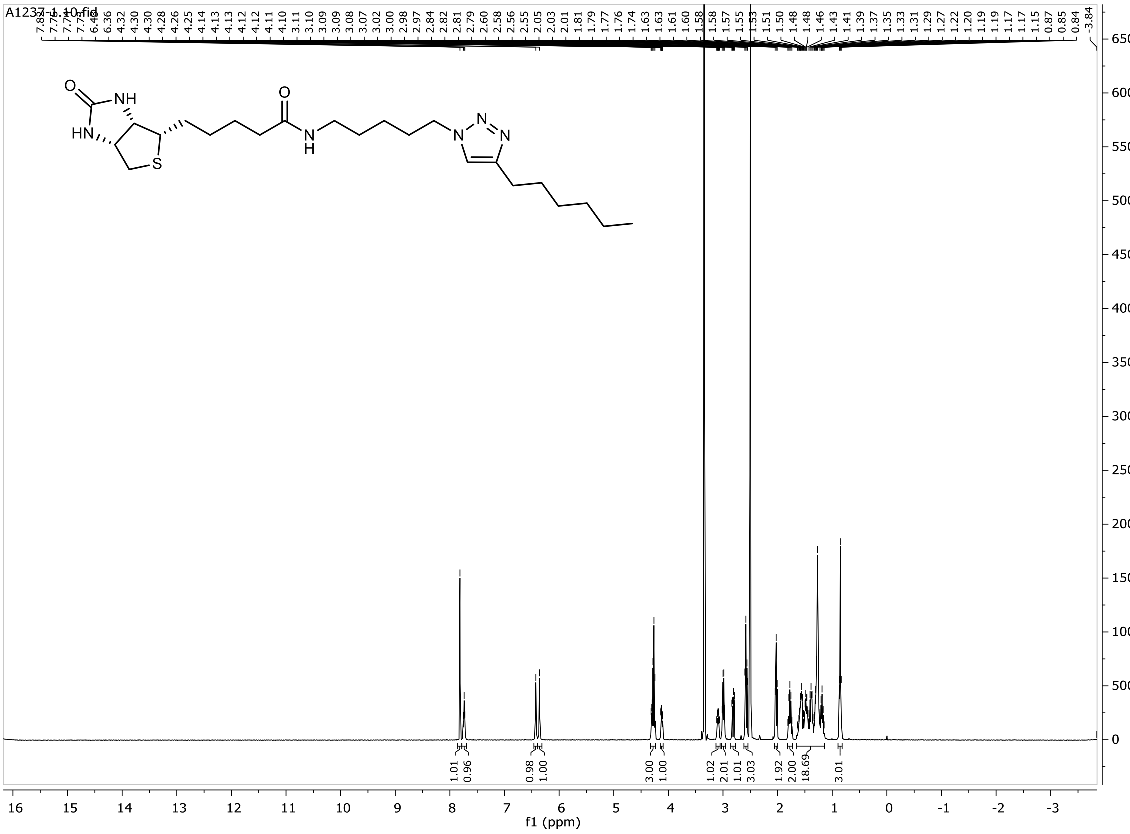
A1237-1.10.fid
6.40
6.36
4.32
4.30
4.30
4.28
4.26
4.25
4.14
4.13
4.13
4.12
4.12
4.11
4.10
3.11
3.10
3.09
3.09
3.08
3.07
3.02
3.00
2.98
2.97
2.84
2.82
2.81
2.79
2.60
2.58
2.56
2.55
2.05
2.03
2.01
1.81
1.79
1.77
1.76
1.74
1.63
1.63
1.61
1.60
1.58
1.58
1.57
1.55
1.53
1.51
1.50
1.48
1.48
1.46
1.43
1.41
1.39
1.37
1.35
1.33
1.31
1.29
1.27
1.22
1.20
1.19
1.19
1.17
1.17
1.15
0.87
0.85
0.84
-3.84
O
NH
HN
S
O
N
H
N
N
N
1.01
0.96
0.98
1.00
3.00
1.00
1.02
2.01
1.01
3.03
1.92
2.00
18.69
3.01
f1 (ppm)

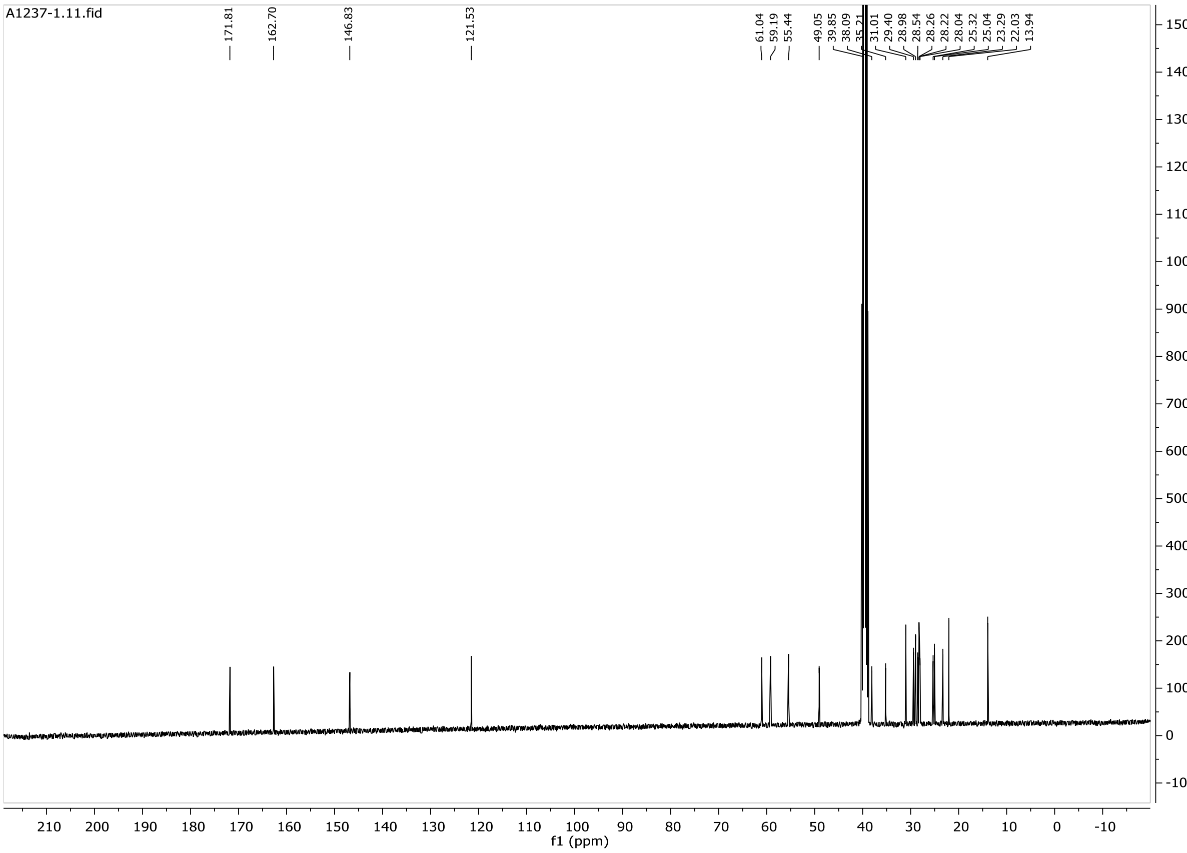
A1237-1.11.fid
171.81
162.70
146.83
121.53
61.04
59.19
55.44
49.05
39.85
38.09
35.21
31.01
29.40
28.98
28.54
28.26
28.22
28.04
25.32
25.04
23.29
22.03
13.94
f1 (ppm)

# B3

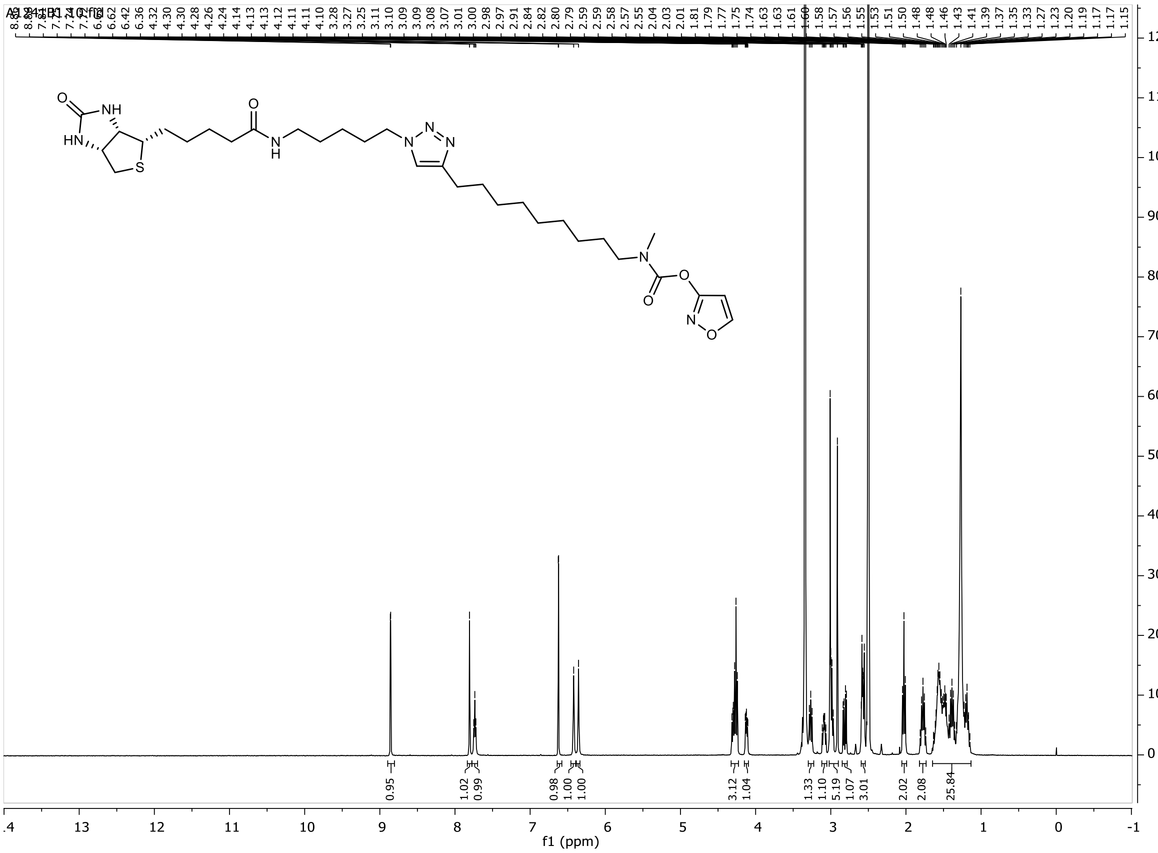
A1241B1.10.fid
6.62
6.42
6.36
4.32
4.30
4.28
4.26
4.24
4.14
4.13
4.13
4.12
4.11
4.11
4.10
3.28
3.27
3.25
3.11
3.10
3.09
3.09
3.08
3.07
3.01
3.00
2.98
2.97
2.91
2.84
2.82
2.80
2.79
2.59
2.59
2.58
2.57
2.55
2.04
2.03
2.01
1.81
1.79
1.77
1.75
1.74
1.63
1.63
1.61
1.60
1.58
1.57
1.56
1.55
1.53
1.51
1.50
1.48
1.48
1.46
1.43
1.41
1.39
1.37
1.35
1.33
1.27
1.23
1.20
1.19
1.17
1.15
0.95
1.02
0.99
0.98
1.00
1.00
3.12
1.04
1.33
1.10
5.19
1.07
3.01
2.02
2.08
25.84
f1 (ppm)

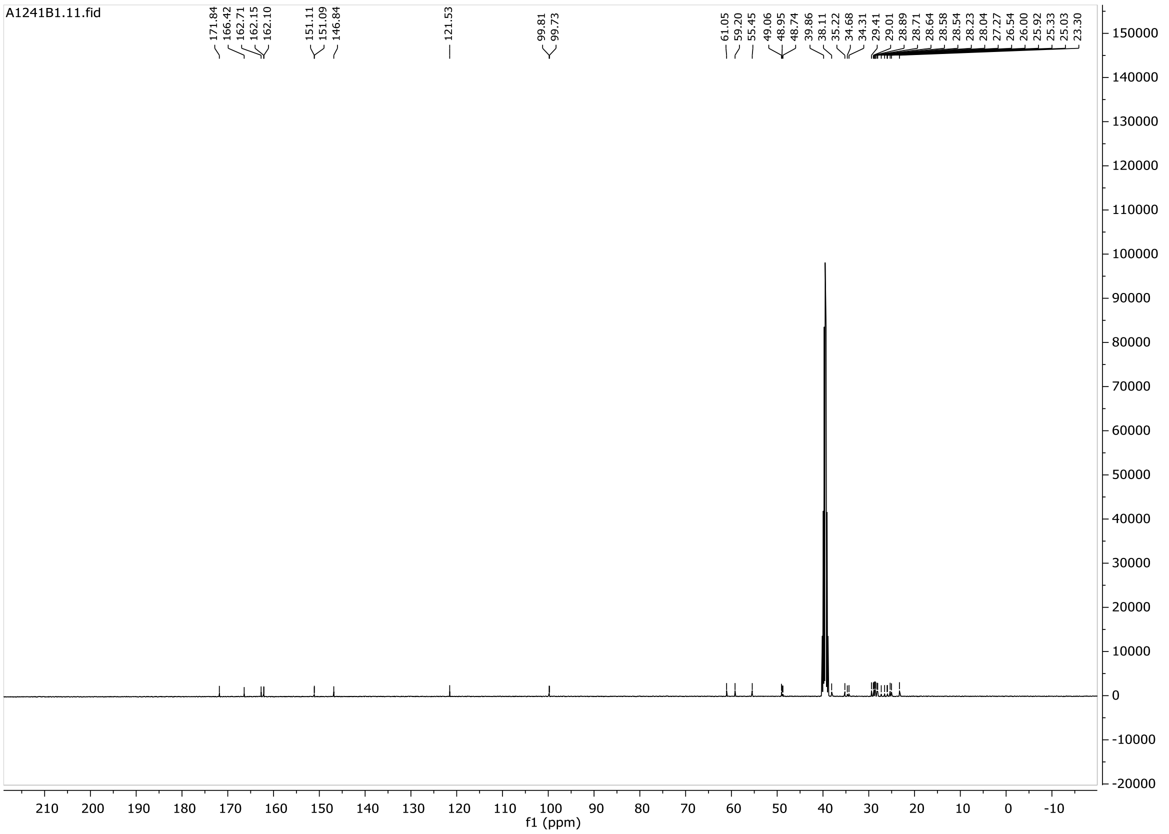
A1241B1.11.fid
171.84
166.42
162.71
162.15
162.10
151.11
151.09
146.84
121.53
99.81
99.73
61.05
59.20
55.45
49.06
48.95
48.74
39.86
38.11
35.22
34.68
34.31
29.41
29.01
28.89
28.71
28.64
28.58
28.54
28.23
28.04
27.27
26.54
26.00
25.92
25.33
25.03
23.30
f1 (ppm)

### UPLC traces

**2**

#### Chromatogram and Results

##### Injection Details

| | | | |
|---|---|---|---|
| *Injection Name:* | **A686-2** | *Run Time (min):* | **8,00** |
| *Vial Number:* | **BD6** | *Injection Volume:* | **2,00** |
| *Injection Type:* | **Unknown** | *Channel:* | **UV_VIS_2** |
| *Calibration Level:* | | *Wavelength:* | **254** |
| *Instrument Method:* | **General method (0.1TFA_MeCN_20-90)_FKKT** | *Bandwidth:* | **16** |
| *Processing Method:* | **Quantitative** | *Dilution Factor:* | **1,0000** |
| *Injection Date/Time:* | **17.jun..20 11:29** | *Sample Weight:* | **1,0000** |

##### Chromatogram

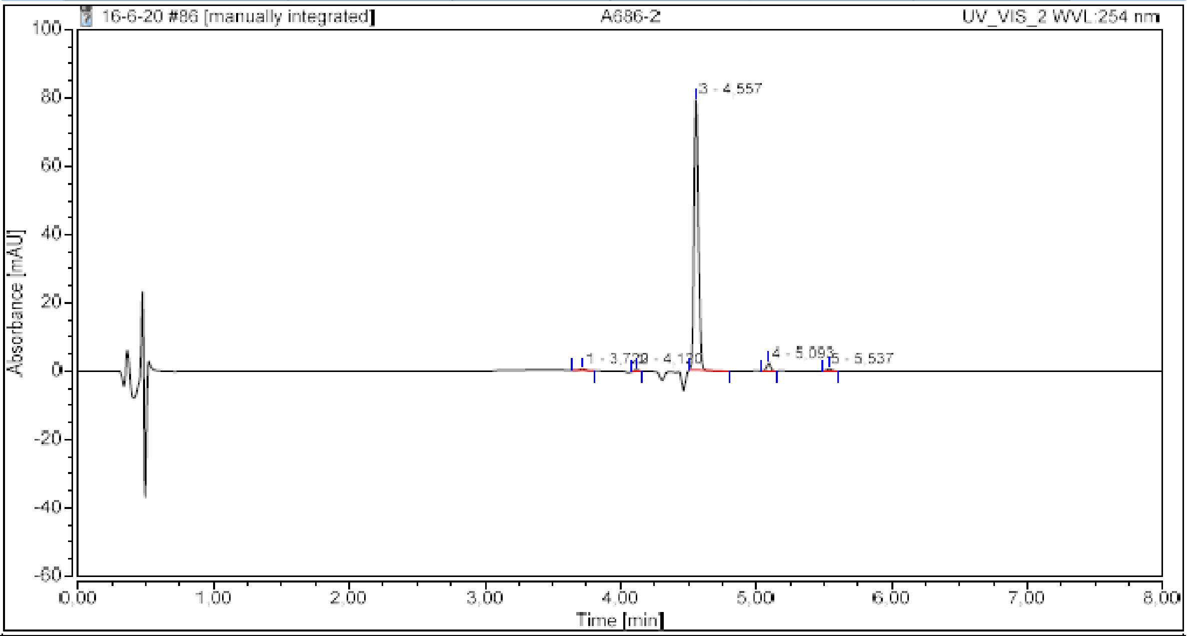

##### Integration Results

| No. | Peak Name | Retention Time min | Area mAU*min | Height mAU | Relative Area % | Relative Height % | Amount n.a. |
|---|---|---|---|---|---|---|---|
| 1 | | 3,720 | 0,018 | 0,403 | 0,59 | 0,49 | n.a. |
| 2 | | 4,120 | 0,020 | 0,515 | 0,66 | 0,62 | n.a. |
| 3 | | 4,557 | 2,931 | 79,002 | 95,47 | 95,24 | n.a. |
| 4 | | 5,093 | 0,076 | 2,341 | 2,47 | 2,82 | n.a. |
| 5 | | 5,537 | 0,025 | 0,692 | 0,81 | 0,83 | n.a. |
| **Total:** | | | **3,070** | **82,953** | **100,00** | **100,00** | |

3

#### Chromatogram and Results

##### Injection Details

| | | | |
|---|---|---|---|
| *Injection Name:* | **A900-1 fr10** | *Run Time (min):* | **8,00** |
| *Vial Number:* | **BA6** | *Injection Volume:* | **5,00** |
| *Injection Type:* | **Unknown** | *Channel:* | **UV_VIS_1** |
| *Calibration Level:* | | *Wavelength:* | **220** |
| *Instrument Method:* | **General method (0.1TFA_MeCN_20-90)_FKKT** | *Bandwidth:* | **8** |
| *Processing Method:* | **Quantitative** | *Dilution Factor:* | **1,0000** |
| *Injection Date/Time:* | **13.jan..21 10:15** | *Sample Weight:* | **1,0000** |

##### Chromatogram

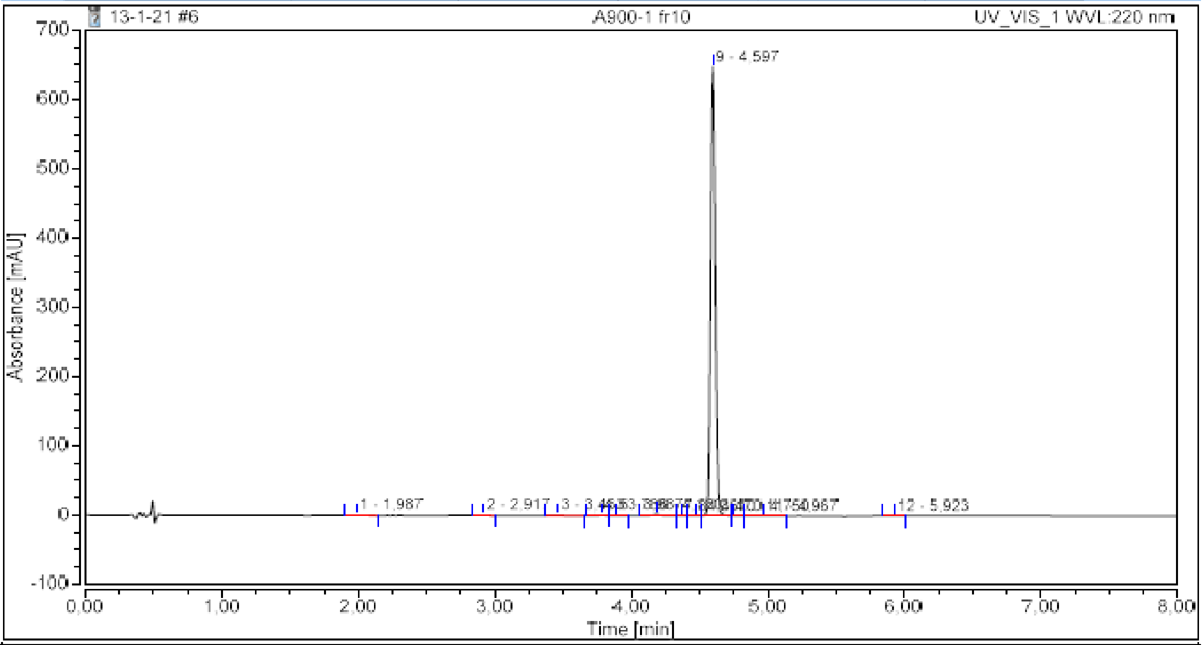

##### Integration Results

| No. | Peak Name | Retention Time min | Area mAU*min | Height mAU | Relative Area % | Relative Height % | Amount n.a. |
|---|---|---|---|---|---|---|---|
| 1 | | 1,987 | 0,018 | 0,167 | 0,07 | 0,03 | n.a. |
| 2 | | 2,917 | 0,017 | 0,198 | 0,06 | 0,03 | n.a. |
| 3 | | 3,463 | 0,032 | 0,277 | 0,12 | 0,04 | n.a. |
| 4 | | 3,783 | 0,048 | 0,578 | 0,18 | 0,09 | n.a. |
| 5 | | 3,887 | 0,041 | 0,436 | 0,15 | 0,07 | n.a. |
| 6 | | 4,180 | 0,143 | 2,071 | 0,52 | 0,32 | n.a. |
| 7 | | 4,367 | 0,010 | 0,258 | 0,04 | 0,04 | n.a. |
| 8 | | 4,470 | 0,022 | 0,293 | 0,08 | 0,04 | n.a. |
| 9 | | 4,597 | 27,060 | 648,451 | 98,56 | 99,21 | n.a. |
| 10 | | 4,750 | 0,019 | 0,390 | 0,07 | 0,06 | n.a. |
| 11 | | 4,967 | 0,038 | 0,423 | 0,14 | 0,06 | n.a. |
| 12 | | 5,923 | 0,006 | 0,086 | 0,02 | 0,01 | n.a. |
| **Total:** | | | **27,455** | **653,628** | **100,00** | **100,00** | |

**4**

#### Chromatogram and Results

| Injection Details | | | |
|---|---|---|---|
| *Injection Name:* | **A1024-2 fr3** | *Run Time (min):* | **8,00** |
| *Vial Number:* | **GB3** | *Injection Volume:* | **5,00** |
| *Injection Type:* | **Unknown** | *Channel:* | **UV_VIS_2** |
| *Calibration Level:* | | *Wavelength:* | **254** |
| *Instrument Method:* | **General method (0.1TFA_MeCN_20-90)_FKKT** | *Bandwidth:* | **16** |
| *Processing Method:* | **Quantitative** | *Dilution Factor:* | **1,0000** |
| *Injection Date/Time:* | **04.jun..21 09:39** | *Sample Weight:* | **1,0000** |

**Chromatogram**

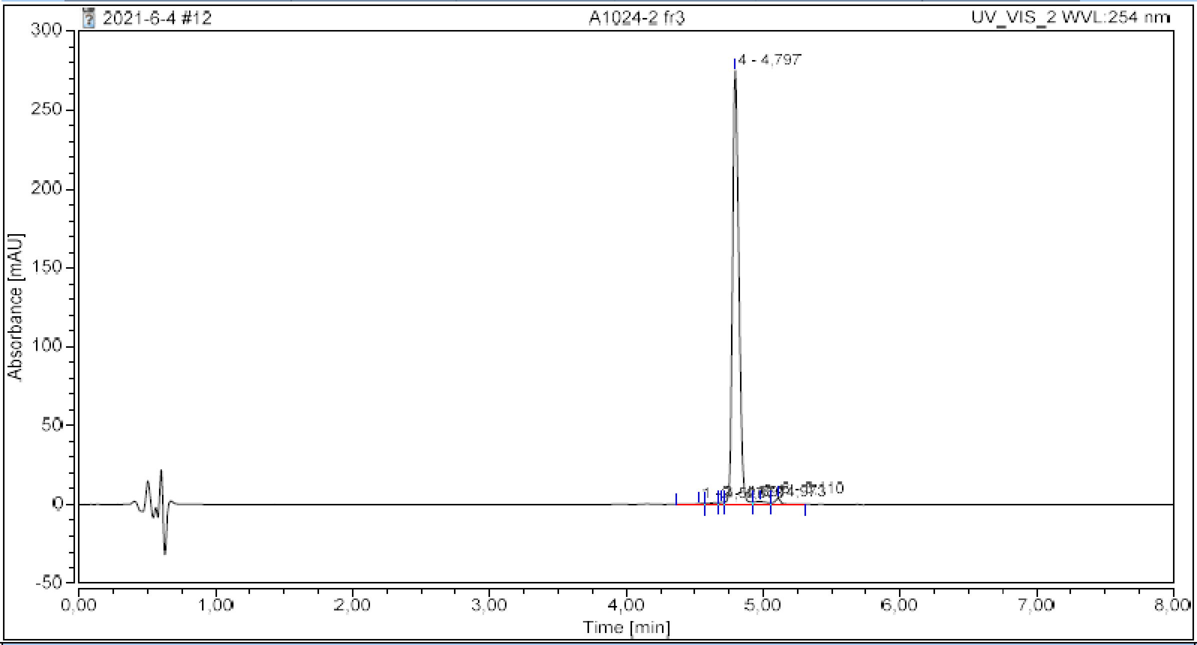

**Integration Results**

| No. | Peak Name | Retention Time<br>min | Area<br>mAU*min | Height<br>mAU | Relative Area<br>% | Relative Height<br>% | Amount<br>n.a. |
|---|---|---|---|---|---|---|---|
| 1 | | 4,527 | 0,043 | 0,464 | 0,28 | 0,16 | n.a. |
| 2 | | 4,670 | 0,085 | 1,062 | 0,55 | 0,37 | n.a. |
| 3 | | 4,697 | 0,053 | 1,261 | 0,34 | 0,44 | n.a. |
| 4 | | 4,797 | 14,742 | 275,212 | 96,14 | 96,97 | n.a. |
| 5 | | 4,973 | 0,206 | 1,848 | 1,34 | 0,65 | n.a. |
| 6 | | 5,110 | 0,205 | 3,952 | 1,34 | 1,39 | n.a. |
| **Total:** | | | **15,333** | **283,799** | **100,00** | **100,00** | |

5

#### Chromatogram and Results

##### Injection Details

| | | | |
|---|---|---|---|
| *Injection Name:* | **A748C** | *Run Time (min):* | **8,00** |
| *Vial Number:* | **BA4** | *Injection Volume:* | **5,00** |
| *Injection Type:* | **Unknown** | *Channel:* | **UV_VIS_2** |
| *Calibration Level:* | | *Wavelength:* | **254** |
| *Instrument Method:* | **General method (0.1TFA_MeCN_20-90)_FKKT** | *Bandwidth:* | **16** |
| *Processing Method:* | **Quantitative** | *Dilution Factor:* | **1,0000** |
| *Injection Date/Time:* | **14.avg..20 11:15** | *Sample Weight:* | **1,0000** |

##### Chromatogram

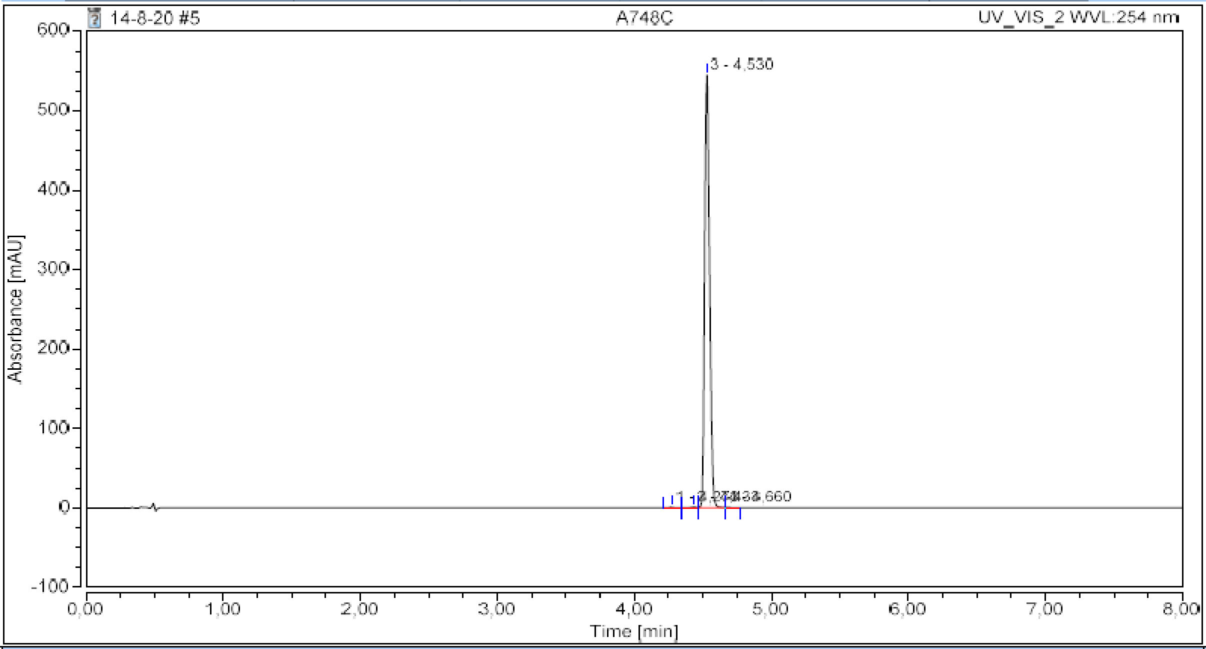

##### Integration Results

| No. | Peak Name | Retention Time min | Area mAU*min | Height mAU | Relative Area % | Relative Height % | Amount n.a. |
|---|---|---|---|---|---|---|---|
| 1 | | 4,273 | 0,031 | 0,859 | 0,13 | 0,16 | n.a. |
| 2 | | 4,433 | 0,040 | 0,879 | 0,17 | 0,16 | n.a. |
| 3 | | 4,530 | 22,873 | 544,939 | 99,59 | 99,59 | n.a. |
| 4 | | 4,660 | 0,024 | 0,498 | 0,11 | 0,09 | n.a. |
| **Total:** | | | **22,968** | **547,175** | **100,00** | **100,00** | |

6

#### Chromatogram and Results

##### Injection Details

| | | | |
|---|---|---|---|
| *Injection Name:* | **A992C fr2** | *Run Time (min):* | **8,00** |
| *Vial Number:* | **BC1** | *Injection Volume:* | **5,00** |
| *Injection Type:* | **Unknown** | *Channel:* | **UV_VIS_1** |
| *Calibration Level:* | | *Wavelength:* | **220** |
| *Instrument Method:* | **General method (0.1TFA_MeCN_20-90)_FKKT** | *Bandwidth:* | **8** |
| *Processing Method:* | **Quantitative** | *Dilution Factor:* | **1,0000** |
| *Injection Date/Time:* | **22.mar..21 15:11** | *Sample Weight:* | **1,0000** |

##### Chromatogram

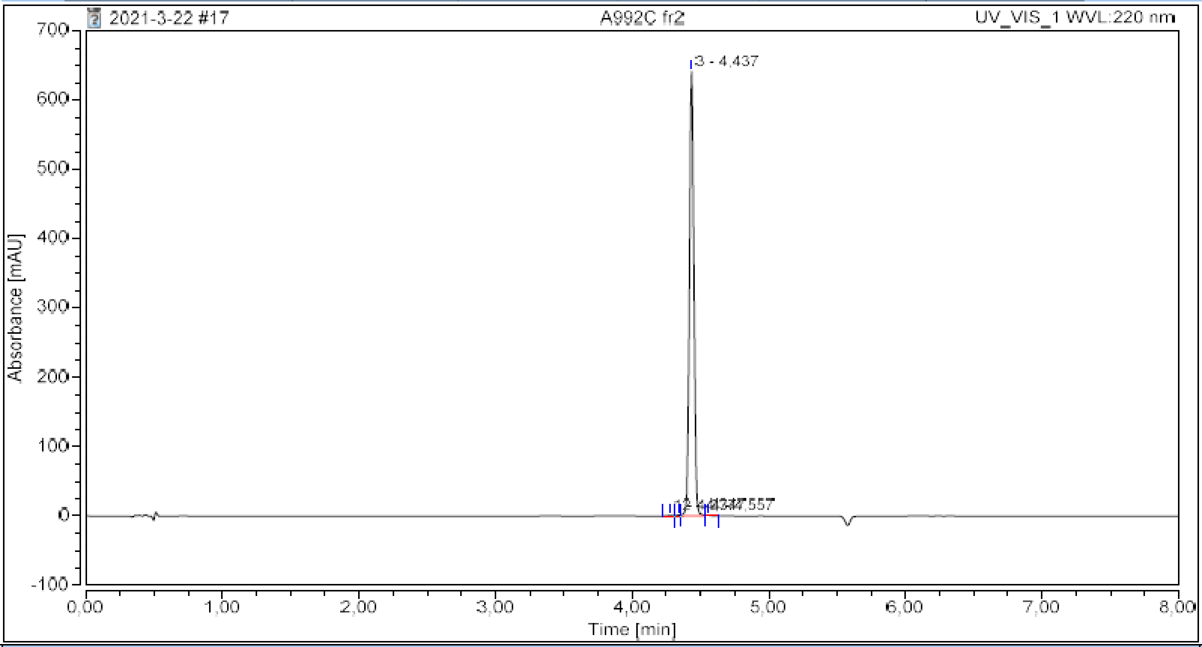

##### Integration Results

| No. | Peak Name | Retention Time<br>min | Area<br>mAU*min | Height<br>mAU | Relative Area<br>% | Relative Height<br>% | Amount<br>n.a. |
|---|---|---|---|---|---|---|---|
| 1 | | 4,277 | 0,033 | 0,677 | 0,13 | 0,10 | n.a. |
| 2 | | 4,347 | 0,054 | 1,517 | 0,22 | 0,24 | n.a. |
| 3 | | 4,437 | 24,600 | 641,457 | 99,45 | 99,50 | n.a. |
| 4 | | 4,557 | 0,050 | 1,049 | 0,20 | 0,16 | n.a. |
| **Total:** | | | **24,736** | **644,700** | **100,00** | **100,00** | |

**6b**

#### Chromatogram and Results

**Injection Details**

| | | | |
|---|---|---|---|
| *Injection Name:* | **A1250** | *Run Time (min):* | **6,00** |
| *Vial Number:* | **GC2** | *Injection Volume:* | **5,00** |
| *Injection Type:* | **Unknown** | *Channel:* | **UV_VIS_1** |
| *Calibration Level:* | | *Wavelength:* | **220** |
| *Instrument Method:* | **Kinetics method 8 per h 0.4 mL** | *Bandwidth:* | **8** |
| *Processing Method:* | **Quantitative** | *Dilution Factor:* | **1,0000** |
| *Injection Date/Time:* | **07.mar..22 11:59** | *Sample Weight:* | **1,0000** |

**Chromatogram**

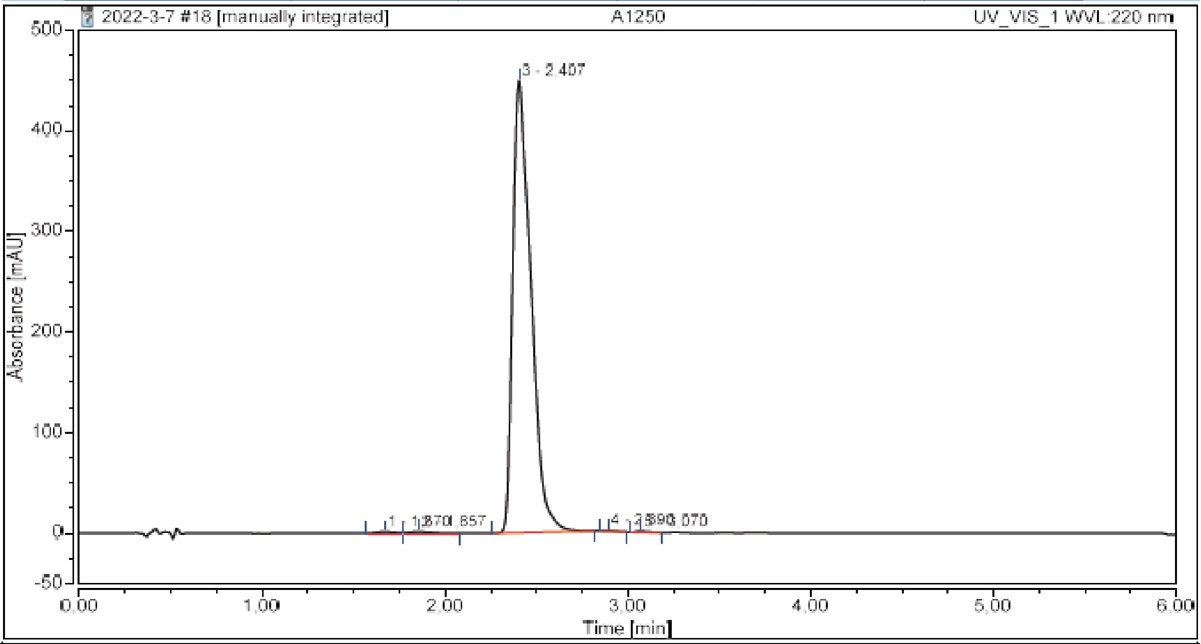

**Integration Results**

| No. | Peak Name | Retention Time min | Area mAU*min | Height mAU | Relative Area % | Relative Height % | Amount n.a. |
|---|---|---|---|---|---|---|---|
| 1 | | 1,670 | 0,157 | 1,435 | 0,30 | 0,32 | n.a. |
| 2 | | 1,857 | 0,191 | 1,520 | 0,37 | 0,33 | n.a. |
| 3 | | 2,407 | 51,862 | 449,274 | 99,14 | 99,02 | n.a. |
| 4 | | 2,890 | 0,034 | 0,548 | 0,07 | 0,12 | n.a. |
| 5 | | 3,070 | 0,067 | 0,959 | 0,13 | 0,21 | n.a. |
| **Total:** | | | **52,311** | **453,735** | **100,00** | **100,00** | |

**6c**

#### Chromatogram and Results

**Injection Details**

| | | | |
|---|---|---|---|
| *Injection Name:* | **A1244** | *Run Time (min):* | **6,00** |
| *Vial Number:* | **GD5** | *Injection Volume:* | **5,00** |
| *Injection Type:* | **Unknown** | *Channel:* | **UV_VIS_1** |
| *Calibration Level:* | | *Wavelength:* | **220** |
| *Instrument Method:* | **Kinetics method 8 per h 0.4 mL** | *Bandwidth:* | **8** |
| *Processing Method:* | **Quantitative** | *Dilution Factor:* | **1,0000** |
| *Injection Date/Time:* | **07.nov..22 16:16** | *Sample Weight:* | **1,0000** |

**Chromatogram**

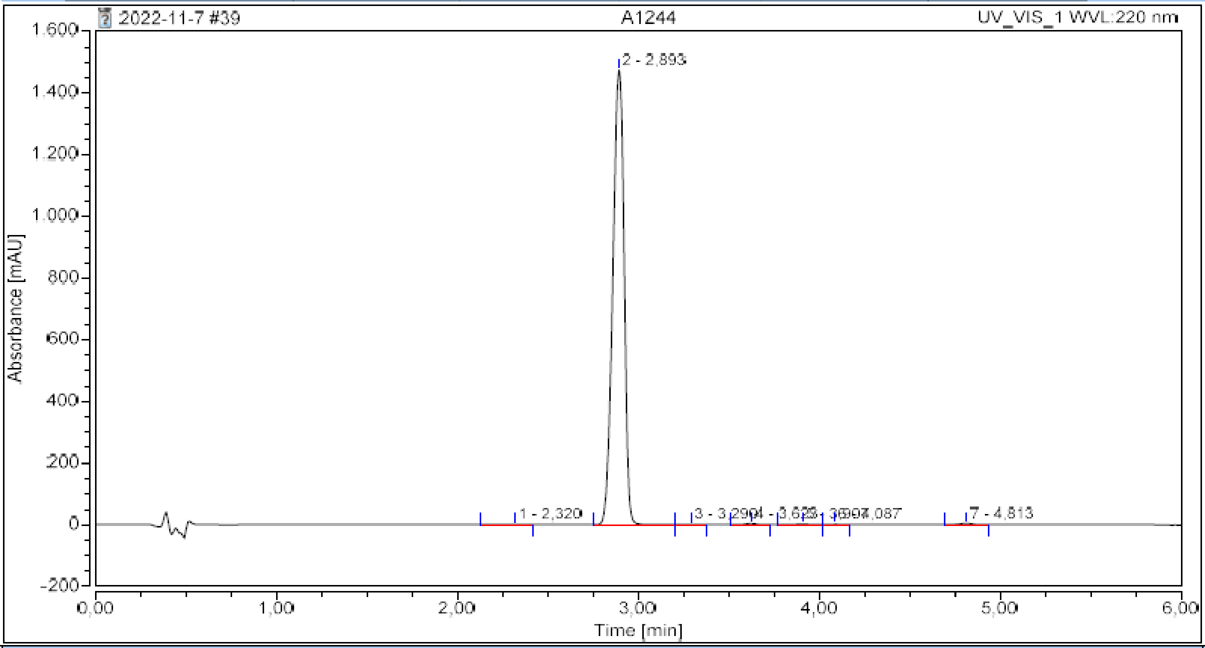

**Integration Results**

| No. | Peak Name | Retention Time min | Area mAU*min | Height mAU | Relative Area % | Relative Height % | Amount n.a. |
|---|---|---|---|---|---|---|---|
| 1 | | 2,320 | 0,077 | 0,487 | 0,07 | 0,03 | n.a. |
| 2 | | 2,893 | 106,985 | 1474,187 | 99,23 | 99,30 | n.a. |
| 3 | | 3,290 | 0,039 | 0,490 | 0,04 | 0,03 | n.a. |
| 4 | | 3,623 | 0,216 | 3,346 | 0,20 | 0,23 | n.a. |
| 5 | | 3,907 | 0,132 | 1,518 | 0,12 | 0,10 | n.a. |
| 6 | | 4,087 | 0,072 | 1,080 | 0,07 | 0,07 | n.a. |
| 7 | | 4,813 | 0,292 | 3,475 | 0,27 | 0,23 | n.a. |
| **Total:** | | | **107,813** | **1484,584** | **100,00** | **100,00** | |

**6d**

#### Chromatogram and Results

**Injection Details**

| | | | |
|---|---|---|---|
| *Injection Name:* | **A1274-1** | *Run Time (min):* | **6,00** |
| *Vial Number:* | **GD3** | *Injection Volume:* | **5,00** |
| *Injection Type:* | **Unknown** | *Channel:* | **UV_VIS_2** |
| *Calibration Level:* | | *Wavelength:* | **254** |
| *Instrument Method:* | **Kinetics method 8 per h 0.4 mL** | *Bandwidth:* | **16** |
| *Processing Method:* | **Quantitative** | *Dilution Factor:* | **1,0000** |
| *Injection Date/Time:* | **07.nov..22 16:01** | *Sample Weight:* | **1,0000** |

**Chromatogram**

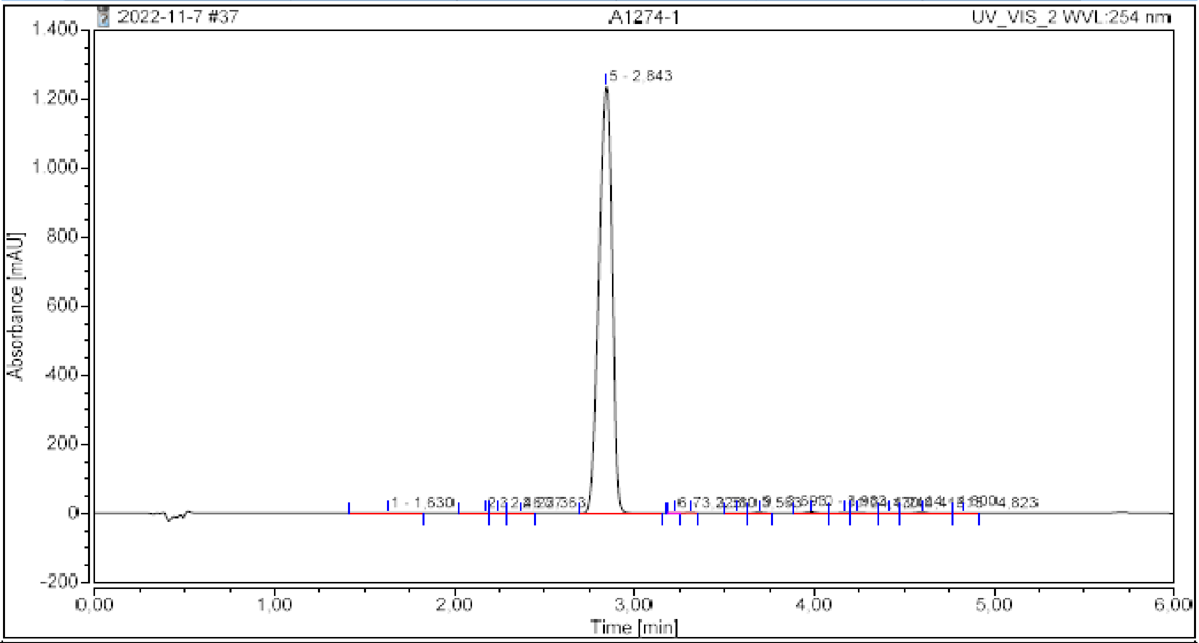

**Integration Results**

| No. | Peak Name | Retention Time min | Area mAU*min | Height mAU | Relative Area % | Relative Height % | Amount n.a. |
|---|---|---|---|---|---|---|---|
| 1 | | 1,630 | 0,117 | 0,849 | 0,11 | 0,07 | n.a. |
| 2 | | 2,167 | 0,057 | 0,823 | 0,05 | 0,07 | n.a. |
| 3 | | 2,237 | 0,093 | 1,428 | 0,09 | 0,11 | n.a. |
| 4 | | 2,363 | 0,058 | 0,648 | 0,05 | 0,05 | n.a. |
| 5 | | 2,843 | 105,491 | 1237,509 | 98,38 | 98,20 | n.a. |
| 6 | | 3,223 | 0,005 | 0,149 | 0,00 | 0,01 | n.a. |
| 7 | | 3,310 | 0,105 | 1,796 | 0,10 | 0,14 | n.a. |
| 8 | | 3,563 | 0,051 | 0,834 | 0,05 | 0,07 | n.a. |
| 9 | | 3,693 | 0,186 | 2,952 | 0,17 | 0,23 | n.a. |
| 10 | | 3,983 | 0,281 | 4,004 | 0,26 | 0,32 | n.a. |
| 11 | | 4,170 | 0,133 | 1,958 | 0,12 | 0,16 | n.a. |
| 12 | | 4,240 | 0,216 | 2,250 | 0,20 | 0,18 | n.a. |
| 13 | | 4,413 | 0,082 | 0,954 | 0,08 | 0,08 | n.a. |
| 14 | | 4,600 | 0,336 | 3,698 | 0,31 | 0,29 | n.a. |
| 15 | | 4,823 | 0,023 | 0,366 | 0,02 | 0,03 | n.a. |
| **Total:** | | | **107,231** | **1260,219** | **100,00** | **100,00** | |

7

#### Chromatogram and Results

**Injection Details**

| | | | |
|---|---|---|---|
| *Injection Name:* | **A878A1 fr5** | *Run Time (min):* | **8,00** |
| *Vial Number:* | **BB7** | *Injection Volume:* | **5,00** |
| *Injection Type:* | **Unknown** | *Channel:* | **UV_VIS_2** |
| *Calibration Level:* | | *Wavelength:* | **254** |
| *Instrument Method:* | **General method (0.1TFA_MeCN_20-90)_FKKT** | *Bandwidth:* | **16** |
| *Processing Method:* | **Quantitative** | *Dilution Factor:* | **1,0000** |
| *Injection Date/Time:* | **22.dec..20 11:30** | *Sample Weight:* | **1,0000** |

**Chromatogram**

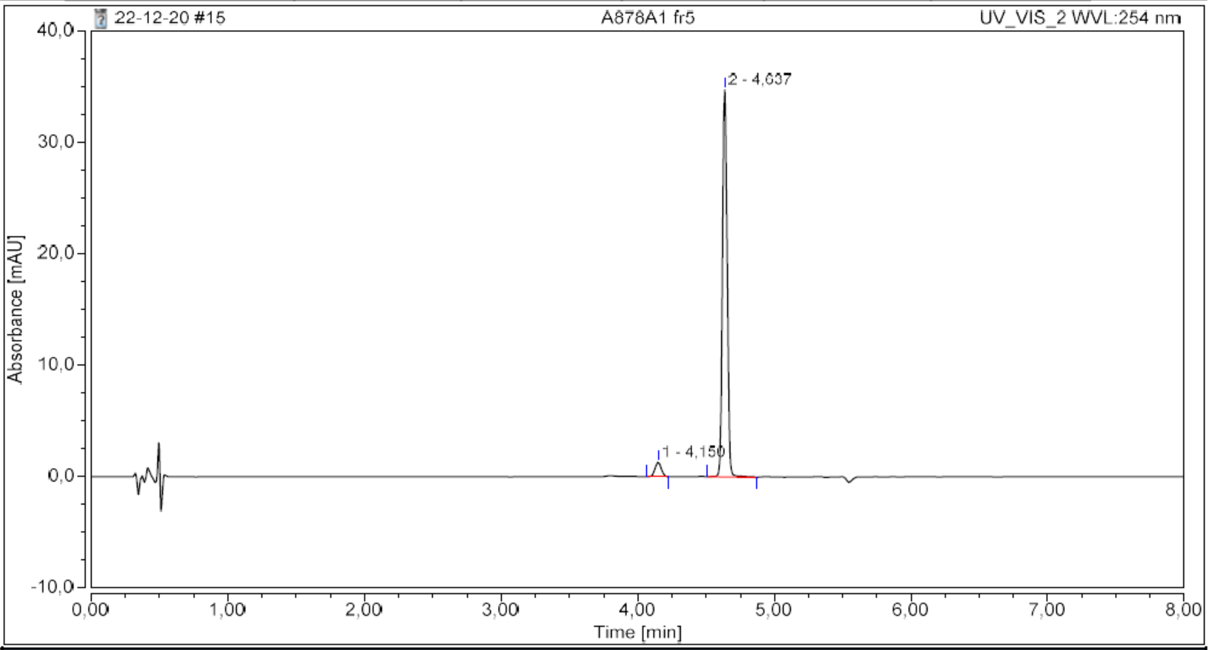

**Integration Results**

| No. | Peak Name | Retention Time min | Area mAU*min | Height mAU | Relative Area % | Relative Height % | Amount n.a. |
|---|---|---|---|---|---|---|---|
| 1 | | 4,150 | 0,069 | 1,306 | 4,68 | 3,62 | n.a. |
| 2 | | 4,637 | 1,409 | 34,761 | 95,32 | 96,38 | n.a. |
| **Total:** | | | **1,478** | **36,067** | **100,00** | **100,00** | |

8

### Chromatogram and Results

#### Injection Details

| | | | |
|---|---|---|---|
| *Injection Name:* | **A992A fr2** | *Run Time (min):* | **8,00** |
| *Vial Number:* | **BA7** | *Injection Volume:* | **5,00** |
| *Injection Type:* | **Unknown** | *Channel:* | **UV_VIS_1** |
| *Calibration Level:* | | *Wavelength:* | **220** |
| *Instrument Method:* | **General method (0.1TFA_MeCN_20-90)_FKKT** | *Bandwidth:* | **8** |
| *Processing Method:* | **Quantitative** | *Dilution Factor:* | **1,0000** |
| *Injection Date/Time:* | **22.mar..21 13:17** | *Sample Weight:* | **1,0000** |

#### Chromatogram

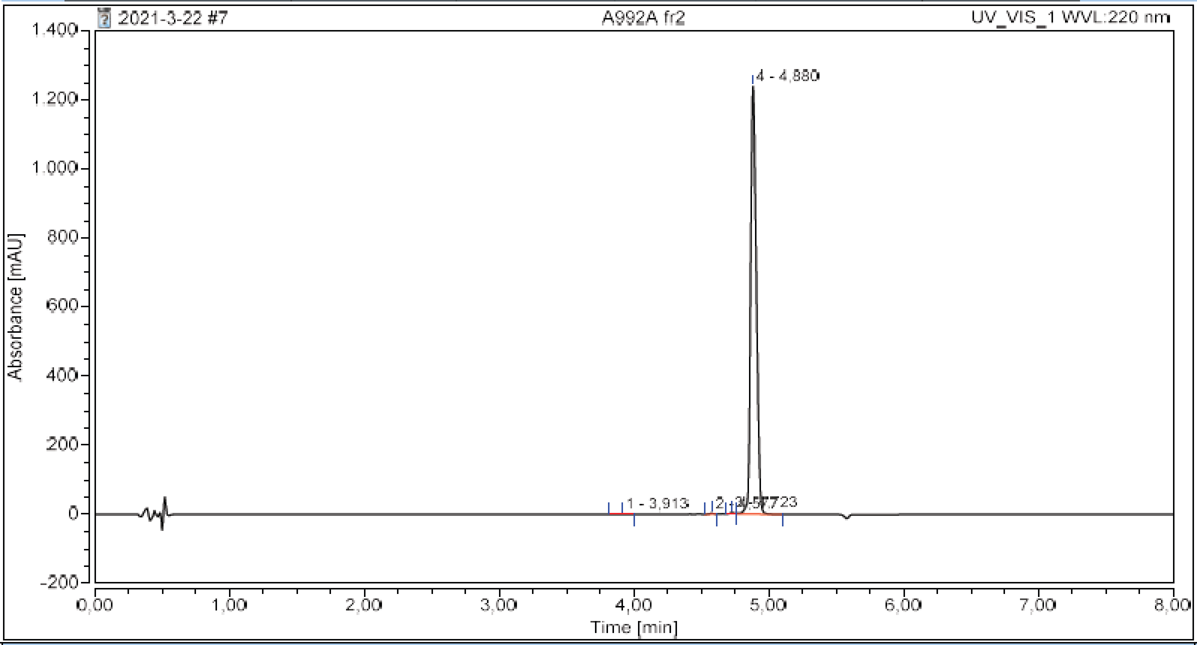

#### Integration Results

| No. | Peak Name | Retention Time min | Area mAU*min | Height mAU | Relative Area % | Relative Height % | Amount n.a. |
|---|---|---|---|---|---|---|---|
| 1 | | 3,913 | 0,025 | 0,359 | 0,04 | 0,03 | n.a. |
| 2 | | 4,577 | 0,089 | 2,597 | 0,15 | 0,21 | n.a. |
| 3 | | 4,723 | 0,168 | 4,027 | 0,28 | 0,32 | n.a. |
| 4 | | 4,880 | 60,530 | 1240,009 | 99,54 | 99,44 | n.a. |
| **Total:** | | | **60,812** | **1246,992** | **100,00** | **100,00** | |

**8b**

#### Chromatogram and Results

**Injection Details**

| | | | |
|---|---|---|---|
| *Injection Name:* | **A1025** | *Run Time (min):* | **6,00** |
| *Vial Number:* | **RE2** | *Injection Volume:* | **5,00** |
| *Injection Type:* | **Unknown** | *Channel:* | **UV_VIS_2** |
| *Calibration Level:* | | *Wavelength:* | **254** |
| *Instrument Method:* | **Kinetics method 8 per h (20-100)** | *Bandwidth:* | **16** |
| *Processing Method:* | **Quantitative** | *Dilution Factor:* | **1,0000** |
| *Injection Date/Time:* | **17.jun..21 12:34** | *Sample Weight:* | **1,0000** |

**Chromatogram**

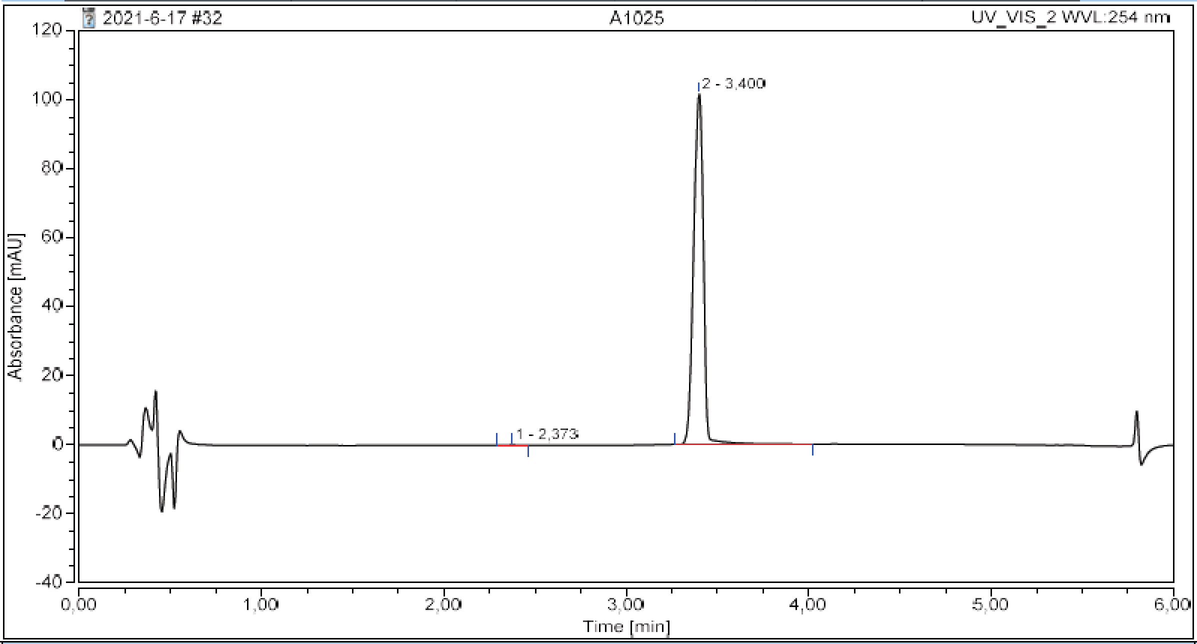

**Integration Results**

| No. | Peak Name | Retention Time min | Area mAU*min | Height mAU | Relative Area % | Relative Height % | Amount n.a. |
|---|---|---|---|---|---|---|---|
| 1 | | 2,373 | 0,007 | 0,115 | 0,11 | 0,11 | n.a. |
| 2 | | 3,400 | 6,314 | 101,606 | 99,89 | 99,89 | n.a. |
| **Total:** | | | **6,321** | **101,722** | **100,00** | **100,00** | |

**9**

#### Chromatogram and Results

##### Injection Details

| | | | |
|---|---|---|---|
| *Injection Name:* | **A1007C 2** | *Run Time (min):* | **8,00** |
| *Vial Number:* | **BA3** | *Injection Volume:* | **5,00** |
| *Injection Type:* | **Unknown** | *Channel:* | **UV_VIS_1** |
| *Calibration Level:* | | *Wavelength:* | **220** |
| *Instrument Method:* | **General method (0.1TFA_MeCN_20-90)_FKKT** | *Bandwidth:* | **8** |
| *Processing Method:* | **Quantitative** | *Dilution Factor:* | **1,0000** |
| *Injection Date/Time:* | **13.apr..21 09:41** | *Sample Weight:* | **1,0000** |

##### Chromatogram

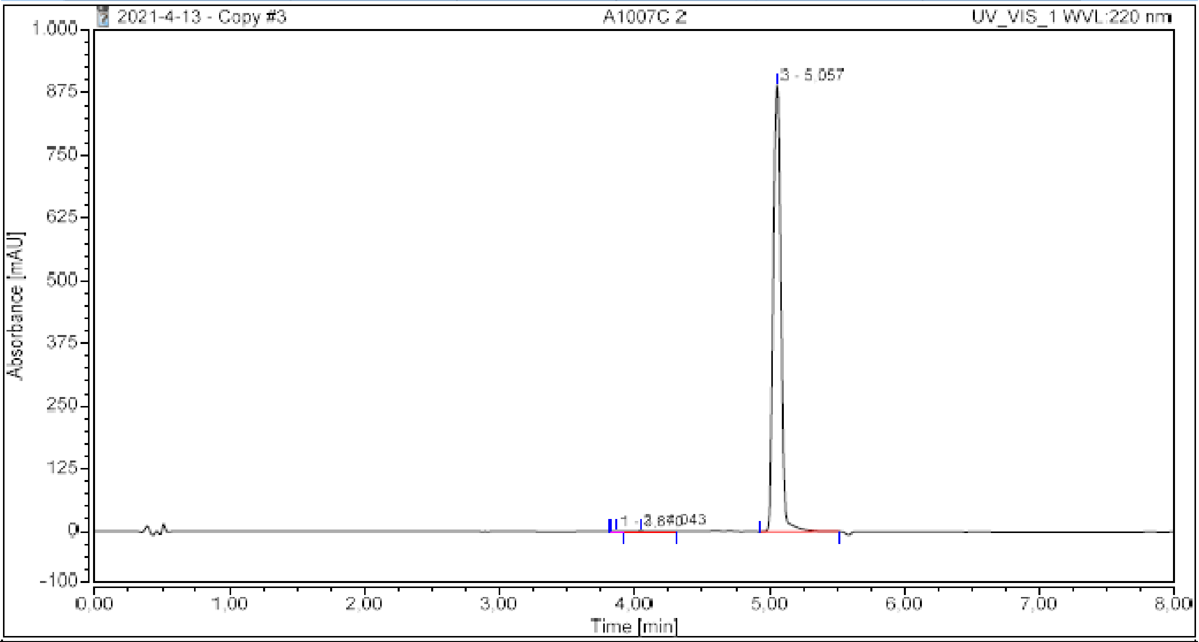

##### Integration Results

| No. | Peak Name | Retention Time min | Area mAU*min | Height mAU | Relative Area % | Relative Height % | Amount n.a. |
|---|---|---|---|---|---|---|---|
| 1 | | 3,870 | 0,005 | 0,139 | 0,01 | 0,02 | n.a. |
| 2 | | 4,043 | 0,151 | 1,263 | 0,26 | 0,14 | n.a. |
| 3 | | 5,057 | 58,855 | 890,033 | 99,74 | 99,84 | n.a. |
| **Total:** | | | **59,011** | **891,435** | **100,00** | **100,00** | |

**10**

#### Chromatogram and Results

##### Injection Details

| | | | |
|---|---|---|---|
| *Injection Name:* | **A886C1 fr3** | *Run Time (min):* | **8,00** |
| *Vial Number:* | **RC1** | *Injection Volume:* | **5,00** |
| *Injection Type:* | **Unknown** | *Channel:* | **UV_VIS_2** |
| *Calibration Level:* | | *Wavelength:* | **254** |
| *Instrument Method:* | **General method (0.1TFA_MeCN_20-90)_FKKT** | *Bandwidth:* | **16** |
| *Processing Method:* | **Quantitative** | *Dilution Factor:* | **1,0000** |
| *Injection Date/Time:* | **05.jan..21 09:08** | *Sample Weight:* | **1,0000** |

##### Chromatogram

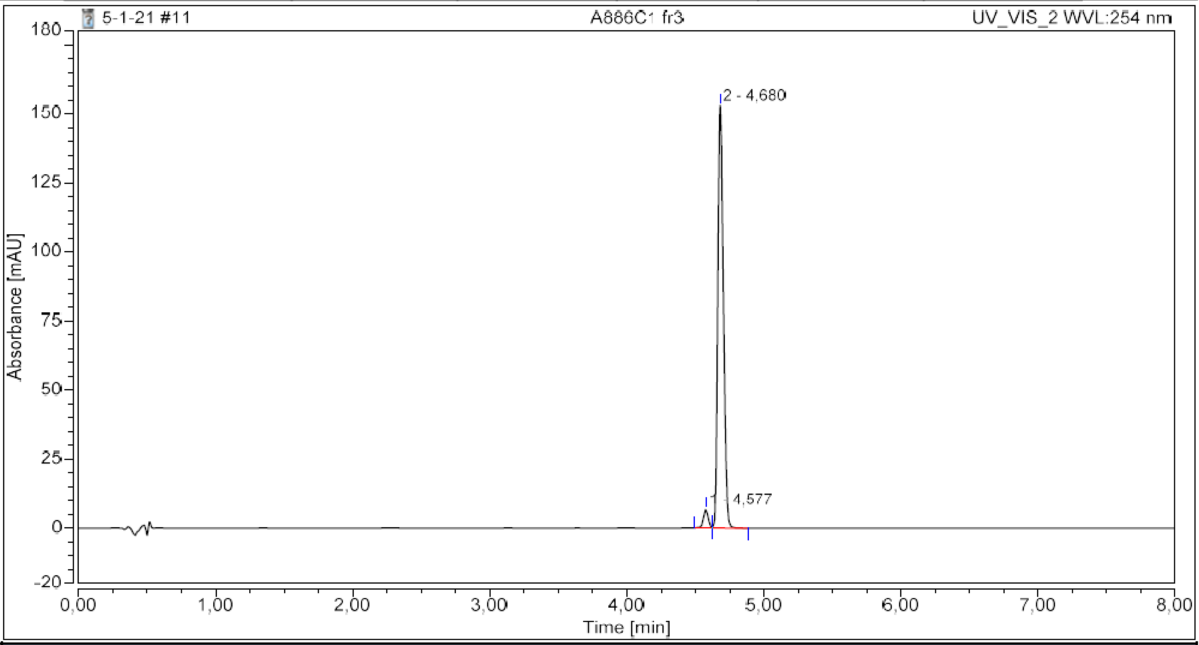

##### Integration Results

| No. | Peak Name | Retention Time min | Area mAU*min | Height mAU | Relative Area % | Relative Height % | Amount n.a. |
|---|---|---|---|---|---|---|---|
| 1 | | 4,577 | 0,275 | 6,742 | 3,67 | 4,22 | n.a. |
| 2 | | 4,680 | 7,206 | 153,044 | 96,33 | 95,78 | n.a. |
| **Total:** | | | **7,480** | **159,786** | **100,00** | **100,00** | |

**11**

#### Chromatogram and Results

##### Injection Details

| | | | |
|---|---|---|---|
| *Injection Name:* | **A920I fr5** | *Run Time (min):* | **8,00** |
| *Vial Number:* | **BA6** | *Injection Volume:* | **5,00** |
| *Injection Type:* | **Unknown** | *Channel:* | **UV_VIS_2** |
| *Calibration Level:* | | *Wavelength:* | **254** |
| *Instrument Method:* | **General method (0.1TFA_MeCN_20-90)_FKKT** | *Bandwidth:* | **16** |
| *Processing Method:* | **Quantitative** | *Dilution Factor:* | **1,0000** |
| *Injection Date/Time:* | **03.feb..21 08:28** | *Sample Weight:* | **1,0000** |

##### Chromatogram

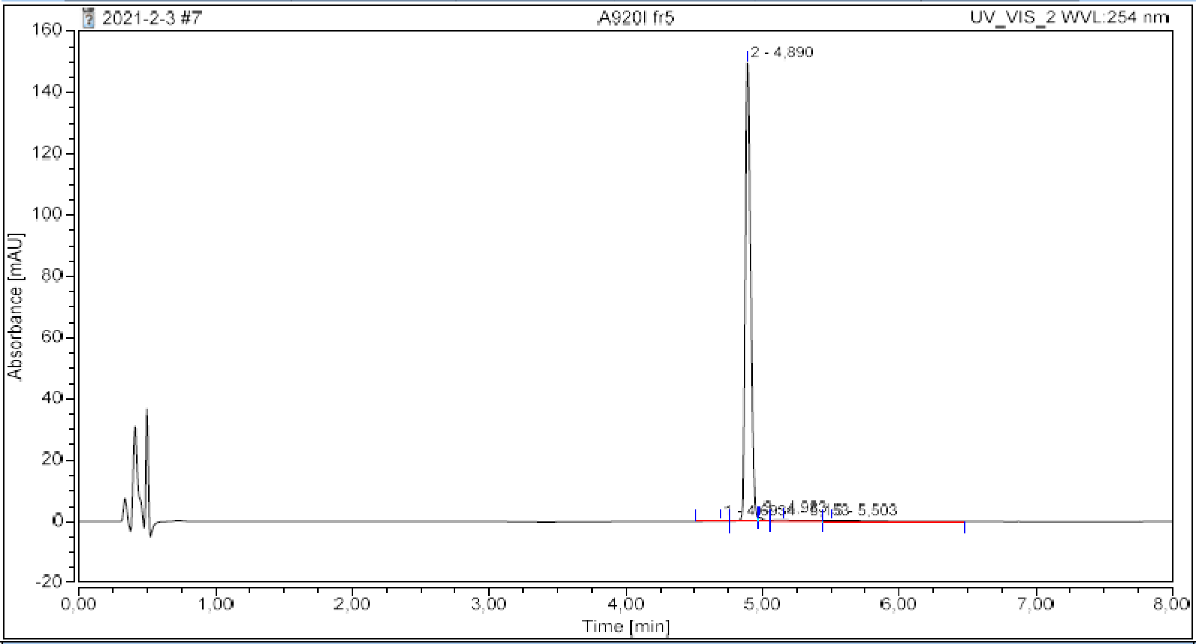

##### Integration Results

| No. | Peak Name | Retention Time min | Area mAU*min | Height mAU | Relative Area % | Relative Height % | Amount n.a. |
|---|---|---|---|---|---|---|---|
| 1 | | 4,693 | 0,021 | 0,123 | 0,30 | 0,08 | n.a. |
| 2 | | 4,890 | 6,865 | 149,612 | 95,83 | 98,66 | n.a. |
| 3 | | 4,983 | 0,056 | 1,270 | 0,78 | 0,84 | n.a. |
| 4 | | 5,153 | 0,100 | 0,366 | 1,39 | 0,24 | n.a. |
| 5 | | 5,503 | 0,122 | 0,275 | 1,71 | 0,18 | n.a. |
| **Total:** | | | **7,164** | **151,647** | **100,00** | **100,00** | |

12

#### Chromatogram and Results

**Injection Details**

| | | | |
|---|---|---|---|
| *Injection Name:* | **A950B fr5** | *Run Time (min):* | **8,00** |
| *Vial Number:* | **RA6** | *Injection Volume:* | **5,00** |
| *Injection Type:* | **Unknown** | *Channel:* | **UV_VIS_2** |
| *Calibration Level:* | | *Wavelength:* | **254** |
| *Instrument Method:* | **General method (0.1TFA_MeCN_20-90)_FKKT** | *Bandwidth:* | **16** |
| *Processing Method:* | **Quantitative** | *Dilution Factor:* | **1,0000** |
| *Injection Date/Time:* | **12.feb..21 08:08** | *Sample Weight:* | **1,0000** |

**Chromatogram**

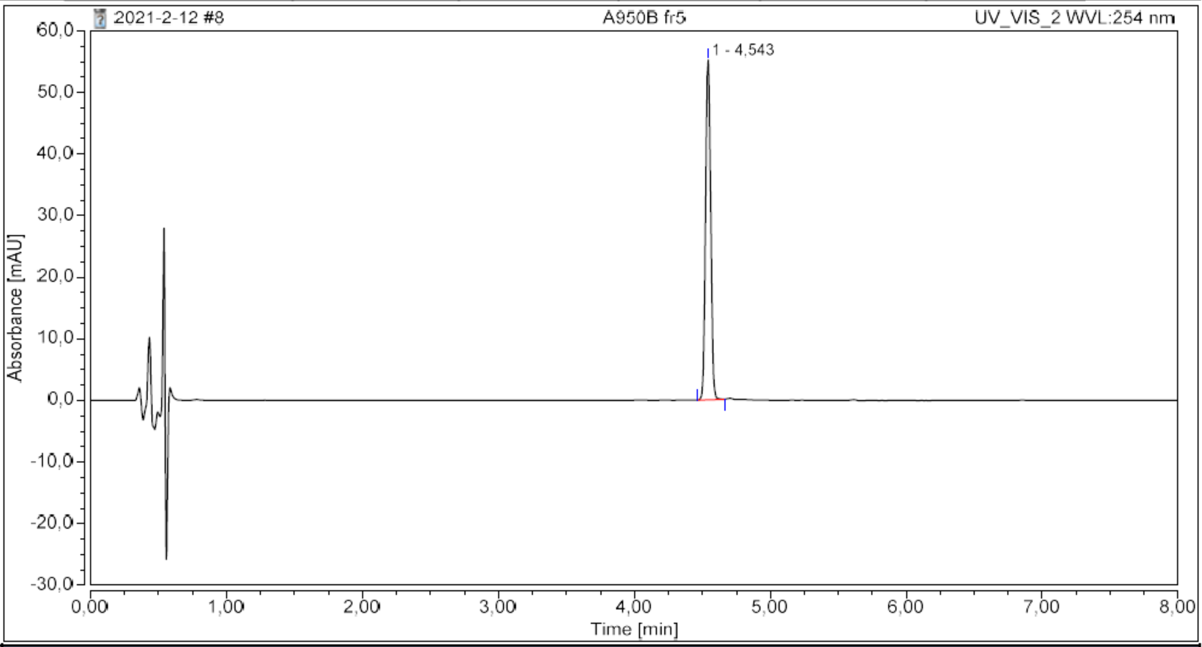

**Integration Results**

| No. | Peak Name | Retention Time min | Area mAU*min | Height mAU | Relative Area % | Relative Height % | Amount n.a. |
|---|---|---|---|---|---|---|---|
| 1 | | 4,543 | 2,425 | 55,205 | 100,00 | 100,00 | n.a. |
| **Total:** | | | **2,425** | **55,205** | **100,00** | **100,00** | |

**13**

#### Chromatogram and Results

**Injection Details**

| | | | |
|---|---|---|---|
| *Injection Name:* | **A950C fr7** | *Run Time (min):* | **8,00** |
| *Vial Number:* | **RA8** | *Injection Volume:* | **5,00** |
| *Injection Type:* | **Unknown** | *Channel:* | **UV_VIS_2** |
| *Calibration Level:* | | *Wavelength:* | **254** |
| *Instrument Method:* | **General method (0.1TFA_MeCN_20-90)_FKKT** | *Bandwidth:* | **16** |
| *Processing Method:* | **Quantitative** | *Dilution Factor:* | **1,0000** |
| *Injection Date/Time:* | **12.feb..21 08:41** | *Sample Weight:* | **1,0000** |

**Chromatogram**

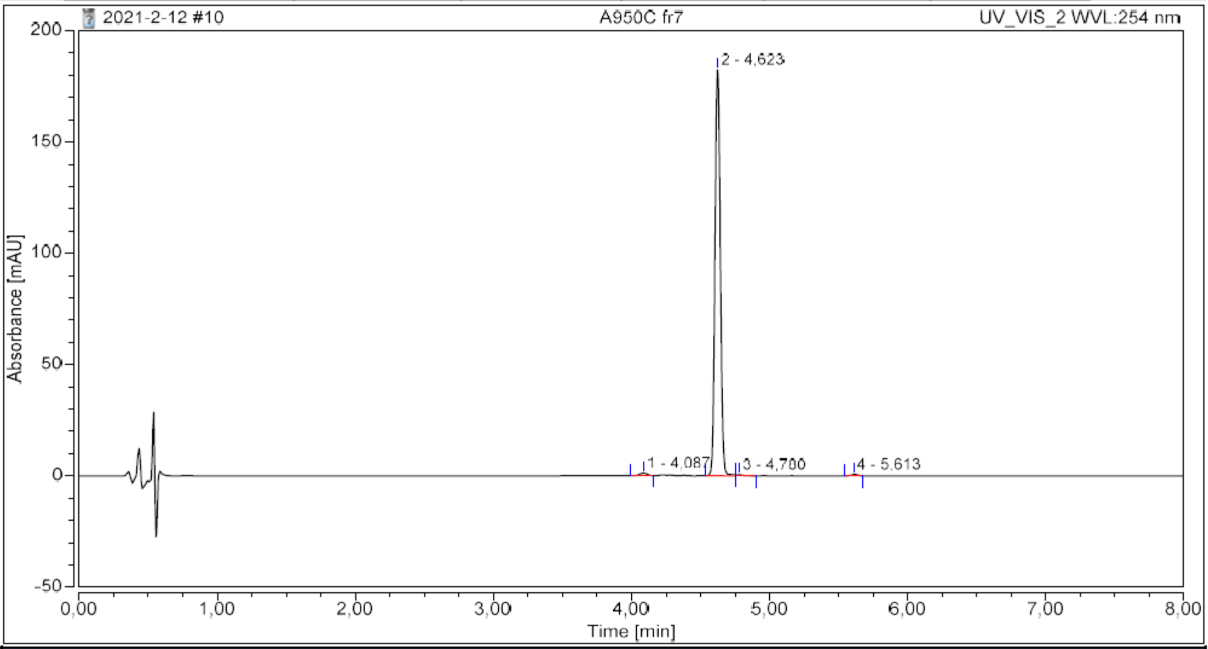

**Integration Results**

| No. | Peak Name | Retention Time min | Area mAU*min | Height mAU | Relative Area % | Relative Height % | Amount n.a. |
|---|---|---|---|---|---|---|---|
| 1 | | 4,087 | 0,078 | 1,250 | 0,94 | 0,68 | n.a. |
| 2 | | 4,623 | 8,180 | 182,469 | 98,38 | 98,71 | n.a. |
| 3 | | 4,780 | 0,021 | 0,423 | 0,26 | 0,23 | n.a. |
| 4 | | 5,613 | 0,035 | 0,707 | 0,42 | 0,38 | n.a. |
| **Total:** | | | **8,314** | **184,848** | **100,00** | **100,00** | |

14

#### Chromatogram and Results

**Injection Details**

| | | | |
|---|---|---|---|
| *Injection Name:* | **A960A fr3** | *Run Time (min):* | **8,00** |
| *Vial Number:* | **BD3** | *Injection Volume:* | **5,00** |
| *Injection Type:* | **Unknown** | *Channel:* | **UV_VIS_1** |
| *Calibration Level:* | | *Wavelength:* | **220** |
| *Instrument Method:* | **General method (0.1TFA_MeCN_20-90)_FKKT** | *Bandwidth:* | **8** |
| *Processing Method:* | **Quantitative** | *Dilution Factor:* | **1,0000** |
| *Injection Date/Time:* | **03.mar..21 09:21** | *Sample Weight:* | **1,0000** |

**Chromatogram**

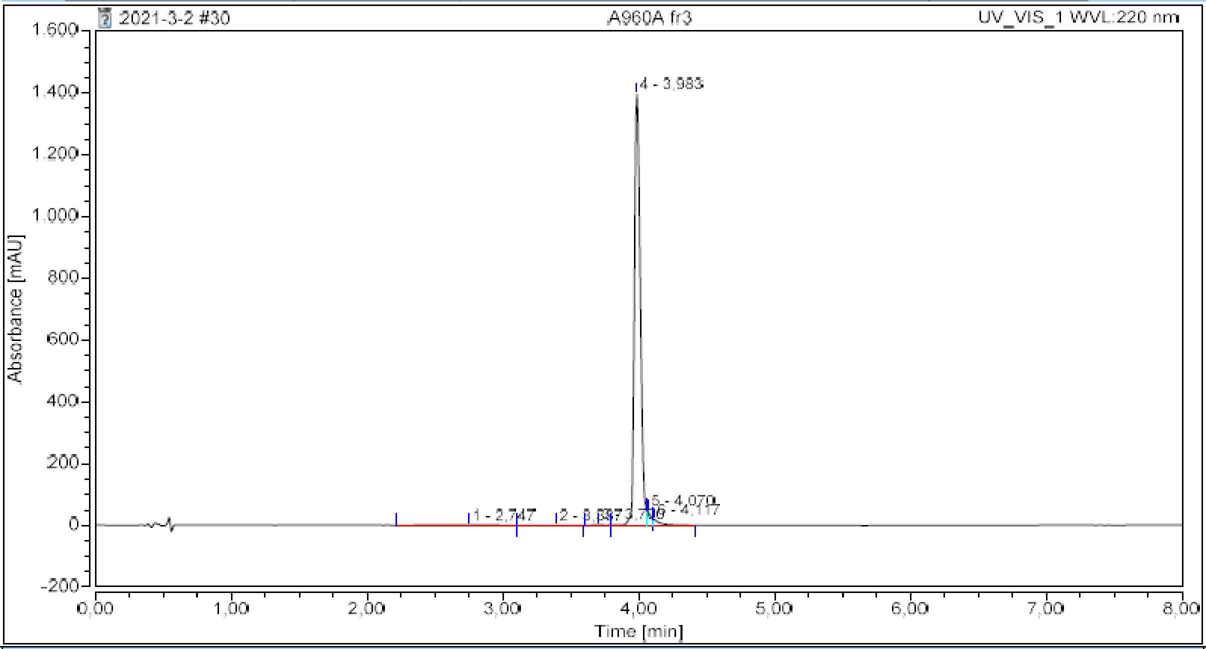

**Integration Results**

| No. | Peak Name | Retention Time min | Area mAU*min | Height mAU | Relative Area % | Relative Height % | Amount n.a. |
|---|---|---|---|---|---|---|---|
| 1 | | 2,747 | 0,211 | 0,388 | 0,28 | 0,03 | n.a. |
| 2 | | 3,387 | 0,103 | 0,352 | 0,14 | 0,02 | n.a. |
| 3 | | 3,700 | 0,107 | 1,086 | 0,14 | 0,07 | n.a. |
| 4 | | 3,983 | 71,312 | 1396,943 | 96,01 | 95,65 | n.a. |
| 5 | | 4,070 | 1,371 | 44,505 | 1,85 | 3,05 | n.a. |
| 6 | | 4,117 | 1,173 | 17,228 | 1,58 | 1,18 | n.a. |
| **Total:** | | | **74,276** | **1460,501** | **100,00** | **100,00** | |

**15**

#### Chromatogram and Results

##### Injection Details

| | | | |
|---|---|---|---|
| *Injection Name:* | **A1007A 2** | *Run Time (min):* | **8,00** |
| *Vial Number:* | **BA7** | *Injection Volume:* | **5,00** |
| *Injection Type:* | **Unknown** | *Channel:* | **UV_VIS_1** |
| *Calibration Level:* | | *Wavelength:* | **220** |
| *Instrument Method:* | **General method (0.1TFA_MeCN_20-90)_FKKT** | *Bandwidth:* | **8** |
| *Processing Method:* | **Quantitative** | *Dilution Factor:* | **1,0000** |
| *Injection Date/Time:* | **13.apr..21 10:27** | *Sample Weight:* | **1,0000** |

##### Chromatogram

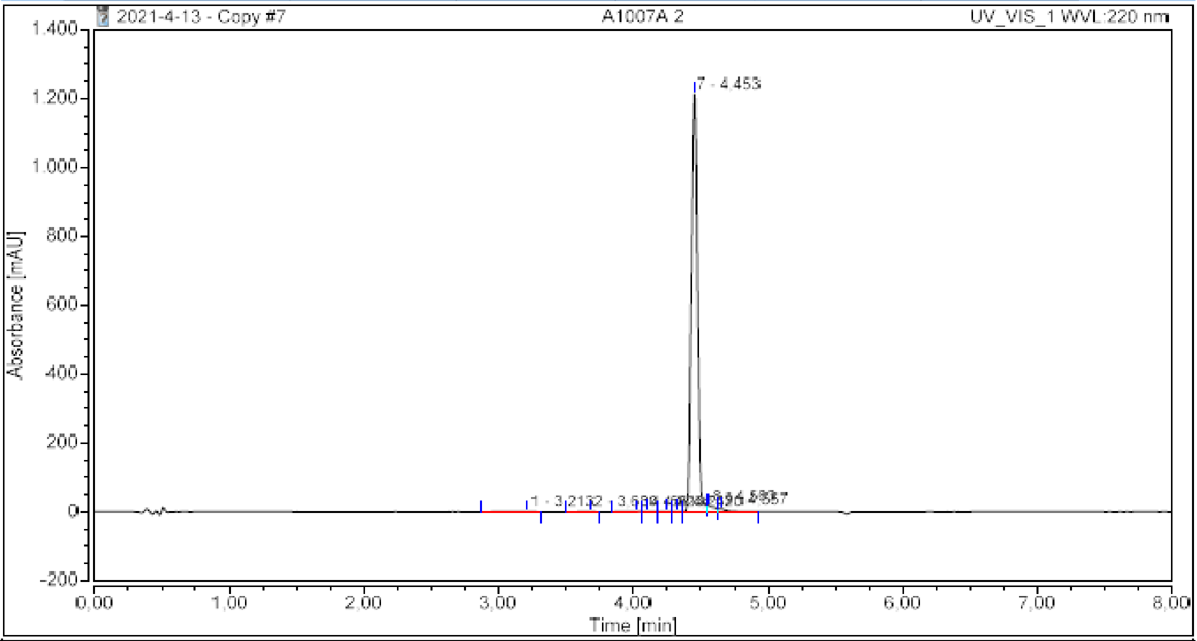

##### Integration Results

| No. | Peak Name | Retention Time min | Area mAU*min | Height mAU | Relative Area % | Relative Height % | Amount n.a. |
|---|---|---|---|---|---|---|---|
| 1 | | 3,213 | 0,058 | 0,249 | 0,09 | 0,02 | n.a. |
| 2 | | 3,680 | 0,021 | 0,226 | 0,03 | 0,02 | n.a. |
| 3 | | 4,023 | 0,044 | 0,534 | 0,07 | 0,04 | n.a. |
| 4 | | 4,107 | 0,049 | 0,940 | 0,07 | 0,08 | n.a. |
| 5 | | 4,247 | 0,048 | 1,080 | 0,07 | 0,09 | n.a. |
| 6 | | 4,320 | 0,040 | 0,761 | 0,06 | 0,06 | n.a. |
| 7 | | 4,453 | 64,736 | 1213,706 | 96,68 | 97,66 | n.a. |
| 8 | | 4,563 | 1,200 | 16,863 | 1,79 | 1,36 | n.a. |
| 9 | | 4,657 | 0,766 | 8,368 | 1,14 | 0,67 | n.a. |
| **Total:** | | | **66,962** | **1242,727** | **100,00** | **100,00** | |

#### Chromatogram and Results

##### Injection Details

| | | | |
|---|---|---|---|
| *Injection Name:* | **A960C fr7** | *Run Time (min):* | **8,00** |
| *Vial Number:* | **BA7** | *Injection Volume:* | **5,00** |
| *Injection Type:* | **Unknown** | *Channel:* | **UV_VIS_2** |
| *Calibration Level:* | | *Wavelength:* | **254** |
| *Instrument Method:* | **General method (0.1TFA_MeCN_20-90)_FKKT** | *Bandwidth:* | **16** |
| *Processing Method:* | **Quantitative** | *Dilution Factor:* | **1,0000** |
| *Injection Date/Time:* | **26.feb..21 09:21** | *Sample Weight:* | **1,0000** |

##### Chromatogram

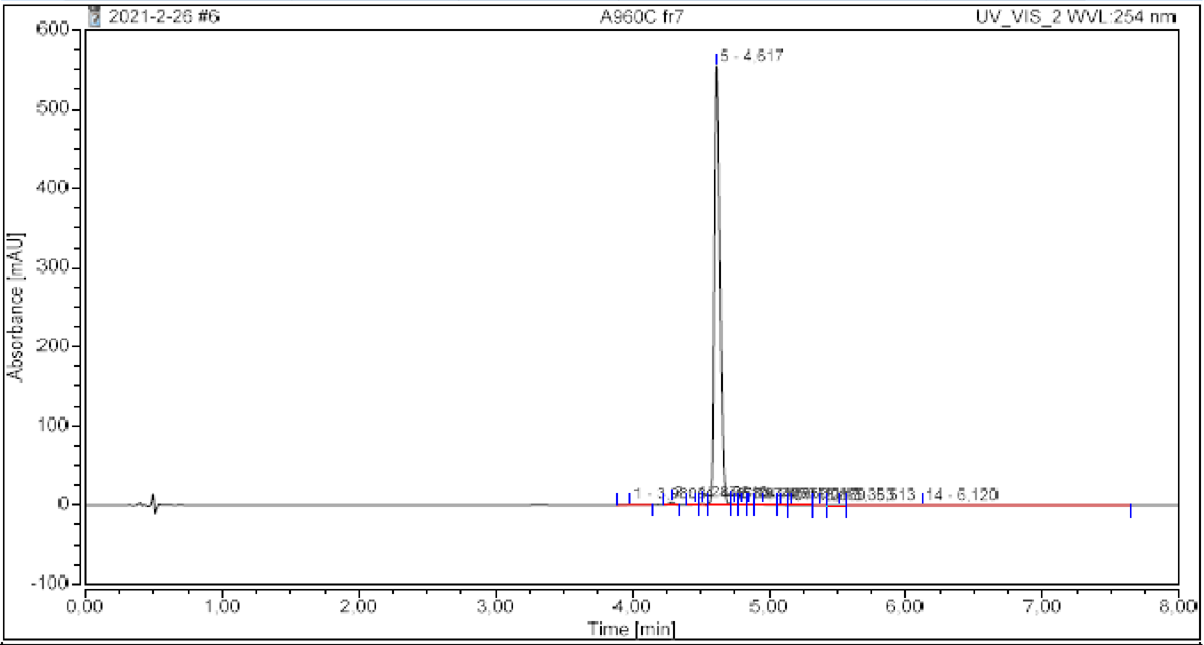

##### Integration Results

| No. | Peak Name | Retention Time min | Area mAU*min | Height mAU | Relative Area % | Relative Height % | Amount n.a. |
|---|---|---|---|---|---|---|---|
| 1 | | 3,980 | 0,047 | 0,296 | 0,17 | 0,05 | n.a. |
| 2 | | 4,287 | 0,157 | 3,830 | 0,56 | 0,68 | n.a. |
| 3 | | 4,457 | 0,036 | 0,888 | 0,13 | 0,16 | n.a. |
| 4 | | 4,510 | 0,084 | 1,874 | 0,30 | 0,33 | n.a. |
| 5 | | 4,617 | 26,750 | 554,514 | 96,08 | 97,74 | n.a. |
| 6 | | 4,740 | 0,061 | 1,484 | 0,22 | 0,26 | n.a. |
| 7 | | 4,797 | 0,063 | 1,070 | 0,22 | 0,19 | n.a. |
| 8 | | 4,853 | 0,046 | 0,865 | 0,17 | 0,15 | n.a. |
| 9 | | 4,950 | 0,088 | 1,041 | 0,32 | 0,18 | n.a. |
| 10 | | 5,087 | 0,021 | 0,295 | 0,08 | 0,05 | n.a. |
| 11 | | 5,160 | 0,045 | 0,230 | 0,16 | 0,04 | n.a. |
| 12 | | 5,363 | 0,027 | 0,296 | 0,10 | 0,05 | n.a. |
| 13 | | 5,513 | 0,042 | 0,350 | 0,15 | 0,06 | n.a. |
| 14 | | 6,120 | 0,375 | 0,281 | 1,35 | 0,05 | n.a. |
| **Total:** | | | **27,843** | **567,313** | **100,00** | **100,00** | |

**17**

#### Chromatogram and Results

**Injection Details**

| | | | |
|---|---|---|---|
| *Injection Name:* | **A920A fr5** | *Run Time (min):* | **8,00** |
| *Vial Number:* | **BB8** | *Injection Volume:* | **5,00** |
| *Injection Type:* | **Unknown** | *Channel:* | **UV_VIS_2** |
| *Calibration Level:* | | *Wavelength:* | **254** |
| *Instrument Method:* | **General method (0.1TFA_MeCN_20-90)_FKKT** | *Bandwidth:* | **16** |
| *Processing Method:* | **Quantitative** | *Dilution Factor:* | **1,0000** |
| *Injection Date/Time:* | **03.feb..21 10:23** | *Sample Weight:* | **1,0000** |

**Chromatogram**

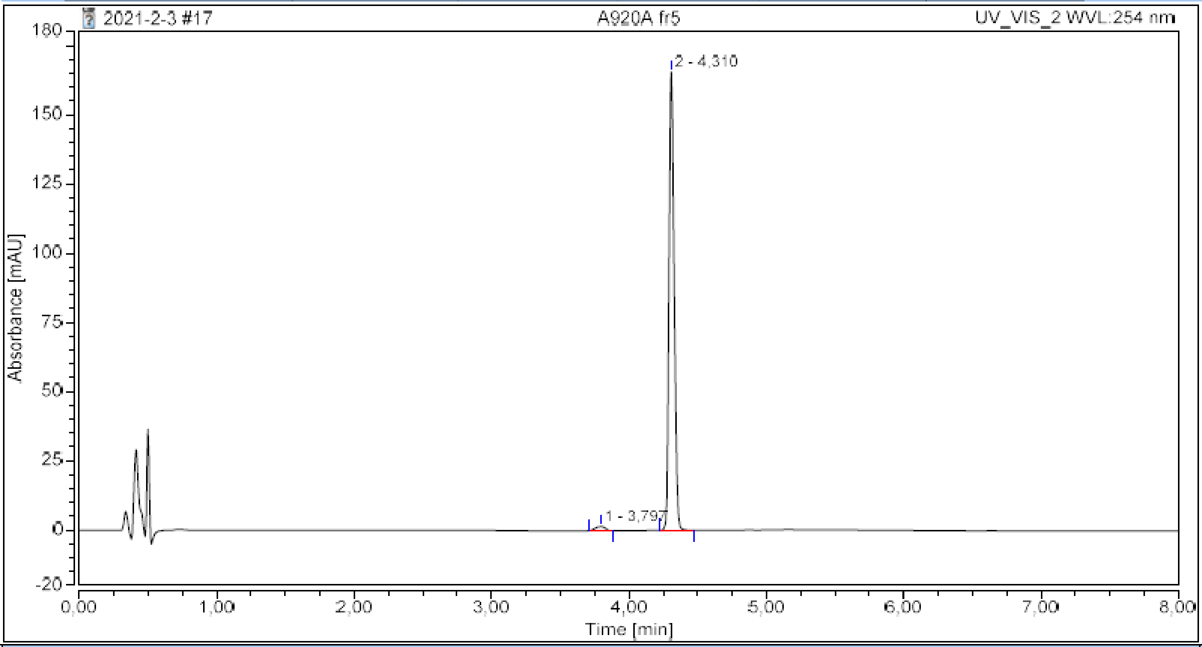

**Integration Results**

| No. | Peak Name | Retention Time min | Area mAU*min | Height mAU | Relative Area % | Relative Height % | Amount n.a. |
|---|---|---|---|---|---|---|---|
| 1 | | 3,797 | 0,124 | 1,534 | 1,69 | 0,92 | n.a. |
| 2 | | 4,310 | 7,184 | 165,456 | 98,31 | 99,08 | n.a. |
| **Total:** | | | **7,308** | **166,990** | **100,00** | **100,00** | |

**17b**

#### Chromatogram and Results

**Injection Details**

| | | | |
|---|---|---|---|
| *Injection Name:* | **A1026** | *Run Time (min):* | **6,00** |
| *Vial Number:* | **GB5** | *Injection Volume:* | **5,00** |
| *Injection Type:* | **Unknown** | *Channel:* | **UV_VIS_1** |
| *Calibration Level:* | | *Wavelength:* | **220** |
| *Instrument Method:* | **Kinetics method 8 per h 0.4 mL** | *Bandwidth:* | **8** |
| *Processing Method:* | **Quantitative** | *Dilution Factor:* | **1,0000** |
| *Injection Date/Time:* | **04.jul..22 17:16** | *Sample Weight:* | **1,0000** |

**Chromatogram**

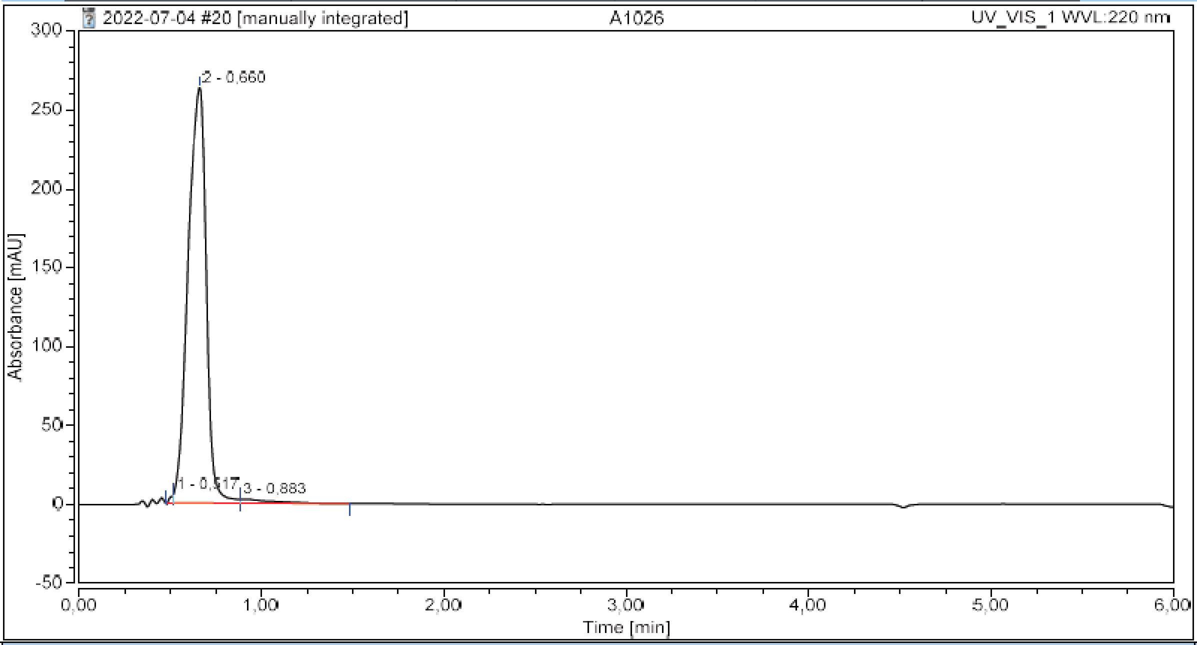

**Integration Results**

| No. | Peak Name | Retention Time min | Area mAU*min | Height mAU | Relative Area % | Relative Height % | Amount n.a. |
|---|---|---|---|---|---|---|---|
| 1 | | 0,517 | 0,099 | 5,495 | 0,32 | 2,03 | n.a. |
| 2 | | 0,660 | 30,817 | 263,244 | 98,13 | 97,09 | n.a. |
| 3 | | 0,883 | 0,489 | 2,393 | 1,56 | 0,88 | n.a. |
| **Total:** | | | **31,405** | **271,132** | **100,00** | **100,00** | |

**18**

#### Chromatogram and Results

##### Injection Details

| | | | |
|---|---|---|---|
| *Injection Name:* | **A878B1 fr4** | *Run Time (min):* | **8,00** |
| *Vial Number:* | **BB1** | *Injection Volume:* | **5,00** |
| *Injection Type:* | **Unknown** | *Channel:* | **UV_VIS_1** |
| *Calibration Level:* | | *Wavelength:* | **220** |
| *Instrument Method:* | **General method (0.1TFA_MeCN_20-90)_FKKT** | *Bandwidth:* | **8** |
| *Processing Method:* | **Quantitative** | *Dilution Factor:* | **1,0000** |
| *Injection Date/Time:* | **22.dec..20 10:21** | *Sample Weight:* | **1,0000** |

##### Chromatogram

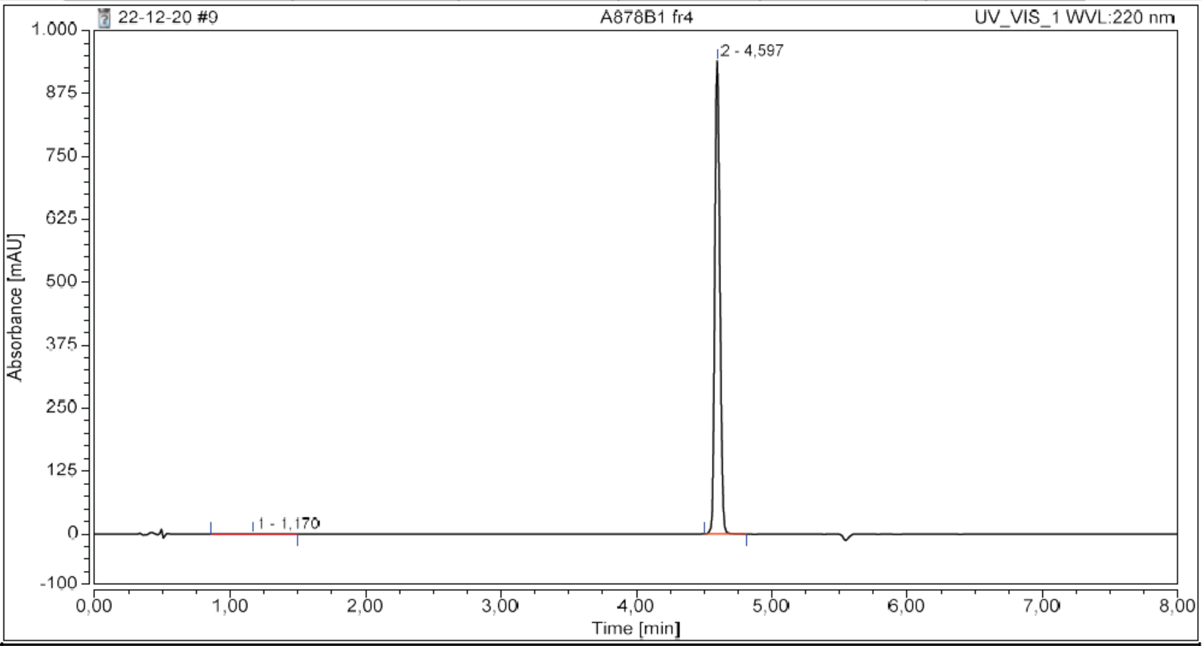

##### Integration Results

| No. | Peak Name | Retention Time min | Area mAU*min | Height mAU | Relative Area % | Relative Height % | Amount n.a. |
|---|---|---|---|---|---|---|---|
| 1 | | 1,170 | 0,059 | 0,201 | 0,14 | 0,02 | n.a. |
| 2 | | 4,597 | 41,016 | 939,691 | 99,86 | 99,98 | n.a. |
| **Total:** | | | **41,075** | **939,892** | **100,00** | **100,00** | |

#### Chromatogram and Results

##### Injection Details

| | | | |
|---|---|---|---|
| *Injection Name:* | **A950A fr7** | *Run Time (min):* | **8,00** |
| *Vial Number:* | **RA3** | *Injection Volume:* | **5,00** |
| *Injection Type:* | **Unknown** | *Channel:* | **UV_VIS_2** |
| *Calibration Level:* | | *Wavelength:* | **254** |
| *Instrument Method:* | **General method (0.1TFA_MeCN_20-90)_FKKT** | *Bandwidth:* | **16** |
| *Processing Method:* | **Quantitative** | *Dilution Factor:* | **1,0000** |
| *Injection Date/Time:* | **12.feb..21 07:33** | *Sample Weight:* | **1,0000** |

##### Chromatogram

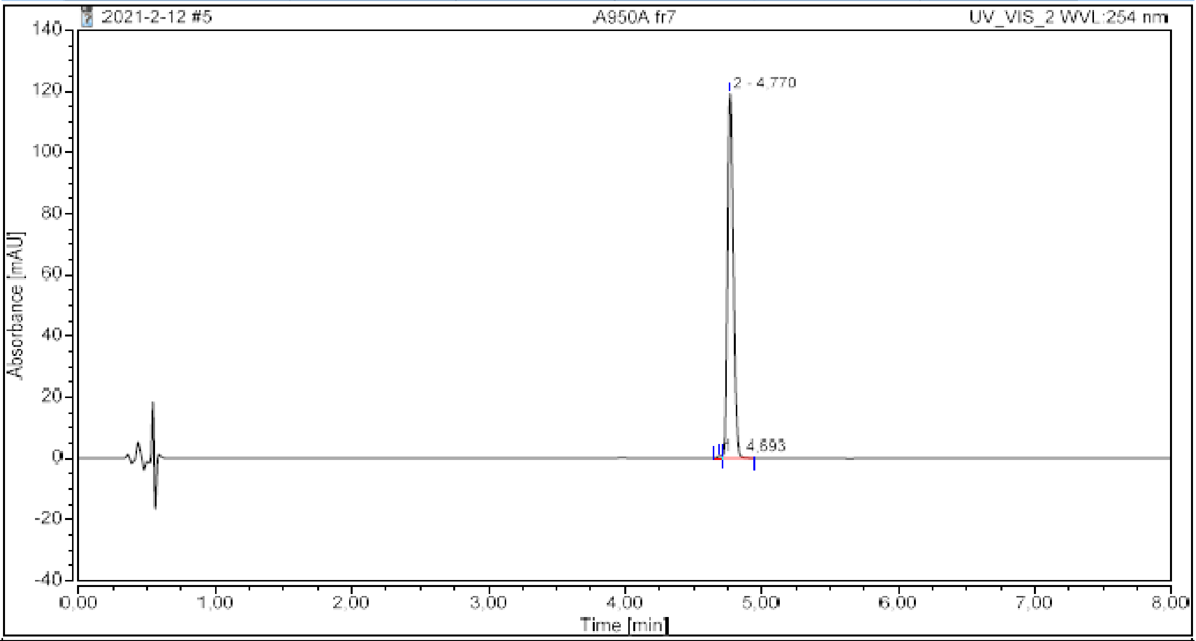

##### Integration Results

| No. | Peak Name | Retention Time<br>min | Area<br>mAU*min | Height<br>mAU | Relative Area<br>% | Relative Height<br>% | Amount<br>n.a. |
|---|---|---|---|---|---|---|---|
| 1 | | 4,693 | 0,023 | 0,692 | 0,39 | 0,58 | n.a. |
| 2 | | 4,770 | 6,020 | 119,232 | 99,61 | 99,42 | n.a. |
| **Total:** | | | **6,043** | **119,924** | **100,00** | **100,00** | |

**19b**

#### Chromatogram and Results

**Injection Details**

| | | | |
|---|---|---|---|
| *Injection Name:* | A1106 | *Run Time (min):* | 6,00 |
| *Vial Number:* | GC5 | *Injection Volume:* | 1,00 |
| *Injection Type:* | Unknown | *Channel:* | UV_VIS_1 |
| *Calibration Level:* | | *Wavelength:* | 220 |
| *Instrument Method:* | Kinetics method 8 per h 0.4 mL | *Bandwidth:* | 8 |
| *Processing Method:* | Quantitative | *Dilution Factor:* | 1,0000 |
| *Injection Date/Time:* | 07.nov..22 17:01 | *Sample Weight:* | 1,0000 |

**Chromatogram**

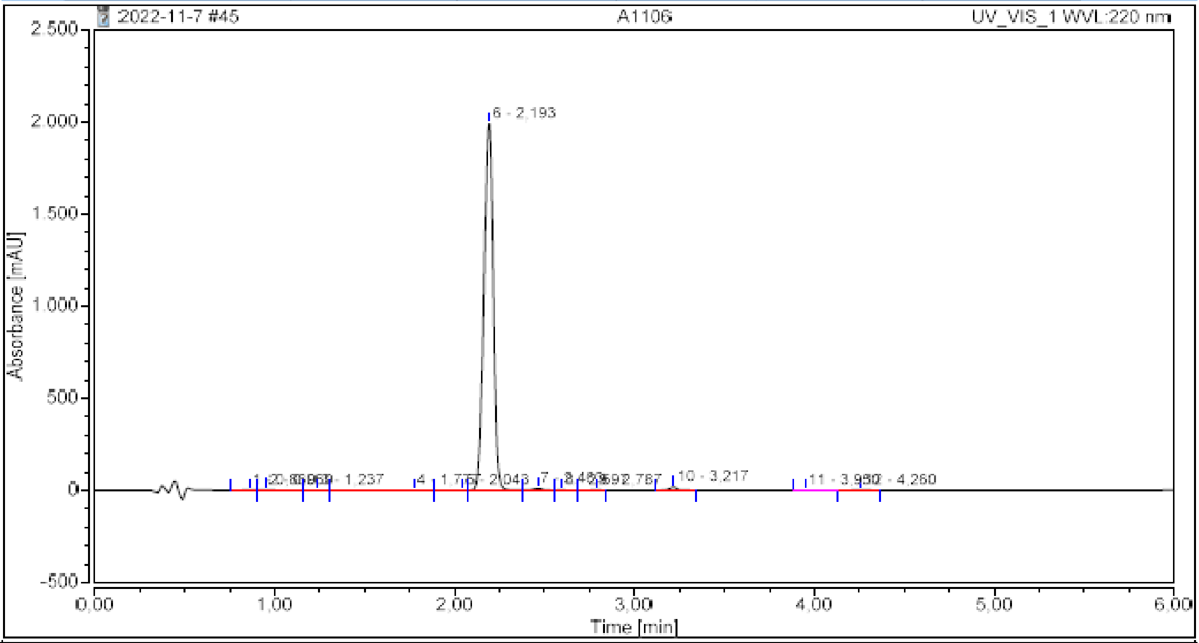

**Integration Results**

| No. | Peak Name | Retention Time min | Area mAU*min | Height mAU | Relative Area % | Relative Height % | Amount n.a. |
|---|---|---|---|---|---|---|---|
| 1 | | 0,860 | 0,159 | 2,324 | 0,13 | 0,11 | n.a. |
| 2 | | 0,950 | 0,252 | 3,070 | 0,21 | 0,15 | n.a. |
| 3 | | 1,237 | 0,043 | 0,408 | 0,04 | 0,02 | n.a. |
| 4 | | 1,777 | 0,199 | 0,471 | 0,16 | 0,02 | n.a. |
| 5 | | 2,043 | 0,042 | 0,367 | 0,03 | 0,02 | n.a. |
| 6 | | 2,193 | 119,470 | 1989,909 | 98,10 | 98,22 | n.a. |
| 7 | | 2,463 | 0,642 | 10,777 | 0,53 | 0,53 | n.a. |
| 8 | | 2,597 | 0,025 | 0,416 | 0,02 | 0,02 | n.a. |
| 9 | | 2,787 | 0,032 | 0,569 | 0,03 | 0,03 | n.a. |
| 10 | | 3,217 | 0,795 | 16,021 | 0,65 | 0,79 | n.a. |
| 11 | | 3,950 | 0,023 | 0,279 | 0,02 | 0,01 | n.a. |
| 12 | | 4,260 | 0,096 | 1,409 | 0,08 | 0,07 | n.a. |
| **Total:** | | | **121,778** | **2026,021** | **100,00** | **100,00** | |

**20**

#### Chromatogram and Results

##### Injection Details

| | | | |
|---|---|---|---|
| *Injection Name:* | **A886D1 fr8** | *Run Time (min):* | **8,00** |
| *Vial Number:* | **RD5** | *Injection Volume:* | **5,00** |
| *Injection Type:* | **Unknown** | *Channel:* | **UV_VIS_2** |
| *Calibration Level:* | | *Wavelength:* | **254** |
| *Instrument Method:* | **General method (0.1TFA_MeCN_20-90)_FKKT** | *Bandwidth:* | **16** |
| *Processing Method:* | **Quantitative** | *Dilution Factor:* | **1,0000** |
| *Injection Date/Time:* | **05.jan..21 10:40** | *Sample Weight:* | **1,0000** |

##### Chromatogram

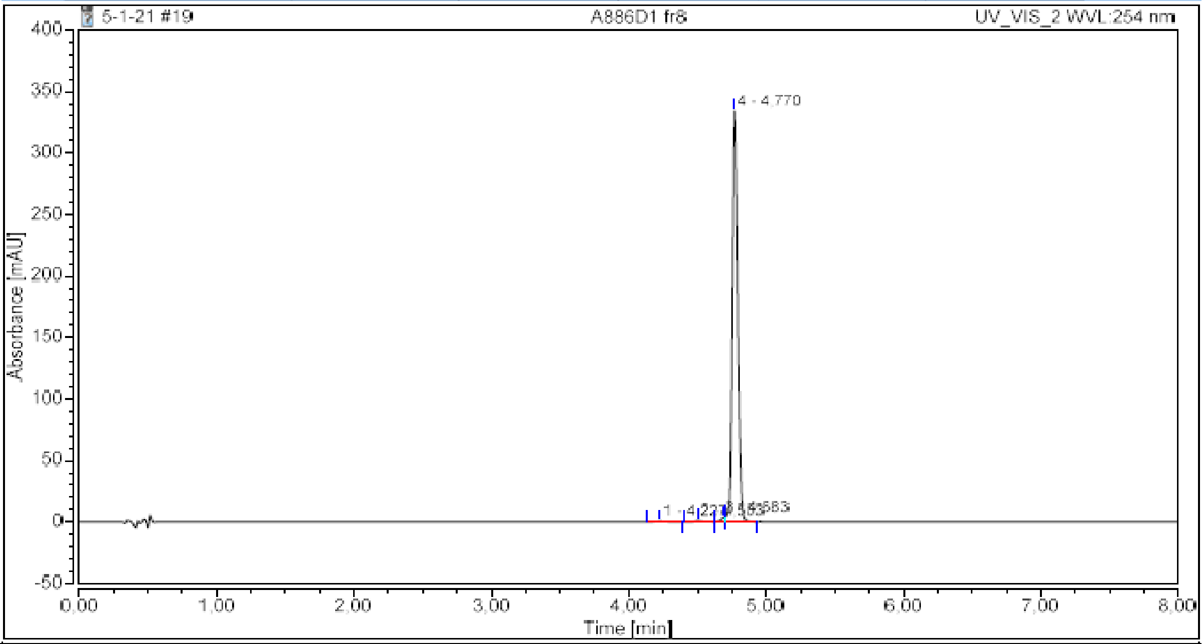

##### Integration Results

| No. | Peak Name | Retention Time min | Area mAU*min | Height mAU | Relative Area % | Relative Height % | Amount n.a. |
|---|---|---|---|---|---|---|---|
| 1 | | 4,227 | 0,050 | 0,608 | 0,30 | 0,18 | n.a. |
| 2 | | 4,503 | 0,067 | 1,100 | 0,40 | 0,32 | n.a. |
| 3 | | 4,683 | 0,120 | 3,053 | 0,71 | 0,90 | n.a. |
| 4 | | 4,770 | 16,522 | 334,147 | 98,58 | 98,60 | n.a. |
| **Total:** | | | **16,759** | **338,908** | **100,00** | **100,00** | |

**21**

#### Chromatogram and Results

**Injection Details**

| | | | |
|---|---|---|---|
| *Injection Name:* | **A920H fr13** | *Run Time (min):* | **8,00** |
| *Vial Number:* | **BC8** | *Injection Volume:* | **5,00** |
| *Injection Type:* | **Unknown** | *Channel:* | **UV_VIS_2** |
| *Calibration Level:* | | *Wavelength:* | **254** |
| *Instrument Method:* | **General method (0.1TFA_MeCN_20-90)_FKKT** | *Bandwidth:* | **16** |
| *Processing Method:* | **Quantitative** | *Dilution Factor:* | **1,0000** |
| *Injection Date/Time:* | **03.feb..21 11:20** | *Sample Weight:* | **1,0000** |

**Chromatogram**

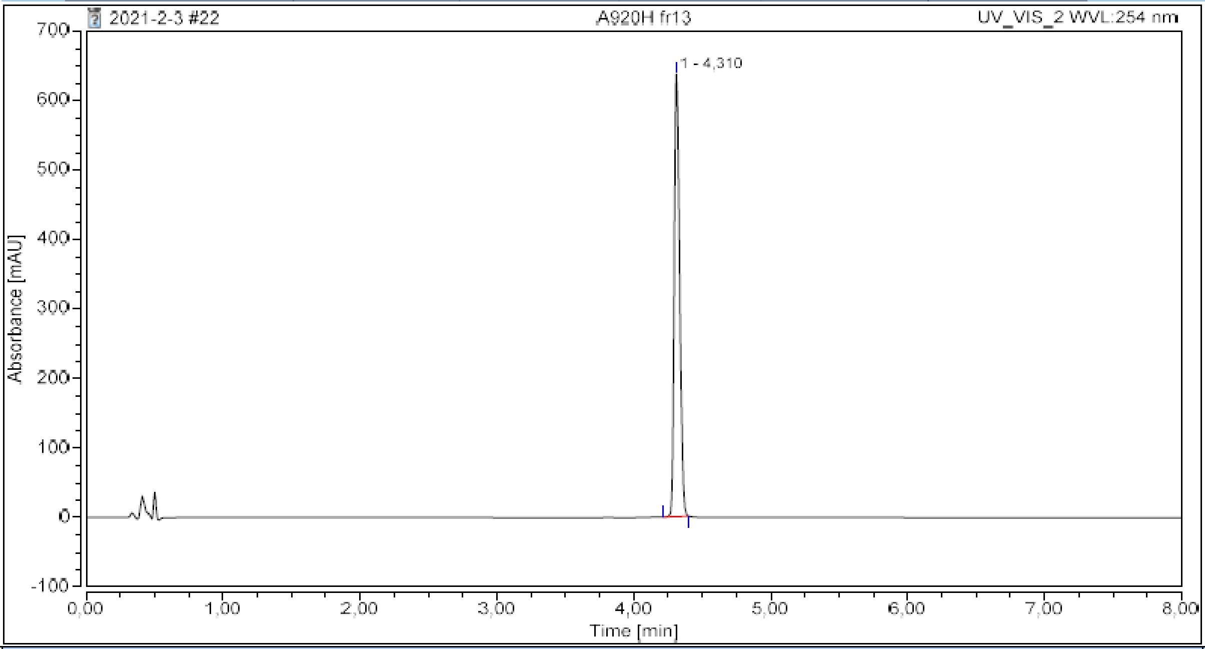

**Integration Results**

| No. | Peak Name | Retention Time min | Area mAU*min | Height mAU | Relative Area % | Relative Height % | Amount n.a. |
|---|---|---|---|---|---|---|---|
| 1 | | 4,310 | 29,213 | 637,960 | 100,00 | 100,00 | n.a. |
| **Total:** | | | **29,213** | **637,960** | **100,00** | **100,00** | |

**21b**

#### Chromatogram and Results

**Injection Details**

| | | | |
|---|---|---|---|
| *Injection Name:* | **A1105** | *Run Time (min):* | **6,00** |
| *Vial Number:* | **GC1** | *Injection Volume:* | **2,00** |
| *Injection Type:* | **Unknown** | *Channel:* | **UV_VIS_2** |
| *Calibration Level:* | | *Wavelength:* | **254** |
| *Instrument Method:* | **Kinetics method 8 per h 0.4 mL** | *Bandwidth:* | **16** |
| *Processing Method:* | **Quantitative** | *Dilution Factor:* | **1,0000** |
| *Injection Date/Time:* | **22.nov..21 11:17** | *Sample Weight:* | **1,0000** |

**Chromatogram**

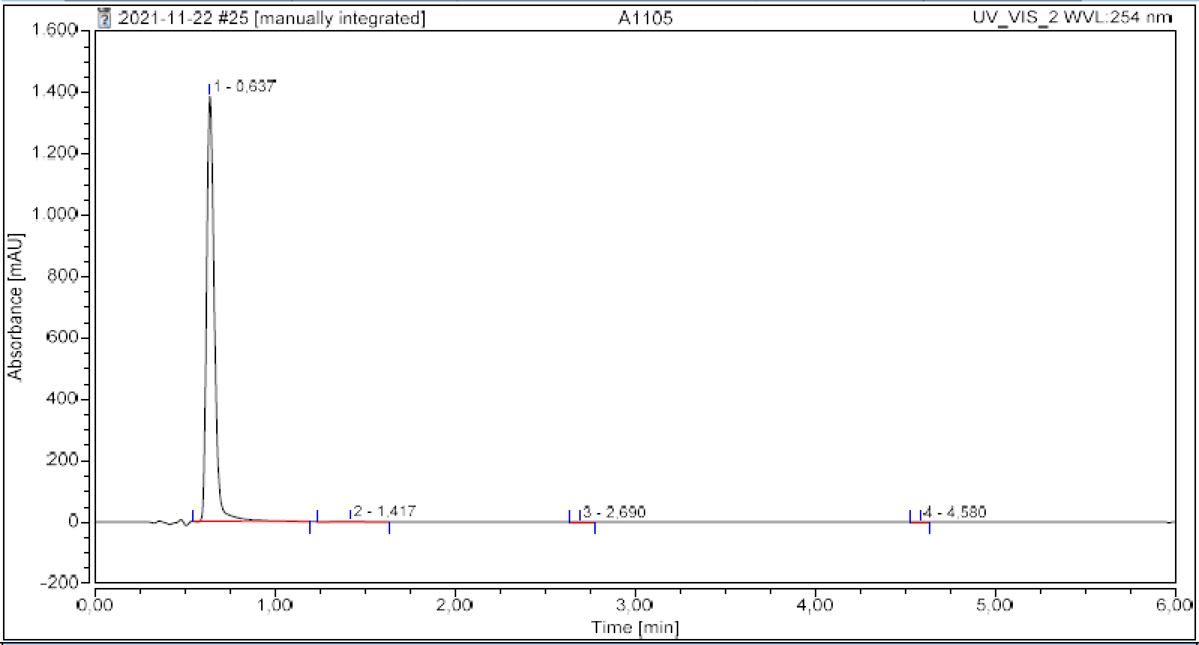

**Integration Results**

| No. | Peak Name | Retention Time min | Area mAU*min | Height mAU | Relative Area % | Relative Height % | Amount n.a. |
|---|---|---|---|---|---|---|---|
| 1 | | 0,637 | 69,963 | 1384,394 | 99,67 | 99,90 | n.a. |
| 2 | | 1,417 | 0,227 | 1,288 | 0,32 | 0,09 | n.a. |
| 3 | | 2,690 | 0,003 | 0,049 | 0,00 | 0,00 | n.a. |
| 4 | | 4,580 | 0,003 | 0,083 | 0,00 | 0,01 | n.a. |
| **Total:** | | | **70,195** | **1385,814** | **100,00** | **100,00** | |

### Chromatogram and Results

#### Injection Details

| | | | |
|---|---|---|---|
| *Injection Name:* | **3** | *Run Time (min):* | **8,00** |
| *Vial Number:* | **BC3** | *Injection Volume:* | **5,00** |
| *Injection Type:* | **Unknown** | *Channel:* | **UV_VIS_1** |
| *Calibration Level:* | | *Wavelength:* | **220** |
| *Instrument Method:* | **General method (0.1TFA_MeCN_20-90)_FKKT** | *Bandwidth:* | **8** |
| *Processing Method:* | **Quantitative** | *Dilution Factor:* | **1,0000** |
| *Injection Date/Time:* | **03.mar..21 08:24** | *Sample Weight:* | **1,0000** |

#### Chromatogram

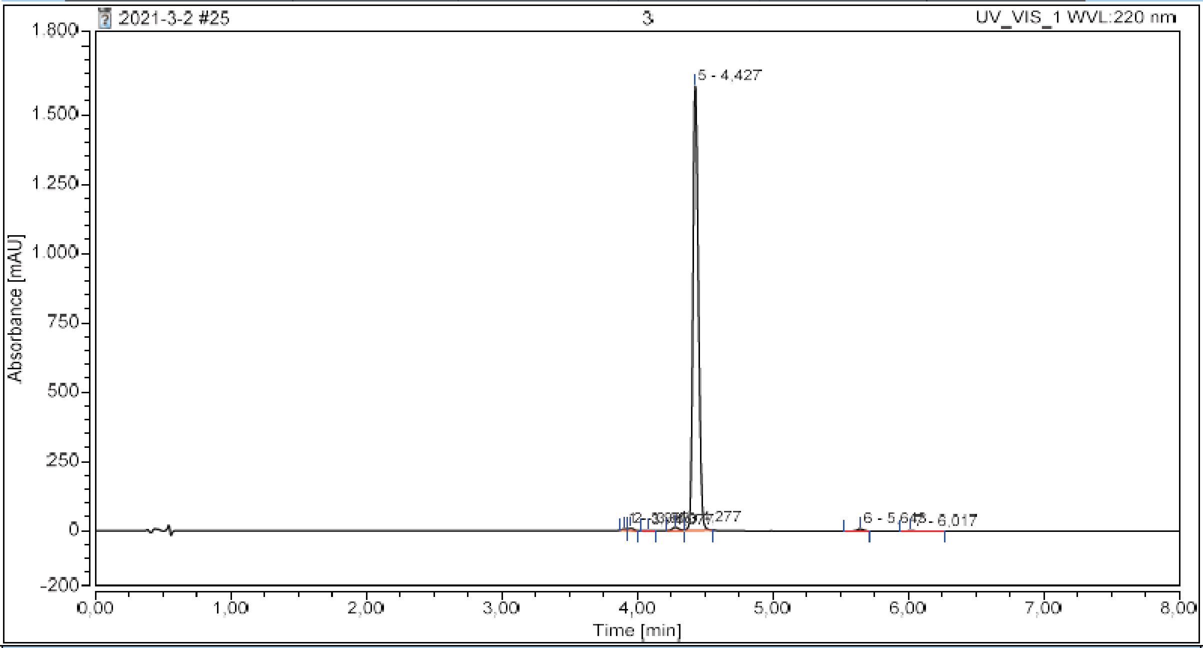

#### Integration Results

| No. | Peak Name | Retention Time min | Area mAU*min | Height mAU | Relative Area % | Relative Height % | Amount n.a. |
|---|---|---|---|---|---|---|---|
| 1 | | 3,903 | 0,164 | 4,611 | 0,21 | 0,28 | n.a. |
| 2 | | 3,950 | 0,378 | 8,535 | 0,49 | 0,52 | n.a. |
| 3 | | 4,077 | 0,038 | 0,823 | 0,05 | 0,05 | n.a. |
| 4 | | 4,277 | 0,674 | 12,176 | 0,87 | 0,74 | n.a. |
| 5 | | 4,427 | 75,525 | 1602,116 | 97,86 | 97,91 | n.a. |
| 6 | | 5,643 | 0,373 | 7,811 | 0,48 | 0,48 | n.a. |
| 7 | | 6,017 | 0,024 | 0,287 | 0,03 | 0,02 | n.a. |
| **Total:** | | | **77,176** | **1636,359** | **100,00** | **100,00** | |

23

#### Chromatogram and Results

**Injection Details**

| | | | |
|---|---|---|---|
| *Injection Name:* | **A1300 = A960E** | *Run Time (min):* | **6,00** |
| *Vial Number:* | **RD2** | *Injection Volume:* | **5,00** |
| *Injection Type:* | **Unknown** | *Channel:* | **UV_VIS_2** |
| *Calibration Level:* | | *Wavelength:* | **254** |
| *Instrument Method:* | **Kinetics method 8 per h 0.4 mL** | *Bandwidth:* | **16** |
| *Processing Method:* | **Quantitative** | *Dilution Factor:* | **1,0000** |
| *Injection Date/Time:* | **28.sep..22 12:07** | *Sample Weight:* | **1,0000** |

**Chromatogram**

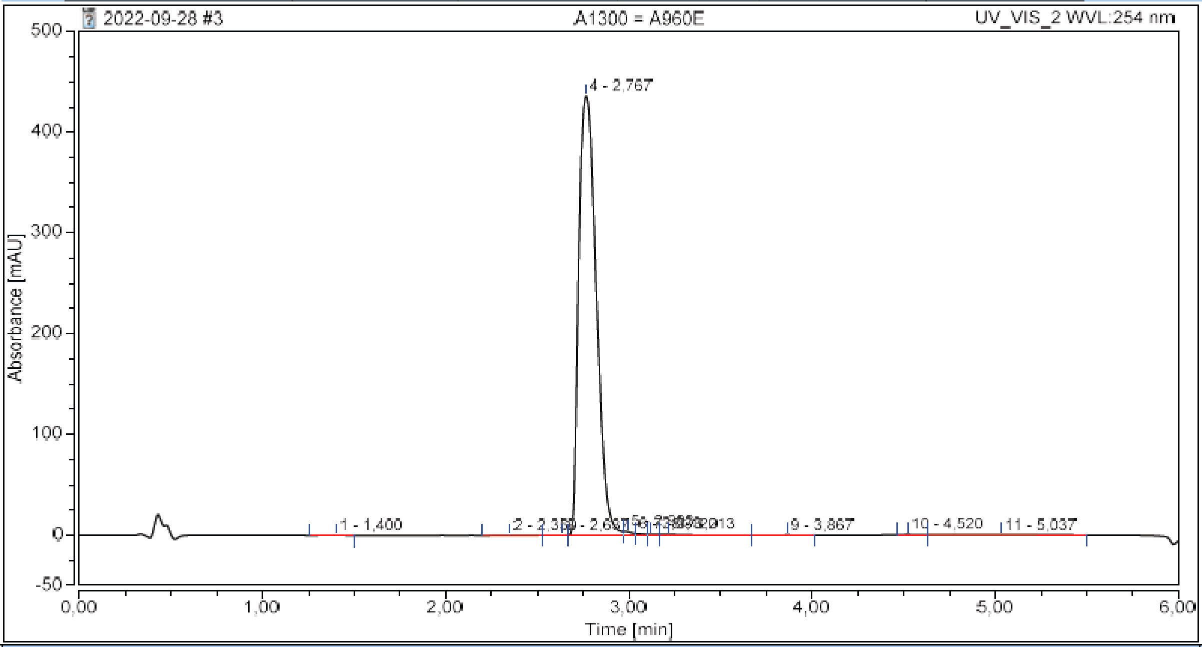

**Integration Results**

| No. | Peak Name | Retention Time min | Area mAU*min | Height mAU | Relative Area % | Relative Height % | Amount n.a. |
|---|---|---|---|---|---|---|---|
| 1 | | 1,400 | 0,029 | 0,270 | 0,06 | 0,06 | n.a. |
| 2 | | 2,350 | 0,032 | 0,263 | 0,07 | 0,06 | n.a. |
| 3 | | 2,637 | 0,021 | 0,385 | 0,04 | 0,09 | n.a. |
| 4 | | 2,767 | 48,310 | 436,438 | 98,65 | 98,00 | n.a. |
| 5 | | 2,993 | 0,187 | 3,564 | 0,38 | 0,80 | n.a. |
| 6 | | 3,033 | 0,094 | 1,718 | 0,19 | 0,39 | n.a. |
| 7 | | 3,120 | 0,063 | 0,979 | 0,13 | 0,22 | n.a. |
| 8 | | 3,213 | 0,137 | 1,068 | 0,28 | 0,24 | n.a. |
| 9 | | 3,867 | 0,012 | 0,165 | 0,03 | 0,04 | n.a. |
| 10 | | 4,520 | 0,017 | 0,365 | 0,03 | 0,08 | n.a. |
| 11 | | 5,037 | 0,069 | 0,124 | 0,14 | 0,03 | n.a. |
| **Total:** | | | **48,972** | **445,339** | **100,00** | **100,00** | |

#### Chromatogram and Results

##### Injection Details

| | | | |
|---|---|---|---|
| *Injection Name:* | **A960G** | *Run Time (min):* | **6,00** |
| *Vial Number:* | **BA2** | *Injection Volume:* | **1,00** |
| *Injection Type:* | **Unknown** | *Channel:* | **UV_VIS_2** |
| *Calibration Level:* | | *Wavelength:* | **254** |
| *Instrument Method:* | **Kinetics method 8 per h 0.4 mL** | *Bandwidth:* | **16** |
| *Processing Method:* | **Quantitative** | *Dilution Factor:* | **1,0000** |
| *Injection Date/Time:* | **30.jun..22 16:57** | *Sample Weight:* | **1,0000** |

##### Chromatogram

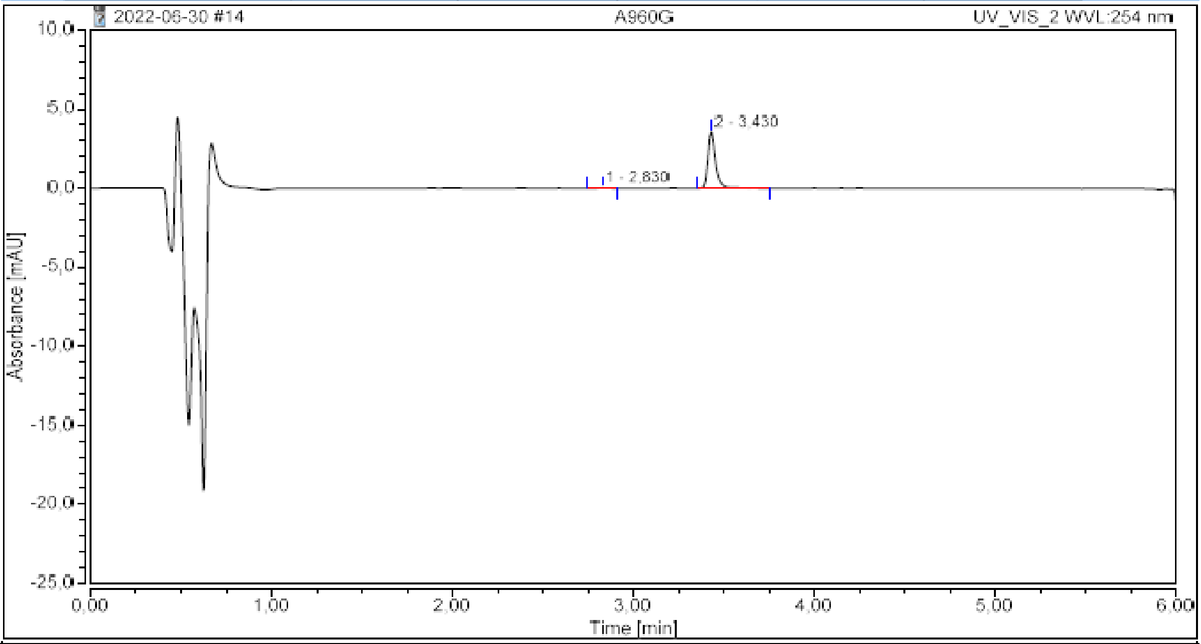

##### Integration Results

| No. | Peak Name | Retention Time<br>min | Area<br>mAU*min | Height<br>mAU | Relative Area<br>% | Relative Height<br>% | Amount<br>n.a. |
|---|---|---|---|---|---|---|---|
| 1 | | 2,830 | 0,002 | 0,030 | 0,93 | 0,83 | n.a. |
| 2 | | 3,430 | 0,171 | 3,575 | 99,07 | 99,17 | n.a. |
| **Total:** | | | **0,172** | **3,605** | **100,00** | **100,00** | |

#### Chromatogram and Results

##### Injection Details

| | | | |
|---|---|---|---|
| *Injection Name:* | **A1009-2** | *Run Time (min):* | **8,00** |
| *Vial Number:* | **BA4** | *Injection Volume:* | **3,00** |
| *Injection Type:* | **Unknown** | *Channel:* | **UV_VIS_1** |
| *Calibration Level:* | | *Wavelength:* | **220** |
| *Instrument Method:* | **General method (0.1TFA_MeCN_20-90)_FKKT** | *Bandwidth:* | **8** |
| *Processing Method:* | **Quantitative** | *Dilution Factor:* | **1,0000** |
| *Injection Date/Time:* | **14.apr..21 11:17** | *Sample Weight:* | **1,0000** |

##### Chromatogram

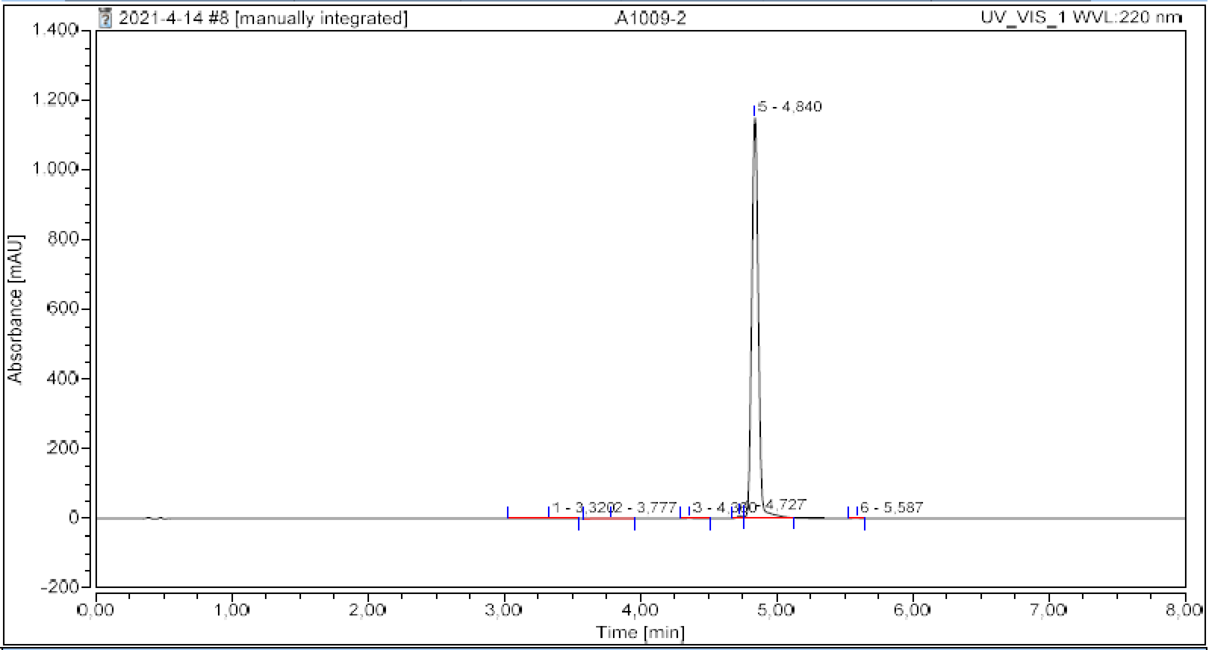

##### Integration Results

| No. | Peak Name | Retention Time min | Area mAU*min | Height mAU | Relative Area % | Relative Height % | Amount n.a. |
|---|---|---|---|---|---|---|---|
| 1 | | 3,320 | 0,094 | 0,330 | 0,14 | 0,03 | n.a. |
| 2 | | 3,777 | 0,045 | 0,235 | 0,07 | 0,02 | n.a. |
| 3 | | 4,360 | 0,044 | 0,708 | 0,07 | 0,06 | n.a. |
| 4 | | 4,727 | 0,315 | 8,025 | 0,48 | 0,69 | n.a. |
| 5 | | 4,840 | 64,848 | 1150,315 | 99,19 | 99,13 | n.a. |
| 6 | | 5,587 | 0,034 | 0,853 | 0,05 | 0,07 | n.a. |
| **Total:** | | | **65,380** | **1160,467** | **100,00** | **100,00** | |

**26**

#### Chromatogram and Results

##### Injection Details

| | | | |
|---|---|---|---|
| *Injection Name:* | **A866-3** | *Run Time (min):* | **8,00** |
| *Vial Number:* | **GA4** | *Injection Volume:* | **5,00** |
| *Injection Type:* | **Unknown** | *Channel:* | **UV_VIS_2** |
| *Calibration Level:* | | *Wavelength:* | **254** |
| *Instrument Method:* | **General method (0.1TFA_MeCN_20-90)_FKKT** | *Bandwidth:* | **16** |
| *Processing Method:* | **Quantitative** | *Dilution Factor:* | **1,0000** |
| *Injection Date/Time:* | **18.dec..20 11:08** | *Sample Weight:* | **1,0000** |

##### Chromatogram

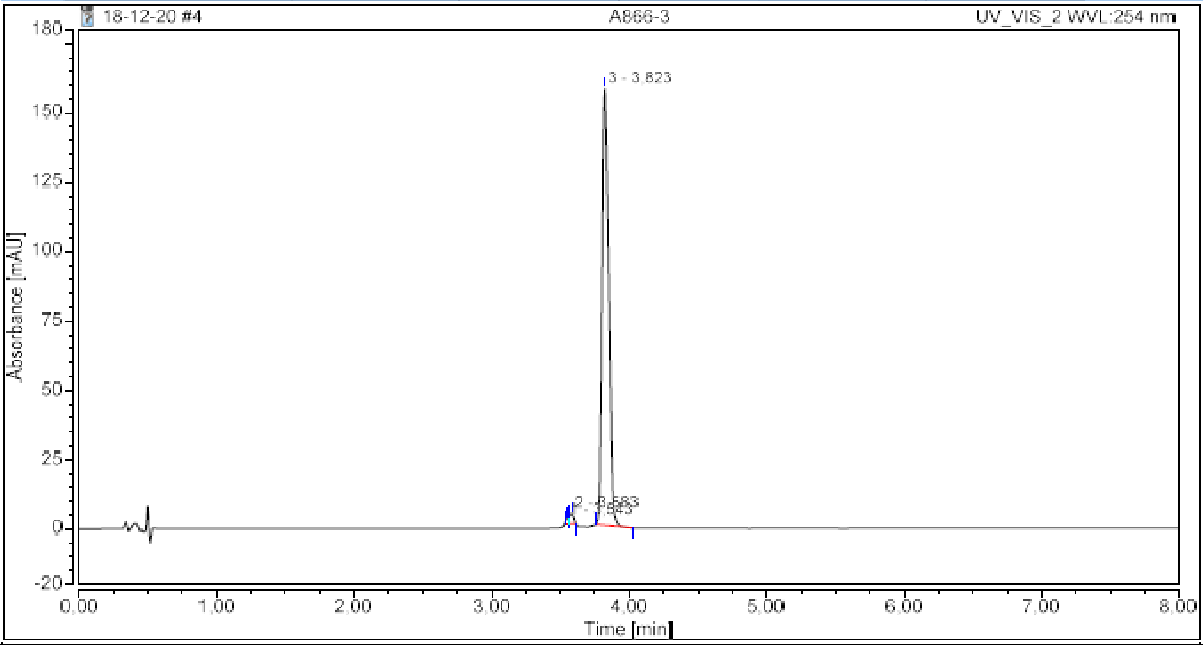

##### Integration Results

| No. | Peak Name | Retention Time min | Area mAU*min | Height mAU | Relative Area % | Relative Height % | Amount n.a. |
|---|---|---|---|---|---|---|---|
| 1 | | 3,543 | 0,044 | 1,524 | 0,49 | 0,94 | n.a. |
| 2 | | 3,583 | 0,134 | 3,619 | 1,49 | 2,22 | n.a. |
| 3 | | 3,823 | 8,829 | 157,601 | 98,02 | 96,84 | n.a. |
| **Total:** | | | **9,008** | **162,743** | **100,00** | **100,00** | |

### Chromatogram and Results

#### Injection Details

| | | | |
|---|---|---|---|
| *Injection Name:* | **A920G f6** | *Run Time (min):* | **8,00** |
| *Vial Number:* | **RA3** | *Injection Volume:* | **5,00** |
| *Injection Type:* | **Unknown** | *Channel:* | **UV_VIS_2** |
| *Calibration Level:* | | *Wavelength:* | **254** |
| *Instrument Method:* | **General method (0.1TFA_MeCN_20-90)_FKKT** | *Bandwidth:* | **16** |
| *Processing Method:* | **Quantitative** | *Dilution Factor:* | **1,0000** |
| *Injection Date/Time:* | **04.feb..21 09:20** | *Sample Weight:* | **1,0000** |

#### Chromatogram

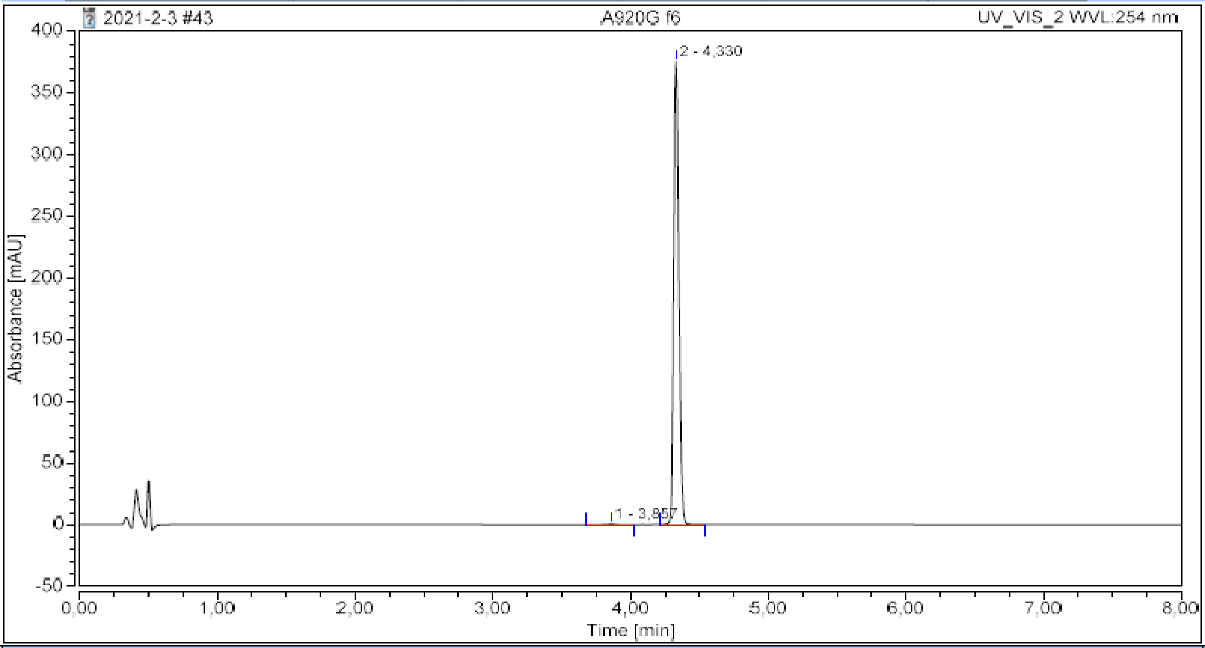

#### Integration Results

| No. | Peak Name | Retention Time min | Area mAU*min | Height mAU | Relative Area % | Relative Height % | Amount n.a. |
|---|---|---|---|---|---|---|---|
| 1 | | 3,857 | 0,056 | 0,634 | 0,34 | 0,17 | n.a. |
| 2 | | 4,330 | 16,540 | 375,543 | 99,66 | 99,83 | n.a. |
| **Total:** | | | **16,596** | **376,177** | **100,00** | **100,00** | |

**28**

#### Chromatogram and Results

##### Injection Details

| | | | |
|---|---|---|---|
| *Injection Name:* | A886B1 fr6 | *Run Time (min):* | 8,00 |
| *Vial Number:* | RC4 | *Injection Volume:* | 5,00 |
| *Injection Type:* | Unknown | *Channel:* | UV_VIS_2 |
| *Calibration Level:* | | *Wavelength:* | 254 |
| *Instrument Method:* | General method (0.1TFA_MeCN_20-90)_FKKT | *Bandwidth:* | 16 |
| *Processing Method:* | Quantitative | *Dilution Factor:* | 1,0000 |
| *Injection Date/Time:* | 05.jan..21 09:43 | *Sample Weight:* | 1,0000 |

##### Chromatogram

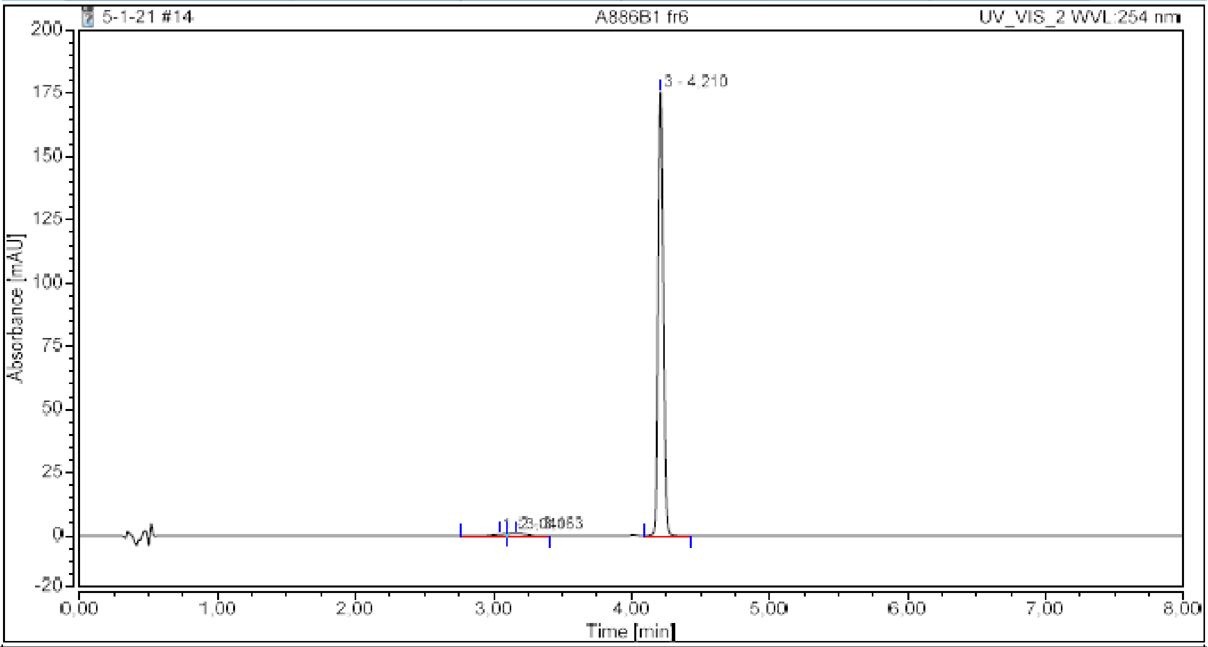

##### Integration Results

| No. | Peak Name | Retention Time<br>min | Area<br>mAU*min | Height<br>mAU | Relative Area<br>% | Relative Height<br>% | Amount<br>n.a. |
|---|---|---|---|---|---|---|---|
| 1 | | 3,040 | 0,081 | 0,617 | 1,04 | 0,35 | n.a. |
| 2 | | 3,163 | 0,137 | 0,967 | 1,76 | 0,55 | n.a. |
| 3 | | 4,210 | 7,560 | 175,751 | 97,20 | 99,11 | n.a. |
| **Total:** | | | **7,778** | **177,336** | **100,00** | **100,00** | |

#### Chromatogram and Results

##### Injection Details

| | | | |
|---|---|---|---|
| *Injection Name:* | **A992B1 fr2** | *Run Time (min):* | **8,00** |
| *Vial Number:* | **BB6** | *Injection Volume:* | **5,00** |
| *Injection Type:* | **Unknown** | *Channel:* | **UV_VIS_1** |
| *Calibration Level:* | | *Wavelength:* | **220** |
| *Instrument Method:* | **General method (0.1TFA_MeCN_20-90)_FKKT** | *Bandwidth:* | **8** |
| *Processing Method:* | **Quantitative** | *Dilution Factor:* | **1,0000** |
| *Injection Date/Time:* | **22.mar..21 14:37** | *Sample Weight:* | **1,0000** |

##### Chromatogram

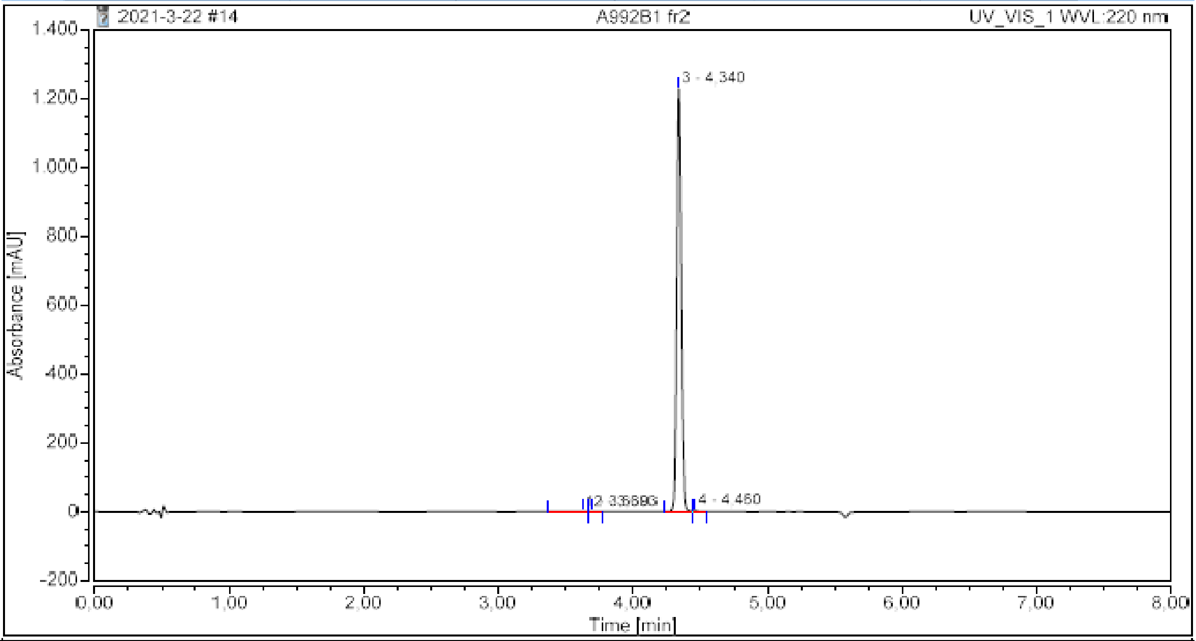

##### Integration Results

| No. | Peak Name | Retention Time min | Area mAU*min | Height mAU | Relative Area % | Relative Height % | Amount n.a. |
|---|---|---|---|---|---|---|---|
| 1 | | 3,630 | 0,097 | 0,784 | 0,20 | 0,06 | n.a. |
| 2 | | 3,693 | 0,052 | 0,832 | 0,11 | 0,07 | n.a. |
| 3 | | 4,340 | 48,587 | 1229,433 | 99,36 | 99,61 | n.a. |
| 4 | | 4,460 | 0,166 | 3,162 | 0,34 | 0,26 | n.a. |
| **Total:** | | | **48,902** | **1234,210** | **100,00** | **100,00** | |

#### Chromatogram and Results

##### Injection Details

| | | | |
|---|---|---|---|
| *Injection Name:* | **A920C resyn fr2** | *Run Time (min):* | **8,00** |
| *Vial Number:* | **GC2** | *Injection Volume:* | **5,00** |
| *Injection Type:* | **Unknown** | *Channel:* | **UV_VIS_2** |
| *Calibration Level:* | | *Wavelength:* | **254** |
| *Instrument Method:* | **General method (0.1TFA_MeCN_20-90)_FKKT** | *Bandwidth:* | **16** |
| *Processing Method:* | **Quantitative** | *Dilution Factor:* | **1,0000** |
| *Injection Date/Time:* | **04.jun..21 10:48** | *Sample Weight:* | **1,0000** |

##### Chromatogram

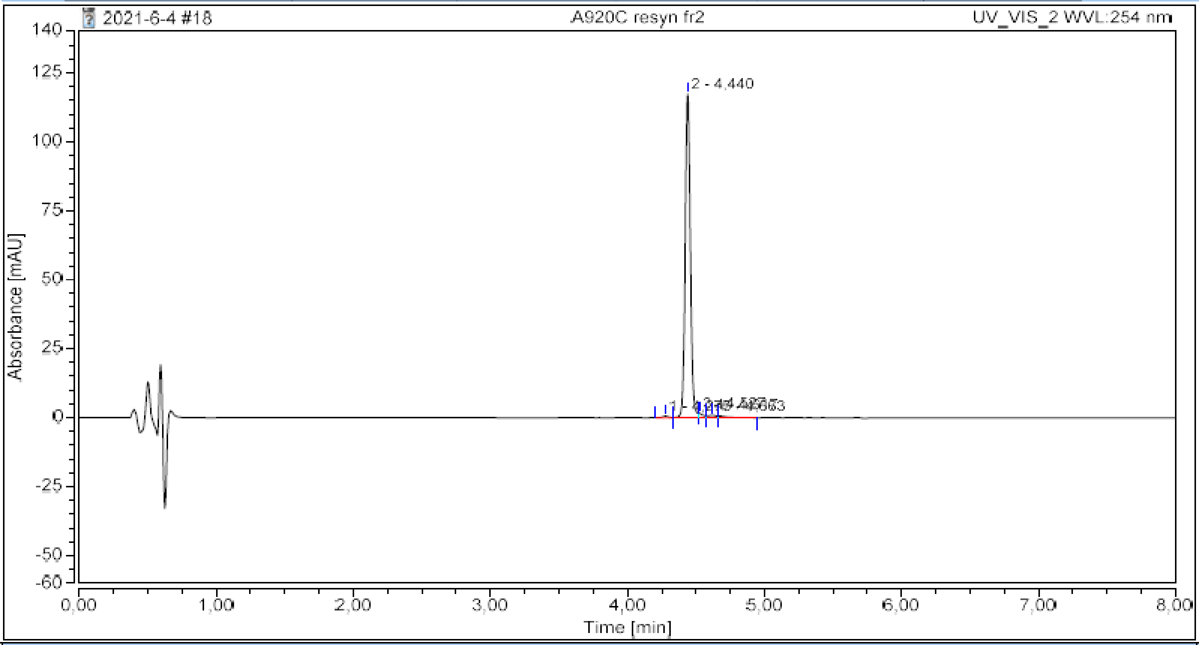

##### Integration Results

| No. | Peak Name | Retention Time min | Area mAU*min | Height mAU | Relative Area % | Relative Height % | Amount n.a. |
|---|---|---|---|---|---|---|---|
| 1 | | 4,277 | 0,034 | 0,628 | 0,64 | 0,52 | n.a. |
| 2 | | 4,440 | 5,113 | 117,272 | 95,41 | 96,81 | n.a. |
| 3 | | 4,527 | 0,076 | 1,635 | 1,42 | 1,35 | n.a. |
| 4 | | 4,617 | 0,072 | 0,973 | 1,34 | 0,80 | n.a. |
| 5 | | 4,663 | 0,064 | 0,625 | 1,19 | 0,52 | n.a. |
| **Total:** | | | **5,359** | **121,134** | **100,00** | **100,00** | |

31

#### Chromatogram and Results

**Injection Details**

| | | | |
|---|---|---|---|
| *Injection Name:* | **A920E fr5** | *Run Time (min):* | **8,00** |
| *Vial Number:* | **BA7** | *Injection Volume:* | **5,00** |
| *Injection Type:* | **Unknown** | *Channel:* | **UV_VIS_2** |
| *Calibration Level:* | | *Wavelength:* | **254** |
| *Instrument Method:* | **General method (0.1TFA_MeCN_20-90)_FKKT** | *Bandwidth:* | **16** |
| *Processing Method:* | **Quantitative** | *Dilution Factor:* | **1,0000** |
| *Injection Date/Time:* | **03.feb..21 08:40** | *Sample Weight:* | **1,0000** |

**Chromatogram**

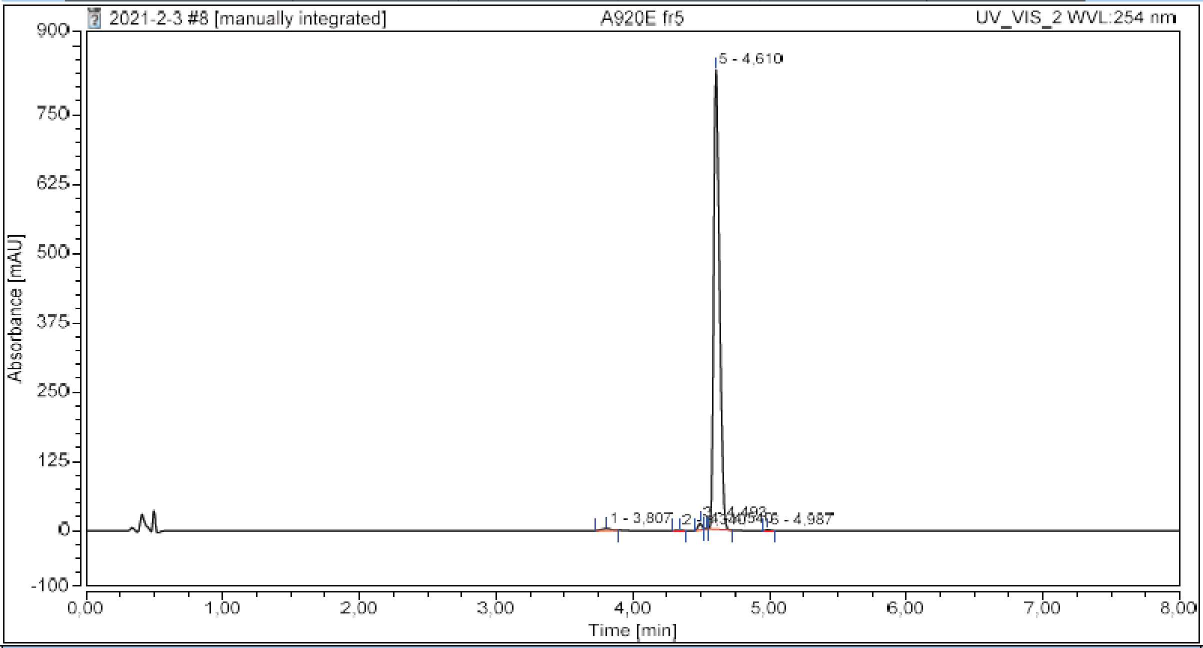

**Integration Results**

| No. | Peak Name | Retention Time<br>min | Area<br>mAU*min | Height<br>mAU | Relative Area<br>% | Relative Height<br>% | Amount<br>n.a. |
|---|---|---|---|---|---|---|---|
| 1 | | 3,807 | 0,251 | 3,454 | 0,61 | 0,41 | n.a. |
| 2 | | 4,340 | 0,041 | 0,945 | 0,10 | 0,11 | n.a. |
| 3 | | 4,493 | 0,398 | 12,294 | 0,96 | 1,45 | n.a. |
| 4 | | 4,540 | 0,023 | 1,074 | 0,06 | 0,13 | n.a. |
| 5 | | 4,610 | 40,730 | 828,878 | 98,15 | 97,73 | n.a. |
| 6 | | 4,987 | 0,056 | 1,514 | 0,13 | 0,18 | n.a. |
| **Total:** | | | **41,499** | **848,159** | **100,00** | **100,00** | |

#### Chromatogram and Results

##### Injection Details

| | | | |
|---|---|---|---|
| *Injection Name:* | **A1308A1 re** | *Run Time (min):* | **6,00** |
| *Vial Number:* | **RE4** | *Injection Volume:* | **5,00** |
| *Injection Type:* | **Unknown** | *Channel:* | **UV_VIS_2** |
| *Calibration Level:* | | *Wavelength:* | **254** |
| *Instrument Method:* | **Kinetics method 8 per h 0.4 mL** | *Bandwidth:* | **16** |
| *Processing Method:* | **Quantitative** | *Dilution Factor:* | **1,0000** |
| *Injection Date/Time:* | **14.nov..22 18:18** | *Sample Weight:* | **1,0000** |

##### Chromatogram

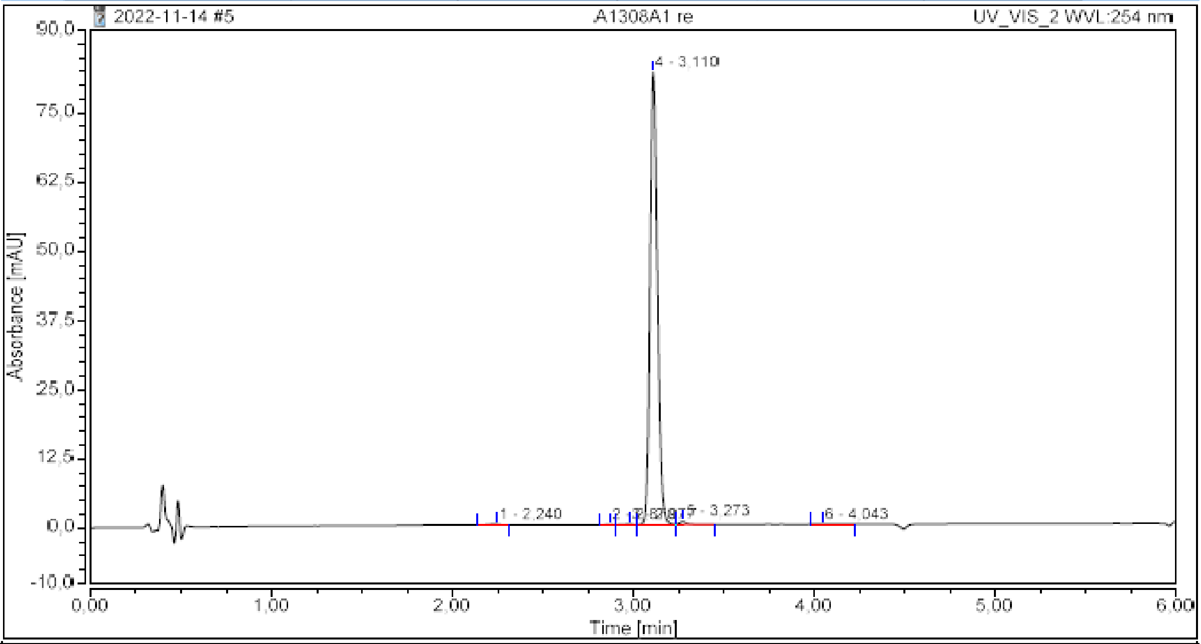

##### Integration Results

| No. | Peak Name | Retention Time min | Area mAU*min | Height mAU | Relative Area % | Relative Height % | Amount n.a. |
|---|---|---|---|---|---|---|---|
| 1 | | 2,240 | 0,011 | 0,148 | 0,26 | 0,18 | n.a. |
| 2 | | 2,870 | 0,003 | 0,058 | 0,08 | 0,07 | n.a. |
| 3 | | 2,977 | 0,007 | 0,118 | 0,17 | 0,14 | n.a. |
| 4 | | 3,110 | 3,918 | 81,717 | 98,71 | 98,88 | n.a. |
| 5 | | 3,273 | 0,027 | 0,557 | 0,69 | 0,67 | n.a. |
| 6 | | 4,043 | 0,004 | 0,040 | 0,09 | 0,05 | n.a. |
| **Total:** | | | **3,969** | **82,639** | **100,00** | **100,00** | |

**33a**

#### Chromatogram and Results

**Injection Details**

| | | | |
|---|---|---|---|
| *Injection Name:* | **A1333** | *Run Time (min):* | **6,00** |
| *Vial Number:* | **GA6** | *Injection Volume:* | **5,00** |
| *Injection Type:* | **Unknown** | *Channel:* | **UV_VIS_1** |
| *Calibration Level:* | | *Wavelength:* | **220** |
| *Instrument Method:* | **Kinetics method 8 per h 0.4 mL** | *Bandwidth:* | **8** |
| *Processing Method:* | **Quantitative** | *Dilution Factor:* | **1,0000** |
| *Injection Date/Time:* | **13.sep..23 09:26** | *Sample Weight:* | **1,0000** |

**Chromatogram**

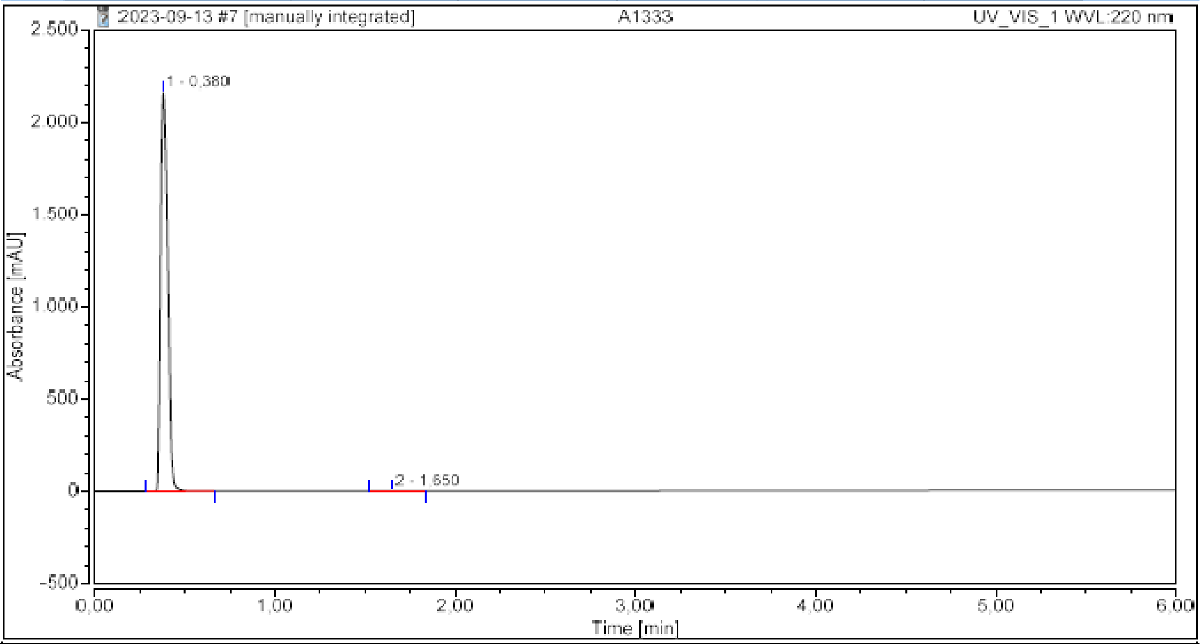

**Integration Results**

| No. | Peak Name | Retention Time min | Area mAU*min | Height mAU | Relative Area % | Relative Height % | Amount n.a. |
|---|---|---|---|---|---|---|---|
| 1 | | 0,380 | 100,204 | 2160,671 | 99,96 | 99,99 | n.a. |
| 2 | | 1,650 | 0,044 | 0,303 | 0,04 | 0,01 | n.a. |
| **Total:** | | | **100,249** | **2160,974** | **100,00** | **100,00** | |

#### Chromatogram and Results

##### Injection Details

| | | | |
|---|---|---|---|
| *Injection Name:* | **A1344** | *Run Time (min):* | **6,00** |
| *Vial Number:* | **GB2** | *Injection Volume:* | **5,00** |
| *Injection Type:* | **Unknown** | *Channel:* | **UV_VIS_1** |
| *Calibration Level:* | | *Wavelength:* | **220** |
| *Instrument Method:* | **Kinetics method 8 per h 0.4 mL** | *Bandwidth:* | **8** |
| *Processing Method:* | **Quantitative** | *Dilution Factor:* | **1,0000** |
| *Injection Date/Time:* | **13.sep..23 09:56** | *Sample Weight:* | **1,0000** |

##### Chromatogram

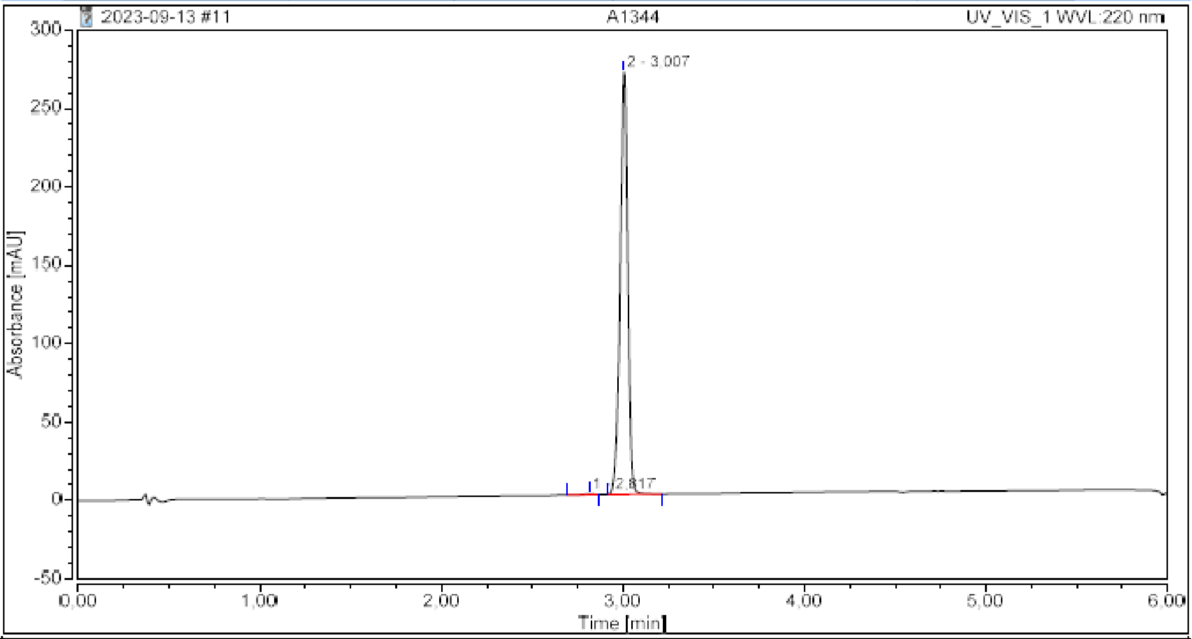

##### Integration Results

| No. | Peak Name | Retention Time min | Area mAU*min | Height mAU | Relative Area % | Relative Height % | Amount n.a. |
|---|---|---|---|---|---|---|---|
| 1 | | 2,817 | 0,031 | 0,454 | 0,23 | 0,17 | n.a. |
| 2 | | 3,007 | 13,189 | 269,407 | 99,77 | 99,83 | n.a. |
| **Total:** | | | **13,220** | **269,862** | **100,00** | **100,00** | |

**P1**

#### Chromatogram and Results

##### Injection Details

| | | | |
|---|---|---|---|
| *Injection Name:* | **A1034E fr19** | *Run Time (min):* | **6,00** |
| *Vial Number:* | **RE3** | *Injection Volume:* | **5,00** |
| *Injection Type:* | **Unknown** | *Channel:* | **UV_VIS_2** |
| *Calibration Level:* | | *Wavelength:* | **254** |
| *Instrument Method:* | **Kinetics method 8 per h (20-100)** | *Bandwidth:* | **16** |
| *Processing Method:* | **Quantitative** | *Dilution Factor:* | **1,0000** |
| *Injection Date/Time:* | **02.jul..21 12:16** | *Sample Weight:* | **1,0000** |

##### Chromatogram

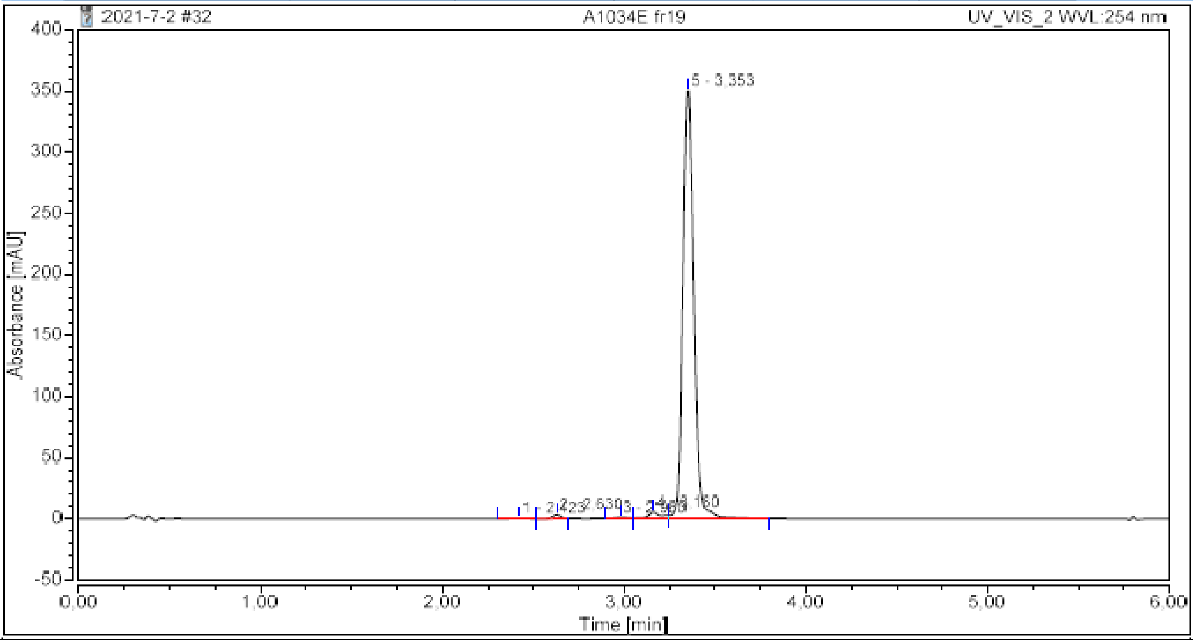

##### Integration Results

| No. | Peak Name | Retention Time min | Area mAU*min | Height mAU | Relative Area % | Relative Height % | Amount n.a. |
|---|---|---|---|---|---|---|---|
| 1 | | 2,423 | 0,025 | 0,261 | 0,10 | 0,07 | n.a. |
| 2 | | 2,630 | 0,158 | 3,356 | 0,64 | 0,93 | n.a. |
| 3 | | 2,980 | 0,081 | 0,997 | 0,33 | 0,28 | n.a. |
| 4 | | 3,160 | 0,424 | 5,550 | 1,72 | 1,54 | n.a. |
| 5 | | 3,353 | 23,976 | 349,971 | 97,21 | 97,18 | n.a. |
| **Total:** | | | **24,664** | **360,134** | **100,00** | **100,00** | |

**P2**

#### Chromatogram and Results

##### Injection Details

| | | | |
|---|---|---|---|
| *Injection Name:* | **A1034F 5** | *Run Time (min):* | **6,00** |
| *Vial Number:* | **RB5** | *Injection Volume:* | **5,00** |
| *Injection Type:* | **Unknown** | *Channel:* | **UV_VIS_2** |
| *Calibration Level:* | | *Wavelength:* | **254** |
| *Instrument Method:* | **Kinetics method 8 per h (20-100)** | *Bandwidth:* | **16** |
| *Processing Method:* | **Quantitative** | *Dilution Factor:* | **1,0000** |
| *Injection Date/Time:* | **02.jul..21 10:10** | *Sample Weight:* | **1,0000** |

##### Chromatogram

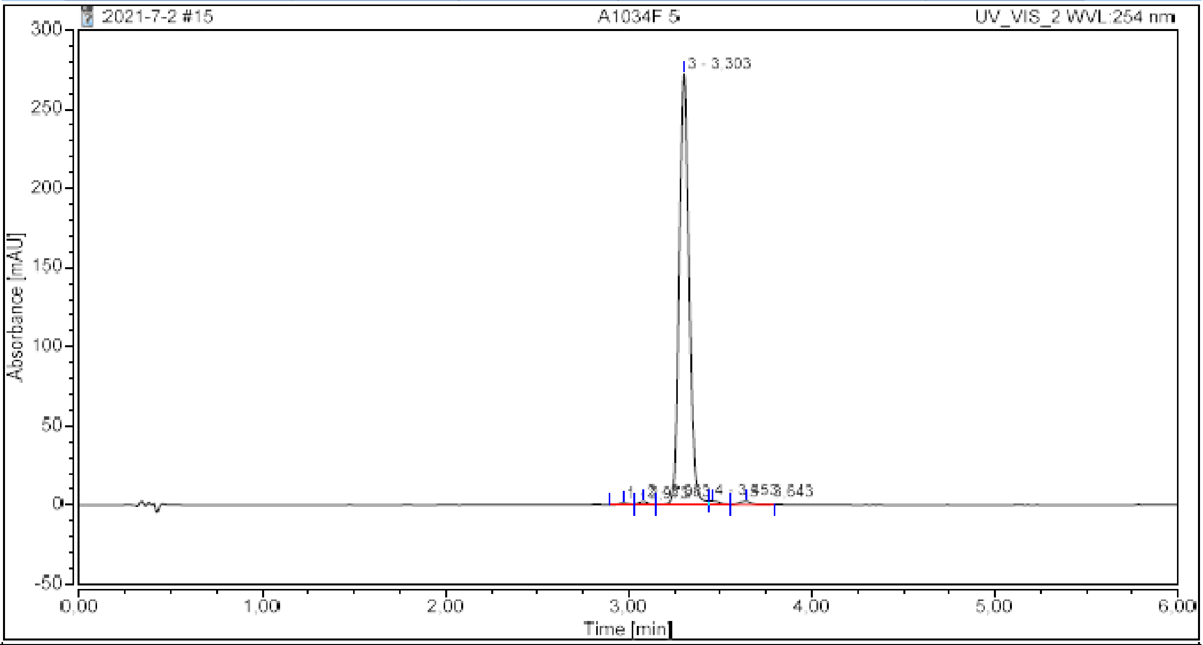

##### Integration Results

| No. | Peak Name | Retention Time min | Area mAU*min | Height mAU | Relative Area % | Relative Height % | Amount n.a. |
|---|---|---|---|---|---|---|---|
| 1 | | 2,973 | 0,071 | 1,142 | 0,40 | 0,41 | n.a. |
| 2 | | 3,083 | 0,116 | 2,131 | 0,66 | 0,76 | n.a. |
| 3 | | 3,303 | 17,107 | 272,135 | 97,04 | 97,24 | n.a. |
| 4 | | 3,453 | 0,168 | 2,351 | 0,96 | 0,84 | n.a. |
| 5 | | 3,643 | 0,166 | 2,102 | 0,94 | 0,75 | n.a. |
| **Total:** | | | **17,628** | **279,862** | **100,00** | **100,00** | |

**P3**

#### Chromatogram and Results

**Injection Details**

| | | | |
|---|---|---|---|
| *Injection Name:* | **A1177 fr5** | *Run Time (min):* | **6,00** |
| *Vial Number:* | **BA5** | *Injection Volume:* | **5,00** |
| *Injection Type:* | **Unknown** | *Channel:* | **UV_VIS_2** |
| *Calibration Level:* | | *Wavelength:* | **254** |
| *Instrument Method:* | **Kinetics method 8 per h 0.4 mL** | *Bandwidth:* | **16** |
| *Processing Method:* | **Quantitative** | *Dilution Factor:* | **1,0000** |
| *Injection Date/Time:* | **27.dec..21 16:37** | *Sample Weight:* | **1,0000** |

**Chromatogram**

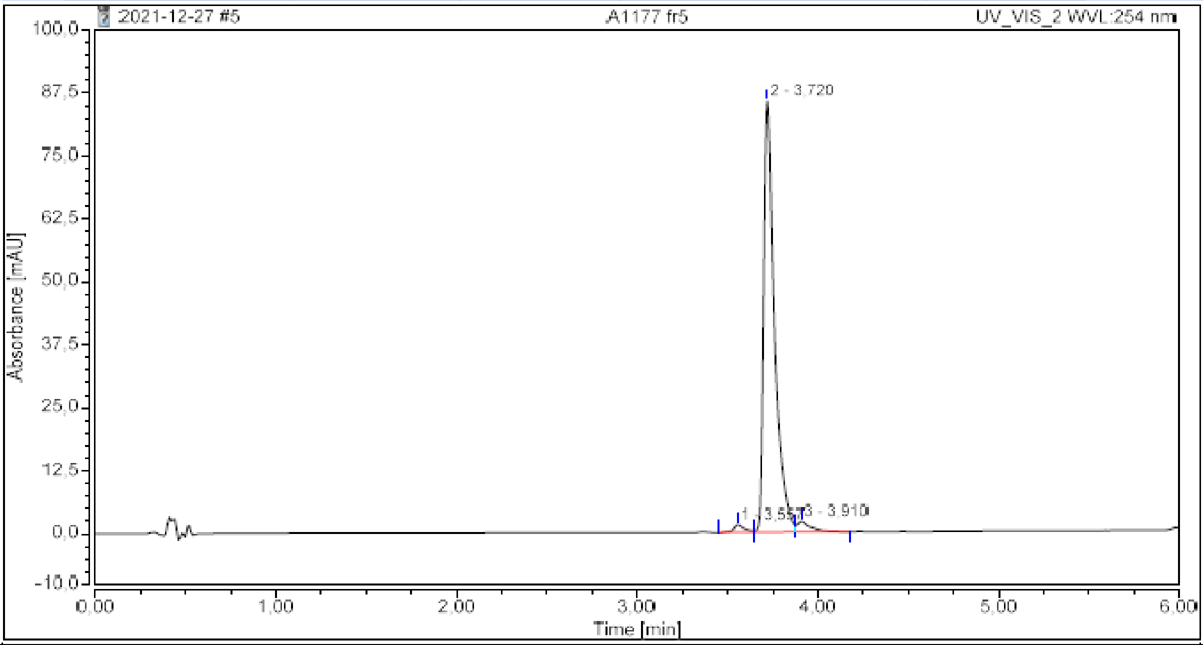

**Integration Results**

| No. | Peak Name | Retention Time<br>min | Area<br>mAU*min | Height<br>mAU | Relative Area<br>% | Relative Height<br>% | Amount<br>n.a. |
|---|---|---|---|---|---|---|---|
| 1 | | 3,557 | 0,102 | 1,470 | 1,65 | 1,65 | n.a. |
| 2 | | 3,720 | 5,926 | 85,546 | 95,48 | 96,00 | n.a. |
| 3 | | 3,910 | 0,178 | 2,096 | 2,87 | 2,35 | n.a. |
| **Total:** | | | **6,206** | **89,112** | **100,00** | **100,00** | |

**P4**

#### Chromatogram and Results

##### Injection Details

| | | | |
|---|---|---|---|
| *Injection Name:* | **A1058-1A** | *Run Time (min):* | **6,00** |
| *Vial Number:* | **RA2** | *Injection Volume:* | **5,00** |
| *Injection Type:* | **Unknown** | *Channel:* | **UV_VIS_4** |
| *Calibration Level:* | | *Wavelength:* | **210** |
| *Instrument Method:* | **Kinetics method 8 per h 0.4 mL** | *Bandwidth:* | **8** |
| *Processing Method:* | **Quantitative(1)** | *Dilution Factor:* | **1,0000** |
| *Injection Date/Time:* | **28.jul..21 07:48** | *Sample Weight:* | **1,0000** |

##### Chromatogram

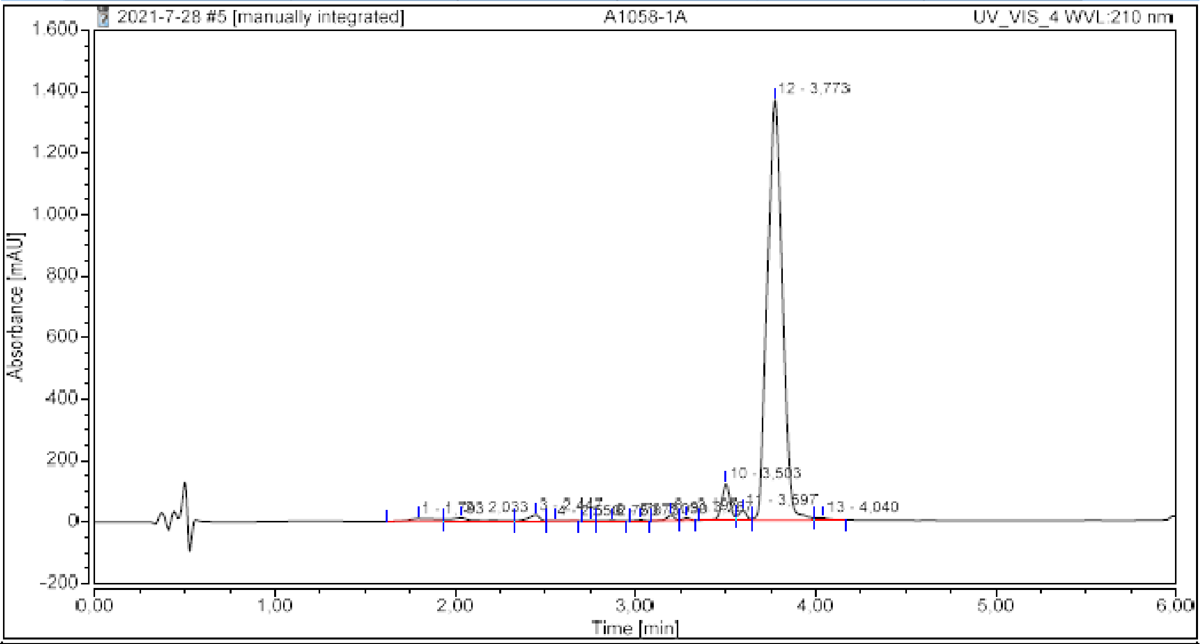

##### Integration Results

| No. | Peak Name | Retention Time min | Area mAU*min | Height mAU | Relative Area % | Relative Height % | Amount n.a. |
|---|---|---|---|---|---|---|---|
| 1 | | 1,793 | 1,452 | 8,225 | 0,99 | 0,51 | n.a. |
| 2 | | 2,033 | 1,651 | 10,731 | 1,12 | 0,67 | n.a. |
| 3 | | 2,447 | 1,410 | 19,551 | 0,96 | 1,22 | n.a. |
| 4 | | 2,550 | 0,110 | 1,167 | 0,08 | 0,07 | n.a. |
| 5 | | 2,750 | 0,023 | 0,615 | 0,02 | 0,04 | n.a. |
| 6 | | 2,873 | 0,126 | 1,669 | 0,09 | 0,10 | n.a. |
| 7 | | 3,030 | 0,138 | 2,926 | 0,09 | 0,18 | n.a. |
| 8 | | 3,197 | 0,830 | 18,318 | 0,56 | 1,15 | n.a. |
| 9 | | 3,287 | 0,362 | 8,539 | 0,25 | 0,53 | n.a. |
| 10 | | 3,503 | 6,275 | 119,561 | 4,27 | 7,48 | n.a. |
| 11 | | 3,597 | 1,650 | 31,244 | 1,12 | 1,95 | n.a. |
| 12 | | 3,773 | 132,280 | 1368,234 | 90,06 | 85,61 | n.a. |
| 13 | | 4,040 | 0,575 | 7,478 | 0,39 | 0,47 | n.a. |
| **Total:** | | | **146,880** | **1598,258** | **100,00** | **100,00** | |

**P5**

#### Chromatogram and Results

##### Injection Details

| | | | |
|---|---|---|---|
| *Injection Name:* | **A1178 fr14** | *Run Time (min):* | **6,00** |
| *Vial Number:* | **BB7** | *Injection Volume:* | **5,00** |
| *Injection Type:* | **Unknown** | *Channel:* | **UV_VIS_2** |
| *Calibration Level:* | | *Wavelength:* | **254** |
| *Instrument Method:* | **Kinetics method 8 per h 0.4 mL** | *Bandwidth:* | **16** |
| *Processing Method:* | **Quantitative** | *Dilution Factor:* | **1,0000** |
| *Injection Date/Time:* | **27.dec..21 18:06** | *Sample Weight:* | **1,0000** |

##### Chromatogram

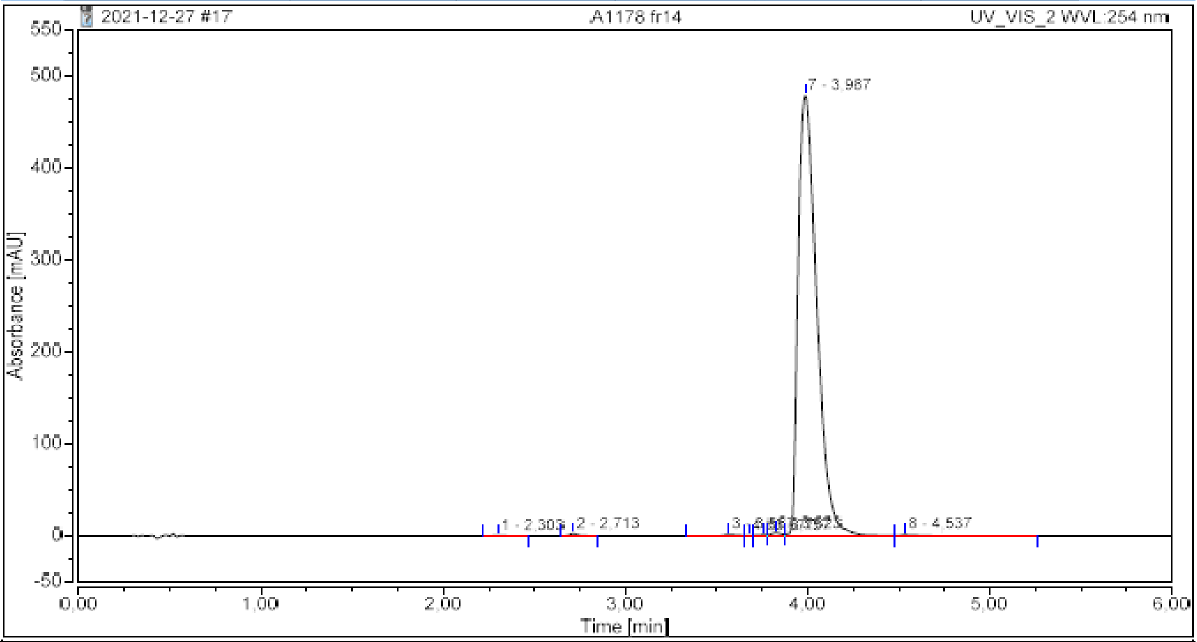

##### Integration Results

| No. | Peak Name | Retention Time<br>min | Area<br>mAU*min | Height<br>mAU | Relative Area<br>% | Relative Height<br>% | Amount<br>n.a. |
|---|---|---|---|---|---|---|---|
| 1 | | 2,303 | 0,029 | 0,382 | 0,05 | 0,08 | n.a. |
| 2 | | 2,713 | 0,111 | 2,076 | 0,19 | 0,42 | n.a. |
| 3 | | 3,567 | 0,129 | 1,595 | 0,22 | 0,33 | n.a. |
| 4 | | 3,677 | 0,022 | 0,495 | 0,04 | 0,10 | n.a. |
| 5 | | 3,757 | 0,073 | 1,190 | 0,13 | 0,24 | n.a. |
| 6 | | 3,823 | 0,220 | 3,283 | 0,38 | 0,67 | n.a. |
| 7 | | 3,987 | 56,652 | 479,166 | 98,73 | 97,91 | n.a. |
| 8 | | 4,537 | 0,142 | 1,217 | 0,25 | 0,25 | n.a. |
| **Total:** | | | **57,379** | **489,405** | **100,00** | **100,00** | |

**B2**

#### Chromatogram and Results

**Injection Details**

| | | | |
|---|---|---|---|
| *Injection Name:* | **A1237-1 fr1** | *Run Time (min):* | **6,00** |
| *Vial Number:* | **BB8** | *Injection Volume:* | **5,00** |
| *Injection Type:* | **Unknown** | *Channel:* | **UV_VIS_1** |
| *Calibration Level:* | | *Wavelength:* | **220** |
| *Instrument Method:* | **Kinetics method 8 per h 0.4 mL** | *Bandwidth:* | **8** |
| *Processing Method:* | **Quantitative** | *Dilution Factor:* | **1,0000** |
| *Injection Date/Time:* | **28.feb..22 11:18** | *Sample Weight:* | **1,0000** |

**Chromatogram**

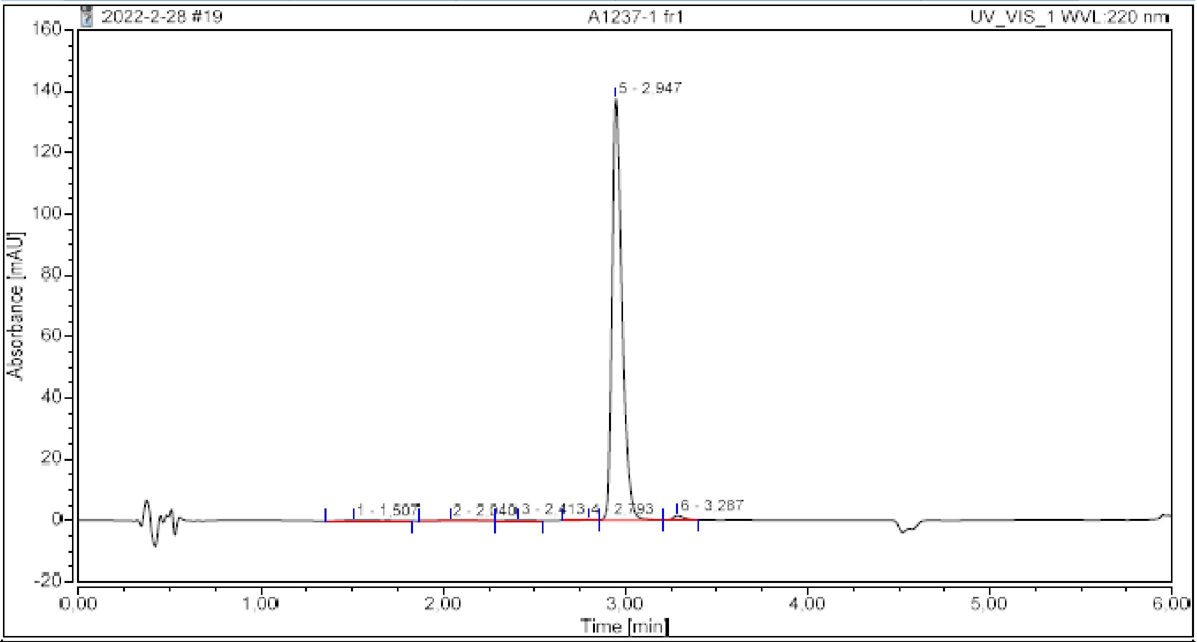

**Integration Results**

| No. | Peak Name | Retention Time min | Area mAU*min | Height mAU | Relative Area % | Relative Height % | Amount n.a. |
|---|---|---|---|---|---|---|---|
| 1 | | 1,507 | 0,049 | 0,214 | 0,54 | 0,15 | n.a. |
| 2 | | 2,040 | 0,033 | 0,175 | 0,36 | 0,12 | n.a. |
| 3 | | 2,413 | 0,024 | 0,186 | 0,26 | 0,13 | n.a. |
| 4 | | 2,793 | 0,026 | 0,319 | 0,29 | 0,23 | n.a. |
| 5 | | 2,947 | 8,837 | 137,529 | 97,52 | 98,35 | n.a. |
| 6 | | 3,287 | 0,094 | 1,406 | 1,03 | 1,01 | n.a. |
| **Total:** | | | **9,062** | **139,829** | **100,00** | **100,00** | |

**B3**

#### Chromatogram and Results

**Injection Details**

| | | | |
|---|---|---|---|
| *Injection Name:* | **A1241B-1 fr1** | *Run Time (min):* | **6,00** |
| *Vial Number:* | **BB2** | *Injection Volume:* | **5,00** |
| *Injection Type:* | **Unknown** | *Channel:* | **UV_VIS_1** |
| *Calibration Level:* | | *Wavelength:* | **220** |
| *Instrument Method:* | **Kinetics method 8 per h 0.4 mL** | *Bandwidth:* | **8** |
| *Processing Method:* | **Quantitative** | *Dilution Factor:* | **1,0000** |
| *Injection Date/Time:* | **28.feb..22 10:11** | *Sample Weight:* | **1,0000** |

**Chromatogram**

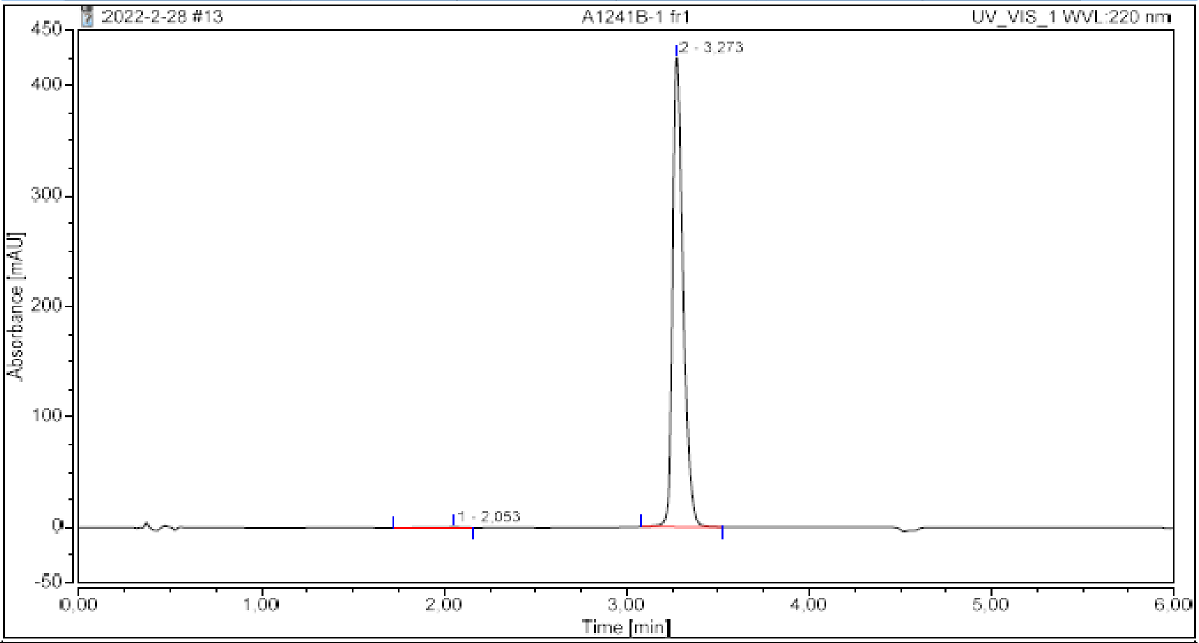

**Integration Results**

| No. | Peak Name | Retention Time min | Area mAU*min | Height mAU | Relative Area % | Relative Height % | Amount n.a. |
|---|---|---|---|---|---|---|---|
| 1 | | 2,053 | 0,072 | 0,406 | 0,25 | 0,10 | n.a. |
| 2 | | 3,273 | 28,690 | 424,674 | 99,75 | 99,90 | n.a. |
| **Total:** | | | **28,762** | **425,080** | **100,00** | **100,00** | |

### Computational studies – atom coordinates of the QM–optimized species (in Å)

#### methoxide

| atom | x | y | z |
|---|---|---|---|
| C1 | -0.0000035865 | -0.0000178545 | 0.6429286411 |
| O2 | 0.0000097628 | 0.0000175994 | -0.6950385718 |
| H3 | 1.0193245162 | -0.0000415486 | 1.1251517002 |
| H4 | -0.5097348028 | 0.8826924892 | 1.1251823975 |
| H5 | -0.5097009521 | -0.8827176654 | 1.1252036312 |

## 1a

| atom | x | y | z |
|---|---|---|---|
| O1 | 0.1043080000 | 0.1190330000 | 0.5095250000 |
| C2 | 1.2679030000 | 0.0106570000 | 0.1331230000 |
| N3 | 2.0774500000 | 1.0223670000 | -0.2241930000 |
| C4 | 3.4132870000 | 0.8447610000 | -0.7901170000 |
| C5 | 1.5452950000 | 2.3770680000 | -0.2663130000 |
| O6 | 1.9030030000 | -1.1965770000 | 0.0353120000 |
| C7 | 1.2349810000 | -2.3504000000 | 0.4211620000 |
| C8 | 1.0318160000 | -3.3331300000 | -0.5538120000 |
| N9 | 1.3178920000 | -3.3377860000 | -1.8973000000 |
| C10 | 0.9376620000 | -4.5535180000 | -2.4212090000 |
| C11 | 0.4080300000 | -5.3464340000 | -1.4346500000 |
| C12 | 0.4561550000 | -4.5843690000 | -0.2165590000 |
| C13 | 0.0792650000 | -4.8332210000 | 1.1188500000 |
| C14 | 0.2870240000 | -3.8456210000 | 2.0695640000 |
| C15 | 0.8696200000 | -2.6046590000 | 1.7283000000 |
| H16 | 3.3961160000 | 1.0412000000 | -1.8694980000 |
| H17 | 3.7721390000 | -0.1649310000 | -0.6124470000 |
| H18 | 4.0961160000 | 1.5552970000 | -0.3135410000 |
| H19 | 1.2984110000 | 2.6631200000 | -1.2960380000 |
| H20 | 0.6497680000 | 2.4392780000 | 0.3501780000 |
| H21 | 2.3016430000 | 3.0650490000 | 0.1217670000 |
| H22 | 1.7359630000 | -2.5792390000 | -2.4275230000 |
| H23 | 1.0803590000 | -4.7582850000 | -3.4737480000 |
| H24 | 0.0299770000 | -6.3528020000 | -1.5561940000 |
| H25 | -0.3623870000 | -5.7862500000 | 1.3995240000 |
| H26 | 0.0066730000 | -4.0229870000 | 3.1039010000 |
| H27 | 1.0379830000 | -1.8454410000 | 2.4872490000 |

## 2a

| atom | x | y | z |
|---|---|---|---|
| O1 | 0.2797290000 | -0.1822640000 | 0.1029920000 |
| C2 | 1.4660600000 | 0.0277900000 | -0.1347380000 |
| N3 | 1.9894150000 | 1.2046970000 | -0.5155440000 |
| C4 | 1.1087250000 | 2.3454640000 | -0.7628650000 |
| C5 | 3.4080300000 | 1.4030250000 | -0.8112410000 |
| S6 | 2.7058740000 | -1.3061590000 | 0.0050900000 |
| C7 | 1.6766230000 | -2.6518740000 | 0.5218240000 |
| C8 | 0.9279580000 | -3.3787850000 | -0.4177380000 |
| N9 | 0.8026020000 | -3.2228270000 | -1.7740590000 |
| C10 | -0.0450500000 | -4.1950360000 | -2.2602840000 |
| C11 | -0.4798360000 | -4.9921380000 | -1.2331990000 |
| C12 | 0.1360300000 | -4.4935920000 | -0.0332710000 |
| C13 | 0.1145600000 | -4.8870340000 | 1.3175580000 |
| C14 | 0.8667200000 | -4.1752610000 | 2.2416880000 |

| | | | |
|---|---|---|---|
| C15 | 1.6399000000 | -3.0630770000 | 1.8481640000 |
| H16 | 0.0758920000 | 2.0559360000 | -0.5794020000 |
| H17 | 1.3819780000 | 3.1701520000 | -0.0972090000 |
| H18 | 1.2176720000 | 2.6720330000 | -1.8023270000 |
| H19 | 3.6640760000 | 1.0273680000 | -1.8085540000 |
| H20 | 3.6117850000 | 2.4752490000 | -0.7782850000 |
| H21 | 4.0382230000 | 0.9214980000 | -0.0595720000 |
| H22 | 1.2790620000 | -2.5312130000 | -2.3444270000 |
| H23 | -0.2698690000 | -4.2372440000 | -3.3179160000 |
| H24 | -1.1528710000 | -5.8346960000 | -1.3172390000 |
| H25 | -0.4788990000 | -5.7430410000 | 1.6305740000 |
| H26 | 0.8646430000 | -4.4718740000 | 3.2865690000 |
| H27 | 2.2229200000 | -2.5182180000 | 2.5856970000 |

**3a**

| atom | x | y | z |
|---|---|---|---|
| O1 | 0.3635120000 | 0.2383890000 | 0.9094000000 |
| C2 | 1.4198560000 | 0.1005230000 | 0.2975880000 |
| N3 | 2.1226710000 | 1.1026440000 | -0.2400600000 |
| C4 | 3.4970120000 | 1.0360610000 | -0.7308330000 |
| C5 | 1.5378960000 | 2.4460550000 | -0.1929390000 |
| Se6 | 2.0931930000 | -1.7846410000 | 0.1086590000 |
| C7 | 2.5103720000 | -1.8298360000 | -1.7875660000 |
| C8 | 1.4790800000 | -1.7660010000 | -2.7367450000 |
| N9 | 0.1206150000 | -1.6476570000 | -2.5769630000 |
| C10 | -0.4804180000 | -1.6423340000 | -3.8179190000 |
| C11 | 0.4732600000 | -1.7370600000 | -4.7977990000 |
| C12 | 1.7453810000 | -1.8232900000 | -4.1319690000 |
| C13 | 3.0741100000 | -1.9701710000 | -4.5702850000 |
| C14 | 4.0884440000 | -2.0622890000 | -3.6275720000 |
| C15 | 3.8105140000 | -1.9894940000 | -2.2460280000 |
| H16 | 3.9838010000 | 0.1293710000 | -0.3782830000 |
| H17 | 4.0423400000 | 1.8983990000 | -0.3361090000 |
| H18 | 3.5216970000 | 1.0649990000 | -1.8251720000 |
| H19 | 0.4510740000 | 2.3776800000 | -0.2410410000 |
| H20 | 1.9095230000 | 3.0104930000 | -1.0513090000 |
| H21 | 1.8282450000 | 2.9589660000 | 0.7320880000 |
| H22 | -0.3779380000 | -1.6721240000 | -1.6941380000 |
| H23 | -1.5571590000 | -1.5674390000 | -3.8971670000 |
| H24 | 0.2971940000 | -1.7473040000 | -5.8654890000 |
| H25 | 3.2995420000 | -2.0211640000 | -5.6332290000 |
| H26 | 5.1186140000 | -2.1875320000 | -3.9484960000 |
| H27 | 4.6268280000 | -2.0570120000 | -1.5307390000 |

**4a**

| atom | x | y | z |
|---|---|---|---|
| C1 | -0.0715960000 | 0.7513550000 | 0.8634400000 |
| N2 | -0.3431280000 | -0.5233540000 | 0.5651370000 |
| C3 | 0.7657930000 | -1.4783580000 | 0.4915540000 |
| C4 | -1.6775850000 | -1.0995070000 | 0.4361140000 |
| Te5 | -1.5428640000 | 2.3556530000 | 0.7921130000 |
| C6 | -3.0524670000 | 1.6246950000 | -0.4980570000 |
| C7 | -4.3201300000 | 1.2747610000 | -0.0020210000 |
| N8 | -4.8131010000 | 1.2873630000 | 1.2803710000 |
| C9 | -6.1190340000 | 0.8506510000 | 1.2686040000 |
| C10 | -6.4946240000 | 0.5391250000 | -0.0129850000 |
| C11 | -5.3568760000 | 0.7991770000 | -0.8519360000 |
| C12 | -5.1071210000 | 0.6690520000 | -2.2305600000 |
| C13 | -3.8584440000 | 1.0171920000 | -2.7242760000 |

| | | | |
|---|---|---|---|
| C14 | -2.8447660000 | 1.4976600000 | -1.8672530000 |
| O15 | 1.0493250000 | 1.1624740000 | 1.1828360000 |
| H16 | 1.6906910000 | -0.9502910000 | 0.2628980000 |
| H17 | 0.8753200000 | -2.0107250000 | 1.4430520000 |
| H18 | 0.5477050000 | -2.1974240000 | -0.3022170000 |
| H19 | -2.4163410000 | -0.4666200000 | 0.9269720000 |
| H20 | -1.9506960000 | -1.2358330000 | -0.6159290000 |
| H21 | -1.6761610000 | -2.0731640000 | 0.9346160000 |
| H22 | -4.3145500000 | 1.5985590000 | 2.1073870000 |
| H23 | -6.6817670000 | 0.7987240000 | 2.1920550000 |
| H24 | -7.4620010000 | 0.1689590000 | -0.3263560000 |
| H25 | -5.8835140000 | 0.3035960000 | -2.8992670000 |
| H26 | -3.6503290000 | 0.9278000000 | -3.7868370000 |
| H27 | -1.8813300000 | 1.7724710000 | -2.2886580000 |

## 5a

| atom | x | y | z |
|---|---|---|---|
| S1 | 3.7393560000 | -1.2454720000 | 0.4418580000 |
| C2 | 2.4507110000 | -0.1592400000 | 0.4687550000 |
| N3 | 2.5324970000 | 1.1372200000 | 0.2062190000 |
| C4 | 1.3395750000 | 1.9921090000 | 0.2999010000 |
| C5 | 3.8273140000 | 1.7338610000 | -0.1246550000 |
| O6 | 1.1804780000 | -0.5182550000 | 0.7690570000 |
| C7 | 0.8803130000 | -1.8369340000 | 1.1023310000 |
| C8 | 0.5358750000 | -2.7421920000 | 0.0935850000 |
| N9 | 0.4952130000 | -2.6065150000 | -1.2719960000 |
| C10 | 0.0955900000 | -3.8069040000 | -1.8203500000 |
| C11 | -0.1278460000 | -4.7257520000 | -0.8269810000 |
| C12 | 0.1480320000 | -4.0668830000 | 0.4211530000 |
| C13 | 0.1027710000 | -4.4631520000 | 1.7733770000 |
| C14 | 0.4393630000 | -3.5449880000 | 2.7559930000 |
| C15 | 0.8268110000 | -2.2277820000 | 2.4239880000 |
| H16 | 0.9854960000 | 2.0375210000 | 1.3337180000 |
| H17 | 0.5416870000 | 1.5961090000 | -0.3313000000 |
| H18 | 1.6095430000 | 2.9908590000 | -0.0385560000 |
| H19 | 4.2621250000 | 1.2260830000 | -0.9885700000 |
| H20 | 3.6740990000 | 2.7870120000 | -0.3556290000 |
| H21 | 4.5156720000 | 1.6373720000 | 0.7197570000 |
| H22 | 0.7162620000 | -1.7721800000 | -1.8072090000 |
| H23 | 0.0003070000 | -3.9081960000 | -2.8933470000 |
| H24 | -0.4511080000 | -5.7495700000 | -0.9619270000 |
| H25 | -0.1960790000 | -5.4737450000 | 2.0414360000 |
| H26 | 0.4060870000 | -3.8329990000 | 3.8028480000 |
| H27 | 1.0904550000 | -1.5118080000 | 3.1977600000 |

## 6a

| atom | x | y | z |
|---|---|---|---|
| C1 | 0.0883180000 | -0.0533520000 | 0.9133850000 |
| N2 | 1.3739970000 | 0.0758170000 | 0.2115090000 |
| C3 | 2.0167660000 | 1.3575760000 | 0.1533510000 |
| C4 | 2.4761060000 | 1.9727010000 | 1.3071080000 |
| C5 | 3.0781030000 | 3.2475040000 | 1.2540310000 |
| C6 | 3.2396640000 | 3.9125290000 | 0.0475790000 |
| C7 | 2.7808580000 | 3.3041490000 | -1.1362420000 |
| C8 | 2.1570530000 | 2.0308400000 | -1.0693490000 |
| N9 | 1.7964850000 | 1.6680480000 | -2.3434080000 |
| C10 | 2.1723130000 | 2.6705440000 | -3.2126600000 |
| C11 | 2.7793440000 | 3.6869760000 | -2.5219630000 |
| C12 | 1.9850660000 | -1.0436660000 | -0.1927940000 |

| | | | |
|---|---|---|---|
| F13 | 3.1810430000 | -0.7878680000 | -0.7685690000 |
| O14 | 1.5948010000 | -2.1852560000 | -0.1123500000 |
| H15 | 0.2461550000 | -0.2895180000 | 1.9710060000 |
| H16 | -0.4419200000 | 0.8953870000 | 0.8220620000 |
| H17 | -0.4969490000 | -0.8461470000 | 0.4466670000 |
| H18 | 2.3720620000 | 1.4557080000 | 2.2580960000 |
| H19 | 3.4273430000 | 3.7026180000 | 2.1762450000 |
| H20 | 3.7132940000 | 4.8906190000 | 0.0154210000 |
| H21 | 1.2851030000 | 0.8336850000 | -2.6148600000 |
| H22 | 1.9699130000 | 2.5737490000 | -4.2712270000 |
| H23 | 3.1782140000 | 4.6005380000 | -2.9432740000 |

**7a**

| atom | x | y | z |
|---|---|---|---|
| O1 | 0.0624150000 | -0.0553170000 | 0.0627370000 |
| C2 | 1.2870210000 | -0.0042980000 | 0.0201630000 |
| O3 | 1.9627680000 | 1.1542560000 | -0.1356020000 |
| C4 | 1.1695750000 | 2.3293890000 | -0.2017680000 |
| C5 | 2.1610580000 | 3.4712010000 | -0.1761150000 |
| F6 | 2.8893190000 | 3.4878330000 | 0.9519980000 |
| F7 | 3.0272750000 | 3.4190570000 | -1.2016230000 |
| F8 | 1.5144180000 | 4.6466280000 | -0.2559110000 |
| N9 | 2.1127150000 | -1.0731800000 | 0.0963170000 |
| C10 | 1.5235500000 | -2.3980300000 | 0.3170660000 |
| C11 | 3.5336950000 | -0.9534950000 | 0.2408280000 |
| C12 | 4.3735060000 | -1.3715760000 | -0.8031020000 |
| N13 | 4.0724230000 | -1.8516720000 | -2.0545370000 |
| C14 | 5.2488280000 | -2.1452550000 | -2.7117820000 |
| C15 | 6.3196170000 | -1.8678780000 | -1.9019260000 |
| C16 | 5.7857890000 | -1.3650490000 | -0.6648710000 |
| C17 | 6.3578700000 | -0.9087030000 | 0.5379290000 |
| C18 | 5.5230850000 | -0.4814460000 | 1.5599380000 |
| C19 | 4.1195420000 | -0.5129950000 | 1.4169550000 |
| H20 | 0.5036560000 | 2.4208530000 | 0.6620760000 |
| H21 | 0.5939280000 | 2.3776960000 | -1.1322050000 |
| H22 | 2.2270500000 | -3.1506540000 | -0.0450360000 |
| H23 | 1.3349780000 | -2.5670020000 | 1.3831670000 |
| H24 | 0.5880190000 | -2.4814840000 | -0.2361200000 |
| H25 | 3.1463560000 | -1.9308110000 | -2.4636120000 |
| H26 | 5.2245920000 | -2.5350260000 | -3.7208330000 |
| H27 | 7.3636680000 | -2.0034910000 | -2.1514250000 |
| H28 | 7.4379740000 | -0.8941360000 | 0.6634430000 |
| H29 | 5.9487730000 | -0.1268100000 | 2.4942670000 |
| H30 | 3.4801630000 | -0.1907010000 | 2.2348950000 |

**8a**

| atom | x | y | z |
|---|---|---|---|
| C1 | 0.0038550000 | 0.1000830000 | 0.1203450000 |
| N2 | 1.4657850000 | 0.1164060000 | 0.2589330000 |
| C3 | 2.1235580000 | 1.3885170000 | 0.1715310000 |
| C4 | 2.4545160000 | 2.1231250000 | 1.2986260000 |
| C5 | 3.0378220000 | 3.4023670000 | 1.1812040000 |
| C6 | 3.2795440000 | 3.9684930000 | -0.0616410000 |
| C7 | 2.9455800000 | 3.2423950000 | -1.2208730000 |
| C8 | 2.3757900000 | 1.9491810000 | -1.0898370000 |
| N9 | 2.1466330000 | 1.4567130000 | -2.3513860000 |
| C10 | 2.5565820000 | 2.3958130000 | -3.2745950000 |
| C11 | 3.0481090000 | 3.5020700000 | -2.6314530000 |
| C12 | 2.0914450000 | -1.0128070000 | 0.6397780000 |

| | | | |
|---|---|---|---|
| O13 | 3.4407510000 | -0.8096850000 | 0.7853070000 |
| C14 | 4.1987880000 | -1.8830620000 | 1.2778270000 |
| C15 | 4.3817730000 | -1.6873340000 | 2.7874310000 |
| F16 | 5.0939880000 | -0.5915040000 | 3.0772060000 |
| F17 | 4.9966660000 | -2.7355680000 | 3.3437640000 |
| F18 | 3.1809800000 | -1.5515330000 | 3.3644800000 |
| C19 | 5.5225850000 | -1.8480040000 | 0.5138160000 |
| F20 | 5.3084830000 | -2.1316580000 | -0.7753350000 |
| F21 | 6.1134950000 | -0.6499000000 | 0.5698840000 |
| F22 | 6.3792100000 | -2.7520830000 | 1.0045280000 |
| O23 | 1.5691690000 | -2.1002100000 | 0.8272050000 |
| H24 | -0.4716960000 | 0.3349440000 | 1.0788350000 |
| H25 | -0.2799420000 | 0.8541170000 | -0.6168010000 |
| H26 | -0.3202150000 | -0.8829120000 | -0.2206620000 |
| H27 | 2.2570310000 | 1.7005400000 | 2.2805800000 |
| H28 | 3.2893000000 | 3.9508300000 | 2.0844070000 |
| H29 | 3.7184920000 | 4.9601200000 | -0.1406770000 |
| H30 | 1.7791310000 | 0.5395440000 | -2.5867610000 |
| H31 | 2.4611480000 | 2.1926030000 | -4.3330030000 |
| H32 | 3.4366620000 | 4.3966120000 | -3.1000080000 |
| H33 | 3.7224700000 | -2.8579760000 | 1.1109110000 |

**9a**

| atom | x | y | z |
|---|---|---|---|
| C1 | -0.2582660000 | 0.0278170000 | -0.4483480000 |
| N2 | 1.1944950000 | 0.0770840000 | -0.6389920000 |
| C3 | 1.9015800000 | 1.2945170000 | -0.3568330000 |
| C4 | 2.0989720000 | 2.2669340000 | -1.3218450000 |
| C5 | 2.8738710000 | 3.4112590000 | -1.0376760000 |
| C6 | 3.4670810000 | 3.5845200000 | 0.2046580000 |
| C7 | 3.2647940000 | 2.6130810000 | 1.2042420000 |
| C8 | 2.4635370000 | 1.4796430000 | 0.9137270000 |
| N9 | 2.4323330000 | 0.6812210000 | 2.0310550000 |
| C10 | 3.1867720000 | 1.2771100000 | 3.0200840000 |
| C11 | 3.7184880000 | 2.4535290000 | 2.5599510000 |
| C12 | 1.8424450000 | -1.0004710000 | -1.1164690000 |
| O13 | 3.1684410000 | -0.6618130000 | -1.3554060000 |
| C14 | 4.2143490000 | -1.5742730000 | -1.3966260000 |
| C15 | 3.9979240000 | -2.7960980000 | -2.3490380000 |
| F16 | 3.4374670000 | -3.8312080000 | -1.7290960000 |
| F17 | 5.1848770000 | -3.2139090000 | -2.8143860000 |
| F18 | 3.2494020000 | -2.4432720000 | -3.3880330000 |
| C19 | 5.3944650000 | -0.7110050000 | -1.9498830000 |
| F20 | 5.2624160000 | -0.5590910000 | -3.2710980000 |
| F21 | 5.3966600000 | 0.4950880000 | -1.3946320000 |
| F22 | 6.5723100000 | -1.2897190000 | -1.7015970000 |
| C23 | 4.5492420000 | -2.0521600000 | 0.0430790000 |
| F24 | 5.0856220000 | -1.0563350000 | 0.7484230000 |
| F25 | 3.4380870000 | -2.4425730000 | 0.6671050000 |
| F26 | 5.4108670000 | -3.0712490000 | 0.0244740000 |
| O27 | 1.3894410000 | -2.1022740000 | -1.3457990000 |
| H28 | -0.7719390000 | 0.4525830000 | -1.3167110000 |
| H29 | -0.5087770000 | 0.6081890000 | 0.4422010000 |
| H30 | -0.5674890000 | -1.0080940000 | -0.3080290000 |
| H31 | 1.6784470000 | 2.1178170000 | -2.3134280000 |
| H32 | 3.0237770000 | 4.1523070000 | -1.8176630000 |
| H33 | 4.0840530000 | 4.4576800000 | 0.4031950000 |
| H34 | 1.8706070000 | -0.1534200000 | 2.1631700000 |
| H35 | 3.2867110000 | 0.8002720000 | 3.9864000000 |
| H36 | 4.3612280000 | 3.1254070000 | 3.1130580000 |

**10a**

| atom | x | y | z |
|---|---|---|---|
| C1 | -0.2071610000 | 0.1086560000 | 0.6151770000 |
| N2 | 1.2547920000 | 0.0876970000 | 0.5519560000 |
| C3 | 1.9340770000 | 1.2119970000 | 0.2253950000 |
| O4 | 3.2714670000 | 0.9902440000 | 0.2841630000 |
| C5 | 4.1762540000 | 2.0195820000 | 0.0612980000 |
| C6 | 5.2193100000 | 1.7378680000 | -0.8145730000 |
| C7 | 6.2276010000 | 2.6846930000 | -0.9926760000 |
| C8 | 6.1803120000 | 3.8968170000 | -0.3030440000 |
| C9 | 5.1240380000 | 4.1587280000 | 0.5725110000 |
| C10 | 4.1130640000 | 3.2175340000 | 0.7680630000 |
| O11 | 1.4296830000 | 2.2774040000 | -0.1041670000 |
| C12 | 1.9323290000 | -1.0949630000 | 0.9965540000 |
| C13 | 2.3962860000 | -2.0308860000 | 0.0605190000 |
| N14 | 2.3405510000 | -2.0253360000 | -1.3108750000 |
| C15 | 2.9572000000 | -3.1635910000 | -1.7845610000 |
| C16 | 3.4166000000 | -3.9182040000 | -0.7359250000 |
| C17 | 3.0679360000 | -3.2125850000 | 0.4674860000 |
| C18 | 3.2597310000 | -3.4563460000 | 1.8413910000 |
| C19 | 2.7863320000 | -2.5331330000 | 2.7618970000 |
| C20 | 2.1276080000 | -1.3570790000 | 2.3423140000 |
| H21 | -0.5802190000 | -0.8640040000 | 0.2883870000 |
| H22 | -0.5487950000 | 0.3021170000 | 1.6376000000 |
| H23 | -0.5871920000 | 0.8852440000 | -0.0481570000 |
| H24 | 5.2380650000 | 0.7818190000 | -1.3299740000 |
| H25 | 7.0512630000 | 2.4702660000 | -1.6675060000 |
| H26 | 6.9671460000 | 4.6322640000 | -0.4409530000 |
| H27 | 5.0892110000 | 5.0979640000 | 1.1167090000 |
| H28 | 3.2978990000 | 3.4040960000 | 1.4601220000 |
| H29 | 1.9152360000 | -1.3112690000 | -1.8938280000 |
| H30 | 3.0178900000 | -3.3446750000 | -2.8494130000 |
| H31 | 3.9415610000 | -4.8613810000 | -0.8069460000 |
| H32 | 3.7731800000 | -4.3552000000 | 2.1748860000 |
| H33 | 2.9280670000 | -2.7068420000 | 3.8244540000 |
| H34 | 1.7759550000 | -0.6342830000 | 3.0742950000 |

**11a**

| atom | x | y | z |
|---|---|---|---|
| O1 | 0.0486280000 | -0.0177600000 | -0.2853300000 |
| C2 | 1.2650810000 | -0.0480640000 | -0.2282990000 |
| O3 | 2.0195280000 | 1.0628220000 | 0.0918710000 |
| C4 | 1.3692750000 | 2.2499050000 | 0.2555120000 |
| C5 | 1.4705700000 | 2.9087090000 | 1.4748460000 |
| F6 | 2.1128520000 | 2.3313900000 | 2.4885610000 |
| C7 | 0.9219130000 | 4.1713380000 | 1.6469720000 |
| F8 | 1.0322190000 | 4.7952090000 | 2.8195380000 |
| C9 | 0.2545510000 | 4.7828300000 | 0.5960480000 |
| F10 | -0.2761810000 | 5.9934560000 | 0.7576670000 |
| C11 | 0.1442160000 | 4.1342320000 | -0.6261840000 |
| F12 | -0.4910860000 | 4.7268910000 | -1.6378200000 |
| C13 | 0.7062100000 | 2.8779130000 | -0.7940440000 |
| F14 | 0.6269830000 | 2.2757340000 | -1.9798400000 |
| N15 | 2.0648770000 | -1.1104050000 | -0.4300290000 |
| C16 | 1.4837060000 | -2.3932550000 | -0.8373860000 |
| C17 | 3.4937270000 | -0.9869330000 | -0.3947690000 |
| C18 | 4.1795590000 | -1.2044040000 | 0.8086820000 |
| N19 | 3.7143770000 | -1.5334770000 | 2.0578760000 |
| C20 | 4.7864030000 | -1.6094050000 | 2.9216260000 |

| | | | |
|---|---|---|---|
| C21 | 5.9505030000 | -1.3380780000 | 2.2511570000 |
| C22 | 5.5901510000 | -1.0737670000 | 0.8846260000 |
| C23 | 6.3118120000 | -0.7307430000 | -0.2749710000 |
| C24 | 5.6259370000 | -0.5318480000 | -1.4637410000 |
| C25 | 4.2222380000 | -0.6574740000 | -1.5251590000 |
| H26 | 1.5179770000 | -2.4990690000 | -1.9268030000 |
| H27 | 2.0621830000 | -3.1966790000 | -0.3760970000 |
| H28 | 0.4501330000 | -2.4454740000 | -0.4959030000 |
| H29 | 2.7469970000 | -1.6965200000 | 2.3210100000 |
| H30 | 4.6268740000 | -1.8570660000 | 3.9622190000 |
| H31 | 6.9457920000 | -1.3242520000 | 2.6750470000 |
| H32 | 7.3930100000 | -0.6211810000 | -0.2381330000 |
| H33 | 6.1696160000 | -0.2670640000 | -2.3653020000 |
| H34 | 3.6943560000 | -0.4793610000 | -2.4591620000 |

## 12a

| atom | x | y | z |
|---|---|---|---|
| C1 | -0.3257510000 | -1.4372360000 | -4.1715740000 |
| N2 | -0.3614550000 | -1.9957510000 | -2.8089530000 |
| C3 | -1.4875330000 | -1.7647860000 | -2.1090700000 |
| O4 | -1.4443350000 | -2.3157150000 | -0.8495470000 |
| C5 | -2.4208570000 | -1.9611710000 | 0.0253200000 |
| N6 | -2.9189150000 | -2.9075340000 | 0.7746720000 |
| O7 | -3.7892110000 | -2.2662510000 | 1.6276810000 |
| C8 | -3.7931120000 | -0.9486540000 | 1.3583600000 |
| C9 | -2.9377300000 | -0.6729650000 | 0.3419730000 |
| O10 | -2.4623690000 | -1.1605330000 | -2.5225940000 |
| C11 | 0.8257090000 | -2.5937810000 | -2.2778590000 |
| C12 | 1.4672580000 | -2.1052160000 | -1.1487370000 |
| C13 | 2.6831460000 | -2.6644250000 | -0.7046800000 |
| C14 | 3.2995920000 | -3.6909470000 | -1.4033910000 |
| C15 | 2.6739450000 | -4.1977660000 | -2.5573360000 |
| C16 | 1.4253400000 | -3.6615490000 | -2.9686130000 |
| N17 | 1.0291770000 | -4.3362970000 | -4.0994740000 |
| C18 | 1.9888540000 | -5.2750950000 | -4.4170210000 |
| C19 | 3.0123150000 | -5.2239960000 | -3.5066930000 |
| H20 | -1.0811920000 | -1.9147440000 | -4.8018570000 |
| H21 | 0.6669930000 | -1.6130560000 | -4.5857430000 |
| H22 | -0.5152390000 | -0.3607580000 | -4.1336260000 |
| H23 | -4.4502360000 | -0.3473650000 | 1.9731050000 |
| H24 | -2.7074540000 | 0.2773610000 | -0.1147810000 |
| H25 | 1.0256090000 | -1.2737930000 | -0.6074360000 |
| H26 | 3.1488310000 | -2.2598360000 | 0.1896400000 |
| H27 | 4.2510050000 | -4.0973090000 | -1.0683850000 |
| H28 | 0.1488760000 | -4.2312230000 | -4.5941110000 |
| H29 | 1.8604970000 | -5.9096910000 | -5.2838110000 |
| H30 | 3.9042280000 | -5.8374070000 | -3.5080820000 |

## 13a

| atom | x | y | z |
|---|---|---|---|
| C1 | 2.8026610000 | 0.0658120000 | -0.0096420000 |
| N2 | 1.8931050000 | -0.4793560000 | -1.0247290000 |
| C3 | 1.1131540000 | 0.3739380000 | -1.7144570000 |
| O4 | 0.2809060000 | -0.3115740000 | -2.5694040000 |
| C5 | -0.5698900000 | 0.3951820000 | -3.3596070000 |
| N6 | -0.1646560000 | 1.2221950000 | -4.2851300000 |
| S7 | -1.5052990000 | 1.7701420000 | -5.0686530000 |
| N8 | -2.6200490000 | 0.9133230000 | -4.2020880000 |
| C9 | -1.9857160000 | 0.2051260000 | -3.2983420000 |

| | | | |
|---|---|---|---|
| O10 | 1.1164930000 | 1.5895720000 | -1.6272680000 |
| C11 | 1.6928790000 | -1.8997310000 | -1.0672910000 |
| C12 | 0.9454630000 | -2.5437940000 | -0.0946490000 |
| C13 | 0.7768290000 | -3.9438820000 | -0.1239340000 |
| C14 | 1.3473750000 | -4.7130060000 | -1.1287840000 |
| C15 | 2.1122320000 | -4.0789700000 | -2.1259960000 |
| C16 | 2.2850420000 | -2.6710630000 | -2.0791300000 |
| N17 | 3.0677970000 | -2.3040960000 | -3.1457170000 |
| C18 | 3.3925090000 | -3.4333390000 | -3.8669610000 |
| C19 | 2.8296920000 | -4.5396530000 | -3.2835060000 |
| H20 | 2.2765410000 | 0.2333890000 | 0.9366390000 |
| H21 | 3.2221920000 | 1.0077500000 | -0.3657010000 |
| H22 | 3.6063970000 | -0.6566920000 | 0.1427290000 |
| H23 | -2.5077830000 | -0.4406120000 | -2.6011670000 |
| H24 | 0.4822140000 | -1.9530030000 | 0.6926620000 |
| H25 | 0.1839360000 | -4.4177850000 | 0.6525070000 |
| H26 | 1.2059250000 | -5.7905110000 | -1.1464110000 |
| H27 | 3.3785270000 | -1.3637350000 | -3.3714690000 |
| H28 | 4.0093250000 | -3.3536520000 | -4.7527800000 |
| H29 | 2.9147460000 | -5.5611500000 | -3.6305330000 |

## 14a

| atom | x | y | z |
|---|---|---|---|
| C1 | 2.1729920000 | 0.6613180000 | -2.9987090000 |
| N2 | 2.3054690000 | 1.4239400000 | -1.7509670000 |
| C3 | 2.2798330000 | 2.8579130000 | -1.8157640000 |
| C4 | 1.0916250000 | 3.5666730000 | -1.7549320000 |
| C5 | 1.0936610000 | 4.9763610000 | -1.8107530000 |
| C6 | 2.2791590000 | 5.6871830000 | -1.9245270000 |
| C7 | 3.4960830000 | 4.9844250000 | -1.9951100000 |
| C8 | 3.4820700000 | 3.5664170000 | -1.9460580000 |
| N9 | 4.7795400000 | 3.1251740000 | -2.0177320000 |
| C10 | 5.6131140000 | 4.2199940000 | -2.1113310000 |
| C11 | 4.8752270000 | 5.3749970000 | -2.0989720000 |
| C12 | 2.3204650000 | 0.7714660000 | -0.5791660000 |
| O13 | 2.3800710000 | 1.6963840000 | 0.4724350000 |
| N14 | 2.4936730000 | 1.1117090000 | 1.6963460000 |
| C15 | 3.6785780000 | 0.9006820000 | 2.3529010000 |
| C16 | 3.2894630000 | 0.4009660000 | 3.5675180000 |
| N17 | 1.9133910000 | 0.3162110000 | 3.6426110000 |
| C18 | 1.4547840000 | 0.7602060000 | 2.4886290000 |
| O19 | 2.2983110000 | -0.4270090000 | -0.3884790000 |
| H20 | 1.1177250000 | 0.5673130000 | -3.2765680000 |
| H21 | 2.7113680000 | 1.1996420000 | -3.7816520000 |
| H22 | 2.6137740000 | -0.3291360000 | -2.8681970000 |
| H23 | 0.1577020000 | 3.0199910000 | -1.6467540000 |
| H24 | 0.1476240000 | 5.5061050000 | -1.7531070000 |
| H25 | 2.2692400000 | 6.7740180000 | -1.9554220000 |
| H26 | 5.0923250000 | 2.1584930000 | -2.0285150000 |
| H27 | 6.6838430000 | 4.0807480000 | -2.1803220000 |
| H28 | 5.2620080000 | 6.3842860000 | -2.1539800000 |
| H29 | 4.6346620000 | 1.1300250000 | 1.9018100000 |
| H30 | 3.9197420000 | 0.1024170000 | 4.3923620000 |
| H31 | 0.4243950000 | 0.8553420000 | 2.1717250000 |

## 15a

| atom | x | y | z |
|---|---|---|---|
| C1 | -0.2243950000 | 0.1596850000 | 0.2288520000 |
| N2 | 1.2292610000 | 0.2129880000 | 0.4277960000 |

| | | | |
|---|---|---|---|
| C3 | 1.8938360000 | 1.4725100000 | 0.2431920000 |
| C4 | 1.8655250000 | 2.4524390000 | 1.2216340000 |
| C5 | 2.5158430000 | 3.6887320000 | 1.0204740000 |
| C6 | 3.2060450000 | 3.9518970000 | -0.1538430000 |
| C7 | 3.2405080000 | 2.9707240000 | -1.1634320000 |
| C8 | 2.5666910000 | 1.7389710000 | -0.9573880000 |
| N9 | 2.7445400000 | 0.9640250000 | -2.0764630000 |
| C10 | 3.5108440000 | 1.6662680000 | -2.9834520000 |
| C11 | 3.8395050000 | 2.8935710000 | -2.4684020000 |
| C12 | 1.8692680000 | -0.8691260000 | 0.8967300000 |
| O13 | 3.2318400000 | -0.5781930000 | 1.0576340000 |
| N14 | 3.9566960000 | -1.6375210000 | 1.5011250000 |
| N15 | 4.0736060000 | -1.8498900000 | 2.8069390000 |
| C16 | 4.9256350000 | -2.8755400000 | 2.8840860000 |
| C17 | 5.3480380000 | -3.2949920000 | 1.6046800000 |
| C18 | 4.6935070000 | -2.4552730000 | 0.7207990000 |
| O19 | 1.4078310000 | -1.9628660000 | 1.1484100000 |
| H20 | -0.4695790000 | 0.7704130000 | -0.6423220000 |
| H21 | -0.7452150000 | 0.5505450000 | 1.1085870000 |
| H22 | -0.5265790000 | -0.8720240000 | 0.0489340000 |
| H23 | 1.3424250000 | 2.2516490000 | 2.1535390000 |
| H24 | 2.4784370000 | 4.4371940000 | 1.8067650000 |
| H25 | 3.7137740000 | 4.9035800000 | -0.2935800000 |
| H26 | 2.3427030000 | 0.0471160000 | -2.2479290000 |
| H27 | 3.7635230000 | 1.2233310000 | -3.9378550000 |
| H28 | 4.4371090000 | 3.6538950000 | -2.9538700000 |
| H29 | 5.2029590000 | -3.2678110000 | 3.8538760000 |
| H30 | 6.0361230000 | -4.0913460000 | 1.3614950000 |
| H31 | 4.6870260000 | -2.3609710000 | -0.3564770000 |

## 16a

| atom | x | y | z |
|---|---|---|---|
| C1 | -0.3371970000 | -0.2731520000 | 0.3444400000 |
| N2 | 1.0416410000 | 0.1530200000 | 0.0603580000 |
| C3 | 1.4034730000 | 1.4058090000 | 0.3567590000 |
| O4 | 2.7903270000 | 1.5737050000 | 0.1403390000 |
| N5 | 3.1388900000 | 2.8748330000 | 0.0533430000 |
| N6 | 3.0915260000 | 3.4654240000 | -1.1321270000 |
| N7 | 3.5233750000 | 4.6767110000 | -0.9996470000 |
| C8 | 3.8657550000 | 4.8760240000 | 0.3180540000 |
| C9 | 3.6183230000 | 3.6927400000 | 1.0294760000 |
| C10 | 3.8596780000 | 3.5605660000 | 2.4063360000 |
| C11 | 4.3650820000 | 4.6892580000 | 3.0232990000 |
| C12 | 4.6221340000 | 5.8990640000 | 2.3172280000 |
| C13 | 4.3804120000 | 6.0152720000 | 0.9644750000 |
| O14 | 0.7136110000 | 2.3162210000 | 0.7623510000 |
| C15 | 1.9613990000 | -0.8171400000 | -0.4644920000 |
| C16 | 1.9474600000 | -1.0978040000 | -1.8389440000 |
| C17 | 2.8062720000 | -2.0799900000 | -2.3956920000 |
| C18 | 2.5287340000 | -2.1112070000 | -3.8065080000 |
| C19 | 1.5504970000 | -1.1796850000 | -4.0390410000 |
| N20 | 1.1978560000 | -0.5611350000 | -2.8567630000 |
| C21 | 3.6959260000 | -2.7728330000 | -1.5526570000 |
| C22 | 3.7029010000 | -2.4864610000 | -0.1952820000 |
| C23 | 2.8329750000 | -1.5188360000 | 0.3503370000 |
| H24 | -0.4026050000 | -0.6836350000 | 1.3568530000 |
| H25 | -0.6102940000 | -1.0432990000 | -0.3795940000 |
| H26 | -1.0099830000 | 0.5789370000 | 0.2422120000 |
| H27 | 3.6637170000 | 2.6380730000 | 2.9436470000 |
| H28 | 4.5755250000 | 4.6581750000 | 4.0881810000 |

| | | | |
|---|---|---|---|
| H29 | 5.0181870000 | 6.7477730000 | 2.8663000000 |
| H30 | 4.5702060000 | 6.9306840000 | 0.4108790000 |
| H31 | 3.0000860000 | -2.7442270000 | -4.5469320000 |
| H32 | 1.0678990000 | -0.8952990000 | -4.9647670000 |
| H33 | 0.4973980000 | 0.1701170000 | -2.7760430000 |
| H34 | 4.3685350000 | -3.5231870000 | -1.9617220000 |
| H35 | 4.3842030000 | -3.0126870000 | 0.4669110000 |
| H36 | 2.8432660000 | -1.3078720000 | 1.4168910000 |

**17a**

| atom | x | y | z |
|---|---|---|---|
| O1 | 1.7083930000 | 2.4238890000 | 1.4294730000 |
| C2 | 1.3678140000 | 1.3243050000 | 1.8156210000 |
| O3 | 0.8221670000 | 0.3718180000 | 0.9454730000 |
| N4 | 0.7198910000 | 0.8307340000 | -0.3276720000 |
| C5 | 1.7592510000 | 0.7146700000 | -1.2415130000 |
| C6 | 1.2447130000 | 1.2726060000 | -2.5475190000 |
| C7 | -0.2052360000 | 1.7117280000 | -2.2656010000 |
| C8 | -0.4439240000 | 1.4166150000 | -0.8039210000 |
| O9 | -1.4245000000 | 1.6170200000 | -0.1196850000 |
| O10 | 2.8443440000 | 0.2489450000 | -0.9690820000 |
| N11 | 1.4231760000 | 0.8420440000 | 3.0662350000 |
| C12 | 2.0238170000 | 1.6813410000 | 4.1138520000 |
| C13 | 1.1062560000 | -0.5173510000 | 3.3954260000 |
| C14 | -0.0650390000 | -0.7945610000 | 4.1153360000 |
| N15 | -1.0605440000 | 0.0377880000 | 4.5646720000 |
| C16 | -2.0017520000 | -0.7190070000 | 5.2314470000 |
| C17 | -1.6316960000 | -2.0391190000 | 5.2246040000 |
| C18 | -0.3860690000 | -2.1185520000 | 4.5115270000 |
| C19 | 0.4815660000 | -3.1716960000 | 4.1658780000 |
| C20 | 1.6360550000 | -2.8882380000 | 3.4503530000 |
| C21 | 1.9526190000 | -1.5661520000 | 3.0734350000 |
| H22 | 1.8930310000 | 2.1031320000 | -2.8433830000 |
| H23 | 1.3182390000 | 0.4929230000 | -3.3116960000 |
| H24 | -0.9421590000 | 1.1540790000 | -2.8518560000 |
| H25 | -0.3729010000 | 2.7793590000 | -2.4350300000 |
| H26 | 1.6165030000 | 1.3654790000 | 5.0762950000 |
| H27 | 3.1117880000 | 1.5587000000 | 4.1230230000 |
| H28 | 1.7695210000 | 2.7266860000 | 3.9363600000 |
| H29 | -1.1233330000 | 1.0416810000 | 4.4218620000 |
| H30 | -2.8731490000 | -0.2441700000 | 5.6626560000 |
| H31 | -2.1773320000 | -2.8596170000 | 5.6719250000 |
| H32 | 0.2494000000 | -4.1934570000 | 4.4571860000 |
| H33 | 2.3158710000 | -3.6897660000 | 3.1764160000 |
| H34 | 2.8645790000 | -1.3560730000 | 2.5198390000 |

**18a**

| atom | x | y | z |
|---|---|---|---|
| O1 | 0.0323190000 | 0.0441780000 | 0.0606390000 |
| C2 | 1.2589500000 | 0.0827090000 | -0.0064700000 |
| S3 | 2.1511510000 | 1.6219800000 | -0.2655770000 |
| C4 | 0.7594580000 | 2.8011480000 | -0.2409600000 |
| C5 | 1.3236780000 | 4.1950470000 | -0.5011950000 |
| N6 | 2.0520860000 | -1.0089130000 | 0.1189640000 |
| C7 | 1.4385240000 | -2.3339940000 | 0.2625870000 |
| C8 | 3.4707630000 | -0.9462720000 | -0.0606780000 |
| C9 | 4.3143300000 | -0.8327270000 | 1.0550820000 |
| N10 | 4.0183980000 | -0.7211970000 | 2.3903210000 |
| C11 | 5.1988490000 | -0.6275200000 | 3.0983520000 |

| | | | |
|---|---|---|---|
| C12 | 6.2661030000 | -0.6809770000 | 2.2397690000 |
| C13 | 5.7260150000 | -0.8115110000 | 0.9136560000 |
| C14 | 6.2899790000 | -0.9086620000 | -0.3723480000 |
| C15 | 5.4512270000 | -1.0234940000 | -1.4717520000 |
| C16 | 4.0497060000 | -1.0466940000 | -1.3171720000 |
| H17 | 0.2702150000 | 2.7451850000 | 0.7334570000 |
| H18 | 0.0434370000 | 2.5104960000 | -1.0118540000 |
| H19 | 1.8061790000 | 4.2541930000 | -1.4815710000 |
| H20 | 2.0537340000 | 4.4814430000 | 0.2620120000 |
| H21 | 0.5098210000 | 4.9239040000 | -0.4801170000 |
| H22 | 1.1903570000 | -2.7544320000 | -0.7184400000 |
| H23 | 2.1546530000 | -2.9875390000 | 0.7649110000 |
| H24 | 0.5320650000 | -2.2534020000 | 0.8637550000 |
| H25 | 3.0895700000 | -0.6784860000 | 2.7998020000 |
| H26 | 5.1795010000 | -0.5301040000 | 4.1754310000 |
| H27 | 7.3116190000 | -0.6334280000 | 2.5134410000 |
| H28 | 7.3694880000 | -0.8947520000 | -0.5021480000 |
| H29 | 5.8731840000 | -1.0994010000 | -2.4695460000 |
| H30 | 3.4017590000 | -1.1360680000 | -2.1855540000 |

**19a**

| atom | x | y | z |
|---|---|---|---|
| C1 | 0.6397120000 | 0.1222420000 | 1.0737350000 |
| N2 | 1.9818140000 | -0.0510270000 | 0.4949200000 |
| C3 | 2.5096630000 | 0.9767690000 | -0.2056370000 |
| S4 | 4.1250250000 | 0.6209040000 | -0.9590730000 |
| C5 | 4.5666510000 | 2.2953710000 | -1.4851550000 |
| C6 | 5.3347140000 | 3.0284000000 | -0.4120640000 |
| F7 | 5.6391030000 | 4.2784650000 | -0.8075840000 |
| F8 | 4.6456730000 | 3.1377980000 | 0.7390260000 |
| F9 | 6.4983570000 | 2.4221310000 | -0.1083960000 |
| O10 | 1.9560060000 | 2.0596250000 | -0.3428530000 |
| C11 | 2.6031060000 | -1.3357180000 | 0.5887010000 |
| C12 | 2.6415870000 | -2.2137150000 | -0.4851670000 |
| C13 | 3.2307840000 | -3.4886610000 | -0.3566590000 |
| C14 | 3.7596370000 | -3.9128420000 | 0.8537160000 |
| C15 | 3.7137950000 | -3.0466730000 | 1.9628730000 |
| C16 | 3.1429710000 | -1.7564490000 | 1.8148560000 |
| N17 | 3.2272060000 | -1.1128280000 | 3.0255010000 |
| C18 | 3.8275140000 | -1.9596300000 | 3.9358130000 |
| C19 | 4.1417060000 | -3.1493720000 | 3.3314140000 |
| H20 | 0.4331700000 | -0.7346030000 | 1.7162780000 |
| H21 | -0.1127840000 | 0.1706670000 | 0.2794880000 |
| H22 | 0.6039510000 | 1.0411120000 | 1.6638310000 |
| H23 | 3.6669630000 | 2.8643810000 | -1.7313240000 |
| H24 | 5.2048340000 | 2.2126810000 | -2.3723990000 |
| H25 | 2.2116500000 | -1.9034070000 | -1.4348030000 |
| H26 | 3.2548360000 | -4.1457850000 | -1.2213200000 |
| H27 | 4.2010100000 | -4.9023180000 | 0.9473920000 |
| H28 | 2.9274450000 | -0.1635380000 | 3.2325490000 |
| H29 | 3.9794980000 | -1.6380910000 | 4.9585380000 |
| H30 | 4.6175360000 | -4.0011240000 | 3.7994670000 |

**20a**

| atom | x | y | z |
|---|---|---|---|
| C1 | 0.1042650000 | -0.0342190000 | -0.0505890000 |
| N2 | 1.5543530000 | 0.0109180000 | 0.1656640000 |
| C3 | 2.2229360000 | 1.1850820000 | 0.0820060000 |
| S4 | 3.9764770000 | 1.0228400000 | 0.5316120000 |

| | | | |
|---|---|---|---|
| C5 | 4.5894610000 | 2.6405530000 | 0.1140040000 |
| C6 | 5.5139870000 | 2.7470580000 | -0.9285460000 |
| C7 | 6.0934600000 | 3.9842450000 | -1.2143120000 |
| C8 | 5.7393850000 | 5.1085320000 | -0.4686660000 |
| C9 | 4.8141550000 | 4.9965310000 | 0.5728160000 |
| C10 | 4.2432400000 | 3.7609810000 | 0.8748300000 |
| O11 | 1.7195070000 | 2.2509850000 | -0.2481010000 |
| C12 | 2.2052530000 | -1.2017520000 | 0.5606200000 |
| C13 | 2.7793110000 | -2.0349990000 | -0.4118770000 |
| N14 | 2.8613140000 | -1.8927560000 | -1.7738570000 |
| C15 | 3.5528320000 | -2.9693130000 | -2.2900490000 |
| C16 | 3.9207430000 | -3.8186760000 | -1.2785700000 |
| C17 | 3.4353240000 | -3.2411190000 | -0.0542590000 |
| C18 | 3.4963620000 | -3.6146930000 | 1.3019340000 |
| C19 | 2.9128880000 | -2.7941850000 | 2.2573280000 |
| C20 | 2.2689490000 | -1.5944270000 | 1.8893180000 |
| H21 | -0.1384470000 | -0.9554620000 | -0.5859710000 |
| H22 | -0.4236900000 | -0.0215650000 | 0.9088860000 |
| H23 | -0.1968240000 | 0.8255450000 | -0.6478070000 |
| H24 | 5.7829660000 | 1.8648370000 | -1.5039360000 |
| H25 | 6.8196900000 | 4.0659660000 | -2.0191160000 |
| H26 | 6.1901030000 | 6.0714210000 | -0.6925380000 |
| H27 | 4.5453500000 | 5.8705930000 | 1.1582520000 |
| H28 | 3.5402840000 | 3.6615160000 | 1.6981110000 |
| H29 | 2.4969370000 | -1.1199790000 | -2.3237750000 |
| H30 | 3.7270350000 | -3.0419410000 | -3.3554740000 |
| H31 | 4.4722370000 | -4.7432030000 | -1.3865560000 |
| H32 | 3.9957640000 | -4.5347310000 | 1.5972220000 |
| H33 | 2.9537300000 | -3.0707190000 | 3.3069160000 |
| H34 | 1.8276720000 | -0.9514720000 | 2.6471910000 |

## 21a

| atom | x | y | z |
|---|---|---|---|
| O1 | 0.2077110000 | -0.1050680000 | -1.1620800000 |
| C2 | 1.3721740000 | -0.0133760000 | -0.8048910000 |
| S3 | 2.1932650000 | 1.6120280000 | -0.6246590000 |
| C4 | 0.9153100000 | 2.6334350000 | -1.2614290000 |
| N5 | 0.7838360000 | 2.9913440000 | -2.5317180000 |
| C6 | -0.2885950000 | 3.8446140000 | -2.5587080000 |
| C7 | -0.7966470000 | 4.0013300000 | -1.2894240000 |
| N8 | -0.0190330000 | 3.2267690000 | -0.4715970000 |
| C9 | -0.1738970000 | 3.0777180000 | 0.9741950000 |
| N10 | 2.1835520000 | -1.0505550000 | -0.5109670000 |
| C11 | 1.6699370000 | -2.4259570000 | -0.5497730000 |
| C12 | 3.5134000000 | -0.8665840000 | -0.0108180000 |
| C13 | 4.6156170000 | -0.9648900000 | -0.8748830000 |
| N14 | 4.6804780000 | -1.1904050000 | -2.2270880000 |
| C15 | 6.0074180000 | -1.2069200000 | -2.6098280000 |
| C16 | 6.8126390000 | -0.9885310000 | -1.5226010000 |
| C17 | 5.9429930000 | -0.8283290000 | -0.3886930000 |
| C18 | 6.1548860000 | -0.5860970000 | 0.9815790000 |
| C19 | 5.0601810000 | -0.4947070000 | 1.8302700000 |
| C20 | 3.7470910000 | -0.6407380000 | 1.3378470000 |
| H21 | -0.6284000000 | 4.3001800000 | -3.4787350000 |
| H22 | -1.6203500000 | 4.5863810000 | -0.9036860000 |
| H23 | 0.7304230000 | 3.4229720000 | 1.4807970000 |
| H24 | -1.0245440000 | 3.6832240000 | 1.2894580000 |
| H25 | -0.3612700000 | 2.0304000000 | 1.2219010000 |
| H26 | 2.4269880000 | -3.0695690000 | -1.0054300000 |
| H27 | 1.4591710000 | -2.7750860000 | 0.4664500000 |

| | | | |
|---|---|---|---|
| H28 | 0.7590450000 | -2.4537890000 | -1.1469800000 |
| H29 | 3.8934580000 | -1.3165920000 | -2.8580080000 |
| H30 | 6.2722710000 | -1.3772610000 | -3.6453520000 |
| H31 | 7.8940740000 | -0.9481130000 | -1.5218490000 |
| H32 | 7.1648130000 | -0.4769610000 | 1.3704830000 |
| H33 | 5.2088850000 | -0.3142130000 | 2.8909780000 |
| H34 | 2.8960970000 | -0.5730780000 | 2.0110640000 |

**22a**

| atom | x | y | z |
|---|---|---|---|
| C1 | 3.6619240000 | 1.4133530000 | -2.8380740000 |
| N2 | 3.6443340000 | 1.5263340000 | -1.3696460000 |
| C3 | 4.3340940000 | 2.6188650000 | -0.7552120000 |
| C4 | 3.6492420000 | 3.7137660000 | -0.2464590000 |
| C5 | 4.3410400000 | 4.7897140000 | 0.3444000000 |
| C6 | 5.7268160000 | 4.8032370000 | 0.3999860000 |
| C7 | 6.4451960000 | 3.7185310000 | -0.1367500000 |
| C8 | 5.7381260000 | 2.6250380000 | -0.7018640000 |
| N9 | 6.6593060000 | 1.7063510000 | -1.1404670000 |
| C10 | 7.9258360000 | 2.1867240000 | -0.8750510000 |
| C11 | 7.8434650000 | 3.4097970000 | -0.2625180000 |
| C12 | 2.8338530000 | 0.6920390000 | -0.6860760000 |
| S13 | 2.9201800000 | 0.9300450000 | 1.1350130000 |
| C14 | 1.8591500000 | -0.3760100000 | 1.6291900000 |
| N15 | 2.1171870000 | -1.6626370000 | 1.5474160000 |
| N16 | 1.0586480000 | -2.3265340000 | 2.1028310000 |
| C17 | 0.2021030000 | -1.4194460000 | 2.5220220000 |
| N18 | 0.6583440000 | -0.1774430000 | 2.2454380000 |
| C19 | 0.0060730000 | 1.0904140000 | 2.5736020000 |
| O20 | 2.1272290000 | -0.1601030000 | -1.1945300000 |
| H21 | 3.8331080000 | 0.3746830000 | -3.1294900000 |
| H22 | 4.4727300000 | 2.0400260000 | -3.2125780000 |
| H23 | 2.7122270000 | 1.7606450000 | -3.2575300000 |
| H24 | 2.5637220000 | 3.7275890000 | -0.3048860000 |
| H25 | 3.7715810000 | 5.6212930000 | 0.7485450000 |
| H26 | 6.2526470000 | 5.6428150000 | 0.8486550000 |
| H27 | 6.4639890000 | 0.8068040000 | -1.5702580000 |
| H28 | 8.7956490000 | 1.6050480000 | -1.1502380000 |
| H29 | 8.6739530000 | 4.0231280000 | 0.0616430000 |
| H30 | -0.7359290000 | -1.6175570000 | 3.0242190000 |
| H31 | -0.9406800000 | 0.8578110000 | 3.0642890000 |
| H32 | 0.6424860000 | 1.6632400000 | 3.2523190000 |
| H33 | -0.1787570000 | 1.6521330000 | 1.6556210000 |

**23a**

| atom | x | y | z |
|---|---|---|---|
| C1 | 1.9301280000 | -0.5445530000 | 2.9293600000 |
| N2 | 1.9019860000 | -0.3242080000 | 1.4771260000 |
| C3 | 2.2316180000 | 0.9715020000 | 0.9714980000 |
| C4 | 3.4863940000 | 1.2547940000 | 0.4539510000 |
| C5 | 3.7863710000 | 2.5377690000 | -0.0416730000 |
| C6 | 2.8488650000 | 3.5565880000 | 0.0026530000 |
| C7 | 1.5763050000 | 3.2994840000 | 0.5431860000 |
| C8 | 1.2751080000 | 1.9965400000 | 1.0145940000 |
| N9 | -0.0159790000 | 1.9997270000 | 1.4758850000 |
| C10 | -0.5420790000 | 3.2635370000 | 1.3056530000 |
| C11 | 0.3932870000 | 4.0883260000 | 0.7436110000 |
| C12 | 1.7118280000 | -1.3808380000 | 0.6659740000 |
| S13 | 1.7963990000 | -0.9318590000 | -1.1210090000 |

| | | | |
|---|---|---|---|
| C14 | 1.2076170000 | -2.4635450000 | -1.7238200000 |
| N15 | 0.0326020000 | -2.6570090000 | -2.2999370000 |
| N16 | 0.0149970000 | -3.9439540000 | -2.6726610000 |
| N17 | 1.1180490000 | -4.5322570000 | -2.3460170000 |
| N18 | 1.8851050000 | -3.6219320000 | -1.7605080000 |
| C19 | 3.2098770000 | -3.9601470000 | -1.2558640000 |
| O20 | 1.4970650000 | -2.5230350000 | 1.0157600000 |
| H21 | 2.9402340000 | -0.8146540000 | 3.2537690000 |
| H22 | 1.6266590000 | 0.3822670000 | 3.4205420000 |
| H23 | 1.2315870000 | -1.3418800000 | 3.1865670000 |
| H24 | 4.2352600000 | 0.4667810000 | 0.4320220000 |
| H25 | 4.7724640000 | 2.7237520000 | -0.4564630000 |
| H26 | 3.0932700000 | 4.5457830000 | -0.3771510000 |
| H27 | -0.5358990000 | 1.1992540000 | 1.8241530000 |
| H28 | -1.5594820000 | 3.4710930000 | 1.6086110000 |
| H29 | 0.2643760000 | 5.1350140000 | 0.5018430000 |
| H30 | 3.8610290000 | -3.0969750000 | -1.4038770000 |
| H31 | 3.1329660000 | -4.2119490000 | -0.1964990000 |
| H32 | 3.5682480000 | -4.8124870000 | -1.8341000000 |

**24a**

| atom | x | y | z |
|---|---|---|---|
| C1 | -0.0589210000 | 0.0367710000 | 0.0347730000 |
| N2 | 1.4103040000 | 0.0764660000 | 0.1242730000 |
| C3 | 2.0238940000 | 1.3693350000 | 0.0548520000 |
| C4 | 2.1662410000 | 2.0339990000 | -1.1541630000 |
| C5 | 2.7164800000 | 3.3304880000 | -1.2083230000 |
| C6 | 3.1300440000 | 3.9804770000 | -0.0537570000 |
| C7 | 2.9741840000 | 3.3351890000 | 1.1874920000 |
| C8 | 2.4004800000 | 2.0364820000 | 1.2325360000 |
| N9 | 2.3234950000 | 1.6523970000 | 2.5492730000 |
| C10 | 2.8358860000 | 2.6637810000 | 3.3351610000 |
| C11 | 3.2488710000 | 3.7073510000 | 2.5489500000 |
| C12 | 2.0393320000 | -1.0247310000 | 0.5954220000 |
| Se13 | 4.0198120000 | -1.1044290000 | 0.6247570000 |
| C14 | 4.6016840000 | -0.2538690000 | -1.0942860000 |
| C15 | 5.4272720000 | 0.9996650000 | -0.8694470000 |
| O16 | 1.4441760000 | -2.0068780000 | 1.0265950000 |
| H17 | -0.5117400000 | 0.2845550000 | 1.0016740000 |
| H18 | -0.3725160000 | 0.7649530000 | -0.7141610000 |
| H19 | -0.3751510000 | -0.9618840000 | -0.2681750000 |
| H20 | 1.8510130000 | 1.5357300000 | -2.0687340000 |
| H21 | 2.8236020000 | 3.8163730000 | -2.1738120000 |
| H22 | 3.5611210000 | 4.9775640000 | -0.1047880000 |
| H23 | 1.9228950000 | 0.7947190000 | 2.9205660000 |
| H24 | 2.8605510000 | 2.5531470000 | 4.4113680000 |
| H25 | 3.6895510000 | 4.6340820000 | 2.8919770000 |
| H26 | 5.1745500000 | -1.0312540000 | -1.6021830000 |
| H27 | 3.6953430000 | -0.0646050000 | -1.6692700000 |
| H28 | 4.8596860000 | 1.7547220000 | -0.3196530000 |
| H29 | 6.3433460000 | 0.7752900000 | -0.3151280000 |
| H30 | 5.7153910000 | 1.4202080000 | -1.8394420000 |

**25a**

| atom | x | y | z |
|---|---|---|---|
| C1 | 0.1092400000 | 0.5631680000 | -0.2779070000 |
| N2 | 1.4766110000 | 0.0394860000 | -0.1137480000 |
| C3 | 2.4758700000 | 0.9445290000 | 0.3635050000 |
| C4 | 3.3090460000 | 1.6296710000 | -0.5062360000 |

| | | | |
|---|---|---|---|
| C5 | 4.2803780000 | 2.5255980000 | -0.0149380000 |
| C6 | 4.4225920000 | 2.7486470000 | 1.3473260000 |
| C7 | 3.5757500000 | 2.0747740000 | 2.2476020000 |
| C8 | 2.5988270000 | 1.1804010000 | 1.7391920000 |
| N9 | 1.9141940000 | 0.6472960000 | 2.8013070000 |
| C10 | 2.4302830000 | 1.1720330000 | 3.9676630000 |
| C11 | 3.4439380000 | 2.0489590000 | 3.6790030000 |
| C12 | 1.7567040000 | -1.1474610000 | -0.6805040000 |
| O13 | 0.9457320000 | -1.8065720000 | -1.3218700000 |
| Se14 | 3.5439280000 | -2.0062430000 | -0.5554770000 |
| C15 | 4.5935510000 | -1.2099510000 | 0.8837780000 |
| C16 | 4.3006000000 | -1.5061110000 | 2.2176330000 |
| C17 | 5.1334530000 | -1.0236450000 | 3.2258260000 |
| C18 | 6.2660190000 | -0.2725170000 | 2.9011130000 |
| C19 | 6.5613610000 | 0.0054750000 | 1.5674320000 |
| C20 | 5.7206100000 | -0.4577780000 | 0.5531530000 |
| H21 | -0.6124610000 | -0.2356600000 | -0.1005810000 |
| H22 | -0.0365970000 | 1.3616170000 | 0.4528810000 |
| H23 | -0.0282650000 | 0.9631190000 | -1.2895830000 |
| H24 | 3.2131810000 | 1.4507740000 | -1.5747540000 |
| H25 | 4.9310970000 | 3.0354890000 | -0.7196370000 |
| H26 | 5.1851010000 | 3.4300380000 | 1.7166470000 |
| H27 | 1.1578340000 | -0.0297770000 | 2.7584520000 |
| H28 | 2.0218490000 | 0.8695230000 | 4.9232910000 |
| H29 | 4.0333790000 | 2.6051520000 | 4.3960330000 |
| H30 | 3.4286770000 | -2.1060620000 | 2.4640330000 |
| H31 | 4.9040610000 | -1.2421290000 | 4.2649750000 |
| H32 | 6.9181120000 | 0.0926420000 | 3.6894850000 |
| H33 | 7.4424570000 | 0.5874090000 | 1.3087720000 |
| H34 | 5.9405470000 | -0.2325170000 | -0.4862540000 |

**26a**

| atom | x | y | z |
|---|---|---|---|
| C1 | -0.1436610000 | 0.0858590000 | -0.4153770000 |
| N2 | 1.3250130000 | 0.0195100000 | -0.3207790000 |
| C3 | 2.1485000000 | 0.9857440000 | 0.1373520000 |
| N4 | 1.5874250000 | 2.2921300000 | 0.2598230000 |
| C5 | 1.9785220000 | 3.1956950000 | 1.2350610000 |
| C6 | 1.3365350000 | 4.3579540000 | 0.9297290000 |
| N7 | 0.5856750000 | 4.2035460000 | -0.2258100000 |
| C8 | 0.7588180000 | 2.9597310000 | -0.6038680000 |
| O9 | 3.3211350000 | 0.8172580000 | 0.4514520000 |
| C10 | 1.9307900000 | -1.2489910000 | -0.6146900000 |
| C11 | 2.2695740000 | -2.1275960000 | 0.4243110000 |
| N12 | 2.1362200000 | -2.0054180000 | 1.7845210000 |
| C13 | 2.6461620000 | -3.1392350000 | 2.3816400000 |
| C14 | 3.1139080000 | -4.0035570000 | 1.4263100000 |
| C15 | 2.8802950000 | -3.3792180000 | 0.1524980000 |
| C16 | 3.1261860000 | -3.7510790000 | -1.1837210000 |
| C17 | 2.7671710000 | -2.8832680000 | -2.2045410000 |
| C18 | 2.1744880000 | -1.6337140000 | -1.9216930000 |
| H19 | -0.4657200000 | 0.2574170000 | -1.4471130000 |
| H20 | -0.5335440000 | 0.8693420000 | 0.2356240000 |
| H21 | -0.5365460000 | -0.8755960000 | -0.0785380000 |
| H22 | 2.6536980000 | 2.9125350000 | 2.0293460000 |
| H23 | 1.3647600000 | 5.2959840000 | 1.4653050000 |
| H24 | 0.3449010000 | 2.4999530000 | -1.4919370000 |
| H25 | 1.7071660000 | -1.2361610000 | 2.2899860000 |
| H26 | 2.6324980000 | -3.2361130000 | 3.4588620000 |

| | | | |
|---|---|---|---|
| H27 | 3.5721960000 | -4.9674750000 | 1.6036060000 |
| H28 | 3.5926470000 | -4.7065420000 | -1.4113930000 |
| H29 | 2.9497770000 | -3.1564190000 | -3.2396800000 |
| H30 | 1.9116810000 | -0.9538370000 | -2.7290210000 |

**27a**

| atom | x | y | z |
|---|---|---|---|
| O1 | -0.3718752705 | 1.5266280765 | -1.6202468407 |
| C2 | 0.2804102339 | 0.5335673269 | -1.3396575319 |
| N3 | -1.9958687933 | -2.2238407148 | -1.5665129964 |
| C4 | -1.1672096801 | -2.6404169383 | -0.5644659958 |
| N5 | -1.3521895232 | -3.9033602097 | 0.0993824466 |
| O6 | -2.2966580552 | -4.5968600254 | -0.2357566306 |
| O7 | -0.5391632981 | -4.2014640814 | 0.9635259307 |
| C8 | -0.1781133148 | -1.7397827188 | -0.2839505225 |
| N9 | -0.4180646775 | -0.7159163401 | -1.1645098886 |
| C10 | -1.5269704291 | -1.0511661962 | -1.9071134712 |
| N11 | 1.6139028351 | 0.5004414328 | -1.1686747032 |
| C12 | 2.3216334678 | 1.7879827436 | -1.2815663123 |
| C13 | 2.3990513548 | -0.6976425035 | -1.2389755885 |
| C14 | 3.2142089392 | -1.0528234704 | -0.1539134889 |
| N15 | 3.3947397693 | -0.4526852361 | 1.0676620716 |
| C16 | 4.3064426719 | -1.1927491864 | 1.7918189649 |
| C17 | 4.7218707148 | -2.2737364262 | 1.0587492196 |
| C18 | 4.0293917937 | -2.2134717578 | -0.1993683453 |
| C19 | 4.0020547405 | -3.0315114129 | -1.3450999154 |
| C20 | 3.1862920288 | -2.6765846754 | -2.4091128375 |
| C21 | 2.3975447857 | -1.5084326118 | -2.3634143341 |
| H22 | 0.6134148921 | -1.7396667739 | 0.4502071622 |
| H23 | -1.9169034474 | -0.3979482819 | -2.6770495589 |
| H24 | 3.3383001496 | 1.6461918242 | -0.9141937373 |
| H25 | 2.3511804601 | 2.1100076499 | -2.3277223622 |
| H26 | 1.8119546741 | 2.5414384321 | -0.6784268781 |
| H27 | 2.9239382577 | 0.3782497559 | 1.4144160808 |
| H28 | 4.5881698505 | -0.8791901746 | 2.7882801113 |
| H29 | 5.4335274483 | -3.0260876914 | 1.3720488367 |
| H30 | 4.6118264803 | -3.9305777491 | -1.3933871817 |
| H31 | 3.1538275469 | -3.2975308152 | -3.2995497469 |
| H32 | 1.7749125802 | -1.2352520032 | -3.2123314081 |

**28a**

| atom | x | y | z |
|---|---|---|---|
| O1 | -0.0216340000 | -0.1024970000 | -0.2588340000 |
| C2 | 1.1931620000 | -0.0546180000 | -0.1164190000 |
| N3 | 1.7936180000 | 1.2387490000 | 0.0527920000 |
| N4 | 2.9846170000 | 1.6097520000 | -0.4843500000 |
| C5 | 3.0436290000 | 2.8908240000 | -0.1981730000 |
| N6 | 1.9608030000 | 3.3684120000 | 0.4894010000 |
| C7 | 1.1892570000 | 2.3086160000 | 0.6165740000 |
| N8 | 2.0289510000 | -1.1064370000 | -0.1013750000 |
| C9 | 3.4390570000 | -1.0974720000 | 0.3327580000 |
| C10 | 1.4741390000 | -2.3789280000 | -0.4691060000 |
| C11 | 0.5605760000 | -3.0287700000 | 0.3725040000 |
| N12 | 0.0250370000 | -2.6534840000 | 1.5780370000 |
| C13 | -0.8485730000 | -3.6317550000 | 2.0002920000 |
| C14 | -0.8912790000 | -4.6487520000 | 1.0819490000 |
| C15 | 0.0120340000 | -4.2897920000 | 0.0225760000 |
| C16 | 0.4155850000 | -4.9087270000 | -1.1761280000 |
| C17 | 1.3374460000 | -4.2676010000 | -1.9899580000 |

| | | | |
|---|---|---|---|
| C18 | 1.8624540000 | -3.0046850000 | -1.6417410000 |
| H19 | 3.8836780000 | 3.5053850000 | -0.4941160000 |
| H20 | 0.2194070000 | 2.2508510000 | 1.0940880000 |
| H21 | 3.6095970000 | -0.2916830000 | 1.0459180000 |
| H22 | 4.1120970000 | -0.9840130000 | -0.5209680000 |
| H23 | 3.6258010000 | -2.0537170000 | 0.8245540000 |
| H24 | 0.2383770000 | -1.8109680000 | 2.1010100000 |
| H25 | -1.3773180000 | -3.5163810000 | 2.9373330000 |
| H26 | -1.4916110000 | -5.5459230000 | 1.1466680000 |
| H27 | 0.0101540000 | -5.8773730000 | -1.4581060000 |
| H28 | 1.6600780000 | -4.7332550000 | -2.9165730000 |
| H29 | 2.5735680000 | -2.5062590000 | -2.2960070000 |

**29a**

| atom | x | y | z |
|---|---|---|---|
| C1 | -0.0582250000 | 0.0074150000 | 0.2104910000 |
| N2 | 1.3963950000 | 0.0092690000 | -0.0310550000 |
| C3 | 2.0494000000 | 1.2880930000 | 0.0285470000 |
| C4 | 2.4573180000 | 1.9353630000 | -1.1256090000 |
| C5 | 3.0239650000 | 3.2259490000 | -1.0674800000 |
| C6 | 3.1671950000 | 3.8910040000 | 0.1413370000 |
| C7 | 2.7574420000 | 3.2516670000 | 1.3259430000 |
| C8 | 2.2146630000 | 1.9415310000 | 1.2592040000 |
| N9 | 1.9076140000 | 1.5446020000 | 2.5378730000 |
| C10 | 2.2417270000 | 2.5611350000 | 3.4092840000 |
| C11 | 2.7593710000 | 3.6232430000 | 2.7152060000 |
| C12 | 2.1532120000 | -1.0709740000 | -0.2879090000 |
| N13 | 1.4221890000 | -2.2589100000 | -0.6650940000 |
| N14 | 1.8863240000 | -3.4674650000 | -0.3589590000 |
| C15 | 1.0653950000 | -4.2981560000 | -0.9822440000 |
| C16 | 0.1120940000 | -3.5326070000 | -1.6893620000 |
| N17 | 0.3648340000 | -2.2508840000 | -1.4742850000 |
| O18 | 3.3719300000 | -1.1296900000 | -0.2413440000 |
| H19 | -0.3682640000 | -0.9421280000 | 0.6475300000 |
| H20 | -0.6058880000 | 0.1911310000 | -0.7177140000 |
| H21 | -0.2695590000 | 0.8098870000 | 0.9209380000 |
| H22 | 2.3255610000 | 1.4366090000 | -2.0826750000 |
| H23 | 3.3359900000 | 3.7051580000 | -1.9904620000 |
| H24 | 3.5891570000 | 4.8924330000 | 0.1743180000 |
| H25 | 1.5609180000 | 0.6353870000 | 2.8250270000 |
| H26 | 2.0794250000 | 2.4393040000 | 4.4720360000 |
| H27 | 3.1026980000 | 4.5592470000 | 3.1361770000 |
| H28 | 1.1801460000 | -5.3705660000 | -0.9062850000 |
| H29 | -0.7026230000 | -3.8486620000 | -2.3230710000 |

**30a**

| atom | x | y | z |
|---|---|---|---|
| O1 | 0.2041810000 | -0.1371150000 | 0.9713130000 |
| C2 | 1.2899690000 | -0.0716030000 | 0.4201540000 |
| N3 | 1.7733510000 | 1.2527750000 | 0.1037420000 |
| C4 | 1.0051770000 | 2.3269880000 | -0.1613530000 |
| N5 | 1.7974610000 | 3.3676240000 | -0.2574960000 |
| N6 | 3.0504410000 | 2.9077280000 | -0.0223490000 |
| N7 | 3.0487900000 | 1.6474560000 | 0.1963830000 |
| N8 | 2.0761610000 | -1.0939730000 | 0.0584580000 |
| C9 | 3.3062450000 | -1.0287990000 | -0.7568540000 |
| C10 | 1.5871780000 | -2.3880000000 | 0.4458610000 |
| C11 | 0.4559140000 | -2.9048050000 | -0.1973780000 |
| N12 | -0.3056540000 | -2.3886790000 | -1.2140410000 |

| | | | |
|---|---|---|---|
| C13 | -1.3552550000 | -3.2469080000 | -1.4594360000 |
| C14 | -1.2888810000 | -4.3216250000 | -0.6105480000 |
| C15 | -0.1243160000 | -4.1305760000 | 0.2121310000 |
| C16 | 0.4824720000 | -4.8574600000 | 1.2552900000 |
| C17 | 1.6279260000 | -4.3556990000 | 1.8563400000 |
| C18 | 2.1774520000 | -3.1159790000 | 1.4624910000 |
| H19 | -0.0697850000 | 2.2996840000 | -0.2731970000 |
| H20 | 3.2378310000 | -0.2251210000 | -1.4918770000 |
| H21 | 4.1838100000 | -0.8799650000 | -0.1234210000 |
| H22 | 3.3859190000 | -1.9828690000 | -1.2794510000 |
| H23 | -0.1205640000 | -1.5457230000 | -1.7468110000 |
| H24 | -2.0738960000 | -3.0108390000 | -2.2330940000 |
| H25 | -1.9847970000 | -5.1483210000 | -0.5703170000 |
| H26 | 0.0539100000 | -5.7988420000 | 1.5894330000 |
| H27 | 2.1035780000 | -4.9085930000 | 2.6608670000 |
| H28 | 3.0526760000 | -2.7135450000 | 1.9670020000 |

## 31a

| atom | x | y | z |
|---|---|---|---|
| O1 | -0.0765480000 | -0.0223840000 | 0.5779290000 |
| C2 | 1.1010660000 | 0.0057440000 | 0.2480200000 |
| N3 | 1.7592670000 | 1.2757800000 | 0.2202250000 |
| C4 | 1.1809230000 | 2.5153780000 | 0.0661240000 |
| C5 | -0.1047230000 | 2.9650340000 | -0.2705720000 |
| C6 | -0.2537950000 | 4.3397600000 | -0.3514890000 |
| C7 | 0.8161640000 | 5.2415260000 | -0.1104600000 |
| C8 | 2.0776940000 | 4.7895100000 | 0.2237790000 |
| C9 | 2.2390230000 | 3.3981050000 | 0.3092390000 |
| N10 | 3.3586160000 | 2.6521150000 | 0.6203980000 |
| N11 | 3.0585210000 | 1.4092620000 | 0.5697640000 |
| N12 | 1.8562350000 | -1.0566200000 | -0.0893310000 |
| C13 | 3.1493600000 | -1.0294470000 | -0.7990380000 |
| C14 | 1.3132080000 | -2.3561380000 | 0.1874490000 |
| C15 | 0.6666690000 | -3.0716630000 | -0.8286880000 |
| N16 | 0.3924880000 | -2.7284290000 | -2.1292710000 |
| C17 | -0.2574000000 | -3.7830020000 | -2.7344810000 |
| C18 | -0.4097790000 | -4.8144040000 | -1.8441280000 |
| C19 | 0.1761830000 | -4.3831430000 | -0.6043290000 |
| C20 | 0.3346110000 | -4.9685860000 | 0.6669010000 |
| C21 | 0.9705120000 | -4.2495350000 | 1.6679930000 |
| C22 | 1.4640380000 | -2.9487990000 | 1.4295710000 |
| H23 | -0.9271190000 | 2.2806080000 | -0.4490890000 |
| H24 | -1.2277060000 | 4.7465500000 | -0.6081120000 |
| H25 | 0.6309450000 | 6.3080390000 | -0.1919500000 |
| H26 | 2.9118480000 | 5.4604870000 | 0.4112540000 |
| H27 | 3.9815710000 | -1.0480600000 | -0.0924610000 |
| H28 | 3.2231180000 | -0.1446300000 | -1.4322340000 |
| H29 | 3.1866980000 | -1.9201400000 | -1.4300520000 |
| H30 | 0.6022180000 | -1.8440960000 | -2.5827500000 |
| H31 | -0.5605950000 | -3.7081870000 | -3.7706210000 |
| H32 | -0.8827410000 | -5.7667050000 | -2.0423870000 |
| H33 | -0.0362190000 | -5.9723720000 | 0.8602320000 |
| H34 | 1.1027580000 | -4.6897930000 | 2.6515910000 |
| H35 | 1.9709890000 | -2.3986940000 | 2.2191620000 |

#### 1a-INT

| atom | x | y | z |
|---|---|---|---|
| O1 | -0.0490320000 | -0.1932820000 | 0.0222140000 |
| C2 | 1.2511770000 | -0.1252030000 | -0.0371110000 |

| | | | |
|---|---|---|---|
| N3 | 1.7888650000 | 1.2474820000 | -0.0145110000 |
| C4 | 3.2459620000 | 1.3171000000 | -0.1294090000 |
| C5 | 1.3503060000 | 1.9900760000 | 1.1640250000 |
| O6 | 1.9178230000 | -0.7770320000 | 1.1232710000 |
| C7 | 1.9991640000 | -2.1230860000 | 1.2518260000 |
| C8 | 2.8973700000 | -2.5804090000 | 2.2405930000 |
| N9 | 3.7044320000 | -1.8518130000 | 3.0874050000 |
| C10 | 4.4220060000 | -2.7235290000 | 3.8734520000 |
| C11 | 4.0973640000 | -4.0175020000 | 3.5484230000 |
| C12 | 3.1151340000 | -3.9522390000 | 2.4988460000 |
| C13 | 2.4049870000 | -4.9154040000 | 1.7471350000 |
| C14 | 1.5144950000 | -4.4654010000 | 0.7871850000 |
| C15 | 1.3022090000 | -3.0879110000 | 0.5352120000 |
| O16 | 1.8572520000 | -0.8004040000 | -1.1450560000 |
| C17 | 1.4237770000 | -0.3490550000 | -2.4141690000 |
| H18 | 3.7523650000 | 0.8455490000 | 0.7277660000 |
| H19 | 3.5847160000 | 0.8225220000 | -1.0402920000 |
| H20 | 3.5505110000 | 2.3678560000 | -0.1745450000 |
| H21 | 1.6506400000 | 3.0374840000 | 1.0623820000 |
| H22 | 0.2648330000 | 1.9457050000 | 1.2518320000 |
| H23 | 1.7966880000 | 1.5926730000 | 2.0906640000 |
| H24 | 3.7510450000 | -0.8405170000 | 3.1454390000 |
| H25 | 5.1168230000 | -2.3477680000 | 4.6131120000 |
| H26 | 4.5071120000 | -4.9113640000 | 4.0011630000 |
| H27 | 2.5565840000 | -5.9781110000 | 1.9166020000 |
| H28 | 0.9566380000 | -5.1851820000 | 0.1938180000 |
| H29 | 0.6089850000 | -2.7734500000 | -0.2349890000 |
| H30 | 1.8686890000 | -1.0175920000 | -3.1561000000 |
| H31 | 0.3337670000 | -0.3886170000 | -2.4962860000 |
| H32 | 1.7598400000 | 0.6768870000 | -2.6083340000 |

#### 2a-INT

| atom | x | y | z |
|---|---|---|---|
| O1 | -0.0746800000 | -0.2507520000 | -0.8809830000 |
| C2 | 1.1053700000 | -0.5350880000 | -0.3811510000 |
| N3 | 1.4607240000 | 0.2355540000 | 0.8011680000 |
| C4 | 1.0756850000 | 1.6368110000 | 0.7208410000 |
| C5 | 2.8277910000 | 0.0993040000 | 1.2836390000 |
| S6 | 1.2567990000 | -2.3454690000 | 0.0706490000 |
| C7 | 0.7579200000 | -2.4581810000 | 1.7719840000 |
| C8 | -0.4032100000 | -1.8408050000 | 2.2708080000 |
| N9 | -1.2916170000 | -1.0129500000 | 1.6368010000 |
| C10 | -2.2646850000 | -0.6371960000 | 2.5307050000 |
| C11 | -2.0422280000 | -1.2363550000 | 3.7467000000 |
| C12 | -0.8442410000 | -2.0199360000 | 3.6074100000 |
| C13 | -0.1081580000 | -2.8561860000 | 4.4700970000 |
| C14 | 1.0402170000 | -3.4627250000 | 3.9853430000 |
| C15 | 1.4747650000 | -3.2581120000 | 2.6561050000 |
| O16 | 2.1894410000 | -0.3565310000 | -1.3390310000 |
| C17 | 1.9727240000 | -0.9279470000 | -2.6186150000 |
| H18 | 1.2554320000 | 2.0986570000 | 1.6972800000 |
| H19 | 1.6572570000 | 2.1957280000 | -0.0333350000 |
| H20 | 0.0159580000 | 1.7264930000 | 0.4849480000 |
| H21 | 2.8902540000 | 0.5250230000 | 2.2922030000 |
| H22 | 3.1057080000 | -0.9542310000 | 1.3444630000 |
| H23 | 3.5610420000 | 0.6164310000 | 0.6426670000 |
| H24 | -1.1246890000 | -0.6238150000 | 0.7034640000 |
| H25 | -3.0537510000 | 0.0370590000 | 2.2250240000 |
| H26 | -2.6544930000 | -1.1349640000 | 4.6328880000 |
| H27 | -0.4339760000 | -3.0197130000 | 5.4943870000 |

| | | | |
|---|---|---|---|
| H28 | 1.6276730000 | -4.1076360000 | 4.6334990000 |
| H29 | 2.3851420000 | -3.7422360000 | 2.3119400000 |
| H30 | 2.8435020000 | -0.6770160000 | -3.2274230000 |
| H31 | 1.8916240000 | -2.0225780000 | -2.5641700000 |
| H32 | 1.0698480000 | -0.5275740000 | -3.0887380000 |

#### 3a-INT

| atom | x | y | z |
|---|---|---|---|
| O1 | 0.4264240000 | 1.6453330000 | -0.1275280000 |
| C2 | 1.0031540000 | 0.4689040000 | -0.1049460000 |
| N3 | 2.4162500000 | 0.5163960000 | -0.4842150000 |
| C4 | 3.2157300000 | 1.3984420000 | 0.3580180000 |
| C5 | 3.0863870000 | -0.7653590000 | -0.6717440000 |
| Se6 | 0.8779460000 | -0.3769420000 | 1.7467960000 |
| C7 | -0.7976240000 | -1.3533200000 | 1.6697350000 |
| C8 | -1.9555730000 | -0.7959250000 | 1.1038520000 |
| N9 | -2.1300450000 | 0.4137660000 | 0.4831080000 |
| C10 | -3.4373770000 | 0.5210200000 | 0.0772860000 |
| C11 | -4.1386600000 | -0.6013270000 | 0.4474020000 |
| C12 | -3.2062120000 | -1.4690880000 | 1.1139420000 |
| C13 | -3.3009980000 | -2.7344320000 | 1.7266010000 |
| C14 | -2.1591690000 | -3.2880730000 | 2.2850830000 |
| C15 | -0.9178850000 | -2.6106260000 | 2.2486850000 |
| O16 | 0.3730420000 | -0.5360020000 | -0.8970550000 |
| C17 | 0.0311610000 | -0.1129390000 | -2.2088560000 |
| H18 | 4.1839760000 | 1.5763360000 | -0.1201810000 |
| H19 | 2.7087170000 | 2.3550020000 | 0.4888260000 |
| H20 | 3.4156340000 | 0.9606510000 | 1.3533670000 |
| H21 | 2.5140550000 | -1.4011870000 | -1.3466520000 |
| H22 | 4.0712940000 | -0.5835950000 | -1.1129090000 |
| H23 | 3.2281210000 | -1.3075660000 | 0.2776550000 |
| H24 | -1.3604640000 | 1.0518700000 | 0.2491260000 |
| H25 | -3.7719290000 | 1.4042690000 | -0.4513870000 |
| H26 | -5.1899410000 | -0.7873110000 | 0.2711960000 |
| H27 | -4.2502430000 | -3.2644900000 | 1.7591760000 |
| H28 | -2.2066780000 | -4.2643810000 | 2.7604440000 |
| H29 | -0.0438000000 | -3.0828270000 | 2.6894350000 |
| H30 | 0.9151560000 | 0.2360400000 | -2.7552980000 |
| H31 | -0.3846070000 | -0.9882210000 | -2.7116900000 |
| H32 | -0.7202860000 | 0.6821040000 | -2.1879120000 |

#### 4a-INT

| atom | x | y | z |
|---|---|---|---|
| O1 | 2.7130360000 | 6.2247710000 | 3.3885750000 |
| C2 | 2.2949600000 | 5.6114380000 | 2.2896790000 |
| N3 | 2.5169100000 | 6.3520930000 | 1.0090040000 |
| C4 | 3.9318550000 | 6.5718080000 | 0.7222450000 |
| C5 | 1.8291920000 | 7.6461130000 | 1.0367000000 |
| Te6 | 3.2171070000 | 3.7372930000 | 2.0193650000 |
| C7 | 1.8690710000 | 2.5165250000 | 3.1020960000 |
| C8 | 1.3446150000 | 2.8854490000 | 4.3508990000 |
| N9 | 1.5035310000 | 4.0517260000 | 5.0570960000 |
| C10 | 0.8013270000 | 3.9580310000 | 6.2346330000 |
| C11 | 0.2003990000 | 2.7267000000 | 6.3279290000 |
| C12 | 0.5330830000 | 2.0127440000 | 5.1257330000 |
| C13 | 0.2427900000 | 0.7259540000 | 4.6329670000 |
| C14 | 0.7483630000 | 0.3586590000 | 3.3950730000 |
| C15 | 1.5452990000 | 1.2453480000 | 2.6359670000 |
| O16 | 0.8770060000 | 5.4106140000 | 2.4238320000 |

| | | | |
|---|---|---|---|
| C17 | 0.1392640000 | 4.8539620000 | 1.3479550000 |
| H18 | 4.0169970000 | 7.1765100000 | -0.1863750000 |
| H19 | 4.4556410000 | 7.0966950000 | 1.5381250000 |
| H20 | 4.4563210000 | 5.6301720000 | 0.5169500000 |
| H21 | 1.9053610000 | 8.1050930000 | 0.0472410000 |
| H22 | 0.7732250000 | 7.5071430000 | 1.2709660000 |
| H23 | 2.2598440000 | 8.3370890000 | 1.7757980000 |
| H24 | 1.9303700000 | 4.9018980000 | 4.6747990000 |
| H25 | 0.7859130000 | 4.7901230000 | 6.9263590000 |
| H26 | -0.4035520000 | 2.3689800000 | 7.1509210000 |
| H27 | -0.3654490000 | 0.0355370000 | 5.2127190000 |
| H28 | 0.5347990000 | -0.6279170000 | 2.9913850000 |
| H29 | 1.9162780000 | 0.9155960000 | 1.6684050000 |
| H30 | 0.2279780000 | 5.4570240000 | 0.4382520000 |
| H31 | 0.4662280000 | 3.8303210000 | 1.1280940000 |
| H32 | -0.9035880000 | 4.8276870000 | 1.6733190000 |

#### 5a-INT

| atom | x | y | z |
|---|---|---|---|
| S1 | -0.0285450000 | -0.1298930000 | -0.0381740000 |
| C2 | 1.8102890000 | -0.0698820000 | -0.0569250000 |
| N3 | 2.4105280000 | 1.2351900000 | 0.1355870000 |
| C4 | 2.1043370000 | 1.8422980000 | 1.4248290000 |
| C5 | 2.1477380000 | 2.1713630000 | -0.9509550000 |
| O6 | 2.2291930000 | -0.9272720000 | 1.0160440000 |
| C7 | 3.5529080000 | -1.0041130000 | 1.3855420000 |
| C8 | 3.7632910000 | -1.3584040000 | 2.7292340000 |
| N9 | 2.8433450000 | -1.5900630000 | 3.7243890000 |
| C10 | 3.5169180000 | -1.8961180000 | 4.8844230000 |
| C11 | 4.8711860000 | -1.8675700000 | 4.6587860000 |
| C12 | 5.0575160000 | -1.5236870000 | 3.2744740000 |
| C13 | 6.1811370000 | -1.3287870000 | 2.4425690000 |
| C14 | 5.9666830000 | -0.9808300000 | 1.1194360000 |
| C15 | 4.6668490000 | -0.8171300000 | 0.5824770000 |
| O16 | 2.3438350000 | -0.5369260000 | -1.2759710000 |
| C17 | 2.1741050000 | -1.9217910000 | -1.5448580000 |
| H18 | 2.7202530000 | 2.7392920000 | 1.5364880000 |
| H19 | 1.0448710000 | 2.1296730000 | 1.5109800000 |
| H20 | 2.3482980000 | 1.1587230000 | 2.2404090000 |
| H21 | 2.7586190000 | 3.0655890000 | -0.7956520000 |
| H22 | 2.4201870000 | 1.7278150000 | -1.9078370000 |
| H23 | 1.0879260000 | 2.4725580000 | -0.9876480000 |
| H24 | 1.8353950000 | -1.5554030000 | 3.6196450000 |
| H25 | 2.9692390000 | -2.1113690000 | 5.7924000000 |
| H26 | 5.6454690000 | -2.0644050000 | 5.3889220000 |
| H27 | 7.1894330000 | -1.4457680000 | 2.8313290000 |
| H28 | 6.8166510000 | -0.8227390000 | 0.4608320000 |
| H29 | 4.5348670000 | -0.5248810000 | -0.4517770000 |
| H30 | 2.5267840000 | -2.0825920000 | -2.5654090000 |
| H31 | 2.7717050000 | -2.5287200000 | -0.8544400000 |
| H32 | 1.1211470000 | -2.2071910000 | -1.4687660000 |

#### 6a-INT

| atom | x | y | z |
|---|---|---|---|
| C1 | 1.8712830000 | -0.6761350000 | 0.9011850000 |
| N2 | 1.5804110000 | -0.0415170000 | -0.3923500000 |
| C3 | 1.8715070000 | 1.3634350000 | -0.3731290000 |
| C4 | 0.9121720000 | 2.2248750000 | 0.1471960000 |
| C5 | 1.1375660000 | 3.6114340000 | 0.2915040000 |

| | | | |
|---|---|---|---|
| C6 | 2.3337700000 | 4.1857930000 | -0.1049320000 |
| C7 | 3.3289780000 | 3.3512250000 | -0.6503080000 |
| C8 | 3.0944960000 | 1.9524660000 | -0.7653520000 |
| N9 | 4.2243890000 | 1.3867820000 | -1.3012730000 |
| C10 | 5.1540730000 | 2.3671400000 | -1.5366790000 |
| C11 | 4.6545490000 | 3.5877470000 | -1.1524960000 |
| C12 | 2.1136250000 | -0.8511650000 | -1.5061390000 |
| F13 | 1.9344460000 | 0.0211990000 | -2.6619890000 |
| O14 | 3.3290620000 | -1.2820040000 | -1.4512580000 |
| O15 | 1.1198120000 | -1.8464260000 | -1.6907790000 |
| C16 | 1.4648700000 | -2.8420870000 | -2.6468720000 |
| H17 | 1.5736360000 | -1.7267210000 | 0.8628870000 |
| H18 | 2.9381480000 | -0.6192230000 | 1.1593870000 |
| H19 | 1.2930820000 | -0.1789870000 | 1.6832910000 |
| H20 | -0.0405960000 | 1.8008400000 | 0.4556060000 |
| H21 | 0.3514020000 | 4.2311180000 | 0.7143610000 |
| H22 | 2.5039820000 | 5.2542970000 | 0.0025220000 |
| H23 | 4.3014870000 | 0.3900050000 | -1.5069670000 |
| H24 | 6.1149480000 | 2.1147700000 | -1.9662200000 |
| H25 | 5.1640550000 | 4.5400730000 | -1.2188780000 |
| H26 | 0.5907830000 | -3.4875630000 | -2.7498110000 |
| H27 | 1.6940800000 | -2.3871820000 | -3.6162210000 |
| H28 | 2.3217630000 | -3.4327260000 | -2.3137920000 |

#### 7a-INT

| atom | x | y | z |
|---|---|---|---|
| O1 | 0.0236080000 | 0.1065240000 | 0.1784250000 |
| C2 | 1.3341250000 | 0.0153510000 | 0.1027330000 |
| O3 | 2.0233380000 | 1.2705460000 | 0.0347740000 |
| C4 | 1.6119150000 | 2.2378410000 | 0.9507400000 |
| C5 | 2.5590150000 | 3.3989400000 | 0.7587540000 |
| F6 | 2.2493250000 | 4.4132020000 | 1.5930480000 |
| F7 | 3.8407580000 | 3.0655530000 | 1.0089860000 |
| F8 | 2.5243270000 | 3.8933330000 | -0.4901240000 |
| N9 | 1.9224400000 | -0.7020280000 | -1.0390640000 |
| C10 | 1.3125000000 | -1.9986460000 | -1.2928280000 |
| C11 | 2.2947130000 | 0.0178280000 | -2.2109630000 |
| C12 | 1.4981820000 | 1.0172120000 | -2.8066480000 |
| N13 | 0.2845240000 | 1.5198190000 | -2.4095480000 |
| C14 | -0.0997120000 | 2.5032560000 | -3.2866840000 |
| C15 | 0.8467910000 | 2.6498720000 | -4.2715590000 |
| C16 | 1.8892170000 | 1.7016260000 | -3.9870320000 |
| C17 | 3.1027190000 | 1.3585280000 | -4.6175940000 |
| C18 | 3.8826320000 | 0.3635290000 | -4.0495370000 |
| C19 | 3.4868480000 | -0.2870060000 | -2.8588660000 |
| O20 | 1.7961170000 | -0.6567580000 | 1.3040960000 |
| C21 | 3.2028710000 | -0.8526180000 | 1.3995930000 |
| H22 | 0.5919010000 | 2.5967220000 | 0.7631550000 |
| H23 | 1.6985950000 | 1.9040300000 | 1.9947960000 |
| H24 | 0.3177570000 | -1.9200300000 | -1.7568520000 |
| H25 | 1.9609390000 | -2.5787580000 | -1.9546310000 |
| H26 | 1.2146860000 | -2.5419040000 | -0.3493850000 |
| H27 | -0.2079760000 | 1.2174980000 | -1.5684120000 |
| H28 | -1.0345730000 | 3.0280870000 | -3.1413520000 |
| H29 | 0.8086900000 | 3.3466240000 | -5.0983190000 |
| H30 | 3.4184110000 | 1.8628450000 | -5.5278460000 |
| H31 | 4.8242670000 | 0.0824370000 | -4.5145590000 |
| H32 | 4.1290530000 | -1.0454670000 | -2.4171700000 |
| H33 | 3.4101100000 | -1.1932860000 | 2.4166970000 |
| H34 | 3.5439740000 | -1.6026970000 | 0.6797580000 |

| | | | |
|---|---|---|---|
| H35 | 3.7389810000 | 0.0818310000 | 1.2092560000 |

**8a-INT**

| atom | x | y | z |
|---|---|---|---|
| C1 | -1.5441890000 | -3.0949550000 | 2.5767810000 |
| C2 | -0.9781060000 | -1.8011320000 | 2.5165490000 |
| C3 | -0.6319220000 | -1.2073230000 | 1.3114200000 |
| C4 | -0.8673190000 | -1.9597840000 | 0.1450790000 |
| C5 | -1.4275440000 | -3.2632130000 | 0.1813090000 |
| C6 | -1.7733890000 | -3.8308990000 | 1.4244970000 |
| C7 | -1.5030710000 | -3.7104810000 | -1.1835000000 |
| C8 | -0.9990630000 | -2.6952590000 | -1.9598710000 |
| N9 | -0.6368910000 | -1.6323020000 | -1.1682160000 |
| N10 | -0.1266820000 | 0.1321040000 | 1.2931800000 |
| C11 | -1.1204210000 | 1.1179040000 | 0.8826570000 |
| C12 | 1.1900100000 | 0.3682580000 | 0.6777320000 |
| O13 | 1.2736370000 | 0.5359630000 | -0.6125990000 |
| O14 | 1.9307750000 | -0.7974210000 | 1.1930700000 |
| C15 | 3.1080870000 | -1.1397780000 | 0.5607270000 |
| C16 | 2.9763950000 | -2.5984950000 | 0.1081890000 |
| C17 | 4.2577630000 | -0.9533810000 | 1.5570270000 |
| F18 | 1.9924770000 | -2.7168940000 | -0.7876890000 |
| F19 | 4.1039190000 | -3.0536050000 | -0.4706650000 |
| F20 | 2.6945500000 | -3.4242800000 | 1.1286680000 |
| F21 | 4.3740800000 | 0.3388700000 | 1.8931890000 |
| F22 | 4.0631380000 | -1.6410160000 | 2.6945060000 |
| F23 | 5.4408620000 | -1.3449270000 | 1.0545240000 |
| O24 | 1.7557320000 | 1.5228620000 | 1.3231050000 |
| C25 | 1.7147200000 | 1.5320560000 | 2.7463140000 |
| H26 | -1.8014750000 | -3.5149410000 | 3.5459580000 |
| H27 | -0.7989490000 | -1.2392020000 | 3.4295380000 |
| H28 | -2.2095390000 | -4.8258430000 | 1.4786910000 |
| H29 | -1.8811260000 | -4.6607420000 | -1.5376240000 |
| H30 | -0.8701570000 | -2.6446330000 | -3.0329760000 |
| H31 | -0.0848110000 | -0.8221450000 | -1.4495960000 |
| H32 | -1.9890840000 | 1.0412630000 | 1.5428130000 |
| H33 | -0.6969510000 | 2.1211120000 | 0.9850270000 |
| H34 | -1.4562900000 | 0.9854820000 | -0.1568640000 |
| H35 | 3.3213400000 | -0.5326960000 | -0.3310700000 |
| H36 | 2.4371660000 | 2.2797140000 | 3.0812020000 |
| H37 | 0.7158150000 | 1.7943700000 | 3.1086560000 |
| H38 | 1.9888030000 | 0.5540480000 | 3.1501630000 |

**9a-INT**

| atom | x | y | z |
|---|---|---|---|
| C1 | 1.1282480000 | -1.3103900000 | -0.6409910000 |
| N2 | 2.2229270000 | -0.5314000000 | -0.0703230000 |
| C3 | 1.9756040000 | 0.8652300000 | 0.0334090000 |
| C4 | 1.7709750000 | 1.5832420000 | -1.1465140000 |
| C5 | 1.5552480000 | 2.9742860000 | -1.1693060000 |
| C6 | 1.5564140000 | 3.7196800000 | -0.0012490000 |
| C7 | 1.7524300000 | 3.0401780000 | 1.2154600000 |
| C8 | 1.9340620000 | 1.6270190000 | 1.2267060000 |
| N9 | 2.0651610000 | 1.2454110000 | 2.5404220000 |
| C10 | 1.9916890000 | 2.3475230000 | 3.3476430000 |
| C11 | 1.7980680000 | 3.4733580000 | 2.5846380000 |
| C12 | 2.9321100000 | -1.2490120000 | 1.0108520000 |
| O13 | 4.0506880000 | -0.4242520000 | 1.3331600000 |
| C14 | 5.2318890000 | -0.3251250000 | 0.6270580000 |

| | | | |
|---|---|---|---|
| C15 | 5.1827540000 | -0.0434160000 | -0.9127390000 |
| F16 | 4.6883960000 | 1.1711750000 | -1.1557880000 |
| F17 | 6.4505530000 | -0.0596130000 | -1.3820640000 |
| F18 | 4.5051390000 | -0.9299090000 | -1.6253120000 |
| C19 | 5.8808970000 | 0.9230090000 | 1.3217430000 |
| F20 | 5.0110910000 | 1.9261430000 | 1.3885960000 |
| F21 | 6.2335430000 | 0.5983490000 | 2.5730420000 |
| F22 | 6.9724920000 | 1.3790920000 | 0.6958800000 |
| C23 | 6.1742260000 | -1.5639300000 | 0.8572900000 |
| F24 | 5.8633800000 | -2.1794490000 | 1.9949880000 |
| F25 | 6.1210050000 | -2.4556970000 | -0.1351820000 |
| F26 | 7.4599070000 | -1.1752210000 | 0.9492230000 |
| O27 | 2.2605950000 | -1.4874820000 | 2.1069300000 |
| O28 | 3.4699600000 | -2.4056500000 | 0.3685050000 |
| C29 | 3.3641350000 | -3.6296220000 | 1.0750480000 |
| H30 | 1.4822950000 | -2.3198850000 | -0.8551560000 |
| H31 | 0.2630700000 | -1.3696490000 | 0.0373320000 |
| H32 | 0.8002090000 | -0.8605660000 | -1.5796600000 |
| H33 | 1.8304900000 | 1.0445180000 | -2.0887990000 |
| H34 | 1.4113310000 | 3.4656190000 | -2.1282340000 |
| H35 | 1.4087820000 | 4.7967040000 | -0.0193430000 |
| H36 | 2.2017210000 | 0.2735400000 | 2.8196720000 |
| H37 | 2.0767420000 | 2.2416040000 | 4.4215390000 |
| H38 | 1.6954430000 | 4.4870050000 | 2.9487770000 |
| H39 | 3.7648180000 | -3.5543160000 | 2.0887580000 |
| H40 | 2.3241160000 | -3.9690990000 | 1.1260400000 |
| H41 | 3.9531410000 | -4.3526570000 | 0.5050390000 |

**10a-INT**

| atom | x | y | z |
|---|---|---|---|
| C1 | 1.0141420000 | 0.8751260000 | 1.1490180000 |
| N2 | 2.3332810000 | 0.3059740000 | 0.8965430000 |
| C3 | 3.3001550000 | 1.3450060000 | 0.4328360000 |
| O4 | 2.9201370000 | 1.8262830000 | -0.8612510000 |
| C5 | 2.5524410000 | 0.9795410000 | -1.8855190000 |
| C6 | 3.2796920000 | -0.1548390000 | -2.2647960000 |
| C7 | 2.8180940000 | -0.9409020000 | -3.3210490000 |
| C8 | 1.6658590000 | -0.5918120000 | -4.0267990000 |
| C9 | 0.9692360000 | 0.5640670000 | -3.6714130000 |
| C10 | 1.4062780000 | 1.3434070000 | -2.6008340000 |
| O11 | 3.3417940000 | 2.3903620000 | 1.2275220000 |
| C12 | 2.3826750000 | -1.0569100000 | 0.4759880000 |
| C13 | 1.3623040000 | -1.6973280000 | -0.2777440000 |
| N14 | 0.1808430000 | -1.2189100000 | -0.8119610000 |
| C15 | -0.4464680000 | -2.2164850000 | -1.5178070000 |
| C16 | 0.2945520000 | -3.3659210000 | -1.4543980000 |
| C17 | 1.4601670000 | -3.0614550000 | -0.6714760000 |
| C18 | 2.5909590000 | -3.8220460000 | -0.3228260000 |
| C19 | 3.5796830000 | -3.2095030000 | 0.4277240000 |
| C20 | 3.4632020000 | -1.8646590000 | 0.8327170000 |
| O21 | 4.5417720000 | 0.6592980000 | 0.3085420000 |
| C22 | 5.5752520000 | 1.4200340000 | -0.2933090000 |
| H23 | 0.5501130000 | 1.3201530000 | 0.2540690000 |
| H24 | 0.3549440000 | 0.0958350000 | 1.5358980000 |
| H25 | 1.0948890000 | 1.6629360000 | 1.8992490000 |
| H26 | 4.1621160000 | -0.4496540000 | -1.7102390000 |
| H27 | 3.3705020000 | -1.8361860000 | -3.5934760000 |
| H28 | 1.3182030000 | -1.2119160000 | -4.8482380000 |
| H29 | 0.0768180000 | 0.8555380000 | -4.2187490000 |
| H30 | 0.8662610000 | 2.2386330000 | -2.3013490000 |

| | | | |
|---|---|---|---|
| H31 | -0.1754440000 | -0.2752920000 | -0.7584750000 |
| H32 | -1.3867670000 | -2.0171130000 | -2.0147510000 |
| H33 | 0.0504710000 | -4.3157140000 | -1.9115870000 |
| H34 | 2.6794170000 | -4.8597620000 | -0.6358850000 |
| H35 | 4.4619410000 | -3.7707430000 | 0.7249280000 |
| H36 | 4.2539810000 | -1.4169440000 | 1.4222420000 |
| H37 | 5.7050080000 | 2.3858920000 | 0.2064710000 |
| H38 | 6.4919650000 | 0.8348890000 | -0.1925260000 |
| H39 | 5.3748760000 | 1.5865890000 | -1.3587430000 |

#### 11a-INT

| atom | x | y | z |
|---|---|---|---|
| O1 | -0.9806820000 | -0.4452130000 | -0.0356630000 |
| C2 | 0.1776740000 | -0.3297850000 | 0.5701020000 |
| O3 | 0.8022860000 | 0.8543360000 | 0.0515260000 |
| C4 | 2.1368580000 | 1.0800040000 | 0.1426760000 |
| C5 | 2.8677940000 | 1.2043830000 | 1.3275960000 |
| F6 | 2.2836880000 | 1.1386130000 | 2.5263430000 |
| C7 | 4.2409920000 | 1.4271330000 | 1.3080290000 |
| F8 | 4.9091590000 | 1.5296770000 | 2.4611380000 |
| C9 | 4.9166850000 | 1.5707150000 | 0.1086220000 |
| F10 | 6.2338590000 | 1.7885330000 | 0.0972440000 |
| C11 | 4.2033630000 | 1.5160380000 | -1.0795720000 |
| F12 | 4.8385860000 | 1.6739240000 | -2.2432830000 |
| C13 | 2.8381690000 | 1.2859750000 | -1.0530710000 |
| F14 | 2.1788420000 | 1.2628870000 | -2.2113960000 |
| N15 | 1.1069610000 | -1.4994800000 | 0.4224030000 |
| C16 | 1.7468760000 | -2.0305470000 | 1.6276070000 |
| C17 | 1.8516570000 | -1.6563790000 | -0.7742830000 |
| C18 | 1.2882700000 | -1.4709740000 | -2.0631630000 |
| N19 | 0.0242360000 | -1.0711130000 | -2.4187070000 |
| C20 | -0.0601970000 | -0.9997090000 | -3.7824560000 |
| C21 | 1.1362030000 | -1.3770340000 | -4.3447430000 |
| C22 | 2.0242850000 | -1.6811180000 | -3.2592710000 |
| C23 | 3.3688400000 | -2.0978090000 | -3.1986190000 |
| C24 | 3.9312120000 | -2.2721960000 | -1.9486830000 |
| C25 | 3.1946390000 | -2.0511910000 | -0.7633700000 |
| O26 | 0.0520790000 | -0.1764950000 | 1.9716190000 |
| C27 | -0.7670110000 | 0.9135710000 | 2.3671240000 |
| H28 | 2.5648060000 | -1.4007660000 | 2.0077120000 |
| H29 | 1.0088740000 | -2.1404590000 | 2.4172330000 |
| H30 | 2.1540860000 | -3.0168780000 | 1.3944720000 |
| H31 | -0.6579970000 | -0.7720020000 | -1.7065490000 |
| H32 | -0.9825310000 | -0.6868920000 | -4.2544600000 |
| H33 | 1.3606840000 | -1.4296400000 | -5.4017660000 |
| H34 | 3.9413950000 | -2.2727230000 | -4.1057820000 |
| H35 | 4.9689010000 | -2.5830100000 | -1.8590730000 |
| H36 | 3.7112530000 | -2.1793330000 | 0.1808870000 |
| H37 | -0.7625010000 | 0.9174690000 | 3.4588090000 |
| H38 | -0.3609350000 | 1.8638470000 | 2.0029320000 |
| H39 | -1.7931060000 | 0.7929110000 | 2.0049060000 |

#### 12a-INT

| atom | x | y | z |
|---|---|---|---|
| C1 | -3.0366890000 | -2.2343920000 | 0.1837500000 |
| C2 | -2.1162660000 | -1.3684350000 | -0.4402440000 |
| C3 | -0.7402510000 | -1.3703840000 | -0.1701940000 |
| C4 | -0.3229800000 | -2.3465750000 | 0.7844830000 |
| C5 | -1.2592880000 | -3.1981790000 | 1.4431170000 |

| | | | |
|---|---|---|---|
| C6 | -2.6328650000 | -3.1466080000 | 1.1373650000 |
| C7 | -0.5191250000 | -3.9962990000 | 2.3785360000 |
| C8 | 0.7965800000 | -3.6272930000 | 2.2545500000 |
| N9 | 0.9137700000 | -2.6430620000 | 1.3144330000 |
| O10 | 4.1556320000 | 0.8675970000 | 2.9851480000 |
| C11 | 4.8776840000 | 0.5991010000 | 1.8851920000 |
| C12 | 4.0705380000 | 0.2715790000 | 0.8481050000 |
| C13 | 2.7551130000 | 0.3681690000 | 1.4160480000 |
| N14 | 2.8032070000 | 0.7170220000 | 2.6871370000 |
| N15 | 0.0697150000 | -0.4135590000 | -0.8331740000 |
| C16 | -0.6255880000 | 0.8425320000 | -1.1493400000 |
| C17 | 1.5115650000 | -0.3422260000 | -0.5699840000 |
| O18 | 2.1528170000 | -1.4467600000 | -0.7330180000 |
| O19 | 1.5614840000 | 0.1780930000 | 0.8708300000 |
| O20 | 2.0569500000 | 0.7806370000 | -1.2461050000 |
| C21 | 2.3673270000 | 0.5402980000 | -2.6108740000 |
| H22 | -4.0842260000 | -2.1588520000 | -0.0967570000 |
| H23 | -2.5097570000 | -0.6707780000 | -1.1693820000 |
| H24 | -3.3409730000 | -3.8005040000 | 1.6398730000 |
| H25 | -0.9225180000 | -4.7426450000 | 3.0500180000 |
| H26 | 1.6757860000 | -3.9938410000 | 2.7679940000 |
| H27 | 1.7762850000 | -2.2895480000 | 0.9098140000 |
| H28 | 5.9501240000 | 0.6882440000 | 1.9972630000 |
| H29 | 4.3371150000 | 0.0097520000 | -0.1628720000 |
| H30 | -1.2723250000 | 1.1583140000 | -0.3206700000 |
| H31 | 0.1052650000 | 1.6226470000 | -1.3372150000 |
| H32 | -1.2440630000 | 0.7364200000 | -2.0462870000 |
| H33 | 2.7570140000 | 1.4791130000 | -3.0078610000 |
| H34 | 3.1249760000 | -0.2413880000 | -2.7083430000 |
| H35 | 1.4704280000 | 0.2527770000 | -3.1733100000 |

#### 13a-INT

| atom | x | y | z |
|---|---|---|---|
| C1 | -0.5945880000 | -0.1565620000 | -0.7605160000 |
| N2 | 0.7759940000 | 0.1514490000 | -0.3602000000 |
| C3 | 1.1656650000 | 1.5301630000 | -0.6619580000 |
| O4 | 0.4534860000 | 2.3524280000 | 0.4196540000 |
| C5 | 0.5196330000 | 1.9755380000 | 1.6932850000 |
| N6 | -0.3348090000 | 2.4793040000 | 2.5563400000 |
| S7 | -0.0019100000 | 1.8571540000 | 4.0544950000 |
| N8 | 1.2743390000 | 0.9057190000 | 3.5908260000 |
| C9 | 1.4603140000 | 1.0531500000 | 2.3056330000 |
| O10 | 0.8076240000 | 2.0571510000 | -1.7864510000 |
| C11 | 1.7053630000 | -0.8937930000 | -0.6480240000 |
| C12 | 1.8710740000 | -1.9303640000 | 0.2614690000 |
| C13 | 2.7266780000 | -3.0245320000 | 0.0001350000 |
| C14 | 3.4549480000 | -3.0984710000 | -1.1759730000 |
| C15 | 3.3166500000 | -2.0614310000 | -2.1214220000 |
| C16 | 2.4344060000 | -0.9828030000 | -1.8508870000 |
| N17 | 2.4640020000 | -0.1328640000 | -2.9288080000 |
| C18 | 3.3403380000 | -0.6284110000 | -3.8638980000 |
| C19 | 3.8860980000 | -1.8068650000 | -3.4167180000 |
| O20 | 2.5331100000 | 1.5884300000 | -0.3303030000 |
| C21 | 3.1956580000 | 2.7783040000 | -0.7292620000 |
| H22 | -0.8898270000 | -1.1084930000 | -0.3130750000 |
| H23 | -1.2640140000 | 0.6228170000 | -0.3873990000 |
| H24 | -0.7078090000 | -0.2242350000 | -1.8517940000 |
| H25 | 2.2502100000 | 0.5277380000 | 1.7865250000 |
| H26 | 1.3146740000 | -1.8881150000 | 1.1950310000 |
| H27 | 2.8149670000 | -3.8125110000 | 0.7433300000 |

| | | | |
|---|---|---|---|
| H28 | 4.1170400000 | -3.9395000000 | -1.3685840000 |
| H29 | 1.9312680000 | 0.7339740000 | -2.9926300000 |
| H30 | 3.5066820000 | -0.0959350000 | -4.7911230000 |
| H31 | 4.6044600000 | -2.4211090000 | -3.9435760000 |
| H32 | 3.2302260000 | 2.8621620000 | -1.8191630000 |
| H33 | 2.7031500000 | 3.6625290000 | -0.3093710000 |
| H34 | 4.2113920000 | 2.7086400000 | -0.3342060000 |

#### 14a-INT

| atom | x | y | z |
|---|---|---|---|
| C1 | 0.0417710000 | -0.0788480000 | 0.5149250000 |
| N2 | 1.3621630000 | -0.0842720000 | -0.1103250000 |
| C3 | 2.0066640000 | 1.1915360000 | -0.0961390000 |
| C4 | 1.7788430000 | 2.0782410000 | -1.1400690000 |
| C5 | 2.3377810000 | 3.3760320000 | -1.1567940000 |
| C6 | 3.1595420000 | 3.8147610000 | -0.1307470000 |
| C7 | 3.4208390000 | 2.9398340000 | 0.9435110000 |
| C8 | 2.8305140000 | 1.6487650000 | 0.9518150000 |
| N9 | 3.2291890000 | 1.0093500000 | 2.0993380000 |
| C10 | 4.0565960000 | 1.8442520000 | 2.8112680000 |
| C11 | 4.2020550000 | 3.0378190000 | 2.1468660000 |
| C12 | 2.1217670000 | -1.2899840000 | 0.2309910000 |
| O13 | 3.4526220000 | -0.9265590000 | -0.4169250000 |
| N14 | 4.3919870000 | -1.8782850000 | -0.2182850000 |
| C15 | 4.6773900000 | -2.9062210000 | -1.0810540000 |
| C16 | 5.7692810000 | -3.5224550000 | -0.5260210000 |
| N17 | 6.1410740000 | -2.8852610000 | 0.6416450000 |
| C18 | 5.2825380000 | -1.8882580000 | 0.7948870000 |
| O19 | 2.2333930000 | -1.6557690000 | 1.4660990000 |
| O20 | 1.6077360000 | -2.3739650000 | -0.5432600000 |
| C21 | 1.3368220000 | -2.1132370000 | -1.9188710000 |
| H22 | 0.0860770000 | 0.1301210000 | 1.5942350000 |
| H23 | -0.5752160000 | 0.6830160000 | 0.0320800000 |
| H24 | -0.4331090000 | -1.0533980000 | 0.3676350000 |
| H25 | 1.1521700000 | 1.7430510000 | -1.9627610000 |
| H26 | 2.1241230000 | 4.0309930000 | -1.9974300000 |
| H27 | 3.5948630000 | 4.8112310000 | -0.1532210000 |
| H28 | 2.9686570000 | 0.0504190000 | 2.3308120000 |
| H29 | 4.4827900000 | 1.5185720000 | 3.7511230000 |
| H30 | 4.7945140000 | 3.8836380000 | 2.4706980000 |
| H31 | 4.1021230000 | -3.0780850000 | -1.9789480000 |
| H32 | 6.3105360000 | -4.3788530000 | -0.9025620000 |
| H33 | 5.2594450000 | -1.1527900000 | 1.5872410000 |
| H34 | 1.2514000000 | -3.0852340000 | -2.4105500000 |
| H35 | 0.4036550000 | -1.5545400000 | -2.0361150000 |
| H36 | 2.1462060000 | -1.5391940000 | -2.3789120000 |

#### 15a-INT

| atom | x | y | z |
|---|---|---|---|
| C1 | 1.6727020000 | 0.0781600000 | -2.6585600000 |
| N2 | 1.8303570000 | -0.5516240000 | -1.3468980000 |
| C3 | 2.4484180000 | 0.2042600000 | -0.3203690000 |
| C4 | 1.9514850000 | 0.3990600000 | 0.9672520000 |
| C5 | 2.5951780000 | 1.2417820000 | 1.8977120000 |
| C6 | 3.7496700000 | 1.9368120000 | 1.5819880000 |
| C7 | 4.3050190000 | 1.7393990000 | 0.3028390000 |
| C8 | 3.6687310000 | 0.8615930000 | -0.6110590000 |
| N9 | 4.4556550000 | 0.8070800000 | -1.7449300000 |
| C10 | 5.5516890000 | 1.6213200000 | -1.5733740000 |

| | | | |
|---|---|---|---|
| C11 | 5.4969690000 | 2.2201850000 | -0.3417560000 |
| C12 | 1.0440450000 | -1.7512470000 | -1.1769660000 |
| O13 | -0.4767820000 | -1.3376520000 | -1.1334260000 |
| N14 | -0.8314830000 | -0.3986600000 | -0.2281990000 |
| N15 | -1.2704750000 | -0.7573590000 | 0.9797500000 |
| C16 | -1.6087620000 | 0.3991140000 | 1.5641740000 |
| C17 | -1.3885140000 | 1.4939570000 | 0.7077820000 |
| C18 | -0.8866390000 | 0.9302460000 | -0.4542700000 |
| O19 | 1.0649460000 | -2.5794710000 | -2.1661720000 |
| O20 | 1.3358950000 | -2.2439470000 | 0.0971940000 |
| C21 | 0.7078580000 | -3.4798610000 | 0.4166570000 |
| H22 | 2.3453950000 | -0.3402450000 | -3.4169380000 |
| H23 | 1.8580110000 | 1.1515710000 | -2.5732940000 |
| H24 | 0.6467050000 | -0.0628270000 | -3.0119610000 |
| H25 | 1.0449850000 | -0.1067780000 | 1.2707090000 |
| H26 | 2.1434820000 | 1.3568640000 | 2.8800180000 |
| H27 | 4.2226830000 | 2.6015280000 | 2.3002120000 |
| H28 | 4.3194670000 | 0.2030330000 | -2.5472310000 |
| H29 | 6.2914730000 | 1.7093290000 | -2.3579580000 |
| H30 | 6.2199470000 | 2.9132170000 | 0.0676960000 |
| H31 | -2.0008220000 | 0.3955350000 | 2.5732810000 |
| H32 | -1.5670180000 | 2.5410790000 | 0.9051780000 |
| H33 | -0.5764880000 | 1.3492360000 | -1.4009140000 |
| H34 | -0.3800970000 | -3.3860580000 | 0.3572820000 |
| H35 | 1.0491520000 | -4.2789000000 | -0.2469270000 |
| H36 | 0.9930530000 | -3.7086210000 | 1.4449010000 |

#### 16a-INT

| atom | x | y | z |
|---|---|---|---|
| C1 | 1.7707540000 | 2.1941620000 | 0.2684060000 |
| N2 | 1.6550010000 | 0.8012240000 | -0.1650630000 |
| C3 | 2.1792490000 | 0.4435030000 | -1.5139330000 |
| O4 | 3.2744970000 | -0.4889300000 | -1.3895980000 |
| N5 | 4.1824510000 | -0.2976590000 | -0.3975180000 |
| N6 | 4.7722220000 | 0.8607650000 | -0.1605910000 |
| N7 | 5.6003290000 | 0.7026290000 | 0.8326190000 |
| C8 | 5.5606770000 | -0.6124570000 | 1.2300040000 |
| C9 | 4.6306300000 | -1.2842830000 | 0.4276050000 |
| C10 | 4.3430500000 | -2.6526500000 | 0.5611530000 |
| C11 | 5.0471880000 | -3.3097610000 | 1.5490000000 |
| C12 | 5.9956870000 | -2.6446470000 | 2.3765040000 |
| C13 | 6.2672020000 | -1.2990070000 | 2.2378810000 |
| O14 | 1.3379810000 | -0.1394870000 | -2.3256840000 |
| C15 | 1.7105960000 | -0.1659020000 | 0.8679040000 |
| C16 | 1.2838150000 | -1.5030810000 | 0.6728030000 |
| C17 | 1.3218210000 | -2.4750060000 | 1.7053250000 |
| C18 | 0.8405890000 | -3.7005260000 | 1.1334020000 |
| C19 | 0.5410740000 | -3.4342490000 | -0.1815300000 |
| N20 | 0.7975400000 | -2.1163920000 | -0.4581580000 |
| C21 | 1.8024720000 | -2.1252350000 | 2.9836860000 |
| C22 | 2.2229010000 | -0.8242700000 | 3.1815000000 |
| C23 | 2.1781120000 | 0.1388510000 | 2.1473620000 |
| O24 | 2.7159420000 | 1.6583270000 | -2.0164800000 |
| C25 | 3.4318280000 | 1.5252450000 | -3.2364610000 |
| H26 | 2.7967980000 | 2.4812850000 | 0.5350490000 |
| H27 | 1.4225880000 | 2.8560600000 | -0.5227120000 |
| H28 | 1.1272290000 | 2.3311970000 | 1.1412230000 |
| H29 | 3.6060580000 | -3.1490400000 | -0.0636500000 |
| H30 | 4.8692640000 | -4.3682620000 | 1.7098760000 |
| H31 | 6.5169840000 | -3.2210450000 | 3.1352310000 |

| | | | |
|---|---|---|---|
| H32 | 6.9912330000 | -0.7865770000 | 2.8653090000 |
| H33 | 0.7363330000 | -4.6535870000 | 1.6346240000 |
| H34 | 0.1555960000 | -4.0875820000 | -0.9536210000 |
| H35 | 0.7383530000 | -1.6433310000 | -1.3631930000 |
| H36 | 1.8495490000 | -2.8611470000 | 3.7823070000 |
| H37 | 2.6116970000 | -0.5206240000 | 4.1500140000 |
| H38 | 2.5459060000 | 1.1367060000 | 2.3627480000 |
| H39 | 3.6052760000 | 2.5382690000 | -3.6059750000 |
| H40 | 4.3969580000 | 1.0345200000 | -3.0644780000 |
| H41 | 2.8558700000 | 0.9594900000 | -3.9746020000 |

#### 17a-INT

| atom | x | y | z |
|---|---|---|---|
| O1 | -0.1556630000 | -0.9989120000 | 0.1191840000 |
| C2 | 1.0929740000 | -0.6477740000 | 0.2800800000 |
| O3 | 1.0793710000 | 0.7940210000 | 0.3632720000 |
| N4 | 2.2789080000 | 1.4503020000 | 0.3035300000 |
| C5 | 2.4548300000 | 2.4285190000 | -0.6870220000 |
| C6 | 3.8194540000 | 3.0394960000 | -0.4658210000 |
| C7 | 4.2574220000 | 2.5264090000 | 0.9162830000 |
| C8 | 3.1227820000 | 1.6422150000 | 1.3942030000 |
| O9 | 2.9496380000 | 1.1944910000 | 2.5086980000 |
| O10 | 1.6165920000 | 2.7236340000 | -1.5081110000 |
| N11 | 2.0923420000 | -1.0763140000 | -0.7330940000 |
| C12 | 1.9530340000 | -2.5151790000 | -1.0039910000 |
| C13 | 2.1878910000 | -0.3406320000 | -1.9628950000 |
| C14 | 1.0920760000 | 0.0031130000 | -2.7786770000 |
| N15 | -0.2519100000 | -0.1741860000 | -2.5722070000 |
| C16 | -0.9412580000 | 0.3419860000 | -3.6414000000 |
| C17 | -0.0609910000 | 0.8507780000 | -4.5658260000 |
| C18 | 1.2585200000 | 0.6507900000 | -4.0329400000 |
| C19 | 2.5539610000 | 0.9672890000 | -4.4854230000 |
| C20 | 3.6357210000 | 0.6394160000 | -3.6827130000 |
| C21 | 3.4492520000 | -0.0041430000 | -2.4398160000 |
| O22 | 1.6917630000 | -1.1391520000 | 1.4778390000 |
| C23 | 0.8591780000 | -1.0433430000 | 2.6215130000 |
| H24 | 4.4796820000 | 2.7036820000 | -1.2724210000 |
| H25 | 3.7383980000 | 4.1278670000 | -0.5225010000 |
| H26 | 5.1694360000 | 1.9217380000 | 0.8797880000 |
| H27 | 4.4089870000 | 3.3273390000 | 1.6447600000 |
| H28 | 2.8107530000 | -2.8484480000 | -1.5935830000 |
| H29 | 1.9393560000 | -3.0640980000 | -0.0606580000 |
| H30 | 1.0341240000 | -2.7459300000 | -1.5606210000 |
| H31 | -0.6307280000 | -0.5370570000 | -1.6940740000 |
| H32 | -2.0227600000 | 0.3026410000 | -3.6631910000 |
| H33 | -0.3192750000 | 1.3129290000 | -5.5094930000 |
| H34 | 2.7016730000 | 1.4597410000 | -5.4439310000 |
| H35 | 4.6459330000 | 0.8743300000 | -4.0080340000 |
| H36 | 4.3108340000 | -0.2553470000 | -1.8244940000 |
| H37 | 1.4857720000 | -1.2773480000 | 3.4841080000 |
| H38 | 0.4656470000 | -0.0285400000 | 2.7369160000 |
| H39 | 0.0275630000 | -1.7517610000 | 2.5670580000 |

#### 18a-INT

| atom | x | y | z |
|---|---|---|---|
| O1 | 0.0655850000 | -0.3629380000 | 0.2216760000 |
| C2 | 1.3461140000 | -0.2340370000 | 0.1637310000 |
| S3 | 1.8743080000 | 1.6810130000 | 0.1047150000 |
| C4 | 1.3482670000 | 2.1128870000 | -1.5922000000 |

| | | | |
|---|---|---|---|
| C5 | 2.4242710000 | 2.9108210000 | -2.3216940000 |
| N6 | 1.9856870000 | -0.9545990000 | -0.9633030000 |
| C7 | 1.0527760000 | -1.6983290000 | -1.8078110000 |
| C8 | 3.1753890000 | -0.5172200000 | -1.5998580000 |
| C9 | 4.3760110000 | -0.2453600000 | -0.8969990000 |
| N10 | 4.6625330000 | -0.3486690000 | 0.4420450000 |
| C11 | 5.9610050000 | 0.0328880000 | 0.6649050000 |
| C12 | 6.5527230000 | 0.3748320000 | -0.5257190000 |
| C13 | 5.5586820000 | 0.1952330000 | -1.5480470000 |
| C14 | 5.5856840000 | 0.3305140000 | -2.9509410000 |
| C15 | 4.4340860000 | 0.0205630000 | -3.6503650000 |
| C16 | 3.2547650000 | -0.3870730000 | -2.9884480000 |
| O17 | 2.0415180000 | -0.7173510000 | 1.3420240000 |
| C18 | 1.6733940000 | -0.1111670000 | 2.5756640000 |
| H19 | 1.1463470000 | 1.1754890000 | -2.1169010000 |
| H20 | 0.4131010000 | 2.6759950000 | -1.5499660000 |
| H21 | 2.1026690000 | 3.1310720000 | -3.3453420000 |
| H22 | 2.6205140000 | 3.8625430000 | -1.8179130000 |
| H23 | 3.3566570000 | 2.3404090000 | -2.3658560000 |
| H24 | 1.6243660000 | -2.4041140000 | -2.4147080000 |
| H25 | 0.3617100000 | -2.2571850000 | -1.1793010000 |
| H26 | 0.4601420000 | -1.0612920000 | -2.4805340000 |
| H27 | 3.9723450000 | -0.6108010000 | 1.1380050000 |
| H28 | 6.3686470000 | 0.0240520000 | 1.6673790000 |
| H29 | 7.5741980000 | 0.7040690000 | -0.6645240000 |
| H30 | 6.4897860000 | 0.6523470000 | -3.4622080000 |
| H31 | 4.4170830000 | 0.1021590000 | -4.7341380000 |
| H32 | 2.3758240000 | -0.5844290000 | -3.5931620000 |
| H33 | 1.9959890000 | -0.7859220000 | 3.3713170000 |
| H34 | 2.1751040000 | 0.8551710000 | 2.7060350000 |
| H35 | 0.5912070000 | 0.0298830000 | 2.6328770000 |

#### 19a-INT

| atom | x | y | z |
|---|---|---|---|
| C1 | 0.4217080000 | 0.1437280000 | -0.1384990000 |
| N2 | 1.2150790000 | 0.1881210000 | -1.3701600000 |
| C3 | 2.1342000000 | 1.3755760000 | -1.4300170000 |
| S4 | 0.9270220000 | 2.6474260000 | -2.0349340000 |
| C5 | 1.8257980000 | 4.1939150000 | -1.7751580000 |
| C6 | 2.5199350000 | 4.6881410000 | -3.0175560000 |
| F7 | 3.1052040000 | 5.8884000000 | -2.8050730000 |
| F8 | 1.6757090000 | 4.8585250000 | -4.0562890000 |
| F9 | 3.4914270000 | 3.8609390000 | -3.4434040000 |
| O10 | 2.7486700000 | 1.7221890000 | -0.3272900000 |
| C11 | 1.7784860000 | -1.0986390000 | -1.6630830000 |
| C12 | 1.0751210000 | -1.9861160000 | -2.4675280000 |
| C13 | 1.5470250000 | -3.2905430000 | -2.7339740000 |
| C14 | 2.7469570000 | -3.7404810000 | -2.2052230000 |
| C15 | 3.4906590000 | -2.8683130000 | -1.3854990000 |
| C16 | 2.9922570000 | -1.5649590000 | -1.1222400000 |
| N17 | 3.8844390000 | -0.9297120000 | -0.2971530000 |
| C18 | 4.9346470000 | -1.7753640000 | -0.0384440000 |
| C19 | 4.7380370000 | -2.9760130000 | -0.6774540000 |
| O20 | 3.1088210000 | 1.1149150000 | -2.4437700000 |
| C21 | 2.6788810000 | 0.8648290000 | -3.7777370000 |
| H22 | 1.0286700000 | -0.1144120000 | 0.7406240000 |
| H23 | -0.0442110000 | 1.1163640000 | 0.0330060000 |
| H24 | -0.3677710000 | -0.6034440000 | -0.2549830000 |
| H25 | 1.1283990000 | 4.9732390000 | -1.4538760000 |
| H26 | 2.5755680000 | 4.0362230000 | -0.9937760000 |

| | | | |
|---|---|---|---|
| H27 | 0.1347810000 | -1.6503440000 | -2.8994160000 |
| H28 | 0.9561830000 | -3.9437990000 | -3.3707540000 |
| H29 | 3.1068960000 | -4.7442030000 | -2.4175560000 |
| H30 | 3.7864750000 | 0.0494390000 | -0.0219260000 |
| H31 | 5.7524880000 | -1.4520350000 | 0.5923860000 |
| H32 | 5.3990190000 | -3.8320810000 | -0.6493680000 |
| H33 | 1.7502740000 | 0.2905510000 | -3.8048660000 |
| H34 | 2.5362270000 | 1.8045220000 | -4.3184900000 |
| H35 | 3.4680680000 | 0.2811250000 | -4.2585760000 |

#### 20a-INT

| atom | x | y | z |
|---|---|---|---|
| C1 | 0.0414220000 | 0.4442620000 | 1.0074020000 |
| N2 | 1.4864640000 | 0.2728840000 | 0.8680080000 |
| C3 | 2.1742220000 | 1.5361400000 | 0.5020120000 |
| S4 | 2.0196680000 | 1.8944470000 | -1.3169500000 |
| C5 | 2.8251890000 | 0.6691920000 | -2.3449740000 |
| C6 | 4.0798570000 | 0.1151570000 | -2.0576100000 |
| C7 | 4.6584370000 | -0.7932000000 | -2.9438720000 |
| C8 | 4.0129030000 | -1.1430960000 | -4.1308250000 |
| C9 | 2.7694160000 | -0.5836900000 | -4.4235670000 |
| C10 | 2.1763650000 | 0.3128630000 | -3.5354420000 |
| O11 | 1.6525740000 | 2.5895690000 | 1.0783410000 |
| C12 | 1.9047040000 | -0.9257730000 | 0.1999580000 |
| C13 | 1.2238700000 | -1.5052940000 | -0.8977210000 |
| N14 | 0.0835350000 | -1.1215290000 | -1.5727580000 |
| C15 | -0.1614720000 | -2.0029340000 | -2.6011160000 |
| C16 | 0.7987930000 | -2.9776750000 | -2.6149880000 |
| C17 | 1.7041820000 | -2.6817540000 | -1.5388140000 |
| C18 | 2.8854070000 | -3.2975200000 | -1.0883120000 |
| C19 | 3.5454720000 | -2.7426340000 | -0.0049920000 |
| C20 | 3.0516120000 | -1.5858360000 | 0.6303640000 |
| O21 | 3.5396180000 | 1.2776610000 | 0.8331240000 |
| C22 | 4.4273790000 | 2.3689090000 | 0.6667580000 |
| H23 | -0.4166990000 | 0.9711800000 | 0.1560130000 |
| H24 | -0.4218300000 | -0.5399830000 | 1.1067470000 |
| H25 | -0.1807250000 | 1.0236160000 | 1.9045170000 |
| H26 | 4.5863610000 | 0.3628700000 | -1.1299800000 |
| H27 | 5.6247430000 | -1.2280690000 | -2.7039880000 |
| H28 | 4.4728100000 | -1.8474760000 | -4.8179440000 |
| H29 | 2.2505260000 | -0.8527330000 | -5.3394010000 |
| H30 | 1.1949000000 | 0.7256390000 | -3.7560490000 |
| H31 | -0.4678640000 | -0.2905430000 | -1.4071030000 |
| H32 | -1.0179170000 | -1.8558040000 | -3.2471680000 |
| H33 | 0.8645090000 | -3.8045930000 | -3.3092880000 |
| H34 | 3.2685030000 | -4.1859910000 | -1.5857990000 |
| H35 | 4.4582660000 | -3.2007650000 | 0.3668500000 |
| H36 | 3.5864490000 | -1.1627820000 | 1.4732390000 |
| H37 | 5.4073790000 | 2.0251550000 | 1.0046110000 |
| H38 | 4.4955630000 | 2.6640620000 | -0.3875370000 |
| H39 | 4.1159410000 | 3.2322410000 | 1.2622570000 |

#### 21a-INT

| atom | x | y | z |
|---|---|---|---|
| O1 | 3.5851270000 | -1.7672790000 | -0.9996960000 |
| C2 | 3.2811510000 | -0.4948830000 | -0.8935720000 |
| S3 | 4.6109060000 | 0.2839760000 | 0.1496280000 |
| C4 | 4.1424760000 | 1.9803590000 | 0.2136420000 |
| N5 | 3.3964640000 | 2.5334900000 | 1.1665050000 |

| | | | |
|---|---|---|---|
| C6 | 3.2964550000 | 3.8629770000 | 0.8377980000 |
| C7 | 3.9916750000 | 4.1158570000 | -0.3183680000 |
| N8 | 4.5306410000 | 2.9162620000 | -0.7032800000 |
| C9 | 5.3652880000 | 2.7329420000 | -1.8836920000 |
| N10 | 3.2334050000 | 0.3036760000 | -2.1381410000 |
| C11 | 4.1805990000 | -0.1596530000 | -3.1448870000 |
| C12 | 1.9497390000 | 0.6129530000 | -2.6830300000 |
| C13 | 0.9970630000 | -0.3754250000 | -2.9942700000 |
| N14 | 1.0537690000 | -1.7316800000 | -2.7929780000 |
| C15 | -0.1223450000 | -2.2949280000 | -3.2288360000 |
| C16 | -0.9523750000 | -1.3194680000 | -3.7259900000 |
| C17 | -0.2545690000 | -0.0693130000 | -3.5839630000 |
| C18 | -0.5613480000 | 1.2751730000 | -3.8840870000 |
| C19 | 0.3725220000 | 2.2532110000 | -3.5811550000 |
| C20 | 1.6111230000 | 1.9236960000 | -2.9816750000 |
| O21 | 2.0400440000 | -0.2043040000 | -0.2652710000 |
| C22 | 1.7140890000 | -1.0506190000 | 0.8180000000 |
| H23 | 2.7417590000 | 4.5618870000 | 1.4493440000 |
| H24 | 4.1572630000 | 5.0217330000 | -0.8865420000 |
| H25 | 5.8718220000 | 1.7721620000 | -1.8051720000 |
| H26 | 4.7513290000 | 2.7449570000 | -2.7883590000 |
| H27 | 6.1034820000 | 3.5366910000 | -1.9257090000 |
| H28 | 5.1618070000 | -0.2996190000 | -2.6815190000 |
| H29 | 3.8814130000 | -1.1125410000 | -3.6059690000 |
| H30 | 4.2699020000 | 0.5986880000 | -3.9283170000 |
| H31 | 1.8207090000 | -2.2126410000 | -2.3244600000 |
| H32 | -0.2783750000 | -3.3628470000 | -3.1498740000 |
| H33 | -1.9412950000 | -1.4706120000 | -4.1379830000 |
| H34 | -1.5132140000 | 1.5395500000 | -4.3388010000 |
| H35 | 0.1530920000 | 3.2954610000 | -3.7979260000 |
| H36 | 2.3255210000 | 2.7048690000 | -2.7336580000 |
| H37 | 1.5500520000 | -2.0787650000 | 0.4830900000 |
| H38 | 2.5063270000 | -1.0389820000 | 1.5765860000 |
| H39 | 0.7943760000 | -0.6556400000 | 1.2544840000 |

**22a-INT**

| atom | x | y | z |
|---|---|---|---|
| C1 | 0.7130890000 | -0.1265910000 | 0.1331970000 |
| N2 | 2.1653560000 | -0.1798050000 | 0.2404020000 |
| C3 | 2.7832120000 | 1.1000470000 | 0.1108310000 |
| C4 | 3.1086910000 | 1.8387250000 | 1.2366040000 |
| C5 | 3.6523940000 | 3.1406350000 | 1.1368330000 |
| C6 | 3.8883450000 | 3.7280830000 | -0.0958050000 |
| C7 | 3.5749230000 | 3.0007800000 | -1.2648200000 |
| C8 | 3.0262460000 | 1.7001050000 | -1.1388540000 |
| N9 | 2.8123790000 | 1.2012010000 | -2.4000000000 |
| C10 | 3.2130900000 | 2.1390450000 | -3.3221130000 |
| C11 | 3.6816640000 | 3.2591190000 | -2.6772160000 |
| C12 | 2.7611270000 | -1.3077320000 | -0.4924940000 |
| S13 | 2.5555190000 | -2.7905660000 | 0.6303120000 |
| C14 | 2.7992460000 | -2.1151990000 | 2.2311290000 |
| N15 | 3.9346190000 | -1.6677910000 | 2.7286220000 |
| N16 | 3.6754570000 | -1.1817150000 | 3.9844590000 |
| C17 | 2.3959620000 | -1.3650850000 | 4.2203950000 |
| N18 | 1.8009780000 | -1.9483340000 | 3.1550030000 |
| C19 | 0.3869550000 | -2.3016240000 | 3.0624280000 |
| O20 | 2.2368020000 | -1.6540550000 | -1.6415110000 |
| O21 | 4.1484770000 | -0.9968090000 | -0.5150640000 |
| C22 | 4.9157200000 | -1.7691730000 | -1.4216530000 |
| H23 | 0.3710540000 | 0.2264640000 | -0.8499130000 |

| | | | |
|---|---|---|---|
| H24 | 0.3041730000 | -1.1273520000 | 0.2979770000 |
| H25 | 0.3230960000 | 0.5417100000 | 0.9049610000 |
| H26 | 2.9348920000 | 1.3887150000 | 2.2103730000 |
| H27 | 3.8916870000 | 3.6820390000 | 2.0482980000 |
| H28 | 4.3091810000 | 4.7286450000 | -0.1608220000 |
| H29 | 2.4831060000 | 0.2541930000 | -2.5892380000 |
| H30 | 3.1298250000 | 1.9330520000 | -4.3822510000 |
| H31 | 4.0575190000 | 4.1574620000 | -3.1491350000 |
| H32 | 1.8725100000 | -1.1104140000 | 5.1332900000 |
| H33 | 0.0469410000 | -2.6194560000 | 4.0497770000 |
| H34 | -0.1928650000 | -1.4358090000 | 2.7316580000 |
| H35 | 0.2730630000 | -3.1227160000 | 2.3533710000 |
| H36 | 5.9574720000 | -1.4729920000 | -1.2779280000 |
| H37 | 4.8168490000 | -2.8408900000 | -1.2081180000 |
| H38 | 4.6179380000 | -1.5769230000 | -2.4564820000 |

#### 23a-INT

| atom | x | y | z |
|---|---|---|---|
| C1 | 0.7119340000 | -0.1133410000 | 0.2008450000 |
| N2 | 2.1687310000 | -0.1756100000 | 0.2565760000 |
| C3 | 2.7873430000 | 1.1070230000 | 0.1272350000 |
| C4 | 3.0574580000 | 1.8704570000 | 1.2535530000 |
| C5 | 3.5916790000 | 3.1764060000 | 1.1538070000 |
| C6 | 3.8788300000 | 3.7446590000 | -0.0774020000 |
| C7 | 3.6234530000 | 2.9936850000 | -1.2438280000 |
| C8 | 3.0764150000 | 1.6912230000 | -1.1208770000 |
| N9 | 2.9150550000 | 1.1772480000 | -2.3827540000 |
| C10 | 3.3452630000 | 2.1050000000 | -3.3000890000 |
| C11 | 3.7853170000 | 3.2343540000 | -2.6527510000 |
| C12 | 2.7431000000 | -1.2914180000 | -0.5055950000 |
| S13 | 2.4841820000 | -2.8207110000 | 0.5999990000 |
| C14 | 2.7635960000 | -2.1282600000 | 2.1736280000 |
| N15 | 3.8894190000 | -1.5807330000 | 2.6228090000 |
| N16 | 3.6060560000 | -1.0799560000 | 3.8304470000 |
| N17 | 2.3761470000 | -1.3070260000 | 4.1499290000 |
| N18 | 1.8264540000 | -1.9586180000 | 3.1293680000 |
| C19 | 0.4430780000 | -2.4139620000 | 3.2155310000 |
| O20 | 2.2207610000 | -1.6130570000 | -1.6542560000 |
| O21 | 4.1300690000 | -1.0207470000 | -0.4986330000 |
| C22 | 4.9025410000 | -1.8336090000 | -1.3637580000 |
| H23 | 0.3391930000 | 0.2232590000 | -0.7770910000 |
| H24 | 0.3005180000 | -1.1063470000 | 0.4021530000 |
| H25 | 0.3565050000 | 0.5748520000 | 0.9724760000 |
| H26 | 2.8446890000 | 1.4414220000 | 2.2302460000 |
| H27 | 3.7831330000 | 3.7371040000 | 2.0649260000 |
| H28 | 4.2921950000 | 4.7484550000 | -0.1428990000 |
| H29 | 2.5747120000 | 0.2355540000 | -2.5759020000 |
| H30 | 3.3030930000 | 1.8838450000 | -4.3585930000 |
| H31 | 4.1783870000 | 4.1274150000 | -3.1204940000 |
| H33 | 0.3782100000 | -3.1821200000 | 3.9896580000 |
| H34 | -0.1868220000 | -1.5582240000 | 3.4679340000 |
| H35 | 0.1601540000 | -2.8307530000 | 2.2487300000 |
| H36 | 5.9440280000 | -1.5447940000 | -1.2079060000 |
| H37 | 4.7822310000 | -2.8952420000 | -1.1128910000 |
| H38 | 4.6261410000 | -1.6737870000 | -2.4093750000 |

#### 24a-INT

| atom | x | y | z |
|---|---|---|---|
| C1 | -0.4737420000 | 0.6315770000 | -2.4384210000 |

| | | | |
|---|---|---|---|
| N2 | 0.9722250000 | 0.5950760000 | -2.2553490000 |
| C3 | 1.4948220000 | 1.8449960000 | -1.7974020000 |
| C4 | 1.9285470000 | 2.7965850000 | -2.7097790000 |
| C5 | 2.3889470000 | 4.0693160000 | -2.2977560000 |
| C6 | 2.4380980000 | 4.4173140000 | -0.9569690000 |
| C7 | 2.0094280000 | 3.4735300000 | 0.0005490000 |
| C8 | 1.5375150000 | 2.2100960000 | -0.4375230000 |
| N9 | 1.1768380000 | 1.4830610000 | 0.6680670000 |
| C10 | 1.4101010000 | 2.2362990000 | 1.7917790000 |
| C11 | 1.9164590000 | 3.4643070000 | 1.4359320000 |
| C12 | 1.4439860000 | -0.6375240000 | -1.5836500000 |
| Se13 | 1.5285410000 | -2.0361820000 | -3.0119840000 |
| C14 | 2.6852660000 | -1.0804070000 | -4.3176760000 |
| C15 | 4.1634240000 | -1.4163770000 | -4.1963210000 |
| O16 | 0.7188590000 | -1.1093370000 | -0.5952080000 |
| O17 | 2.7972250000 | -0.3084540000 | -1.2620810000 |
| C18 | 3.4272990000 | -1.1973870000 | -0.3655540000 |
| H19 | -0.7282180000 | 1.4446150000 | -3.1230280000 |
| H20 | -0.7980010000 | -0.3123620000 | -2.8830010000 |
| H21 | -1.0180720000 | 0.7791210000 | -1.4944160000 |
| H22 | 1.9059030000 | 2.5435770000 | -3.7673910000 |
| H23 | 2.7151800000 | 4.7793910000 | -3.0533830000 |
| H24 | 2.7998560000 | 5.3957150000 | -0.6495440000 |
| H25 | 0.8491110000 | 0.5162160000 | 0.6224880000 |
| H26 | 1.1936140000 | 1.8332530000 | 2.7726430000 |
| H27 | 2.1927520000 | 4.2628370000 | 2.1120820000 |
| H28 | 2.3038360000 | -1.3087870000 | -5.3148170000 |
| H29 | 2.4864070000 | -0.0262700000 | -4.1034550000 |
| H30 | 4.7451510000 | -0.8218180000 | -4.9124080000 |
| H31 | 4.5279630000 | -1.1941150000 | -3.1895850000 |
| H32 | 4.3500970000 | -2.4744580000 | -4.4058310000 |
| H33 | 3.4125570000 | -2.2211880000 | -0.7609140000 |
| H34 | 4.4627630000 | -0.8633570000 | -0.2691080000 |
| H35 | 2.9454460000 | -1.1819780000 | 0.6164250000 |

#### 25a-INT

| atom | x | y | z |
|---|---|---|---|
| C1 | -0.2699780000 | 0.3727490000 | 0.3956200000 |
| N2 | 1.1814140000 | 0.3782540000 | 0.5558470000 |
| C3 | 1.7421260000 | 1.6936850000 | 0.4916190000 |
| C4 | 1.7606420000 | 2.4701620000 | 1.6444190000 |
| C5 | 2.2509360000 | 3.7952120000 | 1.6533120000 |
| C6 | 2.7626660000 | 4.3769630000 | 0.5046940000 |
| C7 | 2.7699660000 | 3.6201810000 | -0.6846750000 |
| C8 | 2.2463820000 | 2.2993040000 | -0.6801120000 |
| N9 | 2.3541330000 | 1.8030600000 | -1.9554100000 |
| C10 | 2.9330870000 | 2.7503910000 | -2.7599630000 |
| C11 | 3.2053520000 | 3.8820180000 | -2.0293730000 |
| C12 | 1.8332300000 | -0.6779920000 | -0.2552750000 |
| O13 | 1.4643060000 | -0.8242650000 | -1.4842750000 |
| Se14 | 1.5779670000 | -2.3738400000 | 0.8594890000 |
| C15 | 0.2145280000 | -3.3797460000 | -0.0774030000 |
| C16 | -0.9920720000 | -3.6765150000 | 0.5680320000 |
| C17 | -1.9527200000 | -4.4669380000 | -0.0655700000 |
| C18 | -1.7210510000 | -4.9504570000 | -1.3542580000 |
| C19 | -0.5245590000 | -4.6402970000 | -2.0065540000 |
| C20 | 0.4439950000 | -3.8616260000 | -1.3703810000 |
| O21 | 3.2241720000 | -0.4315270000 | -0.0191040000 |
| C22 | 4.0971620000 | -1.0756830000 | -0.9268650000 |
| H23 | -0.7223270000 | 1.0373440000 | 1.1357020000 |

| | | | |
|---|---|---|---|
| H24 | -0.5832090000 | 0.6915280000 | -0.6087930000 |
| H25 | -0.6426160000 | -0.6418620000 | 0.5650610000 |
| H26 | 1.3844650000 | 2.0258540000 | 2.5629910000 |
| H27 | 2.2345740000 | 4.3552400000 | 2.5845500000 |
| H28 | 3.1499410000 | 5.3929900000 | 0.5176580000 |
| H29 | 2.0520950000 | 0.8622310000 | -2.2145860000 |
| H30 | 3.1083330000 | 2.5409590000 | -3.8072610000 |
| H31 | 3.6610750000 | 4.7902860000 | -2.4008510000 |
| H32 | -1.1731230000 | -3.2979940000 | 1.5707500000 |
| H33 | -2.8799210000 | -4.7020660000 | 0.4488350000 |
| H34 | -2.4654670000 | -5.5699120000 | -1.8463620000 |
| H35 | -0.3377580000 | -5.0177620000 | -3.0085920000 |
| H36 | 1.3852160000 | -3.6351820000 | -1.8670070000 |
| H37 | 5.1102180000 | -0.9086250000 | -0.5547520000 |
| H38 | 3.8925000000 | -2.1529860000 | -0.9616150000 |
| H39 | 4.0021630000 | -0.6586810000 | -1.9335960000 |

#### 26a-INT

| atom | x | y | z |
|---|---|---|---|
| C1 | 0.0834390000 | 0.1452060000 | 0.3492320000 |
| N2 | 1.4545110000 | -0.0166450000 | -0.1350350000 |
| C3 | 2.2383780000 | 1.2326570000 | -0.0091200000 |
| N4 | 1.6341060000 | 2.1091340000 | -1.1223860000 |
| C5 | 1.4739300000 | 1.7279340000 | -2.4307210000 |
| C6 | 0.9409560000 | 2.8133360000 | -3.0773700000 |
| N7 | 0.7729450000 | 3.8557640000 | -2.1859820000 |
| C8 | 1.2073290000 | 3.3827110000 | -1.0211860000 |
| O9 | 2.2304100000 | 1.8604870000 | 1.1326030000 |
| C10 | 2.0511640000 | -1.2301610000 | 0.3008210000 |
| C11 | 1.6559400000 | -2.4213880000 | -0.3433630000 |
| N12 | 0.7775080000 | -2.6026990000 | -1.3886460000 |
| C13 | 0.7124040000 | -3.9437010000 | -1.6928040000 |
| C14 | 1.5372190000 | -4.6504770000 | -0.8548360000 |
| C15 | 2.1593690000 | -3.6935770000 | 0.0218390000 |
| C16 | 3.1128740000 | -3.7830880000 | 1.0570940000 |
| C17 | 3.5167910000 | -2.6145520000 | 1.6830070000 |
| C18 | 2.9862030000 | -1.3553850000 | 1.3211970000 |
| O19 | 3.5210780000 | 0.8680280000 | -0.5242740000 |
| C20 | 4.4592730000 | 1.9293530000 | -0.5394590000 |
| H21 | -0.4926710000 | -0.7551170000 | 0.1229670000 |
| H22 | 0.0469220000 | 0.3193410000 | 1.4338760000 |
| H23 | -0.3861190000 | 0.9916430000 | -0.1596500000 |
| H24 | 1.7396190000 | 0.7353290000 | -2.7628130000 |
| H25 | 0.6667850000 | 2.9022930000 | -4.1199910000 |
| H26 | 1.2310170000 | 3.9198300000 | -0.0844000000 |
| H27 | 0.2898590000 | -1.8713800000 | -1.8942130000 |
| H28 | 0.0742020000 | -4.2902980000 | -2.4951380000 |
| H29 | 1.6878830000 | -5.7219100000 | -0.8604400000 |
| H30 | 3.5252400000 | -4.7453120000 | 1.3504480000 |
| H31 | 4.2548530000 | -2.6573560000 | 2.4796180000 |
| H32 | 3.3231630000 | -0.4590240000 | 1.8320570000 |
| H33 | 5.3801130000 | 1.5269260000 | -0.9666340000 |
| H34 | 4.1053650000 | 2.7576000000 | -1.1682290000 |
| H35 | 4.6583630000 | 2.3040210000 | 0.4688770000 |

#### 27a-INT

atom x y z

| | | | |
|---|---|---|---|
| O1 | 1.6981620000 | -1.8107670000 | -0.1203800000 |
| C2 | 2.3138630000 | -0.7031220000 | 0.1665380000 |
| N3 | -0.2314730000 | 1.6723850000 | -1.0901540000 |
| C4 | 0.7085010000 | 2.5085610000 | -0.5607320000 |
| N5 | 0.5834160000 | 3.9329970000 | -0.6268840000 |
| O6 | -0.4052080000 | 4.4088820000 | -1.1677410000 |
| O7 | 1.4852520000 | 4.6039810000 | -0.1359680000 |
| C8 | 1.7542150000 | 1.8210350000 | 0.0112650000 |
| N9 | 1.4399000000 | 0.5151690000 | -0.1845530000 |
| C10 | 0.2507430000 | 0.4693980000 | -0.8444930000 |
| N11 | 3.6215960000 | -0.4235660000 | -0.4865600000 |
| C12 | 4.4770960000 | -1.6169930000 | -0.4644080000 |
| C13 | 3.5925010000 | 0.1914780000 | -1.7871320000 |
| C14 | 2.7957580000 | -0.2575560000 | -2.8613710000 |
| N15 | 1.9055760000 | -1.3020940000 | -2.9326250000 |
| C16 | 1.3308580000 | -1.3170710000 | -4.1799970000 |
| C17 | 1.8476210000 | -0.3016730000 | -4.9465840000 |
| C18 | 2.7965680000 | 0.3963970000 | -4.1232760000 |
| C19 | 3.6420660000 | 1.5036580000 | -4.3323000000 |
| C20 | 4.4447400000 | 1.9313480000 | -3.2873270000 |
| C21 | 4.4090510000 | 1.2873090000 | -2.0312260000 |
| O22 | 2.6240830000 | -0.5217880000 | 1.5539110000 |
| C23 | 1.5410010000 | -0.8156820000 | 2.4256250000 |
| H24 | 2.6567530000 | 2.1486880000 | 0.5055640000 |
| H25 | -0.2137730000 | -0.4678960000 | -1.1155370000 |
| H26 | 5.4875830000 | -1.3302040000 | -0.7652060000 |
| H27 | 4.5130470000 | -2.0171210000 | 0.5515390000 |
| H28 | 4.1158810000 | -2.4034280000 | -1.1409000000 |
| H29 | 1.6594690000 | -1.8915130000 | -2.1371430000 |
| H30 | 0.5878720000 | -2.0638780000 | -4.4280980000 |
| H31 | 1.5858460000 | -0.0747580000 | -5.9714400000 |
| H32 | 3.6610380000 | 2.0119010000 | -5.2936790000 |
| H33 | 5.1044030000 | 2.7840970000 | -3.4237550000 |
| H34 | 5.0262310000 | 1.6549780000 | -1.2149180000 |
| H35 | 1.8802190000 | -0.5806040000 | 3.4364450000 |
| H36 | 0.6699790000 | -0.1937290000 | 2.1841260000 |
| H37 | 1.2552940000 | -1.8691370000 | 2.3666280000 |

#### 28a-INT

| atom | x | y | z |
|---|---|---|---|
| O1 | 0.7918020000 | -2.8397310000 | -2.4527000000 |
| C2 | -0.0663190000 | -2.9699380000 | -1.4856870000 |
| N3 | 0.6607220000 | -3.4678970000 | -0.2116700000 |
| N4 | -0.0042120000 | -3.8556920000 | 0.8972880000 |
| C5 | 0.9618140000 | -4.0085050000 | 1.7877620000 |
| N6 | 2.2080590000 | -3.7303530000 | 1.3166860000 |
| C7 | 1.9677270000 | -3.3855790000 | 0.0506850000 |
| N8 | -0.7878180000 | -1.7155540000 | -1.0853280000 |
| C9 | -2.1981480000 | -1.8708390000 | -0.7069200000 |
| C10 | -0.0537220000 | -0.8345540000 | -0.2312700000 |
| C11 | 1.1730140000 | -0.2531520000 | -0.6336970000 |
| N12 | 1.8779380000 | -0.3975330000 | -1.8059210000 |
| C13 | 3.0165370000 | 0.3666700000 | -1.7453860000 |
| C14 | 3.0589830000 | 1.0468240000 | -0.5535860000 |
| C15 | 1.8851090000 | 0.6656650000 | 0.1820910000 |
| C16 | 1.3733490000 | 1.0329330000 | 1.4424940000 |
| C17 | 0.1809050000 | 0.4625540000 | 1.8471930000 |
| C18 | -0.5131570000 | -0.4586570000 | 1.0308650000 |
| O19 | -1.0872840000 | -3.9247940000 | -1.7034790000 |
| C20 | -0.6058380000 | -5.1895180000 | -2.1423500000 |

| | | | |
|---|---|---|---|
| H21 | 0.7640850000 | -4.3315930000 | 2.8013870000 |
| H22 | 2.6954120000 | -3.0802520000 | -0.6882810000 |
| H23 | -2.6119730000 | -0.8758440000 | -0.5278860000 |
| H24 | -2.7475870000 | -2.3314770000 | -1.5250060000 |
| H25 | -2.3410060000 | -2.4801400000 | 0.1951220000 |
| H26 | 1.6616050000 | -1.1302200000 | -2.4831540000 |
| H27 | 3.7168840000 | 0.3671390000 | -2.5698840000 |
| H28 | 3.8288370000 | 1.7370540000 | -0.2343280000 |
| H29 | 1.9007730000 | 1.7451880000 | 2.0724960000 |
| H30 | -0.2436690000 | 0.7180310000 | 2.8147000000 |
| H31 | -1.4356060000 | -0.8800530000 | 1.4138300000 |
| H32 | 0.0988100000 | -5.6081730000 | -1.4128380000 |
| H33 | -1.4796810000 | -5.8409810000 | -2.2163290000 |
| H34 | -0.1150880000 | -5.1126460000 | -3.1173320000 |

#### 29a-INT

| atom | x | y | z |
|---|---|---|---|
| C1 | 0.2702900000 | -0.5805940000 | -0.4790290000 |
| N2 | 1.6014690000 | -0.0405520000 | -0.1931950000 |
| C3 | 1.7063330000 | 1.3731950000 | -0.3305710000 |
| C4 | 0.7210150000 | 2.1437200000 | -0.9495550000 |
| C5 | 0.8000640000 | 3.5524370000 | -1.0267740000 |
| C6 | 1.8744070000 | 4.2493680000 | -0.5060780000 |
| C7 | 2.9003170000 | 3.5084110000 | 0.1148900000 |
| C8 | 2.8020490000 | 2.0954350000 | 0.2016210000 |
| N9 | 3.9009900000 | 1.6429670000 | 0.8926590000 |
| C10 | 4.7050830000 | 2.7084010000 | 1.2050470000 |
| C11 | 4.1283950000 | 3.8742940000 | 0.7649530000 |
| C12 | 2.7735270000 | -0.8563830000 | -0.6534400000 |
| N13 | 3.3144660000 | -0.1362050000 | -1.9321370000 |
| N14 | 4.5186480000 | 0.3928620000 | -2.0171350000 |
| C15 | 4.5353490000 | 1.0003080000 | -3.2080150000 |
| C16 | 3.2876630000 | 0.8040370000 | -3.8119590000 |
| N17 | 2.5361310000 | 0.0823180000 | -2.9762250000 |
| O18 | 3.7272770000 | -1.0153510000 | 0.2109810000 |
| O19 | 2.2122760000 | -2.0542880000 | -1.1572780000 |
| C20 | 3.1771240000 | -2.9920510000 | -1.6167950000 |
| H21 | 0.0210330000 | -0.5642590000 | -1.5483150000 |
| H22 | 0.2032170000 | -1.6060620000 | -0.1220720000 |
| H23 | -0.4645830000 | 0.0208360000 | 0.0613830000 |
| H24 | -0.1389160000 | 1.6609750000 | -1.3988650000 |
| H25 | -0.0097770000 | 4.0877030000 | -1.5161260000 |
| H26 | 1.9273800000 | 5.3335200000 | -0.5676280000 |
| H27 | 4.1449850000 | 0.6505220000 | 0.9255240000 |
| H28 | 5.6363300000 | 2.5518220000 | 1.7332730000 |
| H29 | 4.5227250000 | 4.8735620000 | 0.8934570000 |
| H30 | 5.4091840000 | 1.5266570000 | -3.5642250000 |
| H31 | 2.9137360000 | 1.1349470000 | -4.7708250000 |
| H32 | 3.7867230000 | -2.5568460000 | -2.4180550000 |
| H33 | 3.8295250000 | -3.3223900000 | -0.8045450000 |
| H34 | 2.6152060000 | -3.8412530000 | -2.0109740000 |

#### 30a-INT

| atom | x | y | z |
|---|---|---|---|
| O1 | 0.3054190000 | -0.3091600000 | 0.2117820000 |
| C2 | 1.5631660000 | -0.1361250000 | -0.0117160000 |
| N3 | 1.8578280000 | 1.2933100000 | -0.5825090000 |
| C4 | 3.0465820000 | 1.8821750000 | -0.7527780000 |
| N5 | 2.8658110000 | 2.9915900000 | -1.4457090000 |

| | | | |
|---|---|---|---|
| N6 | 1.5409390000 | 3.0496670000 | -1.6943870000 |
| N7 | 0.9361750000 | 2.0306730000 | -1.1819070000 |
| N8 | 2.2581440000 | -1.0059050000 | -0.9938040000 |
| C9 | 1.9642070000 | -2.4218440000 | -0.7347270000 |
| C10 | 2.0829650000 | -0.6622960000 | -2.3853460000 |
| C11 | 0.8289500000 | -0.5560890000 | -3.0263040000 |
| N12 | -0.4422620000 | -0.6756280000 | -2.5210620000 |
| C13 | -1.3489120000 | -0.4538890000 | -3.5304620000 |
| C14 | -0.6894530000 | -0.2161350000 | -4.7099230000 |
| C15 | 0.7166910000 | -0.2666450000 | -4.4133770000 |
| C16 | 1.8811560000 | -0.0717910000 | -5.1808920000 |
| C17 | 3.1130170000 | -0.1558850000 | -4.5514520000 |
| C18 | 3.2065950000 | -0.4445720000 | -3.1719420000 |
| O19 | 2.4071510000 | -0.1806010000 | 1.1345430000 |
| C20 | 1.9044950000 | 0.5760360000 | 2.2344610000 |
| H21 | 3.9814900000 | 1.4909760000 | -0.3791330000 |
| H22 | 2.6218890000 | -3.0301870000 | -1.3602360000 |
| H23 | 2.1734700000 | -2.6481610000 | 0.3140330000 |
| H24 | 0.9219320000 | -2.6867970000 | -0.9559610000 |
| H25 | -0.6414340000 | -0.7047370000 | -1.5212910000 |
| H26 | -2.4108690000 | -0.4852740000 | -3.3245700000 |
| H27 | -1.1450480000 | -0.0236060000 | -5.6724210000 |
| H28 | 1.8116370000 | 0.1505350000 | -6.2432020000 |
| H29 | 4.0255770000 | 0.0049390000 | -5.1194780000 |
| H30 | 4.1820860000 | -0.4999680000 | -2.6947270000 |
| H31 | 2.6898320000 | 0.5807640000 | 2.9910770000 |
| H32 | 1.6958670000 | 1.6079710000 | 1.9262010000 |
| H33 | 0.9930400000 | 0.1324390000 | 2.6427010000 |

**31a-INT**

| atom | x | y | z |
|---|---|---|---|
| O1 | 0.5509070000 | 0.3819070000 | 0.0670380000 |
| C2 | 1.1566800000 | -0.7411560000 | 0.2960380000 |
| N3 | 2.5439260000 | -0.7726830000 | -0.4716150000 |
| C4 | 3.4596380000 | -1.7849480000 | -0.4967400000 |
| C5 | 3.5164310000 | -3.0665220000 | 0.0868110000 |
| C6 | 4.6418940000 | -3.8156430000 | -0.1953180000 |
| C7 | 5.6897820000 | -3.3262130000 | -1.0260960000 |
| C8 | 5.6361390000 | -2.0688610000 | -1.5911830000 |
| C9 | 4.4918100000 | -1.2941940000 | -1.3104880000 |
| N10 | 4.1387100000 | -0.0311150000 | -1.7182150000 |
| N11 | 2.9732450000 | 0.2429980000 | -1.1965010000 |
| N12 | 0.5437690000 | -1.9877400000 | -0.1657930000 |
| C13 | -0.0367470000 | -1.8888180000 | -1.5004440000 |
| C14 | -0.1925600000 | -2.7806690000 | 0.7588350000 |
| C15 | -1.1156360000 | -2.2519280000 | 1.6829880000 |
| N16 | -1.4877130000 | -0.9503420000 | 1.9111600000 |
| C17 | -2.4040400000 | -0.9205210000 | 2.9336800000 |
| C18 | -2.6484420000 | -2.1954060000 | 3.3814920000 |
| C19 | -1.8337010000 | -3.0756880000 | 2.5888600000 |
| C20 | -1.6551920000 | -4.4739740000 | 2.5460820000 |
| C21 | -0.7727430000 | -5.0012640000 | 1.6167060000 |
| C22 | -0.0452400000 | -4.1626930000 | 0.7434840000 |
| O23 | 1.5392400000 | -1.0113630000 | 1.6291160000 |
| C24 | 2.1119230000 | 0.0947690000 | 2.3089010000 |
| H25 | 2.7149520000 | -3.4326060000 | 0.7178360000 |
| H26 | 4.7362780000 | -4.8124210000 | 0.2267250000 |
| H27 | 6.5484820000 | -3.9642930000 | -1.2134880000 |
| H28 | 6.4311370000 | -1.6862050000 | -2.2252000000 |
| H29 | 0.6835450000 | -1.4172850000 | -2.1767840000 |

| | | | |
|---|---|---|---|
| H30 | -0.9681270000 | -1.3045540000 | -1.5189690000 |
| H31 | -0.2436800000 | -2.8945700000 | -1.8726580000 |
| H32 | -1.1222520000 | -0.1442470000 | 1.4065700000 |
| H33 | -2.8175540000 | 0.0225320000 | 3.2657030000 |
| H34 | -3.3248630000 | -2.4760010000 | 4.1773810000 |
| H35 | -2.2008040000 | -5.1240720000 | 3.2255260000 |
| H36 | -0.6197060000 | -6.0757630000 | 1.5611440000 |
| H37 | 0.6543000000 | -4.5967620000 | 0.0326150000 |
| H38 | 2.4340580000 | -0.2781630000 | 3.2827610000 |
| H39 | 2.9851800000 | 0.4812600000 | 1.7667070000 |
| H40 | 1.3834170000 | 0.8992140000 | 2.4466790000 |

**1a-LG**

| atom | x | y | z |
|---|---|---|---|
| O1 | 2.8899970000 | 2.8568130000 | -1.4941230000 |
| C2 | 1.7381120000 | 2.2209710000 | -1.5647850000 |
| C3 | 0.4786720000 | 2.8157640000 | -1.3546300000 |
| C4 | -0.7350020000 | 2.0910250000 | -1.4388870000 |
| C5 | -0.7710560000 | 0.7388360000 | -1.7357970000 |
| C6 | 0.4678510000 | 0.0892650000 | -1.9583630000 |
| C7 | 1.6697680000 | 0.8279290000 | -1.8705450000 |
| N8 | 2.7160370000 | -0.0353680000 | -2.1293180000 |
| C9 | 2.2124130000 | -1.2893120000 | -2.3750770000 |
| C10 | 0.8405210000 | -1.2622490000 | -2.2816010000 |
| H11 | 0.4477750000 | 3.8771840000 | -1.1178530000 |
| H12 | -1.6686470000 | 2.6222930000 | -1.2625210000 |
| H13 | -1.7111950000 | 0.1958800000 | -1.7961130000 |
| H14 | 3.6978720000 | 0.2179930000 | -2.1361780000 |
| H15 | 2.8744460000 | -2.1152210000 | -2.6003940000 |
| H16 | 0.1742420000 | -2.1039350000 | -2.4256190000 |

**2a-LG**

| atom | x | y | z |
|---|---|---|---|
| S1 | 3.1507030000 | 0.6324760000 | 0.7783470000 |
| C2 | 1.6051500000 | -0.2249520000 | 0.6851930000 |
| C3 | 1.5606550000 | -1.6031790000 | 0.3820530000 |
| N4 | 2.6045310000 | -2.4630640000 | 0.1250430000 |
| C5 | 2.1033970000 | -3.7192720000 | -0.1230780000 |
| C6 | 0.7334910000 | -3.6977590000 | -0.0314340000 |
| C7 | 0.3583380000 | -2.3477140000 | 0.2924200000 |
| C8 | -0.8736940000 | -1.6962770000 | 0.5147990000 |
| C9 | -0.8496040000 | -0.3432670000 | 0.8133800000 |
| C10 | 0.3638250000 | 0.3766120000 | 0.8966190000 |
| H11 | 3.5847380000 | -2.2011540000 | 0.1193790000 |
| H12 | 2.7719210000 | -4.5396760000 | -0.3483090000 |
| H13 | 0.0689880000 | -4.5394350000 | -0.1766830000 |
| H14 | -1.8123750000 | -2.2412320000 | 0.4532920000 |
| H15 | -1.7816540000 | 0.1884480000 | 0.9899980000 |
| H16 | 0.3335670000 | 1.4372270000 | 1.1334500000 |

**3a-LG**

| atom | x | y | z |
|---|---|---|---|
| Se1 | 3.3572380000 | 3.2118780000 | -1.4468340000 |
| C2 | 1.6767110000 | 2.2520620000 | -1.5551630000 |
| C3 | 0.4346530000 | 2.8468110000 | -1.3449650000 |
| C4 | -0.7744030000 | 2.1206410000 | -1.4298370000 |
| C5 | -0.7885780000 | 0.7669110000 | -1.7289830000 |
| C6 | 0.4467520000 | 0.1236240000 | -1.9499050000 |
| C7 | 1.6445930000 | 0.8783320000 | -1.8583700000 |

| | | | |
|---|---|---|---|
| N8 | 2.6941410000 | 0.0236900000 | -2.1147880000 |
| C9 | 2.2024400000 | -1.2358710000 | -2.3624610000 |
| C10 | 0.8323230000 | -1.2238460000 | -2.2730230000 |
| H11 | 0.3988750000 | 3.9066520000 | -1.1073510000 |
| H12 | -1.7104060000 | 2.6456030000 | -1.2543670000 |
| H13 | -1.7245740000 | 0.2173580000 | -1.7912900000 |
| H14 | 3.6711530000 | 0.2992270000 | -2.1163370000 |
| H15 | 2.8765510000 | -2.0526740000 | -2.5861240000 |
| H16 | 0.1757530000 | -2.0722130000 | -2.4195210000 |

## 4a-LG

| atom | x | y | z |
|---|---|---|---|
| Te1 | 2.1853750000 | -1.1399360000 | -4.6801550000 |
| C2 | 1.6978230000 | 0.3953940000 | -3.2860020000 |
| C3 | 2.0406300000 | 1.7339750000 | -3.4640260000 |
| C4 | 1.7029460000 | 2.7315380000 | -2.5211680000 |
| C5 | 1.0071230000 | 2.4262540000 | -1.3615400000 |
| C6 | 0.6390460000 | 1.0837900000 | -1.1393840000 |
| C7 | 0.9923970000 | 0.1034160000 | -2.1036910000 |
| N8 | 0.5207250000 | -1.1087500000 | -1.6497450000 |
| C9 | -0.1129290000 | -0.9281870000 | -0.4431970000 |
| C10 | -0.0660210000 | 0.3978120000 | -0.0900150000 |
| H11 | 2.5865210000 | 2.0233320000 | -4.3580810000 |
| H12 | 2.0006340000 | 3.7583010000 | -2.7204650000 |
| H13 | 0.7518150000 | 3.2008810000 | -0.6426460000 |
| H14 | 0.6321290000 | -1.9898240000 | -2.1405740000 |
| H15 | -0.5524860000 | -1.7711640000 | 0.0742600000 |
| H16 | -0.4824520000 | 0.8347230000 | 0.8084310000 |

## 6a-LG

| atom | x | y | z |
|---|---|---|---|
| F1 | 0.0000000000 | 0.0000000000 | 0.0000000000 |

## 7a-LG

| atom | x | y | z |
|---|---|---|---|
| O1 | 2.0437010000 | 1.0743060000 | -0.2524460000 |
| C2 | 2.5378530000 | -0.2094790000 | -0.2514570000 |
| C3 | 4.0575400000 | -0.2670090000 | -0.2524390000 |
| F4 | 4.5129090000 | -1.5442270000 | -0.2535420000 |
| F5 | 4.6085120000 | 0.3296590000 | -1.3325370000 |
| F6 | 4.6096450000 | 0.3289420000 | 0.8272110000 |
| H7 | 2.2515880000 | -0.8145700000 | -1.1335890000 |
| H8 | 2.2527230000 | -0.8123950000 | 0.6325190000 |

## 8a-LG

| atom | x | y | z |
|---|---|---|---|
| O1 | 2.0813630000 | -0.6508770000 | 1.2834480000 |
| C2 | 2.5508640000 | 0.0135940000 | 0.2009150000 |
| C3 | 4.0905070000 | 0.0162250000 | 0.1519570000 |
| F4 | 4.5506040000 | -1.2479160000 | 0.1441240000 |
| F5 | 4.5923140000 | 0.6159070000 | -0.9487260000 |
| F6 | 4.6513850000 | 0.6262670000 | 1.2126030000 |
| C7 | 2.0404360000 | 1.4663930000 | 0.1521200000 |
| F8 | 0.6955680000 | 1.4792360000 | 0.1412560000 |
| F9 | 2.4275100000 | 2.1985020000 | 1.2131540000 |
| F10 | 2.4397690000 | 2.1390260000 | -0.9483060000 |
| H11 | 2.2533190000 | -0.4074870000 | -0.7835480000 |

## 9a-LG

| atom | x | y | z |
|---|---|---|---|
| O1 | 0.0019230000 | 0.0045460000 | -1.8745380000 |
| C2 | 0.0005250000 | 0.0010830000 | -0.5433550000 |
| C3 | -1.4554550000 | -0.0457640000 | 0.0342250000 |
| F4 | -2.2042230000 | -0.9264210000 | -0.6356450000 |
| F5 | -1.5140050000 | -0.3873220000 | 1.3362750000 |
| F6 | -2.0519620000 | 1.1528470000 | -0.0858350000 |
| C7 | 0.7674060000 | -1.2374850000 | 0.0356210000 |
| F8 | 0.0207750000 | -2.3503900000 | -0.0724970000 |
| F9 | 1.8975590000 | -1.4561690000 | -0.6429360000 |
| F10 | 1.1015870000 | -1.1125910000 | 1.3344560000 |
| C11 | 0.6886600000 | 1.2833530000 | 0.0386870000 |
| F12 | 0.3066020000 | 2.3731990000 | -0.6336750000 |
| F13 | 0.4183160000 | 1.5058370000 | 1.3397120000 |
| F14 | 2.0257600000 | 1.1960110000 | -0.0768620000 |

## 10a-LG

| atom | x | y | z |
|---|---|---|---|
| O1 | 5.4839940000 | -0.6951060000 | 1.4705280000 |
| C2 | 4.3427430000 | -0.3744610000 | 0.9000960000 |
| C3 | 3.2176140000 | -1.2447060000 | 0.9200320000 |
| C4 | 2.0106010000 | -0.9010130000 | 0.3142310000 |
| C5 | 1.8555840000 | 0.3232670000 | -0.3424240000 |
| C6 | 2.9451650000 | 1.1983320000 | -0.3774440000 |
| C7 | 4.1564450000 | 0.8639450000 | 0.2251310000 |
| H8 | 3.3200600000 | -2.2024650000 | 1.4261510000 |
| H9 | 1.1756060000 | -1.5978200000 | 0.3537480000 |
| H10 | 0.9133240000 | 0.5877310000 | -0.8129160000 |
| H11 | 2.8466700000 | 2.1568330000 | -0.8833280000 |
| H12 | 4.9938940000 | 1.5575640000 | 0.1870470000 |

## 11a-LG

| atom | x | y | z |
|---|---|---|---|
| O1 | 0.0001690000 | -0.0001470000 | 2.8121200000 |
| C2 | -0.0000270000 | 0.0000550000 | 1.5195060000 |
| C3 | -0.0000840000 | 1.1833750000 | 0.7375240000 |
| F4 | -0.0000510000 | 2.3731300000 | 1.3751760000 |
| C5 | -0.0000320000 | 1.1905970000 | -0.6473770000 |
| F6 | 0.0000420000 | 2.3564310000 | -1.3137820000 |
| C7 | -0.0000150000 | -0.0000140000 | -1.3598300000 |
| F8 | 0.0000490000 | 0.0000050000 | -2.7050220000 |
| C9 | -0.0000430000 | -1.1906700000 | -0.6474390000 |
| F10 | 0.0000480000 | -2.3564940000 | -1.3138590000 |
| C11 | -0.0000750000 | -1.1833240000 | 0.7374480000 |
| F12 | -0.0000550000 | -2.3729620000 | 1.3752940000 |

## 12a-LG

| atom | x | y | z |
|---|---|---|---|
| O1 | 1.6160790000 | 1.4125920000 | 0.1316590000 |
| C2 | 2.2188320000 | 0.2807320000 | 0.0889580000 |
| N3 | 2.2396890000 | -0.5723570000 | 1.1229400000 |
| O4 | 2.9912920000 | -1.6915150000 | 0.7006100000 |
| C5 | 3.3995560000 | -1.4873380000 | -0.5647380000 |
| C6 | 2.9680160000 | -0.2950170000 | -1.0215180000 |
| H7 | 3.9892230000 | -2.2866750000 | -0.9973380000 |
| H8 | 3.1326630000 | 0.1485030000 | -1.9921240000 |

**13a-LG**

| atom | x | y | z |
|---|---|---|---|
| O1 | 3.5025840000 | 0.1694980000 | -1.9286960000 |
| C2 | 2.8317820000 | -0.3197690000 | -0.9565460000 |
| N3 | 1.5539840000 | -0.0307910000 | -0.6796180000 |
| S4 | 1.0720570000 | -0.8839520000 | 0.6660350000 |
| N5 | 2.5236980000 | -1.6665130000 | 0.9392060000 |
| C6 | 3.3623370000 | -1.2866370000 | 0.0209110000 |
| H7 | 4.3774420000 | -1.6716190000 | 0.0002690000 |

**14a-LG**

| atom | x | y | z |
|---|---|---|---|
| O1 | 2.8969370000 | -1.4251000000 | -1.0208460000 |
| N2 | 2.8851220000 | -0.3030310000 | -0.2679620000 |
| C3 | 2.8653920000 | 0.9817640000 | -0.7429800000 |
| C4 | 2.8609600000 | 1.7872060000 | 0.3765070000 |
| N5 | 2.8766060000 | 1.0172960000 | 1.5127080000 |
| C6 | 2.8916430000 | -0.2470350000 | 1.0776580000 |
| H7 | 2.8567980000 | 1.1898600000 | -1.8029340000 |
| H8 | 2.8477190000 | 2.8678990000 | 0.4184210000 |
| H9 | 2.9070380000 | -1.1408390000 | 1.6859700000 |

**15a-LG**

| atom | x | y | z |
|---|---|---|---|
| O1 | 1.5735520000 | -1.7364230000 | 0.1821430000 |
| N2 | 2.1680210000 | -0.5377050000 | 0.1781790000 |
| N3 | 1.4325500000 | 0.5774800000 | 0.1869190000 |
| C4 | 2.3276120000 | 1.5876150000 | 0.1818220000 |
| C5 | 3.6368960000 | 1.0997450000 | 0.1691380000 |
| C6 | 3.4884630000 | -0.2903870000 | 0.1671050000 |
| H7 | 1.9782480000 | 2.6116210000 | 0.1872920000 |
| H8 | 4.5584350000 | 1.6634830000 | 0.1625850000 |
| H9 | 4.1982020000 | -1.1047290000 | 0.1592150000 |

**16a-LG**

| atom | x | y | z |
|---|---|---|---|
| O1 | 0.2143590000 | 0.8705790000 | -1.3004690000 |
| N2 | 1.2354200000 | 1.3350140000 | -0.6063100000 |
| N3 | 1.1489190000 | 2.4573120000 | 0.0799050000 |
| N4 | 2.3006670000 | 2.7022330000 | 0.6847060000 |
| C5 | 3.1531730000 | 1.6888710000 | 0.3644870000 |
| C6 | 2.4745210000 | 0.7899290000 | -0.4752400000 |
| C7 | 3.0639790000 | -0.3772580000 | -0.9952930000 |
| C8 | 4.3749970000 | -0.5962780000 | -0.6283690000 |
| C9 | 5.0827360000 | 0.3055190000 | 0.2237670000 |
| C10 | 4.4975990000 | 1.4462030000 | 0.7290290000 |
| H11 | 2.5113840000 | -1.0521410000 | -1.6417400000 |
| H12 | 4.8913260000 | -1.4784680000 | -0.9944020000 |
| H13 | 6.1140410000 | 0.0772800000 | 0.4763160000 |
| H14 | 5.0346950000 | 2.1333700000 | 1.3774320000 |

**17a-LG**

| atom | x | y | z |
|---|---|---|---|
| O1 | -0.0010200000 | 0.0000730000 | 1.9530140000 |
| N2 | -0.0003730000 | -0.0000130000 | 0.6128020000 |
| C3 | 0.0001030000 | 1.1492220000 | -0.1453770000 |
| C4 | -0.0001270000 | 0.7677920000 | -1.6080310000 |
| C5 | -0.0000720000 | -0.7678020000 | -1.6080300000 |

| | | | |
|---|---|---|---|
| C6 | 0.0001350000 | -1.1492400000 | -0.1453900000 |
| O7 | 0.0007320000 | -2.2788440000 | 0.3336760000 |
| O8 | 0.0006300000 | 2.2788030000 | 0.3337440000 |
| H9 | 0.8843240000 | 1.1983470000 | -2.0869980000 |
| H10 | -0.8846570000 | 1.1985300000 | -2.0867370000 |
| H11 | 0.8843140000 | -1.1984810000 | -2.0870440000 |
| H12 | -0.8846910000 | -1.1983900000 | -2.0866820000 |

**18a-LG**

| atom | x | y | z |
|---|---|---|---|
| S1 | 1.7817570000 | 1.7669360000 | 0.3245810000 |
| C2 | 2.4498750000 | 0.0472190000 | 0.3286110000 |
| C3 | 3.9730710000 | 0.0004530000 | 0.3289890000 |
| H4 | 2.0694960000 | -0.4874560000 | -0.5474990000 |
| H5 | 2.0686580000 | -0.4839220000 | 1.2065750000 |
| H6 | 4.3316000000 | -1.0347410000 | 0.3277820000 |
| H7 | 4.3778540000 | 0.5043590000 | -0.5534490000 |
| H8 | 4.3777150000 | 0.5015570000 | 1.2132530000 |

**19a-LG**

| atom | x | y | z |
|---|---|---|---|
| S1 | 1.2641550000 | 1.6558080000 | -0.0008360000 |
| C2 | -0.4280790000 | 0.9617560000 | 0.0004910000 |
| C3 | -0.4570050000 | -0.5421270000 | 0.0003690000 |
| F4 | -1.7331970000 | -1.0040050000 | -0.0007610000 |
| F5 | 0.1359370000 | -1.0912600000 | -1.0802790000 |
| F6 | 0.1342460000 | -1.0915730000 | 1.0818100000 |
| H7 | -0.9937340000 | 1.2748260000 | -0.8832780000 |
| H8 | -0.9924450000 | 1.2749050000 | 0.8850590000 |

**20a-LG**

| atom | x | y | z |
|---|---|---|---|
| S1 | 6.0864280000 | -1.2751080000 | 0.1333040000 |
| C2 | 4.5375520000 | -0.4109510000 | 0.0668370000 |
| C3 | 4.4843540000 | 0.9954010000 | 0.0729040000 |
| C4 | 3.2674860000 | 1.6743940000 | 0.0215260000 |
| C5 | 2.0631080000 | 0.9696550000 | -0.0379630000 |
| C6 | 2.0980350000 | -0.4265960000 | -0.0453680000 |
| C7 | 3.3149560000 | -1.1054840000 | 0.0061290000 |
| H8 | 5.4125860000 | 1.5583670000 | 0.1185660000 |
| H9 | 3.2634510000 | 2.7615470000 | 0.0280490000 |
| H10 | 1.1155680000 | 1.4983100000 | -0.0780180000 |
| H11 | 1.1725380000 | -0.9952180000 | -0.0916670000 |
| H12 | 3.3249140000 | -2.1919990000 | -0.0005460000 |

**21a-LG**

| atom | x | y | z |
|---|---|---|---|
| S1 | 1.9748720000 | 0.3281810000 | -0.0000590000 |
| C2 | 0.3368560000 | -0.2730660000 | -0.0002060000 |
| N3 | -0.0420830000 | -1.5576030000 | -0.0006370000 |
| C4 | -1.4241340000 | -1.5470700000 | 0.0001020000 |
| C5 | -1.8856780000 | -0.2604300000 | 0.0005650000 |
| N6 | -0.7605970000 | 0.5398350000 | 0.0002810000 |
| C7 | -0.7523210000 | 1.9930540000 | 0.0000750000 |
| H8 | -1.9979330000 | -2.4633150000 | -0.0000590000 |
| H9 | -2.8823380000 | 0.1587150000 | 0.0007500000 |
| H10 | -1.7845120000 | 2.3471150000 | -0.0010990000 |
| H11 | -0.2392030000 | 2.3650790000 | 0.8898810000 |

| | | | |
|---|---|---|---|
| H12 | -0.2375510000 | 2.3644700000 | -0.8890150000 |

**22a-LG**

| atom | x | y | z |
|---|---|---|---|
| S1 | 2.8408520000 | 2.5301970000 | 0.2150640000 |
| C2 | 2.2413700000 | 0.9075170000 | 0.2165210000 |
| N3 | 0.9695830000 | 0.5154900000 | 0.2167210000 |
| N4 | 0.9473370000 | -0.8669590000 | 0.2175650000 |
| C5 | 2.1909080000 | -1.2733100000 | 0.2170540000 |
| N6 | 3.0356490000 | -0.2121810000 | 0.2170620000 |
| C7 | 4.4891790000 | -0.2437720000 | 0.2166420000 |
| H8 | 2.5226750000 | -2.3026380000 | 0.2170930000 |
| H9 | 4.8099720000 | -1.2866800000 | 0.2216270000 |
| H10 | 4.8632210000 | 0.2594840000 | -0.6779480000 |
| H11 | 4.8630380000 | 0.2679860000 | 1.1064530000 |

**23a-LG**

| atom | x | y | z |
|---|---|---|---|
| S1 | 2.0227630000 | 0.2586640000 | 0.0009180000 |
| C2 | 0.3792630000 | -0.2439780000 | -0.0001080000 |
| N3 | -0.0837480000 | -1.5063800000 | -0.0007700000 |
| N4 | -1.4241050000 | -1.4329220000 | -0.0006260000 |
| N5 | -1.8175270000 | -0.2084160000 | -0.0002310000 |
| N6 | -0.7188020000 | 0.5497880000 | -0.0001350000 |
| C7 | -0.8339080000 | 1.9969270000 | -0.0002020000 |
| H8 | 0.1765660000 | 2.4046700000 | -0.0013290000 |
| H9 | -1.3717680000 | 2.3064810000 | -0.9002980000 |
| H10 | -1.3699410000 | 2.3075090000 | 0.9006420000 |

**24a-LG**

| atom | x | y | z |
|---|---|---|---|
| Se1 | 1.0787550000 | 1.2258610000 | 0.3087690000 |
| C2 | 3.0750110000 | 1.4717870000 | 0.3080880000 |
| C3 | 3.8238510000 | 0.1492400000 | 0.3116860000 |
| H4 | 3.3517210000 | 2.0610500000 | 1.1861790000 |
| H5 | 3.3519310000 | 2.0561690000 | -0.5732040000 |
| H6 | 4.9087620000 | 0.3050750000 | 0.3115780000 |
| H7 | 3.5650380000 | -0.4422090000 | 1.1942680000 |
| H8 | 3.5654320000 | -0.4466240000 | -0.5680000000 |

**25a-LG**

| atom | x | y | z |
|---|---|---|---|
| Se1 | 0.2101490000 | -0.1574600000 | 1.3141100000 |
| C2 | 2.1499630000 | -0.0338200000 | 1.3159030000 |
| C3 | 2.8003980000 | 1.2105400000 | 1.3159490000 |
| C4 | 4.1927910000 | 1.3001160000 | 1.3172260000 |
| C5 | 4.9762490000 | 0.1446600000 | 1.3185580000 |
| C6 | 4.3443590000 | -1.1004030000 | 1.3184390000 |
| C7 | 2.9519850000 | -1.1865850000 | 1.3171960000 |
| H8 | 2.2043910000 | 2.1187730000 | 1.3149530000 |
| H9 | 4.6647720000 | 2.2795260000 | 1.3171780000 |
| H10 | 6.0599140000 | 0.2131160000 | 1.3196740000 |
| H11 | 4.9358800000 | -2.0126060000 | 1.3192990000 |
| H12 | 2.4744010000 | -2.1622750000 | 1.3172250000 |

**26a-LG**

| atom | x | y | z |
|---|---|---|---|

| | | | |
|---|---|---|---|
| N1 | 1.7581660000 | -2.0043180000 | 1.1619870000 |
| C2 | 0.5838970000 | -1.3445110000 | 0.8832890000 |
| C3 | 0.4315210000 | -1.2570000000 | -0.4856260000 |
| N4 | 1.5075310000 | -1.8593480000 | -1.0950100000 |
| C5 | 2.2557660000 | -2.2813850000 | -0.0578130000 |
| H6 | -0.0694770000 | -0.9791420000 | 1.6652650000 |
| H7 | -0.3717620000 | -0.8058380000 | -1.0538110000 |
| H8 | 3.1933790000 | -2.8076620000 | -0.1955500000 |

## 27a-LG

| atom | x | y | z |
|---|---|---|---|
| N1 | -0.4705110000 | 1.6519310000 | -0.0004890000 |
| C2 | 0.2746210000 | 0.5093180000 | -0.0003230000 |
| N3 | 1.6906550000 | 0.5214900000 | 0.0000180000 |
| O4 | 2.2807460000 | 1.5998760000 | 0.0002340000 |
| O5 | 2.2778720000 | -0.5636540000 | 0.0003260000 |
| C6 | -0.5436400000 | -0.6145880000 | -0.0000590000 |
| N7 | -1.8245970000 | -0.1836610000 | 0.0000320000 |
| C8 | -1.7133010000 | 1.1736020000 | 0.0001030000 |
| H9 | -0.2640090000 | -1.6593640000 | 0.0000460000 |
| H10 | -2.5816960000 | 1.8210730000 | 0.0004820000 |

## 28a-LG

| atom | x | y | z |
|---|---|---|---|
| N1 | 1.8157040000 | -1.0605970000 | 1.0786670000 |
| C2 | 1.1488470000 | -2.2082470000 | 1.0100220000 |
| N3 | 0.6206430000 | -2.4563520000 | -0.2080200000 |
| C4 | 1.0141090000 | -1.3635480000 | -0.8974250000 |
| N5 | 1.7272600000 | -0.5050440000 | -0.1757540000 |
| H6 | 1.0462290000 | -2.8726750000 | 1.8578280000 |
| H7 | 0.7777360000 | -1.1915850000 | -1.9392190000 |

## 29a-LG

| atom | x | y | z |
|---|---|---|---|
| N1 | 0.0000390000 | -0.0000070000 | -1.1197690000 |
| N2 | -0.0001580000 | 1.0967460000 | -0.3593070000 |
| C3 | -0.0000070000 | 0.6931810000 | 0.9259880000 |
| C4 | -0.0000050000 | -0.6931600000 | 0.9259600000 |
| N5 | 0.0001260000 | -1.0967600000 | -0.3593360000 |
| H6 | 0.0001290000 | 1.3938200000 | 1.7463480000 |
| H7 | -0.0000960000 | -1.3937800000 | 1.7463450000 |

## 30a-LG

| atom | x | y | z |
|---|---|---|---|
| N1 | 0.4542450000 | -0.8242760000 | 0.0741020000 |
| C2 | 1.3911280000 | -1.4624260000 | 0.7766670000 |
| N3 | 1.8627930000 | -2.5185330000 | 0.1122410000 |
| N4 | 1.1899470000 | -2.5217670000 | -1.0443680000 |
| N5 | 0.3545960000 | -1.5170000000 | -1.0665560000 |
| H6 | 1.7238940000 | -1.1634290000 | 1.7565450000 |

## 31a-LG

| atom | x | y | z |
|---|---|---|---|
| N1 | 3.3035390000 | -1.5219990000 | -0.1352580000 |
| C2 | 1.9920540000 | -1.1501650000 | -0.1199090000 |
| C3 | 0.8043990000 | -1.9139320000 | -0.1125240000 |
| C4 | -0.3849030000 | -1.2149710000 | -0.0963330000 |

| | | | |
|---|---|---|---|
| C5 | -0.4178550000 | 0.2109560000 | -0.0879680000 |
| C6 | 0.7376070000 | 0.9647300000 | -0.0955900000 |
| C7 | 1.9594320000 | 0.2571300000 | -0.1115630000 |
| N8 | 3.2523310000 | 0.6894530000 | -0.1217330000 |
| N9 | 4.0125210000 | -0.3992910000 | -0.1325430000 |
| H10 | 0.8358140000 | -3.0007890000 | -0.1196630000 |
| H11 | -1.3270580000 | -1.7556450000 | -0.0899310000 |
| H12 | -1.3841860000 | 0.7070000000 | -0.0754300000 |
| H13 | 0.7183350000 | 2.0518360000 | -0.0897620000 |

**26a-TS**

| atom | x | y | z |
|---|---|---|---|
| C1 | -0.5308325679 | 0.3403357899 | -0.0654062200 |
| N2 | 0.8930853953 | 0.1097294839 | 0.1875051498 |
| C3 | 1.8629589225 | 0.8461951844 | -0.4679089422 |
| N4 | 1.4447818381 | 2.2137538428 | -0.7910116976 |
| C5 | 0.5186449804 | 2.7073997449 | -1.6855725679 |
| C6 | 0.5270077710 | 4.0633498094 | -1.5107108254 |
| N7 | 1.4337672846 | 4.4159400815 | -0.5272088049 |
| C8 | 1.9647042624 | 3.2789513990 | -0.1221045079 |
| O9 | 3.0722518207 | 0.6841062805 | -0.2467828854 |
| C10 | 1.2732003797 | -1.1392634802 | 0.7675300816 |
| C11 | 1.9973427372 | -2.1077569586 | 0.0498383020 |
| N12 | 2.4444015948 | -2.0669945296 | -1.2397055892 |
| C13 | 3.1060788191 | -3.2361740999 | -1.5070861000 |
| C14 | 3.1007904471 | -4.0530069741 | -0.3977081057 |
| C15 | 2.3783595000 | -3.3488294215 | 0.6262096763 |
| C16 | 1.9840840361 | -3.6398990835 | 1.9476281507 |
| C17 | 1.2425199864 | -2.6971820745 | 2.6447218207 |
| C18 | 0.9003786997 | -1.4541426038 | 2.0664171105 |
| O19 | 1.5443389492 | 0.1552594203 | -2.3621278369 |
| C20 | 2.4985408202 | 0.7484015322 | -3.1900114503 |
| H21 | -1.1025771989 | -0.3620686732 | 0.5409756128 |
| H22 | -0.8153297409 | 1.3570606677 | 0.2160986246 |
| H23 | -0.7736780523 | 0.1763645384 | -1.1212406958 |
| H24 | -0.0158487070 | 2.0598942452 | -2.3621605554 |
| H25 | -0.0558507954 | 4.8043232722 | -2.0390964580 |
| H26 | 2.7188219292 | 3.1600780289 | 0.6445605722 |
| H27 | 2.1999111304 | -1.2501314817 | -1.8623420726 |
| H28 | 3.5417179377 | -3.4047299246 | -2.4842283056 |
| H29 | 3.5511768555 | -5.0336256327 | -0.3219531219 |
| H30 | 2.2471365746 | -4.5883390628 | 2.4096613478 |
| H31 | 0.9237093344 | -2.9055769649 | 3.6620397423 |
| H32 | 0.3362720423 | -0.7236198580 | 2.6414767604 |
| H33 | 2.6954130275 | 0.1441342526 | -4.0894912878 |
| H34 | 2.1905435113 | 1.7511318751 | -3.5330721446 |
| H35 | 3.4643166108 | 0.8729119896 | -2.6689360447 |

**28a-TS**

| atom | x | y | z |
|---|---|---|---|
| C1 | -2.4518110456 | -0.1812068377 | -0.9900672787 |
| C2 | -2.4126817921 | -1.0573231509 | 0.1084435651 |
| C3 | -3.1539847551 | -2.2676040286 | 0.1326363142 |
| C4 | -3.9861148407 | -2.5849815982 | -0.9589392827 |
| C5 | -4.0339886859 | -1.7148920482 | -2.0378811525 |
| C6 | -3.2696248781 | -0.5258004753 | -2.0540691757 |
| C7 | -2.7996310045 | -2.9221463888 | 1.3629685241 |
| C8 | -1.8857334698 | -2.1053270131 | 1.9904707560 |

| | | | |
|---|---|---|---|
| N9 | -1.6693834707 | -0.9715068445 | 1.2491031590 |
| N10 | -1.7645526435 | 1.0818333158 | -1.0044364050 |
| C11 | -2.4987969499 | 2.1665846243 | -0.3465869559 |
| C12 | -0.3846432245 | 1.2322286601 | -1.0412713851 |
| O13 | 0.1432899225 | 2.3371584906 | -1.2812039497 |
| O14 | 0.0796651758 | 0.9456772160 | 0.8534314241 |
| C15 | 1.4283825513 | 0.5968105613 | 0.9655985078 |
| N16 | 1.6814348950 | -1.0259952306 | -2.9204710677 |
| C17 | 1.0120543422 | -1.8401332929 | -2.0482795771 |
| N18 | 0.1883326328 | -1.2116450113 | -1.2391803818 |
| N19 | 0.3256260509 | 0.0801258889 | -1.6092415090 |
| C20 | 1.2243241832 | 0.1761109393 | -2.6086516101 |
| H21 | -4.5743247549 | -3.4999150193 | -0.9608168160 |
| H22 | -4.6665551236 | -1.9451060439 | -2.8907048252 |
| H23 | -3.3179800477 | 0.1425414941 | -2.9098619384 |
| H24 | -3.1694937684 | -3.8709253169 | 1.7293758741 |
| H25 | -1.3702817378 | -2.2501337724 | 2.9317739409 |
| H26 | -0.9032201662 | -0.2440537901 | 1.3128747839 |
| H27 | -3.5106059042 | 2.2038952245 | -0.7549639071 |
| H28 | -2.5549459337 | 2.0004067060 | 0.7363393700 |
| H29 | -2.0003252625 | 3.1155491499 | -0.5409123075 |
| H30 | 1.8251404275 | 0.8339801021 | 1.9634254723 |
| H31 | 1.5930253516 | -0.4798214231 | 0.7938674094 |
| H32 | 2.0483055687 | 1.1409669211 | 0.2324614745 |
| H33 | 1.1423582647 | -2.9137297898 | -2.0124440596 |
| H34 | 1.4952966263 | 1.1173760871 | -3.0650604092 |

#### 1a-4a_common-END

| atom | x | y | z |
|---|---|---|---|
| O1 | 1.2572790000 | -1.0054500000 | -0.0004250000 |
| C2 | 0.2287250000 | -0.3229330000 | 0.0000220000 |
| N3 | 0.1962220000 | 1.0254300000 | -0.0000150000 |
| C4 | -1.0256930000 | 1.8234250000 | -0.0001130000 |
| C5 | 1.4377350000 | 1.7867170000 | -0.0000470000 |
| O6 | -1.0000230000 | -0.8791050000 | 0.0007060000 |
| C7 | -1.0467770000 | -2.3141450000 | -0.0000840000 |
| H8 | -1.9035180000 | 1.1820770000 | -0.0000460000 |
| H9 | -1.0458850000 | 2.4632160000 | 0.8899720000 |
| H10 | -1.0458170000 | 2.4627200000 | -0.8905320000 |
| H11 | 2.2874280000 | 1.1067870000 | 0.0003140000 |
| H12 | 1.4824250000 | 2.4235510000 | -0.8909230000 |
| H13 | 1.4823100000 | 2.4241370000 | 0.8904250000 |
| H14 | -2.1075550000 | -2.5649150000 | -0.0001190000 |
| H15 | -0.5621980000 | -2.7105040000 | -0.8949200000 |
| H16 | -0.5621130000 | -2.7116000000 | 0.8942230000 |

#### 5a-END

| atom | x | y | z |
|---|---|---|---|
| S1 | 0.0179010000 | 0.1655160000 | -0.1100710000 |
| C2 | 0.9436060000 | -1.2590470000 | -0.1103500000 |
| N3 | 2.2751420000 | -1.3112160000 | -0.1104430000 |
| C4 | 3.0314150000 | -2.5667870000 | -0.1105750000 |
| C5 | 3.1237110000 | -0.1233450000 | -0.1110410000 |
| O6 | 0.3843050000 | -2.4704840000 | -0.1106840000 |
| C7 | -1.0442260000 | -2.6207690000 | -0.1117460000 |
| H8 | 2.3634020000 | -3.4223160000 | -0.1085920000 |
| H9 | 3.6655580000 | -2.5950730000 | -1.0025060000 |
| H10 | 3.6686470000 | -2.5933140000 | 0.7792070000 |
| H11 | 2.5125740000 | 0.7745830000 | -0.1118110000 |

| | | | |
|---|---|---|---|
| H12 | 3.7591470000 | -0.1383810000 | 0.7809390000 |
| H13 | 3.7593310000 | -0.1398790000 | -1.0028710000 |
| H14 | -1.2062430000 | -3.6982480000 | -0.1138980000 |
| H15 | -1.4728730000 | -2.1658570000 | 0.7827870000 |
| H16 | -1.4719890000 | -2.1624830000 | -1.0049530000 |

**Table1_common-END**

| atom | x | y | z |
|---|---|---|---|
| O1 | -1.4089750000 | 0.8300980000 | 1.0661720000 |
| C2 | -0.8356740000 | -0.2536340000 | 0.9734720000 |
| C3 | 1.1892930000 | -1.6291540000 | 0.8069230000 |
| C4 | 1.7937990000 | -2.1772580000 | -0.3356710000 |
| N5 | 1.7842490000 | -1.7500450000 | -1.6415280000 |
| C6 | 2.5464830000 | -2.6156640000 | -2.3975910000 |
| C7 | 3.0605540000 | -3.6056630000 | -1.6002530000 |
| C8 | 2.5895750000 | -3.3498980000 | -0.2651860000 |
| C9 | 2.7539820000 | -3.9973220000 | 0.9746620000 |
| C10 | 2.1386780000 | -3.4628490000 | 2.0974040000 |
| C11 | 1.3724820000 | -2.2801030000 | 2.0174830000 |
| N12 | 0.4890300000 | -0.3856720000 | 0.7037450000 |
| C13 | 1.2979770000 | 0.8336050000 | 0.5943710000 |
| O14 | -1.4643430000 | -1.4315590000 | 1.1065290000 |
| C15 | -2.8696990000 | -1.3639280000 | 1.4020690000 |
| H16 | 1.2620090000 | -0.9635860000 | -2.0155370000 |
| H17 | 2.6607010000 | -2.4508690000 | -3.4609420000 |
| H18 | 3.6982900000 | -4.4200050000 | -1.9194460000 |
| H19 | 3.3551390000 | -4.9007960000 | 1.0495640000 |
| H20 | 2.2559630000 | -3.9483830000 | 3.0621260000 |
| H21 | 0.9176930000 | -1.8633470000 | 2.9126390000 |
| H22 | 2.2314650000 | 0.5822830000 | 0.0864590000 |
| H23 | 0.7618780000 | 1.5837930000 | 0.0117540000 |
| H24 | 1.5280610000 | 1.2393360000 | 1.5865000000 |
| H25 | -3.1946040000 | -2.3999520000 | 1.4925330000 |
| H26 | -3.0338070000 | -0.8299810000 | 2.3403830000 |
| H27 | -3.4044990000 | -0.8661860000 | 0.5907690000 |

**1a_ M06-2X/LACV3P++****

| atom | x | y | z |
|---|---|---|---|
| O1 | 0.0298796183 | -0.0338724258 | 0.1666226995 |
| C2 | 1.2408711604 | -0.0488059292 | 0.0117675396 |
| N3 | 2.0292099013 | 1.0000707563 | -0.2614996907 |
| C4 | 3.4627925297 | 0.8530260886 | -0.5181393817 |
| C5 | 1.4302963199 | 2.2973620299 | -0.5539005687 |
| O6 | 1.9759988131 | -1.1987806320 | 0.1050126944 |
| C7 | 1.3124866415 | -2.3702382871 | 0.4449432404 |
| C8 | 0.9927559476 | -3.2777248590 | -0.5664942005 |
| N9 | 1.1669222082 | -3.2091890046 | -1.9260651326 |
| C10 | 0.6934652919 | -4.3748683745 | -2.4870473403 |
| C11 | 0.2111531678 | -5.2036818985 | -1.5104492861 |
| C12 | 0.3925703549 | -4.5227765904 | -0.2576838037 |
| C13 | 0.1166527751 | -4.8418520276 | 1.0848437868 |
| C14 | 0.4425657209 | -3.9291766129 | 2.0722534022 |
| C15 | 1.0426032562 | -2.6926617515 | 1.7564532502 |
| H16 | 3.6474013301 | 0.4766121772 | -1.5293779360 |
| H17 | 3.9129868801 | 0.1755866308 | 0.2045511431 |
| H18 | 3.9244482649 | 1.8348904555 | -0.4156445321 |
| H19 | 1.5125647325 | 2.5207777773 | -1.6219180736 |
| H20 | 0.3816657939 | 2.2866985646 | -0.2688296306 |
| H21 | 1.9508278929 | 3.0716474257 | 0.0138883678 |

| | | | |
|---|---|---|---|
| H22 | 1.5902842643 | -2.4462510813 | -2.4437154868 |
| H23 | 0.7406997428 | -4.5195501637 | -3.5566009032 |
| H24 | -0.2210972539 | -6.1824137110 | -1.6596997357 |
| H25 | -0.3404318499 | -5.7921941040 | 1.3417081153 |
| H26 | 0.2421245645 | -4.1617539012 | 3.1122523875 |
| H27 | 1.3034628269 | -1.9866127270 | 2.5373519210 |

#### 1a-TS_ M06-2X/LACV3P++**

| atom | x | y | z |
|---|---|---|---|
| C1 | -2.0753321692 | -0.5535238365 | 1.1094673399 |
| C2 | -2.9812999634 | -0.1499776201 | 0.1179956797 |
| C3 | -4.3532940601 | -0.4971522829 | 0.1776367767 |
| C4 | -4.8276073315 | -1.2487116058 | 1.2707994264 |
| C5 | -3.9306704705 | -1.6293762212 | 2.2514720398 |
| C6 | -2.5612341793 | -1.2911460989 | 2.1705149308 |
| C7 | -4.9563468435 | 0.0828586565 | -0.9901729646 |
| C8 | -3.9562666006 | 0.7379574521 | -1.6683027234 |
| N9 | -2.7677575868 | 0.5986532380 | -1.0030391578 |
| O10 | -0.7522290492 | -0.1700928596 | 1.1115260538 |
| C11 | 0.0433086175 | -0.5202256566 | 0.0002160137 |
| N12 | 1.3856570236 | -0.5059696958 | 0.3987612949 |
| C13 | 1.9017425794 | 0.6775134169 | 1.0712017097 |
| C14 | 2.3480501115 | -1.1277799704 | -0.5023312480 |
| O15 | -0.1571586948 | 1.2083648013 | -0.8317470929 |
| O16 | -0.3732393921 | -1.3529007934 | -0.8166759411 |
| C17 | 0.5121458067 | 1.2061226930 | -2.0567846114 |
| H18 | -5.8765089517 | -1.5167971116 | 1.3454895073 |
| H19 | -4.2750738550 | -2.2017521356 | 3.1056148825 |
| H20 | -1.8692181161 | -1.5993766844 | 2.9466386292 |
| H21 | -5.9936718240 | 0.0224813132 | -1.2848629424 |
| H22 | -4.0036901987 | 1.3000520182 | -2.5901957137 |
| H23 | -1.8207407637 | 0.9816968001 | -1.1946405817 |
| H24 | 2.7905060958 | 0.4016278379 | 1.6454299158 |
| H25 | 2.1754507199 | 1.4588732429 | 0.3508422437 |
| H26 | 1.1553069351 | 1.0776703695 | 1.7537686822 |
| H27 | 3.2430248657 | -1.3795588526 | 0.0721838169 |
| H28 | 1.9341873468 | -2.0457418146 | -0.9147003007 |
| H29 | 2.6450325699 | -0.4687841942 | -1.3270455487 |
| H30 | 0.0348550003 | 1.8805755353 | -2.7828695796 |
| H31 | 1.5632613292 | 1.5227391923 | -1.9639943521 |
| H32 | 0.5156331043 | 0.2034205232 | -2.5116097050 |

#### 2a_ M06-2X/LACV3P++**

| atom | x | y | z |
|---|---|---|---|
| O1 | 0.3244211629 | -0.0781394626 | 0.3348812479 |
| C2 | 1.4697701895 | 0.0827219389 | -0.0574609370 |
| N3 | 2.0032756374 | 1.2383808219 | -0.4794720537 |
| C4 | 1.2021141234 | 2.4627261561 | -0.4566613623 |
| C5 | 3.3785111634 | 1.3494424461 | -0.9687476403 |
| S6 | 2.6626806134 | -1.3035537300 | -0.0937469284 |
| C7 | 1.6308633836 | -2.6331700121 | 0.4614509080 |
| C8 | 0.9554206127 | -3.4346325891 | -0.4691300423 |
| N9 | 0.8951373276 | -3.3605881877 | -1.8362861520 |
| C10 | 0.1055762009 | -4.3877385677 | -2.3053012963 |
| C11 | -0.3578729438 | -5.1355506601 | -1.2576580694 |
| C12 | 0.1749379883 | -4.5464399227 | -0.0600676891 |
| C13 | 0.0802566054 | -4.8508887155 | 1.3075299649 |
| C14 | 0.7549166561 | -4.0604163417 | 2.2238338897 |
| C15 | 1.5276977973 | -2.9613726106 | 1.8044479479 |

| | | | |
|---|---|---|---|
| H16 | 0.2505466953 | 2.2664904923 | 0.0288307945 |
| H17 | 1.7396766057 | 3.2346696663 | 0.0990520885 |
| H18 | 1.0254127277 | 2.8105594802 | -1.4763968730 |
| H19 | 3.5592552515 | 0.6550871993 | -1.7934422737 |
| H20 | 3.5230114929 | 2.3634590144 | -1.3384583399 |
| H21 | 4.1002841580 | 1.1651824940 | -0.1680807034 |
| H22 | 1.3787106624 | -2.6840592207 | -2.4163405880 |
| H23 | -0.0622363293 | -4.5010467514 | -3.3665951180 |
| H24 | -0.9992592183 | -6.0020673700 | -1.3244450853 |
| H25 | -0.5084980992 | -5.6998917187 | 1.6414600349 |
| H26 | 0.6965653980 | -4.2886168908 | 3.2823324186 |
| H27 | 2.0574532140 | -2.3605408795 | 2.5358716558 |

**2a-TS_ M06-2X/LACV3P++****

| atom | x | y | z |
|---|---|---|---|
| O1 | 0.1876314841 | -0.1095753120 | -0.9936888223 |
| C2 | 1.1038645153 | -0.4701009537 | -0.2506624482 |
| N3 | 1.5950544340 | 0.3182634308 | 0.7498166655 |
| C4 | 1.2638888573 | 1.7347862434 | 0.6461998122 |
| C5 | 2.9015223843 | 0.0482664422 | 1.3303187349 |
| S6 | 1.2443593629 | -2.3164658672 | 0.0614126456 |
| C7 | 0.7174220461 | -2.4485204631 | 1.7557590299 |
| C8 | -0.4867793914 | -1.9011885688 | 2.2291261845 |
| N9 | -1.4473440102 | -1.1670549614 | 1.5808263976 |
| C10 | -2.4459974286 | -0.8450657253 | 2.4720465947 |
| C11 | -2.1547894915 | -1.3667016618 | 3.7031215538 |
| C12 | -0.9005858668 | -2.0559715121 | 3.5760558272 |
| C13 | -0.1037946748 | -2.8004493080 | 4.4627897033 |
| C14 | 1.0726197015 | -3.3561347631 | 3.9936418318 |
| C15 | 1.4842634591 | -3.1716734573 | 2.6587181672 |
| O16 | 2.8105349469 | -0.5152111518 | -1.5098619025 |
| C17 | 2.3095597911 | -0.9581441318 | -2.7270925809 |
| H18 | 1.5690907989 | 2.2248366990 | 1.5718862648 |
| H19 | 1.7817032948 | 2.2072213057 | -0.1988707973 |
| H20 | 0.1889132985 | 1.8572926456 | 0.5169845463 |
| H21 | 3.0837434031 | -1.0220642750 | 1.3629115792 |
| H22 | 3.6891502890 | 0.5211055947 | 0.7339146492 |
| H23 | 2.9280655978 | 0.4422229673 | 2.3488782839 |
| H24 | -1.4193938390 | -0.8907202685 | 0.6062975497 |
| H25 | -3.2963755145 | -0.2625636385 | 2.1491382929 |
| H26 | -2.7567649078 | -1.2762027630 | 4.5956749081 |
| H27 | -0.4097960611 | -2.9370317450 | 5.4953576687 |
| H28 | 1.7018118025 | -3.9374658464 | 4.6587093611 |
| H29 | 2.4205229844 | -3.6040614041 | 2.3221021001 |
| H30 | 3.0626279620 | -0.9382486004 | -3.5306090369 |
| H31 | 1.9363415697 | -1.9999417667 | -2.6838503888 |
| H32 | 1.4582924880 | -0.3548730420 | -3.0836987696 |

**3a_M06-2X/LACV3P++****

| atom | x | y | z |
|---|---|---|---|
| O1 | 1.5489600573 | -2.3931017636 | -0.2240572393 |
| C2 | 0.9561990494 | -1.3287627321 | -0.3199713403 |
| N3 | 0.2323327384 | -0.9468797522 | -1.3756821067 |
| C4 | 0.0313469462 | -1.9196652876 | -2.4546892563 |
| C5 | -0.2387287705 | 0.4075783906 | -1.6608981617 |
| Se6 | 1.0936080684 | -0.1116295360 | 1.2699383181 |
| C7 | -0.7455536300 | 0.4905623065 | 1.4142350255 |
| C8 | -1.7679136151 | -0.4288315570 | 1.6785731169 |
| N9 | -1.7279985574 | -1.7875454932 | 1.8653792071 |

| | | | |
|---|---|---|---|
| C10 | -3.0060669815 | -2.2470683603 | 2.0950133601 |
| C11 | -3.8899130027 | -1.2046643487 | 2.0497117464 |
| C12 | -3.1217004430 | -0.0188058376 | 1.7884125699 |
| C13 | -3.4384061588 | 1.3430037188 | 1.6583546677 |
| C14 | -2.4189013485 | 2.2522526378 | 1.4314541250 |
| C15 | -1.0810987043 | 1.8301226264 | 1.3057803995 |
| H18 | 0.8289328223 | -1.8345179981 | -3.1989558824 |
| H19 | 0.0235029630 | -2.9281174161 | -2.0467649601 |
| H20 | -0.9284343352 | -1.7091063697 | -2.9275731367 |
| H21 | 0.3016910408 | 1.1339930826 | -1.0605997181 |
| H22 | -0.0448634559 | 0.6166141247 | -2.7154923794 |
| H23 | -1.3114104648 | 0.4965521326 | -1.4684892031 |
| H24 | -0.8914033898 | -2.3562317670 | 1.9164614308 |
| H25 | -3.1785238319 | -3.2980595872 | 2.2749504367 |
| H26 | -4.9592122915 | -1.2666410779 | 2.1892975746 |
| H27 | -4.4679737494 | 1.6769635397 | 1.7429472797 |
| H28 | -2.6448048578 | 3.3088660380 | 1.3379393581 |
| H29 | -0.3048371505 | 2.5617044163 | 1.1065419382 |

**3a-TS_ M06-2X/LACV3P++****

| atom | x | y | z |
|---|---|---|---|
| O1 | 1.3782556116 | -2.3400728488 | -0.1695063186 |
| C2 | 1.0821898109 | -1.1583355750 | -0.3887020193 |
| N3 | 0.0885415835 | -0.8072040909 | -1.2657495048 |
| C4 | -0.3453794744 | -1.8474077621 | -2.1875599609 |
| C5 | 0.0150281382 | 0.5394644954 | -1.8077208720 |
| Se6 | 1.2262492275 | 0.0672589021 | 1.1921502552 |
| C7 | -0.6438182848 | 0.5584506126 | 1.3870316619 |
| C8 | -1.6517534583 | -0.4042006509 | 1.5379926268 |
| N9 | -1.5803550359 | -1.7738249836 | 1.5633160138 |
| C10 | -2.8463394660 | -2.2900285733 | 1.7233776678 |
| C11 | -3.7543435876 | -1.2703154463 | 1.8109550657 |
| C12 | -3.0138700368 | -0.0438360763 | 1.6985732366 |
| C13 | -3.3660548928 | 1.3160926650 | 1.7313677770 |
| C14 | -2.3693457941 | 2.2664655345 | 1.5999165800 |
| C15 | -1.0217891567 | 1.8912440400 | 1.4213183146 |
| O16 | 2.6960370264 | -0.3838671481 | -1.3543413986 |
| C17 | 3.8036263394 | -0.8212583478 | -0.6359331696 |
| H18 | 0.4187165557 | -2.0623162977 | -2.9453412132 |
| H19 | -0.5692561931 | -2.7617749134 | -1.6398408302 |
| H20 | -1.2523784510 | -1.5019242442 | -2.6849768288 |
| H21 | 0.4582782667 | 1.2480447816 | -1.1149119457 |
| H22 | 0.5569976144 | 0.6007740522 | -2.7580117048 |
| H23 | -1.0316804526 | 0.8082575376 | -1.9683989139 |
| H24 | -0.7330938304 | -2.3192403216 | 1.4627133587 |
| H25 | -2.9971577028 | -3.3587608342 | 1.7654004383 |
| H26 | -4.8215489461 | -1.3745770946 | 1.9435225066 |
| H27 | -4.4025401531 | 1.6129663824 | 1.8591860939 |
| H28 | -2.6188930698 | 3.3216521275 | 1.6247065254 |
| H29 | -0.2657869292 | 2.6603213121 | 1.3030220354 |
| H30 | 3.7624428901 | -1.9007401867 | -0.4157996599 |
| H31 | 4.7478807552 | -0.6459712640 | -1.1728192582 |
| H32 | 3.9102996547 | -0.3117962920 | 0.3412522733 |

**4a_ M06-2X/LACV3P++****

| atom | x | y | z |
|---|---|---|---|
| O1 | 1.3891279940 | 6.4871720258 | 0.9738165890 |
| C2 | 2.1531519244 | 5.6271566539 | 1.4069451737 |
| N3 | 3.3152802162 | 5.8920881439 | 2.0132626390 |

| | | | |
|---|---|---|---|
| C4 | 4.3685671670 | 4.9226136969 | 2.2990451163 |
| C5 | 3.6579596890 | 7.2936998851 | 2.2760911687 |
| Te6 | 1.4256951813 | 3.5873190332 | 1.1094986249 |
| C7 | 2.2438852710 | 2.5619533580 | 2.7723285126 |
| C8 | 1.7851147598 | 2.8032975918 | 4.0752337255 |
| N9 | 0.8413196579 | 3.6825477675 | 4.5445341294 |
| C10 | 0.7345988996 | 3.5470038942 | 5.9101295304 |
| C11 | 1.6095832220 | 2.5934409547 | 6.3522490962 |
| C12 | 2.2984161492 | 2.0952044353 | 5.1946342752 |
| C13 | 3.2805127903 | 1.1130101261 | 4.9930203412 |
| C14 | 3.7199676607 | 0.8580103998 | 3.7047660137 |
| C15 | 3.2069570990 | 1.5757189928 | 2.6068741752 |
| H18 | 5.3274612561 | 5.3733840199 | 2.0353206001 |
| H19 | 4.3803058627 | 4.6529074402 | 3.3584540000 |
| H20 | 4.2337001679 | 4.0281270691 | 1.6987268040 |
| H21 | 4.2288918050 | 7.7122958909 | 1.4418358804 |
| H22 | 2.7499386146 | 7.8755702385 | 2.4196879687 |
| H23 | 4.2632674659 | 7.3315387297 | 3.1820705542 |
| H24 | 0.2575401494 | 4.2813682877 | 3.9739024478 |
| H25 | 0.0300670781 | 4.1529640834 | 6.4603555712 |
| H26 | 1.7501733009 | 2.2786648582 | 7.3761996255 |
| H27 | 3.6824678649 | 0.5578646216 | 5.8355866957 |
| H28 | 4.4726872237 | 0.0973752477 | 3.5280168840 |
| H29 | 3.5817386502 | 1.3570792615 | 1.6122757236 |

#### 4a-TS_ M06-2X/LACV3P++**

| atom | x | y | z |
|---|---|---|---|
| O1 | 3.3918180981 | 5.6606137502 | 3.5825400183 |
| C2 | 2.7707026384 | 5.5874798001 | 2.4807182481 |
| N3 | 3.0164264764 | 6.5239526038 | 1.4809009758 |
| C4 | 2.8450734215 | 6.2810602583 | 0.0566394064 |
| C5 | 3.6352573786 | 7.7826756948 | 1.8621232140 |
| Te6 | 2.5267860753 | 3.6683495402 | 1.6484436513 |
| C7 | 2.2685754689 | 2.5141623511 | 3.3940992122 |
| C8 | 1.1941371535 | 2.7215498970 | 4.2709269504 |
| N9 | 0.3049366661 | 3.7604134575 | 4.3178151001 |
| C10 | -0.5987032227 | 3.5150378613 | 5.3211263111 |
| C11 | -0.3287226241 | 2.3114824190 | 5.9227095927 |
| C12 | 0.8342515264 | 1.7791269603 | 5.2693096629 |
| C13 | 1.5956873215 | 0.6048376194 | 5.4053453924 |
| C14 | 2.6803564218 | 0.4120447716 | 4.5670809681 |
| C15 | 3.0125352826 | 1.3549987325 | 3.5703656927 |
| O16 | 1.0546936500 | 6.0232599233 | 3.0459702937 |
| C17 | 0.1889500494 | 6.3560645597 | 2.0007217411 |
| H18 | 2.6750579843 | 7.2342636250 | -0.4483025035 |
| H19 | 3.7258325659 | 5.8089498226 | -0.4000124220 |
| H20 | 1.9658481949 | 5.6667436335 | -0.1525691897 |
| H21 | 3.3083646005 | 8.5609549185 | 1.1688643019 |
| H22 | 3.3224773401 | 8.0617675815 | 2.8672014105 |
| H23 | 4.7308711396 | 7.7314901307 | 1.8369088950 |
| H24 | 0.4702393974 | 4.6687812550 | 3.8199534568 |
| H25 | -1.3820735939 | 4.2293017197 | 5.5323921983 |
| H26 | -0.8881503733 | 1.8596729907 | 6.7286484151 |
| H27 | 1.3358694181 | -0.1386785915 | 6.1527719449 |
| H28 | 3.2841932405 | -0.4844633869 | 4.6603286924 |
| H29 | 3.8547378557 | 1.1526825320 | 2.9168754197 |
| H30 | 0.5732525246 | 7.1945749805 | 1.4020852248 |
| H31 | 0.0341018782 | 5.5062602353 | 1.3141519523 |
| H32 | -0.7972912752 | 6.6469960037 | 2.3847230333 |

**$MeO^-$_ M06-2X/LACV3P++****

| atom | x | y | z |
|---|---|---|---|
| C1 | 0.0000128532 | 0.0003502358 | 0.6799681450 |
| O2 | -0.0000145356 | -0.0002965139 | -0.7215255720 |
| H3 | 1.0133700166 | 0.0003604029 | 1.1183589606 |
| H4 | -0.5064874091 | 0.8777665943 | 1.1186866852 |
| H5 | -0.5068039568 | -0.8775913033 | 1.1178369705 |

#### Supplementary References

1. Hoffmann, M. *et al.* Highly Selective Butyrylcholinesterase Inhibitors with Tunable Duration of Action by Chemical Modification of Transferable Carbamate Units Exhibit Pronounced Neuroprotective Effect in an Alzheimer's Disease Mouse Model. *J. Med. Chem.* **62**, 9116–9140 (2019).

2. Groner, E. *et al.* The Kinetics of Inhibition of Human Acetylcholinesterase and Butyrylcholinesterase by Two Series of Novel Carbamates. *Mol. Pharmacol.* **71**, 1610–1617 (2007).

3. Scheiner, M. *et al.* Photoswitchable Pseudoirreversible Butyrylcholinesterase Inhibitors Allow Optical Control of Inhibition in Vitro and Enable Restoration of Cognition in an Alzheimer's Disease Mouse Model upon Irradiation. *J. Am. Chem. Soc.* **144**, 3279–3284 (2022).

4. Darras, F. H., Kling, B., Sawatzky, E., Heilmann, J. & Decker, M. Cyclic acyl guanidines bearing carbamate moieties allow potent and dirigible cholinesterase inhibition of either acetyl- or butyrylcholinesterase. *Bioorg. Med. Chem.* **22**, 5020–5034 (2014).

5. Sawatzky, E. *et al.* Discovery of Highly Selective and Nanomolar Carbamate-Based Butyrylcholinesterase Inhibitors by Rational Investigation into Their Inhibition Mode. *J. Med. Chem.* **59**, 2067–2082 (2016).

6. Yanovsky, I. *et al.* Carbamate Derivatives of Indolines as Cholinesterase Inhibitors and Antioxidants for the Treatment of Alzheimer's Disease. *J. Med. Chem.* **55**, 10700–10715 (2012).

7. Venkatasubban, K. S. *et al.* Decarbamoylation of acetylcholinesterases is markedly slowed as carbamoyl groups increase in size. *Arch. Biochem. Biophys.* **655**, 67–74 (2018).

8. Perola, E., Cellai, L., Lamba, D., Filocamo, L. & Brufani, M. Long chain analogs of physostigmine as potential drugs for Alzheimer's disease: new insights into the mechanism of action in the inhibition of acetylcholinesterase. *Biochim. Biophys. Acta BBA - Protein Struct. Mol. Enzymol.* **1343**, 41–50 (1997).

9. Scheindlin, S. Episodes in the Story of Physostigmine. *Mol. Interv.* **10**, 4 (2010).

10. Bartolucci, C., Stojan, J., Yu, Q., Greig, N. H. & Lamba, D. Kinetics of Torpedo californica acetylcholinesterase inhibition by bisnorcymserine and crystal structure of the complex with its leaving group. *Biochem. J.* **444**, 269–277 (2012).

11. Darvesh, S. *et al.* Carbamates with Differential Mechanism of Inhibition Toward Acetylcholinesterase and Butyrylcholinesterase. *J. Med. Chem.* **51**, 4200–4212 (2008).

12. Kamal, M. A. *et al.* Kinetics of Human Serum Butyrylcholinesterase Inhibition by a Novel Experimental Alzheimer Therapeutic, Dihydrobenzodioxepine Cymserine. *Neurochem. Res.* **33**, 745–753 (2008).

13. Bolognesi, M. L. *et al.* Design, Synthesis, and Biological Evaluation of Conformationally Restricted Rivastigmine Analogues. *J. Med. Chem.* **47**, 5945–5952 (2004).

14. Yu, Q. *et al.* Anticholinesterase Activity of Compounds Related to Geneserine Tautomers. N-Oxides and 1,2-Oxazines. *J. Med. Chem.* **45**, 3684–3691 (2002).

15. Zhang, H. *et al.* Recent advance on carbamate-based cholinesterase inhibitors as potential multifunctional agents against Alzheimer's disease. *Eur. J. Med. Chem.* **240**, 114606 (2022).

16. Lieske, C. N. *et al.* Anticholinesterase Activity of Potential Therapeutic 5-(L,3,3-Trimethylindolinyl) Carbamates. *J. Enzym. Inhib.* **5**, 215–223 (1991).

17. Jiang, X. *et al.* Novel cannabidiol−carbamate hybrids as selective BuChE inhibitors: Docking-based fragment reassembly for the development of potential therapeutic agents against Alzheimer's disease. *Eur. J. Med. Chem.* **223**, 113735 (2021).

18. Mazej, T., Knez, D., Meden, A., Gobec, S. & Sova, M. 4-Phenethyl-1-Propargylpiperidine-Derived Dual Inhibitors of Butyrylcholinesterase and Monoamine Oxidase B. *Molecules* **26**, 4118 (2021).

19. Wang, Y. *et al.* Discovery of carbamate-based N-salicyloyl tryptamine derivatives as novel pleiotropic agents for the treatment of Alzheimer's disease. *Bioorganic Chem.* **127**, 105993 (2022).

20. Rudolph, S. *et al.* Aryl N-[ω-(6-Fluoroindol-1-yl)alkyl]carbamates as Inhibitors of Fatty Acid Amide Hydrolase, Monoacylglycerol Lipase, and Butyrylcholinesterase: Structure–Activity Relationships and Hydrolytic Stability. *ACS Omega* **6**, 13466–13483 (2021).

21. Verma, A. *et al.* 3-Oxoisoxazole-2(3H)-carboxamides and isoxazol-3-yl carbamates: Resistance-breaking acetylcholinesterase inhibitors targeting the malaria mosquito, Anopheles gambiae. *Bioorg. Med. Chem.* **23**, 1321–1340 (2015).

22. Bajda, M., Łątka, K., Hebda, M., Jończyk, J. & Malawska, B. Novel carbamate derivatives as selective butyrylcholinesterase inhibitors. *Bioorganic Chem.* **78**, 29–38 (2018).

23. Metcalf, R. L. Structure—activity relationships for insecticidal carbamates*. *Bull. World Health Organ.* **44**, 43–78 (1971).

24. Krátký, M., Štěpánková, Š., Vorčáková, K., Švarcová, M. & Vinšová, J. Novel Cholinesterase Inhibitors Based on O-Aromatic N,N-Disubstituted Carbamates and Thiocarbamates. *Molecules* **21**, 191 (2016).

25. Buccafusco, J. J. & Smith, M. D. In vivo and in vitro cholinesterase inhibitor property of the antitumor agent caracemide. *Res. Commun. Chem. Pathol. Pharmacol.* **67**, 219–227 (1990).

26. Fukuto, T. R., Metcalf, R. L., Jones, R. L. & Myers, R. O. Structure, reactivity, and biological activity of O-(diethyl phosphoryl)oximes and O-(methylcarbamoyl)oximes of substituted acetophenones and .alpha.-substituted benzaldehydes. *J. Agric. Food Chem.* **17**, 923–930 (1969).

27. Myers, D. K. & Kemp, A. Inhibition of Esterases by the Fluorides of Organic Acids. *Nature* **173**, 33–34 (1954).

28. Bieth, J., Vratsanos, S. M., Wassermann, N. & Erlanger, B. F. Photoregulation of biological activity by photocromic reagents, ii. inhibitors of acetylcholinesterase*†. *Proc. Natl. Acad. Sci.* **64**, 1103–1106 (1969).

29. Metzger, H. P. & Wilson, I. B. Evidence for an Electrophilic Mechanism in Catalysis by Hydrolytic Enzymes*. *Biochemistry* **3**, 926–931 (1964).

30. Loudwig, S. *et al.* Photoreversible Inhibition of Cholinesterases: Catalytic Serine-Labeled Caged Butyrylcholinesterase. *ChemBioChem* **4**, 762–767 (2003).

31. Pejchal, V. *et al.* Synthesis, structural characterization, docking, lipophilicity and cytotoxicity of 1-[(1R)-1-(6-fluoro-1,3-benzothiazol-2-yl)ethyl]-3-alkyl carbamates, novel acetylcholinesterase and butyrylcholinesterase pseudo-irreversible inhibitors. *Bioorg. Med. Chem.* **24**, 1560–1572 (2016).

32. Carolan, C. G. *et al.* Isosorbide-2-benzyl Carbamate-5-salicylate, A Peripheral Anionic Site Binding Subnanomolar Selective Butyrylcholinesterase Inhibitor. *J. Med. Chem.* **53**, 1190–1199 (2010).

33. Gialih Lin, Gan-Hong Chen, & Hong-Chi Ho. Conformationally restricted carbamate inhibitors of horse serum butyrylcholinesterase. *Bioorg. Med. Chem. Lett.* **8**, 2747–2750 (1998).

34. Kovářová, M., Komers, K., Štěpánková, Š. & Čegan, A. Inhibition of acetylcholinesterase by 14 achiral and five chiral imidazole derivates. *Bioresour. Technol.* **101**, 6281–6283 (2010).

35. Matošević, A. *et al.* Quinuclidine-Based Carbamates as Potential CNS Active Compounds. *Pharmaceutics* **13**, 420 (2021).

36. Lock, E. A. Inhibition of rat brain and human red cell acetylcholinesterase by thiocarbamate herbicides. *Toxicol. Res.* **9**, 591–600 (2020).

37. Fallah, A., Mohanazadeh, F. & Safavi, M. Design, synthesis, and in vitro evaluation of novel 1,3,4-oxadiazolecarbamothioate derivatives of Rivastigmine as selective inhibitors of BuChE. *Med. Chem. Res.* **29**, 341–355 (2020).

38. Kitz, R. & Wilson, I. B. Esters of Methanesulfonic Acid as Irreversible Inhibitors of Acetylcholinesterase. *J. Biol. Chem.* **237**, 3245–3249 (1962).

39. Moss, D. E. Improving Anti-Neurodegenerative Benefits of Acetylcholinesterase Inhibitors in Alzheimer's Disease: Are Irreversible Inhibitors the Future? *Int. J. Mol. Sci.* **21**, 3438 (2020).

40. Moss, D. E., Perez, R. G. & Kobayashi, H. Cholinesterase Inhibitor Therapy in Alzheimer's Disease: The Limits and Tolerability of Irreversible CNS-Selective Acetylcholinesterase Inhibition in Primates. *J. Alzheimers Dis.* **55**, 1285–1294 (2017).

41. Moss, D. E. *et al.* A randomized phase I study of methanesulfonyl fluoride, an irreversible cholinesterase inhibitor, for the treatment of Alzheimer's disease. *Br. J. Clin. Pharmacol.* **75**, 1231–1239 (2013).

42. Kraut, D. *et al.* Inactivation Studies of Acetylcholinesterase with Phenylmethylsulfonyl Fluoride. *Mol. Pharmacol.* **57**, 1243–1248 (2000).

43. Worek, F. *et al.* Kinetic analysis of reactivation and aging of human acetylcholinesterase inhibited by different phosphoramidates. *Biochem. Pharmacol.* **73**, 1807–1817 (2007).

44. Charpentier, A., Menozzi, P., Marcel, V., Villatte, F. & Fournier, D. A Method to Estimate Acetylcholinesterase-Active Sites and Turnover in Insects. *Anal. Biochem.* **285**, 76–81 (2000).

45. Baici, A., Schenker, P., Wächter, M. & Rüedi, P. 3-Fluoro-2,4-dioxa-3-phosphadecalins as Inhibitors of Acetylcholinesterase. A Reappraisal of Kinetic Mechanisms and Diagnostic Methods. *Chem. Biodivers.* **6**, 261–282 (2009).

46. KOVARIK, Z. *et al.* Acetylcholinesterase active centre and gorge conformations analysed by combinatorial mutations and enantiomeric phosphonates. *Biochem. J.* **373**, 33–40 (2003).

47. Krátký, M. *et al.* 2-Hydroxy-N-phenylbenzamides and Their Esters Inhibit Acetylcholinesterase and Butyrylcholinesterase. *Biomolecules* **9**, 698 (2019).

48. Berman, H. A. & Taylor, P. Fluorescent phosphonate label for serine hydrolases, pyrenebutyl methylphosphonofluoridate: reaction with acetylcholinesterase. *Biochemistry* **17**, 1704–1713 (1978).

49. Kaur, J. & Singh, P. K. Enzyme-based optical biosensors for organophosphate class of pesticide detection. *Phys. Chem. Chem. Phys.* **22**, 15105–15119 (2020).

50. Liu, W. & Tsou, C. Determination of rate constant for the irreversible inhibition of acetylcholine esterase by continuously monitoring the substrate reaction in the presence of the inhibitor. *Biochim. Biophys. Acta BBA - Protein Struct. Mol. Enzymol.* **870**, 185–190 (1986).

51. Gholivand, K. *et al.* Acetylcholinesterase Inhibition by Diaza- and Dioxophosphole Compounds: Synthesis and Determination of IC50 Values. *J. Enzyme Inhib. Med. Chem.* **19**, 403–407 (2004).

52. Fiore, M. F. *et al.* Guanitoxin, re-naming a cyanobacterial organophosphate toxin. *Harmful Algae* **92**, 101737 (2020).

53. Jokanovic, M. & Johnson, M. K. Interactions in vitro of some organophosphoramidates with neuropathy target esterase and acetylcholinesterase of hen brain. *J. Biochem. Toxicol.* **8**, 19–31 (1993).

54. Costanzi, S., Machado, J.-H. & Mitchell, M. Nerve Agents: What They Are, How They Work, How to Counter Them. *ACS Chem. Neurosci.* **9**, 873–885 (2018).

55. Haux, J. E., Quistad, G. B. & Casida, J. E. Phosphobutyrylcholinesterase: Phosphorylation of the Esteratic Site of Butyrylcholinesterase by Ethephon [(2-Chloroethyl)phosphonic Acid] Dianion. *Chem. Res. Toxicol.* **13**, 646–651 (2000).

56. Quistad, G. B., Zhang, N., Sparks, S. E. & Casida, J. E. Phosphoacetylcholinesterase: Toxicity of Phosphorus Oxychloride to Mammals and Insects That Can Be Attributed to Selective Phosphorylation of Acetylcholinesterase by Phosphorodichloridic Acid. *Chem. Res. Toxicol.* **13**, 652–657 (2000).

57. Fukuto, T. R. Relationships between the structure of organophosphorus compounds and their activity as acetylcholinesterase inhibitors. *Bull. World Health Organ.* **44**, 31–42 (1971).

58. Makhaeva, G. F. *et al.* Kinetics and mechanism of inhibition of serine esterases by fluorinated aminophosphonates. *Chem. Biol. Interact.* **187**, 177–184 (2010).

59. Malla, R. K., Bandyopadhyay, S., Spilling, C. D., Dutta, S. & Dupureur, C. M. The First Total Synthesis of (±)-Cyclophostin and (±)-Cyclipostin P: Inhibitors of the Serine Hydrolases Acetyl Cholinesterase and Hormone Sensitive Lipase. *Org. Lett.* **13**, 3094–3097 (2011).

60. Wandhammer, M. *et al.* Structural Study of the Complex Stereoselectivity of Human Butyrylcholinesterase for the Neurotoxic V-agents*. *J. Biol. Chem.* **286**, 16783–16789 (2011).

61. Ashani, Y., Snyder, S. L. & Wilson, I. B. Inhibition of cholinesterase by 1,3,2-dioxaphosphorinane 2-oxide derivatives. *Biochemistry* **11**, 3518–3523 (1972).

62. Saeidian, H., Hosseini, S. E., Amoozadeh, A., Naseri, M. T. & Babri, M. Investigation of sarin(Se) reactivity against human plasma proteins using liquid chromatography–tandem mass spectrometry. *J. Mass Spectrom.* **53**, 138–145 (2018).

63. Béchet, J.-J., Dupaix, A., Roucous, C. & Bonamy, A.-M. Inactivation of proteases and esterases by halomethylated derivatives of dihydrocoumarins. *Biochimie* **59**, 241–246 (1977).

64. Mmutle, T. B. & Whiteley, C. G. Structural Requirements, Synthesis and Interaction of Secondary Nitrosamine Inhibitors with Acetylcholinesterase from Torpedo Fuscomaculata. *J. Enzym. Inhib.* **8**, 1–15 (1994).

65. Eid, P., Goeldner, M. P., Hirth, C. G. & Jost, P. Photosuicide inactivation of acetylcholinesterase by nitrosamine derivatives. *Biochemistry* **20**, 2251–2256 (1981).

66. BEDDOE, F., NICHOLLS, P. J. & SMITH, H. J. Potential irreversible inhibitors of acetylcholinesterase: N-trimethyl-N'-iodoacetyl diaminoalkane iodides. *J. Pharm. Pharmacol.* **23**, 865–867 (1971).

67. Beddoe, (miss) F. & Smith, H. J. Inhibition of acetylcholinesterase by dibenamine and dibenzyline. *J. Pharm. Pharmacol.* **23**, 37–49 (1971).

68. Cohen, S. G., Lieberman, D. L., Hasan, F. B. & Cohen, J. B. 1-Bromopinacolone, an active site-directed covalent inhibitor for acetylcholinesterase. *J. Biol. Chem.* **257**, 14087–14092 (1982).

69. Patockal, J., Jun, D. & Kucal, K. In vitro inactivation of rat brain acetylcholinesterase by DSP-4 and its derivatives OS-21 and OS-23 and protective activity of tacrine (9-amino-1,2,3,4-tetrahydroacridine). *Acta Medica (Hradec Kralove)* **48**, 145–148 (2005).

70. Belleau, B. & Tani, H. A Novel Irreversible Inhibitor of Acetylcholinesterase Specifically Directed at the Anionic Binding Site: Structure—Activity Relationships. *Mol. Pharmacol.* **2**, 411–422 (1966).

71. O'Brien, R. D. Binding sites of cholinesterases. Alkylation by an aziridinium derivative. *Biochem. J.* **113**, 713–719 (1969).

72. Cengiz, D., Çokugraş, A. N., Kilinç, K. & Tezcan, E. F. Histidine Modification of Human Serum Butyrylcholinesterase. *Biochem. Mol. Med.* **61**, 52–57 (1997).

73. Westkaemper, R. B. & Abeles, R. H. Novel inactivators of serine proteases based on 6-chloro-2-pyrone. *Biochemistry* **22**, 3256–3264 (1983).

74. Bakry, N. M., Sherby, S. M., Eldefrawi, A. T. & Eldefrawi, M. E. Oxadiazolidinones: irreversible inhibition of cholinesterases and effects on acetylcholine receptors. *Neurotoxicology* **7**, 1–10 (1986).

75. Mittag, T. W. & Tormay, A. Effects of isoxazolium cations on some isolated muscle preparations. *Br. J. Pharmacol.* **43**, 604–611 (1971).

76. Segal, D. & Shalitin, Y. The interaction of acetylcholinesterase with carbodiimides. *FEBS Lett.* **147**, 197–200 (1982).

77. Trujillo-Ferrara, J. *et al.* Reversible and irreversible inhibitory activity of succinic and maleic acid derivatives on acetylcholinesterase. *Eur. J. Pharm. Sci.* **18**, 313–322 (2003).

78. Correa-Basurto, J. *et al.* Inhibition of acetylcholinesterase by two arylderivatives: 3a-Acetoxy-5H-pyrrolo(1,2-a) (3,1)benzoxazin-1,5-(3aH)-dione and cis-N-p-Acetoxy-phenylisomaleimide. *J. Enzyme Inhib. Med. Chem.* **21**, 133–138 (2006).

79. Zueva, I. V., Lushchekina, S. V., Pottie, I. R., Darvesh, S. & Masson, P. 1-(3-Tert-Butylphenyl)-2,2,2-Trifluoroethanone as a Potent Transition-State Analogue Slow-Binding Inhibitor of Human Acetylcholinesterase: Kinetic, MD and QM/MM Studies. *Biomolecules* **10**, 1608 (2020).

80. Brodbeck, U., Schweikert, K., Gentinetta, R. & Rottenberg, M. Fluorinated aldehydes and ketones acting as quasi-substrate inhibitors of acetylcholinesterase. *Biochim. Biophys. Acta BBA - Enzymol.* **567**, 357–369 (1979).

81. Voss, G. & Neumann, R. Diethylmesoxalate hydrate, a new irreversible inhibitor of cholinesterases. *Experientia* **35**, 583–584 (1979).

82. Camerino, E. *et al.* Difluoromethyl ketones: Potent inhibitors of wild type and carbamate-insensitive G119S mutant Anopheles gambiae acetylcholinesterase. *Bioorg. Med. Chem. Lett.* **25**, 4405–4411 (2015).

83. Wilson, I. B. & Silman, I. Effects of quaternary ligands on the inhibition of acetylcholinesterase by arsenite. *Biochemistry* **16**, 2701–2708 (1977).

84. Anikienko, K. A., Bychikhin, E. A., Reznik, V. S., Akamsin, D. & Galyametdinova, I. V. Compounds with the dioxopyrimidine cycle inhibit cholinesterases from different groups of animals. *Chem. Biol. Interact.* **175**, 286–292 (2008).

85. Kumar, A. & Darreh-Shori, T. DMSO: A Mixed-Competitive Inhibitor of Human Acetylcholinesterase. *ACS Chem. Neurosci.* **8**, 2618–2625 (2017).

86. Chen, J.-W., Luo, Y.-L., Hwang, M.-J., Peng, F.-C. & Ling, K.-H. Territrem B, a Tremorgenic Mycotoxin That Inhibits Acetylcholinesterase with a Noncovalent yet Irreversible Binding Mechanism*. *J. Biol. Chem.* **274**, 34916–34923 (1999).

87. Al-Rashid, Z. F. & Hsung, R. P. (+)-Arisugacin A—Computational evidence of a dual binding site covalent inhibitor of acetylcholinesterase. *Bioorg. Med. Chem. Lett.* **21**, 2687–2691 (2011).

88. Pang, Y.-P. *et al.* Selective and Irreversible Inhibitors of Mosquito Acetylcholinesterases for Controlling Malaria and Other Mosquito-Borne Diseases. *PLOS ONE* **4**, e6851 (2009).

89. Gorecki, L. *et al.* Cysteine-Targeted Insecticides against A. gambiae Acetylcholinesterase Are Neither Selective nor Reversible Inhibitors. *ACS Med. Chem. Lett.* **11**, 65–71 (2020).

90. Dou, D. *et al.* Novel Selective and Irreversible Mosquito Acetylcholinesterase Inhibitors for Controlling Malaria and Other Mosquito-Borne Diseases. *Sci. Rep.* **3**, 1068 (2013).

91. Tomlinson, G., Mutus, B., Mclennan, I. & Mooibroek, M. J. Activation and inactivation of purified acetylcholinesterase from Electrophorus electricus by lanthanum(III). *Biochim. Biophys. Acta BBA - Protein Struct. Mol. Enzymol.* **703**, 142–148 (1982).

92. Tomlinson, G., Mutus, B. & McLennan, I. Activation and inactivation of acetylcholinesterase by metal ions. *Can. J. Biochem.* **59**, 728–735 (1981).

93. Frasco, M. F. *et al.* Mechanisms of cholinesterase inhibition by inorganic mercury. *FEBS J.* **274**, 1849–1861 (2007).

94. Hastings, F. L., Lucier, G. W. & Klein, R. Methylmercury-cholinesterase interactions in rats. *Environ. Health Perspect.* **12**, 127–130 (1975).

95. Shinar, E., Navok, T. & Chevion, M. The analogous mechanisms of enzymatic inactivation induced by ascorbate and superoxide in the presence of copper. *J. Biol. Chem.* **258**, 14778–14783 (1983).

96. den Hartog, G. J. M., Vegt, E., van der Vijgh, W. J. F., Haenen, G. R. M. M. & Bast, A. Hypochlorous Acid Is a Potent Inhibitor of Acetylcholinesterase. *Toxicol. Appl. Pharmacol.* **181**, 228–232 (2002).

97. Schallreuter, K. U. & Elwary, S. Hydrogen peroxide regulates the cholinergic signal in a concentration dependent manner. *Life Sci.* **80**, 2221–2226 (2007).

98. Kanazawa, H., Fujimoto, S. & Ohara, A. Inactivation of Cholinesterase by Ascorbic Acid in the Presence of Cupric Ions : A Possible Mechanism for the Inactivation of an Enzyme by the Metal-Catalyzed Oxidation System. *Biol. Pharm. Bull.* **18**, 1179–1183 (1995).

99. Mir, Y., Houde, D. & van Lier, J. E. Photodynamic inhibition of acetylcholinesterase after two-photon excitation of copper tetrasulfophthalocyanine. *Lasers Med. Sci.* **23**, 19–25 (2008).

100. Herz, F. Stability of Membrane Acetylcholinesterase in Human Erythrocytes Treated with Tetraphenylboron. *Enzyme* **20**, 151–155 (1975).

101. Tomlinson, G., Cummings, M. D. & Hryshko, L. Photoinactivation of acetylcholinesterase by erythrosin B and related compounds. *Biochem. Cell Biol.* **64**, 515–522 (1986).

102. Tacal, O., Li, B., Lockridge, O. & Schopfer, L. M. Resistance of Human Butyrylcholinesterase to Methylene Blue-Catalyzed Photoinactivation; Mass Spectrometry Analysis of Oxidation Products. *Photochem. Photobiol.* **89**, 336–348 (2013).

103. Stengelin, S., Walther, C. & Hucho, F. Azidophenantridinium compounds as photoaffinity labels of cholinergic proteins. *Biochim. Biophys. Acta BBA - Gen. Subj.* **542**, 107–114 (1978).

104. Ehret-Sabatier, L., Goeldner, M. P. & Hirth, C. G. p-Butyroxybenzenediazonium fluoroborate, substrate of acetylcholinesterase and butyrylcholinesterase, discriminates between the two enzymes by a specific affinity labelling. *Biochim. Biophys. Acta BBA - Protein Struct. Mol. Enzymol.* **1076**, 137–142 (1991).

105. Ehret-Sabatier, L., Schalk, I., Goeldner, M. & Hirth, C. Photoaffinity labelling of cholinesterases. *Eur. J. Biochem.* **203**, 475–481 (1992).

106. Nachon, F. *et al.* Trp82 and Tyr332 Are Involved in Two Quaternary Ammonium Binding Domains of Human Butyrylcholinesterase as Revealed by Photoaffinity Labeling with [3H]DDF. *Biochemistry* **37**, 10507–10513 (1998).

107. Coleman, B. A. & Oswald, R. E. Identification of the benzomorphan opiate binding site on the catalytic subunit of acetylcholinesterase. *Mol. Pharmacol.* **43**, 217–225 (1993).

108. Mautner, H. G. & Bartels, E. Interactions of p-Nitrobenzene Diazonium Fluoroborate and Analogs with the Active Sites of Acetylcholine-Receptor and -Esterase*. *Proc. Natl. Acad. Sci.* **67**, 74–78 (1970).

109. Kiefer, H., Lindstrom, J., Lennox, E. S. & Singer, S. J. Photo-Affinity Labeling of Specific Acetylcholine-Binding Sites on Membranes. *Proc. Natl. Acad. Sci.* **67**, 1688–1694 (1970).

110. Meden, A. *et al.* Pseudo-irreversible butyrylcholinesterase inhibitors: Structure–activity relationships, computational and crystallographic study of the N-dialkyl O-arylcarbamate warhead. *Eur. J. Med. Chem.* **247**, 115048 (2023).

111. Stojan, J. Rapid Mechanistic Evaluation and Parameter Estimation of Putative Inhibitors in a Single-Step Progress-Curve Analysis: The Case of Horse Butyrylcholinesterase. *Molecules* **22**, 1248 (2017).

112. The UniProt Consortium. UniProt: the universal protein knowledgebase in 2021. *Nucleic Acids Res.* **49**, D480–D489 (2021).

113. Bevc, S. *et al.* ENZO: A Web Tool for Derivation and Evaluation of Kinetic Models of Enzyme Catalyzed Reactions. *PLOS ONE* **6**, e22265 (2011).

114. Janssen, A. P. A. *et al.* Development of a Multiplexed Activity-Based Protein Profiling Assay to Evaluate Activity of Endocannabinoid Hydrolase Inhibitors. *ACS Chem. Biol.* **13**, 2406–2413 (2018).

115. Sobotič, B. *et al.* Proteomic Identification of Cysteine Cathepsin Substrates Shed from the Surface of Cancer Cells *. *Mol. Cell. Proteomics* **14**, 2213–2228 (2015).

116. Cox, J. & Mann, M. MaxQuant enables high peptide identification rates, individualized p.p.b.-range mass accuracies and proteome-wide protein quantification. *Nat. Biotechnol.* **26**, 1367–1372 (2008).

117. Tyanova, S., Temu, T. & Cox, J. The MaxQuant computational platform for mass spectrometry-based shotgun proteomics. *Nat. Protoc.* **11**, 2301–2319 (2016).

118. Cox, J. *et al.* Andromeda: A Peptide Search Engine Integrated into the MaxQuant Environment. *J. Proteome Res.* **10**, 1794–1805 (2011).

119. Brazzolotto, X., Igert, A., Guillon, V., Santoni, G. & Nachon, F. Bacterial Expression of Human Butyrylcholinesterase as a Tool for Nerve Agent Bioscavengers Development. *Molecules* **22**, 1828 (2017).

120. Chambers, M. C. *et al.* A cross-platform toolkit for mass spectrometry and proteomics. *Nat. Biotechnol.* **30**, 918–920 (2012).

121. Wenig, P. & Odermatt, J. OpenChrom: a cross-platform open source software for the mass spectrometric analysis of chromatographic data. *BMC Bioinformatics* **11**, 405 (2010).

122. Marty, M. T. *et al.* Bayesian Deconvolution of Mass and Ion Mobility Spectra: From Binary Interactions to Polydisperse Ensembles. *Anal. Chem.* **87**, 4370–4376 (2015).

123. Nachon, F. *et al.* Engineering of a monomeric and low-glycosylated form of human butyrylcholinesterase. *Eur. J. Biochem.* **269**, 630–637 (2002).

124. Brazzolotto, X. *et al.* Human butyrylcholinesterase produced in insect cells: huprine-based affinity purification and crystal structure. *FEBS J.* **279**, 2905–2916 (2012).

125. Vonrhein, C. *et al.* Data processing and analysis with the autoPROC toolbox. *Acta Crystallogr. Sect. D* **67**, 293–302 (2011).

126. Incardona, M.-F. *et al.* EDNA: a framework for plugin-based applications applied to X-ray experiment online data analysis. *J. Synchrotron Radiat.* **16**, 872–879 (2009).

127. Liebschner, D. *et al.* Macromolecular structure determination using X-rays, neutrons and electrons: recent developments in Phenix. *Acta Crystallogr. Sect. Struct. Biol.* **75**, 861–877 (2019).

128. Moriarty, N. W., Grosse-Kunstleve, R. W. & Adams, P. D. electronic Ligand Builder and Optimization Workbench (eLBOW): a tool for ligand coordinate and restraint generation. *Acta Crystallogr. Sect. D* **65**, 1074–1080 (2009).

129. Emsley, P., Lohkamp, B., Scott, W. G. & Cowtan, K. Features and development of ıt Coot. *Acta Crystallogr. Sect. D* **66**, 486–501 (2010).

130. Bowers, K. J. *et al.* Scalable algorithms for molecular dynamics simulations on commodity clusters. in *Proceedings of the 2006 ACM/IEEE conference on Supercomputing* 84-es (Association for Computing Machinery, New York, NY, USA, 2006). doi:10.1145/1188455.1188544.

131. Madhavi Sastry, G., Adzhigirey, M., Day, T., Annabhimoju, R. & Sherman, W. Protein and ligand preparation: parameters, protocols, and influence on virtual screening enrichments. *J. Comput. Aided Mol. Des.* **27**, 221–234 (2013).

132. Jacobson, M. P. *et al.* A hierarchical approach to all-atom protein loop prediction. *Proteins Struct. Funct. Bioinforma.* **55**, 351–367 (2004).

133. Shelley, J. C. *et al.* Epik: a software program for pKaprediction and protonation state generation for drug-like molecules. *J. Comput. Aided Mol. Des.* **21**, 681–691 (2007).

134. Koellner, G. *et al.* Active-site gorge and buried water molecules in crystal structures of acetylcholinesterase from Torpedo californica11Edited by R. Huber. *J. Mol. Biol.* **296**, 713–735 (2000).

135. Olsson, M. H. M., Søndergaard, C. R., Rostkowski, M. & Jensen, J. H. PROPKA3: Consistent Treatment of Internal and Surface Residues in Empirical pKa Predictions. *J. Chem. Theory Comput.* **7**, 525–537 (2011).

136. Radic, Z. *et al.* Expression of recombinant acetylcholinesterase in a baculovirus system: kinetic properties of glutamate 199 mutants. *Biochemistry* **31**, 9760–9767 (1992).

137. Gibney, G., Camp, S., Dionne, M., MacPhee-Quigley, K. & Taylor, P. Mutagenesis of essential functional residues in acetylcholinesterase. *Proc. Natl. Acad. Sci.* **87**, 7546–7550 (1990).

138. Vagedes, P., Rabenstein, B., Åqvist, J., Marelius, J. & Knapp, E.-W. The Deacylation Step of Acetylcholinesterase: Computer Simulation Studies. *J. Am. Chem. Soc.* **122**, 12254–12262 (2000).

139. Zhang, Y., Kua, J. & McCammon, J. A. Role of the Catalytic Triad and Oxyanion Hole in Acetylcholinesterase Catalysis: An ab initio QM/MM Study. *J. Am. Chem. Soc.* **124**, 10572–10577 (2002).

140. Zhou, Y., Wang, S. & Zhang, Y. Catalytic Reaction Mechanism of Acetylcholinesterase Determined by Born−Oppenheimer Ab Initio QM/MM Molecular Dynamics Simulations. *J. Phys. Chem. B* **114**, 8817–8825 (2010).

141. Driant, T., Nachon, F., Ollivier, C., Renard, P.-Y. & Derat, E. On the Influence of the Protonation States of Active Site Residues on AChE Reactivation: A QM/MM Approach. *ChemBioChem* **18**, 666–675 (2017).

142. Wang, J. *et al.* The protonation state of Glu202 in acetylcholinesterase. *Proteins Struct. Funct. Bioinforma.* **90**, 485–492 (2022).

143. Suárez, D. & Field, M. J. Molecular dynamics simulations of human butyrylcholinesterase. *Proteins Struct. Funct. Bioinforma.* **59**, 104–117 (2005).

144. Wan, X., Yao, Y., Fang, L. & Liu, J. Unexpected protonation state of Glu197 discovered from simulations of tacrine in butyrylcholinesterase. *Phys. Chem. Chem. Phys.* **20**, 14938–14946 (2018).

145. Zlobin, A., Smirnov, I. & Golovin, A. Dynamic interchange between two protonation states is characteristic of active sites of cholinesterases. *Protein Sci.* **33**, e5100 (2024).

146. Lu, C. *et al.* OPLS4: Improving Force Field Accuracy on Challenging Regimes of Chemical Space. *J. Chem. Theory Comput.* **17**, 4291–4300 (2021).

147. Friesner, R. A. *et al.* Extra precision glide: Docking and scoring incorporating a model of hydrophobic enclosure for protein-ligand complexes. *J. Med. Chem.* **49**, 6177–6196 (2006).

148. Zhu, K. *et al.* Docking covalent inhibitors: A parameter free approach to pose prediction and scoring. *J. Chem. Inf. Model.* **54**, 1932–1940 (2014).

149. Jorgensen, W. L., Chandrasekhar, J., Madura, J. D., Impey, R. W. & Klein, M. L. Comparison of simple potential functions for simulating liquid water. *J. Chem. Phys.* **79**, 926–935 (1983).

150. Jorgensen, W. L., Maxwell, D. S. & Tirado-Rives, J. Development and Testing of the OPLS All-Atom Force Field on Conformational Energetics and Properties of Organic Liquids. *J. Am. Chem. Soc.* **118**, 11225–11236 (1996).

151. Banks, J. L. *et al.* Integrated Modeling Program, Applied Chemical Theory (IMPACT). *J. Comput. Chem.* **26**, 1752–1780 (2005).

152. Bochevarov, A. D. *et al.* Jaguar: A high-performance quantum chemistry software program with strengths in life and materials sciences. *Int. J. Quantum Chem.* **113**, 2110–2142 (2013).

153. Yu, H. S., Watson, M. A. & Bochevarov, A. D. Weighted Averaging Scheme and Local Atomic Descriptor for pKa Prediction Based on Density Functional Theory. *J. Chem. Inf. Model.* **58**, 271–286 (2018).

154. Hermann, M. R., Pautsch, A., Grundl, M. A., Weber, A. & Tautermann, C. S. Covalent inhibitor reactivity prediction by the electrophilicity index—in and out of scope. *J. Comput. Aided Mol. Des.* **35**, 531–539 (2021).

155. Keeley, A., Ábrányi-Balogh, P. & Keserű, G. M. Design and characterization of a heterocyclic electrophilic fragment library for the discovery of cysteine-targeted covalent inhibitors. *MedChemComm* **10**, 263–267 (2019).

156. Electrophilicity Index | Chemical Reviews. https://pubs.acs.org/doi/10.1021/cr040109f.

157. Politzer, P. & Murray, J. S. An Occam's razor approach to chemical hardness: lex parsimoniae. *J. Mol. Model.* **24**, 332 (2018).

158. Shokhen, M. *et al.* Differentiating Serine and Cysteine Protease Mechanisms by New Covalent QSAR Descriptors. *ChemBioChem* **12**, 1023–1026 (2011).

159. Karelson, M., Lobanov, V. S. & Katritzky, A. R. Quantum-Chemical Descriptors in QSAR/QSPR Studies. *Chem. Rev.* **96**, 1027–1044 (1996).

160. Zhang, Y., Liu, H. & Yang, W. Free energy calculation on enzyme reactions with an efficient iterative procedure to determine minimum energy paths on a combined ab initio QM/MM potential energy surface. *J. Chem. Phys.* **112**, 3483–3492 (2000).

161. Sørensen, A., Rasmussen, B. & Pittelkow, M. 77Se NMR Spectroscopy As a Sensitive Probe for Hammett σ Constants. *J. Org. Chem.* **80**, 3852–3857 (2015).

162. Duddeck, H. & Biallaβ, A. Substituent effects and stereochemistry in 125Te NMR spectroscopy. Diorganyltellurium dihalides and some tellurides and ditellurides. *Magn. Reson. Chem.* **32**, 303–311 (1994).

163. Meden, A., Knez, D. & Gobec, S. Facile Synthesis of Carbamoyl Fluorides via N-Carbamoylimidazole Activation. *ACS Omega* **10**, 6908–6917 (2025).

164. Turksoy, A., Scattolin, T., Bouayad-Gervais, S. & Schoenebeck, F. Facile Access to AgOCF3 and Its New Applications as a Reservoir for OCF2 for the Direct Synthesis of N−CF3, Aryl or Alkyl Carbamoyl Fluorides. *Chem. – Eur. J.* **26**, 2183–2186 (2020).

165. Bonnefoy, C. *et al.* Study of Carbamoyl Fluoride: Synthesis, Properties and Applications. *Chem. – Eur. J.* **28**, e202201589 (2022).

166. Zhen, L., Fan, H., Wang, X. & Jiang, L. Synthesis of Thiocarbamoyl Fluorides and Isothiocyanates Using CF3SiMe3 and Elemental Sulfur or AgSCF3 and KBr with Amines. *Org. Lett.* **21**, 2106–2110 (2019).

167. Liu, L., Ran, L.-Y., Gu, Y. & Zhang, C.-P. Facile synthesis of selenocarbamyl fluorides, selenoureas and their derivatives with [Me4N][SeCF3]. *Org. Chem. Front.* **8**, 5736–5743 (2021).

168. Dürr, A. B., Fisher, H. C., Kalvet, I., Truong, K.-N. & Schoenebeck, F. Divergent Reactivity of a Dinuclear (NHC)Nickel(I) Catalyst versus Nickel(0) Enables Chemoselective Trifluoromethylselenolation. *Angew. Chem. Int. Ed.* **56**, 13431–13435 (2017).

169. Santhosh, U. *et al.* Development of a Safe, Scalable Process for the Preparation of an Oxaisoxazolidinone. *Org. Process Res. Dev.* **18**, 1802–1806 (2014).

170. Weinstock, L. M., Davis, P., Handelsman, B. & Tull, R. A general synthetic system for 1,2,5-thiadiazoles. *Tetrahedron Lett.* **7**, 1263–1268 (1966).

171. Eriksen, B. L., Vedsø, P., Morel, S. & Begtrup, M. Synthesis of 2-Substituted 1-Hydroxyimidazoles through Directed Lithiation. *J. Org. Chem.* **63**, 12–16 (1998).

172. Duspara, P. A., Islam, Md. S., Lough, A. J. & Batey, R. A. Synthesis and Reactivity of N-Alkyl Carbamoylimidazoles: Development of N-Methyl Carbamoylimidazole as a Methyl Isocyanate Equivalent. *J. Org. Chem.* **77**, 10362–10368 (2012).
